# Delineating the functional activity of antibodies with cross-reactivity to SARS-CoV-2, SARS-CoV-1 and related sarbecoviruses

**DOI:** 10.1101/2024.04.24.590836

**Authors:** Felicitas Ruiz, Will Foreman, Michelle Lilly, Viren A. Baharani, Delphine M. Depierreux, Vrasha Chohan, Ashley L. Taylor, Jamie Guenthoer, Duncan Ralph, Frederick A. Matsen, Helen Y. Chu, Paul D. Bieniasz, Marceline Côté, Tyler N. Starr, Julie Overbaugh

## Abstract

The recurring spillover of pathogenic coronaviruses and demonstrated capacity of sarbecoviruses, such SARS-CoV-2, to rapidly evolve in humans underscores the need to better understand immune responses to this virus family. For this purpose, we characterized the functional breadth and potency of antibodies targeting the receptor binding domain (RBD) of the spike glycoprotein that exhibited cross-reactivity against SARS-CoV-2 variants, SARS-CoV-1 and sarbecoviruses from diverse clades and animal origins with spillover potential. One neutralizing antibody, C68.61, showed remarkable neutralization breadth against both SARS-CoV-2 variants and viruses from different sarbecovirus clades. C68.61, which targets a conserved RBD class 5 epitope, did not select for escape variants of SARS-CoV-2 or SARS-CoV-1 in culture nor have predicted escape variants among circulating SARS-CoV-2 strains, suggesting this epitope is functionally constrained. We identified 11 additional SARS-CoV-2/SARS-CoV-1 cross-reactive antibodies that target the more sequence conserved class 4 and class 5 epitopes within RBD that show activity against a subset of diverse sarbecoviruses with one antibody binding every single sarbecovirus RBD tested. A subset of these antibodies exhibited Fc-mediated effector functions as potent as antibodies that impact infection outcome in animal models. Thus, our study identified antibodies targeting conserved regions across SARS-CoV-2 variants and sarbecoviruses that may serve as therapeutics for pandemic preparedness as well as blueprints for the design of immunogens capable of eliciting cross-neutralizing responses.

**AUTHOR SUMMARY:** There is a large collection of sarbecoviruses related to SARS-CoV-2 circulating in animal reservoirs with the potential to spillover into humans. Neutralizing antibodies have the potential to protect against infection, although viral escape is common. In this study, we isolated several monoclonal antibodies that show broad activity against different sarbecoviruses. The antibodies target epitopes in the core of the receptor binding domain that are highly conserved in sequence across sarbecoviruses and emerging SARS-CoV-2 variants. One antibody showed remarkable breadth against both SARS-CoV-1 variants as well as diverse sarbecoviruses. The results of deep mutational scanning suggest that mutations at these predicted sites of escape may functionally constrain viral fitness. Our functional profiling of cross-reactive antibodies highlights vulnerable sites of sarbecoviruses, with some antibodies poised as broadly neutralizing candidates for therapeutic use against future sarbecovirus emergence.

## INTRODUCTION

In the past two decades, zoonotic betacoronaviruses, most notably SARS-CoV-1 and SARS-CoV-2 (sarbecoviruses), have infected humans and caused large-scale outbreaks with significant associated morbidity and mortality (1–4). Humans are permissive to many members of the sarbecovirus family because they encode a functional cellular receptor that permits viral entry. Both SARS-CoV-1 and SARS-CoV-2 initiate entry by binding to the angiotensin-converting enzyme 2 (ACE2) receptor via the receptor-binding domain (RBD) in the spike glycoprotein (5,6). Many bat sarbecoviruses that utilize human ACE2 for entry *in vitro* have been identified through epidemiological surveillance of animal reservoirs in East and Southeast Asia, suggesting the potential for more introduction of sarbecoviruses into humans (7–11). Most bat isolates from Africa and Europe cannot infect human cells and exhibit more narrow host species ACE2 usage (11,12). Recently, one of these clade 3 sarbecovirus has been found to utilize human ACE2 to infect cells, and human ACE2-independent clade 3 sarbecoviruses can acquire mutations that permit binding to human ACE2 with just two substitutions thus potentially broadening their host cell range to include humans as well (11,13). The COVID-19 pandemic has demonstrated the ability of the SARS-CoV-2 RBD to tolerate mutations (14,15), and evidence of recombinants (16) highlights the diversity of RBD that can be generated within a single sarbecovirus lineage that has circulated in the human population in just a few years. The large number of sarbecoviruses in animal reservoirs that can use the human ACE2 receptor for entry, coupled with the pathways for diversity and mutational tolerance among these viruses, underscores the urgent need to understand the immune responses that may be useful for eliciting in future spillover events.

Because of its role in viral entry, the RBD is a major target of neutralizing antibodies and many RBD-specific monoclonal antibodies (mAbs) have been characterized, providing insights into the major antigenic sites in the SARS-CoV-2 RBD (17–20). Several of the potent neutralizing mAbs identified to date target residues overlapping the ACE2 footprint (receptor binding motif; RBD class 1/2 mAbs based on Barnes et al (20) classification) and directly interfere with RBD-ACE2 binding (21,22). However, this binding interface is highly variable not only within the SARS-CoV-2 lineage but also across more diverse sarbecoviruses; thus class 1/2 mAbs typically show limited breadth against SARS-CoV-2 variants and limited cross-reactivity against other sarbecoviruses (23–25). In contrast, neutralizing antibodies that target the RBD core (class 4-5) target regions with high sequence conservation, which is thought to be driven by functional constraints in this region (26,27). These antibodies have been found to exhibit breadth in binding and neutralization across SARS-CoV-2 variants as well as some cross-reactive activity against animal sarbecoviruses indicating that broadly active RBD-targeting mAbs can be elicited by SARS-CoV-2 infections (21,25,28–30). However, few mAbs with cross-reactivity across the diverse classes of sarbecoviruses have been comprehensively characterized despite the potential utility of these for future spillover events (25,28–33).

mAbs that cross-react with diverse sarbecoviruses have been isolated from both SARS-CoV-1 and SARS-CoV-2 infected individuals (25,28–34). S309, which is a class 3 neutralizing mAb isolated from a SARS-CoV-1 infected individual (29), previously received emergency authorization for the treatment of SARS-CoV-2 infection (35). S309 not only showed neutralizing activity, but also the ability to kill infected cells via antibody-dependent cellular cytotoxicity (ADCC), and both neutralization and ADCC have been implicated in protection from infection and disease progression (36,37). Several class 4 cross-reactive mAbs have demonstrated protection against SARS-CoV-2 in animal models (28,33,38) although they have more limited clinical value because they show reduced breadth across SARS-CoV-2 variants (39,40). One class 5 mAb, S2H97, which has broad neutralization activity against SARS-CoV-2 Omicron variants and some animal sarbecoviruses, showed protection against SARS-CoV-2 challenge in animal models (25). Studies have suggested that S2H97 class 5-like mAbs contribute to less than ∼17% of infection- and vaccine-elicited antibody responses and are thus understudied (41,42).

We previously described a class 5-like neutralizing antibody C68.61, isolated from an individual with a Delta breakthrough infection, that binds a conserved epitope in the core of RBD and demonstrates neutralization breadth across SARS-CoV-2 variants as well as cross reactivity with SARS-CoV-1 (31). In this study, we demonstrate cross-reactivity of the C68.61 mAb across sarbecovirus clades and show that it is refractory to viral escape both in cell culture and in nature. We further isolate additional mAbs from this individual, including several class-4 and class-5 RBD-specific mAbs that display neutralization activity across multiple sarbecoviruses. One such mAb, C68.185, displayed extensive binding and retained potent neutralization across tested bat-derived sarbecoviruses. Some mAbs also displayed potent ADCC activity in-line with the activity of therapeutic antibodies shown to reduce SARS-CoV-2 viral load in animal models (37). Taken together, these cross-reactive mAbs with differential functional activity and distinct epitope footprints in the RBD may contribute to pan-coronavirus pandemic preparedness and inform vaccine efforts.

## RESULTS

### Isolation of C68 cross-reactive antibodies targeting the RBD

We reported the identification of four potent neutralizing mAbs from an individual (C68) at 30 days after a SARS-CoV-2 Delta breakthrough infection, including C68.61, which showed consistent activity against SARS-CoV-2 variants as well as SARS-CoV-1 (31). These mAbs were reconstructed from memory B cells that bound the SARS-CoV-2 Delta prefusion-stabilized spike trimer and/or the Wuhan-Hu-1 S2 spike subunit. To explore the range of RBD cross-reactive neutralizing antibodies elicited in this Delta breakthrough infection case, we recovered additional antibodies from this 30-day post-infection time point. We identified 42 antibodies that bound the SARS-CoV-2 spike trimer and RBD protein in an enzyme-linked immunosorbent assay (ELISA). All C68 antibodies bound the SARS-CoV-2 Wuhan-Hu-1 (WH-1) vaccine strain trimer and RBD subunit to comparable levels (**Figs 1A, C**) when tested at a fixed concentration (500 ng/ml) and to higher levels than the negative control Fl6v3 (43), an influenza-specific mAb. Of the newly isolated C68 antibodies, we identified 11 additional antibodies that exhibited cross-reactive binding between SARS-CoV-2 and SARS-CoV-1 (**Figs 1A, C**). Only 3/31 (10%) of the SARS-CoV-2 specific mAbs retained activity against the highly mutated SARS-CoV-2 Omicron XBB trimer, whereas 10/11 (91%) of the cross-reactive mAbs exhibited activity. Among these 42 mAbs that bound the WH-1 spike, we observed variation in neutralizing activities against the corresponding SARS-CoV-2 WH-1 pseudovirus, ranging from an IC50 of 0.01 to >20 μg/mL (**Fig 1C**). Generally, SARS-CoV-2 specific mAbs were more potent neutralizers (median IC50 = 0.05 μg/mL; IC50 range 0.01 μg/mL to >20 μg/mL) than cross-reactive mAbs (median IC50 = 5.3 μg/mL; IC50 range 0.06 μg/mL to >20 μg/mL; **Fig 1C**).

**Fig 1.**
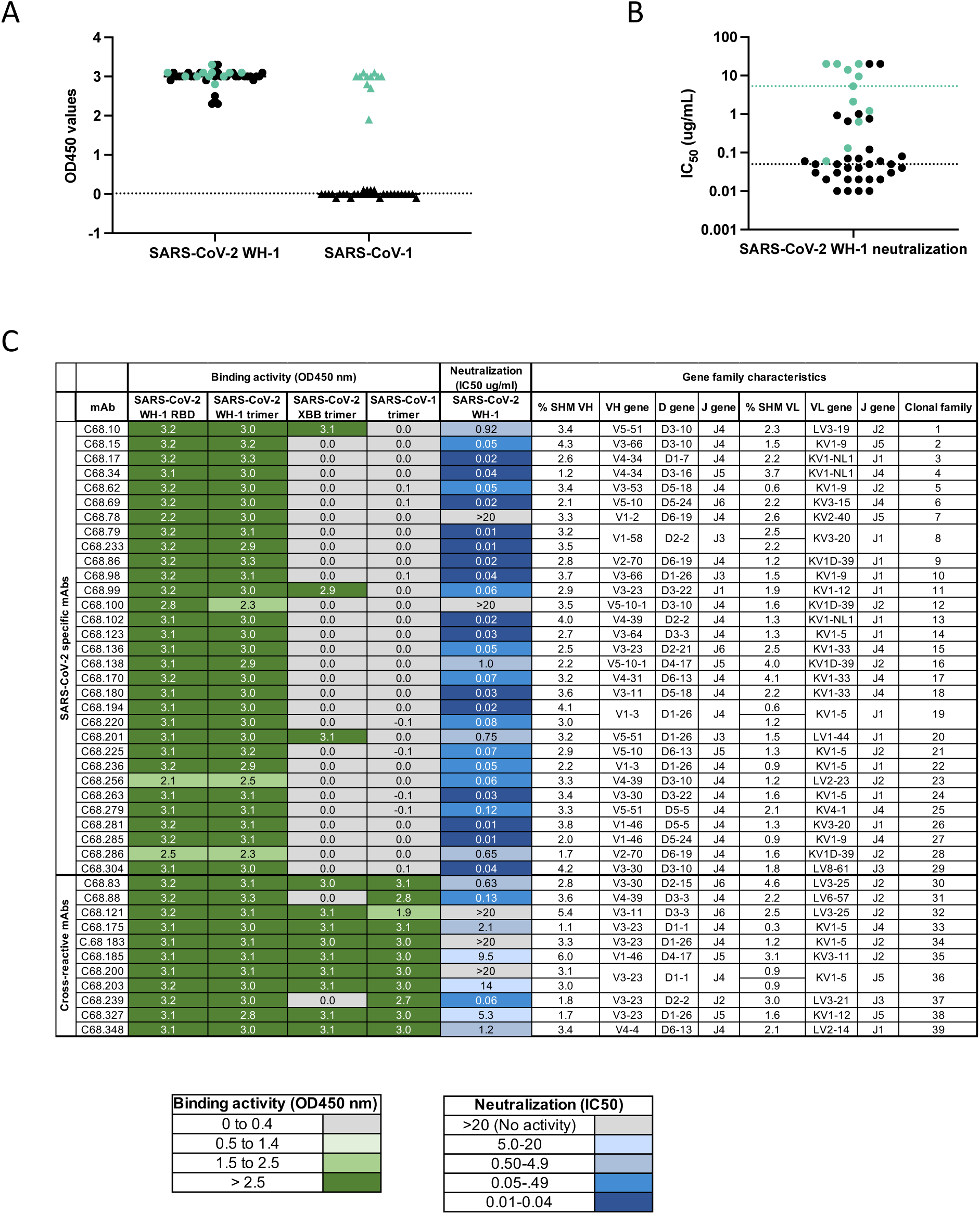
Identification and molecular characteristics of SARS-CoV-1 and SARS-CoV-2 cross-reactive mAbs. **(A)** C68 mAbs binding to SARS-CoV-2 or SARS-CoV-1 recombinant spike glycoprotein by ELISA (OD450nm values). C68 antibodies that bound SARS-CoV-2 trimer and SARS-CoV-1 trimer (designated by green closed circles; SARS-CoV-2 trimer-specific mAbs shown in black closed circles). Antibodies tested at 500 ng/ml, represents the average of two technical replicates shown as background-corrected OD450nm values. Dotted line represents 3 standard deviations above the negative control Fl6v3, (43) an influenza-specific mAb. **(B)** Neutralization of spike-pseudotyped lentiviruses by C68 mAbs against SARS-CoV-2 WH-1. C68 antibodies that bound SARS-CoV-2 trimer and SARS-CoV-1 trimer are indicated (in green). IC50 values (μg/mL) were calculated by nonlinear regression analysis in the statistical software package PRISM from at least two independent experiments. Black dotted line represents the median IC50 value for the SARS-CoV-2 specific mAbs. Green line represents the median IC50 value for the cross-reactive mAbs. **(C)** Summary table representing gene family usage, somatic hypermutation percent (% SHM relative to germline), and a heatmap of functional activity of C68 mAbs. OD450 values indicate absorbance at OD450nm values where each mAb was tested at 500 ng/ml from technical replicates and background-corrected.

We next evaluated the genotypic properties of these mAbs, including somatic hypermutation (SHM), V(D)J gene usage and clonal diversity. We used partis, a computational tool designed to assess levels of SHM using a distance-based clustering on the inferred naive sequence, along with a hidden Markov model to infer clonality (44,45). There was a high degree of clonal diversity among these 42 mAbs with antibodies representing 39 clonal lineages (**Fig 1C, S1 Fig; S1 Table**). Low levels of SHM were observed among SARS-CoV-2-specific and cross-reactive mAbs (geomean 2.9% and 3.3% SHM relative to germline in the heavy chain genes, respectively). We observed overrepresentation of V_H_3-23 among cross-reactive mAbs, and this has been shown to be a common gene family elicited from COVID-19 convalescent individuals (46–49).

### Cross-reactive mAbs exhibit binding across SARS-CoV-2 variants and human and animal sarbecoviruses

To determine whether the 11 cross-reactive mAbs that bound both SARS-CoV-1 and SARS-CoV-2 exhibited breadth across SARS-CoV-2 variants, we tested binding to various recombinant SARS-CoV-2 spike proteins. For comparison, we also included the previously characterized cross-reactive mAb C68.61 that had been tested against SARS-CoV-1 and SARS-CoV-2 (31) but not more diverse sarbecoviruses. All antibodies bound SARS-CoV-2 WH-1 and the Delta variant proteins with half-maximal effective concentrations (EC50s) ranging from 6 to 21 ng/mL and 5 to 28 ng/ml, respectively (**Fig 2A, S2 Fig**). Of these, ten mAbs, including C68.61, retained similar binding breadth across SARS-CoV-2 variants including Omicron XBB and BQ.1.1 variants (EC50s 9 to 45 ng/ml and 11 to 62 ng/ml; **Fig 2A, S2 Fig**); only two mAbs, C68.88 and C68. 239, did not bind Omicron variants. Together, these 12 mAbs bound SARS-CoV-1 spike glycoprotein with EC50s ranging from 12 to 237 ng/ml (**Fig 2A**), with ten mAbs displaying EC50s less than 25 ng/ml, which is comparable to what has previously been reported for the SARS-CoV-1 cross-reactive mAb S309 (31).

**Fig 2.**
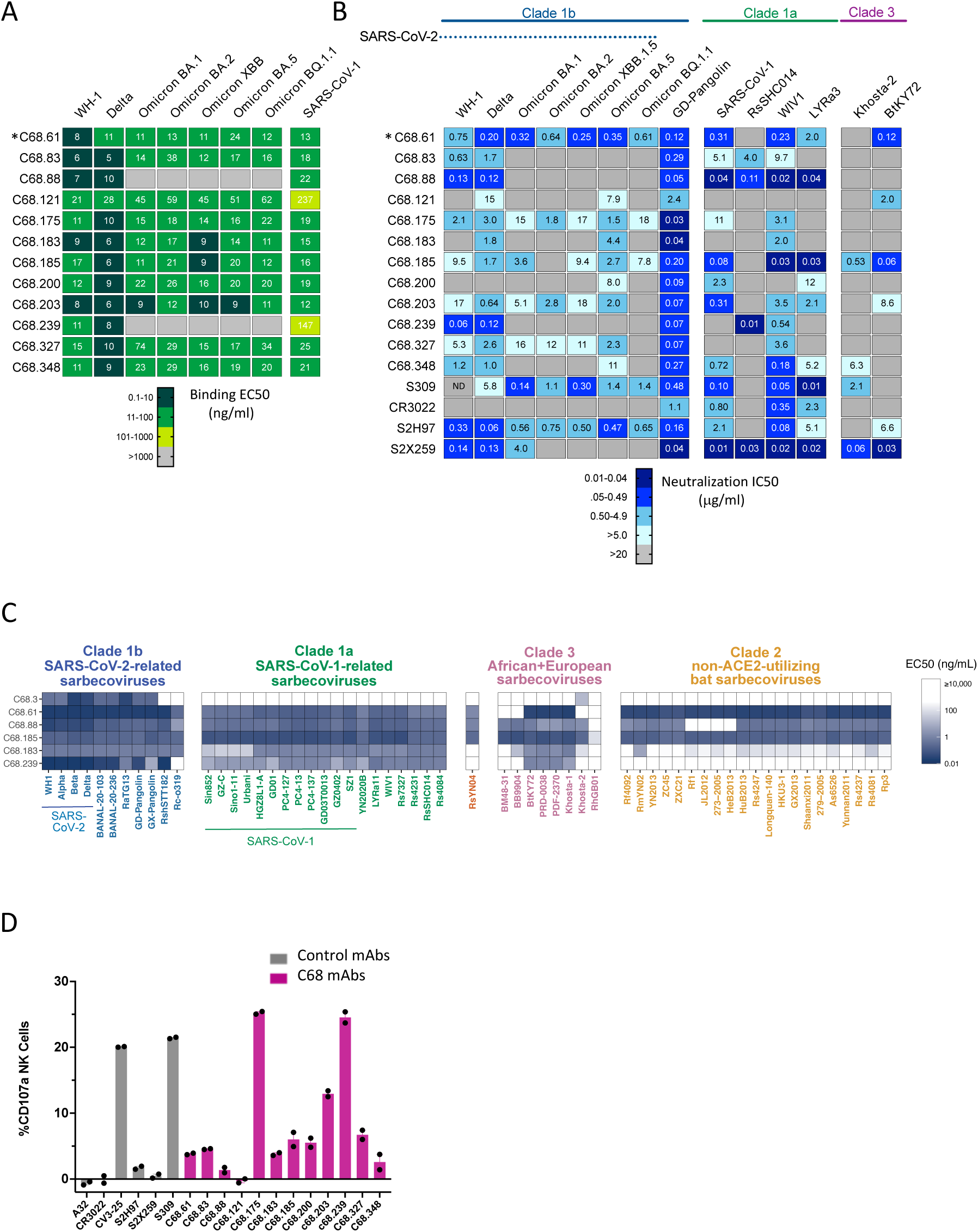
Comprehensive functional characterization of C68 cross-reactive mAbs. **(A)** C68 mAbs binding to SARS-CoV-2 variants and SARS-CoV-1 recombinant spike glycoprotein by ELISA (OD450nm values). Half-maximal effective concentrations (EC50 values) calculated by nonlinear regression analysis from at least two independent replicates. *Binding activity for C68.61 from (31). **(B)** SARS-CoV-2 variants and sarbecovirus neutralization activity. IC50 values (μg/mL) were calculated by nonlinear regression analysis from at least two independent experiments with technical replicates. Samples with IC50 values above 20 μg/mL were plotted at 20 μg/mL (no activity). *Neutralization activity for C68.61 and S309 against SARS-CoV-2 variants and SARS-CoV-1 from (31). S309 neutralizing activity against WH-1 was not determined (ND). **(C)** Pan-sarbecovirus RBD yeast display to assess sarbecovirus binding breadth. EC50 (ng/ml) value represents the geometric mean across the independent barcodes for the corresponding sarbecovirus RBD. **(D)** C68 mAbs can mediate antibody dependent cellular cytotoxicity. Cells expressing SARS-CoV-2 D614G spike cells were incubated with C68 mAbs and effector cells (PBMC) for 4h. NK cell degranulation (% CD107a+) was measured as a proxy for ADCC. The assay was run in technical duplicate with two independent PBMC donors. Background subtracted values (target cells + PBMC only) from a single donor are shown.

To determine the functional activity of these cross-reactive mAbs, we measured neutralization against a panel of SARS-CoV-2 variants and bat SARS-like sarbecovirus strains known to utilize human ACE2 to infect cells: Pangolin GD, WIV1, RsSHC014, LYRa3 (8). WIV-1 and LyRa3 are 96% and 95% sequence similar to SARS-CoV-1 (Urbani strain) RBD whereas Pangolin GD is closely related to SARS-CoV-2 WH-1 (96% sequence similarity to RBD; **S3A Fig**). Among the SARS-CoV-1 related bat-derived isolates, RsSHC014 shares 82% amino acid similarity in the RBD to SARS-CoV-1 and 77% similarity to SARS-CoV-2 WH-1 RBD. We also examined neutralization against two representative clade 3 viruses that have been shown to utilize human ACE2 to infect cells *in vitro*, including Khosta-2 and the BtKY72 (K493Y/T498W mutant that confers human ACE2 binding) which share 70% and 74% similarity to SARS-CoV-1 (7,11). We included four previously described cross-reactive mAbs with functionally defined properties and epitopes for comparison: S309, S2H97, S2X259 and CR3022 (21,25,29,50).

C68.61, which was previously described as having breadth against SARS-CoV-2 variants ((31) and **Fig 2B**), also showed breadth against SARS-CoV-1-related sarbecoviruses WIV1 (IC50s 0.23 μg/mL) and LyRa3 (IC50s 2.0 μg/mL) (**Fig 2B**). C68.61 did not neutralize the more sequence divergent RsSHC014. We further examined neutralization against the more distantly-related clade 3 sarbecoviruses Khosta-2 and the BtKY72 (7,11). C68.61 neutralized BtKY72 at an IC50 of 0.12 μg/mL but did not neutralize Khosta-2. S2H97 (28), which partially overlaps the epitope escape profile of C68.61 defined by deep mutational scanning (DMS) (31), exhibited similar cross-neutralization but reduced potency against more diverse sarbecoviruses tested in parallel.

Among the 11 other newly isolated cross-reactive C68 mAbs, we observed limited breadth and potency against SARS-CoV-2 Omicron variants. These 11 mAbs had geometric mean IC50s ranging from 4.2 to 18 μg/mL among SARS-CoV-2 viruses (**S3B Fig**). Among animal sarbecovirus strains, all mAbs tested were able to potently neutralize Pangolin-GD, which is highly similar in sequence to SARS-CoV-2 WH-1 RBD and is generally an easy to neutralize SARS-CoV-2-related strain, with IC50s ranging from 0.03 to 2.4 μg/mL (**Fig 2B**). Seven of these eleven mAbs neutralized SARS-CoV-1 with IC50s ranging from 0.04 to 11 μg/mL. Several of these mAbs also neutralized various clade 1a viruses, including some mAbs that did not neutralize SARS-CoV-1, such as C68.239. Most notably, C68.88 neutralized all four SARS-CoV-1-related viruses tested (IC50s ranging from 0.02 to 0.11 μg/mL), which was similar to some of the most potent cross-reactive mAbs described to date, such as S2X259 (32) and S309 (29,51) tested here in parallel. C68.185 also showed similar potency against select SARS-CoV-1-related strains as S309, and neither neutralized RsSHC014 (31,52). C68.239 neutralized RsSHC014 with IC50 of 0.01 μg/mL, effectively complementing the activity of C68.185 against clade 1a viruses. There was more limited activity against clade 3 viruses among these 11 mAbs; only C68.185 neutralized both Khosta-2 and BtKY72.

Thus, this set of 12 cross-reactive mAbs included examples with: 1) activity against nearly all SARS-CoV-1 and SARS-CoV-2 variants tested (C68.61), 2) activity against all SARS-CoV-1 clade 1a variants, but limited breadth against SARS-CoV-2 (C68.88) and 3) potent activity against select non-SARS-CoV-2 sarbecoviruses (C68.185 and C68.239). While C68.61 showed the greatest overall breadth against the full virus panel tested (neutralization of 12/14 viruses tested with geometric mean IC50 across the 12 viruses of 0.62 μg/mL; **S3C Fig**), the previously described S2X259 showed breadth against SARS-CoV-1-related viruses (geomean IC50 of 0.32 μg/mL against human and animal sarbecovirus strains), although it loses neutralization activity against the SARS-CoV-2 Omicron variants as previously described (21,32).

### Exceptional binding breadth of C68 mAbs across sarbecoviruses

Given the cross neutralization profiles we observed for some mAbs, we explored the extent of their cross reactivity using a yeast library displaying diverse sarbecovirus RBD proteins from across the known evolutionary clades (25). The clades, which were defined based on phylogenetic classification of their RBD sequence, share ∼62-96% sequence similarity to SARS-CoV-2 RBD (25). We tested four mAbs displaying high cross-neutralization activity (C68.61, C68.88, C68.185, and C68.239) as well as one of the mAbs with more limited activity (C68.183) and the RBD mAb C68.3, which we identified previously from C68 and has broad SARS-CoV-2 activity but does not bind SARS-CoV-1 (31). C68.3 displayed a narrow binding profile with functional activity mostly restricted to the SARS-CoV-2-related sarbecoviruses as well as weak binding to Khosta-2 (**Fig 2C**). The weak, limited binding of C68.3 is consistent with other SARS-CoV-2-specific mAbs that lack SARS-CoV-1 cross-reactivity (25,34).

C68.61, C68.88 and C68.185 exhibited broad binding against the more sequence divergent sarbecovirus clades, whereas C68.183 and C68.239 showed more restricted binding. Both C68.183 and C68.239 displayed restricted or weak binding against clade 2 viruses, which are ACE2-independent and display low sequence similarity (∼62-65%) to SARS-CoV-2 RBD (**Fig 2C**). C68.61, C68.88 and C68.185 not only bound to clade 2 viruses, but also clade 3 viruses, in which only Khosta-2 strain can bind human ACE2 (**Fig 2C**; **S4A Fig**). These mAbs showed distinct binding cross-reactivity for specific RBDs and a pattern that suggests differential preference for binding to certain clades. For example, C68.61 exhibited higher binding to clade 1b RBDs (geomean EC50 0.25 ng/ml across clade 1b sarbecoviruses) than clade 1a (geomean EC50 1.9 ng/ml) RBDs whereas the opposite was true for C68.185, which bound with a higher affinity to the clade 1a RBDs than clade 1b RBDs (geomean EC50 0.66 ng/ml and 1.3 ng/ml, respectively) consistent with their differential neutralization profiles (**Fig 2B; S4A-B Fig**). Both C68.61 and C68.185 displayed the broadest binding profiles; they potently bound nearly all RBDs evaluated (EC50 values less than 10,000 ng/ml), including RBDs from ACE2-independent clade 2 viruses, suggesting they are targeting an epitope that is highly conserved across sarbecoviruses. Of note, C68.61, C68.88, C68.183, C68.185, and C68.239 all showed cross-reactive binding to the RsSHC014 RBD in the yeast assay, but only C68.88 and C68.239 showed detectable neutralization of RsSHC014 spike-pseudotyped lentivirus. The two RsSHC014-neutralizing antibodies bind to a similar class 4 epitope while the three non-neutralizers bind to the class 5 epitope (see below), suggesting that RsSHC014 might exhibit conformational dynamics that limit access to the class 5 epitope in the full spike context.

### Several cross-reactive C68 mAbs display Fc-mediated effector function

Given that ADCC, an Fc-dependent effector function largely mediated by natural killer (NK) cells, has been associated with protection from SARS-CoV-2 infection in animal models and human studies (36,37,53,54), we tested the ability of the 12 cross-reactive mAbs to trigger ADCC in primary human NK cells in the presence of SARS-CoV-2 spike expressing cells (target cells). We adapted a validated SARS-CoV-2 specific ADCC assay (gating strategy shown in **S5 Fig**) (55–57) to quantify NK cell activation (surface CD107a (a proxy for degranulation) and intracellular IFN-γ) against target cells expressing SARS-CoV-2 D614G spike protein. We included two spike-specific mAbs previously shown to mediate ADCC activity, CV3-25 and S309, which induced robust NK cell activation in this assay (20% and 21% CD107a+ expression, respectively; **Fig 2D**) (36,37). The negative control mAbs, A32 (HIV-specific mAb) (58) and S2X259 (a SARS-CoV-2 mAb previously reported to lack ADCC activity) (28), did not show appreciable activity in this assay with less than 0.5% NK activation. Two C68 mAbs, C68.175 and C68.239, induced the most robust NK cells CD107a expression (**Fig 2D**) to similar levels to CV3-25 and S309, the latter of which has been shown to mediate ADCC and improve infection outcome in animal studies (37,57). Most of the C68 mAbs demonstrated intermediate levels of CD107a expression, that were above the negative control mAbs, suggesting low level ADCC activity. These data were observed for two independent donors (**S6A Fig**) indicating that this phenotype is not donor dependent. CD107a expression also correlated with NK cell production of intracellular IFN-γ (**S6D-E Fig**), another marker for NK cell activation, and cell death of SARS-CoV-2 spike expressing target cells (**S6F-G Fig**). All antibodies bound target cells expressing the D614G spike above the negative control mAb (A32), except CR3022 and C68.121. Among the antibodies that could bind target cells, mAbs like C68.88 did not mediate ADCC functions, suggesting that this was not due to absence of binding ability (**S6B Fig**). Consistent with this, CD107a expression did not correlate with binding activity (**S6C Fig**). Overall, these data demonstrate that in addition to neutralization, some C68 RBD antibodies induce robust NK cell-mediated ADCC.

### Epitope profiling of cross-reactive antibodies suggest they cluster within two major epitope domains

To identify mutations that facilitate escape from antibody binding and assess the potential for antigenic variation at these sites, we employed a yeast display deep mutational scanning library encoding all possible amino-acid mutations in the RBD (14,25,59). This approach defines the functional epitope and generally captures key residues important for antibody-RBD interaction while also highlighting the mutations that are tolerant for viral functional activity. This approach has been used to map escape from structurally defined classes and these escape maps do tend to cluster by RBD antibody class (25,32,59,60). Using this approach, we classified C68 mAbs into two epitope profile groups: one which most resembles class 4-like mAbs and another which resembles class 5-like mAbs. The class 4-like mAbs include C68.88 and C68.239, which show activity against clade 1a sarbecoviruses (**Fig 2C**), while the class 5-like mAbs includes C68.61 and C68.185, which shows breadth across clade 1a/b sarbecoviruses and clade 3 viruses (**Fig 2C**).

Among the class-4 like mAbs, there is heterogeneity in the escape pathways. For example, both C68.88 and C68.239 share escape mutations centered on sites that include 374 - 377 (**Fig 3A**) but C68.239 has additional sites that confer escape from binding. Additionally, these escape profiles generally aligned well with the functional data. For example, substitutions at sites S375 and T376 occurred early in Omicron variants (**Fig 3B**) and are predicted to facilitate escape from C68.88 and C68.239, which is consistent with the observed losses in binding and neutralization against Omicron variants (**Fig 2A-B**). For this reason, the epitope of these two mAbs were profiled in SARS-CoV-2 WH-1 RBD backgrounds as they retain binding activity against this viral library. C68.88 appears to target residues that are relatively conserved across SARS-CoV-1-related sarbecoviruses (**Fig 3B**), consistent with the observed neutralization of these viruses (**Fig 2C**). C68.83, C68.121 and C68.348 show more limited overlap in specific amino acids that drive escape but in general, the mAbs in this class target epitopes distal from the ACE2 binding footprint (26).

**Fig 3.**
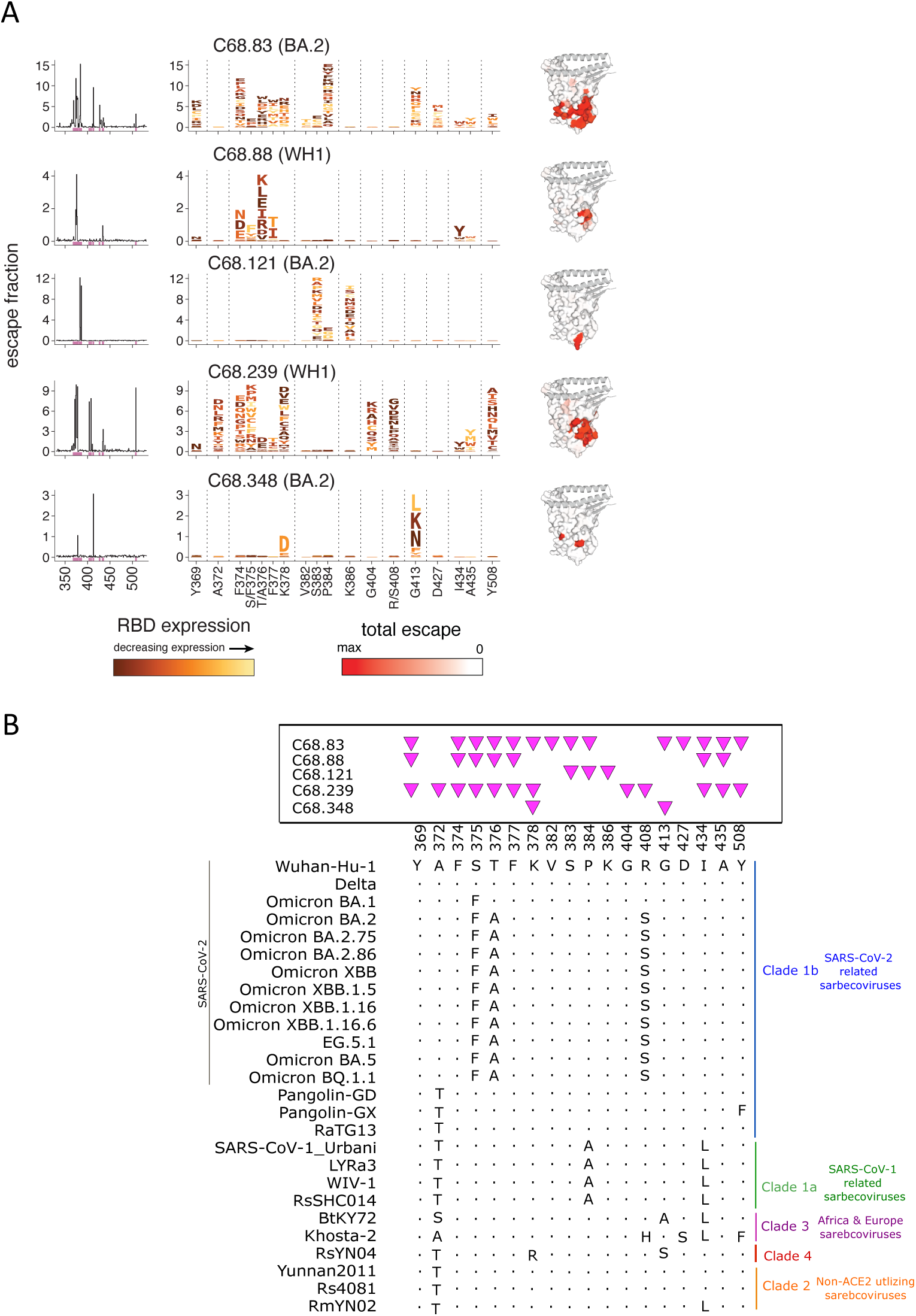
Epitope profiling by deep mutational scanning identifies a group of C68 mAbs targeting class 4 epitopes. **(A)** Complete escape maps of antibody-binding escape mutations for C68 mAbs using a yeast-displayed SARS-CoV-2 RBD deep mutational scanning system (Wuhan-Hu-1 RBD DMS for C68.88 and C68.239; Omicron BA.2 RBD DMS for C68. 83, C68.121, C68.348). Residue colors are assigned based on effect these mutations have on RBD expression (with yellow indicating mutations deleterious for RBD expression). The height of letters in the logo plots indicate level of escape by that amino acid at that site. Logo plot residue numbering is based on SARS-CoV-2 WH-1 (/Omicron BA.2 numbering). Experiments were performed in biological duplicate using independent mutant RBD libraries (14) that correlate well (**S9B Fig**) so escape fractions represent the average of these two independent biological replicates. On the right, sites of escape for C68 mAbs are mapped onto the RBD structure and gradient of red coloring indicates magnitude of escape fraction at each indicated site. **(B)** Multiple sequence alignments of select SARS-CoV-2 variants and sarbecoviruses are shown corresponding to sites of escape. Residue numbering based on SARS-CoV-2 Wuhan-Hu-1 sequence with shared amino acids denoted by dots (.) and differences across sarbecoviruses relative to the reference sequence (SARS-CoV-2 Wuhan-Hu-1) indicated. Pink triangles depict the sites of escape identified by the escape maps (**Fig 3A**).

The second cluster of C68 mAbs target regions that are spatially distinct from the first mAb group. Many mutations at these regions are not well tolerated, indicated by decreasing RBD expression at predicted escape pathways (**Fig 4A**). For example, some of the mutations that allow escape from binding at sites 464-466 reduce RBD expression (yellow and orange colors; **Fig 4A**) and this is true for many of the predicted escape mutations for class 5-like mAbs. Whereas the class 4-like mAbs are escaped by at least some mutations that have minimal effects on SARS-CoV-2 RBD functional properties (**Fig 3A**; see the volume of dark-colored letters, though positions in both epitopes may experience further constraint beyond these isolated-RBD measurements due to quaternary packing in full spike trimers). As previously described, C68.61 mAb escape mutations included sites K462, E465, R466, and I468, which are conserved sites across all SARS-CoV-2 variants and some sarbecoviruses (**Fig 4B**).

**Fig 4.**
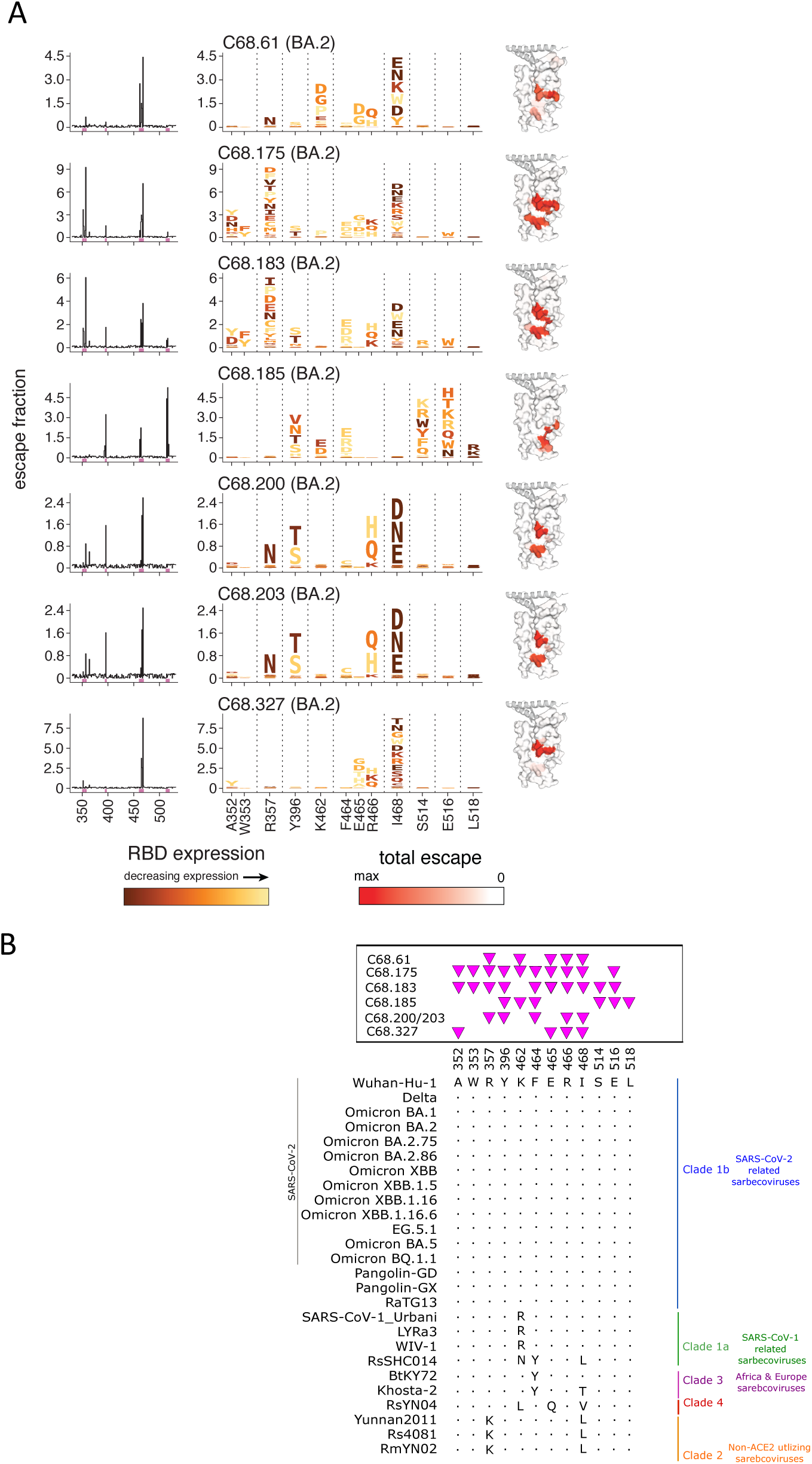
C68 mAbs target class 5 epitopes that are relatively invariant regions on the Spike Protein. **(A)** Complete escape maps of antibody-binding escape mutations for C68 mAbs using a yeast-displayed SARS-CoV-2 RBD deep mutational scanning system (Omicron BA.2 RBD DMS for all class 5 mAbs). Residue colors are assigned based on effect these mutations have on RBD expression. The height of letters in the logo plots indicate escape scores. Logo plot residue numbering is based on SARS-CoV-2 Omicron BA.2. Experiments were performed in biological duplicate using independent mutant RBD libraries (14) that correlate well (**S9B Fig**) so escape fractions represent the average of these two independent biological replicates. On the right, sites of escape for C68 mAbs are mapped onto the RBD structure and gradient of red coloring indicates magnitude of escape fraction at each indicated site. **(B)** Multiple sequence alignments of select SARS-CoV-2 variants and sarbecoviruses are shown corresponding to sites of escape. Residue numbering based on SARS-CoV-2 Wuhan-Hu-1 sequence with shared amino acids denoted by dots (.) and differences across sarbecoviruses relative to the reference sequence (SARS-CoV-2 Wuhan-Hu-1) indicated. Pink triangles depict the sites of escape identified by the escape maps (**Fig 3A**).

The key C68.185 mAb-escape mutations included sites I396, K462, F464, S514, E516, and L518, and, with the exception of K462, are all highly conserved among all sarbecoviruses (**Fig 4A-B**). Inspection of the logo plots of mutational effects indicates that these mutations are largely not tolerated as they would be deleterious for SARS-CoV-2 RBD expression. In some instances where we observed substitutions, there was a differential effect on neutralization in different sarbecoviruses. SARS-CoV-1-related sarbecoviruses encode an arginine at position 462 as opposed to lysine in SARS-CoV-2, and we observed increased binding affinity (**S4A-B Fig**) and neutralization potency (**Fig 2B**) to the SARS-CoV-1 viruses compared to SARS-CoV-2 viruses, suggesting that the arginine at this position may be preferred for C68.185 binding. The K462N mutation found in RsHC014 may account for a loss in neutralization activity as C68.185 bound RsSHC014 RBD ∼10-fold lower than SARS-CoV-1 RBD (**Fig 2C; S4 Fig**) and studies have suggested that SARS-CoV-2 RBD binding is correlated with neutralization potency (61–64). Overall, we see that these core-RBD-directed mAbs target epitopes that are mutationally constrained with respect to RBD expression and are highly conserved across the sarbecovirus family.

We additionally chose to epitope profile a selection of C68 mAbs that broadly bind SARS-CoV-2 variants but not SARS-CoV-1 spike trimer. We selected 2 additional mAbs to epitope profile, C68.10 and C68.201, which share the heavy chain variable gene usage with C68.61 and retain XBB spike trimer binding activity (**Fig 1C**). Interestingly, these mAbs are not clonally related (do not share V(D)J gene rearrangement) to C68.61, yet the key alterations in the spike protein that facilitate escape for these mAbs are centered around K462 similar to C68.61 (**S7A Fig**). These sites are mutationally intolerant as substitutions would functionally constrain RBD expression (**Fig 4B; S7A Fig**). Because these mAbs are targeting the RBD core, we wanted to assess their pan-sarbecovirus RBD binding breadth. Despite overlapping escape maps from SARS-CoV-2, these mAbs display different binding profiles across sarbecoviruses. Both mAbs exhibit limited SARS-CoV-1-related protein reactivity (**Fig 1C; S7B Fig**) but C68.201 can additionally bind the sequence divergent clade 2 and clade 3 sarbecoviruses.

### Resilience and exceptional mutation tolerance of C68 mAbs

To further assess the potential for escape from C68 cross-reactive mAbs, we employed a replication competent recombinant vesicular stomatitis virus system (rVSV) where viruses bearing spike in the presence of antibody select for antibody escape mutants (65,66). rVSV with either SARS-CoV-2 or SARS-CoV-1 spikes were cultured in the presence of antibody to select for mutations, which were identified by next-generation sequencing. C68.185 drove selection for a single substitution at site L504R (SARS-CoV-1 numbering) in rVSV-SARS-CoV-1 at 10 μg/mL (**Fig 5A**), which is a site and residue that led to loss of antibody binding in DMS experiments (**Fig 4**; L518R in SARS-CoV-2 WH-1 numbering). At a lower concentration (1 μg/mL), selection occurred at additional sites centered at Y383H, R449G and Q523R (SARS-CoV-1 numbering). Two of these escape mutations were also identified by the SARS-CoV-2 DMS (Y396, K462 in SARS-CoV-2 numbering). The third substitution identified is outside of the RBD and thus not selected for in the RBD DMS experiments. For this reason, we cannot determine if this mutation directly affects mAb binding or it was selected for other stochastic reasons, or is a stochastic ‘passenger’ mutation in this single culture (67,68).

**Fig 5.**
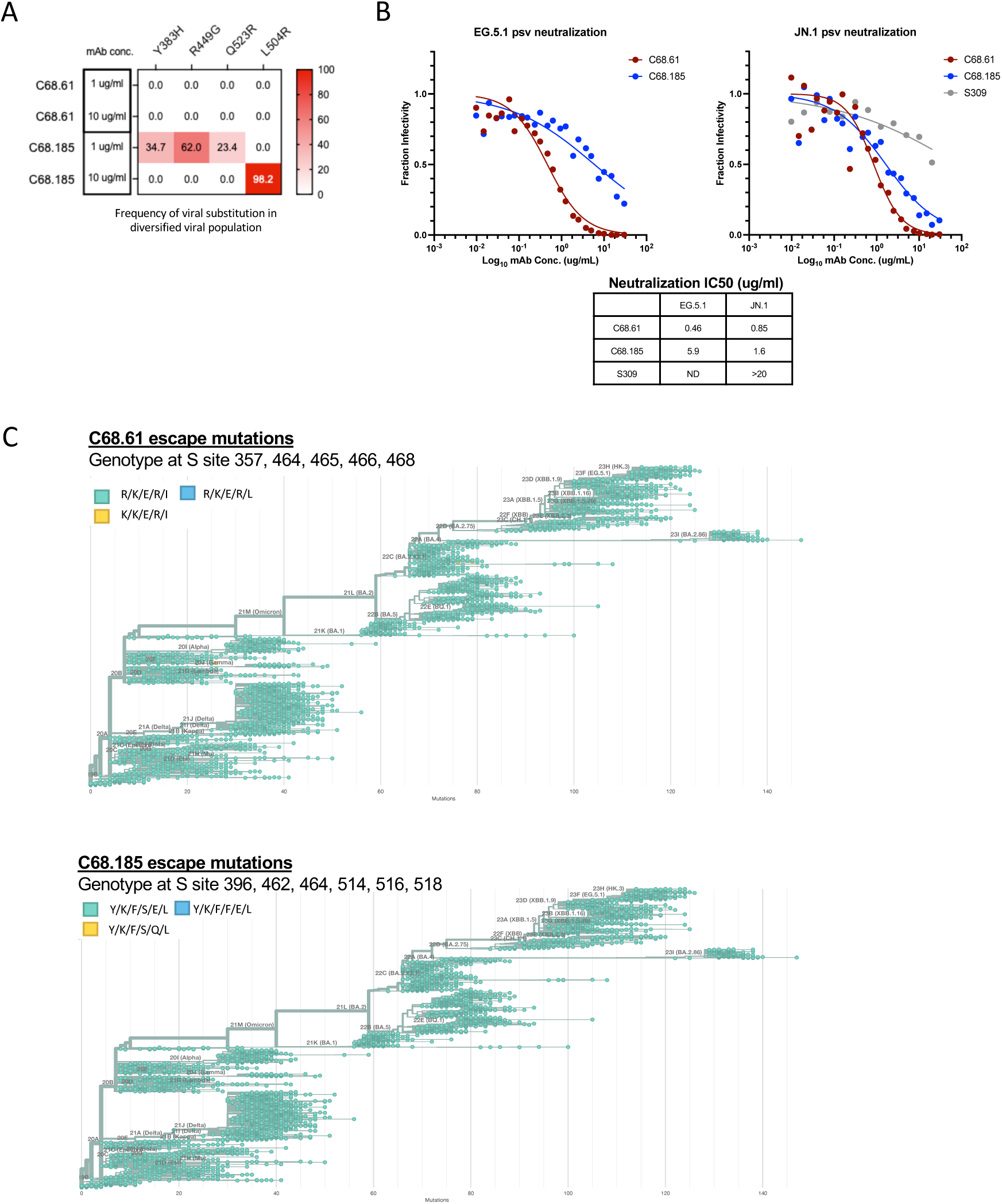
Selection experiments to identify escape mutation in the presence of SARS-Cov-1 and SARS-CoV-2 encoding VSV in cell culture and SARS-CoV-2 evolution observed in nature. **(A)** C68 mAbs selection experiments in the context of SARS-CoV-1 replication competent VSV-G. Frequency of variants in viral population in the presence of C68 mAbs (variant defined as occurring at a frequency greater than 3%). **(B)** Neutralization activity of C68 mAbs (C68.61 in red and C68.185 in blue) and S309 (in grey) against recently circulating and dominant SARS-CoV-2 variant spike-pseudotyped lentiviruses. Antibodies were two-fold serially diluted and tested starting at 20 and/or 30 μg/mL. Samples with IC50 values above the tested concentration were plotted at that concentration (>20 μg/mL is no activity). **(C)** Phylogeny of SARS-CoV-2 Pangolin clade sequences containing C68 mAb RBD binding escape mutations (from Figure 4A) (Nextstrain) for C68.61 (**top panel**) and C68.185 (**bottom panel**). Circles in teal are the residues found in SARS-CoV-2 Wuhan-Hu-1 whereas viral genomes containing escape mutations are indicated by yellow/blue circles. Depicts 3900 SARS-CoV-2 viral genomes that are sampled between December 2019 and March 2024. (Nextstrain: https://nextstrain.org, CC-BY-4.0 license)

Remarkably, C68.61 did not drive escape in this replicating virus system bearing SARS-CoV-1 or SARS-CoV-2 spikes (**Fig 5A**). C68.61 was then tested against three SARS-CoV-2 variants spikes (WH-1, Omicron BA.2 and XBB.1.5) that it neutralized in single-cycle infection assays. This included a more recent Omicron variant XBB.1.5, which was tested because previous DMS and replication competent VSV selection studies suggest there may broadening of pathways of escape with later SARS-CoV-2 variants (65,69). Thus, while there are amino acid mutations that are predicted to disrupt C68.61 binding to RBD (by DMS experiments), viruses with these substitutions were not selected in a replicating virus system, suggesting they may be functionally constrained sites.

Given the lack of predicted escape in cell culture among SARS-CoV-2 variants, we next evaluated the ability of two C68 broadly cross-reactive mAbs to neutralize the most recent dominant circulating variants, JN.1 and EG.5.1 (70). C68.185 exhibited moderate neutralization activity against both SARS-CoV-2 variants (EG.5.1 IC50s of 5.9 μg/mL and JN.1 IC50s of 1.6 μg/mL; **Fig 5B**). C68.61 retained potent neutralization activity at <1 μg/mL against both viruses with minimal fold differences in activity compared to SARS-CoV-2 WH-1 neutralization, emphasizing this antigenic site is highly conserved. We also observed dramatic losses in neutralization activity by S309 against JN.1 (IC50s >20 μg/mL), as has been observed by other studies (71), suggesting this antigenic interface is sequence diverse (72).

### Antibody-escape mutations are present at a low frequency in SARS-CoV-2 evolution

To examine antigenic variation in SARS-CoV-2 spike residues that disrupt antibody binding, we used the bioinformatics tool Nextstrain to assess mutation frequency of SARS-CoV-2 spike residues (73,74). This tool has been used throughout the pandemic to understand SARS-CoV-2 viral evolution by sampling temporal viral sequences from diverse geographic locations (**Fig 5C, top panel**; **S8 Fig**; 3900 genomes). Viral sequences that contained the C68.61 RBD binding escape mutations at sites 357, 462, 465, 466, and 468 amounted to low viral diversity with substitutions accounting for < 2% frequencies at a given time. For instance, a R357K substitution occurred but did not increase in frequency, further suggesting that this mutation may have a fitness disadvantage. We observed a similar lack of viral diversity in these regions for C68.185 (**Fig 5C, bottom panel; S8 Fig**). These epitope targets are not sites that are commonly mutated throughout SARS-CoV-2 evolution (42,75,76) have also not been observed in wastewater sequencing where “cryptic” mutated lineages have been found (77,78) demonstrating that the C68.61 and C68.185 epitope has remained relatively unchanged.

## DISCUSSION

Here, we have identified a collection of cross-reactive mAbs from a Delta breakthrough infection that display cross-reactive neutralizing activity against diverse sarbecoviruses with high spillover risk. Some mAbs displayed potent neutralization (half maximal inhibitory concentration <0.05 μg/mL) against SARS-CoV-1 as well as more divergent animal sarbecoviruses from Africa and Europe. Notably, many of the isolated mAbs demonstrated ADCC activity against SARS-CoV-2 spike and two mAbs exhibited activity on par with another cross-reactive mAb, S309, which has been shown to provide *in vivo* protective efficacy due to its neutralizing and non-neutralizing effector functions (37). One mAb, C68.61, showed unattenuated neutralization against all SARS-CoV-2 variants and several SARS-CoV-1 related sarbecoviruses as well as a more divergent clade 3 bat sarbecovirus. Remarkably, C68.61 did not select for escape mutants in the presence of replication competent virus and SARS-CoV-2 strains harboring C68.61 escape mutations have not increased in prevalence in nature indicating constraints on escape from this mAb. While deep mutational scanning indicates that all these mAbs target one of two regions in the RBD core, previously defined as class 4 and 5 epitopes (27,70), there is heterogeneity in the escape pathways even among mAbs targeting a similar region.

While there was some overlap in the escape mutations among class 4 mAbs, each mAb had a unique escape and neutralization profile. For instance, C68.88 exhibits a narrowly focused epitope that partially overlaps that of C68.239, yet it shows greater cross-reactivity. While these mAbs target epitopes distal to the ACE2 binding site, these antibodies exhibit potent neutralization of a subset of SARS-CoV-2 and SARS-CoV-1 variants. Potent neutralization has been observed by other class 4 mAbs whose angle of approach partially blocks ACE2 access as well as induces S1 shedding (28,79). Differences in gene usage, antigen affinity and angle of approach could also contribute to differential functional activities including C68.239’s ability to mediate potent ADCC against SARS-CoV-2. While class 4 targeting mAbs were posited as potential SARS-CoV-2 broadly neutralizing antibodies (28,52,80), we and others have shown that these mAbs lose activity against SARS-CoV-2 Omicron variants (21,79). Still, their epitope is conserved across SARS-CoV-1-related and ACE2-independent sarbecoviruses tested here. If these findings extend to a larger virus panel, these mAbs could hold promise as broad-spectrum antibodies against potential spillover sarbecoviruses (26,28,79).

The second cluster of C68 mAbs recognize a highly conserved cryptic epitope in RBD designated class 5. While few class 5 mAbs have been comprehensively characterized, its known that these mAbs target a conserved quaternary epitope that is accessible when spike has one-RBD-up open conformation, while occupying the space of the N-terminal domain in an adjacent protomer (25,27,30,81,82). Upon binding, tight packing of RBD-mAb interactions induces further opening of the RBD protomer (25,82) leading to conformational changes that induce S1 shedding. Many of the identified escape mutations were predicted to be functionally intolerant, which is in agreement with published studies (21,30,31,81), and may also be structurally intolerant as these residues are involved in RBD-NTD interactions (27). DMS experiments suggested that C68 mAbs bind to a similar cryptic antigenic region on the spike glycoprotein shared with the class 5 antibody S2H97, which has been shown to reduce viral load in animal studies when given prior to exposure (25). Additionally, some C68 class 5 mAbs displayed cross-reactivity with SARS-CoV-1-related sarbecoviruses and can also neutralize the more divergent African and European animal sarbecoviruses (25,32). This breadth in sarbecovirus activity is exemplified by C68.185, which shows notable binding activity against a diverse array of sarbecovirus RBDs, as well as neutralization of clade 1a and clade 3 ACE2-dependent sarbecoviruses, suggesting the sequence of this epitope is highly conserved among members of these viral clades. However, neutralization is weak against SARS-CoV-2 variants, perhaps because the mechanism of neutralization for this class of mAbs lacks ACE2 blocking.

While binding to spike trimer and high sequence conservation of the binding residues were largely predictive of neutralization activity, this was not always the case. For example, four of twelve mAbs that were cross reactive based on binding to SARS-CoV-1 spike trimer did not neutralize the corresponding virus. This discrepancy in binding and neutralization could be due to differences in RBD dynamics between sarbecoviruses, as has been suggested for the class 4 mAb CR3022, which binds the one-RBD-up open conformation in SARS-CoV-1 but would sterically clash with a neighboring RBD in SARS-CoV-2 (50). This may explain the ability of mAb C68.121 to bind but not neutralize sarbecoviruses, since it overlaps in contact residues with CR3022. While SARS-CoV-2, SARS-CoV-1 and other sarbecoviruses require one RBD in an “up” conformation to bind host cell receptor (5,83,84), studies have indicated that SARS-CoV-2 variants also alter RBD conformational dynamics (85–87). This could indirectly affect binding affinity and neutralization of mAbs whose epitopes are highly sequence conserved but may be occluded in the closed RBD conformation (88), such as C68.175 and C68.203. Another explanation for this differential effect in neutralization despite sequence conservation across sarbecoviruses may be due to epistatic shifts as the accumulation of mutations outside the antibody binding site may lead to shifts in the effect of residues on antibody function, which has been observed in other viral contexts (89,90) as well as in SARS-CoV-2 variants (65,69). While we performed our DMS experiments in the genetic background of the SARS-CoV-2 WH-1 or Omicron BA.2, the effects of mutations may differ in different viral contexts (15,65).

SARS-CoV-2 evolution is likely shaped by both mutation and positive selection to evade immunity, yet the capacity to escape antibody recognition can be limited by selection to maintain functional capabilities (41). To examine this, we used a replicating virus system to select for escape for two of the mAbs that neutralized SARS-CoV-1 and SARS-CoV-2 and show activity against more diverse sarbecoviruses. In the case of C68.185, which neutralized several clade 1a and clade 3 sarbecoviruses, escape mutants against SARS-CoV-1 virus were selected. Some of the selected mutations are highly sequence conserved across sarbecoviruses including SARS-CoV-2 emerging variants. Remarkably, there was no evidence of escape from C68.61 for either SARS-CoV-1 or SARS-CoV-2 in this same assay, including different SARS-CoV-2 variants. In agreement with this finding, SARS-CoV-2 variants that contain these spike substitutions occur at very low frequency in nature and there is no evidence of expansion of these rare mutants. Thus, mAbs like C68.61 that can functionally constrain sarbecovirus spike evolution could lead to broadly protective interventions for current and emerging SARS-CoV-1 and SARS-CoV-2 variants and inform pan-sarbecovirus vaccine strategies.

This study identified new antibodies with a range of cross-reactivity that target diverse epitopes within the core of RBD, including functionally constrained epitopes. Several of these mAbs have complementary neutralization and DMS epitope profiles that suggest they have minimal or non-overlapping escape mutations. Thus, a combination of these mAbs could form the basis for an antibody cocktail that would show activity across very diverse sarbecoviruses to be used in the event of another pandemic (91,92).

## MATERIALS AND METHODS

### Human PBMC specimen

Peripheral blood mononuclear cell (PBMC) sample was collected at 30 days after a breakthrough infection from an individual, C68, who received two doses of the Pfizer-BioNTech mRNA vaccine and was subsequently infected with the Delta variant. C68 was enrolled in the study and followed prospectively as part of the Hospitalized or Ambulatory Adults with Respiratory Viral Infections cohort (study approved by the University of Washington Protocol #STUDY00000959 and Fred Hutch Cancer Center Institutional Review Boards).

### SARS-CoV-2 memory B cell isolation and antibody reconstruction

The methods of isolation of SARS-CoV-2 memory B cells from C68 were previously described (31,93). In brief, cells were incubated with a cocktail of anti-CD3-BV711 (BD Biosciences, clone UCHT1), anit-CD14-BV711 (BD Biosciences, clone MφP9), anti-CD16-BV711 (BD Biosciences, clone 3G8), anti-CD19-BV510 (BD Biosciences, clone SJ25C1), anti-IgM-FITC (BD Biosciences, clone G20-127), anti-IgD-FITC (BD Biosciences, clone IA6-2) along with APC/PE-labeled Delta HexaPro spike protein (a generous gift from David Veesler) and spike S2 peptide (Acro Biosystems, cat. S2N-C52E8) for 30 minutes on ice. Single B cells defined as CD3^−^, CD14^−^, CD16^−^, CD19^+^, IgD^−^, IgM^−^, PE^+^, APC^+^ and live cells (Ghost Dye Red 780; Tonbo Biosciences cat: 13-0865-T500) were sorted (BD Biosciences, FACS AriaII) into a total of four 96-well PCR plates containing RNA storage buffer and stored at −80C. RNA storage buffer consists of RNAse OUT, 5X first-strand buffer, 0.1 M DTT, Igepal, carrier RNA, ultrapure water.

A total of 384 SARS-CoV-2 Delta trimer and/or S2 subunit specific B cells were recovered from C68. Reconstruction of antibodies was conducted as previously described (31,93–96). RT-PCR amplification of IgG antibody heavy and light chain variable regions was performed followed by nested chain-specific amplification PCRs (31,94,97). Two different primer sets were used to maximize recovery of antibody chain DNA (98,99). Amplicons were sequenced and the sequences were initially assessed for the presence of an intact open-reading frame (in-frame junction and no stop codons) by comparing the sequences to IMGT V-QUEST (100,101). Functional sequences were then subcloned into their respective IgK or IgL expression vectors and all heavy chain were expressed as IgG1 (95,96). To determine levels of somatic hypermutation, clonality, and germline inference were performed with the partis software package (https://github.com/psathyrella/partis/ and https://arxiv.org/abs/2203.11367). Of the 384 B cells recovered, 118 had paired functional heavy and light chain sequences and of these 45 (42 mAbs characterized here and 3 mAbs in (31)) were found to be RBD specific based on binding to the RBD subunit. The remaining mAbs have been characterized here (31,93) and were found to be NTD (17), CTD (3), S2 (29) or not binding reactive to SARS-CoV-2 antigen.

### Antibody binding profiling by enzyme-linked immunosorbent assays

Antibody binding to SARS-CoV-2 or SARS-CoV-1 recombinant spike protein was performed using ELISA as previously shown (31,102). Briefly, 384-well maxisorp plates (Nunc, Thermo Fisher Scientific) were coated with SARS-CoV-2 or SARS-CoV-1 antigen 25 uL at 1 µg/ml in 1x phosphate buffered saline (PBS) coating buffer overnight at 4°C. The recombinant spike protein included: Wuhan-Hu-1 (Sino Biologics; cat. 40589-V08H4), Delta (Sino Biologics; cat. 40589-V08H10), Omicron BA.1 (Sino Biologics; cat. 40589-V08H26), Omicron BA.2 (Sino Biologics; cat. 40589-V08H28), Omicron BA.4/BA.5 (Sino Biologics; cat. 40589-V08H32), Omicron XBB (Sino Biologics; cat. 40589-V08H40), Omicron BQ.1.1 (Sino Biologics; cat. 40589-V08H41), and SARS-CoV-1 (Acro Biosystems, CUHK-W1 strain) as well as the SARS-CoV-2 RBD subunit (Sino Biologics; cat. 40592-V08H). The following day, plates were washed 4x with 100 uL of PBS-T (1x PBS, 0.05% Tween-20) using a plate washer. Plates were then blocked in 50 uL of 3% non-fat dry milk in PBS-T for 1 hour at ambient temperature. Blocking buffer was removed and plates were incubated with 25uL monoclonal antibody dilutions for 1 hour at 37 °C. Plates were then washed and incubated with HRP-conjugated goat anti-human IgG antibody (Sigma-Aldrich) (diluted 1:2500 in PBS-T) for 1 hour at ambient temperature. Plates were washed and incubated with TMB substrate (Thermo Fisher Scientific) for ∼3-5 minutes until the reaction was quenched with 1N sulfuric acid. Finally, plates were read at absorption 450 nm (BMG Labtech). For ELISAs that were used to screen for SARS-CoV-2 specificity (**Fig 1A**), the binding assay was performed at a single antibody dilution (500 ng/ml) and background in the 1x PBS control was subtracted to determine binding activity (OD450). Results presented were the average of technical duplicates. For experiments to determine the 50% effective concentration (EC50), ELISAs were performed at a starting concentration of 1000 ng/ml antibody and serially diluted 2-fold for at least 12 dilutions. OD450 nm values minus background from 1x PBS only wells were averaged from technical duplicates and the final results presented are the average of two biological replicates in technical duplicate. EC50 was calculated using GraphPad Prism software (v10) by fitting a four-parameter (agonist vs response) nonlinear regression curve with the bottom fixed at 0 and the top constrained to the highest OD450 nm value observed.

### Spike-pseudotyped lentiviral particle production and neutralization assays

Spike-pseudotyped lentiviral particle production was performed in 293T cells expressing high levels of ACE2 as previously described (31,102–104). Sarbecovirus spike plasmids with spike genes specific for SARS-CoV-2 variants WH-1, Delta, Omicron BA.1, Omicron BA.2, Omicron BA.4/BA.5, as well as SARS-CoV-1 (Urbani strain), BtKY72 (K493Y/T498W mutant), Khosta-2, WIV1, LYRa3, Pangolin-GD, RsSHC014 were provided as follows: BA.4/5, XBB.1.5, BQ.1.1, EG.5.1, JN.1 and SARS-CoV-1 were generated as previously described (105); WH-1, BA.1, BA.2 were provided by Jesse Bloom (103,104); Delta was a gift from Amit Sharma; BtKY72, Khosta-2, WIV1, LYRa3, Pangolin-GD, RsSHC014 were gifts from Pamela Bjorkman (96). Supernatant from transfected 293T cells was harvested 72-hour post-transfection and sterile filtered through a 0.2-μm filter and concentrated using Amicon Ultra Centrifugal Filters (Sigma-Aldrich). Pseudovirus was then used to infect 293T cells constitutively expressing human ACE2 (generously provided by Jesse Bloom) (103), and viral titer was determined by measuring luminescence using Bright-Glo Luciferase Assay System (Promega) at 48 hours post-infection.

Spike-pseudotyped neutralization assays was performed as previously described (31,102,103). Neutralization assays were carried out by adding pseudotyped lentiviral particles to obtain 2-5 × 10^5^ relative luciferase units (RLUs) of luminescence signal per well to an equal volume of serially diluted antibody sample. To determine the inhibitory concentration at 50% (IC50), antibody dilutions were prepared at a starting concentration of 20 or 30 μg/mL and two-fold mAb dilution curves were assessed in a 384-well format. Antibody-pseudovirus mix were then added to a pre-seeded plate of 293T-ACE2 cells. Plates were read two days later by measuring luminescence using Bright-Glo Luciferase Assay System (Promega). Background values were subtracted using the “cells only” condition as background signal. Percent neutralization was then calculated as a reduction in RLUs of each antibody-pseudovirus mix compared to cells infected with virus in the absence of antibody (cells and virus only). If the “cells and virus only” condition was 1.3x higher than the average luminescence signal from the non-neutralizing wells (highest diluted mAb plus virus condition), then the non-neutralizing wells were defined as the background signal to minimize potential variability due to edge effects. Spike-pseudotyped neutralization assays were done in technical duplicate and replicate experiments were averaged, and the fraction of infectivity was calculated. We calculated the IC50 using GraphPad Prism software by fitting a four-parameter (agonist vs response) nonlinear regression curve with the bottom fixed at 0, the top constrained to 1 and HillSlope > 1.

### SARS-CoV-2 spike antibody-dependent cellular cytotoxicity assay

To measure Fc-mediated function of C68 mAbs, we adapted a validated SARS-CoV-2 specific ADCC assay (56,57,93,106). CEM.NKr cells stably expressing a full length GFP-tagged SARS-CoV-2 spike (variant D614G; target cells) were generously provided by Andrés Finzi. Cryopreserved PBMCs (effector cells; acquired from Bloodworks Northwest or kindly given from Andrés Finzi) from healthy donors were thawed and rested overnight in RPMI (Gibco) supplemented with 10% FBS (Gibco), 4.0mM Glutamax (Gibco) and 1% antibiotic-antimycotic (Life Technologies). The following day, effector cells were mixed at a 10:1 ratio with target cells with C68 mAbs in a 96-well v-bottom non-TC treated plates. All C68 monoclonal antibodies were diluted in 1x PBS and tested at 5000ng/ml. This concentration of 5000ng/ml was chosen because it was the highest activity observed before reaching the prozone effect based on previous studies (57) and in-house experiments comparing ADCC activity at different dilutions (not shown).

Effector cells, target cells and antibodies were mixed and incubated with anti-CD107a-APC (clone H4A3, BioLegend) and protein transport inhibitor cocktail (1:500 dilution in 1x PBS, eBioscience). The plates were subsequently centrifuged for 1 min at 100xg, and incubated at 37°C, 5% CO_2_ for 4 hrs. After incubation, cells were washed and centrifuged at 400xg for 3 minutes. Cells were stained with an antibody cocktail consisting of: viability dye (Viability Dye eFluor™ 780; Thermo Fisher Scientific), anti-CD3-BV711 (Clone UCHT1, Biolegend), anti-CD56-BV605 (clone 5.1H11, BioLegend), and anti-CD16-BUV395 (clone 3G8, BD Horizon) on ice for 30 minutes. Cells were washed, fixed and permeabilized (eBioscience™ Intracellular Fixation & Permeabilization Buffer Set) before staining for intracellular IFN-γ-PE (clone 4S.B3, BioLegend) for 20 minutes at room temperature. Cells were acquired on a Fortessa LSR instrument (Fortessa X50, BD Biosciences). ADCC was measured as the percentage of NK cells (defined as live, singlets, GFP-, CD3-, CD56+ cells) positive for CD107a or intracellular IFN-γ (gating strategy shown in S5 Fig.). Target cell death was measured as percentage of spike expressing target cells positive for cell death marker (gating strategy shown S5 Fig.). Data analysis was performed using FlowJo v10.7.1. Each experiment was performed in technical duplicates and with two independent PBMC donors. Experimental values shown are background subtracted (background defined as spike expressing target cells and effector cells in the absence of antibodies).

### Epitope mapping and escape profiling by yeast display RBD DMS

To determine the epitope targets and the escape profiles of mAbs, we used a yeast library displaying RBD proteins with nearly every single amino acid mutation in the RBD of Wuhan-Hu-1 or Omicron BA.2, as previously described (25,31,59). First, a yeast strain expressing the unmutated RBD of Wuhan-Hu-1 or Omicron BA.2 and flow cytometry were used to identify an EC90 concentration for antibody binding to the wildtype yeast-displayed RBD construct. Then, approximately 5 OD units of yeast libraries were incubated with that EC90 concentration of mAb. Cells were washed and incubated with 1:200 PE-conjugated goat anti-human-IgG antibody (Jackson ImmunoResearch; clone 109-115-098), and 1:100 FITC-conjugated chicken anti-Myc-tag (Immunology Consultants Lab; clone CYMC-45F). A FACS-based approach was used to identify antibody escape mutants by gating on yeast mutants with reduced antibody binding, with gates drawn on wildtype control cells labeled at 10% of the library selection mAb concentration to approximate an escape bin of 10x or greater loss of binding (**S9 Fig**). Approximately, 4 million RBD+ cells were sorted on a FACS AriaII.

Pre-sort and sorted antibody-escape cells were sequenced using an Illumina NovaSeq and a 16-nucleotide barcode linked to each RBD mutant was used to identify the mutant sequence. Raw sequence reads are available on NCBI Sequence Read Archive (SRA), BioProject PRJNA770094, BioSample SAMN40905401. Sequencing reads were then compiled and compared to the pre-sort population frequencies to generate the “escape fraction” visualized on logoplots. Experiments were performed in biological duplicate using independent mutant RBD libraries (14) that correlate well (**S9B Fig**) so escape fractions represent the average of these two independent biological replicates. Wuhan-Hu-1 RBD library backgrounds (used to profile C68.88 and C68.239) and Omicron BA.2 (used to profile C68.61, C68.83, C68.121, C68.175, C68.183, C68.185, C68.200, C68.203, C68.327, C68.348) were chosen based on initial binding experiments assessing ability to bind either Wuhan-Hu-1 RBD or Omicron BA.2 RBD expressed on the surface of yeast (not shown but aligns with ELISA binding experiments, Fig 2A). Final escape fraction measurements averaged across two replicates are available from GitHub: https://github.com/tstarrlab/SARS-CoV-2-RBD_Omicron_MAP_Overbaugh_v2/tree/main/results/supp_data. The entire pipeline for epitope escape profiling is available from GitHub: https://github.com/tstarrlab/SARS-CoV-2-RBD_Omicron_MAP_Overbaugh_v2.

### Antibody selection experiments using replication competent VSV expressing sarbecovirus spike

To assess what escape mutants arise in the presence of C68 mAbs in the context of virus replication, we used a recombinant replication-competent vesicular stomatitis virus (rVSV) expressing the SARS-CoV-2 variants (rVSV/SARS-CoV-2/GFP) or rVSV/SARS-CoV-1/GFP spike as previously described (60,65). Generation and titering of rVSV/SARS-CoV-2/GFP chimeric viruses using 293T cells expressing ACE2 has been previously described (65,66).

Antibodies were incubated with rVSV at 1 million infectious units and tested at either 1 or 10 µg/mL with concentration determined by potency of a given mAb. The antibody-virus mix was incubated for 1 hour at 37°C before infecting ACE2 expressing 293Ts (Passage #1). Twenty-four hours later, supernatant was removed and replenished with fresh media containing mAb at the same antibody concentration. After an additional 24 hours, 100 uL of the virus-containing supernatant was filtered and incubated with antibody at the same concentration for 1 hour at 37 C. This virus-antibody mixture was then used to inoculate fresh 293T-ACE2 cells for a second passage with the above passaging steps repeated. After two passages, viral supernatant was harvested and viral RNA was extracted. RT-PCR was used to amplify the RBD and RBD amplicons were then tagged for sequencing analysis using Nextera TDE1 Tagment enzyme (Illumina) and then i5/i7 barcoded primers were added to tagmented cDNA using Illumina Nextera XT Index Kit v2 and KAPA HiFi HotStart ReadyMix (Roche) to barcode amplicons. The barcoded library was then pooled and sequenced. Sequencing reads were filtered and compared to the parental spike sequence with mutation counts compiled. A mutation was defined as a true variant if substitution occurred at a frequency greater than 3%.

### Yeast display of sarbecovirus RBDs binding assays

To evaluate the ability to bind a panel of sarbecovirus RBDs, we used yeast libraries displaying RBDs from sarbecoviruses representing different clades as previously described (11,25). A full list of all RBDs and sequence accession numbers is available on GitHub: https://github.com/tstarrlab/SARSr-CoV_mAb-breadth_Overbaugh/blob/main/data/data-supp_RBD-sequences.csv. Briefly, sarbecovirus clades were phylogenetically defined based on RBD sequence, with the Hibecovirus RBD sequence (GenBank: KF636752) used to root the sarbecovirus phylogeny tree. Each sarbecovirus RBD is represented by >100 barcodes such that technical pseudo-replicates can be ascertained within each binding experiment. Antibodies were incubated with sarbecovirus libraries at five concentration points (10,000 ng/mL and four serial 25-fold dilutions to 0.0256 ng/mL; plus zero mAb condition). Approximately 1 OD units of yeast libraries were incubated with serially diluted mAb. Cells were washed and incubated with 1:200 PE-conjugated goat anti-human-IgG antibody (Jackson ImmunoResearch; clone 109-115-098), and 1:100 FITC-conjugated chicken anti-Myc-tag (Immunology Consultants Lab; clone CYMC-45F). At each concentration, approximately 1 million RBD+ cells were sorted and binned representing 4 cell populations of low and high mAb binding. Barcodes were counted within each FACS bin via Illumina sequencing, with raw sequence reads available on NCBI SRA, BioProject PRJNA714677, BioSample SAMN40905419. EC50s were calculated using the sequence data of each serially diluted bin allowing us to visualize approximate differences in affinity between different sarbecovirus RBDs. EC50 scores are the geometric mean across the independent barcodes. Violin plots show distribution of Log10 EC50 values from independent barcodes. The computational pipeline for computing sarbecovirus mAb-binding breadth is available on GitHub: https://github.com/tstarrlab/SARSr-CoV_mAb-breadth_Overbaugh.

## ACKNOWLEDGEMENTS

We thank the Flow Cytometry Core Facility at the University of Utah Health Sciences Campus (supported by NIH 5P30CA042014-24) and the University of Utah Center for High Performance Computing (supported by NIH 1S10OD021644-01A1) for experimental support. We thank Amit Sharma (Ohio State University), Jesse Bloom (Fred Hutchinson Cancer Center), David Veesler (University of Washington) and Pamela Bjorkman (Caltech) for generously providing spike plasmids for the production of spike-pseudotyped lentiviruses. We thank all the researchers who were involved in sequencing and submitting data to GISAID. We also thank David Veesler (University of Washington) for providing Delta spike trimer to use as bait to capture spike-specific B-cells and Andrés Finzi (Université de Montréal) for the CEM.NKr CCR5+ cells. We also would like to thank the participants and the study staff of the Hospitalized or Ambulatory Adults with Respiratory Viral Infections (HAARVI) study.

## SUPPORTING INFORMATION CAPTIONS

**S1 Fig. Antibody heavy and light chain V-gene family characteristics.** Counts for the heavy and light chain V-gene segments among C68 RBD mAbs (related to **Fig 1C**) based on *partis* computational software analysis. Antibody heavy (left panel) and light (right panel) chain amino acid sequences found in **S1 Table**.

**S2 Fig. Binding curves for C68 mAbs to SARS-CoV-2 variants and SARS-CoV-1 recombinant spike glycoprotein by ELISA.** Binding of C68 mAbs to SARS-CoV-2 WH-1, SARS-CoV-2 variants (Delta, Omicron BA.1, Omicron BA.2, Omicron XBB, Omicron BA.4/BA.5, Omicron BQ.1.1) or SARS-CoV-1 recombinant spike trimers. The concentration of each mAb (ng/mL) is plotted on a log10 scale versus absorbance (OD450nm values). Half-maximal effective concentrations (EC_50_ values) calculated by nonlinear regression analysis from at least two independent technical replicates. Each mAb was serially diluted 2-fold for 11-12 total dilutions. Curves represent nonlinear regression fits with error bars that indicate SEM.

**S3 Fig. Sarbecovirus percent sequence identity relative to SARS-CoV-2 or SARS-CoV-1 RBD and neutralization statistics. (A)** Sarbecovirus percent sequence identity to SARS-CoV-2 WH-1 RBD or SARS-CoV-1 RBD. **(B)** Geometric means (geomeans) of the IC50s and 95% confidence interval (95% CI) across SARS-CoV-2 variants only (left panel) and all tested sarbecoviruses (right panel) tested in **Fig 2B**.

**S4 Fig. Violin plots showing Log10 EC50 values from pan-coronavirus RBD yeast display assay. (A)** We assessed the cross-reactivity of C68.61 (upper panel) and C68.185 (lower panel) (data from **Fig 2C**) with 61 RBDs from sarbecovirus clades (indicated by different colors where clade 1b SARS-CoV-2-related sarbecoviruses are in blue, clade 1a SARS-CoV-1-related sarbecoviruses are in green, clade 2 ACE2-independent sarbecoviruses in yellow, clade 3 African/European sarbecoviruses in purple, and clade 4 sarbecoviruses in orange) analyzed by flow cytometry. The lower the EC50 (log10 ng/ml), the stronger binding response. Represents data from at least two independent experiments. (**B**) Geometric means of the EC50s (ng/ml; data from **Fig 2C**) and 95% confidence interval (95% CI) across clade 1b SARS-CoV-2-related sarbecoviruses and clade 1a SARS-CoV-1-related sarbecoviruses tested.

**S5 Fig. Gating strategy for ADCC. (A)** The ability of candidate mAbs to trigger ADCC was determined by flow cytometry. NK cell (gated as live, single cells, CEM.NKr.spike^-^, CD3^-^, CD56^+^ cells) activation was measured by surface expression of CD107a (a proxy for degranulation) and intracellular IFN-γ (93). **(B)** Representative gating strategy for CEM.NKr.spike cell death.

**S6 Fig. Assessment of Fc-mediated functions of C68 mAbs. (A)** Correlation plot of the ability of C68 mAbs and control mAbs (control mAbs: S309, CV3-25, A32, S2H97, S2H97) to trigger NK cell activation (%CD107a+) from two independent PBMC donors using Pearson’s correlation. **(B)** Measurement of the degree of binding to CEM.NKr cells expressing D614G SARS-CoV-2 spike (target cells). Control mAbs are shown in grey and C68 mAbs in pink. **(C)** Correlation plot between the degree of binding to target cells and the ability to trigger NK cell activation (%CD107a+). Pearson’s correlation analysis showed no association between NK cell activation (%CD107a+) and MFI (r = 0.3, p = 0.2). **(D)** Assessment of the ability of C68 mAbs to trigger NK cell activation (%intracellular IFN-γ). **(E)** Correlation between the ability of C68 mAbs to trigger NK cell activation as measured by percent cell surface CD107a or percent intracellular IFN-γ. **(F)** Percent of cell death within the target cell population. **(G)** Correlation plot between target cell death and %CD107a expression. Correlation plots between NK activation measurements (%CD107a, %intracellular IFN-γ, %target cell death) and binding of targets cells by mAbs (MFI) were assessed by Pearson correlation using PRISM. Results represent assays run in technical duplicate with background subtracted values, data shown from a single donor (for B-G).

**S7 Fig. Epitope profiling and pan-sarbecovirus RBD binding activity for C68 mAbs. (A)** Complete escape maps of antibody-binding escape mutations for C68 mAbs using a yeast-displayed SARS-CoV-2 RBD deep mutational scanning system (Omicron BA.2 RBD). Line plots (left panel) show escape at each site in RBD. Residue colors are assigned based on effect these mutations have on RBD expression (with yellow indicating mutations deleterious for RBD expression). The height of letters in the logo plots indicate level of escape by that amino acid at that site. Logo plot residue numbering is based on SARS-CoV-2 Omicron BA.2 **(B)** Pan-sarbecovirus RBD yeast display to assess C68 mAbs sarbecovirus RBD binding breadth. EC50 value represents the geometric mean across the independent barcodes for corresponding sarbecovirus RBD.

**S8 Fig. Frequency of viral genomes containing C68 RBD binding escape mutants.** Area plot indicating the frequency of RBD binding escape mutants (from Fig 4A) found in SARS-CoV-2 sequences (Nextstrain) for C68.61 (**top panel**) and C68.185 (**bottom panel**) over time. Circles in teal are the residues found in the SARS-CoV-2 reference sequence (Wuhan-Hu-1) whereas mutations or deletions are indicated by yellow/blue circles. Depicts 3900 SARS-CoV-2 viral genomes sampled between Dec 2019 and Mar 2024 (**S2 Table**). (Nextstrain: https://nextstrain.org, CC-BY-4.0 license)

**S9 Fig. Yeast displaying SARS-CoV-2 RBD deep mutational scanning library gating strategy and correlation between screens using independently generated libraries.** (A) Gating strategy to identify antibody escape mutants by gating on yeast mutants with reduced antibody binding. A yeast strain expressing the unmutated RBD of Wuhan-Hu-1 or Omicron BA.2 and flow cytometry were used to identify an EC90 and then yeast libraries were incubated with that EC90 concentration of mAb. Cells are incubated with PE-conjugated 1:200 goat anti-human-IgG and 1:100 FITC-conjugated chicken anti-Myc-tag. Cells are then gated on unmutated WH-1 or Omicron BA.2 control cells to identify escape mutants with >10x loss in antibody binding. (B) Correlation of mutation-level escape from independently generated mutant RBD libraries.

**S1 Table. Amino acid sequences for C68 cross-reactive antibodies characterized in this study.**

**S2 Table. SARS-CoV-2 sequence accession IDs and acknowledgements table**. Sequences used by Nextstrain (https://nextstrain.org) to generate **S8 Fig** and **Fig 5C**.

