## Supplemental figures for "Delineating the functional activity of antibodies with cross-reactivity to SARS-CoV-2, SARS-CoV-1 and related sarbecoviruses"

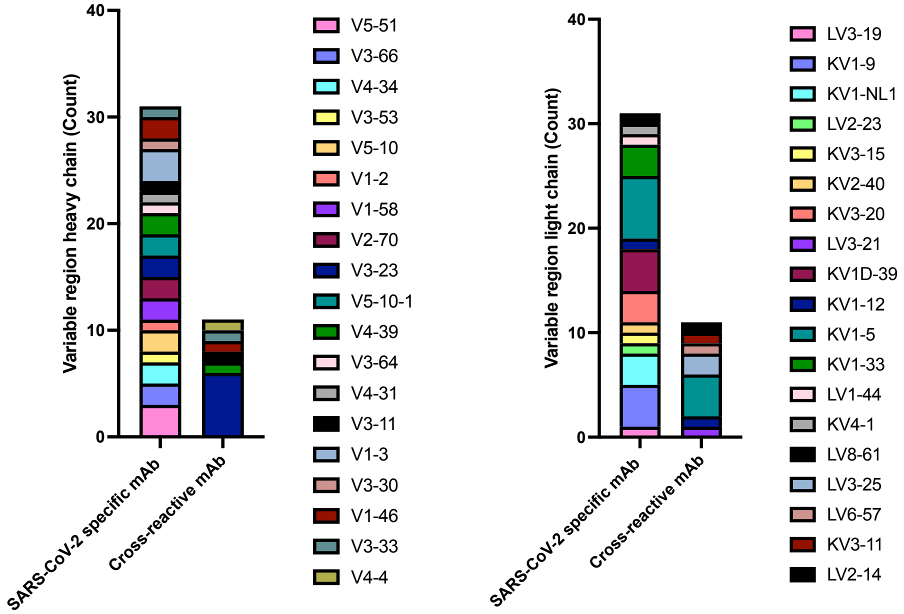
Variable region heavy chain (Count)
40
30
20
10
0
SARS-CoV-2 specific mAb
Cross-reactive mAb
V5-51
V3-66
V4-34
V3-53
V5-10
V1-2
V1-58
V2-70
V3-23
V5-10-1
V4-39
V3-64
V4-31
V3-11
V1-3
V3-30
V1-46
V3-33
V4-4
Variable region light chain (Count)
40
30
20
10
0
SARS-CoV-2 specific mAb
Cross-reactive mAb
LV3-19
KV1-9
KV1-NL1
LV2-23
KV3-15
KV2-40
KV3-20
LV3-21
KV1D-39
KV1-12
KV1-5
KV1-33
LV1-44
KV4-1
LV8-61
LV3-25
LV6-57
KV3-11
LV2-14

### S1 Fig.

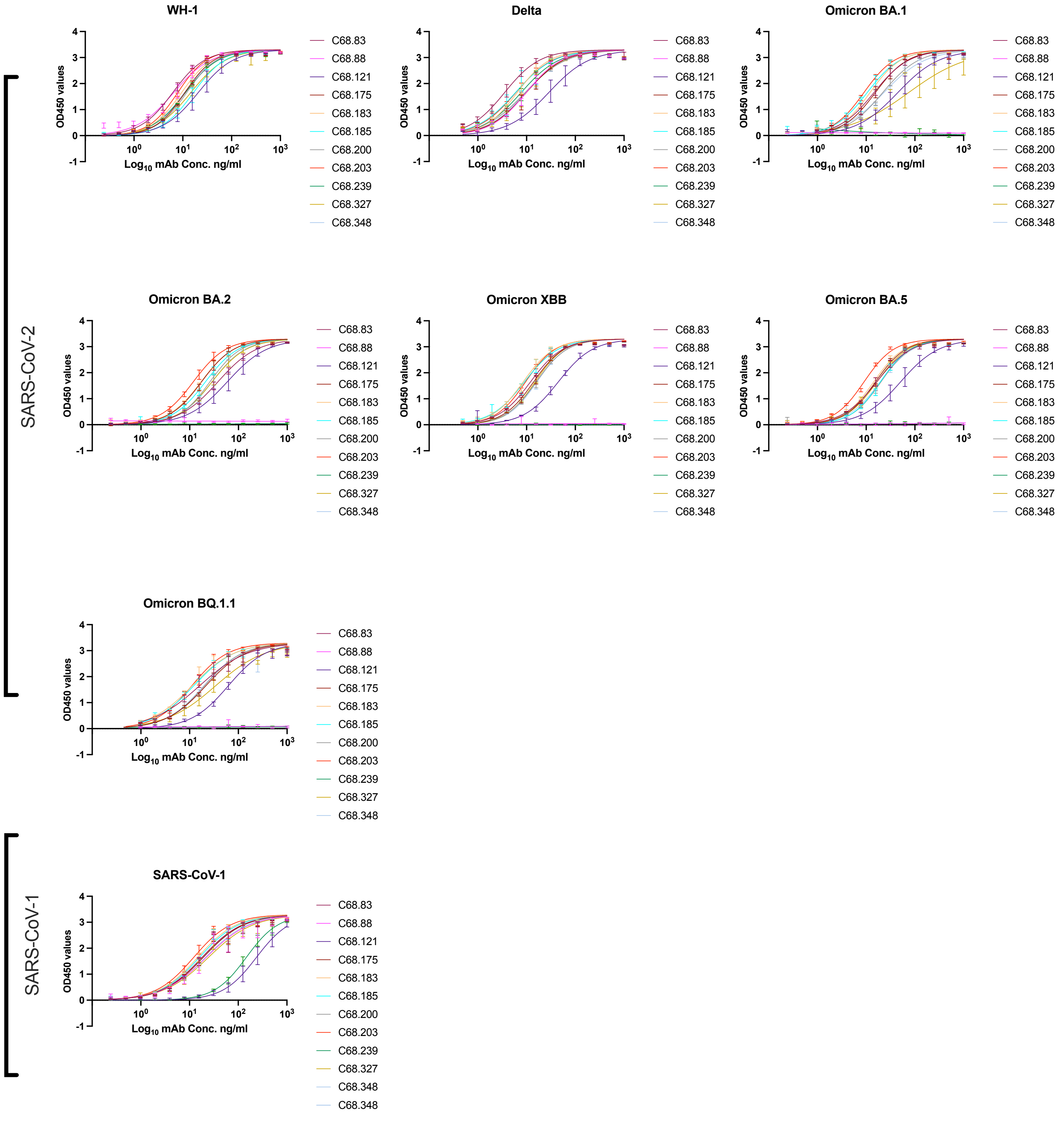
SARS-CoV-2
SARS-CoV-1
WH-1
Delta
Omicron BA.1
Omicron BA.2
Omicron XBB
Omicron BA.5
Omicron BQ.1.1
SARS-CoV-1
OD450 values
Log10 mAb Conc. ng/ml
C68.83
C68.88
C68.121
C68.175
C68.183
C68.185
C68.200
C68.203
C68.239
C68.327
C68.348

### S2 Fig.

## A

| | WH-1 | Delta | Omicron BA.1 | Omicron BA.2 | Omicron XBB.1.5 | Omicron BA.5 | Omicron BQ.1.1 | GD-Pangolin | SARS-CoV-1 | RsSHC014 | WIV1 | LYRa3 | Khosta-2 | BtKY72 |
|---|---|---|---|---|---|---|---|---|---|---|---|---|---|---|
| % Sequence identity to SARS-CoV-2 WH-1 RBD | - | 99 | 93 | 92 | 89 | 92 | 90 | 96 | 74 | 77 | 77 | 75 | 68 | 73 |
| % Sequence identity to SARS-CoV-1 RBD | 74 | 75 | 72 | 71 | 71 | 71 | 72 | 75 | - | 82 | 96 | 95 | 70 | 74 |

## B

| Statistics for SARS-CoV-2 variants neutralization | | |
|---|---|---|
| | Geomean | 95% CI |
| C68.61 | 0.40 | (0.25, 0.64) |
| C68.83 | 8.6 | (2.2, 34) |
| C68.88 | 4.7 | (0.47, 46) |
| C68.121 | 17 | (12, 23) |
| C68.175 | 5.0 | (1.7, 14) |
| C68.183 | 11 | (4.6, 29) |
| C68.185 | 5.8 | (2.6, 13) |
| C68.200 | 18 | (13, 24) |
| C68.203 | 5.3 | (1.6, 18) |
| C68.239 | 4.2 | (0.35, 50) |
| C68.327 | 7.5 | (3.4, 17) |
| C68.348 | 8.0 | (2.2, 29) |
| S309 | 1.4 | (0.3, 6.6) |
| CR3022 | 20 | (20, 20) |
| S2H97 | 0.39 | (0.17, 0.86) |
| S2X259 | 3.8 | (0.43, 34) |

| Statistics for all sarbecovirus neutralization | | |
|---|---|---|
| | Geomean | 95% CI |
| C68.61 | 0.62 | (0.24, 1.6) |
| C68.83 | 7.4 | (3.2, 17) |
| C68.88 | 1.1 | (0.19, 6.4) |
| C68.121 | 13 | (8.4, 21) |
| C68.175 | 5.3 | (1.9, 15) |
| C68.183 | 8.2 | (3, 23) |
| C68.185 | 1.1 | (0.26, 4.3) |
| C68.200 | 11 | (4.4, 25) |
| C68.203 | 3.6 | (1.3, 9.9) |
| C68.239 | 2.7 | (0.52, 14) |
| C68.327 | 7.2 | (2.9, 18) |
| C68.348 | 4.4 | (1.6, 12) |
| S309 | 0.90 | (0.23, 3.5) |
| CR3022 | 8.3 | (3.5, 20) |
| S2H97 | 0.95 | (0.32, 2.8) |
| S2X259 | 0.32 | (0.05, 1.9) |

### S3 Fig.

A

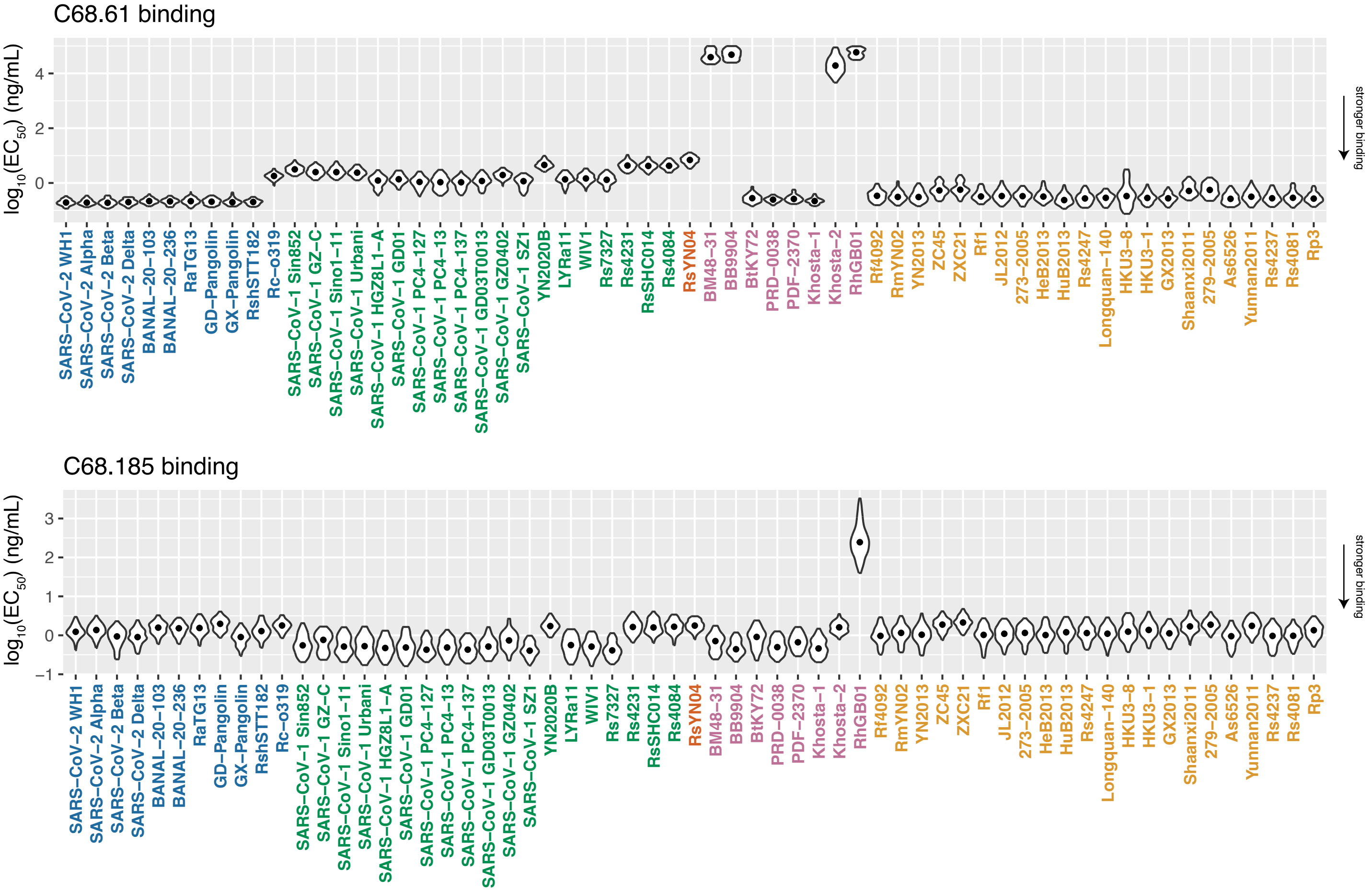

B

| | Clade 1b SARS-CoV-2 related sarbecoviruses | |
|---|---|---|
| | Geomean | 95% CI |
| **C68.61** | 0.25 | (0.39,0.16) |
| **C68.185** | 1.3 | (1.6,1.1) |
| | **Clade 1a SARS-CoV-2 related sarbecoviruses** | |
| | **Geomean** | **95% CI** |
| **C68.61** | 1.9 | (2.5,1.4) |
| **C68.185** | 0.66 | (0.85,0.52) |

**S4 Fig.**

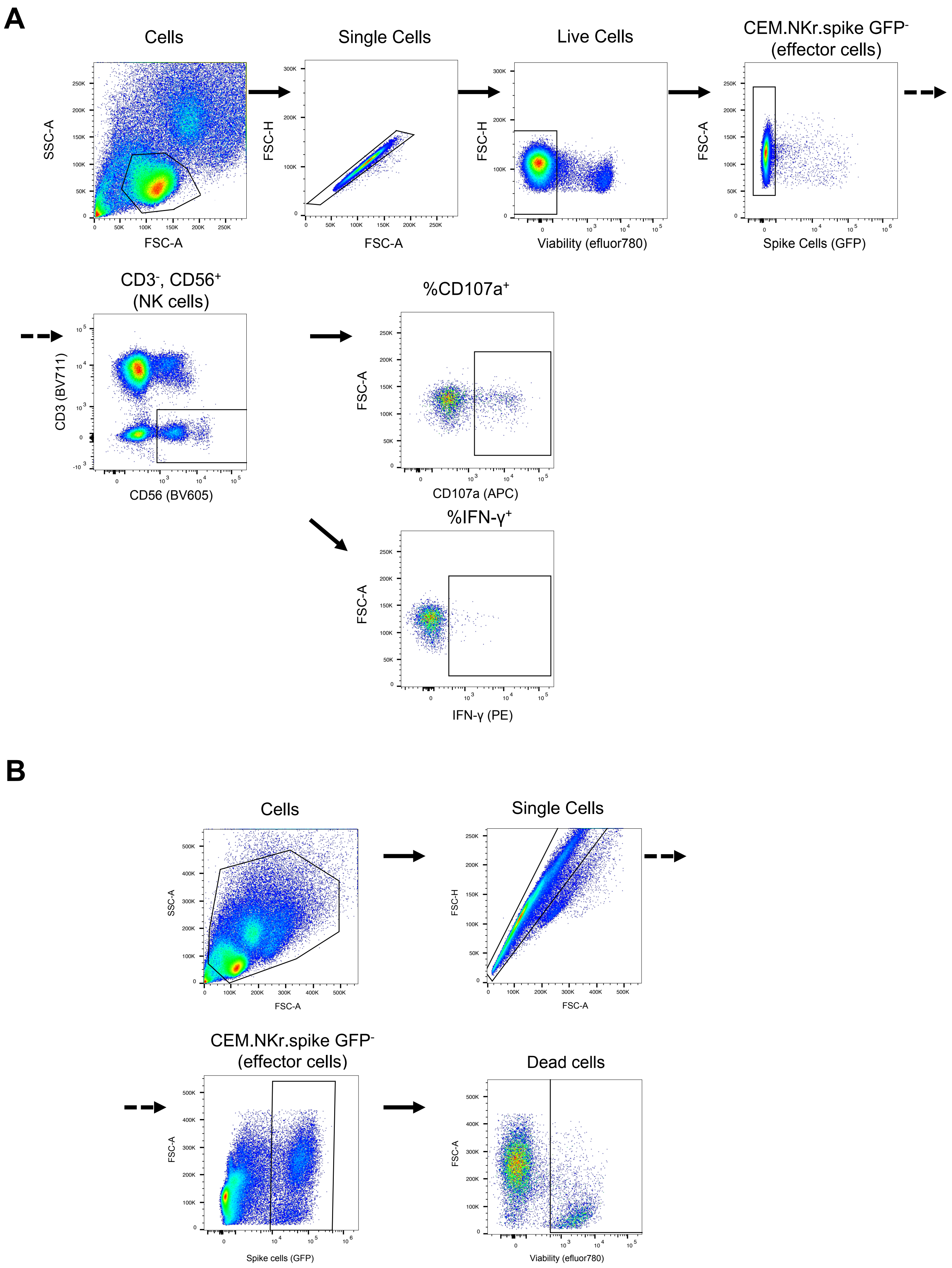
A
Cells
SSC-A
FSC-A
Single Cells
FSC-H
FSC-A
Live Cells
FSC-H
Viability (efluor780)
CEM.NKr.spike GFP-
(effector cells)
FSC-A
Spike Cells (GFP)
CD3-, CD56+
(NK cells)
CD3 (BV711)
CD56 (BV605)
%CD107a+
FSC-A
CD107a (APC)
%IFN-γ+
FSC-A
IFN-γ (PE)
B
Cells
SSC-A
FSC-A
Single Cells
FSC-H
FSC-A
CEM.NKr.spike GFP-
(effector cells)
FSC-A
Spike cells (GFP)
Dead cells
FSC-A
Viability (efluor780)

**S5 Fig.**

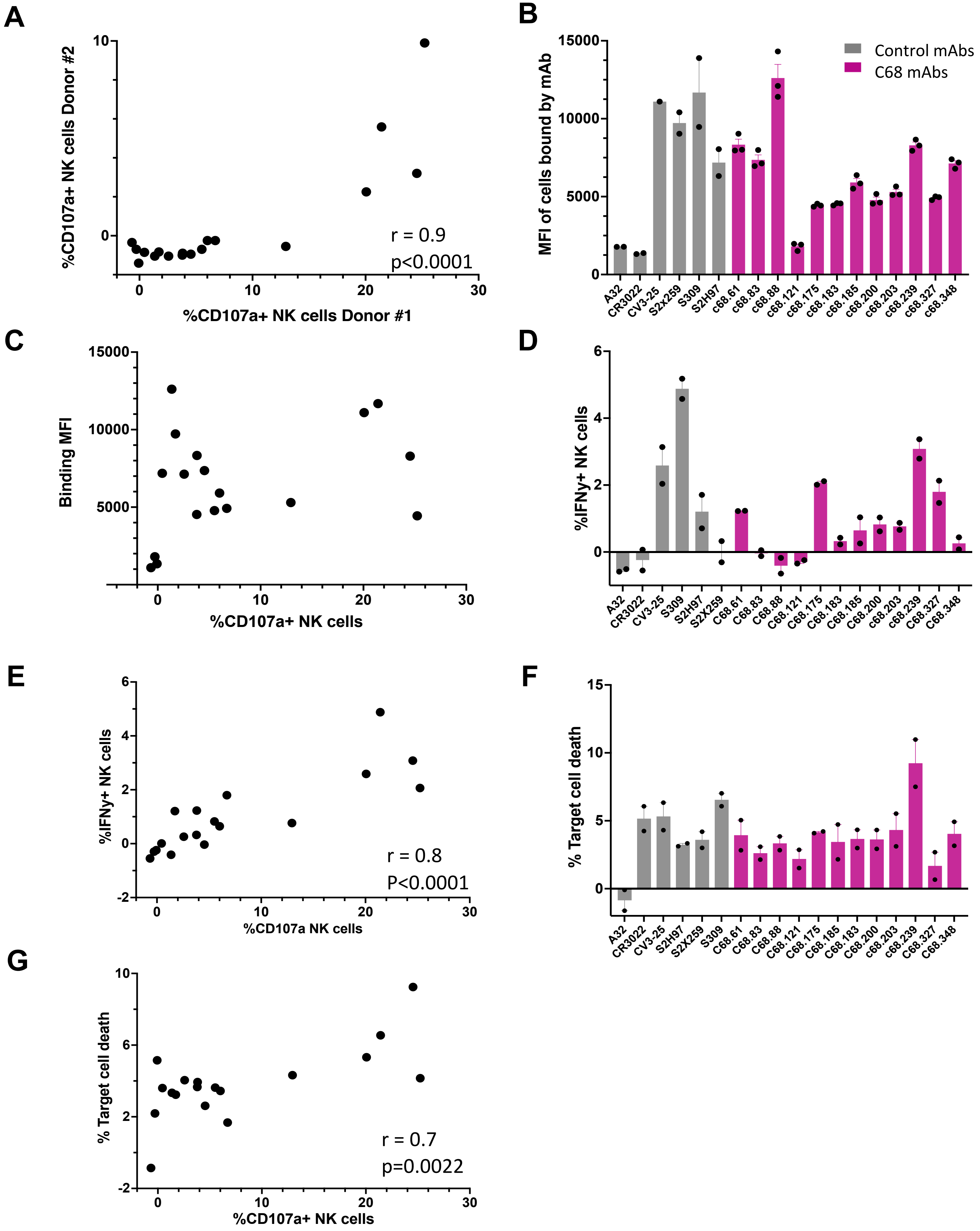

A
%CD107a+ NK cells Donor #2
%CD107a+ NK cells Donor #1
r = 0.9
p<0.0001
B
MFI of cells bound by mAb
Control mAbs
C68 mAbs
A32
CR3022
CV3-25
S2x259
S309
S2H97
c68.61
c68.83
c68.88
c68.121
c68.175
c68.183
c68.185
c68.200
c68.203
c68.239
c68.327
c68.348
C
Binding MFI
%CD107a+ NK cells
D
%IFNy+ NK cells
E
%IFNy+ NK cells
%CD107a NK cells
r = 0.8
P<0.0001
F
% Target cell death
G
% Target cell death
%CD107a+ NK cells
r = 0.7
p=0.0022

S6 Fig.

A

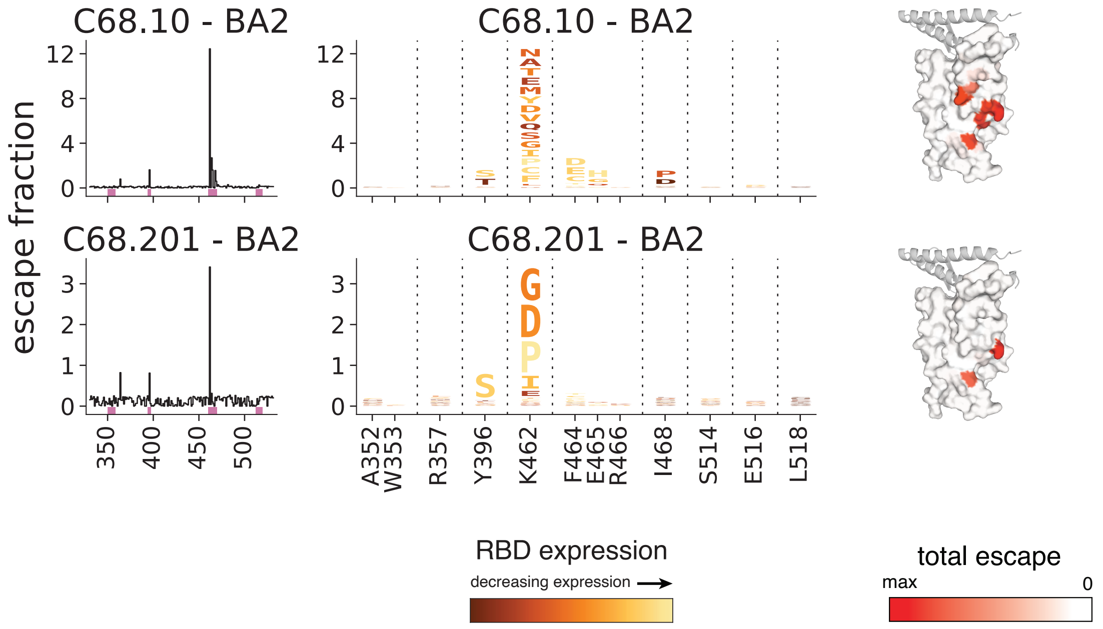
C68.10 - BA2
C68.201 - BA2
escape fraction
RBD expression
decreasing expression
total escape
max
0
A352
W353
R357
Y396
K462
F464
E465
R466
I468
S514
E516
L518

B

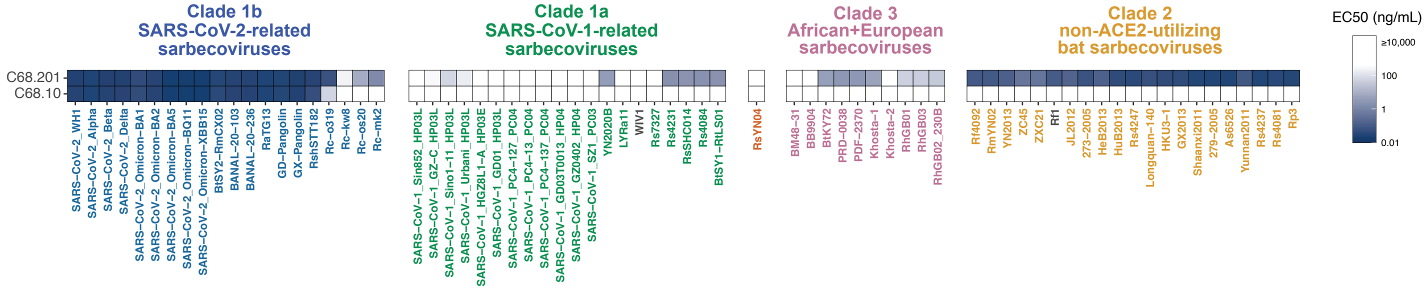
Clade 1b
SARS-CoV-2-related
sarbecoviruses
Clade 1a
SARS-CoV-1-related
sarbecoviruses
Clade 3
African+European
sarbecoviruses
Clade 2
non-ACE2-utilizing
bat sarbecoviruses
EC50 (ng/mL)
C68.201
C68.10

**S7 Fig.**

**C68.61 escape mutations**

Genotype at S site 357, 464, 465, 466, 468

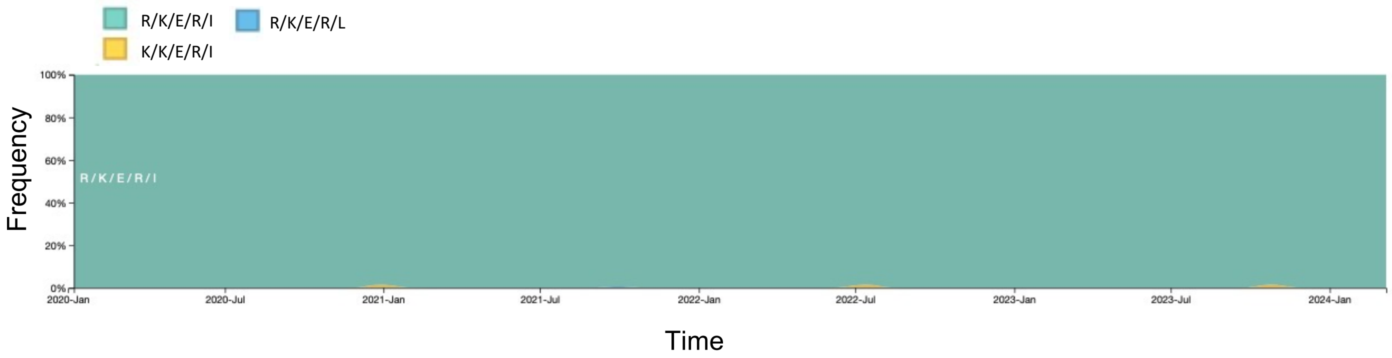

**C68.185 escape mutations**

Genotype at S site 396, 462, 464, 514, 516, 518

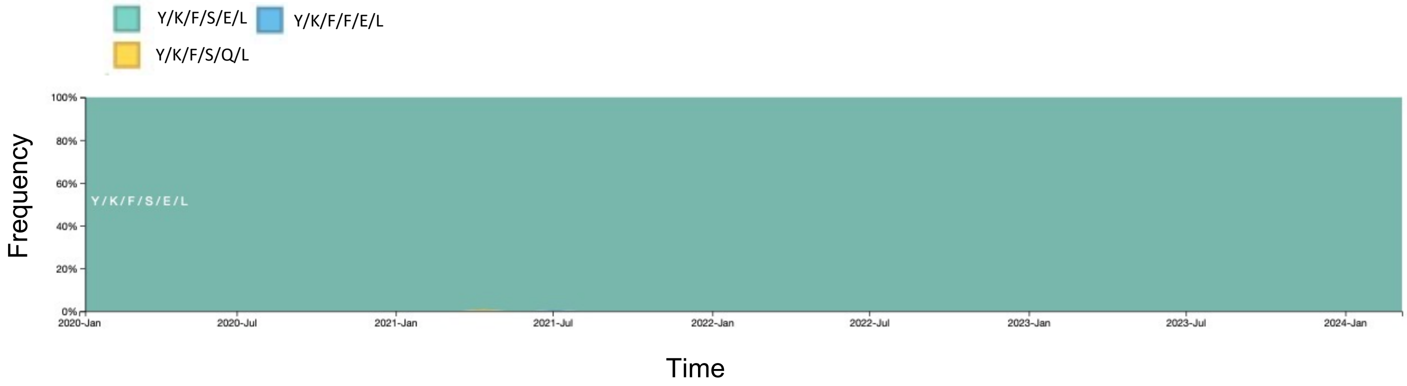

### S8 Fig.

A

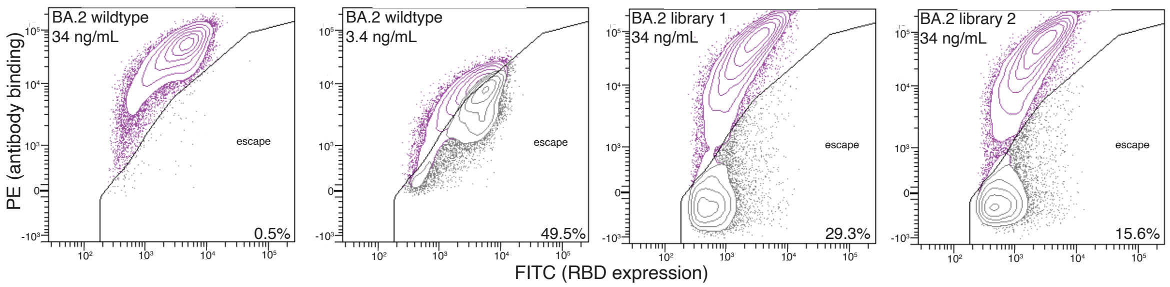
BA.2 wildtype
34 ng/mL
escape
0.5%
BA.2 wildtype
3.4 ng/mL
escape
49.5%
BA.2 library 1
34 ng/mL
escape
29.3%
BA.2 library 2
34 ng/mL
escape
15.6%
PE (antibody binding)
FITC (RBD expression)

B

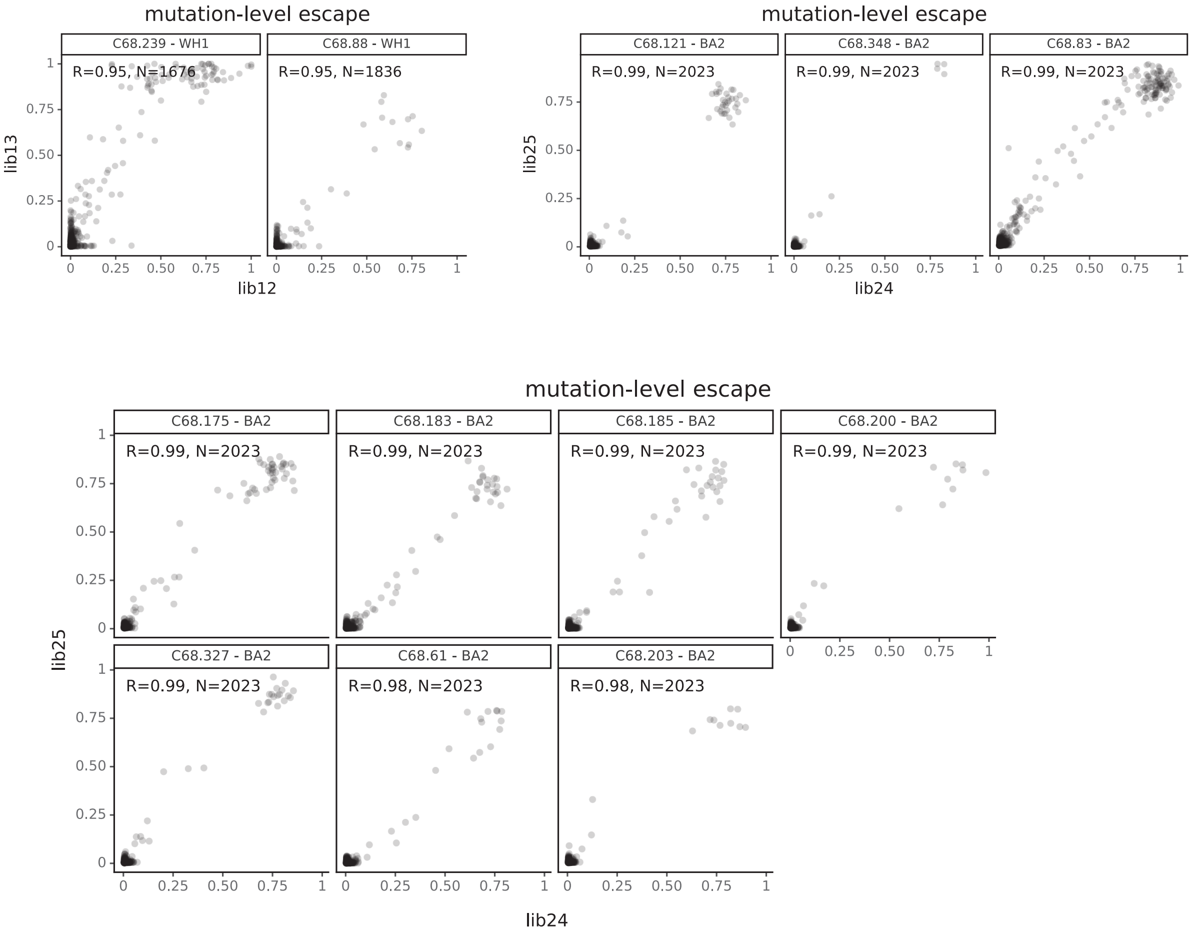
mutation-level escape
C68.239 - WH1
R=0.95, N=1676
C68.88 - WH1
R=0.95, N=1836
lib13
lib12
mutation-level escape
C68.121 - BA2
R=0.99, N=2023
C68.348 - BA2
R=0.99, N=2023
C68.83 - BA2
R=0.99, N=2023
lib25
lib24
mutation-level escape
C68.175 - BA2
R=0.99, N=2023
C68.183 - BA2
R=0.99, N=2023
C68.185 - BA2
R=0.99, N=2023
C68.200 - BA2
R=0.99, N=2023
C68.327 - BA2
R=0.99, N=2023
C68.61 - BA2
R=0.98, N=2023
C68.203 - BA2
R=0.98, N=2023
lib25
lib24

**S9 Fig.**
