## Supplementary material for "Delineating the functional activity of antibodies with cross-reactivity to SARS-CoV-2, SARS-CoV-1 and related sarbecoviruses": S1 Table

| C68 mAb ID | Sequence (Amino acids) |
|---|---|
| C68.61_IgH | EVQLVQSGAEVKKPGESLKISCKGSAYIFTRYWIGWVRQMPGKGLEWMGIIYPGDSDTRYSPSFQGQVTISADKSISTAYLQWSSLKASDTAMYYCARSGTTNYFDYWGQGTLVTVSS |
| C68.61_IgK | DIQLTQSPSTLSASVGDRVTITCRASQSISSWLAWYQQKPGKAPKLLIYDASSLDSGVPSRFSGSGSGTDFTLTISSLQPDDFATYYCQQFNSYWTFGQGTKVEIK |
| C68.83_IgH | EVQLVESGGGVVQPGRSLRLSCAASGFIFSSYGMHWVRQAPGKGLEWVAVIWYDGSNKYYADSVKGRFTISRDNSKNTLYLQMNSLRAEDTAVYYCARDDPEVGCSGGSCYYYYDMDVWGQGTTVTVSS |
| C68.83_IgL | SYELTQPPSVSVSPGQTARITCSGDALPKQHAHWYQQKPGQAPVLVIYKDIERPSGIPERFSGSSSGTTVTLTISGVQAEDEADYYCQSADSSVTYVVFGGGTKLTVL |
| C68.88_IgH | QVQLQESGPGLVKPSETLSLTCSVSGGPISSSRYYWGWIRQSPGEGLEWIGTIYYSGSTYYNPSFKSRVTISVDTSKNQFSLKLSSVTAADTAVYYCATRCYDFWSGYYNREMLWFDCWGQGTLVTVSS |
| C68.88_IgL | NFMLTQPHSVSESPGKTVTISCTRSSGSIASNYVQWYQQRPGSSPTTVIYENNQTPSGVPDRFSGSIDSSSNSASLTISGLKTEDEADYYCQSYDSSVVVFGGGTKLTVL |
| C68.121_IgH | EVQLVESGGGLVKPGGSLRLSCAASGFTFSDYYMTWIRQAPGKGLEWVSKITNSNSYTHYADSVKGRFTISRDNAKNSLFLHMNSLRAEDTAVYYCARAIWIGYPDDYYMDVWGKGTTVTVSS |
| C68.121_IgL | SYELTQPPSVSVSPGQTARITCSGDTLPNQYVYWYQQKPGQAPVLLIYKDTERPSGIPERVSGSTSGTTVTLTISGVQAEDEADYYCQSADSSGTSFGGGTKLTVL |
| C68.175_IgH | EVQLLESGGGLVQPGGSLRLSCAASGFTFSSYAMSWVRQAPGKGLEWVSGISGSGGSTYYADSVKGRFTISRDNSKNTLHLQMNSLRAEDTAVYYCAKGSRWNPDYFDYWGQGTLVTVSS |
| C68.175_IgK | DIQLTQSPSTLSASVGDRVTITCRASQSISSWLAWYQQKPGKAPKLLIYDASSLESGVPSRFSGSGSGTEFTLTISSLQPDDFATYYCQQYNSYSPLTFGGGTKVEIK |
| C68.183_IgH | EVQLLESGGGLVQPGGSLRLSCVASGFTFRNFAMSWVRQAPGKGLEWVSAISGSGDSTYYADSVKGRFTISRDNSKNTLYLQVNSLRVEDTAIYYCAKGSRGSTDYFDSWGQGTLVTVSS |
| C68.183_IgK | DIQMTQSPSTLSASIGDRVTITCRASQSISSWLAWYQQKPGKAPKLLIYDASSLESGVPSRFSGSGSGTEFTLTISSLQPDDFATYYCQQYNGYSMYTFGQGTKLEIK |
| C68.185_IgH | QVQLVQSGAEVKKPGASVKVSCKTSGYAFTDHYVHWVRQAPGQGLEWMGVINPSGVGTSYAQMFQDRVTLTSDTSTSTVYMVLSSLRSEDTAVYYCVRDDYGVPARGWFDPWGQGTLVTVSS |
| C68.185_IgK | EVVLTQSPATLSLSPGERATLSCRASQSISSYLAWYQQKPGQAPRLLIYDASNRATGVPARFSGSGSGADFTLTISSLEAEDFALYYCQHRSNWLYTFGQGTKLEIK |
| C68.200_IgH | EVQLLESGGGLVQPGGSLRLSCTASGFTFSRYALTWVRQAPGKGLEWVSAISGSGGNTNYADSVKGRFTISRDNSKNTLYLQMNSLRAEDTAVYYCAKSPRYNNDYFDFWGQGTLVTVSS |

| | |
|---|---|
| C68.200_IgK | DIQLTQSPSTLSASVGDRVTITCRASQSISSWLAWYQQKPGKAPKLLIYDASSLESGVPSRFSGSGSGTEFTLTISSLQPDDFATYYCQQYNTFITFGHGTRLEIK |
| C68.203_IgH | EVQLLESGGGLVQPGGSLRLSCTASGFTFSRYALTWVRQAPGKGLECVSAFSGSGDSTYYADSVKGRFTISRDNSKNTLYLQMNSLRAEDTAVYYCAKSPRYNSDYFDFWGQGTLVTVSS |
| C68.203_IgK | DIQLTQSPSTLSASVGDRVTITCRASQSISSWLAWYQQKPGKAPKLLIYDASSLESGVPSRFSGSGSGTEFTLTISSLQPDDFATYYCQQYNSFITFGQGTRLEIK |
| C68.239_IgH | EVQLLESGGGLVQPGGSLRLSCAASGFTFSSYAMSWVRQAPGKGLEWVSAISGGGDNTYYADSVKGRFTISRDNSKNTLYLQMNSLRAEDTAVYFCAKNPITVVPAAWDRGYFDLWGRGTLVTVSS |
| C68.239_IgL | SYELTQPPSVSVAPRKTARITCGGNNIGSYSVHWYQQRPGQAPVLVVHDDSDRPSGIPERFSGSNSGNTATLTISRVEAGDEADYYCQVWDSSSDHRGVFGGGTKLTVL |
| C68.327_IgH | EVQLLESGGGLVQPGGSLRLSCAASGFTFGSYAMSWVRQAPGKGLEWVSAISGSGGSTYYADSVKGRFTISRDNSKNTLYLQMNSLRAEDTAVYNCAKDLWDGFHWFDSWGQGTLVTVSS |
| C68.327_IgK | DIQMTQSPSSVSASVGDRVTITCRASQGISSWLAWYQQKPGKAPNLLIYSASSLQSGVPSRFSGSGSGTDFTLTISSLQPEDFATYYCQQANSFPITFGQGTRLEIK |
| C68.348_IgH | QVQLQESGPRLVKSSGTLSLTCAVSGGSISSSKWWSWVRQPPGKGLEWIGEVYHGGSTNYNPSLKSRVTISVDKAKNQFSLKLSSVTAADSAVYYCASLDSAAVFDYWGQGTLVTVSS |
| C68.348_IgL | QSALTQPASVSGSPGQSITISCTGTSSDVGRFNYVSWYQQHPGKAPKLLIYDVSNRPSGVSNRFSGSKSGNTASLTISGLQAEDEADYYCNSYTSSSTLYVFGTGTKVTVL |
