## Supplementary material for "Delineating the functional activity of antibodies with cross-reactivity to SARS-CoV-2, SARS-CoV-1 and related sarbecoviruses": S2 Table

| strain | gisaid_epi_isl | genbank_accession | originating_lab | submitting_lab | author |
|---|---|---|---|---|---|
| hCoV-19/Wuhan/Hu-1/2019 | EPI_ISL_402125 | MN908947.3 | National Institute for Communicable Disease Control and Prevention (ICDC) Chinese Center for Disease Control and Prevention (China CDC) | National Institute for Communicable Disease Control and Prevention (ICDC) Chinese Center for Disease Control and Prevention (China CDC) | Zhang et al A (https://dx.doi.org/10.1038/s41586-020-2008-3) |
| hCoV-19/Guangdong/GD2020139-P0007/2020 | EPI_ISL_413882 | | Guangdong Provincial Institution of Public Health, Guangdong Provinical Center for Disease Control and Prevention | Guangdong Provincial Institution of Public Health | Jing Lu et al (https://dx.doi.org/10.1016/j.cell.2020.04.023) |
| hCoV-19/Zhejiang/WZ-01/2020 | EPI_ISL_404227 | | Zhejiang Provincial Center for Disease Control and Prevention | Department of Microbiology, Zhejiang Provincial Center for Disease Control and Prevention | Yin Chen et al |
| hCoV-19/Wuhan/IME-WH04/2019 | EPI_ISL_529216 | MT291829.1 | Beijing Institute of Microbiology and Epidemiology | Beijing Institute of Microbiology and Epidemiology | Fan et al |
| hCoV-19/Taiwan/2/2020 | EPI_ISL_406031 | | Centers for Disease Control, R.O.C. (Taiwan) | Centers for Disease Control, R.O.C. (Taiwan) | Ji-Rong Yang et al A (https://dx.doi.org/10.1099/acmi.0.000133) |
| hCoV-19/Guangxi/NAN.20-4/2020 | EPI_ISL_6951027 | | Molecular Diagnostics Lab, Affiliated Infectious Disease Hospital of Nanning, Guangxi Medical University | Key Laboratory of Infectious Diseases,Nanning Health Commission. | JianYan Lin et al |
| hCoV-19/Jiangxi/JX2019/2020 | EPI_ISL_455507 | | Jiangxi Province Center for Disease Control and Prevention | Jiangxi Province Center for Disease Control and Prevention | JianXiong Li et al |
| hCoV-19/Anhui/29-2/2020 | EPI_ISL_16138317 | | College of life sciences, Anhui Medical University | College of life sciences, Anhui Medical University | Cheng et al |
| hCoV-19/Australia/NSW03/2020 | EPI_ISL_408977 | | Serology, Virology and OTDS Laboratories (SAViD), NSW Health Pathology Randwick | NSW Health Pathology - Institute of Clinical Pathology and Medical Research; Centre for Infectious Diseases and Microbiology Laboratory Services; Westmead Hospital; University of Sydney | Eden J-S et al (https://www.nature.com/articles/s41591-020-1000-7) |
| hCoV-19/Vietnam/KH-01/2020 | EPI_ISL_17373276 | | CDC Khánh Hòa, Vietnam | Department of Microbiology and Immunology, Pasteur Institute in Nha Trang | Trinh Hoang Long et al |
| hCoV-19/Vietnam/PIHCM-0002/2020 | EPI_ISL_17454569 | | Pasteur Institute in Ho Chi Minh city Department of Microbiology and Immunology | Saint Petersburg Pasteur Institute, Molecular Genetic Monitoring Group | Thang Minh Cao et al |
| hCoV-19/Wuhan/0125-A171/2020 | EPI_ISL_493162 | | National Virus Resource Center, Chinese Academy of Sciences, Wuhan 430071, China | Computational Virology Group, Center for Bacteria and Viruses Resources and Bioinformation, Wuhan Institute of Virology, Chinese Academy of Sciences, Wuhan 430071, China | Jianjun Chen et al |
| hCoV-19/Wuhan/HB-WH3-188/2020 | EPI_ISL_455383 | | Wuhan Chain Medical Labs (CMLabs) | State Key Laboratory of Biotherapy of Sichuan University | Baowen Du et al |
| hCoV-19/Thailand/NIH-162/2020 | EPI_ISL_434694 | | Bamrasnaradura hospital | National Institute of Health. Department of medical Sciences, Ministry of Public Health, Thailand | Pilailuk et al |
| hCoV-19/Malaysia/6359/2020 | EPI_ISL_501220 | | Department of Medical Microbiology, University Malaya Medical Centre | Department of Medical Microbiology, Faculty of Medicine, University of Malaya | Yoong Min CHONG et al |
| hCoV-19/UnitedArabEmirates/skmc-004/2020 | EPI_ISL_582613 | | Sheikh Khalifa Medical City | Molecular/Surveillance lab Sheikh Khalifa Medical City | Amirtharaj Francis et al |
| hCoV-19/Lu-an/425/2020 | EPI_ISL_1081260 | | Microbiology Laboratory,Lu`an Center for Disease Control and Prevention | Microbiology Laboratory,Lu`an Center for Disease Control and Prevention | Yang Wei et al |
| hCoV-19/Shanghai/SH0091/2020 | EPI_ISL_416388 | | Shanghai Public Health Clinical Center, Shanghai Medical College, Fudan University | National Research Center for Translational Medicine (Shanghai), Ruijin Hospital affiliated to Shanghai Jiao Tong University School of Medicine & Shanghai Public Health Clinical Center | Shengyue Wang et al |
| hCoV-19/Fujian/13/2020 | EPI_ISL_411066 | | Fujian Center for Disease Control and Prevention | Fujian Center for Disease Control and Prevention | Chen Wei et al |
| hCoV-19/Hunan/HN-CIDC-P5/2020 | EPI_ISL_539334 | | Institute of Disease Control and Prevention, People's Liberation Army | Institute of Disease Control and Prevention, People's Liberation Army | Qiu et al |
| hCoV-19/Jiangsu/JS03/2020 | EPI_ISL_411953 | | NHC Key laboratory of Enteric Pathogenic Microbiology, Institute of Pathogenic Microbiology | Jiangsu Provincial Center for Disease Control & Prevention | Kangchen Zhao et al |
| hCoV-19/Sichuan/SC-WCH4-289/2020 | EPI_ISL_451391 | | West China Hospital of Sichuan University | State Key Laboratory of Biotherapy of Sichuan University | Baowen Du et al |
| hCoV-19/Guangdong/20SF665/2020 | EPI_ISL_428469 | | Guangdong Provincial Center for Diseases Control and Prevention;Guangdong Provincial Institute of Public Health | School of Public Health, The University of Hong Kong | Bosheng Li et al |
| hCoV-19/Wuhan/DT-WH03/2020 | EPI_ISL_453782 | | Laboratory of Infectious Diseases Center of Beijing Ditan Hospital | Laboratory of Infectious Diseases Center of Beijing Ditan Hospital | Siyuan Yang et al |
| hCoV-19/Yunnan/0306-466/2020 | EPI_ISL_429239 | MT396241.1 | Department of Clinical Laboratory, the First People's Hospital of Yunnan Province | Department of Clinical Laboratory, the First People's Hospital of Yunnan Province | Yi Sun et al |
| hCoV-19/USA/WV-QDX-1284/2020 | EPI_ISL_572053 | MW065351.1 | Quest Diagnostics | Quest Diagnostics | Rosenthal et al |
| hCoV-19/Oman/RESP-20-B-6588/2020 | EPI_ISL_525429 | | Oman-National Influenza Center | Biotechnology & OMICs Laboratory | Samira Al-Mahruqi et al |
| hCoV-19/HongKong/HKPU-0248/2020 | EPI_ISL_610172 | | Department of Health Technology and Informatics, The Hong Kong Polytechnic University | Department of Health Technology and Informatics, The Hong Kong Polytechnic University | Siu et al |
| hCoV-19/Bulgaria/15/2020 | EPI_ISL_480298 | | National Reference Laboratory "Influenza and acute respiratory diseases" | NRL-HIV | Ivan Ivanov et al |
| hCoV-19/Pakistan/Gilgit1/2020 | EPI_ISL_417444 | MT240479.1 | Department of Healthcare Biotechnology, National University of Sciences and Technology (NUST) | Department of Healthcare Biotechnology, National University of Sciences and Technology (NUST) | Javed et al (http://biorxiv.org/lookup/doi/10.1101/2020.04.01.019463) |
| hCoV-19/Oman/RESP-20-837/2020 | EPI_ISL_457704 | | Oman-NIC | Oman-NIC | Samira Al-Maruqi et al |
| hCoV-19/Oman/RESP-20-797/2020 | EPI_ISL_457701 | | Oman-NIC | Oman-NIC | Samira Al-Maruqi et al |
| hCoV-19/SriLanka/CDR503/2020 | EPI_ISL_525479 | | Centre for Dengue Research | Centre for Dengue Research | Chandima Jeewandara et al |
| hCoV-19/Georgia/Tb-82/2020 | EPI_ISL_415644 | | R. G. Lugar Center for Public Health Research, National Center for Disease Control and Public Health (NCDC) of Georgia. | R. G. Lugar Center for Public Health Research, National Center for Disease Control and Public Health (NCDC) of Georgia. | Nato Kotaria et al |
| hCoV-19/Bahrain/BAH-25/2020 | EPI_ISL_483639 | | Kingdom of Bahrein Ministry of Health | Erasmus Medical Center | Bas Oude Munnink et al |
| hCoV-19/Bahrain/BAH-27/2020 | EPI_ISL_483565 | | Kingdom of Bahrein Ministry of Health | Erasmus Medical Center | Bas Oude Munnink et al |
| hCoV-19/Taiwan/TSGH-26/2020 | EPI_ISL_447252 | | TSGH-CP molecular lab | TSGH-CP molecular lab | Cherng-Lih Perng et al |
| hCoV-19/Tunisia/7899/2020 | EPI_ISL_763067 | | 1-Laboratory of Microbiology, National Reference Lab, Charles Nicolle Hospital; 2-University of Tunis ElManar, Faculty of Medicine of Tunis, LR99ES09, Tunis, Tunisia | 1-Clinical and Experimental Pharmacology Lab, LR16SP02, National Center of Pharmacovigilance, University of Tunis El Manar, Tunis, Tunisia. 2- Neurodegenerative diseases and psychiatric troubles, LR18SP03, Razi Hospital, University of Tunis El Manar, Tunis, Tunisia. 3- Ministry of Health, National Observatory of New and Emerging Diseases, 1006, Tunis, Tunisia | Ilhem Boutiba-Ben Boubaker et al |
| hCoV-19/Georgia/Tb-7851/2020 | EPI_ISL_477169 | MT786438.1 | Department for Virology, Molecular Biology and Genome Research, R. G. Lugar Center for Public Health Research, National Center for Disease Control and Public Health (NCDC) of Georgia. | Department for Virology, Molecular Biology and Genome Research, R. G. Lugar Center for Public Health Research, National Center for Disease Control and Public Health (NCDC) of Georgia. | Tata Imnadze et al |
| hCoV-19/Iceland/13/2020 | EPI_ISL_417765 | OU161584.1 | The National University Hospital of Iceland | deCODE genetics | Daniel F Gudbjartsson et al A (https://dx.doi.org/10.1056/NEJMoa2006100) |
| hCoV-19/Georgia/MM202054/2020 | EPI_ISL_2896259 | MT231246.1 | Department of Virology, Molecular Biology and Genome Research | Department of Virology, Molecular Biology and Genome Research | Kotorashvili et al |
| hCoV-19/Georgia/Tb-54/2020 | EPI_ISL_415641 | | R. G. Lugar Center for Public Health Research, National Center for Disease Control and Public Health (NCDC) of Georgia. | R. G. Lugar Center for Public Health Research, National Center for Disease Control and Public Health (NCDC) of Georgia. | Nato Kotaria et al |
| hCoV-19/Aruba/AW-RIVM-10122/2020 | EPI_ISL_547449 | | National Institute for Public Health and the Environment (RIVM) | National Institute for Public Health and the Environment (RIVM) | Adam Meijer et al |
| hCoV-19/Qatar/QA-WCMQ_FD15540717/2020 | EPI_ISL_1714064 | | Ministry of Public Health / Hamad Medical Corporation | Weill Cornell Medical College - Qatar (WCM-Q), Genomics Core Laboratory / Qatar Genome Project (QGP) | WCMQ: Ayeda A. Ahmed et al |
| hCoV-19/Jordan/SR-057/2020 | EPI_ISL_430015 | MT811565.1 | Biolab Diagnostic Laboratories | Andersen lab at Scripps Research | Issa Abu-Dayyeh et al |
| hCoV-19/Anhui/909/2020 | EPI_ISL_16138285 | | College of life sciences, Anhui Medical University | College of life sciences, Anhui Medical University | Cheng et al |
| hCoV-19/Japan/FHU-CS108-0222/2020 | EPI_ISL_728156 | | Fujita Health University Okazaki Medical Center | Department of Virology and Parasitology, Fujita Health University School of Medicine | Takayuki Murata et al |
| hCoV-19/India/JK-ICMR-229/2020 | EPI_ISL_1072989 | | VRDL, Sher-I-Kashmir Institute of Medical Sciences, Srinagar, Jammu & Kashmir | Indian Council of Medical Research-National Institute of Virology, Microbial Containment Complex | Pragya D. Yadav et al |
| hCoV-19/Zambia/ZMB-319/2020 | EPI_ISL_2803448 | | University of Zambia, School of Veterinary Medicine | UNZAVET and PATH | Mulenga Mwenda-Chimfwembe et al |
| hCoV-19/Brunei/3/2020 | EPI_ISL_435675 | | National Virology Reference Laboratory | National Public Health Laboratory, National Centre for Infectious Diseases | Mak Tze Minn et al |
| hCoV-19/SaudiArabia/KAIMRC01/2020 | EPI_ISL_507010 | MT820461.1 | | Infectious Diseases Research, King Abdullah International Medical Research Center (KAIMRC) | Alghoribi et al |
| hCoV-19/India/DL-MaxCov0044-CSIR-IGIB/2020 | EPI_ISL_459946 | | Devki Devi Foundation, a unit of Max Healthcare | CSIR-IGIB/Max | Rajesh Pandey# et al |
| hCoV-19/Philippines/GS/2020 | EPI_ISL_15592185 | | VIROLOGY, AFRIMS | VIROLOGY, AFRIMS | Velasco et al |
| hCoV-19/India/AS-563/2020 | EPI_ISL_11448682 | | State Level VRDL GMCH | State Level VRDL GMCH | Ajanta Sharma et al |
| hCoV-19/India/AS-22580/2020 | EPI_ISL_11448683 | | State Level VRDL GMCH | State Level VRDL GMCH | Ajanta Sharma et al |
| hCoV-19/India/MP-ICMR-NIV-INSACOG-SEQ-3717/2020 | EPI_ISL_15038584 | | ICMR-National Institute of Virology - INSACOG | NIV Influenza | Dr. Varsha Potdar et al |
| hCoV-19/India/TG-GMC-RK1090/2020 | EPI_ISL_438139 | | Department of Microbiology,Gandhi Medical College and Hospital,Hyderabad | Virus Research Laboratory, Department of Zoology, Osmania University, Hyderabad, India | Muttineni Radhakrishna et al |
| hCoV-19/India/RJ-NCDC-2522/2020 | EPI_ISL_436420 | | National Centre for Disease control (NCDC) | NCDC/CSIR-IGIB | Pramod Kumar# et al (https://dx.doi.org/10.1101/2020.06.04.128751) |
| hCoV-19/India/TN-NCDC-02327/2020 | EPI_ISL_435094 | | National Centre for Disease control (NCDC), CSIR-Institute of Genomics and Integrative Biology (CSIR-IGIB) | NCDC/CSIR-IGIB | Pramod Kumar et al (https://dx.doi.org/10.1101/2020.06.04.128751) |
| hCoV-19/Brunei/4/2020 | EPI_ISL_435676 | | National Virology Reference Laboratory | National Public Health Laboratory, National Centre for Infectious Diseases | Mak Tze Minn et al |
| hCoV-19/Malaysia/2101/2020 | EPI_ISL_501191 | | Department of Medical Microbiology, University Malaya Medical Centre | Department of Medical Microbiology, Faculty of Medicine, University of Malaya | Yoong Min CHONG et al |
| hCoV-19/Malaysia/2065/2020 | EPI_ISL_501189 | | Department of Medical Microbiology, University Malaya Medical Centre | Department of Medical Microbiology, Faculty of Medicine, University of Malaya | Yoong Min CHONG et al |
| hCoV-19/Timor-Leste/TL07/2020 | EPI_ISL_456606 | MT612289.1 | National Health Laboratory, Timor-Leste | Microbiological Diagnostic Unit Public Health Laboratory, The Peter Doherty Institute for Infection and Immunity | Soares da Silva et al |
| hCoV-19/Singapore/236/2020 | EPI_ISL_462354 | | National Public Health Laboratory, National Centre for Infectious Diseases | National Public Health Laboratory, National Centre for Infectious Diseases | Mak TM et al |
| hCoV-19/India/MH-NCDC-02330/2020 | EPI_ISL_435077 | | National Centre for Disease control (NCDC), CSIR-Institute of Genomics and Integrative Biology (CSIR-IGIB) | NCDC/CSIR-IGIB | Pramod Kumar et al (https://dx.doi.org/10.1101/2020.06.04.128751) |
| hCoV-19/India/TG-CCMB-J043/2020 | EPI_ISL_447560 | | CSIR-Centre for Cellular and Molecular Biology | CSIR-Centre for Cellular and Molecular Biology | Payel Mukherjee et al A (https://dx.doi.org/10.1101/2020.05.31.126136) |
| hCoV-19/Philippines/PH-RITM-0065/2020 | EPI_ISL_3105872 | | Taguig City Health Office | Research Institute for Tropical Medicine | Lei Lanna Dancel et al |
| hCoV-19/India/HR-THSTI-BAL-912/2020 | EPI_ISL_454864 | MZ356528.1 | Translational Health Science and Technology Institute -ESIC medical college and hospital, Faridabad | THSTI Bioassay laboratory | Saurabh Kumar et al |
| hCoV-19/Malaysia/MGI-MAEPS54/2020 | EPI_ISL_528743 | | Malaysia Genome Institute | Malaysia Genome Institute | Mohd Noor Mat Isa et al |
| hCoV-19/India/AP-CS0330/2020 | EPI_ISL_862518 | | Kurnool Medical College (KMC) | CSIR Institute of Genomics and Integrative Biology | Pallavali Roja Rani et al |
| hCoV-19/Sichuan/SC-NC-071/2020 | EPI_ISL_451330 | | West China Hospital of Sichuan University | State Key Laboratory of Biotherapy of Sichuan University | Baowen Du et al |
| hCoV-19/Zhejiang/Shaoxing-10/2020 | EPI_ISL_464102 | | Shaoxing Center for Disease Control and Prevention | Department of Pathology and Laboratory Medicine, University of California Los Angeles | Jinkun Chen et al |
| hCoV-19/Nanchang/JX14/2020 | EPI_ISL_421243 | | Jiangxi Province Center for Disease Control and Prevention | Jiangxi Province Center for Disease Control and Prevention | JianXiong Li et al |
| hCoV-19/Canada/BC_83109/2020 | EPI_ISL_415588 | | BCCDC Public Health Laboratory | BCCDC Public Health Laboratory | Harrigan et al |
| hCoV-19/Chongqing/YC01/2020 | EPI_ISL_408478 | | Yongchuan District Center for Disease Control and Prevention | Chongqing Municipal Center for Disease Control and Prevention | Ye Sheng et al |
| hCoV-19/Yunnan/01/2020 | EPI_ISL_16138328 | | Yunnan Center for Disease Control Prevention, Institute for Acute Communicable Disease Prevention and Control | Yunnan Center for Disease Control Prevention, Institute for Acute Communicable Disease Prevention and Control | Fu et al |
| hCoV-19/Cambodia/Kunming_kms-2/2020 | EPI_ISL_682298 | MW341443.1 | Respiratory virus Laboratory, Chinese Academy of Medical Science | Respiratory virus Laboratory, Chinese Academy of Medical Science | Li et al |
| hCoV-19/Beijing/BJ752/2020 | EPI_ISL_430724 | | Chinese PLA Institute for Disease Control and Prevention | Chinese PLA Institute for Disease Control and Prevention | Peng Li et al |
| hCoV-19/Henan/fahzu0018/2020 | EPI_ISL_1040039 | | PCR Laboratory, The First Affiliated Hospital of Zhengzhou University, Zhengzhou, Henan, China | PCR Laboratory, The First Affiliated Hospital of Zhengzhou University, Zhengzhou, Henan, China | Guanglin Cui et al |
| hCoV-19/SouthKorea/KCDC07/2020 | EPI_ISL_412871 | MT134008.1 | Division of Viral Diseases, Center for Laboratory Control of Infectious Diseases, Korea Centers for Diseases Control and Prevention | Division of Viral Diseases, Center for Laboratory Control of Infectious Diseases, Korea Centers for Diseases Control and Prevention | Jeong-Min Kim et al |
| hCoV-19/SouthKorea/KCDC06/2020 | EPI_ISL_412870 | MT134007.1 | Division of Viral Diseases, Center for Laboratory Control of Infectious Diseases, Korea Centers for Diseases Control and Prevention | Division of Viral Diseases, Center for Laboratory Control of Infectious Diseases, Korea Centers for Diseases Control and Prevention | Jeong-Min Kim et al |
| hCoV-19/Beijing/105/2020 | EPI_ISL_413518 | MT135041.1 | Infectious Disease Control Center, Center for Disease Control and Prevention of PLA | Infectious Disease Control Center, Center for Disease Control and Prevention of PLA | Li et al |
| hCoV-19/Shandong/2020C1240428/2020 | EPI_ISL_962547 | | Infectious Disease Control and Prevention Institute | Infectious Disease Control and Prevention Institute | Xiaolin Jiang et al |
| hCoV-19/Beijing/DT-travelES02/2020 | EPI_ISL_452344 | | Laboratory of Infectious Diseases Center of Beijing Ditan Hospital | Laboratory of Infectious Diseases Center of Beijing Ditan Hospital | Siyuan Yang et al |
| hCoV-19/Spain/VC-FISABIO-4/2020 | EPI_ISL_416483 | MT233520.1 | Servicio de Microbiología. Consorcio Hospital General Universitario de Valencia | Sequencing and Bioinformatics Service and Molecular Epidemiology Research Group. FISABIO-Public Health | Maria Alma Bracho et al |
| hCoV-19/EquatorialGuinea/253/2020 | EPI_ISL_648318 | | Laboratorio de Investigaciones de Baney | University Hospital Basel, Clinical Bacteriology | Carlos Cortes et al |
| hCoV-19/Kazakhstan/7341/2020 | EPI_ISL_454572 | | National Center of Expertise | National Center for Expertise, Kazakhstan National Center for Biotechnology, Kazakhstan | Abdaliyev Askar et al |
| hCoV-19/Kazakhstan/17920/2020 | EPI_ISL_454577 | | Laboratory of virology, National Center of Expertise | Laboratory of molecular-genetic research, National Center of Expertise, Kazakhstan National Center for Biotechnology, Kazakhstan | Abdaliyev Askar et al |
| hCoV-19/Kazakhstan/18243/2020 | EPI_ISL_454500 | | RSE "National Center for Biotechnology" | RSE "National Center for Biotechnology" | Alexandr Shevtsov et al |
| hCoV-19/Panama/GMI-PA357073/2020 | EPI_ISL_1225389 | | Gorgas Memorial Laboratory of Health Studies | Gorgas Memorial Laboratory of Health Studies | Yamilka Diaz et al |
| hCoV-19/Panama/GMI-PA398110/2020 | EPI_ISL_1225489 | | Gorgas Memorial Laboratory of Health Studies | Gorgas Memorial Laboratory of Health Studies | Yamilka Diaz et al |
| hCoV-19/Panama/GMI-PA676305/2021 | EPI_ISL_1503129 | | Hospital Regional Anita Moreno Los Santos | Gorgas Memorial Laboratory of Health Studies | Gonzalez Claudia et al |
| hCoV-19/Panama/GMI-PA641589/2021 | EPI_ISL_1502957 | | Gorgas Memorial Laboratory of Health Studies | Gorgas Memorial Laboratory of Health Studies | Gonzalez Claudia et al |
| hCoV-19/Paraguay/WOR12903A869/2021 | EPI_ISL_6944628 | ON017309.1 | Laboratorio Central de Salud Publica | Viral Special Pathogens Branch, Centers for Disease Control and Prevention | Cynthia Vázquez et al |
| hCoV-19/CostaRica/INC-0486/2021 | EPI_ISL_1827516 | | HOSPITAL MONSEÑOR SANABRIA | Inciensa, Instituto Costarricense de Investigación y Enseñanza en Nutrición y Salud | Francisco Duarte et al |
| hCoV-19/Panama/GMI-PACNB25262/2021 | EPI_ISL_11681785 | | Gorgas Memorial Institute of Health Studies | Gorgas Memorial Institute of Health Studies | Claudia Gonzalez et al |
| hCoV-19/Panama/GMI-PA713890/2021 | EPI_ISL_4744544 | | Gorgas Memorial Laboratory of Health Studies | Gorgas Memorial Laboratory of Health Studies | Gonzalez Claudia et al |
| hCoV-19/Panama/GMI-PA677127/2021 | EPI_ISL_1503132 | | Gorgas Memorial Laboratory of Health Studies | Gorgas Memorial Laboratory of Health Studies | Gonzalez Claudia et al |
| hCoV-19/Belize/CML-77/2021 | EPI_ISL_4296381 | | Central Medical Laboratory | Baylor College of Medicine | Alexander Kneubehl et al |
| hCoV-19/Nicaragua/CNDR-130414/2021 | EPI_ISL_17065503 | | CENTRO DE SALUD HEROES Y MARTIRES-15 DE JULIO | CNDR, Departamento de Virología | Angel Balmaseda et al |
| hCoV-19/India/MH-1-31/2020 | EPI_ISL_413523 | | Indian Council of Medical Research-National Institute of Virology | National Influenza Center, Indian Council of Medical Research-National Institute of Virology | Potdar et al A (https://dx.doi.org/10.4103/ijmr.IJMR_663_20) |
| hCoV-19/Guangdong/GDIPH-GDPCC-03/2020 | EPI_ISL_18934698 | | Guangdong Provincial Institute of Public Health, Guangdong Provincial Institute of Public Health | Guangdong Provincial Institute of Public Health, Guangdong Provincial Institute of Public Health | Zhou et al |
| hCoV-19/India/un-ILS21/2020 | EPI_ISL_15591022 | | Infectious Disease Biology, Institute of Life Sciences | Infectious Disease Biology, Institute of Life Sciences | Syed et al |
| hCoV-19/Niger/202034/2020 | EPI_ISL_2281811 | | Centre de Recherche Médicale et Sanitaire (CERMES) | Institut Pasteur de Dakar | Lagare Adamou et al |
| hCoV-19/Niger/202029/2020 | EPI_ISL_2281810 | | Centre de Recherche Médicale et Sanitaire (CERMES) | Institut Pasteur de Dakar | Lagare Adamou et al |
| hCoV-19/Togo/BMC-16649-20/2020 | EPI_ISL_1416192 | | Institut National d'hygiène | "Unité Mixte Internationale TransVIHMI (UMI 233 IRD – U1175 INSERM - Université de Montpellier) IRD (Institut de recherche pour le développement)" | Mounerou SALOU et al |
| hCoV-19/Coted-Ivoire/BKE1661-bc58/2021 | EPI_ISL_3545645 | | Molecular diagnostic unit for viral haemorrhagic fevers and emerging viruses, Bouake CHU Laboratory | Molecular diagnostic unit for viral haemorrhagic fevers and emerging viruses, Bouake CHU Laboratory | Chantal Akoua-Koffi et al |
| hCoV-19/Niger/Y/2020 | EPI_ISL_2281821 | | Centre de Recherche Médicale et Sanitaire (CERMES) | Institut Pasteur de Dakar | Lagare Adamou et al |
| hCoV-19/Coted-Ivoire/COV-BKE1450-bc27/2020 | EPI_ISL_1662511 | | Molecular diagnostic unit for viral haemorrhagic fevers and emerging viruses, Bouaké CHU Laboratory | Molecular diagnostic unit for viral haemorrhagic fevers and emerging viruses, Bouaké CHU Laboratory | Chantal Akoua-Koffi et al |
| hCoV-19/Coted-Ivoire/BKE0366/2020 | EPI_ISL_681834 | | Molecular diagnostic unit for viral haemorrhagic fevers and emerging viruses, Bouaké CHU Laboratory | Project group Epidemology of Highly Pathogenic Microorganisms, Robert Koch-Institute | Chantal Akoua-Koffi et al |
| hCoV-19/BurkinaFaso/588/2020 | EPI_ISL_660475 | | Laboratoire de Microbiologie CHU Souro Sanou | Centre Muraz | Abdoul-Salam Ouedraogo et al |
| hCoV-19/BurkinaFaso/3952/2020 | EPI_ISL_660489 | | Laboratoire de Microbiologie CHU Souro Sanou | Centre Muraz | Abdoul-Salam Ouedraogo et al |
| hCoV-19/Togo/BMC-98-20/2020 | EPI_ISL_1293351 | | Institut National d'Hygiène (INH) | Unité Mixte Internationale TransVIHMI (UMI 233 IRD – U1175 INSERM - Université de Montpellier)IRD (Institut de recherche pour le développement) | Mounerou SALOU et al |
| hCoV-19/Coted-Ivoire/BKE0965/2020 | EPI_ISL_614393 | | Molecular diagnostic unit for viral haemorrhagic fevers and emerging viruses, Bouaké CHU Laboratory | Project group Epidemology of Highly Pathogenic Microorganisms, Robert Koch-Institute | Chantal Akoua-Koffi et al |
| hCoV-19/BurkinaFaso/3570/2020 | EPI_ISL_660468 | | Laboratoire de Microbiologie CHU Souro Sanou | Centre Muraz | Abdoul-Salam Ouedraogo et al |
| hCoV-19/Sudan/SS1708/2020 | EPI_ISL_14620348 | | Port Sudan RT PCR lab | KEMRI-Wellcome Trust Research Programme,Kilifi | Githinji G. et al |
| hCoV-19/Uganda/UG383/2020 | EPI_ISL_2346396 | | MRC/UVRI & LSHTM Uganda Research Unit, Central Public Health Laboratories | MRC/UVRI & LSHTM Uganda Research Unit, Central Public Health Laboratories | Matthew Cotten et al |
| hCoV-19/Uganda/MAKCHScov6/2020 | EPI_ISL_1652054 | | Integrated Biorepository of H3Africa Uganda – IBRH3AU | Molecular Biology Laboratory | Savannah Mwesigwa et al |
| hCoV-19/CentralAfricanRepublic/24014/2020 | EPI_ISL_2281233 | | Insitut Pasteur Bangui | Institut Pasteur de Dakar | Marie-Astrid Vernet et al |
| hCoV-19/Uganda/050/2020 | EPI_ISL_13860467 | | International Organisation for Migration | Mulago Molecular Biology Laboratory | Stephen Kanyerezi et al |
| hCoV-19/Uganda/UG494/2021 | EPI_ISL_2690464 | | MRC/UVRI & LSHTM Uganda Research Unit, Central Public Health Laboratories, Rakai Health Sciences Program | MRC/UVRI & LSHTM Uganda Research Unit, Rakai Health Sciences Program | Matthew Cotten et al |
| hCoV-19/Vietnam/HCMC-05046-02/2021 | EPI_ISL_1366744 | | OUCRU | OUCRU | Nguyen Van Vinh Chau et al |
| hCoV-19/Vietnam/HCMC-05044-02/2021 | EPI_ISL_1366743 | | OUCRU | OUCRU | Nguyen Van Vinh Chau et al |
| hCoV-19/Shanghai/SH0093/2020 | EPI_ISL_416389 | | Shanghai Public Health Clinical Center, Shanghai Medical College, Fudan University | National Research Center for Translational Medicine (Shanghai), Ruijin Hospital affiliated to Shanghai Jiao Tong University School of Medicine & Shanghai Public Health Clinical Center | Shengyue Wang et al |
| hCoV-19/Hangzhou/HZCDC6706/2020 | EPI_ISL_421222 | | Hangzhou Center for Diseases Control and Prevention | Hangzhou Center for Diseases Control and Prevention | Jun Li et al |
| hCoV-19/India/RJ-SMSCOV141/2020 | EPI_ISL_454831 | | SMS Medical College, Jaipur | CSIR Institute of Genomics and Integrative Biology | Sudhir Bhandari et al |
| hCoV-19/Peru/CAL-INS-1997/2021 | EPI_ISL_3376185 | | Laboratorio de Referencial Nacional de Virus Respiratorios | Laboratorio de Referencial Nacional de Virus Respiratorios | Carlos Padilla Rojas et al |
| hCoV-19/India/MP-NCDC-1009329/2021 | EPI_ISL_2459677 | | National Centre for Disease Control (NCDC) Biotechnology Division, Delhi | NCDC Delhi, Biotechnology Division INSACOG | Kalaiarasan Ponnusamy et al |
| hCoV-19/India/MH-CCMB-NIV5/2020 | EPI_ISL_450325 | | NIV Pune | CSIR-Centre for Cellular and Molecular Biology | Dr V A Potdar et al (https://dx.doi.org/10.1101/2020.05.31.126136) |
| hCoV-19/Luxembourg/LNS8881447/2020 | EPI_ISL_744819 | | Laboratoire national de santé, Microbiology, Virology | Laboratoire national de santé, Microbiology, Epidemiology and Microbial Genomics | Anke Wienecke-Baldacchino et al |
| hCoV-19/Germany/BY-MVP-0271/2020 | EPI_ISL_466914 | | Max von Pettenkofer Institute, Virology, National Reference Center for Retroviruses, LMU München | Laboratory for Functional Genome Analysis, Dept. Genomics, Gene Center of the LMU Munich | Max Muenchhoff et al |
| hCoV-19/Denmark/SSI-02/2020 | EPI_ISL_416143 | | Department of Virus and Microbiological Special diagnostics, Statens Serum Institut, Copenhagen, Denmark. | ViFU | Morten Rasmussen et al |
| hCoV-19/Mexico/COA-InDRE-IBT-28845-NC/2020 | EPI_ISL_3463608 | | Instituto de Diagnostico y Referencia Epidemiologicos InDRE_RNLSP | Instituto de Biotecnología de la UNAM | Authors from IBT et al |
| hCoV-19/USA/WY-WYPHL-20037582/2020 | EPI_ISL_2727997 | | Wyoming Public Health Laboratory | Wyoming Public Health Laboratory | Jim Mildenberger et al |
| hCoV-19/India/GJ-GBRC316b/2020 | EPI_ISL_495074 | | Department of MicroBiology, Government Medical College, Surat | Gujarat Biotechnology Research Centre | Pinal Trivedi et al |
| hCoV-19/India/HP-NCDC-609354/2021 | EPI_ISL_2458655 | | National Centre for Disease Control (NCDC) Biotechnology Division, Delhi | NCDC Delhi, Biotechnology Division INSACOG | Priyanka Singh et al |
| hCoV-19/India/UT-NCDC-120991/2021 | EPI_ISL_2448147 | | National Centre for Disease Control (NCDC) Biotechnology Division, Delhi | NCDC Delhi, Biotechnology Division INSACOG | Robin Marwal et al |
| hCoV-19/Austria/Graz-MUG9/2020 | EPI_ISL_437299 | | Diagnostic- and Research Institute of Pathology, Medical University of Graz | Diagnostic- and Research Institute of Pathology, Medical University of Graz | Karl Kashofer et al (https://www.biorxiv.org/content/10.1101/2020.07.15.204339v1) |
| hCoV-19/Jordan/SR-0335/2020 | EPI_ISL_450187 | | Biolab Diagnostic Laboratories | Andersen lab at Scripps Research | Issa Abu-Dayyeh et al |
| hCoV-19/India/OR-RMRC5/2020 | EPI_ISL_455782 | | REGIONAL VRDL,ICMR-RMRC BBSR | Immunogenomics lab, Institute of Life Sciences, Bhubaneswar | Sunil Raghav et al |
| hCoV-19/Indonesia/JI-ITD-5820N/2020 | EPI_ISL_17323235 | | Islamic Hospital Jemursari Surabaya | Institute of Tropical Disease, Universitas Airlangga | Jezzy R Dewantari et al |
| hCoV-19/Indonesia/JB-TFRIC19-R49344/2020 | EPI_ISL_528751 | | Santo Borromeus Hospital | Molecular Genetics Laboratory-Faculty of Medicine-Universitas Padjadjaran; School of Life Sciences and Technology & School of Pharmacy-Institut Teknologi Bandung; Laboratorium Kesehatan Provinsi Jawa Barat | Yunia Sribudiani et al |
| hCoV-19/Timor-Leste/TL31/2020 | EPI_ISL_4761500 | | National Health Laboratory, Timor-Leste | Microbiological Diagnostic Unit - Public Health Laboratory (MDU-PHL) | Soares da Silva et al |
| hCoV-19/Indonesia/JB-EIJK108/2020 | EPI_ISL_889015 | | RS Anna | Eijkman Institute for Molecular Biology, Ministry of Research and Technology/National Agency for Research and Innovation | Iskandar Adnan et al |
| hCoV-19/Indonesia/BA-EIJK-UNWAR-336/2021 | EPI_ISL_2101309 | | RSUD Mangusada | Eijkman Institute for Molecular Biology, National Agency for Research and Innovation; Molecular Biology Laboratory, Faculty Medicine and Health Sciences, Warmadewa University | Hidayat Trimarsanto et al |
| hCoV-19/Indonesia/JK-NIHRD-WGS04064/2021 | EPI_ISL_2854771 | | RSPI Prof. DR. Sulianti Saroso | National Institute of Health Research and Development | Hana Apsari Pawestri et al |
| hCoV-19/Indonesia/JB-GS-WJHL-ITB-WGS049/2021 | EPI_ISL_8435342 | | Bandung Public Health Office | West Java Health Laboratory; School of Life Sciences and Technology, Institut Teknologi Bandung | Azzania Fibriani et al |

| | | | | | |
|---|---|---|---|---|---|
| **hCoV-19/Indonesia/JK-NIHRD-WGS04775/2021** | EPI_ISL_3138830 | | GSI LAB | National Institute of Health Research and Development | Arie Ardiansyah Nugraha et al |
| **hCoV-19/Timor-Leste/TL592/2021** | EPI_ISL_4761638 | | National Health Laboratory, Timor-Leste | Microbiological Diagnostic Unit - Public Health Laboratory (MDU-PHL) | Soares da Silva et al |
| **hCoV-19/Timor-Leste/TL479/2021** | EPI_ISL_4761588 | | National Health Laboratory, Timor-Leste | Microbiological Diagnostic Unit - Public Health Laboratory (MDU-PHL) | Soares da Silva et al |
| **hCoV-19/India/HR-NCDC-909217/2020** | EPI_ISL_2459428 | | National Centre for Disease Control (NCDC) Biotechnology Division, Delhi | NCDC Delhi, Biotechnology Division INSACOG | Robin Marwal et al |
| **hCoV-19/Cambodia/VIR2122581/2021** | EPI_ISL_1096347 | | Cambodian National Public Health Laboratory, National Institute of Public Health | Virology Unit, Institut Pasteur du Cambodge | Sokhoun Yann et al |
| **hCoV-19/Indonesia/SU-USU-02NT/2020** | EPI_ISL_1335896 | | Columbia Asia Hospital Medan | Faculty of Medicine, Universitas Sumatera Utara; Institute of Tropical Disease, Universitas Airlangga | Aldise M Nastri et al |
| **hCoV-19/Afghanistan/IMB07966/2020** | EPI_ISL_1001002 | | Bundeswehr Institute of Microbiology | Bundeswehr Institute of Microbiology | Markus Antwerpen et al |
| **hCoV-19/Jordan/AM-ALSR-2586/2020** | EPI_ISL_635782 | | Biolab Diagnostic Laboratories | Andersen lab at Scripps Research | Issa Abu-Dayyeh et al |
| **hCoV-19/Egypt/CPHL-S8/2021** | EPI_ISL_3274155 | | CPHL/MOH/EGYPT | CPHL/MOH/EGYPT | Wael H. Roshdy/ Mohamed Kamal / Shymaa s. Ahmed/ Ramy Galal/Nancy el guindy/ Amelia nagiub/salma sayed/ Abd Monaem Adel/Galal Mahmoud/Dalia Ramadan/Rabeh ..R. El/Shesheny/ Mohamed A Ali/Mohamed Hassany et al |
| **hCoV-19/Jordan/AM-ALSR-2585/2020** | EPI_ISL_635781 | | Biolab Diagnostic Laboratories | Andersen lab at Scripps Research | Issa Abu-Dayyeh et al |
| **hCoV-19/Indonesia/JB-TFRIC19-R42550/2020** | EPI_ISL_511878 | | Santosa Hospital Bandung Kopo | Laboratorium Kesehatan Provinsi Jawa Barat; Molecular Genetics Laboratory-Faculty of Medicine-Universitas Padjadjaran; School of Life Sciences and Technology & School of Pharmacy-Institut Teknologi Bandung | Ema Rahmawati et al |
| **hCoV-19/SaudiArabia/KAUST-JEDDAH1291/2020** | EPI_ISL_677922 | OV079401.1 | Pathogen Genomics Lab King Abdullah University of Science and Technology(KAUST) | Pathogen Genomics Lab King Abdullah University of Science and Technology(KAUST) | Muhammad Shuaib et al |
| **hCoV-19/SaudiArabia/KAUST-JEDDAH473/2020** | EPI_ISL_512927 | OU994351.1 | Pathogen Genomics Lab King Abdullah University of Science and Technology(KAUST) | Pathogen Genomics Lab King Abdullah University of Science and Technology(KAUST) | Fadwa Alofi et al |
| **hCoV-19/Pakistan/JRCGR-KHI41/2020** | EPI_ISL_779287 | MW447642.1 | Jamil-ur-Rahman Center for Genome Research, Dr. Panjwani Center for Molecular Medicine and Drug Research | Jamil-ur-Rahman Center for Genome Research, Dr. Panjwani Center for Molecular Medicine and Drug Research | Shakeel et al |
| **hCoV-19/SaudiArabia/KAUST-RIYADH1262/2020** | EPI_ISL_678185 | OV090862.1 | Pathogen Genomics Lab King Abdullah University of Science and Technology(KAUST) | Pathogen Genomics Lab King Abdullah University of Science and Technology(KAUST) | Sara Mfarrej et al |
| **hCoV-19/Tunisia/35507/2020** | EPI_ISL_1116464 | | Laboratoire de Microbiologie- CHU Habib Bourguiba – Sfax | Laboratoire des Procédés de Criblage Moléculaire et Cellulaire-Centre de Biotechnologie de Sfax | Souissi et al |
| **hCoV-19/Sweden/20VPC572808/2020** | EPI_ISL_3792227 | | The Public Health Agency of Sweden | The Public Health Agency of Sweden | Maximilian Riess et al |
| **hCoV-19/Austria/CeMM2652/2020** | EPI_ISL_934587 | | Department of Laboratory Medicine, Division of Clinical Virology, University of Medicine, Vienna | Bergthaler laboratory, CeMM Research Center for Molecular Medicine of the Austrian Academy of Sciences | Lukas Endler et al |
| **hCoV-19/Hungary/HM-0201-71/2021** | EPI_ISL_1041211 | | Hungarian Defence Forces Military Medical Centre | National Laboratory of Virology, Szentágothai Research Centre | Endre Gábor Tóth et al |
| **hCoV-19/England/MILK-A1D464/2020** | EPI_ISL_2735307 | | Respiratory Virus Unit, Microbiology Services Colindale, Public Health England | COVID-19 Genomics UK (COG-UK) Consortium | PHE Covid Sequencing Team et al |
| **hCoV-19/Tunisia/Q-6811/2021** | EPI_ISL_10101207 | | Institut Pasteur de Tunis | Institut Pasteur de Dakar | Chouikha Anissa et al |
| **hCoV-19/Tunisia/1821/2021** | EPI_ISL_803120 | | Laboratory of Microbiology, National Reference Lab, Charles Nicolle Hospital; 2-University of Tunis ElManar, Faculty of Medicine of Tunis, LR99ES09, Tunis, Tunisia | Clinical and Experimental Pharmacology Lab, LR16SP02, National Center of Pharmacovigilance, University of Tunis El Manar, Tunis, Tunisia. 2- Neurodegenerative diseases and psychiatric troubles, LR18SP03, Razi Hospital, University of Tunis El Manar, Tunis, Tunisia. 3- Ministry of Health, National Observatory of New and Emerging Diseases, 1006, Tunis, Tunisia | Ilhem Boutiba-Ben Boubaker et al |
| **hCoV-19/Tunisia/H-4648/2021** | EPI_ISL_10101353 | | Institut Pasteur de Tunis | Institut Pasteur de Dakar | Chouikha Anissa et al |
| **hCoV-19/Tunisia/6841/2021** | EPI_ISL_11266008 | | Laboratory of Clinical Virology, Institut Pasteur de Tunis | The African Centre of Excellence for Genomics of Infectious Diseases (ACEGID) | Anissa Chouikha et al |
| **hCoV-19/Singapore/1388/2020** | EPI_ISL_645115 | | National Public Health Laboratory, National Centre for Infectious Diseases | National Public Health Laboratory, National Centre for Infectious Diseases | Tze Minn Mak et al |
| **hCoV-19/CentralAfricanRepublic/23749/2020** | EPI_ISL_2281229 | | Insitut Pasteur Bangui | Institut Pasteur de Dakar | Marie-Astrid Vernet et al |
| **hCoV-19/Bulgaria/21BG-NC_A000031/2020** | EPI_ISL_2081898 | | National Center of Infectious and Parasitic Diseases | National Center of Infectious and Parasitic Diseases | Alexiev et al |
| **hCoV-19/CaboVerde/PortoNovo_13453/2020** | EPI_ISL_1711657 | | Virology Laboratory of São Vicente | Institut Pasteur de Dakar | Nadia Rodrigues et al |
| **hCoV-19/Tahiti/HCL021013320801/2021** | EPI_ISL_1371901 | | C H DE LA POLYNESIE FRANCAISE | CNR Virus des Infections Respiratoires - France SUD | Antonin Bal et al |
| **hCoV-19/Tunisia/S-0010/2021** | EPI_ISL_17018814 | | Laboratoire de Virologie Clinique, Institut Pasteur de Tunis | Laboratoire de Virologie Clinique, Institut Pasteur de Tunis | Samar Ben Halima et al |
| **hCoV-19/Indonesia/JB-GS-WJHL-ITB-WGS067/2021** | EPI_ISL_8435356 | | Bandung Barat Public Health Office | West Java Health Laboratory; School of Life Sciences and Technology, Institut Teknologi Bandung | Azzania Fibriani et al |
| **hCoV-19/Indonesia/JBUA-2-FKUA/2020** | EPI_ISL_1533097 | | Diagnostic and Research Center of Infectious Diseases, Medical Faculty, Andalas University | Diagnostic and Research Center of Infectious Diseases, Medical Faculty, Andalas University | Andani Eka Putra et al |
| **hCoV-19/TrinidadandTobago/1116/2020** | EPI_ISL_2230686 | | Trinidad Public Health Laboratory | Carrington Lab, Department of PreClinical Sciences, Faculty of Medical Sciences, The University of the West Indies | Nikita S. D. Sahadeo et al |
| **hCoV-19/Myanmar/99790-20/2020** | EPI_ISL_3252980 | | National Health Laboratory, Ministry Of Health And Sports, The Republic Of The Union Of Myanmar | Pathogen Genomics Center, National Institute of Infectious Diseases | Htay Htay Tin et al |
| **hCoV-19/Yunnan/03/2020** | EPI_ISL_632935 | | Department of Acute Infectious Diseases Control and Prevention, Yunnan Provincial Center for Disease Control and Prevention | Department of Acute Infectious Diseases Control and Prevention, Yunnan Provincial Center for Disease Control and Prevention | Meiling Zhang et al |
| **hCoV-19/Thailand/SamutSakhon-CONI-0370/2020** | EPI_ISL_733564 | ON879051.1 | Institute for Urban Disease Control and Prevention | COVID-19 Network Investigations (CONI) Alliance | Kamolthip Atsawawaranunt et al |
| **hCoV-19/Myanmar/DSMRC-002/2021** | EPI_ISL_849736 | MW491244.1 | Special Operations Medical Research Division, Defence Services Medical Research Centre | Special Operations Medical Research Division, Defence Services Medical Research Centre | Oo et al |
| **hCoV-19/Myanmar/DSMRC-001/2021** | EPI_ISL_833041 | MW491243.1 | Defence Services Medical Research Center, Biological Research Laboratory | Defence Services Medical Research Center, Biological Research Laboratory | Khine Zaw Oo et al |
| **hCoV-19/Yunnan/RLC21/2021** | EPI_ISL_13005084 | | Department of Acute Infectious Diseases Control and Prevention, Yunnan Provincial Center for Disease Control and Prevention | Department of Acute Infectious Diseases Control and Prevention, Yunnan Provincial Center for Disease Control and Prevention | Meiling Zhang et al |
| **hCoV-19/Yunnan/RLC68/2021** | EPI_ISL_13005132 | | Department of Acute Infectious Diseases Control and Prevention, Yunnan Provincial Center for Disease Control and Prevention | Department of Acute Infectious Diseases Control and Prevention, Yunnan Provincial Center for Disease Control and Prevention | Meiling Zhang et al |
| **hCoV-19/UnitedArabEmirates/3357/2020** | EPI_ISL_859750 | | BTC, Khalifa University | BTC, Khalifa University | Al Safar et al |
| **hCoV-19/UnitedArabEmirates/3314/2020** | EPI_ISL_859732 | | BTC, Khalifa University | BTC, Khalifa University | Al Safar et al |
| **hCoV-19/India/OR-ILSGS00117/2020** | EPI_ISL_1662256 | | Virology Lab, Ispat General Hospital, Rourkela | Institute of Life Sciences - INSACOG | Sunil K. Raghav et al |
| **hCoV-19/India/HR-IMT-CB130/2020** | EPI_ISL_547592 | | Civil Hospital, Panchkula | CSIR-Institute of Microbial Technology | Kanika Bansal et al |
| **hCoV-19/UnitedArabEmirates/3746/2020** | EPI_ISL_859834 | | BTC, Khalifa University | BTC, Khalifa University | Al Safar et al |
| **hCoV-19/HongKong/HKPU-05444/2020** | EPI_ISL_1019750 | | Department of Health Technology and Informatics, The Hong Kong Polytechnic University | Department of Health Technology and Informatics, The Hong Kong Polytechnic University | Gilman Kit-Hang Siu et al |
| **hCoV-19/HongKong/CUHK-NC7806/2020** | EPI_ISL_3066198 | | Department of Microbiology | Department of Microbiology | Zigui Chen et al |
| **hCoV-19/HongKong/HKPU-09606/2021** | EPI_ISL_2713095 | | Department of Health Technology and Informatics, The Hong Kong Polytechnic University | Department of Health Technology and Informatics, The Hong Kong Polytechnic University | Gilman Kit-Hang Siu et al |
| **hCoV-19/HongKong/HKPU-11254/2021** | EPI_ISL_2860793 | | Department of Health Technology and Informatics, The Hong Kong Polytechnic University | Department of Health Technology and Informatics, The Hong Kong Polytechnic University | Gilman Kit-Hang Siu et al |
| **hCoV-19/India/MP-ICMR-NIV-INSACOG-SEQ-3187/2020** | EPI_ISL_15038582 | | ICMR-National Institute of Virology - INSACOG | NIV Influenza | Dr. Varsha Potdar et al |
| **hCoV-19/India/HP-170/2020** | EPI_ISL_2099616 | | National Centre for Disease Control | CDFD -INSACOG | To be addded later et al |
| **hCoV-19/India/PB-NCDC-430923/2021** | EPI_ISL_4493008 | | National Centre for Disease Control (NCDC) Biotechnology Division, Delhi - INSACOG | NCDC Delhi, Biotechnology Division -INSACOG | Kalaiarasan Ponnusamy et al |
| **hCoV-19/India/GJ-NCCS-VOC-435/2021** | EPI_ISL_1533742 | | National Centre For Cell Science | National Centre For Cell Science – INSACOG | Dhiraj Paul et al |
| **hCoV-19/SaudiArabia/KAUST-MADINAH1269/2020** | EPI_ISL_678004 | OV090036.1 | Pathogen Genomics Lab King Abdullah University of Science and Technology(KAUST) | Pathogen Genomics Lab King Abdullah University of Science and Technology(KAUST) | Sara Mfarrej et al |
| **hCoV-19/India/KL-GENESCOV00659/2020** | EPI_ISL_7212435 | | INSACOG-IGIB and Kerala State SARS-COV-2 Genome Surveillance Programme (GENESCOV2) | INSACOG and GENESCOV2 at CSIR Institute of Genomics and Integrative Biology | INSACOG-IGIB et al |
| **hCoV-19/India/KA-041628_2_S54_R1_001/2020** | EPI_ISL_1708366 | | Thindlu PHC | inStem NCBS - INSACOG | Uma Ramakrishnan Dasaradhi Palakodeti Aswin SaiNarain et al |
| **hCoV-19/India/DL-60881-CSIR-IGIB/2020** | EPI_ISL_636769 | | National Centre for Disease control (NCDC) | NCDC/CSIR-IGIB | Mahesh S. Dhar1* et al |
| **hCoV-19/Iran/Ghilan-M30-241/2020** | EPI_ISL_1014681 | | National Influenza Center, Virology Department | National Influenza Center | V Salimi et al |
| **hCoV-19/Norway/7260/2020** | EPI_ISL_796651 | | Medical Microbiology Unit, Department for Laboratory Medicine, Drammen Hospital, Vestre Viken Health Trust, | Norwegian Institute of Public Health, Department of Virology | Kathrine Stene-Johansen et al |
| **hCoV-19/India/OR-ILSGS00118/2021** | EPI_ISL_1662257 | | Virology Lab, Ispat General Hospital, Rourkela | Institute of Life Sciences - INSACOG | Sunil K. Raghav et al |
| **hCoV-19/Pakistan/JRCGR_SCV-61_IBGE-13/2020** | EPI_ISL_2151357 | MW960285.1 | Jamil-ur-Rahman Center for Genome Research, Dr. Panjwani Center for Molecular Medicine and Drug Research | Jamil-ur-Rahman Center for Genome Research, Dr. Panjwani Center for Molecular Medicine and Drug Research | Irfan et al |
| **hCoV-19/Romania/DJ50176/2020** | EPI_ISL_6436483 | | Medical Genetics Laboratory, Regional Centre of Medical Genetics, Emergency County Hospital Craiova | Medical Genetics Laboratory, Regional Centre of Medical Genetics, Emergency County Hospital Craiova | Anca-Lelia (Riza) Costache et al |
| **hCoV-19/SaudiArabia/KAUST-MADINAH1116/2020** | EPI_ISL_513120 | OV081787.1 | Pathogen Genomics Lab King Abdullah University of Science and Technology(KAUST) | Pathogen Genomics Lab King Abdullah University of Science and Technology(KAUST) | Fathia Ben Rached et al |
| **hCoV-19/India/TG-CCMB-CIA7834/2021** | EPI_ISL_3935245 | | Mapmygenome | CSIR-Centre for Cellular and Molecular Biology - INSACOG | Sumedha Avadhanula et al |
| **hCoV-19/India/MZ-ZMC264C/2020** | EPI_ISL_1419077 | | INSACOG-Mizoram | National Institute of Biomedical Genomics – INSACOG | Arindam Maitra et al A |
| **hCoV-19/India/KA-NIMH-SEQ-201/2021** | EPI_ISL_1385818 | | BBMP Urban PHC | INSACOG-KA, NIMHANS | Chitra Pattabiraman et al |
| **hCoV-19/India/TG-CCMB-AZ523/2020** | EPI_ISL_1380374 | | CSIR-Centre for Cellular and Molecular Biology | CSIR-Centre for Cellular and Molecular Biology-INSACOG | Payel Mukherjee et al B |
| **hCoV-19/India/AP-CCMB-CIA2372/2021** | EPI_ISL_2441094 | | AP SSO | CSIR-Centre for Cellular and Molecular Biology-INSACOG | Sofia Banu et al |
| **hCoV-19/India/TG-CCMB-AW553/2020** | EPI_ISL_1380292 | | CSIR-Centre for Cellular and Molecular Biology | CSIR-Centre for Cellular and Molecular Biology-INSACOG | Payel Mukherjee et al B |
| **hCoV-19/India/KA-043885_2_S101_R1_001/2020** | EPI_ISL_1708395 | | Kodigehalli PHC | inStem NCBS - INSACOG | Uma Ramakrishnan Dasaradhi Palakodeti Aswin SaiNarain et al |
| **hCoV-19/India/TG-RFCH00250_CIB0660/2021** | EPI_ISL_6696788 | | Osmania Medical College | CSIR-Centre for Cellular and Molecular Biology | Lamuk Zaveri et al |
| **hCoV-19/India/AP-CS0845/2020** | EPI_ISL_862304 | | Kurnool Medical College (KMC) | CSIR Institute of Genomics and Integrative Biology | Pallavali Roja Rani et al |
| **hCoV-19/India/ILSGS00532/2020** | EPI_ISL_1663257 | | BRLSABVM Government Medical College, Rajnandgaon, Chhattisgarh | Institute of Life Sciences - INSACOG | Sunil K. Raghav et al |
| **hCoV-19/India/ILSGS00576/2021** | EPI_ISL_1663277 | | CIIMS, Bilaspur, Chhattisgarh | Institute of Life Sciences - INSACOG | Sunil K. Raghav et al |
| **hCoV-19/India/AP-CS0789/2020** | EPI_ISL_862276 | | Kurnool Medical College (KMC) | CSIR Institute of Genomics and Integrative Biology | Pallavali Roja Rani et al |
| **hCoV-19/India/KA-045947_2_S114_R1_001/2020** | EPI_ISL_1708408 | | MS Palya PHC | inStem NCBS - INSACOG | Uma Ramakrishnan Dasaradhi Palakodeti Aswin SaiNarain et al |
| **hCoV-19/India/KA-NIMH-SEQ-277/2021** | EPI_ISL_1595894 | | BBMP Urban PHC | INSACOG-KA, NIMHANS | Chitra Pattabiraman et al |
| **hCoV-19/India/UT-AR34/2020** | EPI_ISL_508182 | | All india institute of Medical Sciences Rishikesh | National Institute of Biomedical Genomics | Arindam Maitra et al A |
| **hCoV-19/India/HR-NIBG-TF98/2020** | EPI_ISL_653913 | | Translational Health Science and Technology Institute | National Institute of Biomedical Genomics | Arindam Maitra et al A |
| **hCoV-19/India/HR-TF7/2020** | EPI_ISL_508508 | | Translational Health Science and Technology Institute | National Institute of Biomedical Genomics | Arindam Maitra et al A |
| **hCoV-19/India/MZ-ZMC3043A/2020** | EPI_ISL_1419081 | | INSACOG-Mizoram | National Institute of Biomedical Genomics – INSACOG | Arindam Maitra et al A |
| **hCoV-19/India/MZ-ZMC3048A/2020** | EPI_ISL_1419083 | | INSACOG-Mizoram | National Institute of Biomedical Genomics – INSACOG | Arindam Maitra et al A |
| **hCoV-19/India/UP-AR20/2020** | EPI_ISL_508168 | | All india institute of Medical Sciences Rishikesh | National Institute of Biomedical Genomics | Arindam Maitra et al A |
| **hCoV-19/India/DL-AIIMS16667/2020** | EPI_ISL_17463917 | | Virology Laboratory, AIIMS New Delhi | Virology Laboratory, AIIMS New Delhi | Aashish Choudhary et al |
| **hCoV-19/India/JK-ICMR-65965/2020** | EPI_ISL_1073014 | | VRDL, Sher-I-Kashmir Institute of Medical Sciences, Srinagar, Jammu & Kashmir | Indian Council of Medical Research-National Institute of Virology, Microbial Containment Complex | Pragya D. Yadav et al |
| **hCoV-19/Bhutan/RCDC0020-SV200576/2020** | EPI_ISL_16984368 | | AFRIMS | AFRIMS | Chonticha Klungthong et al |
| **hCoV-19/India/UP-MRU70695/2020** | EPI_ISL_862815 | | Multidisciplinary Research Unit, DHR-ICMR, Institute of Medical Sciences, Banaras Hindu University, Varanasi | Multidisciplinary Research Unit, DHR-ICMR, Institute of Medical Sciences, Banaras Hindu University, Varanasi | Royana Singh et al |
| **hCoV-19/India/HR-NCDC-909237/2020** | EPI_ISL_2459332 | | National Centre for Disease Control (NCDC) Biotechnology Division, Delhi | NCDC Delhi, Biotechnology Division INSACOG | Robin Marwal et al |
| **hCoV-19/India/RJ-CCMB-CIA6185/2020** | EPI_ISL_3453815 | | RUHS College of Medical Sciences | CSIR-Centre for Cellular and Molecular Biology - INSACOG | Dr. Gaurav Dalela et al |
| **hCoV-19/India/PB-ICMR-NIV-INSACOG-GSEQ-572/2021** | EPI_ISL_1704658 | | ICMR-National Institute of Virology - INSACOG | NIV Influenza | Potdar et al B |
| **hCoV-19/India/MZ-ZMC2889J/2020** | EPI_ISL_1419054 | | INSACOG-Mizoram | National Institute of Biomedical Genomics – INSACOG | Arindam Maitra et al A |
| **hCoV-19/India/JH-ICMR-NIV-INSACOG-SEQ-2572/2020** | EPI_ISL_15038482 | | ICMR-National Institute of Virology - INSACOG | NIV Influenza | Dr. Varsha Potdar et al |
| **hCoV-19/India/un-SRIHER-wuhan_IME-WH05/2021** | EPI_ISL_2350916 | MZ305017.1 | msc medical bioinformatics, sri ramachandra institute of higher education and research | msc medical bioinformatics, sri ramachandra institute of higher education and research | Monisha et al |
| **hCoV-19/India/GJ-ICMR-NIV-INSACOG-SEQ-1793/2020** | EPI_ISL_15038469 | | ICMR-National Institute of Virology - INSACOG | NIV Influenza | Dr. Varsha Potdar et al |
| **hCoV-19/India/OR-ILSGS00286/2021** | EPI_ISL_1662265 | | REGIONAL VRDL, ICMR-RMRC BBSR | Institute of Life Sciences - INSACOG | Sunil K. Raghav et al |
| **hCoV-19/India/MZ-ZMC6345G/2021** | EPI_ISL_2521627 | | INSACOG-Mizoram | National Institute of Biomedical Genomics – INSACOG | Arindam Maitra et al A |
| **hCoV-19/India/HP-168/2020** | EPI_ISL_2099613 | | National Centre for Disease Control | CDFD -INSACOG | To be addded later et al |
| **hCoV-19/India/GJ-ICMR-NIV-INSACOG-VOC-653/2020** | EPI_ISL_1704290 | | ICMR-National Institute of Virology - INSACOG | NIV Influenza | Potdar et al B |
| **hCoV-19/India/AN-ICMR-NIV-INSACOG-SEQ-2742/2020** | EPI_ISL_15038550 | | ICMR-National Institute of Virology - INSACOG | NIV Influenza | Dr. Varsha Potdar et al |
| **hCoV-19/India/LEH-NCDC-1009107/2021** | EPI_ISL_2459881 | | National Centre for Disease Control (NCDC) Biotechnology Division, Delhi | NCDC Delhi, Biotechnology Division INSACOG | Mahesh S Dhar et al |
| **hCoV-19/India/LA-NCDC-609208/2021** | EPI_ISL_2458629 | | National Centre for Disease Control (NCDC) Biotechnology Division, Delhi | NCDC Delhi, Biotechnology Division INSACOG | Priyanka Singh et al |
| **hCoV-19/India/UP-NBRI-N55-TEH861/2020** | EPI_ISL_516983 | | King Georges Medical University | CSIR-National Botanical Research Institute | Priti Prasad et al |
| **hCoV-19/India/MN-ICMR-NIV-INSACOG-SEQ-3110/2020** | EPI_ISL_15038507 | | ICMR-National Institute of Virology - INSACOG | NIV Influenza | Dr. Varsha Potdar et al |
| **hCoV-19/Canada/BC-BCCDC-7666/2020** | EPI_ISL_974784 | | BCCDC Public Health Laboratory | BCCDC Public Health Laboratory | Prystajecky Natalie et al |
| **hCoV-19/India/JK-ICMR-172729/2020** | EPI_ISL_1073016 | | VRDL, Sher-I-Kashmir Institute of Medical Sciences, Srinagar, Jammu & Kashmir | Indian Council of Medical Research-National Institute of Virology, Microbial Containment Complex | Pragya D. Yadav et al |
| **hCoV-19/India/MN-ICMR-NIV-INSACOG-SEQ-3120/2020** | EPI_ISL_15038508 | | ICMR-National Institute of Virology - INSACOG | NIV Influenza | Dr. Varsha Potdar et al |
| **hCoV-19/Bhutan/RCDC0019-SV200575/2020** | EPI_ISL_16984365 | | AFRIMS | AFRIMS | Chonticha Klungthong et al |
| **hCoV-19/India/un-SMSMC-791502/2021** | EPI_ISL_857395 | | S.M.S.Medical College, Jaipur, Rajasthan | S.M.S.Medical College, Jaipur, Rajasthan | Dr. Bharti Malhotra et al |
| **hCoV-19/India/TR-ICMR-NIV-INSACOG-SEQ-1390/2020** | EPI_ISL_15038536 | | ICMR-National Institute of Virology - INSACOG | NIV Influenza | Dr. Varsha Potdar et al |
| **hCoV-19/Bhutan/RCDC0051-SV200607/2020** | EPI_ISL_16984395 | | AFRIMS | AFRIMS | Chonticha Klungthong et al |
| **hCoV-19/India/ML-ICMR-NIV-INSACOG-SEQ-2421/2020** | EPI_ISL_15038498 | | ICMR-National Institute of Virology - INSACOG | NIV Influenza | Dr. Varsha Potdar et al |
| **hCoV-19/India/ML-ICMR-NIV-INSACOG-SEQ-2484/2020** | EPI_ISL_15038502 | | ICMR-National Institute of Virology - INSACOG | NIV Influenza | Dr. Varsha Potdar et al |
| **hCoV-19/India/AS-CSIR-NEIST-INSACOG-NEIST15-CIDIN0941/2020** | EPI_ISL_8889204 | | Centre for Infectious Diseases, CSIR-NEIST | Centre for Infectious Diseases, CSIR-NEIST | Romi Wahengbam et al |
| **hCoV-19/DRC/236/2020** | EPI_ISL_420032 | | Viral Respiratory Lab, National Institute for Biomedical Research (INRB) | Pathogen Sequencing Lab, National Institute for Biomedical Research (INRB) | Placide Mbala-Kingebeni et al A (https://virological.org/t/phylogenetic-analysis-of-sars-cov-2-in-drc/528) |
| **hCoV-19/Canada/UN-CRCHUM-PreFreeze-9/2020** | EPI_ISL_667766 | | Microbiology, Infectious Diseases and Immunology, Centre de Recherche du Centre Hospitalier de l'Universite de Montreal | Microbiology, Infectious Diseases and Immunology, Centre de Recherche du Centre Hospitalier de l'Universite de Montreal | Benoit et al |
| **hCoV-19/Peru/ICA-INS-108/2020** | EPI_ISL_536506 | | Instituto Nacional de Salud | Laboratorio de Infecciones Respiratorias Agudas | Eduardo Juscamayta Lopez et al |
| **hCoV-19/Peru/LIM-INS-458/2020** | EPI_ISL_1111356 | | Laboratorio de Referencia Nacional de Virus Respiratorio. Instituto Nacional de Salud Perú | Laboratorio de Referencia Nacional de Enteropatógenos. Instituto Nacional de Salud del Perú | Ronnie Gavilan Chavez et al |
| **hCoV-19/Croatia/52123_ID1647_40/2020** | EPI_ISL_994739 | | "Dr. Andrija Stampar" Teaching Institute of Public Health, Department of Clinical Microbiology | Institute of Applied Genomics | Jasmina Vranes et al |
| **hCoV-19/Ecuador/USFQ-196/2020** | EPI_ISL_527812 | | Institute of Microbiology, Universidad San Francisco de Quito | Institute of Microbiology, Universidad San Francisco de Quito | Belén Prado-Vivar et al |
| **hCoV-19/Maldives/MAV11950/2020** | EPI_ISL_3452727 | | Indira Gandhi Memorial Hospital | Indira Gandhi Memorial Hospital | Dr. Milza Abdul Muhsin et al |
| **hCoV-19/Maldives/MAV09481/2020** | EPI_ISL_3275392 | | Indira Gandhi Memorial Hospital | Indira Gandhi Memorial Hospital | Dr. Milza Abdul Muhsin et al |
| **hCoV-19/Maldives/MAV04305/2020** | EPI_ISL_3275389 | | Indira Gandhi Memorial Hospital | Indira Gandhi Memorial Hospital | Dr. Milza Abdul Muhsin et al |
| **hCoV-19/Maldives/MAV01979/2020** | EPI_ISL_3275385 | | Indira Gandhi Memorial Hospital | Indira Gandhi Memorial Hospital | Dr. Milza Abdul Muhsin et al |
| **hCoV-19/USA/IA-CDC-2-4044433/2020** | EPI_ISL_1446728 | MW850137.1 | IA State Hygienic Laboratory | Centers for Disease Control and Prevention Division of Viral Diseases, Pathogen Discovery | Mili Sheth et al |
| **hCoV-19/Kuwait/JAH3092399/2021** | EPI_ISL_2557309 | | MOH - Jaber Al-Ahmad Hospital (Innovation Research Laboratory) | MOH - Jaber Al-Ahmad Hospital (Innovation Research Laboratory) | Salman Al-Sabah et al |
| **hCoV-19/Nigeria/BCVL-21954/2021** | EPI_ISL_4026347 | | Biorepository and Clinical Virology Laboratory | Northwestern University - Center for Pathogen Genomics and Microbial Evolution | Ramon Lorenzo-Redondo et al |
| **hCoV-19/SouthSudan/UG433/2021** | EPI_ISL_2450805 | | National Public Health Laboratory, Ministry of Health, Ministry of Health, Republic of South Sudan | South Sudan Ministry of Health, WHO South Sudan, MRC/UVRI & LSHTM Uganda Research Unit | Joseph Francis Wamala et al |
| **hCoV-19/Ghana/WACCBIP-GS1040/2021** | EPI_ISL_3869495 | | WACCBIP, University of Ghana, Accra, Ghana | WACCBIP, University of Ghana, Volta Road, Legon-Accra, Ghana | Collins M. Morang'a et al |
| **hCoV-19/Coted-Ivoire/IPCI-cv347/2021** | EPI_ISL_10070656 | | Environment and Health Department, Institut Pasteur of Côte d'Ivoire, Abidjan, Côte d'Ivoire | Molecular Genetics Platform, Institut Pasteur of Côte d'Ivoire, Abidjan, Côte d'Ivoire | Aristide Berenger Ako AKO et al |
| **hCoV-19/Sudan/ILRI_M03144/2021** | EPI_ISL_17971660 | | University of Kassala | International Livestock Research Institute | Samuel O. Oyola et al |
| **hCoV-19/India/GJ-ICMR-NIV-INSACOG-SEQ-1771/2020** | EPI_ISL_15038565 | | ICMR-National Institute of Virology - INSACOG | NIV Influenza | Dr. Varsha Potdar et al |
| **hCoV-19/India/OR-ILSGS00011/2020** | EPI_ISL_1063809 | | REGIONAL VRDL, ICMR-RMRC BBSR | Immunogenomics lab, Institute of Life Sciences, Bhubaneswar | Sunil K. Raghav et al |
| **hCoV-19/Mexico/QUE-InDRE_401/2020** | EPI_ISL_1054971 | | Instituto de Diagnostico y Referencia Epidemiologicos INDRE_RNLSP | Instituto de Diagnostico y Referencia Epidemiologicos (INDRE) | Claudia Wong-Arambula et al |
| **hCoV-19/DRC/ILRI_COVM00357/2020** | EPI_ISL_12257144 | | Molecular Biology laboratory, Université Evangélique en Afrique | International Livestock Research Institute | Samuel O. Oyola et al |
| **hCoV-19/Bhutan/RCDC0002-SV210018/2020** | EPI_ISL_16984321 | | AFRIMS | AFRIMS | Chonticha Klungthong et al |
| **hCoV-19/Kenya/IPRCOV02788/2020** | EPI_ISL_17500845 | | IPR COVID-19 TESTING AND RESEARCH CENTER | IPR COVID-19 TESTING AND RESEARCH CENTER | Kamau J et al |
| **hCoV-19/Mexico/QUE-IBT-IMSS-436/2020** | EPI_ISL_1301626 | | Laboratorio Central de Epidemiología IMSS | Instituto de Biotecnología de la UNAM | Authors from IBT et al |
| **hCoV-19/Nigeria/LA-CV370/2020** | EPI_ISL_730031 | | Nigeria Centre for Disease Control (NCDC) | African Centre of Excellence for Genomics of Infectious Diseases (ACEGID), Redeemer's University, Ede, Osun State, Nigeria | Oluniyi P.E. et al |
| **hCoV-19/USA/MI-MDHHS-SC20511/2020** | EPI_ISL_460559 | OL756646.1 | Michigan Department of Health and Human Services, Bureau of Laboratories | Michigan Department of Health and Human Services, Bureau of Laboratories | Blankenship HM et al |
| **hCoV-19/India/MP-NCDC-4859/2020** | EPI_ISL_436457 | | National Centre for Disease control (NCDC) | NCDC/CSIR-IGIB | Pramod Kumar# et al (https://dx.doi.org/10.1101/2020.06.04.128751) |
| **hCoV-19/SouthAfrica/KRISP-0035/2020** | EPI_ISL_467441 | | NHLS-IALCH | KRISP, KZN Research Innovation and Sequencing Platform | Giandhari J et al |
| **hCoV-19/Bolivia/26932/2020** | EPI_ISL_837555 | | Centro Nacional de Enfermedades Tropicales (CENETROP) | Laboratory of Respiratory Viruses and Measles, Oswaldo Cruz Institute, FIOCRUZ | Paola Resende et al |
| **hCoV-19/Bolivia/SAN-UPCH-1067/2020** | EPI_ISL_5147843 | | Laboratorio de Diagnóstico e Investigación BIOSCIENCE SRL | Laboratorio de Genómica Microbiana, Universidad Peruana Cayetano Heredia | Pablo Tsukayama et al |
| **hCoV-19/Hungary/MH-13211/2020** | EPI_ISL_526225 | | Hungarian Defence Forces Military Medical Centre | National Laboratory of Virology, Szentágothai Research Centre | Endre Gábor Tóth et al |
| **hCoV-19/Mexico/AGU-InDRE_F11767B_S1089/2020** | EPI_ISL_1516781 | | LESP Aguascalientes | Instituto de Diagnostico y Referencia Epidemiologicos (INDRE) | Claudia Wong-Arambula et al |
| **hCoV-19/Germany/HH-HPI-p2859/2020** | EPI_ISL_959946 | | University Medical Center Hamburg Eppendorf | Heinrich Pette Institute, Leibniz Institute for Experimental Virology | Alexis Robitaille et al |
| **hCoV-19/Kosovo/ChVir23117/2020** | EPI_ISL_1027642 | | Department of Microbiology, National Institute for Public Health of Kosova | Charité Universitätsmedizin Berlin, Institut für Virologie | Victor M Corman et al |
| **hCoV-19/Mexico/VER-InDRE-IBT-96/2020** | EPI_ISL_1301514 | | Instituto de Diagnostico y Referencia Epidemiologicos InDRE_RNLSP | Instituto de Biotecnología de la UNAM | Authors from IBT et al |
| **hCoV-19/Belize/CML-7/2021** | EPI_ISL_4296316 | | Central Medical Laboratory | Baylor College of Medicine | Alexander Kneubehl et al |
| **hCoV-19/Belize/CML-17/2021** | EPI_ISL_4296325 | | Central Medical Laboratory | Baylor College of Medicine | Alexander Kneubehl et al |
| **hCoV-19/Belize/CML-36/2021** | EPI_ISL_4296344 | | Central Medical Laboratory | Baylor College of Medicine | Alexander Kneubehl et al |
| **hCoV-19/Malaysia/UNIMAS-15693R/2020** | EPI_ISL_2367475 | | Ministry of Health Hospitals | Institute of Health and Community Medicine | Chan Chia Jui et al |
| **hCoV-19/Malaysia/IMR_WC365760/2020** | EPI_ISL_944100 | | Institute for Medical Research, Infectious Disease Research Centre, National Institutes of Health, Ministry of Health Malaysia | Institute for Medical Research, Infectious Disease Research Centre, National Institutes of Health, Ministry of Health Malaysia | Suppiah J et al |
| **hCoV-19/Malaysia/4867/2021** | EPI_ISL_2769425 | | Department of Medical Microbiology, University Malaya Medical Centre | Department of Medical Microbiology, Faculty of Medicine, University of Malaya | Yoong Min CHONG et al |
| **hCoV-19/Romania/DJ60465/2021** | EPI_ISL_6195914 | | Medical Genetics Laboratory, Regional Centre of Medical Genetics, Emergency County Hospital Craiova | Medical Genetics Laboratory, Regional Centre of Medical Genetics, Emergency County Hospital Craiova | Anca-Lelia (Riza) Costache et al |
| **hCoV-19/India/TG-CCMB-CIA5118/2021** | EPI_ISL_2775125 | | Mapmygenome | CSIR-Centre for Cellular and Molecular Biology-INSACOG | Payel Mukherjee et al B |
| **hCoV-19/Honduras/19984/2021** | EPI_ISL_4029400 | | Laboratorio Nacional de Vigilancia de la Salud – Sección de Virologia | Genomics and Proteomics Departament, Gorgas Memorial Institute For Health Studies | Mitzi Castro et al |

| | | | | | |
|---|---|---|---|---|---|
| **hCoV-19/ElSalvador/69396-LNSP/2021** | EPI_ISL_18902319 | | Region Oriental | Laboratorio Nacional de Salud Publica | Laboratorio de Secuenciacion y genomica del Laboratorio Nacional de Salud Publica de El Salvador - Instituto Nacional de Salud et al |
| **hCoV-19/Sweden/20-53086/2020** | EPI_ISL_648136 | | The Public Health Agency of Sweden | The Public Health Agency of Sweden | Anna-Malin Linde et al |
| **hCoV-19/Togo/BMC-4-20/2020** | EPI_ISL_1499727 | | Institut National d'Hygiène (INH) | Unité Mixte Internationale TransVIHMI (UMI 233 IRD – U1175 INSERM - Université de Montpellier)IRD (Institut de recherche pour le développement) | Mounerou SALOU et al |
| **hCoV-19/Turkey/CTF_17032020_003/2020** | EPI_ISL_2015129 | | IU-Cerrahpasa, Cerrahpasa School of Medicine, COVID-19 Lab | IU-Cerrahpasa, Cerrahpasa School of Medicine, COVID-19 Lab | Mert Kuskucu et al |
| **hCoV-19/Afghanistan/IMB07960/2020** | EPI_ISL_1000999 | | Bundeswehr Institute of Microbiology | Bundeswehr Institute of Microbiology | Markus Antwerpen et al |
| **hCoV-19/Turkey/DEN-PAU02/2020** | EPI_ISL_707939 | | Pamukkale University Hospital | Pamukkale University Department of Medical Genetics | Onur TOKGUN et al. |
| **hCoV-19/Kashgar/IVDC-01-01/2020** | EPI_ISL_850950 | | National Institute for Viral Disease Control and Prevention, China CDC | National Institute for Viral Disease Control and Prevention, China CDC | Xiang Zhao et al |
| **hCoV-19/USA/OR-OHSU-0179/2020** | EPI_ISL_468122 | | OHSU Lab Services Molecular Microbiology Lab | Oregon SARS-CoV-2 Genome Sequencing Center | Brendan L. O'Connell et al |
| **hCoV-19/USA/NY-NYCPHL-000746/2020** | EPI_ISL_632127 | | Flushing Hospital Medical Center | New York City Public Health Laboratory | Jade Wang et al |
| **hCoV-19/Egypt/C-CEIRS-19MOH/2020** | EPI_ISL_605780 | | CEIRS Data Processing and Coordinating Center, St. Jude Center of Excellence for Influenza Research and Surveillance (CEIRS) | CEIRS Data Processing and Coordinating Center, St. Jude Center of Excellence for Influenza Research and Surveillance (CEIRS) | Roshdy et al |
| **hCoV-19/Chile/RM-29191/2020** | EPI_ISL_445325 | | HOSPITAL DR.SOTERO DEL RIO | Instituto de Salud Publica de Chile | Andrés E Castillo et al |
| **hCoV-19/Venezuela/Tru723/2020** | EPI_ISL_15104901 | | IVIC | Laboratorio de Virologia Molecular | Rossana C Jaspe et al |
| **hCoV-19/Curacao/CW-RIVM-11654/2021** | EPI_ISL_1014581 | | National Institute for Public Health and the Environment (RIVM) | National Institute for Public Health and the Environment (RIVM) | Adam Meijer et al |
| **hCoV-19/Curacao/CW-RIVM-21729/2020** | EPI_ISL_1014554 | | National Institute for Public Health and the Environment (RIVM) | National Institute for Public Health and the Environment (RIVM) | Adam Meijer et al |
| **hCoV-19/USA/ND-NDDH-0097/2020** | EPI_ISL_1035868 | | North Dakota Department of Health, Public Health Laboratory | North Dakota Department of Health, Public Health Laboratory | Lisa Wingerter et al |
| **hCoV-19/Taiwan/KMU-1/2020** | EPI_ISL_4169856 | | Dr.Chiaho Shih | Dr.Chiaho Shih | An-Ting Liou et al |
| **hCoV-19/Canada/PE-NML-3090/2020** | EPI_ISL_15184927 | | PE-QEH Dept of Lab Medicine | National Microbiology Laboratory (NML) | Vanessa Arseneau et al |
| **hCoV-19/Iran/GRC-T11763/2021** | EPI_ISL_2365354 | | Kariminejad-Najmabadi Pathology & Genetics Center, Tehran, Iran | Genetics Research Center, University of Social Welfare and Rehabilitation Sciences | Zohreh Fattahi et al |
| **hCoV-19/Haiti/49220162/2020** | EPI_ISL_14186946 | | LABORATOIRE NATIONAL DE SANTÉ PUBLIQUE | Laboratory of Parasitic Diseases, Systems Genomics Section, National Institute of Allergy and Infectious Diseases, National Institutes of Health | Samson Marseille et al |
| **hCoV-19/Togo/BMC-15540-20/2020** | EPI_ISL_2278896 | | Institut National d'Hygiène (INH) | Unité Mixte Internationale TransVIHMI (UMI 233 IRD – U1175 INSERM - Université de Montpellier)IRD (Institut de recherche pour le développement) | Mounerou SALOU et al |
| **hCoV-19/India/GJ-GBRC316a/2020** | EPI_ISL_495073 | | Department of MicroBiology, Government Medical College, Surat | Gujarat Biotechnology Research Centre | Monika Gandhi et al |
| **hCoV-19/Qatar/QA-QU_12-10-D10/2020** | EPI_ISL_1713214 | | Ministry of Public Health / Hamad Medical Corporation | Biomedical Research Center (BRC), Qatar University / Qatar Genome Project (QGP) | BRC: Fatiha M. Benslimane et al |
| **hCoV-19/Morocco/FMP-76/2021** | EPI_ISL_1116555 | | Laboratoire central de Virologie | Laboratoire de Biotechnologie | Ghizlane EL Amin et al |
| **hCoV-19/Peru/ANC-INS-2907/2021** | EPI_ISL_3769567 | | Laboratorio de Referencia Nacional de Virus Respiratorios. Centro Nacional de Salud Publica. Instituto Nacional de Salud Peru. | Laboratorio de Referencia Nacional de Virus Respiratorios. Centro Nacional de Salud Publica. Instituto Nacional de Salud Peru. | Carlos Padilla Rojas et al |
| **hCoV-19/USA/IL-IDPH-SAN-S-000418/2020** | EPI_ISL_8009216 | | Illinois Department of Public Health | Gagnon Lab, Southern Illinois University | Keith Gagnon et al |
| **hCoV-19/CentralAfricanRepublic/RCA-050/2022** | EPI_ISL_10854848 | | Laboratoire National de Biologie Clinique et de Santé Publique, Bangui, République Centrafricaine | Pathogen Genomics Lab, National Institute for Biomedical Research (INRB) | Placide Mbala-Kingebeni et al B |
| **hCoV-19/Ghana/TRA-2159/2021** | EPI_ISL_6944033 | | National Influenza Centre | National Influenza Centre | William K. Ampofo et al |
| **hCoV-19/Colombia/ATL-I61/2021** | EPI_ISL_4417643 | | LSP de Atlantico | Centro de Investigaciones en Microbiología y Biotecnología-UR (CIMBIUR), Facultad de Ciencias Naturales, Universidad del Rosario, Bogotá, Colombia | Marina Muñoz et al |
| **hCoV-19/Panama/GMI-PAM34722/2021** | EPI_ISL_11695786 | | Gorgas Memorial Institute of Health Studies | Gorgas Memorial Institute of Health Studies | Claudia Gonzalez et al |
| **hCoV-19/Venezuela/Bol7115/2021** | EPI_ISL_16895336 | | Instituto Nacional de Higiene Rafael Rangel | Laboratorio de Virología Molecular | Rossana C Jaspe et al |
| **hCoV-19/Aruba/AW-RIVM-38124/2021** | EPI_ISL_2476286 | | National Institute for Public Health and the Environment (RIVM) | National Institute for Public Health and the Environment (RIVM) | Adam Meijer et al |
| **hCoV-19/Venezuela/Lar644/2021** | EPI_ISL_11629599 | | IVIC | Laboratorio de Virologia Molecular | Rossana C Jaspe et al |
| **hCoV-19/CostaRica/CABANA-CR-UCR-INCIENSA-ITV-UBA-Uda-UGA-IMSS-UPC-2103-697384/2021** | EPI_ISL_7711707 | | Area De Salud Heredia Cubujuchi | Universidad de Costa Rica e Instituto Costarricense de Investigacion y Ensenanza en Nutricion y Salud | Brenes Hebleen et al |
| **hCoV-19/Colombia/VAC-ICE-SCOV-042/2021** | EPI_ISL_4963428 | | Laboratorio COLCAN | Laboratorio de biotecnología, Universidad Icesi | Maria I. Gutierrez Lopez et al |
| **hCoV-19/Ecuador/NIC-INSPI-353708/2021** | EPI_ISL_6208263 | | INSPI-CRN DE INFLUENZA Y OTROS VIRUS RESPIRATORIOS | INSPI-CRN DE INFLUENZA Y OTROS VIRUS RESPIRATORIOS | Alfredo Bruno et al |
| **hCoV-19/SaintVincentandtheGrenadines/101240/2021** | EPI_ISL_6941451 | | The Caribbean Public Health Agency | Carrington Lab, Department of Preclinical Sciences, Faculty of Medical Sciences, The University of the West Indies, St Augustine Campus | Nikita S. D. Sahadeo et al |
| **hCoV-19/USA/MA-CDC-QDX25472477/2021** | EPI_ISL_4370327 | OK250604.1 | Quest Diagnostics Incorporated | Centers for Disease Control and Prevention Division of Viral Diseases, Pathogen Discovery | Dakota Howard et al |
| **hCoV-19/Chile/AR-34396/2020** | EPI_ISL_447119 | | HOSPITAL DR.HERNAN HENRIQUEZ ARAVENA | Instituto de Salud Publica de Chile | Andrés E Castillo et al |
| **hCoV-19/Chile/MA-CADIUMAG-18/2020** | EPI_ISL_681686 | | Molecular Medicine Laboratory, University of Magallanes | Centro Asistencial Docente y de Investigacion, Universidad de Magallanes | Jorge González et al |
| **hCoV-19/USA/IA-Mayo21806/2020** | EPI_ISL_13687359 | | Dept. Clinical Microbiology/Mayo Clinic | Dept. Clinical Microbiology/Mayo Clinic | Pritt Bobbi Norgan Andrew et al |
| **hCoV-19/Turkey/HSGM-4807/2021** | EPI_ISL_1171933 | | Ministry of Health Turkey | Ministry of Health Turkey | Fatma Bayrakdar et al |
| **hCoV-19/Lebanon/LAU-R19/2020** | EPI_ISL_8311716 | | LAUMC-Rizk Hospital | Multi-Omics Laboratory - School of Pharmacy - Lebanese American University | Nancy Fayad et al |
| **hCoV-19/Madagascar/IP-01861/2020** | EPI_ISL_508863 | | Virology Unit, Institut Pasteur de Madagascar | Virology Unit, Institut Pasteur de Madagascar | Christian Ranaivoson et al |
| **hCoV-19/Canada/AB-10548/2020** | EPI_ISL_854465 | | Alberta Precision Labs (APL) | Alberta Precision Labs (APL) | Gordon P et al |
| **hCoV-19/Canada/AB-28007/2020** | EPI_ISL_806198 | | Alberta Precision Labs (APL) | Alberta Precision Labs (APL) | Gordon P et al |
| **hCoV-19/Canada/AB-ABPHL-02446/2020** | EPI_ISL_2162998 | | Alberta Precision Labs (APL) | Public Health Agency of Canada (PHAC) National Microbiology Laboratory | Buss et al |
| **hCoV-19/Croatia/52114_ID1647_31/2020** | EPI_ISL_994740 | | "Dr. Andrija Stampar" Teaching Institute of Public Health, Department of Clinical Microbiology | Institute of Applied Genomics | Jasmina Vranes et al |
| **hCoV-19/Bahrain/BAH-22/2020** | EPI_ISL_483563 | | Kingdom of Bahrein Ministry of Health | Erasmus Medical Center | Bas Oude Munnink et al |
| **hCoV-19/Algeria/48642/2020** | EPI_ISL_15928078 | | NIC, Viral Respiratory Unit | NIC, Viral Respiratory Unit | Mohamed Amine Beloufa et al |
| **hCoV-19/Malta/BAL-Sliema-5/2020** | EPI_ISL_576121 | MW079422.1 | Laboratory, The Bio Arte Limited | Laboratory, The Bio Arte Limited | Biazzo et al |
| **hCoV-19/Gibraltar/203360727/2020** | EPI_ISL_547442 | | Microbiology, Department of Pathology, St. Bernard's Hospital, Gibraltar Health Authority | Respiratory Virus Unit, Microbiology Services Colindale, Public Health England | PHE Covid Sequencing Team et al |
| **hCoV-19/GuineaBissau/51LNSPGB/2020** | EPI_ISL_2158597 | | Laboratório de Biologia Molecular Jean Piaget | MRCG at LSHTM, Genomics lab | Aladje Balde et al |
| **hCoV-19/France/IDF-HB-112006085806/2020** | EPI_ISL_833192 | | Hôpital Bichat Claude Bernard, Laboratoire de Virologie | IAME UMR1137 Inserm, Université de Paris, Hôpital Bichat | Antoine Bridier et al |
| **hCoV-19/Benin/418200/2021** | EPI_ISL_4566852 | | Laboratoire des Fièvres Hémorragiques Virales du Benin | Institut für Virologie - Institute of Virology - Charite | Anges Yadouleton et al |
| **hCoV-19/Togo/C4528/2020** | EPI_ISL_1575299 | | Laboratoire Biolim/FSS/UL | Unité Mixte Internationale TransVIHMI (UMI 233 IRD – U1175 INSERM - Université de Montpellier) IRD (Institut de recherche pour le développement) | Mounerou SALOU et al |
| **hCoV-19/Zambia/ZMB-9163/2020** | EPI_ISL_977469 | | University of Zambia, School of Veterinary Medicine | UNZAVET and PATH | Mulenga Mwenda-Chimfwembe et al |
| **hCoV-19/Ethiopia/AHRI-002/2020** | EPI_ISL_3071136 | | Armauer Hansen Research Institute | Armauer Hansen Research Institute | Mr. Dawit Hailu Alemayehu et al |
| **hCoV-19/Ethiopia/CERI-KRISP-K025614/2021** | EPI_ISL_5429015 | | Addis Ababa University | CERI, Centre for Epidemic Response and Innvoation, Stellenbosch University and KRISP, KZN Research Innovation and Sequencing Platform, UKZN. | Abay Sisay et al |
| **hCoV-19/Ethiopia/AH-001/2020** | EPI_ISL_2970353 | MZ172407.1 | Bacterial and Viral Diseases & Biotechnology and Bioinformatics, Armauer Hansen Research Institute | Bacterial and Viral Diseases & Biotechnology and Bioinformatics, Armauer Hansen Research Institute | Alemayehu et al |
| **hCoV-19/Somalia/CV1238/2020** | EPI_ISL_2308259 | | National Institute of Health (NIH) - Federal Government of Somalia | African Centre of Excellence for Genomics of Infectious Diseases (ACEGID), Redeemer's University | Olawoye et al |
| **hCoV-19/Niger/28/2020** | EPI_ISL_2281807 | | Centre de Recherche Médicale et Sanitaire (CERMES) | Institut Pasteur de Dakar | Lagare Adamou et al |
| **hCoV-19/Norway/3617/2020** | EPI_ISL_2491573 | | Medical Microbiology Unit, Department for Laboratory Medicine, Drammen Hospital, Vestre Viken Health Trust, | Norwegian Institute of Public Health, Department of Virology | Kathrine Stene-Johansen et al |
| **hCoV-19/NewZealand/20CV0310/2020** | EPI_ISL_622800 | | Canterbury Health Laboratories | Institute of Environmental Science and Research (ESR) | Xiaoyun Ren et al |
| **hCoV-19/Singapore/117/2020** | EPI_ISL_443209 | | National Public Health Laboratory, National Centre for Infectious Diseases | National Public Health Laboratory, National Centre for Infectious Diseases | Mak Tze Minn et al |
| **hCoV-19/Togo/BMC-1014-20/2020** | EPI_ISL_1415425 | | Institut National d'hygiène | "Unité Mixte Internationale TransVIHMI (UMI 233 IRD – U1175 INSERM - Université de Montpellier) IRD (Institut de recherche pour le développement)" | Mounerou SALOU et al |
| **hCoV-19/Tunisia/TUN_ADAGE_EZ504/2020** | EPI_ISL_7055354 | MT955170.1 | Laboratoire de microbiologie | Molecular and Cellular Screening Process Laboratory | Souissi et al |
| **hCoV-19/Coted-Ivoire/BKE0992/2020** | EPI_ISL_614394 | | Molecular diagnostic unit for viral haemorrhagic fevers and emerging viruses, Bouaké CHU Laboratory | Project group Epidemology of Highly Pathogenic Microorganisms, Robert Koch-Institute | Chantal Akoua-Koffi et al |
| **hCoV-19/Israel/990307712/2020** | EPI_ISL_447263 | | Microbiology laboratory, Assuta Ashdod University-Affiliated Hospital | Stern Lab | Stern Lab et al (https://dx.doi.org/10.1101/2020.05.21.20104521) |
| **hCoV-19/NanChang/JX216/2020** | EPI_ISL_417420 | | Jiangxi province Center for Disease Control and Prevention | Jiangxi province Center for Disease Control and Prevention | Li jian Xiong et al |
| **hCoV-19/Kazakhstan/38716/2020** | EPI_ISL_454575 | | Laboratory of virology, National Center of Expertise | Laboratory of molecular-genetic research, National Center for Expertise, Kazakhstan National Center for Biotechnology, Kazakhstan | Abdaliyev Askar et al |
| **hCoV-19/SouthKorea/KDCA0465/2020** | EPI_ISL_850181 | | Division of Emerging Infectious Diseases, Bureau of Infectious Diseases Diagnosis Control, Korea Disease Control and Prevention Agency | Division of Emerging Infectious Diseases, Bureau of Infectious Diseases Diagnosis Control, Korea Disease Control and Prevention Agency | Ae Kyung Park et al |
| **hCoV-19/Argentina/PAIS-A0359/2020** | EPI_ISL_1396163 | | Laboratorio de Virología del Hospital de Niños Dr. Ricardo Gutierrez | Área de Secuenciación del Laboratorio de Virología del Hospital de Niños Dr. Ricardo Gutierrez on behalf of 'Proyecto Argentino Interinstitucional de genomica de SARS-CoV-2' (PAIS Consortium) | Alexay et al |
| **hCoV-19/Israel/ISR_NH_GD1_0714/2020** | EPI_ISL_594159 | | Israel Institute for Biological Research | Israel Institute for Biological Research | Galia Zaide et al |
| **hCoV-19/SouthKorea/KDCA0098/2020** | EPI_ISL_747335 | | Division of Emerging Infectious Diseases, Bureau of Infectious Diseases Diagnosis Control, Korea Disease Control and Prevention Agency | Division of Emerging Infectious Diseases, Bureau of Infectious Diseases Diagnosis Control, Korea Disease Control and Prevention Agency | Ae Kyung Park et al |
| **hCoV-19/SouthKorea/KCDC2364/2020** | EPI_ISL_514767 | | Division of Viral Diseases, Center for Laboratory Control of Infectious Diseases, Korea Centers for Diseases Control and Prevention | Division of Viral Diseases, Center for Laboratory Control of Infectious Diseases, Korea Centers for Diseases Control and Prevention | Jeong-Min Kim et al |
| **hCoV-19/SouthKorea/KDCA0489/2020** | EPI_ISL_850201 | | Division of Emerging Infectious Diseases, Bureau of Infectious Diseases Diagnosis Control, Korea Disease Control and Prevention Agency | Division of Emerging Infectious Diseases, Bureau of Infectious Diseases Diagnosis Control, Korea Disease Control and Prevention Agency | Ae Kyung Park et al |
| **hCoV-19/SouthKorea/KDCA0765/2020** | EPI_ISL_850442 | | Division of Emerging Infectious Diseases, Bureau of Infectious Diseases Diagnosis Control, Korea Disease Control and Prevention Agency | Division of Emerging Infectious Diseases, Bureau of Infectious Diseases Diagnosis Control, Korea Disease Control and Prevention Agency | Ae Kyung Park et al |
| **hCoV-19/SouthKorea/KDCA0946/2020** | EPI_ISL_959535 | | Division of Emerging Infectious Diseases, Bureau of Infectious Diseases Diagnosis Control, Korea Disease Control and Prevention Agency | Division of Emerging Infectious Diseases, Bureau of Infectious Diseases Diagnosis Control, Korea Disease Control and Prevention Agency | Ae Kyung Park et al |
| **hCoV-19/SouthKorea/KDCA0992/2020** | EPI_ISL_994708 | | Division of Emerging Infectious Diseases, Bureau of Infectious Diseases Diagnosis Control, Korea Disease Control and Prevention Agency | Division of Emerging Infectious Diseases, Bureau of Infectious Diseases Diagnosis Control, Korea Disease Control and Prevention Agency | Ae Kyung Park et al |
| **hCoV-19/SouthKorea/KDCA2144/2021** | EPI_ISL_1319000 | | Division of Emerging Infectious Diseases, Bureau of Infectious Diseases Diagnosis Control, Korea Disease Control and Prevention Agency | Division of Emerging Infectious Diseases, Bureau of Infectious Diseases Diagnosis Control, Korea Disease Control and Prevention Agency | Ae Kyung Park et al |
| **hCoV-19/Senegal/003/2020** | EPI_ISL_418206 | | Institut Pasteur Dakar | Institut Pasteur de Dakar | Ndongo Dia et al |
| **hCoV-19/Israel/2089852/2020** | EPI_ISL_447362 | | Clinical Virology Unit, Hadassah Hebrew University Medical Center | Stern Lab | Stern Lab et al (https://dx.doi.org/10.1101/2020.05.21.20104521) |
| **hCoV-19/Harbin/IVDC-01-01/2020** | EPI_ISL_850946 | | National Institute for Viral Disease Control and Prevention, China CDC | National Institute for Viral Disease Control and Prevention, China CDC | Xiang Zhao et al |
| **hCoV-19/USA/WA-UW-8873/2020** | EPI_ISL_570224 | | UW Virology Lab | UW Virology Lab | Pavitra Roychoudhury et al |
| **hCoV-19/Botswana/BOT2283/2020** | EPI_ISL_560385 | | National Health Laboratory | Botswana Institute for Technology Research and Innovation | Kefentse Arnold Tumedi et al |
| **hCoV-19/NewZealand/20CV0298/2020** | EPI_ISL_622795 | | Canterbury Health Laboratories | Institute of Environmental Science and Research (ESR) | Xiaoyun Ren et al |
| **hCoV-19/USA/CO-CDPHE-2006260794/2020** | EPI_ISL_6602612 | OL641037.1 | Colorado Department of Public Health and Environment | Colorado Department of Public Health and Environment | Laura Bankers et al |
| **hCoV-19/CostaRica/INC-0096/2020** | EPI_ISL_682238 | | AREA DE SALUD LA CRUZ | Inciensa, Instituto Costarricense de Investigación y Enseñanza en Nutrición y Salud | Francisco Duarte et al |
| **hCoV-19/USA/MS-BIE-CAT00302/2020** | EPI_ISL_6582805 | OL589102.1 | Ochsner Health | BioInfoExperts | Amy Feehan et al |
| **hCoV-19/USA/OH-21AM-032CO01329/2021** | EPI_ISL_2312846 | | The Ohio State University Applied Microbiology Services Laboratory | The Ohio State University Applied Microbiology Services Laboratory | Seth A. Faith PhD et al |
| **hCoV-19/Greece/6899/2020** | EPI_ISL_2301777 | | Laboratory of Molecular Biology, Mamatsio General Hospital of Kozani | Greek Genome Center, Biomedical Research Foundation of the Academy of Athens (BRFAA) | Emmanouil Athanasiadis et al |
| **hCoV-19/USA/MA-CDC-STM-A100363/2021** | EPI_ISL_876700 | MW523817.1 | Helix/Illumina | Genomics and Discovery, Respiratory Viruses Branch, Division of Viral Diseases, Centers for Disease Control and Prevention | Peter W. Cook et al |
| **hCoV-19/USA/AL-Limestone-JDXC/2020** | EPI_ISL_872465 | MW861622.1 | Eurofins Diatherix | Hudsonalpha Genome Sequencing Center | Jane Grimwood et al |
| **hCoV-19/USA/MO-CDC-3953597-001/2020** | EPI_ISL_2422254 | MZ348327.1 | Springfield Greene County Health Dept. | Genomics and Discovery, Respiratory Viruses Branch, Division of Viral Diseases, Centers for Disease Control and Prevention | Anna Kelleher et al |
| **hCoV-19/Belize/CML-131/2021** | EPI_ISL_4296435 | | Central Medical Laboratory | Baylor College of Medicine | Alexander Kneubehl et al |
| **hCoV-19/USA/AR-CDC-2-3693771/2020** | EPI_ISL_747068 | MW406520.1 | Respiratory Viruses Branch, Centers for Disease Control and Prevention | Respiratory Viruses Branch, Centers for Disease Control and Prevention | Queen et al |
| **hCoV-19/Guyana/50832/2021** | EPI_ISL_2230700 | | The Caribbean Public Health Agency | Carrington Lab, Department of PreClinical Sciences, Faculty of Medical Sciences, The University of the West Indies | Nikita S. D. Sahadeo et al |
| **hCoV-19/Indonesia/BA-EIJK-UNWAR-033/2021** | EPI_ISL_2101054 | | Molecular Biology Laboratory, Faculty Medicine and Health Sciences, Warmadewa University | Eijkman Institute for Molecular Biology, National Agency for Research and Innovation; Molecular Biology Laboratory, Faculty Medicine and Health Sciences, Warmadewa University | Sukma Oktavianthi et al |
| **hCoV-19/Ecuador/USFQ-1384/2021** | EPI_ISL_2361463 | | Institute of Microbiology, Universidad San Francisco de Quito | Institute of Microbiology, Universidad San Francisco de Quito | Sully Márquez et al |
| **hCoV-19/USA/NM-UNM-EDB00180/2021** | EPI_ISL_1064968 | ON612204.1 | University of New Mexico Hospital | Center for Global Health, University of New Mexico Health Sciences Center | Daryl Domman et al |
| **hCoV-19/USA/HI-H200015/2020** | EPI_ISL_752611 | | State Laboratories Division, Hawaii State Department of Health | State Laboratories Division, Hawaii State Department of Health | Pamela O'Brien et al |
| **hCoV-19/CaboVerde/Praia_0338/2021** | EPI_ISL_1711661 | | Virology Laboratory of Praia | Institut Pasteur de Dakar | Sanders Kevin et al |
| **hCoV-19/Israel/CVL5786/2020** | EPI_ISL_516919 | | Israel Central Virology laboratory | Israel Central Virology laboratory | Neta Zuckerman et al |
| **hCoV-19/Israel/CVL_40898/2020** | EPI_ISL_3002494 | | Infection Prevention & Control Unit | Central Virology laboratory, Ministry of Health | Neta Zuckerman et al |
| **hCoV-19/Israel/CVL_40903/2020** | EPI_ISL_3003644 | | Infection Prevention & Control Unit | Central Virology laboratory, Ministry of Health | Neta Zuckerman et al |
| **hCoV-19/Denmark/DCGC-6735/2020** | EPI_ISL_622355 | | Department of Virus and Microbiological Special Diagnostics, Statens Serum Institut, Denmark | Albertsen lab, Department of Chemistry and Bioscience, Aalborg University, Denmark | Thomas Yssing Michaelsen on behalf of the Danish Covid-19 Genome Consortium et al |
| **hCoV-19/Taiwan/CGMH-CGU-34/2020** | EPI_ISL_667807 | | Laboratory Medicine | Department of Laboratory Medicine, Lin-Kou Chang Gung Memorial Hospital, Taoyuan, Taiwan | Kuo-Chien Tsao et al |
| **hCoV-19/Colombia/ANT-GUV-92256/2020** | EPI_ISL_447768 | | Instituto Nacional de Salud, Bogotá, Colombia | Grupo de Investigaciones Microbiológicas-UR (GIMUR), Departamento de Biología, Facultad de Ciencias Naturales, Universidad del Rosario, Bogotá, Colombia Instituto Nacional de Salud, Bogotá, Colombia Icahn School of Medicine at Mount Sinai, New York, USA | Juan David Ramírez et al (https://www.medrxiv.org/content/10.1101/2020.06.11.20125799v1) |
| **hCoV-19/Niger/66-20/2020** | EPI_ISL_2281783 | | Centre de Recherche Médicale et Sanitaire (CERMES) | Institut Pasteur de Dakar | Lagare Adamou et al |
| **hCoV-19/USA/MT-Walk-777/2020** | EPI_ISL_10834885 | | Montana Public Health Laboratory | Walk Lab Montana State University | Seth Walk et al |
| **hCoV-19/Somalia/CV1236/2020** | EPI_ISL_2308257 | | National Institute of Health (NIH) - Federal Government of Somalia | African Centre of Excellence for Genomics of Infectious Diseases (ACEGID), Redeemer's University | Olawoye et al |
| **hCoV-19/Kenya/C53548/2020** | EPI_ISL_3049397 | OK487780.1 | KEMRI-Wellcome Trust Research Programme,Kilifi | KEMRI-Wellcome Trust Research Programme,Kilifi | Githinji G. et al |
| **hCoV-19/Haiti/34957/2020** | EPI_ISL_15279685 | | Laboratoire National de Santé Publique | Laboratory of Respiratory Viruses and Measles, Oswaldo Cruz Institute, FIOCRUZ | Paola Resende et al |
| **hCoV-19/USA/AR-20U122MD00289/2020** | EPI_ISL_17130200 | | Department of Pathology and Laboratory Services, University of Arkansas for Medical Sciences | Department of Pathology and Laboratory Services, University of Arkansas for Medical Sciences | "Harley et al |
| **hCoV-19/Georgia/Tb-7856/2020** | EPI_ISL_481380 | MT786442.1 | Department for Virology, Molecular Biology and Genome Research, R. G. Lugar Center for Public Health Research, National Center for Disease Control and Public Health (NCDC) of Georgia. | Department for Virology, Molecular Biology and Genome Research, R. G. Lugar Center for Public Health Research, National Center for Disease Control and Public Health (NCDC) of Georgia. | Ana Papkiauri et al |
| **hCoV-19/Austria/CeMM0617/2020** | EPI_ISL_583616 | | Institut für Virologie am Department für Hygiene, Mikrobiologie und Public Health | Bergthaler laboratory, CeMM Research Center for Molecular Medicine of the Austrian Academy of Sciences | Alexandra Popa et al |
| **hCoV-19/Lebanon/LAU-R16/2020** | EPI_ISL_8317207 | | LAUMC-Rizk Hospital | Multi-Omics Laboratory - School of Pharmacy - Lebanese American University | Nancy Fayad et al |
| **hCoV-19/USA/TN-CDC-4023988-001/2020** | EPI_ISL_1373885 | MW863944.1 | Vanderbilt University Medical Center | Pathogen Discovery, Respiratory Viruses Branch, Division of Viral Diseases, Centers for Disease Control and Prevention | Jing Zhang et al |
| **hCoV-19/USA/MD-HP09774-PIDBIQAEDK/2020** | EPI_ISL_5412311 | | Johns Hopkins Hospital Department of Pathology | Johns Hopkins Hospital Department of Pathology | C. Paul Morris et al |
| **hCoV-19/Guatemala/ASI01026/2020** | EPI_ISL_3446724 | | Laboratorio Nacional de Salud | Asociación de Salud Integral/Clínica Familiar "Luis Ángel García" | Eduardo Arathoon et al |
| **hCoV-19/Thailand/BIDI-63039985__2019/2020** | EPI_ISL_12717927 | | Bamrasnaradura Infectious Disease Institute | Molecular HIV and Emerging Infectious Diseases Laboratory, Bamrasnaradura Infectious Diseases Institute (BIDI), Department of Disease Control (DDC), Ministry of Public Health (MOPH) | Sumonmal Uttayamakul et al |
| **hCoV-19/USA/DC-DFS-PHL-0086/2020** | EPI_ISL_804926 | MZ488256.1 | DC Public Health Lab/ Dept. of Forensic Sciences | DC Public Health Lab/ Dept. of Forensic Sciences | Scott Nguyen et al |
| **hCoV-19/Lithuania/MR-LUHS-Eilnr174/2020** | EPI_ISL_636840 | | Lithuanian University of Health Sciences Hospital, Department of Laboratory Medicine | Lithuanian University of Health Sciences, Molecular cardiology lab. | Lukas Zemaitis et al |
| **hCoV-19/USA/DC-DFS-PHL-0248/2020** | EPI_ISL_1295621 | MZ487710.1 | DC Public Health Lab/ Dept. of Forensic Sciences | DC Public Health Lab/ Dept. of Forensic Sciences | Scott Nguyen et al |
| **hCoV-19/USA/IA-10859/2020** | EPI_ISL_475190 | | Nebraska Public Health Laboratory | UNMC COVID-19 Response Team | UNMC COVID-19 Response Team et al |
| **hCoV-19/Bahamas/18013-GMI/2021** | EPI_ISL_11680121 | | Reference Laboratory of the Ministry of Health Royal Victoria Gardens | Genomics and Proteomics Departament, Gorgas Memorial Institute For Health Studies | Indira Martins et al |
| **hCoV-19/DominicanRepublic/202102998_602399/2021** | EPI_ISL_6437538 | ON022760.1 | Laboratorio nacional de salud publica | Viral Special Pathogens Branch, Centers for Disease Control and Prevention | Isaac Miguel et al |
| **hCoV-19/Qatar/QA-WCMQ_FD18163171/2020** | EPI_ISL_1714439 | | Ministry of Public Health / Hamad Medical Corporation | Weill Cornell Medical College - Qatar (WCM-Q), Genomics Core Laboratory / Qatar Genome Project (QGP) | WCMQ: Ayeda A. Ahmed et al |
| **hCoV-19/Qatar/QA-QU_12-10-C3/2020** | EPI_ISL_1713209 | | Ministry of Public Health / Hamad Medical Corporation | Biomedical Research Center (BRC), Qatar University / Qatar Genome Project (QGP) | BRC: Fatiha M. Benslimane et al |
| **hCoV-19/SriLanka/Genelabs-CA4755/2020** | EPI_ISL_1185245 | | Genelabs Medical (Pvt) Ltd | Genelabs Medical (Pvt) Ltd | Chandanamali Punchihewa et al |
| **hCoV-19/SriLanka/DRVL0073/2020** | EPI_ISL_18108258 | | Diagnostic and Research Virology Laboratory | Diagnostic and Research Virology Laboratory | Bushran N Iqbal et al |
| **hCoV-19/SriLanka/CMC70852/2021** | EPI_ISL_872394 | | Centre for Dengue Research and AICBU, Department of Immunology and Molecular Medicine | Centre for Dengue Research and AICBU, Department of Immunology and Molecular Medicine | Chandima Jeewandara et al |
| **hCoV-19/SriLanka/CMC68909/2021** | EPI_ISL_1717044 | | Centre for Dengue Research and AICBU, Department of Immunology and Molecular Medicine | Centre for Dengue Research and AICBU, Department of Immunology and Molecular Medicine | Chandima Jeewandara et al |
| **hCoV-19/CaboVerde/Santiago_31612/2021** | EPI_ISL_8350856 | | Virology Laboratory of Praia | Institut Pasteur de Dakar | Sanders Kevin et al |
| **hCoV-19/CaboVerde/Santiago_32025/2021** | EPI_ISL_8350865 | | Virology Laboratory of Praia | Institut Pasteur de Dakar | Sanders Kevin et al |
| **hCoV-19/USA/MT-USAFSAM-S1853/2021** | EPI_ISL_2383345 | | US Air Force School of Aerospace Medicine | US Air Force School of Aerospace Medicine | Anthony Fries et al |
| **hCoV-19/Cameroon/Yaounde-21V-06070/2021** | EPI_ISL_1790107 | | Centre Pasteur du Cameroun | Institut Pasteur de Dakar | Njouom Richard et al |
| **hCoV-19/Guam/GU-CDC-2-3906254/2021** | EPI_ISL_1273061 | MW766880.1 | Guam Public Health Laboratory | Centers for Disease Control and Prevention Division of Viral Diseases, Pathogen Discovery | Krista Queen et al |
| **hCoV-19/USA/CA-CZB-14885/2020** | EPI_ISL_739556 | MW564883.1 | Orange County Public Health Lab | Chan-Zuckerberg Biohub | CZB Cliahub Consortium et al |
| **hCoV-19/USA/WV-WVU-WV031830/2021** | EPI_ISL_1423094 | MZ500021.1 | WVU Rapid Development Laboratory | WVU and Marshall University Combined Genomics Core Facilities | "James Denvir et al |
| **hCoV-19/Cambodia/VIR2107865/2021** | EPI_ISL_1096140 | | Cambodian National Public Health Laboratory, National Institute of Public Health | Virology Unit, Institut Pasteur du Cambodge | Sokhoun Yann et al |
| **hCoV-19/Cambodia/VIR2107864/2021** | EPI_ISL_1096139 | | Cambodian National Public Health Laboratory, National Institute of Public Health | Virology Unit, Institut Pasteur du Cambodge | Sokhoun Yann et al |
| **hCoV-19/Botswana/R8B61_BHP_AAB26174/2020** | EPI_ISL_2233463 | | Botswana Harvard HIV Reference Laboratory | Botswana Harvard HIV Reference Laboratory | Sikhulile Wonderful T. Choga et al |
| **hCoV-19/Angola/CERI-KRISPK013576/2021** | EPI_ISL_2609550 | | Instituto Nacional de Investigação em Saúde | KRISP, KZN Research Innovation and Sequencing Platform | Morais J et al |
| **hCoV-19/Angola/CERI-KRISP-K013577/2021** | EPI_ISL_2492885 | | Instituto Nacional de Investigação em Saúde | CERI, Centre for Epidemic Response and Innvoation, Stellenbosch University and KRISP, KZN Research Innovation and Sequencing Platform, UKZN. | Morais J et al |
| **hCoV-19/SouthAfrica/VIDA-KRISP-K012974/2020** | EPI_ISL_2494752 | | Vaccines and Infectious Diseases Analytics Research Unit (VIDA) | KRISP, KZN Research Innovation and Sequencing Platform | Baillie Vicky et al |
| **hCoV-19/Angola/CERI-KRISP-K013424/2021** | EPI_ISL_2492895 | | Instituto Nacional de Investigação em Saúde | CERI, Centre for Epidemic Response and Innvoation, Stellenbosch University and KRISP, KZN Research Innovation and Sequencing Platform, UKZN. | Morais J et al |
| **hCoV-19/Angola/CERI-KRISP-K013467/2020** | EPI_ISL_2492785 | | Instituto Nacional de Investigação em Saúde | CERI, Centre for Epidemic Response and Innvoation, Stellenbosch University and KRISP, KZN Research Innovation and Sequencing Platform, UKZN. | Morais J et al |
| **hCoV-19/Qatar/QA-QU_18-8-H10/2021** | EPI_ISL_1713817 | | Ministry of Public Health / Hamad Medical Corporation | Biomedical Research Center (BRC), Qatar University / Qatar Genome Project (QGP) | BRC: Fatiha M. Benslimane et al |
| **hCoV-19/Zambia/SP113/2021** | EPI_ISL_6768069 | | Macha Research Trust | Macha Research Trust | Katowa B et al |
| **hCoV-19/Zambia/ZMB-93772/2021** | EPI_ISL_2803632 | | University of Zambia, School of Veterinary Medicine | UNZAVET and PATH | Mulenga Mwenda-Chimfwembe et al |

| | | | | | |
|---|---|---|---|---|---|
| **hCoV-19/SouthSudan/UG570/2021** | EPI_ISL_2928017 | | South Sudan Ministry of Health, WHO South Sudan, MRC/UVRI & LSHTM Uganda Research Unit | MRC/UVRI & LSHTM Uganda Research Unit, South Sudan Ministry of Health, WHO South Sudan | Joseph Francis Wamala et al |
| **hCoV-19/Zambia/ZMB-70145/2020** | EPI_ISL_977290 | | University of Zambia, School of Veterinary Medicine | UNZAVET and PATH | Mulenga Mwenda-Chimfwembe et al |
| **hCoV-19/Lesotho/N4813/2020** | EPI_ISL_13037722 | | National Reference Laboratories | National Institute for Communicable Diseases of the National Health Laboratory Service | Gorova V et al |
| **hCoV-19/Mozambique/MZ-L-IBV-97025357/2021** | EPI_ISL_2442118 | | Centro de Investigação em Saúde de Manhiça (CISM) & ISGlobal, Institut de Salut Global de Barcelona | Instituto de Biomedicina de València - CSIC | Alfredo Mayor et al |
| **hCoV-19/Lesotho/N4820/2020** | EPI_ISL_13037729 | | National Reference Laboratories | National Institute for Communicable Diseases of the National Health Laboratory Service | Gorova V et al |
| **hCoV-19/Mozambique/COV-2297-A1945/2021** | EPI_ISL_5822661 | | Instiuto Nacional de Saude, Mozambique | Forschungszentrum Borstel | Christian Utpatel et al |
| **hCoV-19/Coted-Ivoire/IPCI-cv323/2021** | EPI_ISL_10077726 | | Environment and Health Department, Institut Pasteur of Côte d'Ivoire, Abidjan, Côte d'Ivoire | Molecular Genetics Platform, Institut Pasteur of Côte d'Ivoire, Abidjan, Côte d'Ivoire | Aristide Berenger Ako AKO et al |
| **hCoV-19/China/AP100/2021** | EPI_ISL_2779639 | | State Key Laboratory of Emerging Infectious Diseases, The University of Hong Kong | The University of Hong Kong | Lifeng Li et al |
| **hCoV-19/Jiangsu/JS02/2021** | EPI_ISL_16405158 | | Jiangsu Provincial Center for Disease Control and Prevention | Jiangsu Provincial Center for Disease Control and Prevention | Lunbiao Cui et al |
| **hCoV-19/Shandong/QD-1/2021** | EPI_ISL_13499640 | | Shandong Center for Disease Control and Prevention | Shandong Center for Disease Control and Prevention | Xiaolin Jiang et al |
| **hCoV-19/Philippines/PH-PGC-45949/2021** | EPI_ISL_5570609 | | Qualimed Hospital Sta. Rosa | Philippine Genome Center | Cynthia P. Saloma et al |
| **hCoV-19/Taiwan/1013/2021** | EPI_ISL_5854267 | | Centers for Disease Control, R.O.C. (Taiwan) | Centers for Disease Control, R.O.C. (Taiwan) | Ji-Rong Yang et al B |
| **hCoV-19/Malawi/CERI-KRISP-K015199/2021** | EPI_ISL_2492734 | | National HIV Reference Laboratory, Ministry of Health, Public Health Institute of Malawi | CERI, Centre for Epidemic Response and Innvoation, Stellenbosch University and KRISP, KZN Research Innovation and Sequencing Platform, UKZN. | Mvula B et al |
| **hCoV-19/SouthAfrica/NICD-A112-002-146/2020** | EPI_ISL_16078824 | | National Institute for Communicable Diseases of the National Health Laboratory Service | National Institute for Communicable Diseases of the National Health Laboratory Service | Everatt J et al |
| **hCoV-19/SouthAfrica/UFS-133/2021** | EPI_ISL_912484 | | NHLS Universitas Academic | UFS Virology | PA Bester et al |
| **hCoV-19/Zambia/ZMB-70147/2020** | EPI_ISL_977274 | | University of Zambia, School of Veterinary Medicine | UNZAVET and PATH | Mulenga Mwenda-Chimfwembe et al |
| **hCoV-19/Zambia/ZMB-93775/2021** | EPI_ISL_2803646 | | University of Zambia, School of Veterinary Medicine | UNZAVET and PATH | Mulenga Mwenda-Chimfwembe et al |
| **hCoV-19/SaudiArabia/KFSHRC_008S/2021** | EPI_ISL_7570637 | | Molecular Virology section, department of infection & immunity, KFSHRC | Molecular Virology section, department of infection & immunity, KFSHRC | Fatimah S. Alhamlan et al |
| **hCoV-19/Mozambique/MZ-L-IBV-97025429/2021** | EPI_ISL_2442175 | | Centro de Investigação em Saúde de Manhiça (CISM) & ISGlobal, Institut de Salut Global de Barcelona | Instituto de Biomedicina de València - CSIC | Alfredo Mayor et al |
| **hCoV-19/Tanzania/KIDH-01B04/2021** | EPI_ISL_16131913 | | National Public Health Laboratory | Kibong'oto Infectious Diseases Hospital Laboratory | Shabani Mziray et al |
| **hCoV-19/Pakistan/NIH-B12-S12/2021** | EPI_ISL_2757745 | | Department of Virology | Department of Virology | Massab Umair et al |
| **hCoV-19/SouthKorea/KDCA5465/2021** | EPI_ISL_2887378 | | Division of Emerging Infectious Diseases, Bureau of Infectious Diseases Diagnosis Control, Korea Disease Control and Prevention Agency | Division of Emerging Infectious Diseases, Bureau of Infectious Diseases Diagnosis Control, Korea Disease Control and Prevention Agency | Jeong-Ah Kim et al |
| **hCoV-19/Eswatini/N2780/2020** | EPI_ISL_1196005 | | RSP Clinic | National Institute for Communicable Diseases of the National Health Laboratory Service | Maphalala GP et al |
| **hCoV-19/Eswatini/N2793/2021** | EPI_ISL_1196013 | | Hlathikhulu Government Hospital | National Institute for Communicable Diseases of the National Health Laboratory Service | Maphalala GP et al |
| **hCoV-19/Eswatini/N2787/2020** | EPI_ISL_1196009 | | Nhlangano Health Centre | National Institute for Communicable Diseases of the National Health Laboratory Service | Maphalala GP et al |
| **hCoV-19/Malawi/COM-SEQ8473/2021** | EPI_ISL_8889566 | | Kamuzu University of Health Sciences, College of Medicine, Department of Pathology | Kamuzu University of Health Sciences, College of Medicine, Department of Pathology | Tonney Nyirenda et al |
| **hCoV-19/Malawi/KRISP-K010154/2020** | EPI_ISL_1407115 | | National HIV Reference Laboratory, Ministry of Health, Public Health Institute of Malawi | KRISP, KZN Research Innovation and Sequencing Platform | Mvula B et al |
| **hCoV-19/Rwanda/NRLNAT4113/2021** | EPI_ISL_2828496 | | Nucleic Acid Testing, National Reference Laboratory | GIGA Medical Genomics | Yvan Butera et al |
| **hCoV-19/Eswatini/N4796/2021** | EPI_ISL_1827699 | | Lobamba | National Institute for Communicable Diseases of the National Health Laboratory Service | Maphalala GP et al |
| **hCoV-19/Guangzhou/ZhangHuan/2020** | EPI_ISL_2536954 | | Guangdong Provincial Center for Disease Control and Prevention,The Biological safety level-3 (BSL-3)Laboratory | Guangdong Provincia Center for Disease Control and Prevention,Institute of Pathogenic Microbiology,Respiratory Pathogens Laboratory. | Zhang Huan et al |
| **hCoV-19/mouse/Beijing/MP7/2021** | EPI_ISL_16138137 | | Key Laboratory of Human Diseases Comparative Medicine, Institute of Laboratory Animal Science | Key Laboratory of Human Diseases Comparative Medicine, Institute of Laboratory Animal Science | Qin et al |
| **hCoV-19/Guangdong/GDIPH-GDPCC-05/2021** | EPI_ISL_18934693 | | Guangdong Provincial Institute of Public Health, Guangdong Provincial Institute of Public Health | Guangdong Provincial Institute of Public Health, Guangdong Provincial Institute of Public Health | Zhou et al |
| **hCoV-19/Eswatini/N10449/2021** | EPI_ISL_9438987 | | Nhlangano Health Centre | National Institute for Communicable Diseases of the National Health Laboratory Service | Maphalala G et al |
| **hCoV-19/Kyrgyzstan/NRL-53480/2021** | EPI_ISL_13283881 | | Republican Reference Laboratory, National TB center, Kyrgyzstan | Republican Reference Laboratory, National TB center, Kyrgyzstan | Gulmira Kalmambetova et al |
| **hCoV-19/Seychelles/G38723/2021** | EPI_ISL_4880534 | OK377351.1 | Seychelles Public Health Laboratory | KEMRI-Wellcome Trust Research Programme,Kilifi | Githinji G. et al |
| **hCoV-19/Seychelles/G40372/2021** | EPI_ISL_4880536 | OK377353.1 | Seychelles Public Health Laboratory | KEMRI-Wellcome Trust Research Programme,Kilifi | Githinji G. et al |
| **hCoV-19/Seychelles/SS459/2021** | EPI_ISL_4880537 | OK377354.1 | Seychelles Public Health Laboratory | KEMRI-Wellcome Trust Research Programme,Kilifi | Githinji G. et al |
| **hCoV-19/Singapore/274/2021** | EPI_ISL_1312385 | | National Public Health Laboratory, National Centre for Infectious Diseases | National Public Health Laboratory, National Centre for Infectious Diseases | Tze Minn Mak et al |
| **hCoV-19/SaudiArabia/KFSHRC_0121J/2021** | EPI_ISL_6641078 | | Molecular Virology section, department of infection & immunity, KFSHRC | Molecular Virology section, department of infection & immunity, KFSHRC | Fatimah S. Alhamlan et al |
| **hCoV-19/Madagascar/IPM-62176/2021** | EPI_ISL_7722848 | | Virology Unit, Institut Pasteur de Madagascar | Virology Unit, Institut Pasteur de Madagascar | Christian Ranaivoson et al |
| **hCoV-19/Spain/CT-HUGTiPM011LA8A11/2021** | EPI_ISL_1661884 | | Microbiology Department, Laboratori Clínic Metropolitana Nord. Hospital Universitari Germans Trias i Pujol. | Can Ruti SARS-CoV-2 Sequencing Hub (HUGTiP/IrsiCaixa/IGTP) | Marc Noguera-Julian et al |
| **hCoV-19/Madagascar/IPM-61943/2021** | EPI_ISL_7722800 | | Virology Unit, Institut Pasteur de Madagascar | Virology Unit, Institut Pasteur de Madagascar | Christian Ranaivoson et al |
| **hCoV-19/France/IdF-IPP14988/2021** | EPI_ISL_2923209 | | Labo Analyses Med | National Reference Center for Viruses of Respiratory Infections, Institut Pasteur, Paris | Marion Barbet et al A |
| **hCoV-19/Luxembourg/LNS6131672/2021** | EPI_ISL_3148335 | | Laboratoires d'analyses medicales - Ketterthill | Laboratoire national de sante, Microbiology, Microbial Genomics Platform | Anke Wienecke-Baldacchino et al |
| **hCoV-19/Namibia/CERI-KRISP-K024883_2/2021** | EPI_ISL_6506435 | | Namibia Institute of Pathology LTD | CERI, Centre for Epidemic Response and Innvoation, Stellenbosch University and KRISP, KZN Research Innovation and Sequencing Platform, UKZN. | Iyaloo Constantinus et al |
| **hCoV-19/EquatorialGuinea/78388/2021** | EPI_ISL_1700675 | | Laboratorio de Investigaciones de Baney | Swiss Tropical and Public Health Institute | Salome Hosch et al |
| **hCoV-19/Djibouti/NAMRU3_ET_126/2021** | EPI_ISL_3920412 | OK018290.1 | Naval Medical Research Unit No. 3 | Naval Medical Research Center Biological Defense Research Directorate | Catherine E. Arnold et al |
| **hCoV-19/Djibouti/NAMRU3_A175/2021** | EPI_ISL_3920396 | OK018237.1 | Naval Medical Research Unit No. 3 | Naval Medical Research Center Biological Defense Research Directorate | Catherine E. Arnold et al |
| **hCoV-19/Uganda/UG019/2020** | EPI_ISL_451201 | | Uganda Virus Research Institute | MRC/UVRI & LSHTM Uganda Research Unit | Dan Lule Bugembe et al (https://dx.doi.org/10.3201/eid2610.202575) |
| **hCoV-19/Oman/520244640/2020** | EPI_ISL_1532284 | | Oman-National Influenza Center | Biotechnology & OMICs Laboratory | Ahmed Al Harrasi et al |
| **hCoV-19/Belgium/ULG-10358/2020** | EPI_ISL_540525 | | Department of Clinical Microbiology | GIGA Medical Genomics | Keith Durkin et al |
| **hCoV-19/Netherlands/DR-RIVM-13027/2021** | EPI_ISL_1035753 | | National Institute for Public Health and the Environment (RIVM) | National Institute for Public Health and the Environment (RIVM) | Adam Meijer et al |
| **hCoV-19/Portugal/PT4377/2021** | EPI_ISL_1399487 | | INSA | Instituto Nacional de Saude (INSA) | Borges et al |
| **hCoV-19/Rwanda/NRLNAT2006/2020** | EPI_ISL_960232 | | Nucleic Acid Testing, National Reference Laboratory | GIGA Medical Genomics | Yvan Butera et al |
| **hCoV-19/Rwanda/AV79004RD/2020** | EPI_ISL_1064022 | | Rwanda National Reference Laboratory | Rwanda National Reference Laboratory | Enatha Mukantwari et al |
| **hCoV-19/Rwanda/NRLNAT2009/2020** | EPI_ISL_960228 | | Nucleic Acid Testing, National Reference Laboratory | GIGA Medical Genomics | Yvan Butera et al |
| **hCoV-19/India/GJ-GBRC624b/2021** | EPI_ISL_2001184 | MZ325344.1 | GMERS, Government Medical College (GMC), Gotri | Gujarat Biotechnology Research Centre | Sonal Sharma et al |
| **hCoV-19/India/GJ-GBRC-377b/2020** | EPI_ISL_524736 | MT953893.1 | Government General Hospital, Jam Khambhaliya, Devbhoomi Dwarka | Gujarat Biotechnology Research Centre | Janvi Raval et al |
| **hCoV-19/Benin/1022/2020** | EPI_ISL_476826 | MT653075.1 | Laboratoire des Fièvres Hémorragiques Virales du Benin | Charité-Universitätsmedizin Berlin | Yadouleton et al (https://www.medrxiv.org/content/10.1101/2020.06.29.20140749v1) |
| **hCoV-19/India/AN-ICMR-NIV-INSACOG-SEQ-PTBR13/2020** | EPI_ISL_15038437 | | ICMR-National Institute of Virology - INSACOG | NIV Influenza | Dr. Varsha Potdar et al |
| **hCoV-19/SriLanka/CDR1885/2020** | EPI_ISL_525476 | | Centre for Dengue Research | Centre for Dengue Research | Chandima Jeewandara et al |
| **hCoV-19/SouthKorea/KDCA8575/2021** | EPI_ISL_3544916 | | Division of Emerging Infectious Diseases, Bureau of Infectious Diseases Diagnosis Control, Korea Disease Control and Prevention Agency | Division of Emerging Infectious Diseases, Bureau of Infectious Diseases Diagnosis Control, Korea Disease Control and Prevention Agency | Ae Kyung Park et al |
| **hCoV-19/SouthKorea/KDCA5261/2021** | EPI_ISL_2754711 | | Division of Emerging Infectious Diseases, Bureau of Infectious Diseases Diagnosis Control, Korea Disease Control and Prevention Agency | Division of Emerging Infectious Diseases, Bureau of Infectious Diseases Diagnosis Control, Korea Disease Control and Prevention Agency | Jeong-Ah Kim et al |
| **hCoV-19/Congo/FCRM-44-A32-07-07/2021** | EPI_ISL_3040131 | | Fondation Congolaise pour la recherche medicale (FCRM), Francine Ntoumi | Fondation Congolaise pour la Recherche Médicale | Mfoutou Mapanguy Claujens Chastel et al |
| **hCoV-19/CentralAfricanRepublic/237/2021** | EPI_ISL_2365411 | | WWF Bayanga field laboratory | Robert Koch Institute | S. Calvignac-Spencer et al |
| **hCoV-19/Guangdong/SYSU-IHV/2020** | EPI_ISL_444969 | | Guangzhou Eighth People's Hospital (Jiahe Sector) | Institute of Human Virology, Zhongshan School of Medicine, Sun Yat-sen University | Junsong Zhang et al |
| **hCoV-19/Suriname/SR-28/2020** | EPI_ISL_517639 | | Academic Hospital Paramaribo | Erasmus Medical Center | Bas Oude Munnink et al |
| **hCoV-19/Suriname/SR-30/2020** | EPI_ISL_517641 | | Academic Hospital Paramaribo | Erasmus Medical Center | Bas Oude Munnink et al |
| **hCoV-19/Suriname/SR-40/2020** | EPI_ISL_517651 | | Academic Hospital Paramaribo | Erasmus Medical Center | Bas Oude Munnink et al |
| **hCoV-19/Suriname/SR-89/2020** | EPI_ISL_2502538 | | Central Laboratory, Bureau of Public Health (BOG) and Academic Hospital Paramaribo | Erasmus Medical Center | Bas B Oude Munnink et al |
| **hCoV-19/Suriname/SR-69/2020** | EPI_ISL_517661 | | Academic Hospital Paramaribo | Erasmus Medical Center | Bas Oude Munnink et al |
| **hCoV-19/Mexico/YUC-InDRE-95/2020** | EPI_ISL_861419 | | Instituto de Diagnostico y Referencia Epidemiologicos (INDRE) | Instituto de Diagnostico y Referencia Epidemiologicos (INDRE) | Fabiola Garces-Ayala et al |
| **hCoV-19/Luxembourg/LNS3257811/2020** | EPI_ISL_429773 | | Laboratoire National de Sante, Microbiology, Virology | Laboratoire National de Sante, Microbiology, Epidemiology and Microbial Genomics | Anke Wienecke-Baldacchino et al |
| **hCoV-19/Senegal/6427/2020** | EPI_ISL_486873 | | Institut Pasteur Dakar | Institut Pasteur de Dakar | Ndongo Dia et al |
| **hCoV-19/India/MH-ACTREC-057/2020** | EPI_ISL_699713 | | Hematopathology Laboratory, ACTREC, TMC | Hematopathology Laboratory, ACTREC, TMC | Hematopathology Laboratory et al |
| **hCoV-19/India/GJ-GBRC170a/2020** | EPI_ISL_467031 | | GMERS Medical College and Hospital, Gandhinagar | Gujarat Biotechnology Research Centre | Bhavesh Modi et al (https://www.biorxiv.org/content/10.1101/2020.07.10.197095v1) |
| **hCoV-19/Cameroon/CV417/2020** | EPI_ISL_845550 | | National Public Health Laboratory, Cameroon | African Centre of Excellence for Genomics of Infectious Diseases (ACEGID), Redeemer's University | Oluniyi P.E. et al |
| **hCoV-19/Morocco/IPM20155760/2020** | EPI_ISL_17650193 | | Institut Pasteur du Maroc, Medical Virology and BSL3 Laboratory | Institut Pasteur du Maroc, Centre de Séquençage Génomique | EL Hamouchi Adil et al |
| **hCoV-19/Guinea/Conakry_IPG14329/2020** | EPI_ISL_2245855 | | Institut Pasteur de Guinée | Institut Pateur de Dakar | Grayo Solene et al |
| **hCoV-19/Shandong/2020C1240537/2020** | EPI_ISL_962535 | | Infectious Disease Control and Prevention Institute | Infectious Disease Control and Prevention Institute | Xiaolin Jiang et al |
| **hCoV-19/Hungary/SRC-00066/2020** | EPI_ISL_435403 | | Virological Research Group, Szentágothai Research Centre | Bioinformatics Research Group, Szentágothai Research Centre | Péter Urbán et al |
| **hCoV-19/Japan/PG-0542/2020** | EPI_ISL_480198 | | Toyama Institute of Health | Pathogen Genomics Center, National Institute of Infectious Diseases | Tsuyoshi Sekizuka et al A (https://www.medrxiv.org/content/10.1101/2020.07.01.20143958v1) |
| **hCoV-19/Senegal/1821336/2020** | EPI_ISL_8525919 | | Institut Pasteur de Dakar | Institut Pasteur de Dakar | Diallo Amadou et al |
| **hCoV-19/Senegal/91093/2020** | EPI_ISL_8525833 | | Institut Pasteur de Dakar | Institut Pasteur de Dakar | Diallo Amadou et al |
| **hCoV-19/Gambia/23736/2020** | EPI_ISL_1234531 | | MRCG at LSHTM Genomics lab | MRCG at LSHTM Genomics lab | Abdul Karim sesay et al |
| **hCoV-19/Gambia/GC192583/2020** | EPI_ISL_561119 | | MRCG at LSHTM Genomics lab | MRCG at LSHTM Genomics lab | Abdul Karim sesay et al |
| **hCoV-19/Mexico/YUC-InDRE_33/2020** | EPI_ISL_493348 | | Instituto de Diagnostico y Referencia Epidemiologicos (INDRE) | Instituto de Diagnostico y Referencia Epidemiologicos (INDRE) | Ernesto Ramirez-Gonzalez et al |
| **hCoV-19/Qatar/QA-WCMQ_FD17438193/2020** | EPI_ISL_1714186 | | Ministry of Public Health / Hamad Medical Corporation | Weill Cornell Medical College - Qatar (WCM-Q), Genomics Core Laboratory / Qatar Genome Project (QGP) | WCMQ: Ayeda A. Ahmed et al |
| **hCoV-19/Morocco/IPM2037181/2020** | EPI_ISL_17650163 | | Institut Pasteur du Maroc, Medical Virology and BSL3 Laboratory | Institut Pasteur du Maroc, Centre de Séquençage Génomique | EL Hamouchi Adil et al |
| **hCoV-19/Guatemala/ASI1119/2020** | EPI_ISL_5926980 | | Laboratorio Nacional de Salud | Asociación de Salud Integral / Clínica Familiar "Luis Ángel García" | Eduardo Arathon et al |
| **hCoV-19/Colombia/DC-INS-027/2020** | EPI_ISL_526969 | | Instituto Nacional de Salud, Bogotá, Colombia | Instituto Nacional de Salud, Bogotá, Colombia | Katherine Laiton-Donato et al |
| **hCoV-19/USA/TX-HMH-MCoV-12293/2020** | EPI_ISL_790285 | | Houston Methodist Hospital | Houston Methodist Hospital | S. Wesley Long et al |
| **hCoV-19/Morocco/CP17545-INH/2020** | EPI_ISL_18210325 | | NIC at National Institute of Hygiene (NIH) | National Institute of Hygiene | Hassan Ihazmade et al |
| **hCoV-19/EquatorialGuinea/3823/2020** | EPI_ISL_648333 | | Laboratorio de Investigaciones de Baney | University Hospital Basel, Clinical Bacteriology | Carlos Cortes et al |
| **hCoV-19/Morocco/CP19147-INH/2020** | EPI_ISL_18210328 | | NIC at National Institute of Hygiene (NIH) | National Institute of Hygiene | Hassan Ihazmade et al |
| **hCoV-19/Romania/SC04741/2020** | EPI_ISL_5917804 | | Medical Genetics Laboratory, Regional Centre of Medical Genetics, Emergency County Hospital Craiova | Medical Genetics Laboratory, Regional Centre of Medical Genetics, Emergency County Hospital Craiova | Anca-Lelia (Riza) Costache et al |
| **hCoV-19/Romania/DJ38004/2020** | EPI_ISL_6194022 | | Medical Genetics Laboratory, Regional Centre of Medical Genetics, Emergency County Hospital Craiova | Medical Genetics Laboratory, Regional Centre of Medical Genetics, Emergency County Hospital Craiova | Anca-Lelia (Riza) Costache et al |
| **hCoV-19/Romania/DJ52080/2020** | EPI_ISL_6436566 | | Medical Genetics Laboratory, Regional Centre of Medical Genetics, Emergency County Hospital Craiova | Medical Genetics Laboratory, Regional Centre of Medical Genetics, Emergency County Hospital Craiova | Anca-Lelia (Riza) Costache et al |
| **hCoV-19/Romania/DJ44415/2020** | EPI_ISL_6710202 | | Medical Genetics Laboratory, Regional Centre of Medical Genetics, Emergency County Hospital Craiova | Medical Genetics Laboratory, Regional Centre of Medical Genetics, Emergency County Hospital Craiova | Anca-Lelia (Riza) Costache et al |
| **hCoV-19/Italy/EMR-UA-02_00187/2020** | EPI_ISL_5736617 | | Centro di riferimento regionale per le emergenze microbiologiche (CRREM) | Laboratory of Medical Microbiology, University of Antwerp | Mathias Smet et al |
| **hCoV-19/Greece/146389/2020** | EPI_ISL_2302043 | | Department of Microbiology, AHEPA University Hospital | Greek Genome Center, Biomedical Research Foundation of the Academy of Athens (BRFAA) | Emmanouil Athanasiadis et al |
| **hCoV-19/Lebanon/QIB-13/2020** | EPI_ISL_3343227 | OU607095.1 | National Influenza Center | Quadram Institute Bioscience | Georgi Merhi et al |
| **hCoV-19/Austria/CeMM4883/2021** | EPI_ISL_11229731 | | ILV Kärnten | Bergthaler laboratory, CeMM Research Center for Molecular Medicine of the Austrian Academy of Sciences | Lukas Endler et al |
| **hCoV-19/Lithuania/VULSK-397/2020** | EPI_ISL_914886 | | Vilnius university hospital Santaros Klinikos, Center of Laboratory Medicine | Vilnius University Hospital Santaros Klinikos | Ingrida Olendraite et al |
| **hCoV-19/Iceland/4100/2020** | EPI_ISL_827796 | OU165224.1 | deCODE genetics | deCODE genetics | Daniel F Gudbjartsson et al B |
| **hCoV-19/Slovenia/4990/2021** | EPI_ISL_2465316 | | Institute of Microbiology and Immunology, Faculty of Medicine, University of Ljubljana | Institute of Microbiology and Immunology, Faculty of Medicine, University of Ljubljana | Alen Suljič et al |
| **hCoV-19/Slovenia/35182/2021** | EPI_ISL_2465409 | | Institute of Microbiology and Immunology, Faculty of Medicine, University of Ljubljana | Institute of Microbiology and Immunology, Faculty of Medicine, University of Ljubljana | Alen Suljič et al |
| **hCoV-19/Liechtenstein/UHB-4168696501/2020** | EPI_ISL_1233663 | | Labormedizinisches Zentrum Dr Risch | Clinical Bacteriology | Tim Roloff et al |
| **hCoV-19/Kosovo/ChVir23111/2021** | EPI_ISL_1027652 | | Department of Microbiology, National Institute for Public Health of Kosova | Charité Universitätsmedizin Berlin, Institut für Virologie | Victor M Corman et al |
| **hCoV-19/HongKong/CUHK-NC1683/2020** | EPI_ISL_3065981 | | Department of Microbiology | Department of Microbiology | Zigui Chen et al |
| **hCoV-19/SouthAfrica/NHLS-UCT-GS-0421/2020** | EPI_ISL_10815274 | ON055794.1 | Victoria Hospital wc VHW | NHLS/UCT | Arash Iranzadeh et al |
| **hCoV-19/USA/AZ-CDC-4063448-001/2020** | EPI_ISL_1626888 | MW958154.1 | WI State Laboratory of Hygiene | Genomics and Discovery, Respiratory Viruses Branch, Division of Viral Diseases, Centers for Disease Control and Prevention | Jing Zhang et al |
| **hCoV-19/USA/MS-UT-1750/2020** | EPI_ISL_1009195 | | American Esoteric Laboratories | Colleen B. Jonsson | Mariah K. Taylor et al |
| **hCoV-19/Serbia/Belgrade-2950122912/2020** | EPI_ISL_1017681 | | Institute of Virology, Vaccines and Sera ''Torlak'' | Institute of microbiology and Immunology, Faculty of Medicine, University of Belgrade | Knezevic et al |
| **hCoV-19/USA/AL-HGSC-JFWK/2020** | EPI_ISL_883687 | MW930539.1 | Eurofins Diatherix | Hudsonalpha Genome Sequencing Center | Jane Grimwood et al |
| **hCoV-19/Aruba/AW-RIVM-10392/2020** | EPI_ISL_636518 | | National Institute for Public Health and the Environment (RIVM) | National Institute for Public Health and the Environment (RIVM) | Adam Meijer et al |
| **hCoV-19/Curacao/CW-RIVM-21703/2020** | EPI_ISL_1014579 | | National Institute for Public Health and the Environment (RIVM) | National Institute for Public Health and the Environment (RIVM) | Adam Meijer et al |
| **hCoV-19/Aruba/AW-RIVM-11761/2021** | EPI_ISL_1013807 | | National Institute for Public Health and the Environment (RIVM) | National Institute for Public Health and the Environment (RIVM) | Adam Meijer et al |
| **hCoV-19/USA/AZ-ASU76467/2020** | EPI_ISL_13065135 | | Arizona State University | Arizona State University | Steven C. Holland et al |
| **hCoV-19/Haiti/56310583/2020** | EPI_ISL_14186976 | | LABORATOIRE NATIONAL DE SANTÉ PUBLIQUE | Laboratory of Parasitic Diseases, Systems Genomics Section, National Institute of Allergy and Infectious Diseases, National Institutes of Health | Samson Marseille et al |
| **hCoV-19/USA/HI-H200471/2020** | EPI_ISL_753014 | | State Laboratories Division, Hawaii State Department of Health | State Laboratories Division, Hawaii State Department of Health | Pamela O'Brien et al |
| **hCoV-19/Lebanon/ML19_S55/2020** | EPI_ISL_12470090 | | Rafik Hariri University Hospital | Khazen Lab | Rola Husni et al |
| **hCoV-19/Lebanon/CMUL-109/2020** | EPI_ISL_13526928 | | Laboratoire Microbiologie Santé et Environnement (LMSE), Lebanese University | Bioinformatics Laboratory / LNCC | Dalal Nour et al |
| **hCoV-19/Lebanon/CMUL-168/2020** | EPI_ISL_13526984 | | Laboratoire Microbiologie Santé et Environnement (LMSE), Lebanese University | Bioinformatics Laboratory / LNCC | Dalal Nour et al |
| **hCoV-19/Lebanon/CMUL-181/2020** | EPI_ISL_13526996 | | Laboratoire Microbiologie Santé et Environnement (LMSE), Lebanese University | Bioinformatics Laboratory / LNCC | Dalal Nour et al |
| **hCoV-19/Lebanon/CMUL-149/2020** | EPI_ISL_13526965 | | Laboratoire Microbiologie Santé et Environnement (LMSE), Lebanese University | Bioinformatics Laboratory / LNCC | Dalal Nour et al |
| **hCoV-19/Lebanon/CMUL-153/2020** | EPI_ISL_13526969 | | Laboratoire Microbiologie Santé et Environnement (LMSE), Lebanese University | Bioinformatics Laboratory / LNCC | Dalal Nour et al |
| **hCoV-19/Morocco/6893/2020** | EPI_ISL_459971 | | Institut Pasteur du Maroc | Institut Pasteur du Maroc | Marion Barbet et al B (https://dx.doi.org/10.1101/2020.06.23.165902) |
| **hCoV-19/SaudiArabia/KAUST-JEDDAH538/2020** | EPI_ISL_512951 | OU996007.1 | Pathogen Genomics Lab King Abdullah University of Science and Technology(KAUST) | Pathogen Genomics Lab King Abdullah University of Science and Technology(KAUST) | Sara Mfarrej et al |
| **hCoV-19/Ecuador/USFQ-020/2020** | EPI_ISL_471267 | | Hospital IESS Babahoyo | Institute of Microbiology, Universidad San Francisco de Quito | Sully Márquez et al |
| **hCoV-19/Panama/GMI-PA392814/2020** | EPI_ISL_1225483 | | Gorgas Memorial Laboratory of Health Studies | Gorgas Memorial Laboratory of Health Studies | Yamilka Diaz et al |
| **hCoV-19/Nigeria/ISTH-E0199/2020** | EPI_ISL_14205937 | | Irrua Specialist Teaching Hospital, ISTH | "Irrua Specialist Teaching Hospital, ISTH, and the Bernhard Nocht Institute for Tropical Medicine, BNITM" | "Sylvanus Okogbenin et al |
| **hCoV-19/India/OR-ICMR-NIV-INSACOG-SEQ-996/2020** | EPI_ISL_15038521 | | ICMR-National Institute of Virology - INSACOG | NIV Influenza | Dr. Varsha Potdar et al |
| **hCoV-19/Ukraine/UA-RIVM-21881/2020** | EPI_ISL_3253267 | | National Institute for Public Health and the Environment (RIVM) | National Institute for Public Health and the Environment (RIVM) | Adam Meijer et al |
| **hCoV-19/Turkey/HSGM-B11140/2021** | EPI_ISL_2158047 | | Ministry of Health Turkey | Ministry of Health Turkey | Fatma Bayrakdar et al |
| **hCoV-19/Spain/MD-ISCIII-2014936/2020** | EPI_ISL_871993 | | Hospital Clínico San Carlos | Instituto de Salud Carlos III | Iglesias-Caballero et al |
| **hCoV-19/Spain/MD-ISCIII-2014662/2020** | EPI_ISL_691717 | | Hospital Clínico San Carlos | Instituto de Salud Carlos III | Iglesias-Caballero et al |
| **hCoV-19/Tunisia/M2355/2020** | EPI_ISL_11172666 | | Laboratory of Clinical Virology, Institut Pasteur de Tunis | Laboratory of Clinical Virology, Institut Pasteur de Tunis | Haifa Jmel et al |
| **hCoV-19/Romania/SV_SJU_26261/2020** | EPI_ISL_6640779 | | Molecular Biology Lab, Suceava County Emergency Hospital | Stefan cel Mare University Metagenomics Lab | Lobiuc Andrei et al |
| **hCoV-19/Vietnam/PY-56/2020** | EPI_ISL_17373283 | | CDC Phú Yên, Vietnam | Department of Microbiology and Immunology, Pasteur Institute in Nha Trang | Trinh Hoang Long et al |
| **hCoV-19/Germany/BY-MVP-000001177/2020** | EPI_ISL_1751462 | | Max von Pettenkofer Institute, Virology, National Reference Center for Retroviruses, LMU Munich | Laboratory for Functional Genome Analysis; Dept. Genomics; Gene Center of the LMU Munich | Max Muenchhoff et al |
| **hCoV-19/Spain/CT-IBV-98014717/2021** | EPI_ISL_1194627 | | Departamento de Microbiología, CDB, Hospital Clínic, Barcelona | SeqCOVID-SPAIN consortium/IBV(CSIC) | Andrea Vergara et al |
| **hCoV-19/Belgium/rega-0804719/2020** | EPI_ISL_734906 | | UZ Leuven, National Reference Laboratory for Coronaviruses, Laboratory Medicine, Leuven, Belgium | KU Leuven, Rega Institute, Clinical and Epidemiological Virology | Tony Wawina-Bokalanga et al |
| **hCoV-19/Gibraltar/204462891/2020** | EPI_ISL_615056 | | Microbiology, Department of Pathology, St. Bernard's Hospital, Gibraltar Health Authority | Respiratory Virus Unit, Microbiology Services Colindale, Public Health England | PHE Covid Sequencing Team et al |
| **hCoV-19/Norway/3377/2020** | EPI_ISL_3801249 | | Vestfold Hospital, Toensberg Department of Microbiology | Norwegian Institute of Public Health, Department of Virology | Kathrine Stene-Johansen et al |
| **hCoV-19/Latvia/126/2020** | EPI_ISL_534215 | | E. Gulbja Laboratorija | Latvian Biomedical Research and Study Centre | Ivars Silamiķelis et al |
| **hCoV-19/Turkey/HSGM-1196/2020** | EPI_ISL_812779 | | Ministry of Health Turkey | Ministry of Health Turkey | Fatma Bayrakdar et al |
| **hCoV-19/Spain/GA-EPI-HULA-921/2020** | EPI_ISL_18630263 | | Servicio de Microbiología. Hospital Lucus Augusti | CINBIO / EPICOVIGAL Consortium | María del Pilar Gallego on behalf of EPICOVIGAL Consortium et al |
| **hCoV-19/Spain/GA-EPI-CHUVI-769/2020** | EPI_ISL_18630490 | | Microbiology Department. Complexo Hospitalario Universitario de Vigo | CiMUS / EPICOVIGAL Consortium | María del Pilar Gallego on behalf of EPICOVIGAL Consortium et al |
| **hCoV-19/Spain/MD-IBV-99008678/2020** | EPI_ISL_654233 | | Hospital General Universitario Gregorio Marañón | SeqCOVID-SPAIN consortium/IBV(CSIC) | Darío García de Viedma et al |
| **hCoV-19/Netherlands/LI-MUMC-360/2021** | EPI_ISL_965633 | | Dutch COVID-19 response team | Medical Microbiology, Maastricht University Medical Centre | Jozef Dingemans" et al |
| **hCoV-19/Ireland/D-NVRL-20IRL77557/2020** | EPI_ISL_732399 | | National Virus Reference Laboratory | National Virus Reference Laboratory | Michael Carr et al |
| **hCoV-19/Thailand/Chonburi-SQ-LN0002/2020** | EPI_ISL_708804 | | Regional medical sciences center 6 chonburi | National Institute of Health, Department of Medical Sciences, Ministry of Public Health, Thailand | Pilailuk Okada et al |
| **hCoV-19/Thailand/A21942/2020** | EPI_ISL_3892157 | | Port Health and Quarantine, Suvarnabhumi Airport | Thai Red Cross Emerging Infectious Diseases Clinical Center and Faculty of Medicine, Chulalongkorn University | Rome Buathong et al |
| **hCoV-19/Cyprus/BMVUCY536/2020** | EPI_ISL_7210724 | | NIPD Genetics Limited | Laboratory of Biotechnology and Molecular Virology, University of Cyprus | Leondios G. Kostrikis et al. |
| **hCoV-19/Turkey/HSGM-B10310/2021** | EPI_ISL_2158060 | | Ministry of Health Turkey | Ministry of Health Turkey | Fatma Bayrakdar et al |
| **hCoV-19/Latvia/299/2021** | EPI_ISL_961836 | | Centrālā Laboratorija | Latvian Biomedical Research and Study Centre | Janis Pjalkovskis et al |
| **hCoV-19/India/AS-CVD-34936/2020** | EPI_ISL_4192891 | | INSACOG-Assam | National Institute of Biomedical Genomics – INSACOG | Arindam Maitra et al A |
| **hCoV-19/India/GA-IGIB-GSEQ_2651/2021** | EPI_ISL_2879601 | | INSACOG Surveillance | INSACOG at CSIR Institute of Genomics and Integrative Biology | INSACOG et al |
| **hCoV-19/India/TG-RFCH00300_CIB0710/2021** | EPI_ISL_6696835 | | Osmania Medical College | CSIR-Centre for Cellular and Molecular Biology | Lamuk Zaveri et al |

| | | | | | |
|---|---|---|---|---|---|
| **hCoV-19/India/WB-1931501069320/2021** | EPI_ISL_2503820 | | INSACOG-WB | National Institute of Biomedical Genomics – INSACOG | Arindam Maitra et al A |
| **hCoV-19/India/TG-CDFD-Xi44/2021** | EPI_ISL_5053395 | | Centre for DNA Fingerprinting and Diagnostics | CDFD-INSACOG | Murali Bashyam et al |
| **hCoV-19/India/TG-CCMB-CIA5219/2021** | EPI_ISL_2775048 | | CSIR-Centre for Cellular and Molecular Biology | CSIR-Centre for Cellular and Molecular Biology-INSACOG | Priya Nurkurthy et al |
| **hCoV-19/Singapore/1214/2021** | EPI_ISL_2508815 | | National Public Health Laboratory, National Centre for Infectious Diseases | National Public Health Laboratory, National Centre for Infectious Diseases | Tze Minn Mak et al |
| **hCoV-19/India/un-1_31/2021** | EPI_ISL_15590992 | | Infection & Immunity, University College London | Infection & Immunity, University College London | Mlcochova et al |
| **hCoV-19/India/ML-CCMB-CIA3186/2021** | EPI_ISL_2660865 | | PASTEUR MOLECULAR LABORATORY - INSACOG MEGHALAYA | CSIR-Centre for Cellular and Molecular Biology-INSACOG | R. Lyngdoh et al |
| **hCoV-19/Guangdong/6486/2021** | EPI_ISL_16138273 | | GDCDC&GDIPH, Guangdong Provincial Center for Diseases Control and Prevention | GDCDC&GDIPH, Guangdong Provincial Center for Diseases Control and Prevention | Li et al |
| **hCoV-19/Guangdong/5648/2021** | EPI_ISL_16138237 | | GDCDC&GDIPH, Guangdong Provincial Center for Diseases Control and Prevention | GDCDC&GDIPH, Guangdong Provincial Center for Diseases Control and Prevention | Li et al |
| **hCoV-19/Maldives/MAV28931/2021** | EPI_ISL_3275166 | | AA. Rasdhoo | Indira Gandhi Memorial Hospital | Dr. Milza Abdul Muhsin et al |
| **hCoV-19/Chongqing/210052/2021** | EPI_ISL_1970349 | | Chongqing International Travel Health Care Center | Chongqing Municipal Center for Disease Control and Prevention | Sheng Ye et al |
| **hCoV-19/India/MZ-INSACOG-1526100785782/2021** | EPI_ISL_8661346 | | INSACOG-Mizoram | National Institute of Biomedical Genomics – INSACOG | Arindam Maitra et al A |
| **hCoV-19/India/MZ-1526100276803/2021** | EPI_ISL_3189471 | | INSACOG-Mizoram | National Institute of Biomedical Genomics – INSACOG | Arindam Maitra et al A |
| **hCoV-19/India/MZ-INSACOG-1526100306322/2021** | EPI_ISL_4312501 | | INSACOG-Mizoram | National Institute of Biomedical Genomics – INSACOG | Arindam Maitra et al A |
| **hCoV-19/India/MZ-INSACOG-1526100339464/2021** | EPI_ISL_4313068 | | INSACOG-Mizoram | National Institute of Biomedical Genomics – INSACOG | Arindam Maitra et al A |
| **hCoV-19/India/CT-ILSGS04068/2021** | EPI_ISL_3015521 | | Pt. Jawahar Lal Nehru Memorial Medical College, Raipur | Institute of Life Sciences - INSACOG | Sunil K. Raghav et al |
| **hCoV-19/India/MH-RFIP03416/2021** | EPI_ISL_6144450 | | Genepath, Pune | RF-IISER Pune | Krishanpal Karmodiya et al |
| **hCoV-19/Mexico/BCN-SEARCH-104125/2021** | EPI_ISL_3215991 | | Genomica Lab Molecular, Mexico | Andersen lab at Scripps Research | SEARCH Alliance San Diego with Jonathan Gonzalez Garcia et al |
| **hCoV-19/India/CH-NCDC-4509141/2021** | EPI_ISL_4460783 | | National Centre for Disease Control (NCDC) Biotechnology Division, Delhi - INSACOG | NCDC Delhi, Biotechnology Division -INSACOG | Kalaiarasan Ponnusamy et al |
| **hCoV-19/India/KL-GENESCOV04192/2021** | EPI_ISL_7221979 | | INSACOG-IGIB and Kerala State SARS-COV-2 Genome Surveillance Programme (GENESCOV2) | INSACOG and GENESCOV2 at CSIR Institute of Genomics and Integrative Biology | INSACOG-IGIB et al |
| **hCoV-19/India/TN-32548_S299_R1_001/2021** | EPI_ISL_3265248 | | JIPMER Puducherry | inStem NCBS – INSACOG | Uma Ramakrishnan Dasaradhi Palakodeti Aswin SaiNarain et al |
| **hCoV-19/Taiwan/16164/2021** | EPI_ISL_16544814 | | Centers for Disease Control, R.O.C. (Taiwan) | Centers for Disease Control, R.O.C. (Taiwan) | Yu-Chi Lin et al |
| **hCoV-19/India/TG-CDFD-102994/2021** | EPI_ISL_3242273 | | Osmania Medical College | CDFD-INSACOG | Murali Bashyam et al |
| **hCoV-19/India/AP-CCMB-CIA7951/2021** | EPI_ISL_4545929 | | AP SSO | CSIR-Centre for Cellular and Molecular Biology - INSACOG | Sumedha Avadhanula et al |
| **hCoV-19/Zhejiang/zhoushan37/2021** | EPI_ISL_1911197 | | Zhejiang Provincial Center for Disease Control and Prevention Zhoushan Center for Disease Prevention and Contol | Zhejiang Province Center of Disease Control and prevention | Yanjun Zhang et al |
| **hCoV-19/Mongolia/UB-373297/2021** | EPI_ISL_9910921 | | National Center for Communicable Diseases (NCCD) National Influenza Center | National Center for Communicable Diseases (NCCD) National Influenza Center | Naranzul Tsedenbal et al |
| **hCoV-19/India/MN-IBSD-IE-P2-239/2021** | EPI_ISL_11803986 | | Babina Diagnostics | INSACOG-IBSD Manipur | Dr. Nanaocha Sharma et al |
| **hCoV-19/India/AN-ICMR-NIV-INSACOG-GSEQ-4941/2021** | EPI_ISL_3532972 | | ICMR-National Institute of Virology - INSACOG | NIV Influenza | Potdar et al B |
| **hCoV-19/India/LEH-NCDC-2709182/2021** | EPI_ISL_2461995 | | National Centre for Disease Control (NCDC) Biotechnology Division, Delhi | NCDC Delhi, Biotechnology Division INSACOG | Kalaiarasan Ponnusamy et al |
| **hCoV-19/India/OR-ILSGS03220/2021** | EPI_ISL_3008124 | | District Headquarter Hospital, Baripada | Institute of Life Sciences - INSACOG | Sunil K. Raghav et al |
| **hCoV-19/Bangladesh/IEDCR-OIS-0079/2021** | EPI_ISL_2284894 | MZ377113.1 | Institute of Epidemiology, Disease Control and Research (IEDCR) | IEDCR-ideSHi-icddr,b | Hassan Afrad et al |
| **hCoV-19/India/WB-267174/2021** | EPI_ISL_7210872 | | ICMR-National Institute of Cholera and Enteric Diseases | DBT-National Institute Of Biomedical Genomics | Arindam Maitra et al A |
| **hCoV-19/Bangladesh/icddrb-GSB14-010/2021** | EPI_ISL_3762673 | | Shaheed Ziaur Rahman Medical College, Bogra | Virology Laboratory, International Centre for Diarrhoeal Disease Research, Bangladesh (ICDDR,B) | Mojnu Miah et al |
| **hCoV-19/India/CG-AIIMS-Raipur-L-936/2021** | EPI_ISL_2307108 | | State Virus Research and Diagnostic Laboratory (VRDL), AIIMS Raipur | State Virus Research and Diagnostic Laboratory (VRDL), AIIMS Raipur | Pushpendra Singh et al |
| **hCoV-19/India/SK-58543/2021** | EPI_ISL_3019004 | | INSACOG-Sikkim | National Institute of Biomedical Genomics – INSACOG | Arindam Maitra et al A |
| **hCoV-19/India/TR-373860489/2021** | EPI_ISL_3982873 | | INSACOG-Tripura | National Institute of Biomedical Genomics – INSACOG | Arindam Maitra et al A |
| **hCoV-19/Yunnan/RLD156/2021** | EPI_ISL_13008919 | | Department of Acute Infectious Diseases Control and Prevention, Yunnan Provincial Center for Disease Control and Prevention | Department of Acute Infectious Diseases Control and Prevention, Yunnan Provincial Center for Disease Control and Prevention | Meiling Zhang et al |
| **hCoV-19/India/DH-ICMR-NIV-INSACOG-GSEQ-5420/2021** | EPI_ISL_4950426 | | ICMR-National Institute of Virology - INSACOG | NIV Influenza | Potdar et al B |
| **hCoV-19/Hangzhou/ZJU-12/2021** | EPI_ISL_3127444 | | State Key Laboratory for Diagnosis and Treatment of Infectious Diseases, National Clinical Research Center for Infectious Diseases, First Affiliated Hospital, Zhejiang University School of Medicine, Hangzhou, China. 310003 | State Key Laboratory for Diagnosis and Treatment of Infectious Diseases, National Clinical Research Center for Infectious Diseases, First Affiliated Hospital, Zhejiang University School of Medicine, Hangzhou, China. 310003 | Hangping Yao et al |
| **hCoV-19/India/MH-NCDC-2909301/2021** | EPI_ISL_2955867 | | National Centre for Disease Control (NCDC) Biotechnology Division, Delhi | NCDC Delhi, Biotechnology Division INSACOG | Priyanka Singh et al |
| **hCoV-19/India/HR-IGIB-60823/2021** | EPI_ISL_2880515 | | INSACOG Surveillance | INSACOG at CSIR Institute of Genomics and Integrative Biology | INSACOG et al |
| **hCoV-19/Yunnan/YN-11/2021** | EPI_ISL_2840857 | | Department of Acute Infectious Diseases Control and Prevention, Yunnan Provincial Center for Disease Control and Prevention | Department of Acute Infectious Diseases Control and Prevention, Yunnan Provincial Center for Disease Control and Prevention | Meiling Zhang et al |
| **hCoV-19/Togo/BL21_C15396/2021** | EPI_ISL_12521506 | | LNR/CHU-SO | Unité Mixte Internationale TransVIHMI (UMI 233 IRD – U1175 INSERM - Université de Montpellier) IRD (Institut de recherche pour le développement) | A.S. COMLAN et al |
| **hCoV-19/India/JK-NCDC-45098/2021** | EPI_ISL_4460837 | | National Centre for Disease Control (NCDC) Biotechnology Division, Delhi - INSACOG | NCDC Delhi, Biotechnology Division -INSACOG | Kalaiarasan Ponnusamy et al |
| **hCoV-19/Congo/FCRM-47-A16-03-07/2021** | EPI_ISL_3040134 | | Fondation Congolaise pour la recherche medicale (FCRM), Francine Ntoumi | Fondation Congolaise pour la Recherche Médicale | Mfoutou Mapanguy Claujens Chastel et al |
| **hCoV-19/SierraLeone/SL39/2021** | EPI_ISL_15283168 | | State Key Laboratory of Pathogen and Biosecurity, Beijing Institute of Microbiology and Epidemiology | State Key Laboratory of Pathogen and Biosecurity, Beijing Institute of Microbiology and Epidemiology | Wurie et al |
| **hCoV-19/SierraLeone/CPHRL-092/2021** | EPI_ISL_4755975 | | China P3 Laboratory | Central Public Health Reference Laboratory | Allan K.O. Campbell et al |
| **hCoV-19/Thailand/DMSc-02753/2021** | EPI_ISL_5065386 | | Plublic Health Center 29 Chuang Nuknet | Division of Genomic Medicine and Innovation support,Department of Medical Sciences, Ministry of Public Health, Thailand | Surakameth Mahasirimongkol et al |
| **hCoV-19/Thailand/Chanthaburi_SEQ18381/2021** | EPI_ISL_5446199 | | Regional Medical Sciences Center 6 Chonburi | National Institute of Health, Department of Medical Sciences, Ministry of Public Health, Thailand | Pilailuk Okada et al |
| **hCoV-19/Cambodia/892332/2021** | EPI_ISL_8067578 | | Virology Unit, Institut Pasteur du Cambodge | Virology Unit, Institut Pasteur du Cambodge | Leakhena Pum et al |
| **hCoV-19/Cambodia/838346/2021** | EPI_ISL_5802954 | | Virology Unit, Institut Pasteur du Cambodge | Virology Unit, Institut Pasteur du Cambodge | Leakhena Pum et al |
| **hCoV-19/Cambodia/843207/2021** | EPI_ISL_5802988 | | Virology Unit, Institut Pasteur du Cambodge | Virology Unit, Institut Pasteur du Cambodge | Leakhena Pum et al |
| **hCoV-19/Cambodia/22004181/2022** | EPI_ISL_13985253 | | National Institute of Public Health | National Institute of Public Health | Chhorvann CHHEA et al |
| **hCoV-19/Cambodia/f1105053/2021** | EPI_ISL_6227879 | | Virology Unit, Institut Pasteur du Cambodge | Virology Unit, Institut Pasteur du Cambodge | Leakhena Pum et al |
| **hCoV-19/Yunnan/RLD124/2021** | EPI_ISL_13008904 | | Department of Acute Infectious Diseases Control and Prevention, Yunnan Provincial Center for Disease Control and Prevention | Department of Acute Infectious Diseases Control and Prevention, Yunnan Provincial Center for Disease Control and Prevention | Meiling Zhang et al |
| **hCoV-19/Yunnan/RLD105/2021** | EPI_ISL_13008884 | | Department of Acute Infectious Diseases Control and Prevention, Yunnan Provincial Center for Disease Control and Prevention | Department of Acute Infectious Diseases Control and Prevention, Yunnan Provincial Center for Disease Control and Prevention | Meiling Zhang et al |
| **hCoV-19/Thailand/TUC-L21-20040/2021** | EPI_ISL_5106617 | | Mae Sot General Hospital | Thailand MOPH - U.S. CDC Collaboration (TUC) | Thidathip Wongsurawat et al |
| **hCoV-19/Shanxi/IVDC-579/2022** | EPI_ISL_11905844 | | Shanxi Center for Disease Control and Prevention | National Institute for Viral Disease Control and Prevention, China CDC | Binzhi Ren et al |
| **hCoV-19/Beijing/BJCDC-112/2022** | EPI_ISL_13858872 | | Beijing Center for Disease Control and Prevention | Beijing Center for Disease Control and Prevention | Yang Pan et al |
| **hCoV-19/India/GJ-INSACOG-GBRC1309/2021** | EPI_ISL_3161864 | | GMERS Medical College, Valsad | Gujarat Biotechnology Research Centre | Twinkle Soni et al |
| **hCoV-19/SierraLeone/SL38/2021** | EPI_ISL_15283167 | | State Key Laboratory of Pathogen and Biosecurity, Beijing Institute of Microbiology and Epidemiology | State Key Laboratory of Pathogen and Biosecurity, Beijing Institute of Microbiology and Epidemiology | Wurie et al |
| **hCoV-19/India/GJ-INSACOG-GBRC2572/2021** | EPI_ISL_6181391 | | Neuberg Supratech Reference Laboratory, Surat | Gujarat Biotechnology Research Centre | Nitin Savaliya et al |
| **hCoV-19/India/DH-ICMR-NIV-INSACOG-GSEQ-4573/2021** | EPI_ISL_2966187 | | ICMR-National Institute of Virology - INSACOG | NIV Influenza | Potdar et al B |
| **hCoV-19/India/HR-IGIB-606700431077/2021** | EPI_ISL_4126180 | | INSACOG Surveillance | INSACOG at CSIR Institute of Genomics and Integrative Biology | INSACOG et al |
| **hCoV-19/India/MN-1425500077457/2021** | EPI_ISL_3021890 | | INSACOG-MANIPUR | National Institute of Biomedical Genomics – INSACOG | Arindam Maitra et al A |
| **hCoV-19/India/SK-1122500046961/2021** | EPI_ISL_2151096 | | INSACOG-Sikkim | National Institute of Biomedical Genomics – INSACOG | Arindam Maitra et al A |
| **hCoV-19/India/GJ-ICMR-NIV-INSACOG-GSEQ-5365/2021** | EPI_ISL_3982572 | | ICMR-National Institute of Virology - INSACOG | NIV Influenza | Potdar et al B |
| **hCoV-19/Indonesia/SN-EIJK-UNSRAT-028/2021** | EPI_ISL_8258170 | | RS Awaloei | Eijkman Research Center for Molecular Biology, National Research and Innovation Agency; RS Awaloei; RSUP Prof. Dr. R. Kandou Manado | Lidwina Priliani et al |
| **hCoV-19/China/GMU-GX-001/2021** | EPI_ISL_17959241 | | Guangxi Medical University | Guangxi Medical University | Liao et al |
| **hCoV-19/Brunei/15112/2021** | EPI_ISL_8172223 | | National Virology Reference Laboratory | Microbial Genomic Services | Zainun Zaini et al |
| **hCoV-19/Coted-Ivoire/347325-cv025/2021** | EPI_ISL_9888101 | | Environment and Health Department, Institut Pasteur of Côte d'Ivoire, Abidjan, Côte d'Ivoire | Molecular Genetics Platform, Institut Pasteur of Côte d'Ivoire, Abidjan, Côte d'Ivoire | Aristide Berenger Ako AKO et al |
| **hCoV-19/SierraLeone/CPHRL-22/2021** | EPI_ISL_4635451 | | Connaught Hospital Laboratory | Central Public Health Reference Laboratory | Doris Harding et al |
| **hCoV-19/Bahrain/342314242/2021** | EPI_ISL_3462101 | MZ620546.1 | Communicable Disease Laboratory, Public Health Directorate | Communicable Disease Laboratory, Public Health Directorate | Alwasti et al |
| **hCoV-19/UnitedArabEmirates/BTC-7699/2021** | EPI_ISL_16804944 | | Khalifa University Center for Biotechnology | Khalifa University Center for Biotechnology | AlSafar et al. |
| **hCoV-19/Mexico/DUR-CABANA-MX-UGA-IMSS_ITV_UBA_UCR_UdA_UPC-3311/2021** | EPI_ISL_7812952 | | Instituto Mexicano del Seguro Social (IMSS) | CABANA: Unidad de Genómica Avanzada del Centro de Investigación y de Estudios Avanzados (UGA-LANGEBIO, CINVESTAV) | Hebleen Brenes et al |
| **hCoV-19/Djibouti/NAMRU3_SOTFEA_46/2021** | EPI_ISL_6693798 | OL640834.1 | Naval Medical Research Unit No. 3 | Naval Medical Research Center Biological Defense Research Directorate | Catherine E. Arnold et al |
| **hCoV-19/India/AS-NEIST-IGIB-FIND-R32-CID1200/2021** | EPI_ISL_14242801 | | JMCH, Jorhat | Centre for Infectious Diseases, CSIR-NEIST | Emon Kalyan et al |
| **hCoV-19/India/HP-NCDC-2809166/2021** | EPI_ISL_2460991 | | National Centre for Disease Control (NCDC) Biotechnology Division, Delhi | NCDC Delhi, Biotechnology Division INSACOG | Priyanka Singh et al |
| **hCoV-19/India/BR-NCDC-490959/2021** | EPI_ISL_11050728 | | National Centre for Disease Control (NCDC) Biotechnology Division, Delhi - INSACOG | NCDC Delhi, Biotechnology Division -INSACOG | Mahesh S Dhar et al |
| **hCoV-19/Liberia/LIB-0076/2021** | EPI_ISL_3547687 | | National Public Health Institute of Liberia Reference Lab | Center for Infection and Immunity, Columbia University | Bode Shobayo et al |
| **hCoV-19/Canada/NS-NML-249201/2021** | EPI_ISL_8711069 | | NS-QEII Health Sciences Centre | National Microbiology Laboratory (NML) | Anna Majer et al |
| **hCoV-19/Heilongjiang/IVDC-SH/313/2021** | EPI_ISL_9910265 | | Heilongjiang Center for Disease Control and Prevention | National Institute for Viral Disease Control and Prevention, China CDC | Lu Tang et al |
| **hCoV-19/Liberia/LIB-0196/2021** | EPI_ISL_4232135 | | National Public Health Institute of Liberia Reference Lab | Center for Infection and Immunity, Columbia University | Bode Shobayo et al |
| **hCoV-19/India/JK-NCDC-3409380/2021** | EPI_ISL_4445007 | | National Centre for Disease Control (NCDC) Biotechnology Division, Delhi - INSACOG | NCDC Delhi, Biotechnology Division -INSACOG | Robin Marwal et al |
| **hCoV-19/Kuwait/KCCC-91045-3336123-2/2021** | EPI_ISL_2833949 | | Adan Hospital | Cancer Control Center Center | Shakir Bahzad et al |
| **hCoV-19/India/PY-COV2-50478_S9_R1_001/2021** | EPI_ISL_5806938 | | JIPMER Puducherry | inStem NCBS – INSACOG | Uma Ramakrishnan Dasaradhi Palakodeti Aswin SaiNarain et al |
| **hCoV-19/UnitedArabEmirates/BTC-6005/2021** | EPI_ISL_16802390 | | Khalifa University Center for Biotechnology | Khalifa University Center for Biotechnology | AlSafar et al. |
| **hCoV-19/Seychelles/SS1003/2021** | EPI_ISL_4880602 | OK377411.1 | Seychelles Public Health Laboratory | KEMRI-Wellcome Trust Research Programme,Kilifi | Githinji G. et al |
| **hCoV-19/Jiangsu/XB/2021** | EPI_ISL_7876605 | | Institute of pathogenic microbiology, Jiangsu Provincial Center for Disease Control and Prevention | Institute of pathogenic microbiology, Jiangsu Provincial Center for Disease Control and Prevention | lunbiao cui et al |
| **hCoV-19/SriLanka/aicbu59/2021** | EPI_ISL_2985004 | | Allergy, Immunology and Cell Biology Unit (AICBU) | Allergy, Immunology and Cell Biology Unit (AICBU) | Chandima Jeewandara et al |
| **hCoV-19/SriLanka/aicbu1925/2021** | EPI_ISL_5494590 | | Allergy, Immunology and Cell Biology Unit (AICBU) | Allergy, Immunology and Cell Biology Unit (AICBU) | Chandima Jeewandara et al |
| **hCoV-19/SriLanka/aicbu2557/2021** | EPI_ISL_7146466 | | Allergy, Immunology and Cell Biology Unit (AICBU) | Allergy, Immunology and Cell Biology Unit (AICBU) | Chandima Jeewandara et al |
| **hCoV-19/SriLanka/idea_uoc_00193/2021** | EPI_ISL_8879855 | | Department of Molecular Biology, Medical Research Institute, Colombo, Sri Lanka | IDEA Lab | Inoka C. Perera et al |
| **hCoV-19/Philippines/PH-RITM-599/2021** | EPI_ISL_9177523 | | SAN PEDRO RURAL HEALTH UNIT I | Research Institute for Tropical Medicine | Francisco Gerardo M. Polotan et al |
| **hCoV-19/Latvia/2110069122/2021** | EPI_ISL_9700285 | | Riga East Clinical University Hospital | Riga East Clinical University Hospital, National Microbiology Reference Laboratory | Reinis Vangravs et al |
| **hCoV-19/Jiangsu/IVDC-07-01/2021** | EPI_ISL_8295277 | | Jiangsu Provincial Center for Disease Control and Prevention | National Institute for Viral Disease Control and Prevention, China CDC | Shenjiao Wang et al |
| **hCoV-19/Philippines/PH-PGC-100950/2021** | EPI_ISL_9305903 | | Bicol South Luzon - Sub National Reference Laboratory | Philippine Genome Center | Cynthia P. Saloma et al |
| **hCoV-19/Fujian/BD-VIR-SPE-0633/2021** | EPI_ISL_15775712 | | Fujian Center for Disease Control and Prevention | Fujian Center for Disease Control and Prevention | Lin Qi et al |
| **hCoV-19/India/AS-1828500219115/2021** | EPI_ISL_2502363 | | INSACOG-Assam | National Institute of Biomedical Genomics – INSACOG | Arindam Maitra et al A |
| **hCoV-19/India/TG-CDFD-103310/2021** | EPI_ISL_3243240 | | Government Fever Hospital | CDFD-INSACOG | Murali Bashyam et al |
| **hCoV-19/India/AS-NEIST-IGIB-FIND-R32-CID1287/2021** | EPI_ISL_14242823 | | JMCH, Jorhat | Centre for Infectious Diseases, CSIR-NEIST | Emon Kalyan et al |
| **hCoV-19/India/AS-1829500233907/2021** | EPI_ISL_2877775 | | INSACOG-Assam | National Institute of Biomedical Genomics – INSACOG | Arindam Maitra et al A |
| **hCoV-19/India/AS-1829500220578/2021** | EPI_ISL_2877787 | | INSACOG-Assam | National Institute of Biomedical Genomics – INSACOG | Arindam Maitra et al A |
| **hCoV-19/India/CT-ILSGS09450/2021** | EPI_ISL_7877764 | | GMC, Raigarh | Institute of Life Sciences-INSACOG | Sunil K. Raghav et al |
| **hCoV-19/Philippines/PH-VSMMC-7-89/2021** | EPI_ISL_17948984 | | Vicente Sotto Memorial Medical Center | Vicente Sotto Memorial Medical Center | Reynette Christine J. Ligaray et al |
| **hCoV-19/Guatemala/ASI4441/2021** | EPI_ISL_7740505 | | Laboratorio Nacional de Salud | Asociación de Salud Integral / Clínica Familiar "Luis Ángel García" | Eduardo Arathon et al |
| **hCoV-19/India/DL-NCDC-480932/2021** | EPI_ISL_11049563 | | National Centre for Disease Control (NCDC) Biotechnology Division, Delhi - INSACOG | NCDC Delhi, Biotechnology Division -INSACOG | Robin Marwal et al |
| **hCoV-19/Belize/CML-193/2021** | EPI_ISL_5887280 | | Central Medical Laboratory | Baylor College of Medicine | Alexander Kneubehl et al |
| **hCoV-19/India/MP-NCDC-3109320/2021** | EPI_ISL_4436135 | | National Centre for Disease Control (NCDC) Biotechnology Division, Delhi - INSACOG | NCDC Delhi, Biotechnology Division -INSACOG | Radhakrishnan V. S et al |
| **hCoV-19/India/TR-INSACOG-1665500077515/2021** | EPI_ISL_9721768 | | INSACOG-Tripura | National Institute of Biomedical Genomics – INSACOG | Arindam Maitra et al A |
| **hCoV-19/India/ML-1727400000293/2021** | EPI_ISL_3019169 | | INSACOG-MEGHALAYA | National Institute of Biomedical Genomics – INSACOG | Arindam Maitra et al A |
| **hCoV-19/Malaysia/TIDREC-CVD-21-3176/2021** | EPI_ISL_8147386 | | Pejabat Kesihatan Daerah Lembah Pantai | Tropical Infectious Diseases Research & Education Centre (TIDREC), Universiti Malaya | Kim-Kee Tan et al |
| **hCoV-19/India/PB-VRDLGMCPTA-ICMR-INSACOG-465109902/2021** | EPI_ISL_7133348 | | VRDL Government Medical College,Patiala Punjab | NIV Influenza | Dr.Rupinder Bakshi et al |
| **hCoV-19/Malaysia/C19UMB27/2021** | EPI_ISL_3858144 | | HPSF MUAR | UKM Medical Molecular Biology Institute (UMBI) | Mira Farzana binti Mohamad Mokhtar et al |
| **hCoV-19/Hungary/mbl_32759/2021** | EPI_ISL_9625994 | | Hungarian Defence Forces Medical Centre COVID Laboratory | Hungarian Defence Forces Medical Centre Epidemiological and Scientific Research Institute | Csaba István Pereszlényi et al |
| **hCoV-19/Aruba/AW-RIVM-46305/2021** | EPI_ISL_3138380 | | National Institute for Public Health and the Environment (RIVM) | National Institute for Public Health and the Environment (RIVM) | Adam Meijer et al |
| **hCoV-19/Palestine/QIBPS-PS-JER117847/2021** | EPI_ISL_10033915 | | Al-Quds University | Quadram Institute Bioscience | Abdelmajeed Nasereddin et al |
| **hCoV-19/Curacao/CW-RIVM-61653/2021** | EPI_ISL_4845988 | | National Institute for Public Health and the Environment (RIVM) | National Institute for Public Health and the Environment (RIVM) | Adam Meijer et al |
| **hCoV-19/Curacao/CW-RIVM-69762/2021** | EPI_ISL_6852723 | | National Institute for Public Health and the Environment (RIVM) | National Institute for Public Health and the Environment (RIVM) | Adam Meijer et al |
| **hCoV-19/Laos/LOMWRU-0467/2022** | EPI_ISL_15656017 | | LOMWRU | LOMWRU/Microbiology Laboratory, Mahosot Hospital | Manila Souksavanh et al |
| **hCoV-19/Cyprus/BMV1849/2021** | EPI_ISL_16372809 | | Department of Molecular Virology, Cyprus Institute of Neurology and Genetics | Laboratory of Biotechnology and Molecular Virology, University of Cyprus | Leondios G. Kostrikis et al. |
| **hCoV-19/Fujian/2021BD-VIR-SPE-0414/2021** | EPI_ISL_2965604 | | Fujian Center for Disease Control and Prevention | Fujian Center for Disease Control and Prevention | Lin Qi et al |
| **hCoV-19/CzechRepublic/NRL_9855/2021** | EPI_ISL_3318869 | | National Institute of Public Health | National Institute of Public Health | Helena Jirincova et al |
| **hCoV-19/Yunnan/RLD44/2021** | EPI_ISL_13008840 | | Department of Acute Infectious Diseases Control and Prevention, Yunnan Provincial Center for Disease Control and Prevention | Department of Acute Infectious Diseases Control and Prevention, Yunnan Provincial Center for Disease Control and Prevention | Meiling Zhang et al |
| **hCoV-19/Yunnan/RLD201/2021** | EPI_ISL_13008952 | | Department of Acute Infectious Diseases Control and Prevention, Yunnan Provincial Center for Disease Control and Prevention | Department of Acute Infectious Diseases Control and Prevention, Yunnan Provincial Center for Disease Control and Prevention | Meiling Zhang et al |
| **hCoV-19/Yunnan/RLD258/2021** | EPI_ISL_13008971 | | Department of Acute Infectious Diseases Control and Prevention, Yunnan Provincial Center for Disease Control and Prevention | Department of Acute Infectious Diseases Control and Prevention, Yunnan Provincial Center for Disease Control and Prevention | Meiling Zhang et al |
| **hCoV-19/Vietnam/HCMC_b01_82.06/2021** | EPI_ISL_5104583 | | Vietnamese-German Center for Medical Research (VG-CARE), Vietnamese Military Medical University, Hanoi, Ho-Chi-Minh-City | Vietnamese-German Center for Medical Research, VG-CARE | Pham Xuan Huy et al |
| **hCoV-19/Vietnam/NHTD-OUCRU1629/2021** | EPI_ISL_7650431 | | National Hospital for Tropical Diseases | Oxford University Clinical Research Unit, Hanoi, Vietnam | Pham Ngoc Thach et al |
| **hCoV-19/Guangxi/IVDC-575/2022** | EPI_ISL_11873938 | | Guangxi Zhuang Autonomous Region Center for Disease Prevention and Control | National Institute for Viral Disease Control and Prevention, China CDC | Yu Ju et al |
| **hCoV-19/SouthKorea/KDCA27309/2022** | EPI_ISL_9511216 | | Division of Emerging Infectious Diseases, Bureau of Infectious Diseases Diagnosis Control, Korea Disease Control and Prevention Agency | Division of Emerging Infectious Diseases, Bureau of Infectious Diseases Diagnosis Control, Korea Disease Control and Prevention Agency | Jeong-Ah Kim et al |
| **hCoV-19/SouthKorea/KDCA16713/2021** | EPI_ISL_6747178 | | Division of Emerging Infectious Diseases, Bureau of Infectious Diseases Diagnosis Control, Korea Disease Control and Prevention Agency | Division of Emerging Infectious Diseases, Bureau of Infectious Diseases Diagnosis Control, Korea Disease Control and Prevention Agency | Ae Kyung Park et al |
| **hCoV-19/India/ML-1727400000217/2021** | EPI_ISL_3019131 | | INSACOG-MEGHALAYA | National Institute of Biomedical Genomics – INSACOG | Arindam Maitra et al A |
| **hCoV-19/India/ML-1727800002486/2021** | EPI_ISL_4192989 | | INSACOG-Meghalaya | National Institute of Biomedical Genomics – INSACOG | Arindam Maitra et al A |
| **hCoV-19/India/KA-SLS-SEQ-5077/2021** | EPI_ISL_11736478 | | Strand Life Sciences (supported by IN COVID Support, FZE LLC) | Strand Life Sciences (supported by IN COVID Support, FZE LLC) | Priyanka Ananta Mulay et al |
| **hCoV-19/Canada/NL-PHML-216796/2021** | EPI_ISL_5596286 | | NL-Dr. Leonard A. Miller Centre for Health Services | Newfoundland and Labrador - Eastern Health | Anna Majer et al |
| **hCoV-19/Iraq/Erbil-01/2022** | EPI_ISL_16279097 | | Rozhawa Emergency Hospital Lab-Erbil | Salahaddin university-Erbil | Rezhna Kheder Ali et al |
| **hCoV-19/India/AS-INSACOG-1861803018349/2021** | EPI_ISL_9056239 | | INSACOG-Assam | National Institute of Biomedical Genomics – INSACOG | Arindam Maitra et al A |
| **hCoV-19/India/AS-INSACOG-1861803018375/2021** | EPI_ISL_9056240 | | INSACOG-Assam | National Institute of Biomedical Genomics – INSACOG | Arindam Maitra et al A |
| **hCoV-19/Bangladesh/TND-icddrb-06-0008/2021** | EPI_ISL_7571085 | | International Centre for Diarrhoeal Disease Research, Bangladesh (icddr,b) | International Centre for Diarrhoeal Disease Research, Bangladesh (icddr,b) | Afrad MH et al |
| **hCoV-19/Turkey/HSGM-C4485/2021** | EPI_ISL_3548915 | | Ministry of Health Turkey | Ministry of Health Turkey | Fatma Bayrakdar et al |
| **hCoV-19/Zimbabwe/CERI-KRISP-K027252/2021** | EPI_ISL_6699667 | | National Microbiology Reference Laboratory, Ministry of Health, Harare, Zimbabwe | CERI, Centre for Epidemic Response and Innvoation, Stellenbosch University and KRISP, KZN Research Innovation and Sequencing Platform, UKZN. | Air Comodor Dr J. Chimedza et al |
| **hCoV-19/Pakistan/NIH-B21-S13/2021** | EPI_ISL_5427749 | | Department of Virology | Department of Virology | Massab Umair et al |
| **hCoV-19/Serbia/J788010-0609/2021** | EPI_ISL_10134145 | | Huo-yan national laboratory for molecular detection of infectious agents Nis | Institute of Molecular Genetics and Genetic Engineering University of Belgrade | Huang J et al |
| **hCoV-19/Montenegro/CCO-00531_MNE000_5436112102/2021** | EPI_ISL_7381447 | | Laboratory for Molecular Diagnostics,IPHMN | Eurofins Genomics Europe Sequencing GmbH | Danijela Vujošević et al |
| **hCoV-19/India/MN-IBSD-IW-P2-188/2021** | EPI_ISL_11803935 | | Babina Diagnostics | INSACOG-IBSD Manipur | Dr. Nanaocha Sharma et al |
| **hCoV-19/India/OR-ILSGS09915/2021** | EPI_ISL_7877892 | | ILS, Bhubaneswar | Institute of Life Sciences-INSACOG | Sunil K. Raghav et al |
| **hCoV-19/India/MZ-INSACOG-1526100415547/2021** | EPI_ISL_8700689 | | INSACOG-Mizoram | National Institute of Biomedical Genomics – INSACOG | Arindam Maitra et al A |
| **hCoV-19/India/TN-CBR-SEQ-7550/2021** | EPI_ISL_9429757 | | CBR,IISc | CBR,IISc-INSACOG | Bratati Kahali et al |
| **hCoV-19/India/AN-ICMR-NIV-INSACOG-GSEQ-4899/2021** | EPI_ISL_3532948 | | ICMR-National Institute of Virology - INSACOG | NIV Influenza | Potdar et al B |
| **hCoV-19/India/MN-IBSD-IE-P2-141/2021** | EPI_ISL_11803889 | | Babina Diagnostics | INSACOG-IBSD Manipur | Dr. Nanaocha Sharma et al |
| **hCoV-19/India/NL-IBSD-KHM-P2-286/2021** | EPI_ISL_11804031 | | Naga Hospital Authority,Kohima | INSACOG-IBSD Manipur | Dr. Nanaocha Sharma et al |
| **hCoV-19/India/MN-INSACOG-1425500154905/2021** | EPI_ISL_9323588 | | INSACOG-MANIPUR | National Institute of Biomedical Genomics – INSACOG | Arindam Maitra et al A |
| **hCoV-19/India/MN-IBSD-IW-P2-127/2021** | EPI_ISL_11803876 | | Babina Diagnostics | INSACOG-IBSD Manipur | Dr. Nanaocha Sharma et al |
| **hCoV-19/India/AS-1828400133681/2021** | EPI_ISL_2877803 | | INSACOG-Assam | National Institute of Biomedical Genomics – INSACOG | Arindam Maitra et al A |
| **hCoV-19/India/LA-NCDC-590994/2021** | EPI_ISL_11053280 | | National Centre for Disease Control (NCDC) Biotechnology Division, Delhi - INSACOG | NCDC Delhi, Biotechnology Division -INSACOG | Kalaiarasan Ponnusamy et al |
| **hCoV-19/India/OR-ILSGS10287/2021** | EPI_ISL_7877966 | | Angul, DHH | Institute of Life Sciences-INSACOG | Sunil K. Raghav et al |
| **hCoV-19/Greece/255481/2021** | EPI_ISL_3689296 | | Hellenic National Blood Transfusion Center - EKEA | Greek Genome Center, Biomedical Research Foundation of the Academy of Athens (BRFAA) | Emmanouil Athanasiadis et al |
| **hCoV-19/HongKong/HKPU-11942/2021** | EPI_ISL_3039609 | | Department of Health Technology and Informatics, The Hong Kong Polytechnic University | Department of Health Technology and Informatics, The Hong Kong Polytechnic University | Gilman Kit-Hang Siu et al |
| **hCoV-19/India/GA-NCCS-169651/2021** | EPI_ISL_2382959 | | National Centre For Cell Science | National Centre For Cell Science-INSACOG | Dhiraj Paul et al |
| **hCoV-19/India/GJ-INSACOG-GBRC2875/2021** | EPI_ISL_6931527 | | Unipath Speciality Laboratory Limited, Ahmedabad | Gujarat Biotechnology Research Centre | Priyank Chavda et al |
| **hCoV-19/India/MH-IISER-Pune-IP-01044/2021** | EPI_ISL_4415614 | | Department of Microbiology, Government Medical College, Baramati | IISER Pune-INSACOG | Krishanpal Karmodiya et al |
| **hCoV-19/Maldives/MAV98816/2022** | EPI_ISL_9607083 | | Kulhudufushi Regional Hospital | Kulhudufushi Regional Hospital | D. Fathmath Nazla Rafeeq et al |
| **hCoV-19/Oman/rega-OM-129/2021** | EPI_ISL_4062922 | | Sultan Qaboos University Hospital, Department of Microbiology & Immunology, Molecular Biology Section | KU Leuven, Rega Institute, Clinical and Epidemiological Virology | Tony Wawina-Bokalanga et al |
| **hCoV-19/India/DD-INSACOG-GBRC2260/2021** | EPI_ISL_4533560 | | SN Genelab Pvt. Ltd, Surat | Gujarat Biotechnology Research Centre | Apurvasinh Puvar et al |
| **hCoV-19/India/GJ-ICMR-NIV-INSACOG-GSEQ-5085/2021** | EPI_ISL_3533043 | | ICMR-National Institute of Virology - INSACOG | NIV Influenza | Potdar et al B |

| | | | | | |
|---|---|---|---|---|---|
| **hCoV-19/India/KL_KZ-NIV-19-TS_RNA_7639/2021** | EPI_ISL_5104541 | | Indian Council of Medical Research-National Institute of Virology, Microbial Containment Complex | Indian Council of Medical Research-National Institute of Virology, Microbial Containment Complex | Pragya D. Yadav et al |
| **hCoV-19/India/MH-RFIP05656/2021** | EPI_ISL_8145600 | | KEM, Pune | RF-IISER Pune | Krishanpal Karmodiya et al |
| **hCoV-19/Brazil/RJ-LNN06612/2021** | EPI_ISL_6691311 | | Unidade de apoio ao diagnostico da COVID - UNADIG | Bioinformatics Laboratory / LNCC | Luiz G P de Almeida et al |
| **hCoV-19/Brazil/SP-IB_131629/2021** | EPI_ISL_4746669 | | AFIP | Instituto Butantan | Dimas Tadeu Covas et al |
| **hCoV-19/Brazil/RS-LMM67162/2021** | EPI_ISL_7898426 | | Laboratório de Microbiologia Molecular - Universidade FEEVALE | Molecular Microbiology Laboratory | Alana Witt Hansen et al |
| **hCoV-19/Brazil/SC-UFSC-40-68/2021** | EPI_ISL_9246495 | | SMS - SAO FRANCISCO DO SUL | Laboratório de Bioinformática - Universidade Federal de Santa Catarina | "Glauber Wagner et al |
| **hCoV-19/Guatemala/ASI4354/2021** | EPI_ISL_7740586 | | Laboratorio Nacional de Salud | Asociación de Salud Integral / Clínica Familiar "Luis Ángel García" | Eduardo Arathon et al |
| **hCoV-19/Nicaragua/MAN-134082/2021** | EPI_ISL_9159812 | | CNDR | MSHS Pathogen Surveillance Program | Aubree Gordon et al |
| **hCoV-19/Guatemala/ASI4654/2021** | EPI_ISL_7740529 | | Laboratorio Nacional de Salud | Asociación de Salud Integral / Clínica Familiar "Luis Ángel García" | Eduardo Arathon et al |
| **hCoV-19/CostaRica/INC-1319-703408/2021** | EPI_ISL_5262731 | | HOSPITAL SAN JUAN DE DIOS | Inciensa, Instituto Costarricense de Investigación y Enseñanza en Nutrición y Salud | Francisco Duarte et al |
| **hCoV-19/CostaRica/INC-2278-721669/2021** | EPI_ISL_8649077 | | AREA DE SALUD SIQUIRRES | Inciensa, Instituto Costarricense de Investigación y Enseñanza en Nutrición y Salud | Francisco Duarte et al |
| **hCoV-19/Guatemala/INC-LNS-140/2021** | EPI_ISL_8056736 | | LNS | Inciensa, Instituto Costarricense de Investigación y Enseñanza en Nutrición y Salud | César Conde et al |
| **hCoV-19/India/GA-CDFD-C61182/2021** | EPI_ISL_5003944 | | Goa Medical College | CDFD-INSACOG | Murali Bashyam et al |
| **hCoV-19/India/GA-NCCS-C59073/2021** | EPI_ISL_8803714 | | National Centre For Cell Science | National Centre For Cell Science – INSACOG | Dhiraj Dhotre et al |
| **hCoV-19/India/AP-CCMB-CIA7641/2021** | EPI_ISL_3935048 | | AP SSO | CSIR-Centre for Cellular and Molecular Biology - INSACOG | Tulasi Nagabandi et al |
| **hCoV-19/India/TG-CCMB-CIA3136/2021** | EPI_ISL_2661184 | | CSIR-Centre for Cellular and Molecular Biology | CSIR-Centre for Cellular and Molecular Biology-INSACOG | B Himasri et al |
| **hCoV-19/Paraguay/294306/2021** | EPI_ISL_9437444 | | Laboratorio Central de Salud Pública | Laboratorio Central de Salud Pública | Sandra Gonzalez et al |
| **hCoV-19/India/MH-IISER-Pune-IP-01109/2021** | EPI_ISL_4415617 | | Virus Research & Diagnostic Laboratory | IISER Pune-INSACOG | Krishanpal Karmodiya et al |
| **hCoV-19/India/TG-CDFD-127869/2021** | EPI_ISL_6032285 | | Government Fever Hospital | CDFD-INSACOG | Murali Bashyam et al |
| **hCoV-19/Mexico/CHH-CABANA-MX-UGA-IMSS_ITV_UBA_UCR_UdA_UPC-3404/2021** | EPI_ISL_7812903 | | Instituto Mexicano del Seguro Social (IMSS) | CABANA: Unidad de Genómica Avanzada del Centro de Investigación y de Estudios Avanzados (UGA-LANGEBIO, CINVESTAV) | Hebleen Brenes et al |
| **hCoV-19/CostaRica/INC-1334-704195/2021** | EPI_ISL_5262745 | | HOSPITAL DR. ENRIQUE BALTODANO BRICEÑO | Inciensa, Instituto Costarricense de Investigación y Enseñanza en Nutrición y Salud | Francisco Duarte et al |
| **hCoV-19/Mexico/GRO-INMEGEN-33-100/2021** | EPI_ISL_5846099 | | Salud Digna | Instituto Nacional de Medicina Genomica | Cedro-Tanda A et al |
| **hCoV-19/Panama/GMI-PASEQ548/2021** | EPI_ISL_11695697 | | Gorgas Memorial Institute of Health Studies | Gorgas Memorial Institute of Health Studies | Claudia Gonzalez et al |
| **hCoV-19/Panama/GMI-PASEQ539/2021** | EPI_ISL_11695689 | | Gorgas Memorial Institute of Health Studies | Gorgas Memorial Institute of Health Studies | Claudia Gonzalez et al |
| **hCoV-19/Israel/SMC-7019056/2021** | EPI_ISL_6205466 | | Shamir Medical Center (Asaf Harofe) | Shamir Medical Center (Asaf Harofe) | Nir Rainy et al |
| **hCoV-19/Guatemala/ASI4664/2021** | EPI_ISL_7729607 | | Laboratorio Nacional de Salud | Asociación de Salud Integral / Clínica Familiar "Luis Ángel García" | Eduardo Arathon et al |
| **hCoV-19/Guatemala/INC-LNS-042/2021** | EPI_ISL_5169135 | | LNS | Inciensa, Instituto Costarricense de Investigación y Enseñanza en Nutrición y Salud | César Conde et al |
| **hCoV-19/Mexico/GRO-INMEGEN-35-145/2021** | EPI_ISL_6181787 | | Salud Digna | Instituto Nacional de Medicina Genomica | Cedro-Tanda A et al |
| **hCoV-19/Mexico/SLP_IBT_IMSS_3060/2021** | EPI_ISL_4879061 | ON484086.1 | Unidad de Investigación Biomédica de Zacatecas (UIBZ) | Instituto de Biotecnología de la UNAM | Consorcio Mexicano de Vigilancia Genómica (CoViGen-Mex). Authors (in alphabetical order): Julio Elias Alvarado-Yaah et al |
| **hCoV-19/India/TR-329622210/2021** | EPI_ISL_3982889 | | INSACOG-Tripura | National Institute of Biomedical Genomics – INSACOG | Arindam Maitra et al A |
| **hCoV-19/Iraq/Erbil-8/2021** | EPI_ISL_16279109 | | Emirates Hospital Lab-Erbil | Emirates Hospital Lab-Erbil | Rezhna Kheder Ali et al |
| **hCoV-19/India/HR-IGIB-0606300670046/2021** | EPI_ISL_10206123 | | INSACOG at CSIR Institute of Genomics and Integrative Biology | INSACOG at CSIR Institute of Genomics and Integrative Biology | INSACOG et al |
| **hCoV-19/Henan/IVDC-546/2022** | EPI_ISL_11873909 | | Henan Center for Disease Control and Prevention | National Institute for Viral Disease Control and Prevention, China CDC | Hongxia Ma et al |
| **hCoV-19/Beijing/BJCDC-090/2021** | EPI_ISL_8582088 | | Beijing Center for Disease Control and Prevention | Beijing Center for Disease Control and Prevention | Zhichao liang et al |
| **hCoV-19/Azerbaijan/un-ChVir28851/2021** | EPI_ISL_15714272 | | Xususi Tehlukeli Infeksiyalara Nezaret Merkezi | Charité Universitätsmedizin Berlin, Institut für Virologie | Victor M Corman et al |
| **hCoV-19/India/HP-NCDC-780938/2021** | EPI_ISL_11059148 | | National Centre for Disease Control (NCDC) Biotechnology Division, Delhi - INSACOG | NCDC Delhi, Biotechnology Division -INSACOG | Kalaiarasan Ponnusamy et al |
| **hCoV-19/India/KA-SEQ-2748_S105_R1_001/2021** | EPI_ISL_3275844 | | Dr. Madhusudhan, MBBD, MD, ICMR VRDL, Wenlock District Hospital | inStem NCBS – INSACOG | Uma Ramakrishnan Dasaradhi Palakodeti Aswin SaiNarain et al |
| **hCoV-19/Guinea/IPG-24294/2021** | EPI_ISL_10312004 | | Institut Pasteur de Guinée | Institut Pasteur de Guinée | Solène Grayo et al |
| **hCoV-19/Guinea/IP-GNC_22718/2021** | EPI_ISL_3988395 | | Institut Pasteur de Guinée | Institut Pateur de Dakar | Grayo Solene et al |
| **hCoV-19/CaboVerde/ST40495/2021** | EPI_ISL_14093693 | | Virology Laboratory of Praia | Instituto Nacional de Saúde Pública (INSP) | Kevin Sanders et al |
| **hCoV-19/India/WB-INSACOG-1931300485234/2021** | EPI_ISL_6885411 | | INSACOG-WB | National Institute of Biomedical Genomics – INSACOG | Arindam Maitra et al A |
| **hCoV-19/India/GA-NCCS-C51024/2021** | EPI_ISL_3315050 | | National Centre For Cell Science | National Centre For Cell Science – INSACOG | Dhiraj Paul et al |
| **hCoV-19/India/MP-NCDC-5309189/2021** | EPI_ISL_11051827 | | National Centre for Disease Control (NCDC) Biotechnology Division, Delhi - INSACOG | NCDC Delhi, Biotechnology Division -INSACOG | Mahesh S Dhar et al |
| **hCoV-19/Sudan/ILRI_M03038/2021** | EPI_ISL_17971563 | | Epidemiology Lab, National Public Health | International Livestock Research Institute | Samuel O. Oyola et al |
| **hCoV-19/India/MZ-INSACOG-1526700038137/2021** | EPI_ISL_9056580 | | INSACOG-Mizoram | National Institute of Biomedical Genomics – INSACOG | Arindam Maitra et al A |
| **hCoV-19/India/OR-ILSGS12178/2021** | EPI_ISL_7878161 | | Nayagarh, DHH | Institute of Life Sciences-INSACOG | Sunil K. Raghav et al |
| **hCoV-19/India/JH-ILSGS12163/2021** | EPI_ISL_7878153 | | RIMS,Ranchi | Institute of Life Sciences-INSACOG | Sunil K. Raghav et al |
| **hCoV-19/India/JH-ILSGS10236/2021** | EPI_ISL_7877950 | | Department of Pathology, Tata Main Hospital, Jamshedpur | Institute of Life Sciences-INSACOG | Sunil K. Raghav et al |
| **hCoV-19/India/JH-ILSGS12088/2021** | EPI_ISL_7878135 | | RIMS,Ranchi | Institute of Life Sciences-INSACOG | Sunil K. Raghav et al |
| **hCoV-19/India/MP-NCDC-560955/2021** | EPI_ISL_11053899 | | National Centre for Disease Control (NCDC) Biotechnology Division, Delhi - INSACOG | NCDC Delhi, Biotechnology Division -INSACOG | Mahesh S Dhar et al |
| **hCoV-19/India/AS-INSACOG-1829700709641/2021** | EPI_ISL_9056375 | | INSACOG-Assam | National Institute of Biomedical Genomics – INSACOG | Arindam Maitra et al A |
| **hCoV-19/India/NL-NIBMG-1324900003201/2021** | EPI_ISL_2675364 | | INSACOG-NAGALAND | National Institute of Biomedical Genomics – INSACOG | Arindam Maitra et al A |
| **hCoV-19/India/NL-1324900004894/2021** | EPI_ISL_3189528 | | INSACOG-NAGALAND | National Institute of Biomedical Genomics – INSACOG | Arindam Maitra et al A |
| **hCoV-19/India/LD-ICMR-NIV-INSACOG-GSEQ-5204/2021** | EPI_ISL_3982438 | | ICMR-National Institute of Virology - INSACOG | NIV Influenza | Potdar et al B |
| **hCoV-19/India/LD-ICMR-NIV-INSACOG-GSEQ-5218/2021** | EPI_ISL_3982452 | | ICMR-National Institute of Virology - INSACOG | NIV Influenza | Potdar et al B |
| **hCoV-19/India/LD-ICMR-NIV-INSACOG-GSEQ-5032/2021** | EPI_ISL_3533002 | | ICMR-National Institute of Virology - INSACOG | NIV Influenza | Potdar et al B |
| **hCoV-19/Slovenia/08-155850-MB/2021** | EPI_ISL_4252757 | | National Laboratory for Health, Environment and Food, OMM, Maribor | NLZOH (National Laboratory for Health, Environment and Food) / CISLD (Clinical Institute of Special Laboratory Diagnostics), University Children's Hospital, University Medical Center Ljubljana | Sandra Janezic et al |
| **hCoV-19/India/TR-AGMC/COV/R47129/2021** | EPI_ISL_2878018 | | INSACOG-Tripura | National Institute of Biomedical Genomics – INSACOG | Arindam Maitra et al A |
| **hCoV-19/Oman/rega-OM-180/2021** | EPI_ISL_4063018 | | Sultan Qaboos University Hospital, Department of Microbiology & Immunology, Molecular Biology Section | KU Leuven, Rega Institute, Clinical and Epidemiological Virology | Tony Wawina-Bokalanga et al |
| **hCoV-19/India/PY-COv2-51704_S15_R1_001/2021** | EPI_ISL_5806966 | | JIPMER Puducherry | inStem NCBS – INSACOG | Uma Ramakrishnan Dasaradhi Palakodeti Aswin SaiNarain et al |
| **hCoV-19/Georgia/Tb-s1049/2021** | EPI_ISL_8885100 | | Department for Virology, Molecular Biology and Genome Research, R. G. Lugar Center for Public Health Research, National Center for Disease Control and Public Health (NCDC) of Georgia. | Department for Virology, Molecular Biology and Genome Research, R. G. Lugar Center for Public Health Research, National Center for Disease Control and Public Health (NCDC) of Georgia. | Giorgi Tomashvili et al |
| **hCoV-19/Sweden/SUS0004971/2021** | EPI_ISL_4762130 | | Clinical Microbiology, Infection Prevention and Control | Section for Molecular Diagnostics | Björn Hallström et al |
| **hCoV-19/Cameroon/CERI-KRISP-K026437/2021** | EPI_ISL_6308498 | | Africa_CDC - Cameroon | KRISP, KZn Research Innovation and Sequencing Platform | Dr Essomba Rene Ghislain et al |
| **hCoV-19/Algeria/79234/2021** | EPI_ISL_8421066 | | NIC, Viral Respiratory Unit | Virology Departement | Sarah Belkalem et al |
| **hCoV-19/Algeria/87743/2021** | EPI_ISL_15928134 | | NIC, Viral Respiratory Unit | NIC, Viral Respiratory Unit | Mohamed Amine Beloufa et al |
| **hCoV-19/Iraq/KR-SEARCH-118887/2021** | EPI_ISL_11999793 | | University of Duhok COVID-19 Center for Research and Diagnosis | Andersen lab at Scripps Research | SEARCH Alliance with Jassim Mohammed Abdo et al |
| **hCoV-19/Lebanon/NIC52/2022** | EPI_ISL_10969489 | | National Influenza Center - Rafic Hariri University Hospital | National Influenza Center Lebanon Rafik Hariri University Hospital | Mona Albuaini et al |
| **hCoV-19/Zhejiang/ZPCDCP-37/2021** | EPI_ISL_17261539 | | Zhejiang Provincial Center for Disease Control and Prevention | Zhejiang Provincial Center for Disease Control and Prevention | Yi Sun et al |
| **hCoV-19/Morocco/20358839/2021** | EPI_ISL_4741171 | | Medical Virology and BSL3 Laboratory | Centre de Séquençage Génomique | EL Hamouchi Adil et al |
| **hCoV-19/Burundi/UG522/2021** | EPI_ISL_2928006 | | National Institute of Public Health Burundi, WHO Burundi, ASLM Burundi, MRC/UVRI & LSHTM Uganda Research Unit | MRC/UVRI & LSHTM Uganda Research Unit, National Institute of Public Health Burundi, WHO Burundi, ASLM Burundi | Jerome Nkurunziza et al |
| **hCoV-19/Australia/NSW-R0377/2021** | EPI_ISL_2274250 | | New South Wales Health Pathology Royal Prince Alfred Hospital | Microbiology RPAH | Foster et al |
| **hCoV-19/Rwanda/NRLNAT4209/2021** | EPI_ISL_4415301 | | Nucleic Acid Testing, National Reference Laboratory | GIGA Medical Genomics | Yvan Butera et al |
| **hCoV-19/Uganda/UG742/2021** | EPI_ISL_6332929 | | MRC/UVRI & LSHTM Uganda Research Unit , Rakai Health Sciences Program | MRC/UVRI & LSHTM Uganda Research Unit , Rakai Health Sciences Program | Matthew Cotten Dan Lule Bugembe et al |
| **hCoV-19/NewZealand/21MV0496/2021** | EPI_ISL_2650017 | | LabPLUS | Institute of Environmental Science and Research (ESR) | Rachel Boyle et al |
| **hCoV-19/SouthSudan/UG688/2021** | EPI_ISL_3546327 | | South Sudan Ministry of Health, WHO South Sudan, MRC/UVRI & LSHTM Uganda Research Unit | MRC/UVRI & LSHTM Uganda Research Unit, South Sudan Ministry of Health, WHO South Sudan | Joseph Francis Wamala et al |
| **hCoV-19/DRC/INRB-RDC-1157/2021** | EPI_ISL_15297199 | | Laboratoire Rodolphe Mérieux (INRB Goma) | Pathogen Genomics Lab, National Institute for Biomedical Research (INRB) | Placide Mbala-Kingebeni et al B |
| **hCoV-19/SouthSudan/UG686/2021** | EPI_ISL_3546325 | | South Sudan Ministry of Health, WHO South Sudan, MRC/UVRI & LSHTM Uganda Research Unit | MRC/UVRI & LSHTM Uganda Research Unit, South Sudan Ministry of Health, WHO South Sudan | Joseph Francis Wamala et al |
| **hCoV-19/SouthSudan/UG1089/2021** | EPI_ISL_10512551 | | South Sudan Ministry of Health, WHO South Sudan, MRC/UVRI & LSHTM Uganda Research Unit | MRC/UVRI & LSHTM Uganda Research Unit, South Sudan Ministry of Health, WHO South Sudan | Abe G Abias et al |
| **hCoV-19/Djibouti/NAMRU3_E65/2021** | EPI_ISL_6693783 | OL640819.1 | Naval Medical Research Unit No. 3 | Naval Medical Research Center Biological Defense Research Directorate | Catherine E. Arnold et al |
| **hCoV-19/SouthSudan/UG1247/2021** | EPI_ISL_10512594 | | South Sudan Ministry of Health, WHO South Sudan, MRC/UVRI & LSHTM Uganda Research Unit | MRC/UVRI & LSHTM Uganda Research Unit, South Sudan Ministry of Health, WHO South Sudan | Abe G Abias et al |
| **hCoV-19/SouthSudan/UG1095/2021** | EPI_ISL_10512529 | | South Sudan Ministry of Health, WHO South Sudan, MRC/UVRI & LSHTM Uganda Research Unit | MRC/UVRI & LSHTM Uganda Research Unit, South Sudan Ministry of Health, WHO South Sudan | Abe G Abias et al |
| **hCoV-19/Guinea/LFHVG-G0470/2021** | EPI_ISL_9569736 | | Laboratoire des Fievres Hemorragiques Virales en Guinee, LFHVG | "Laboratoire des Fievres Hemorragiques Virales en Guinee, LFHVG, and the Bernhard Nocht Institute for Tropical Medicine, BNITM" | "N'Faly Magassouba et al |
| **hCoV-19/Chad/TCH-30/2021** | EPI_ISL_9274867 | | Ministère de Santé Publique et de la Solidarité Nationale | Pathogen Genomics Lab, National Institute for Biomedical Research (INRB) | Placide Mbala-Kingebeni et al B |
| **hCoV-19/Israel/CVL-22972/2021** | EPI_ISL_7133825 | | Israel Central Virology laboratory | Israel Central Virology laboratory | Neta Zuckerman et al |
| **hCoV-19/Ukraine/Kyiv-10/2021** | EPI_ISL_5852911 | | L.V. Gromashevsky Institute of Epidemiology and Infectious Diseases of the NAMS of Ukraine | L.V. Gromashevsky Institute of Epidemiology and Infectious Diseases of the NAMS of Ukraine | Anna Yakovleva et al |
| **hCoV-19/EquatorialGuinea/C21093/2021** | EPI_ISL_10525211 | | Laboratorio de Investigaciones de Baney | Swiss Tropical and Public Health Institute | Salome Hosch et al |
| **hCoV-19/Benin/NMIMR-LCA5328/2021** | EPI_ISL_7139829 | | National Reference Laboratory, Laboratories Department, Cotonou, Benin | Noguchi Memorial Institute for Medical Research, University of Ghana, Legon, Ghana | Rene K. Keke et al |
| **hCoV-19/Mexico/TLA_IBT_IMSS_4794/2021** | EPI_ISL_7969909 | | Laboratorio Central de Epidemiología (LCE) | Instituto de Biotecnología de la UNAM | Consorcio Mexicano de Vigilancia Genómica (CoViGen-Mex). Authors (in alphabetical order): Julio Elias Alvarado-Yaah et al |
| **hCoV-19/Mexico/ZAC_InDRE_FB39240_S7224/2021** | EPI_ISL_4603188 | | LESP Zacatecas | Instituto de Diagnostico y Referencia Epidemiologicos (INDRE) | Claudia Wong-Arambula et al |
| **hCoV-19/Mexico/AGU-InDRE_FB18596_S4465/2021** | EPI_ISL_2937902 | | LESP Aguascalientes | Instituto de Diagnostico y Referencia Epidemiologicos (INDRE) | Claudia Wong-Arambula et al |
| **hCoV-19/Mexico/TLA_InDRE_FB38243_S7066/2021** | EPI_ISL_4603024 | | LESP Tlaxcala | Instituto de Diagnostico y Referencia Epidemiologicos (INDRE) | Claudia Wong-Arambula et al |
| **hCoV-19/Mexico/GUA_InDRE_FB41977_S7607/2021** | EPI_ISL_4918515 | | LESP Guanajuato | Instituto de Diagnostico y Referencia Epidemiologicos (INDRE) | Claudia Wong-Arambula et al |
| **hCoV-19/Mexico/TLA_InDRE_FB46175_S8530/2021** | EPI_ISL_6097659 | | LESP Tlaxcala | Instituto de Diagnostico y Referencia Epidemiologicos (INDRE) | Claudia Wong-Arambula et al |
| **hCoV-19/Colombia/TOL-INS-VG-11844/2021** | EPI_ISL_8562354 | | ANALICEMOS | Instituto Nacional de Salud- Dirección de Investigación en Salud Pública | Katherine Laiton-Donato et al |
| **hCoV-19/Mexico/MIC-InDRE_FB50352_S9431/2021** | EPI_ISL_6810254 | | LESP Michoacan | Instituto de Diagnostico y Referencia Epidemiologicos (INDRE) | Claudia Wong-Arambula et al |
| **hCoV-19/Malta/MDxMDH680/2021** | EPI_ISL_7458716 | | Molecular Diagnostics Pathology Department Mater Dei Hospital Malta | Molecular Diagnostics Pathology Department Mater Dei Hospital Malta | G Zahra et al |
| **hCoV-19/Kenya/C101636/2021** | EPI_ISL_3049666 | OK017218.1 | KEMRI-Wellcome Trust Research Programme,Kilifi | KEMRI-Wellcome Trust Research Programme,Kilifi | Githinji G. et al |
| **hCoV-19/Zambia/ZMB-121239/2021** | EPI_ISL_8025550 | | Right to Care Zambia | UNZAVET and PATH | Mulenga Mwenda et al |
| **hCoV-19/Zambia/ZMB-116115/2021** | EPI_ISL_4513737 | | University of Zambia, School of Veterinary Medicine | UNZAVET and PATH | Mulenga Mwenda-Chimfwembe et al |
| **hCoV-19/India/AS-1828000314208/2021** | EPI_ISL_3189064 | | INSACOG-Assam | National Institute of Biomedical Genomics – INSACOG | Arindam Maitra et al A |
| **hCoV-19/NorthMacedonia/16TE/2021** | EPI_ISL_3545709 | | Laboratory of virology and molecular diagnostics, Institute of Public Health | Laboratory of virology and molecular diagnostics, Institute of Public Health | Kuzmanovska M et al |
| **hCoV-19/Belgium/UZA-UA-1501510800/2021** | EPI_ISL_6135635 | | Algemeen Klinisch Labo | Labo Klinische Biologie, UZA | Marie Le Mercier et al |
| **hCoV-19/Iran/AK241/2021** | EPI_ISL_10586571 | | COVID-19 National Reference Laboratory, Pasteur Institute of Iran | COVID-19 National Reference Laboratory, Pasteur Institute of Iran | Zahra Hosseini et al |
| **hCoV-19/India/TR-INSACOG-565336602/2021** | EPI_ISL_8661424 | | INSACOG-Tripura | National Institute of Biomedical Genomics – INSACOG | Arindam Maitra et al A |
| **hCoV-19/India/TR-INSACOG-465441737/2021** | EPI_ISL_6395387 | | INSACOG-Tripura | National Institute of Biomedical Genomics – INSACOG | Arindam Maitra et al A |
| **hCoV-19/India/JH-ILSGS01707/2021** | EPI_ISL_2342720 | | MGM Medical College & Hospital, Jamshedpur | Institute of Life Sciences - INSACOG | Sunil K. Raghav et al |
| **hCoV-19/UnitedArabEmirates/BTC-5785/2021** | EPI_ISL_16802272 | | Khalifa University Center for Biotechnology | Khalifa University Center for Biotechnology | AlSafar et al. |
| **hCoV-19/Beijing/BJCDC-105/2021** | EPI_ISL_8582065 | | Beijing Center for Disease Control and Prevention | Beijing Center for Disease Control and Prevention | Yang Pan et al |
| **hCoV-19/Andorra/AND-251_220121429201_COV-GC/2021** | EPI_ISL_10122608 | | Laboratori d'anàlisis clíniques, Hospital Nostra Senyora de Meritxell | LBM de CHU de Toulouse, Hôpitaux de Toulouse | Rendon M et al |
| **hCoV-19/Andorra/AND-241_213431458901_COV-GC/2021** | EPI_ISL_8185531 | | Laboratori d'anàlisis clíniques, Hospital Nostra Senyora de Meritxell | LBM de CHU de Toulouse, Hôpitaux de Toulouse | Rendon M et al |
| **hCoV-19/Guadeloupe/IPP30329/2021** | EPI_ISL_5718395 | | Outre Mer | National Reference Center for Viruses of Respiratory Infections, Institut Pasteur, Paris | Marion Barbet et al A |
| **hCoV-19/Zhejiang/ZPCDCP-23/2021** | EPI_ISL_17261525 | | Zhejiang Provincial Center for Disease Control and Prevention | Zhejiang Provincial Center for Disease Control and Prevention | Yi Sun et al |
| **hCoV-19/Colombia/DC-INS-VG-9772/2021** | EPI_ISL_7314412 | | ALIFE HEALTH S.A.S. | Instituto Nacional de Salud- Dirección de Investigación en Salud Pública | Katherine Laiton-Donato et al |
| **hCoV-19/Colombia/VAC-LS-14134303293/2021** | EPI_ISL_8173240 | | Laboratorio Angel Diagnostica sede 100 ( Synlab) | CIAT, Laboratorio de Virologia | Ana M. Leiva et al |
| **hCoV-19/Egypt/CCHE57357_Wave_4_070/2021** | EPI_ISL_6011741 | OL351370.1 | Genomics Program, Children Cancer Hospital | Genomics Program, Children Cancer Hospital | Hatem et al |
| **hCoV-19/Djibouti/NAMRU3_D464/2021** | EPI_ISL_6693747 | OL640784.1 | Naval Medical Research Unit No. 3 | Naval Medical Research Center Biological Defense Research Directorate | Catherine E. Arnold et al |
| **hCoV-19/India/GA-CDFD-C61282/2021** | EPI_ISL_5003952 | | Goa Medical College | CDFD-INSACOG | Murali Bashyam et al |
| **hCoV-19/India/MN-1425500077560/2021** | EPI_ISL_3024886 | | INSACOG-MANIPUR | National Institute of Biomedical Genomics – INSACOG | Arindam Maitra et al A |
| **hCoV-19/Kazakhstan/NLA-16-57/2021** | EPI_ISL_18595947 | | Center for Life Sciences, National Laboratory Astana | Center for Life Sciences, National Laboratory Astana | Kairov et al |
| **hCoV-19/India/UT-NCDC-4609145/2021** | EPI_ISL_11049265 | | National Centre for Disease Control (NCDC) Biotechnology Division, Delhi - INSACOG | NCDC Delhi, Biotechnology Division -INSACOG | Robin Marwal et al |
| **hCoV-19/Nepal/EPN-K2066/2021** | EPI_ISL_12956274 | | Koshi Hospital | Center for Molecular Dynamics Nepal | Epidemic Intelligence Team et al |
| **hCoV-19/India/TR-AGMC/COV/R36187/2021** | EPI_ISL_2877959 | | INSACOG-Tripura | National Institute of Biomedical Genomics – INSACOG | Arindam Maitra et al A |
| **hCoV-19/Mozambique/COV-2258-A1906/2021** | EPI_ISL_5822698 | | Instiuto Nacional de Saude, Mozambique | Forschungszentrum Borstel | Christian Utpatel et al |
| **hCoV-19/India/WB-1931501931042/2021** | EPI_ISL_4193366 | | INSACOG-WB | National Institute of Biomedical Genomics – INSACOG | Arindam Maitra et al A |
| **hCoV-19/Lebanon/LAU-CVD-258/2021** | EPI_ISL_11327145 | | Laboratory of Molecular Biology and Cancer Immunology, Faculty of Sciences, Lebanese University | Microbial Pathogenomics Lab - LAU | Sima tokajian et al |
| **hCoV-19/Mozambique/CERI-KRISP-K027537/2021** | EPI_ISL_9084391 | | Instituto Nacional de Saude (INS), Mozambique | CERI, Centre for Epidemic Response and Innvoation, Stellenbosch University and KRISP, KZN Research Innovation and Sequencing Platform, UKZN. | Nalia Ismael et al |
| **hCoV-19/Fujian/2021BD-VIR-SPE-1269/2021** | EPI_ISL_15756110 | | Fujian Center for Disease Control and Prevention | Fujian Center for Disease Control and Prevention | Lin Qi et al |
| **hCoV-19/India/LA-NCDC-5909103/2021** | EPI_ISL_11053288 | | National Centre for Disease Control (NCDC) Biotechnology Division, Delhi - INSACOG | NCDC Delhi, Biotechnology Division -INSACOG | Kalaiarasan Ponnusamy et al |
| **hCoV-19/Togo/8280/2021** | EPI_ISL_12521340 | | LNR/CHU-SO | Unité Mixte Internationale TransVIHMI (UMI 233 IRD – U1175 INSERM - Université de Montpellier) IRD (Institut de recherche pour le développement) | A.S. COMLAN et al |
| **hCoV-19/Togo/NMIMR-21-TGS-269/2021** | EPI_ISL_3914994 | | Institut National D'hygiene, Lome, Togo | Noguchi Memorial Institute for Medical Research, University of Ghana, Legon, Ghana | Afiwa W. Halatoko et al |
| **hCoV-19/Tunisia/S-1563/2021** | EPI_ISL_11334075 | | Laboratoire de Microbiologie, Hôpital Charles Nicolle de Tunis | Laboratory of Clinical Virology, Institut Pasteur de Tunis | Boutiba Ilhem et al |
| **hCoV-19/Tunisia/F-0672/2021** | EPI_ISL_8298390 | | Laboratoire de Virologie Clinique, Institut Pasteur de Tunis | Laboratory of Clinical Virology, Institut Pasteur de Tunis | Haddad-Boubaker Sondes et al |
| **hCoV-19/Tunisia/Tunis-13845/2021** | EPI_ISL_18516117 | | Laboratoire de Microbiologie, HOPITAL D'ENFANTS BECHIR HAMZA DE TUNIS | Laboratoire de Virologie Clinique, Institut Pasteur de Tunis | Smaoui Hanene et al |
| **hCoV-19/Tunisia/S-1319/2021** | EPI_ISL_11333966 | | Laboratoire de Microbiologie, Hôpital Charles Nicolle de Tunis | Laboratory of Clinical Virology, Institut Pasteur de Tunis | Boutiba Ilhem et al |
| **hCoV-19/Madagascar/LA2M-142003/2021** | EPI_ISL_13729314 | | Laboratoire d'Analyses Médicales Malagasy (LA2M) | Virology Unit, Institut Pasteur de Madagascar | Vololoniaina Raharinosy et al |
| **hCoV-19/Madagascar/LA2M-165254/2021** | EPI_ISL_13729331 | | Laboratoire d'Analyses Médicales Malagasy (LA2M) | Virology Unit, Institut Pasteur de Madagascar | Vololoniaina Raharinosy et al |
| **hCoV-19/Finland/10459/2021** | EPI_ISL_3644828 | | Department of Virology and Immunology, University of Helsinki and Helsinki University Hospital, Huslab Finland | Department of Virology, Faculty of Medicine, University of Helsinki, Helsinki, Finland | Teemu Smura et al |
| **hCoV-19/Turkey/HSGM-F6003/2021** | EPI_ISL_7741101 | | Ministry of Health Turkey | Ministry of Health Turkey | Fatma Bayrakdar et al |
| **hCoV-19/Guinea/CERFIG-00010/2021** | EPI_ISL_4219403 | | Clinical specimen | Centre de Recherche et de Formation en Infectiologie Guinée | Alpha Kabinet KEITA et al |
| **hCoV-19/Benin/439661/2021** | EPI_ISL_4566887 | | Laboratoire des Fièvres Hémorragiques Virales du Benin | Institut für Virologie - Institute of Virology - Charite | Anges Yadouleton et al |
| **hCoV-19/Shandong/Yantai_03/2021** | EPI_ISL_8525419 | | Yantai Center for Disease Control and Prevention | Yantai Center for Disease Control and Prevention | Yulou Sun et al |
| **hCoV-19/Armenia/IMB6-6/2021** | EPI_ISL_3543622 | | National Center of Disease Control and Prevention of the Republic of Armenia, Davidyants Laboratories, Yerevan, Armenia | Institute of Molecular Biology NAS RA, Republic of Armenia, Department of Bioengineering, BioinformaticsInstitute and Molecular Biology IBMPh RAU, Republic of Armenia | Arsen Arakelyan et al |
| **hCoV-19/Armenia/IMB11-12/2021** | EPI_ISL_9018786 | | National Center of Disease Control and Prevention of the Republic of Armenia, Davidyants Laboratories, Yerevan, Armenia | Institute of Molecular Biology NAS RA, Republic of Armenia, Department of Bioengineering, BioinformaticsInstitute and Molecular Biology IBMPh RAU, Republic of Armenia | Arsen Arakelyan et al |
| **hCoV-19/Seychelles/SS3591/2021** | EPI_ISL_5943375 | OL352005.1 | Seychelles Public Health Laboratory | KEMRI-Wellcome Trust Research Programme,Kilifi | Githinji G. et al |
| **hCoV-19/Cyprus/BMV1598/2021** | EPI_ISL_16372659 | | Department of Molecular Virology, Cyprus Institute of Neurology and Genetics | Laboratory of Biotechnology and Molecular Virology, University of Cyprus | Leondios G. Kostrikis et al. |
| **hCoV-19/Jiangsu/XSG/2021** | EPI_ISL_7876606 | | Institute of pathogenic microbiology, Jiangsu Provincial Center for Disease Control and Prevention | Institute of pathogenic microbiology, Jiangsu Provincial Center for Disease Control and Prevention | lunbiao cui et al |
| **hCoV-19/Azerbaijan/un-ChVir28860/2021** | EPI_ISL_15714280 | | Xususi Tehlukeli Infeksiyalara Nezaret Merkezi | Charité Universitätsmedizin Berlin, Institut für Virologie | Victor M Corman et al |
| **hCoV-19/Moldova/D0074/2021** | EPI_ISL_10512081 | | MedExpert | MedExpert | Vadim Nirca et al |
| **hCoV-19/Curacao/CW-RIVM-43129/2021** | EPI_ISL_3056751 | | National Institute for Public Health and the Environment (RIVM) | National Institute for Public Health and the Environment (RIVM) | Adam Meijer et al |
| **hCoV-19/Israel/SMC-7004858/2021** | EPI_ISL_3019823 | | Shamir Medical Center (Asaf Harofe) | Shamir Medical Center (Asaf Harofe) | Nir Rainy et al |
| **hCoV-19/Heilongjiang/IVDC-551/2022** | EPI_ISL_11873914 | | HeiLongjiang Provincial Center For Disease Control And Prevention | National Institute for Viral Disease Control and Prevention, China CDC | Qi Liang et al |
| **hCoV-19/Hainan/IVDC-609/2022** | EPI_ISL_12030374 | | Hainan Provincial Center for Disease Control and Prevention | National Institute for Viral Disease Control and Prevention, China CDC | Chuyang Sun et al |
| **hCoV-19/Netherlands/NB-RIVM-45037/2021** | EPI_ISL_3137109 | | National Institute for Public Health and the Environment (RIVM) | National Institute for Public Health and the Environment (RIVM) | Adam Meijer et al |
| **hCoV-19/Mongolia/10011820/2021** | EPI_ISL_10899724 | OM961246.1 | Biotechnology Core Facility Branch, Centers for Disease Control and Prevention | Centers for Disease Control and Prevention Division of Viral Diseases, Pathogen Discovery | Anir Enkhbat et al |
| **hCoV-19/InnerMongolia/IVDC-597/2022** | EPI_ISL_12030362 | | The center for disease control and prevention of Inner Mongolia | National Institute for Viral Disease Control and Prevention, China CDC | Yan Hai et al |
| **hCoV-19/Moldova/A0047_2021_10_22/2021** | EPI_ISL_7677990 | | Moldova, Chisinau, MedExpert | Moldova, Chisinau, MedExpert | Morozov Alexandr et al |
| **hCoV-19/Heilongjiang/IVDC-HH-455/2021** | EPI_ISL_10925363 | | Heilongjiang Center for Disease Control and Prevention | National Institute for Viral Disease Control and Prevention, China CDC | Xinfa Pei et al |
| **hCoV-19/USA/NE-NPHL-S428156/2021** | EPI_ISL_6261209 | | Nebraska Public Health Laboratory | Nebraska Public Health Laboratory | NPHL COVID-19 Response Team et al |
| **hCoV-19/Azerbaijan/un-ChVir28862/2021** | EPI_ISL_15714282 | | Xususi Tehlukeli Infeksiyalara Nezaret Merkezi | Charité Universitätsmedizin Berlin, Institut für Virologie | Victor M Corman et al |

| | | | | | |
|---|---|---|---|---|---|
| **hCoV-19/Bonaire/BQ-RIVM-73762/2021** | EPI_ISL_7785046 | | National Institute for Public Health and the Environment (RIVM) | National Institute for Public Health and the Environment (RIVM) | Adam Meijer et al |
| **hCoV-19/Crimea/CRIE-L219D0006/2021** | EPI_ISL_3996687 | | Molecular Diagnostic Laboratory of Federal Budget Institution of Science at the Central Research Institute of Epidemiology of the Federal Service on Customers' Rights Protection and Human Well-being Surveillance | Group of Genomics and Postgenomic Technologies of Central Research Institute of Epidemiology | Samoilov A.E. et al |
| **hCoV-19/Hunan/CS-LSI005/2021** | EPI_ISL_16138063 | | Life Sciences Institute, Sansure | Life Sciences Institute, Sansure | He et al |
| **hCoV-19/Shandong/Yantai_01/2021** | EPI_ISL_8525417 | | Yantai Center for Disease Control and Prevention | Yantai Center for Disease Control and Prevention | Yulou Sun et al |
| **hCoV-19/Anhui/WHCDC-001/2021** | EPI_ISL_12901913 | | Wuhu Center for Disease Control and Prevention | Wuhu Center for Disease Control and Prevention | Gongfei Wang et al |
| **hCoV-19/Liaoning/IVDC-SY-220/2021** | EPI_ISL_9301323 | | Liaoning Center for Disease Control and Prevention | National Institute for Viral Disease Control and Prevention, China CDC | Haibo Sun et al |
| **hCoV-19/Russia/KO-3433/2021** | EPI_ISL_14842468 | | Molecular Genetic Monitoring Group | Molecular Genetic Monitoring Group | Anna S. Gladkikh et al |
| **hCoV-19/Jiangxi/IVDC-10-10/2021** | EPI_ISL_8295291 | | Jiangxi Provincial Center for Disease Control and Prevention | National Institute for Viral Disease Control and Prevention, China CDC | Tian Gong et al |
| **hCoV-19/Ecuador/NIC-INSPI-15380/2021** | EPI_ISL_6569632 | | INSPI-CRN DE INFLUENZA Y OTROS VIRUS RESPIRATORIOS | INSPI-CRN DE INFLUENZA Y OTROS VIRUS RESPIRATORIOS | Alfredo Bruno et al |
| **hCoV-19/Fujian/2021KT-VIR-SPE-0004/2021** | EPI_ISL_15756126 | | Fujian Center for Disease Control and Prevention | Fujian Center for Disease Control and Prevention | Lin Qi et al |
| **hCoV-19/Kazakhstan/NLA-04-80/2021** | EPI_ISL_18595431 | | Center for Life Sciences, National Laboratory Astana | Center for Life Sciences, National Laboratory Astana | Kairov et al |
| **hCoV-19/Kazakhstan/NLA-06-33/2021** | EPI_ISL_18595469 | | Center for Life Sciences, National Laboratory Astana | Center for Life Sciences, National Laboratory Astana | Kairov et al |
| **hCoV-19/Sweden/4054049264/2021** | EPI_ISL_5103969 | | Sweden/Sodermanland/Unilabs | Unilabs/Eskilstuna/Sweden | Emma Arvidsson et al |
| **hCoV-19/Montenegro/CO-00421_MNE000_5300242102/2021** | EPI_ISL_7241383 | | Laboratory for Molecular Diagnostics,IPHMN | Eurofins Genomics Europe Sequencing GmbH | Danijela Vujošević et al |
| **hCoV-19/Chad/TCH-29/2021** | EPI_ISL_9274866 | | Ministère de Santé Publique et de la Solidarité Nationale | Pathogen Genomics Lab, National Institute for Biomedical Research (INRB) | Placide Mbala-Kingebeni et al B |
| **hCoV-19/Germany/HH-hpi-p9401/2021** | EPI_ISL_4939621 | | Labor Dr. Fenner und Kollegen | Heinrich Pette Institute, Leibniz Institute for Experimental Virology | Alexis Robitaille et al |
| **hCoV-19/Denmark/DCGC-197525/2021** | EPI_ISL_5916383 | OW397069.1 | Department of Bacteria, Parasites and Fungi, Statens Serum Institut, Copenhagen, Denmark | Statens Serum Institut Bioinformatics and Microbial Genomics | Povilas Matusevicius on behalf of the Danish Covid-19 Genome Consortium et al |
| **hCoV-19/Heilongjiang/IVDC-HH-381/2021** | EPI_ISL_10925289 | | Heilongjiang Center for Disease Control and Prevention | National Institute for Viral Disease Control and Prevention, China CDC | Lu Tang et al |
| **hCoV-19/Belarus/BR-RRPCEM-VIS_10057/2021** | EPI_ISL_11063110 | | Laboratory for HIV and opportunistic infections diagnosis The Republican Research and Practical Center for Epidemiology and Microbiology (RRPCEM) | Laboratory for HIV and opportunistic infections diagnosis The Republican Research and Practical Center for Epidemiology and Microbiology (RRPCEM) | Elena Gasich et al |
| **hCoV-19/Liaoning/IVDC-DL-188/2021** | EPI_ISL_8896680 | | Dalian Municipal Center for Disease Control and Prevention | National Institute for Viral Disease Control and Prevention, China CDC | Xingying Lang et al |
| **hCoV-19/Heilongjiang/IVDC-HEB-534/2021** | EPI_ISL_10925442 | | Heilongjiang Center for Disease Control and Prevention | National Institute for Viral Disease Control and Prevention, China CDC | Quanbo Hu et al |
| **hCoV-19/Nepal/EPN-T171/2021** | EPI_ISL_13331533 | | Sukraraj Tropical and Infectious Disease Hospital | Center for Molecular Dynamics Nepal | Epidemic Intelligence Team et al |
| **hCoV-19/Scotland/QEUH-17B538F/2021** | EPI_ISL_2808009 | OU379351.1 | Lighthouse Lab in Glasgow | Wellcome Sanger Institute for the COVID-19 Genomics UK (COG-UK) Consortium | Harper VanSteenhouse et al |
| **hCoV-19/Belgium/rega-16254/2021** | EPI_ISL_4572026 | | KU Leuven, Rega Institute, Clinical and Epidemiological Virology | KU Leuven, Rega Institute, Clinical and Epidemiological Virology | Tony Wawina-Bokalanga et al |
| **hCoV-19/Kosovo/CO-00367-XXK000_9797/2021** | EPI_ISL_4545806 | | Department of Microbiology, National Institute of Public Health of Kosovo | Department of Microbiology, National Institute of Public Health of Kosovo | Xhevat Jakupi et al |
| **hCoV-19/Italy/SIC-AOUP-UNIPA_86103/2021** | EPI_ISL_3280870 | | Laboratorio di Riferimento Regionale della Sicilia Occidentale per l'Emergenza COVID-19 | Laboratorio di Riferimento Regionale della Sicilia Occidentale per l'Emergenza COVID-19 | Fabio Tramuto et al |
| **hCoV-19/India/AR-CSIR-NEIST-INSACOG-NEIST16-CID0142/2021** | EPI_ISL_8889296 | | RT-PCR Lab, TRIHMS, Naharlagun | Centre for Infectious Diseases, CSIR-NEIST | Romi Wahengbam et al |
| **hCoV-19/India/KL-GENESCOV07179/2021** | EPI_ISL_4749210 | | Kerala State SARS-COV-2 Genome Surveillance Programme (GENESCOV2) | GENESCOV2 at CSIR Institute of Genomics and Integrative Biology | Kerala State SARS-COV-2 Genome Surveillance Programme (GENESCOV2) et al |
| **hCoV-19/India/MZ-INSACOG-1526100531479/2021** | EPI_ISL_8700820 | | INSACOG-Mizoram | National Institute of Biomedical Genomics – INSACOG | Arindam Maitra et al A |
| **hCoV-19/India/CT-ILSGS11382/2021** | EPI_ISL_7877853 | | Pt. JNM Medical College, Raipur | Institute of Life Sciences-INSACOG | Sunil K. Raghav et al |
| **hCoV-19/Kuwait/KCCC-4250468-W037396/2021** | EPI_ISL_4085144 | | WARA LAB | KUWAIT CANCER CONTROL CENTER | SHAKIR BAHZAD et al |
| **hCoV-19/Jordan/PHBC10/2021** | EPI_ISL_2932618 | | Jordan University of Science Technology/ Irbid filed Hospital/ MOH | Princess Haya Biotechnology Center/ Jordan University of Science Technology | Saied Jaradat et al |
| **hCoV-19/India/MZ-INSACOG-587704719/2021** | EPI_ISL_9017990 | | INSACOG-Mizoram | National Institute of Biomedical Genomics – INSACOG | Arindam Maitra et al A |
| **hCoV-19/India/MZ-INSACOG-1526800051416/2021** | EPI_ISL_9721279 | | INSACOG-Mizoram | National Institute of Biomedical Genomics – INSACOG | Arindam Maitra et al A |
| **hCoV-19/Seychelles/SS3903/2021** | EPI_ISL_6705023 | OL691194.1 | Seychelles Public Health Laboratory | KEMRI-Wellcome Trust Research Programme,Kilifi | Githinji G. et al |
| **hCoV-19/India/RJ-IGIB-135585/2021** | EPI_ISL_2880131 | | INSACOG Surveillance | INSACOG at CSIR Institute of Genomics and Integrative Biology | INSACOG et al |
| **hCoV-19/Peru/LIM-INS-12362/2021** | EPI_ISL_9053600 | | Laboratorio de Referencia Nacional de Virus Respiratorios. Centro Nacional de Salud Publica. Instituto Nacional de Salud Peru. | Laboratorio de Referencia Nacional de Virus Respiratorios. Centro Nacional de Salud Publica. Instituto Nacional de Salud Peru. | Carlos Padilla Rojas et al |
| **hCoV-19/Fujian/2021BD-VIR-SPE-0579/2021** | EPI_ISL_15756978 | | Fujian Center for Disease Control and Prevention | Fujian Center for Disease Control and Prevention | Lin Qi et al |
| **hCoV-19/Indonesia/SA-NIHRD-ES1717207091W/2021** | EPI_ISL_11864536 | | RSUD Manembo-nembo Bitung | National Institute of Health Research and Development | Hana Apsari Pawestri et al |
| **hCoV-19/Indonesia/JB-KWG-LIPI-NIHRD-0026/2021** | EPI_ISL_2631467 | | RS Bayukarta | Indonesian Institute of Sciences (LIPI)/National Agency of Research and Innovation (BRIN) and National Institute of Health Research and Development (NIHRD) | Nova D. Yanthi et al |
| **hCoV-19/France/NAQ-ALPIGENE-2107220277/2021** | EPI_ISL_11448565 | | Synlab | Laboratoire de Biologie Médicale (LBM) Alpigene | Stéphanie Ducreux et al |
| **hCoV-19/Timor-Leste/TL969/2021** | EPI_ISL_4761812 | | National Health Laboratory, Timor-Leste | Microbiological Diagnostic Unit - Public Health Laboratory (MDU-PHL) | Soares da Silva et al |
| **hCoV-19/Malaysia/UNIMAS-B12577/2021** | EPI_ISL_8782810 | | Bintulu Medical Centre (Bintulu) | Institute of Health and Community Medicine | Chan Chia Jui et al |
| **hCoV-19/Zhejiang/ZPCDCP-15/2021** | EPI_ISL_17261517 | | Zhejiang Provincial Center for Disease Control and Prevention | Zhejiang Provincial Center for Disease Control and Prevention | Yi Sun et al |
| **hCoV-19/Singapore/3176/2021** | EPI_ISL_3307980 | | National Public Health Laboratory, National Centre for Infectious Diseases | National Public Health Laboratory, National Centre for Infectious Diseases | Tze Minn Mak et al |
| **hCoV-19/Fujian/2021KT-VIR-SPE-0023/2021** | EPI_ISL_15756114 | | Fujian Center for Disease Control and Prevention | Fujian Center for Disease Control and Prevention | Lin Qi et al |
| **hCoV-19/Singapore/6979/2021** | EPI_ISL_5421878 | | National Public Health Laboratory, National Centre for Infectious Diseases | National Public Health Laboratory, National Centre for Infectious Diseases | Grace Ngan et al |
| **hCoV-19/Singapore/6943/2021** | EPI_ISL_5421303 | | National Public Health Laboratory, National Centre for Infectious Diseases | National Public Health Laboratory, National Centre for Infectious Diseases | Katherine Ching et al |
| **hCoV-19/Singapore/5631/2021** | EPI_ISL_4341980 | | National Public Health Laboratory, National Centre for Infectious Diseases | National Public Health Laboratory, National Centre for Infectious Diseases | Katherine Ching et al |
| **hCoV-19/Singapore/5947/2021** | EPI_ISL_4557887 | | National Public Health Laboratory, National Centre for Infectious Diseases | National Public Health Laboratory, National Centre for Infectious Diseases | Katherine Ching et al |
| **hCoV-19/Singapore/7198/2021** | EPI_ISL_5722288 | | National Public Health Laboratory, National Centre for Infectious Diseases | National Public Health Laboratory, National Centre for Infectious Diseases | Grace Ngan et al |
| **hCoV-19/Indonesia/JK-NIHRD-WGS12643/2021** | EPI_ISL_6846823 | | Smartco Laboratorium | National Institute of Health Research and Development | Hana Apsari Pawestri et al |
| **hCoV-19/India/UT-NCDC-5809206/2021** | EPI_ISL_11052991 | | National Centre for Disease Control (NCDC) Biotechnology Division, Delhi - INSACOG | NCDC Delhi, Biotechnology Division -INSACOG | Mahesh S Dhar et al |
| **hCoV-19/India/KA-NIMH-SEQ-2334/2021** | EPI_ISL_3233027 | | VICTORIA HOSPITAL | INSACOG-KA, NIMHANS | Chitra Pattabiraman et al |
| **hCoV-19/India/MN-INSACOG-1425500153394/2021** | EPI_ISL_9116457 | | INSACOG-MANIPUR | National Institute of Biomedical Genomics – INSACOG | Arindam Maitra et al A |
| **hCoV-19/India/AS-CSIR-NEIST-INSACOG-NEIST3-CIDIN0233/2021** | EPI_ISL_10955255 | | COVID-19 Testing Laboratory, CSIR-NEIST | Centre for Infectious Diseases, CSIR-NEIST | Romi Wahengbam et al |
| **hCoV-19/India/NL-IBSD-DMP-P11-43/2021** | EPI_ISL_8890615 | | DSO, DIMAPUR | INSACOG-IBSD Manipur | Dr. Nanaocha Sharma et al |
| **hCoV-19/India/LD-ICMR-NIV-INSACOG-GSEQ-4651/2021** | EPI_ISL_3244604 | | ICMR-National Institute of Virology - INSACOG | NIV Influenza | Potdar et al B |
| **hCoV-19/India/GJ-INSACOG-GBRC2387/2021** | EPI_ISL_5023021 | | Toprani Advanced Lab system,Vadodara | Gujarat Biotechnology Research Centre | Ramesh Pandit et al |
| **hCoV-19/India/SK-INSACOG-1122500105794/2021** | EPI_ISL_9323734 | | INSACOG-Sikkim | National Institute of Biomedical Genomics – INSACOG | Arindam Maitra et al A |
| **hCoV-19/Croatia/6721/2021** | EPI_ISL_3061307 | | Nastavni zavod za javno zdravstvo Primorsko- Goranske Županije | Hrvatski zavod za javno zdravstvo | Irena Tabain et al |
| **hCoV-19/USA/OK-CDC-ASC210154154/2021** | EPI_ISL_2875844 | MZ533421.1 | Aegis Sciences Corporation | Centers for Disease Control and Prevention Division of Viral Diseases, Pathogen Discovery | Dakota Howard et al |
| **hCoV-19/Aruba/AW-RIVM-52600/2021** | EPI_ISL_3733655 | | National Institute for Public Health and the Environment (RIVM) | National Institute for Public Health and the Environment (RIVM) | Adam Meijer et al |
| **hCoV-19/USA/AL-ADPH-ADPH_0896/2021** | EPI_ISL_5887329 | | ADPH | ADPH | Adrianna Walker et al |
| **hCoV-19/USA/ME-HETL-J6895/2021** | EPI_ISL_4084577 | | Maine Health and Environmental Testing Laboratory | Tewhey Lab, The Jackson Laboratory | Matluk et al |
| **hCoV-19/USA/OK-OKPHLCOV0002834/2021** | EPI_ISL_15568281 | | Oklahoma Department of Health | Oklahoma Department of Health | Grimaldo et al |
| **hCoV-19/TrinidadandTobago/117319/2021** | EPI_ISL_8936904 | | Trinidad Public Health Laboratory | Carrington Lab, Department of Preclinical Sciences, Faculty of Medical Sciences, The University of the West Indies, St Augustine Campus | Nikita S. D. Sahadeo et al |
| **hCoV-19/Pakistan/NIH-B15-S9/2021** | EPI_ISL_3827531 | | Department of Virology | Department of Virology | Massab Umair et al |
| **hCoV-19/India/KL-GENESCOV07241/2021** | EPI_ISL_4749273 | | Kerala State SARS-COV-2 Genome Surveillance Programme (GENESCOV2) | GENESCOV2 at CSIR Institute of Genomics and Integrative Biology | Kerala State SARS-COV-2 Genome Surveillance Programme (GENESCOV2) et al |
| **hCoV-19/Azerbaijan/un-ChVir28816/2021** | EPI_ISL_15714251 | | Xususi Tehlukeli Infeksiyalara Nezaret Merkezi | Charité Universitätsmedizin Berlin, Institut für Virologie | Victor M Corman et al |
| **hCoV-19/Madagascar/LA2M-140588/2021** | EPI_ISL_13729350 | | Laboratoire d'Analyses Médicales Malagasy (LA2M) | Virology Unit, Institut Pasteur de Madagascar | Vololoniaina Raharinosy et al |
| **hCoV-19/Senegal/260959/2021** | EPI_ISL_8526935 | | Institut Pasteur de Dakar | Institut Pasteur de Dakar | Diallo Amadou et al |
| **hCoV-19/Senegal/285627/2021** | EPI_ISL_8526886 | | Institut Pasteur de Dakar | Institut Pasteur de Dakar | Diallo Amadou et al |
| **hCoV-19/Mauritanie/395964/2021** | EPI_ISL_12316584 | | INRSP-Mauritania | INRSP-Mauritania | Sidi mohamed Ahmed et al |
| **hCoV-19/Ireland/G-NVRL-M35IRL21378/2021** | EPI_ISL_3979949 | | National Virus Reference Laboratory | National Virus Reference Laboratory | Zoe Yandle et al |
| **hCoV-19/Austria/CeMM13452/2021** | EPI_ISL_3230954 | | Austrian Agency for Health and Food Safety (AGES) | Bergthaler laboratory, CeMM Research Center for Molecular Medicine of the Austrian Academy of Sciences | Lukas Endler et al |
| **hCoV-19/Gambia/71846/2021** | EPI_ISL_8401735 | | MRCG at LSHTM Genomics lab | MRCG at LSHTM Genomics lab | Abdul Karim sesay et al |
| **hCoV-19/India/LA-NCDC-380980/2021** | EPI_ISL_4446788 | | National Centre for Disease Control (NCDC) Biotechnology Division, Delhi - INSACOG | NCDC Delhi, Biotechnology Division -INSACOG | Mahesh S Dhar et al |
| **hCoV-19/Japan/RB21-005-72/2021** | EPI_ISL_5403066 | | Microbiology laboratory, National Center for Global Health and Medicine Hospital | Pathogen Genomics Center, National Institute of Infectious Diseases | Tsuyoshi Sekizuka et al B |
| **hCoV-19/Japan/PG-174444/2021** | EPI_ISL_9223376 | | Chiba Prefectural Institute of Public Health | Division of Virology and Zoology, Chiba Prefectural Institute of Public Health | Yuki Fujinuma et al |
| **hCoV-19/Japan/PG-130081/2021** | EPI_ISL_5769938 | | Fukuoka Institute of Health and Environmental Sciences | Fukuoka Institute of Health and Environmental Sciences | Yoshiki Etoh et al |
| **hCoV-19/Panama/GMI-PASEQ576/2021** | EPI_ISL_11695721 | | Gorgas Memorial Institute of Health Studies | Gorgas Memorial Institute of Health Studies | Claudia Gonzalez et al |
| **hCoV-19/Mexico/ZAC-InDRE_FB17737_S4371/2021** | EPI_ISL_2937837 | | LESP Zacatecas | Instituto de Diagnostico y Referencia Epidemiologicos (INDRE) | Claudia Wong-Arambula et al |
| **hCoV-19/HongKong/VM21035701/2021** | EPI_ISL_3547106 | | Home Quarantine Taskforce | Hong Kong Department of Health | Alan K.L. Tsang et al |
| **hCoV-19/USA/MT-MTPHL-3883367/2021** | EPI_ISL_4417449 | | Montana Public Health Laboratory | Montana Public Health Laboratory | Joy Ritter et al |
| **hCoV-19/USA/OH-CDC-MMB09548667/2021** | EPI_ISL_3604224 | MZ875475.1 | Mako Medical | Centers for Disease Control and Prevention Division of Viral Diseases, Pathogen Discovery | Dakota Howard et al |
| **hCoV-19/USA/MO-CDC-ASC210424594/2021** | EPI_ISL_7177349 | OL750160.1 | Aegis Sciences Corporation | Centers for Disease Control and Prevention Division of Viral Diseases, Pathogen Discovery | Dakota Howard et al |
| **hCoV-19/USA/MO-CDC-ASC210097787/2021** | EPI_ISL_4378855 | OK223161.1 | Aegis Sciences Corporation | Centers for Disease Control and Prevention Division of Viral Diseases, Pathogen Discovery | Dakota Howard et al |
| **hCoV-19/India/PB-NCDC-370959/2021** | EPI_ISL_4473803 | | National Centre for Disease Control (NCDC) Biotechnology Division, Delhi - INSACOG | NCDC Delhi, Biotechnology Division -INSACOG | Robin Marwal et al |
| **hCoV-19/India/SK-INSACOG-1122500110493/2021** | EPI_ISL_9323782 | | INSACOG-Sikkim | National Institute of Biomedical Genomics – INSACOG | Arindam Maitra et al A |
| **hCoV-19/India/HR-IGIB-60553/2021** | EPI_ISL_2880363 | | INSACOG Surveillance | INSACOG at CSIR Institute of Genomics and Integrative Biology | INSACOG et al |
| **hCoV-19/India/UK-NCDC-2609374/2021** | EPI_ISL_2461514 | | National Centre for Disease Control (NCDC) Biotechnology Division, Delhi | NCDC Delhi, Biotechnology Division INSACOG | Mahesh S Dhar et al |
| **hCoV-19/Qatar/QA.QU.18.25.F11/2021** | EPI_ISL_5473757 | | Ministry of Public Health / Hamad Medical Corporation | Biomedical Research Center (BRC), Qatar University / Qatar Genome Project (QGP) | BRC: Fatiha M. Benslimane et al |
| **hCoV-19/Qatar/QA.QU.18.28.H1/2021** | EPI_ISL_7167064 | | Ministry of Public Health / Hamad Medical Corporation | Biomedical Research Center (BRC), Qatar University / Qatar Genome Project (QGP) | BRC: Heba A. Al-Khatib et al |
| **hCoV-19/India/OR-ILSGS08201/2021** | EPI_ISL_7873902 | | Srirama Chandra Bhanja Medical College and Hospital, Cuttack | Institute of Life Sciences-INSACOG | Sunil K. Raghav et al |
| **hCoV-19/Mongolia/202107070/2021** | EPI_ISL_8482978 | OM182163.1 | Biotechnology Core Facility Branch, Centers for Disease Control and Prevention | Centers for Disease Control and Prevention Division of Viral Diseases, Pathogen Discovery | Anir Enkhbat et al |
| **hCoV-19/Bonaire/BQ-RIVM-60465/2021** | EPI_ISL_4839549 | | National Institute for Public Health and the Environment (RIVM) | National Institute for Public Health and the Environment (RIVM) | Adam Meijer et al |
| **hCoV-19/Maldives/MAV32203/2021** | EPI_ISL_3275173 | | F. Feeali | Indira Gandhi Memorial Hospital | Dr. Milza Abdul Muhsin et al |
| **hCoV-19/SouthAfrica/Tygerberg_2221/2021** | EPI_ISL_3856187 | | Division of Medical Virology, National Health Laboratory Service (NHLS), Tygerberg Hospital / Stellenbosch University | Division of Medical Virology, National Health Laboratory Service (NHLS), Tygerberg Hospital / Stellenbosch University | Susan Engelbrecht et al |
| **hCoV-19/Maldives/MAV61973/2021** | EPI_ISL_3540058 | | Indira Gandhi Memorial Hospital | Indiar Gandhi Memorial Hospital | D. Fathmath Nazla Rafeeq et al |
| **hCoV-19/Maldives/MAV62487/2021** | EPI_ISL_3540059 | | Indira Gandhi Memorial Hospital | Indiar Gandhi Memorial Hospital | D. Fathmath Nazla Rafeeq et al |
| **hCoV-19/Maldives/MAV73473/2021** | EPI_ISL_3540064 | | Indira Gandhi Memorial Hospital | Indiar Gandhi Memorial Hospital | D. Fathmath Nazla Rafeeq et al |
| **hCoV-19/Maldives/MAV80940/2021** | EPI_ISL_4104420 | | Indira Gandhi Memorial Hospital | Indira Gandhi Memorial Hospital | D. Fathmath Nazla Rafeeq et al |
| **hCoV-19/Maldives/MAV86594/2021** | EPI_ISL_5985316 | | Indira Gandhi Memorial Hospital | Indira Gandhi Memorial Hospital | D. Fathmath Nazla Rafeeq et al |
| **hCoV-19/Maldives/MAV86520/2021** | EPI_ISL_5985306 | | Indira Gandhi Memorial Hospital | Indira Gandhi Memorial Hospital | D. Fathmath Nazla Rafeeq et al |
| **hCoV-19/Qatar/QA.QU_18.19.B10/2021** | EPI_ISL_2843253 | | Ministry of Public Health / Hamad Medical Corporation | Biomedical Research Center (BRC), Qatar University / Qatar Genome Project (QGP) | BRC: Fatiha M. Benslimane et al |
| **hCoV-19/Latvia/5479/2021** | EPI_ISL_9030777 | | BIOR, Institute of Food Safety Animal health and Environment | Latvian Biomedical Research and Study Centre | Alise Jakovele et al |
| **hCoV-19/India/MN-INSACOG-1425500164004/2021** | EPI_ISL_9721181 | | INSACOG-Manipur | National Institute of Biomedical Genomics | Arindam Maitra et al A |
| **hCoV-19/Romania/IS-475441/2021** | EPI_ISL_5241256 | | Cantacuzino National Military-Medical Institute, Viral Respiratory Infections Laboratory | Cantacuzino National Military-Medical Institute, Viral Respiratory Infections Laboratory | Luiza Ustea et al |
| **hCoV-19/Romania/B-38400/2021** | EPI_ISL_7890559 | | "Victor Babes" Hospital | National Institute of Infectious Diseases-Prof. Dr. Matei Bals Molecular Diagnostics Laboratory | Corina Casangiu et al |
| **hCoV-19/Ireland/D-NVRL-M21IRL00293036/2021** | EPI_ISL_7370630 | | National Virus Reference Laboratory | National Virus Reference Laboratory | Zoe Yandle et al |
| **hCoV-19/Lebanon/NIC680/2022** | EPI_ISL_11209683 | | National Influenza Center - Rafic Hariri University Hospital | National Influenza Center Lebanon Rafik Hariri University Hospital | Mona Albuaini et al |
| **hCoV-19/India/AR-CSIR-NEIST-INSACOG-NEIST13-CID0084/2021** | EPI_ISL_5433667 | | Microbiology Department, BPGHTC | Centre for Infectious Diseases, CSIR-NEIST | Romi Wahengbam et al |
| **hCoV-19/India/AR-CSIR-NEIST-INSACOG-NEIST13-CID0086/2021** | EPI_ISL_5433669 | | Microbiology Department, BPGHTC | Centre for Infectious Diseases, CSIR-NEIST | Romi Wahengbam et al |
| **hCoV-19/India/AR-CSIR-NEIST-INSACOG-5-CID0077/2021** | EPI_ISL_5064588 | | Microbiology Department, BPGHTC | Centre for Infectious Diseases, CSIR-NEIST | Romi Wahengbam et al |
| **hCoV-19/Fujian/BD-VIR-SPE-0486/2021** | EPI_ISL_15775696 | | Fujian Center for Disease Control and Prevention | Fujian Center for Disease Control and Prevention | Lin Qi et al |
| **hCoV-19/USA/CA-CDC-FG-149301/2021** | EPI_ISL_5579564 | OK614591.1 | Fulgent Genetics | Centers for Disease Control and Prevention Division of Viral Diseases, Pathogen Discovery | Dakota Howard et al |
| **hCoV-19/Paraguay/247711/2021** | EPI_ISL_6973929 | | Hospital Nacional de Itagua | Laboratorio Central de Salud Pública | Sandra Gonzalez et al |
| **hCoV-19/Congo/RC-383/2021** | EPI_ISL_10023512 | | Molecular Lab, National Laboratory of Public Health | Pathogen Genomics Lab, National Institute for Biomedical Research (INRB) | Placide Mbala-Kingebeni et al B |
| **hCoV-19/Madagascar/LA2M-134600/2021** | EPI_ISL_13729368 | | Laboratoire d'Analyses Médicales Malagasy (LA2M) | Virology Unit, Institut Pasteur de Madagascar | Vololoniaina Raharinosy et al |
| **hCoV-19/Luxembourg/LNS1568144/2021** | EPI_ISL_7791287 | | BioneXt Lab | Laboratoire national de sante, Microbiology, Microbial Genomics Platform | Anke Wienecke-Baldacchino et al |
| **hCoV-19/Bulgaria/21BG-NC_003600_R14/2021** | EPI_ISL_2967192 | | National Center of Infectious and Parasitic Diseases | National Center of Infectious and Parasitic Diseases | Alexiev et al |
| **hCoV-19/India/MZ-INSACOG-1526100740898/2021** | EPI_ISL_9323792 | | INSACOG-Mizoram | National Institute of Biomedical Genomics – INSACOG | Arindam Maitra et al A |
| **hCoV-19/Iraq/KR-SEARCH-119884/2021** | EPI_ISL_12277868 | | University of Duhok COVID-19 Center for Research and Diagnosis | Andersen lab at Scripps Research | SEARCH Alliance with Jassim Mohammed Abdo et al |
| **hCoV-19/Aruba/AW-RIVM-77691/2021** | EPI_ISL_8586738 | | National Institute for Public Health and the Environment (RIVM) | National Institute for Public Health and the Environment (RIVM) | Adam Meijer et al |
| **hCoV-19/Palestine/QIBPS-PS-JER123611/2021** | EPI_ISL_10033926 | | Arab American University | Quadram Institute Bioscience | Amer Al-Jewabreh et al |
| **hCoV-19/Jordan/RMSR11S83/2021** | EPI_ISL_6075200 | | Jordan Royal Medical Services Mobil Biological Lab | Jordan Royal Medical Services Genomics Core Lab | Dr. Rame Hamdi khasawneh et al |
| **hCoV-19/Italy/EMR_AUSLRomagna_C091-21-24/2021** | EPI_ISL_5580546 | | U.O. Microbiologia Laboratorio Unico Centro Servizi - AUSL della Romagna | U.O. Microbiologia, Laboratorio Unico Centro Servizi - AUSL della Romagna | Giorgio Dirani et al |
| **hCoV-19/CanaryIslands/HUGCDN-02541500/2021** | EPI_ISL_11755426 | | Servicio Microbiología H.U. Dr. Negrín | Servicio Microbiología H.U. Dr. Negrín | Francisco Javier Chamizo López et al |
| **hCoV-19/Lebanon/HCL022120860701/2021** | EPI_ISL_17551924 | | Center For Infectious Diseases Research | CNR Virus des Infections Respiratoires, France SUD | Antonin Bal et al |
| **hCoV-19/India/ML-1727400000241/2021** | EPI_ISL_3019145 | | INSACOG-MEGHALAYA | National Institute of Biomedical Genomics – INSACOG | Arindam Maitra et al A |
| **hCoV-19/India/TG-CDFD-130759/2021** | EPI_ISL_6643059 | | Government Fever Hospital | CDFD-INSACOG | Murali Bashyam et al |
| **hCoV-19/Lebanon/8470/2021** | EPI_ISL_13829481 | | National Influenza Center, Rafic Hariri University Hospital | The Francis Crick Institute | Jammal N et al |
| **hCoV-19/Lebanon/LAU-CVD-518/2021** | EPI_ISL_11327206 | | Laboratory of Molecular Biology and Cancer Immunology, Faculty of Sciences, Lebanese University | Microbial Pathogenomics Lab - LAU | Sima tokajian et al |
| **hCoV-19/Spain/CL-COV09047/2021** | EPI_ISL_3645481 | | Hospital Clínico Universitario de Valladolid (HCUV) | Hospital Clínico Universitario de Valladolid (HCUV) | Antonio Orduña-Domingo et al |
| **hCoV-19/Chile/AN-CC356/2022** | EPI_ISL_10487171 | | Laboratorio Hospital del Cobre | Laboratorio de Infectologia y Virologia Molecular_Red SaludUC_Christus. | Constanza Martinez-Valdebenito et al |
| **hCoV-19/Norway/18636/2021** | EPI_ISL_4203038 | | Medical Microbiology Unit, Department for Laboratory Medicine, Drammen Hospital, Vestre Viken Health Trust, | Norwegian Institute of Public Health, Department of Virology | Kathrine Stene-Johansen et al |
| **hCoV-19/France/un-HCL021131235901/2021** | EPI_ISL_3205799 | | AX BIO OCEAN | CNR Virus des Infections Respiratoires - France SUD | Antonin Bal et al |
| **hCoV-19/Bonaire/BQ-RIVM-55857/2021** | EPI_ISL_4076998 | | National Institute for Public Health and the Environment (RIVM) | National Institute for Public Health and the Environment (RIVM) | Adam Meijer et al |
| **hCoV-19/Switzerland/GE-DIA-011421086301/2021** | EPI_ISL_3160487 | | Dianalabs SA | Genesupport | Tanguy ARAUD et al |
| **hCoV-19/India/AN-ICMR-NIV-INSACOG-GSEQ-7565/2021** | EPI_ISL_9503679 | | ICMR-National Institute of Virology - INSACOG | NIV Influenza | Dr. Varsha Potdar et al |
| **hCoV-19/India/WB-INSACOG-1931300481780/2021** | EPI_ISL_6886468 | | INSACOG-WB | National Institute of Biomedical Genomics – INSACOG | Arindam Maitra et al A |
| **hCoV-19/Somalia/0080/2021** | EPI_ISL_14236398 | | Puntland-Public Health Lab | Puntland-Public Health Lab | Mahamoud Hussein et al |
| **hCoV-19/Djibouti/NAMRU3_E295/2021** | EPI_ISL_6693770 | OL640806.1 | Naval Medical Research Unit No. 3 | Naval Medical Research Center Biological Defense Research Directorate | Catherine E. Arnold et al |
| **hCoV-19/Djibouti/NAMRU3_E303/2021** | EPI_ISL_6693775 | OL640811.1 | Naval Medical Research Unit No. 3 | Naval Medical Research Center Biological Defense Research Directorate | Catherine E. Arnold et al |
| **hCoV-19/Djibouti/NAMRU3_IT002/2021** | EPI_ISL_6693792 | OL640828.1 | Naval Medical Research Unit No. 3 | Naval Medical Research Center Biological Defense Research Directorate | Catherine E. Arnold et al |
| **hCoV-19/India/WB-INSACOG-1930400717267/2021** | EPI_ISL_6885588 | | INSACOG-WB | National Institute of Biomedical Genomics – INSACOG | Arindam Maitra et al A |
| **hCoV-19/India/JK-NCDC-6009179/2021** | EPI_ISL_11054364 | | National Centre for Disease Control (NCDC) Biotechnology Division, Delhi - INSACOG | NCDC Delhi, Biotechnology Division -INSACOG | Kalaiarasan Ponnusamy et al |
| **hCoV-19/India/RJ-SMS-ICMR-INSACOG-S-720/2021** | EPI_ISL_12587751 | | SMS MEDICAL COLLEGE,JAIPUR | NIV Influenza | Bharti Malhotra et al |
| **hCoV-19/India/MH-NEERI-CIB5110/2021** | EPI_ISL_7468456 | | CSIR-National Environmental Engineering Research Institute | CSIR-Centre for Cellular and Molecular Biology-INSACOG | Lamuk Zaveri et al |
| **hCoV-19/India/WB-INSACOG-1931502338881/2021** | EPI_ISL_9018102 | | INSACOG-WB | National Institute of Biomedical Genomics – INSACOG | Arindam Maitra et al A |
| **hCoV-19/India/TN-CBR-SEQ-10644/2021** | EPI_ISL_9366540 | | "CBR,IISc" | CBR,IISc | "Bratati Kahali" et al |
| **hCoV-19/USA/MO-MSPHL-002391/2021** | EPI_ISL_2635131 | | Missouri State Public Health Laboratory | Missouri State Public Health Laboratory | Matthew Sinn et al |
| **hCoV-19/Bonaire/BQ-RIVM-66249/2021** | EPI_ISL_5968086 | | National Institute for Public Health and the Environment (RIVM) | National Institute for Public Health and the Environment (RIVM) | Adam Meijer et al |
| **hCoV-19/Bonaire/BQ-RIVM-62985/2021** | EPI_ISL_5210145 | | National Institute for Public Health and the Environment (RIVM) | National Institute for Public Health and the Environment (RIVM) | Adam Meijer et al |
| **hCoV-19/SouthKorea/KDCA13652/2021** | EPI_ISL_5634340 | | Division of Emerging Infectious Diseases, Bureau of Infectious Diseases Diagnosis Control, Korea Disease Control and Prevention Agency | Division of Emerging Infectious Diseases, Bureau of Infectious Diseases Diagnosis Control, Korea Disease Control and Prevention Agency | Ae Kyung Park et al |
| **hCoV-19/Andorra/AND-242_212721322001_COV-GC/2021** | EPI_ISL_5684469 | | Laboratori d'anàlisis clíniques, Hospital Nostra Senyora de Meritxell | LBM de CHU de Toulouse, Hôpitaux de Toulouse | Rendon M et al |
| **hCoV-19/USA/NV-CDC-LC0336403/2021** | EPI_ISL_5871604 | OL324030.1 | Laboratory Corporation of America | Centers for Disease Control and Prevention Division of Viral Diseases, Pathogen Discovery | Dakota Howard et al |
| **hCoV-19/Nepal/K2050/2021** | EPI_ISL_9907706 | | Koshi Hospital | Center for Molecular Dynamics Nepal | Epidemic Intelligence Team et al |
| **hCoV-19/Jordan/Biolab0379/2021** | EPI_ISL_7979253 | | Biolab Diagnostic Laboratories | Biolab Diagnostic Laboratories | Issa Abu-Dayyeh et al |
| **hCoV-19/Lesotho/N22937/2021** | EPI_ISL_13037946 | | National Reference Laboratories | National Institute for Communicable Diseases of the National Health Laboratory Service | Gorova V et al |
| **hCoV-19/Chad/TCH-28/2021** | EPI_ISL_9274865 | | Ministère de Santé Publique et de la Solidarité Nationale | Pathogen Genomics Lab, National Institute for Biomedical Research (INRB) | Placide Mbala-Kingebeni et al B |

| | | | | | |
|---|---|---|---|---|---|
| **hCoV-19/Algeria/36421/2021** | EPI_ISL_3161806 | | NIC, Viral Respiratory Unit | Virology Departement | Alia Grad et al |
| **hCoV-19/India/ML-CCMB-CIA3245/2021** | EPI_ISL_2660889 | | PASTEUR MOLECULAR LABORATORY - INSACOG MEGHALAYA | CSIR-Centre for Cellular and Molecular Biology-INSACOG | R. Lyngdoh et al |
| **hCoV-19/Jordan/Biolab0209/2021** | EPI_ISL_5529842 | | Biolab Diagnostic Laboratories | Biolab Diagnostic Laboratories | Issa Abu-Dayyeh et al |
| **hCoV-19/Uzbekistan/Tashkent-CGB-44/2021** | EPI_ISL_3673673 | | Private clinic of Biogen Med, Tashkent, Uzbekistan | Center of Genomics and bioinformatics, Bioinformatics laboratory | Mirzakamol S Ayubov et al |
| **hCoV-19/Jordan/Biolab0392/2021** | EPI_ISL_7979257 | | Biolab Diagnostic Laboratories | Biolab Diagnostic Laboratories | Issa Abu-Dayyeh et al |
| **hCoV-19/UnitedArabEmirates/BTC-9975/2021** | EPI_ISL_16814565 | | Khalifa University Center for Biotechnology | Khalifa University Center for Biotechnology | AlSafar et al. |
| **hCoV-19/InnerMongolia/IVDC-10-02/2021** | EPI_ISL_8295282 | | Inner Mongolia Comprehensive Center for Disease Control and Prevention | National Institute for Viral Disease Control and Prevention, China CDC | Xiaoling Tian et al |
| **hCoV-19/HongKong/VM21033536/2021** | EPI_ISL_3019496 | | HOME QUARANTINE TASKFORCE | Hong Kong Department of Health | Alan K.L. Tsang et al |
| **hCoV-19/Iran/AH08481NGS/2021** | EPI_ISL_15756475 | | Hamedan University of Medical Sciences | COVID-19 National Reference Laboratory, Pasteur Institute of Iran | Zahra Fereydouni et al |
| **hCoV-19/Portugal/PT9090/2021** | EPI_ISL_2694950 | | H Fernando Fonseca | Instituto Nacional de Saude (INSA) | Borges et al |
| **hCoV-19/Australia/NSW-ICPMR-9959/2021** | EPI_ISL_5306343 | | Histopath | NSW Health Pathology - Institute of Clinical Pathology and Medical Research; Westmead Hospital; University of Sydney | Arnott A. et al |
| **hCoV-19/Australia/TAS000294/2021** | EPI_ISL_3247060 | | Royal Hobart Hospital | Royal Hobart Hospital | Dr L. Cooley et al |
| **hCoV-19/Australia/ACT1729/2021** | EPI_ISL_7569702 | | Capital Pathology | Schwessinger Lab | Austin Bird et al |
| **hCoV-19/Australia/NSW-ICPMR-11508/2021** | EPI_ISL_5533616 | | Southern. IML Pathology | NSW Health Pathology - Institute of Clinical Pathology and Medical Research; Westmead Hospital; University of Sydney | Arnott A. et al |
| **hCoV-19/NewZealand/21CV3349/2021** | EPI_ISL_7377195 | | MedLab South Ltd Nelson | Institute of Environmental Science and Research (ESR) | Sarah Cockerton et al |
| **hCoV-19/NewZealand/22CV0056/2021** | EPI_ISL_8648725 | | Hawkes Bay Hospital | Institute of Environmental Science and Research (ESR) | Sarah Cockerton et al |
| **hCoV-19/Australia/ACT0423/2021** | EPI_ISL_4108126 | | ACT Pathology | Schwessinger Lab | Carolina Correa Ospina et al |
| **hCoV-19/Australia/NT392/2021** | EPI_ISL_12275598 | | Royal Darwin Hospital Pathology | Microbiological Diagnostic Unit - Public Health Laboratory (MDU-PHL) | Meumann et al |
| **hCoV-19/India/SK-INSACOG-1122500113875/2021** | EPI_ISL_8661030 | | INSACOG-Sikkim | National Institute of Biomedical Genomics – INSACOG | Arindam Maitra et al A |
| **hCoV-19/India/AS-INSACOG-1861803648163/2021** | EPI_ISL_9720972 | | INSACOG-Assam | National Institute of Biomedical Genomics – INSACOG | Arindam Maitra et al A |
| **hCoV-19/India/OR-ILSGS12973/2021** | EPI_ISL_10398741 | | FMFMCH, Balasore | Institute of Life Sciences-INSACOG | Sunil K. Raghav et al |
| **hCoV-19/India/MN-INSACOG-1425500154511/2021** | EPI_ISL_9323583 | | INSACOG-MANIPUR | National Institute of Biomedical Genomics – INSACOG | Arindam Maitra et al A |
| **hCoV-19/Guangdong/GDIPH-GDPCC-06/2022** | EPI_ISL_18934697 | | Guangdong Provincial Institute of Public Health, Guangdong Provincial Institute of Public Health | Guangdong Provincial Institute of Public Health, Guangdong Provincial Institute of Public Health | Zhou et al |
| **hCoV-19/Laos/LOMWRU-1447/2021** | EPI_ISL_18001431 | | NCLE | LOMWRU/Microbiology Laboratory, Mahosot Hospital | Malavanh Vongsouvath et al |
| **hCoV-19/Vietnam/QT2021_182/2021** | EPI_ISL_9691653 | | CDC Quảng Trị, Vietnam | Department of Microbiology and Immunology, Pasteur Institute in Nha Trang | Trinh Hoang Long et al |
| **hCoV-19/India/TR-360548177/2021** | EPI_ISL_3982885 | | INSACOG-Tripura | National Institute of Biomedical Genomics – INSACOG | Arindam Maitra et al A |
| **hCoV-19/India/AP-CCMB-CIA4533/2021** | EPI_ISL_2775325 | | AP SSO | CSIR-Centre for Cellular and Molecular Biology-INSACOG | Priya Nurkurthy et al |
| **hCoV-19/India/MP-NCDC-590931/2021** | EPI_ISL_11053239 | | National Centre for Disease Control (NCDC) Biotechnology Division, Delhi - INSACOG | NCDC Delhi, Biotechnology Division -INSACOG | Kalaiarasan Ponnusamy et al |
| **hCoV-19/Jordan/RMSR12S7/2021** | EPI_ISL_7547533 | | Jordan Royal Medical Services Mobile biological lab | Jordan Royal Medical Services Genomics Core Lab | Dr. Rame Hamdi khasawneh et al |
| **hCoV-19/Chad/20220316_49744/2021** | EPI_ISL_18731712 | | Department of Laboratories, Ministry of Public Health, Chad | Department of Laboratories, Ministry of Public Health, Chad | Mathiew Hota et al |
| **hCoV-19/Pakistan/DUHS30/2021** | EPI_ISL_8329847 | | Provincial Public Health Reference Laboratory Sindh, Dow University of Health Sciences. | Genomic Surveillance Lab, Provincial Public Health Reference Laboratory Sindh, Dow University of Health Sciences | Saaed Khan et al |
| **hCoV-19/Spain/CT-HUGTiPM071XR5E12/2022** | EPI_ISL_9012529 | | Microbiology Department, Laboratori Clínic Metropolitana Nord. Hospital Universitari Germans Trias i Pujol | Can Ruti SARS-CoV-2 Sequencing Hub (HUGTiP/IrsiCaixa/IGTP) | Marc Noguera-Julian et al |
| **hCoV-19/Libya/86574a-37/2021** | EPI_ISL_14014997 | | Public Health Reference Laboratory NCDC | Public Health Reference Laboratory NCDC | Abdulla Bashein et al |
| **hCoV-19/Lithuania/LTU000_KNS_ANT_07_26_158/2021** | EPI_ISL_3340962 | | UAB Diagnostikos laboratorija | National Public Health Surveillance Laboratory | Lukas Zemaitis et al |
| **hCoV-19/Lithuania/LTU000_REZUS_12_30_387/2021** | EPI_ISL_9611488 | | UAB "Rezus.lt" | National Public Health Surveillance Laboratory | Lukas Zemaitis et al |
| **hCoV-19/Belgium/AZDelta-2139-16659/2021** | EPI_ISL_5315086 | | AZDelta | AZ Delta Medical Laboratories in Roeselare, Belgium | Geert Martens et al |
| **hCoV-19/Ghana/TRA-2118/2021** | EPI_ISL_6944012 | | National Influenza Centre | National Influenza Centre | William K. Ampofo et al |
| **hCoV-19/England/BRBR-2BE5E7F/2021** | EPI_ISL_7188677 | OV299370.1 | Berkshire and Surrey Pathology Services Lighthouse Laboratory | Wellcome Sanger Institute for the COVID-19 Genomics UK (COG-UK) Consortium | Berkshire et al |
| **hCoV-19/Cameroon/C132/2021** | EPI_ISL_13296379 | | Laboratoire Nationale de Santé Publique | Institut Pasteur de Dakar | Essomba René Ghislain et al |
| **hCoV-19/Romania/Calarasi_464873/2021** | EPI_ISL_3924090 | | Cantacuzino National Military-Medical Institute, Viral Respiratory Infections Laboratory | Cantacuzino Institute Virology | Carmen Chiriclu et al |
| **hCoV-19/Wales/PHWC-PYP77W/2021** | EPI_ISL_3004614 | | Originating lab: Wales Specialist Virology Centre Sequencing lab: Pathogen Genomics Unit | Public Health Wales Microbiology Cardiff Wales Specialist Virology Centre | Catherine Moore et al |
| **hCoV-19/Norway/17512/2021** | EPI_ISL_4080376 | | Vestfold Hospital, Toensberg Department of Microbiology | Norwegian Institute of Public Health, Department of Virology | Kathrine Stene-Johansen et al |
| **hCoV-19/Kosovo/CO-00533-XXK000_5046/2021** | EPI_ISL_6965639 | | Department of Microbiology, National Institute of Public Health of Kosovo | Department of Microbiology, National Institute of Public Health of Kosovo | Xhevat Jakupi et al |
| **hCoV-19/Kosovo/CO-00397-XXK000_7417/2021** | EPI_ISL_4550717 | | Department of Microbiology, National Institute of Public Health of Kosovo | Department of Microbiology, National Institute of Public Health of Kosovo | Xhevat Jakupi et al |
| **hCoV-19/Ireland/KY-NVRL-M37IRL14160/2021** | EPI_ISL_5302797 | | National Virus Reference Laboratory | National Virus Reference Laboratory | Zoe Yandle et al |
| **hCoV-19/Israel/CVL-6000401/2021** | EPI_ISL_7134120 | | Israel Central Virology laboratory | Israel Central Virology laboratory | Neta Zuckerman et al |
| **hCoV-19/Moldova/E0042/2021** | EPI_ISL_10637863 | | MedExpert | MedExpert | Vadim Nirca et al |
| **hCoV-19/Libya/59772-19/2022** | EPI_ISL_14014989 | | Janzor Surveillance | Public Health Reference Laboratory NCDC | Abdulla Bashein et al |
| **hCoV-19/Poland/4588WSN2021_wsseol/2021** | EPI_ISL_8476533 | | WSSE w Olsztynie | Wojewodzka Stacja Sanitarno-Epidemiologiczna w Olsztynie, Laboratorium Badan Epidemiologiczno-Klinicznych | Sylwia Krzelowska et al |
| **hCoV-19/India/GA-ICMR-NIV-NCCS-INSACOG-GSEQ-6064/2021** | EPI_ISL_5535983 | | ICMR-National Institute of Virology - INSACOG | NIV Influenza | Potdar et al B |
| **hCoV-19/Maldives/MAV96128/2021** | EPI_ISL_8886556 | | Indira Gandhi Memorial Hospital | Indira Gandhi Memorial Hospital | D. Fathmath Nazla Rafeeq et al |
| **hCoV-19/India/AN-ICMR-NIV-INSACOG-GSEQ-4944/2021** | EPI_ISL_3532974 | | ICMR-National Institute of Virology - INSACOG | NIV Influenza | Potdar et al B |
| **hCoV-19/India/AS-DIB-312667/2021** | EPI_ISL_14917916 | | ICMR - Regional Medical Research Centre | ICMR - Regional Medical Research Centre | B Borkakoty et al |
| **hCoV-19/SriLanka/DRVL0028/2021** | EPI_ISL_18118741 | | Diagnostic and Research Virology Laboratory | Diagnostic and Research Virology Laboratory | Bushran N Iqbal et al |
| **hCoV-19/India/AP-CCMB-CIA7040/2021** | EPI_ISL_3689798 | | AP SSO | CSIR-Centre for Cellular and Molecular Biology - INSACOG | Jandhyala Sai Krishna et al |
| **hCoV-19/USA/MD-MDH-3642/2021** | EPI_ISL_3304360 | MZ726930.1 | MD PHL | Maryland Department of Health Laboratories Administration | Eric N. Keller et al |
| **hCoV-19/USA/TX-DSHS-8208/2021** | EPI_ISL_4000748 | | TXDSHS | TXDSHS | Rashmi Tuladhar et al |
| **hCoV-19/Canada/UN-146164/2021** | EPI_ISL_12388748 | | Switch Health | Public Health Agency of Canada (PHAC) National Microbiology Laboratory | Anna Majer et al |
| **hCoV-19/Guam/GU-CDC-2-4918948/2021** | EPI_ISL_4539506 | OK300476.1 | Guam Public Health Laboratory | Centers for Disease Control and Prevention Division of Viral Diseases, Pathogen Discovery | Mili Sheth et al |
| **hCoV-19/USA/DC-CDC-LC0350943/2021** | EPI_ISL_6078411 | OL391606.1 | Laboratory Corporation of America | Centers for Disease Control and Prevention Division of Viral Diseases, Pathogen Discovery | Dakota Howard et al |
| **hCoV-19/Argentina/INEI114692/2021** | EPI_ISL_9189383 | | Servicio Virosis Respiratorias-Departamento Virología-INEI | Instituto Nacional Enfermedades Infecciosas C.G.Malbran | Baumeister E. et al |
| **hCoV-19/USA/NE-COPH-S9081/2021** | EPI_ISL_7800916 | | Nebraska Medicine | UNMC COVID-19 Response Team | Catherine B. Pratt et al |
| **hCoV-19/Canada/AB-ABPHL-26751/2021** | EPI_ISL_3911230 | | Alberta Precision Labs (APL) | Public Health Agency of Canada (PHAC) National Microbiology Laboratory | Buss et al |
| **hCoV-19/USA/RI-RISHL-082706/2021** | EPI_ISL_16705771 | | Rhode Island State Health Laboratory | Rhode Island State Health Laboratory | Richard C. Huard et al |
| **hCoV-19/Canada/AB-ABPHL-32659/2021** | EPI_ISL_4561268 | | Alberta Precision Labs (APL) | Public Health Agency of Canada (PHAC) National Microbiology Laboratory | Buss et al |
| **hCoV-19/Colombia/ANT-CWOHC-VG-SEC02502F/2021** | EPI_ISL_8519359 | | SYNLAB | Universidad Nacional de Colombia - Laboratorio Genómico One Health | Andres F. Cardona-Rios et al |
| **hCoV-19/CostaRica/INC-1478-708814/2021** | EPI_ISL_5999289 | | AREA DE SALUD COLORADO | Inciensa, Instituto Costarricense de Investigación y Enseñanza en Nutrición y Salud | Francisco Duarte et al |
| **hCoV-19/Canada/SK-RRPL-275002/2021** | EPI_ISL_7985295 | | SK-Roy Romanow Provincial Laboratory | Saskatchewan - Roy Romanow Provincial Laboratory (RRPL) | Ryan McDonald et al |
| **hCoV-19/Canada/NL-PHML-216815/2021** | EPI_ISL_5596340 | | NL-Dr. Leonard A. Miller Centre for Health Services | Newfoundland and Labrador - Eastern Health | Anna Majer et al |
| **hCoV-19/Aruba/AW-RIVM-62874/2021** | EPI_ISL_4993233 | | National Institute for Public Health and the Environment (RIVM) | National Institute for Public Health and the Environment (RIVM) | Adam Meijer et al |
| **hCoV-19/Chile/LJ-PUC_MVL_1630/2021** | EPI_ISL_5144660 | | Laboratorio de Infectología y Virología Molecular | Laboratory of Molecular Virology, School of Medicine, Pontificia Universidad Católica de Chile | Catalina Pardo-Roa et al |
| **hCoV-19/USA/MA-CDCBI-CRSP_HKYHU23LZG7RFIE/2021** | EPI_ISL_3407133 | MZ709813.1 | Broad Institute Clinical Research Sequencing Platform | Infectious Disease Program, Broad Institute of Harvard and MIT | Siddle et al |
| **hCoV-19/Peru/CAJ-INS-6027/2021** | EPI_ISL_5115797 | | Laboratorio de Referencia Nacional de Virus Respiratorios. Centro Nacional de Salud Publica. Instituto Nacional de Salud Peru. | Laboratorio de Referencia Nacional de Virus Respiratorios. Centro Nacional de Salud Publica. Instituto Nacional de Salud Peru. | Carlos Padilla Rojas et al |
| **hCoV-19/USA/NE-USAFSAM-S9278/2021** | EPI_ISL_7831293 | | US Air Force School of Aerospace Medicine | US Air Force School of Aerospace Medicine | Anthony Fries et al |
| **hCoV-19/Mexico/CHP_LANGEBIO_IMSS_4258/2021** | EPI_ISL_8251417 | | Laboratorio Central de Epidemiología (LCE) | Unidad de Genomica Avanzada | Consorcio Mexicano de Vigilancia Genomica (CoViGen-Mex). Authors (in alphabetical order): Julio Elias Alvarado-Yaah et al |
| **hCoV-19/Djibouti/NAMRU3_E932/2021** | EPI_ISL_6693786 | OL640822.1 | Naval Medical Research Unit No. 3 | Naval Medical Research Center Biological Defense Research Directorate | Catherine E. Arnold et al |
| **hCoV-19/Ethiopia/ET2108059/2021** | EPI_ISL_6266217 | | Ethiopian Public Health Institute | EPHI Pathogen Genomic Sequence Lab | Abebe G Weldemariam et al |
| **hCoV-19/India/AN-ICMR-NIV-INSACOG-GSEQ-5006/2021** | EPI_ISL_3532997 | | ICMR-National Institute of Virology - INSACOG | NIV Influenza | Potdar et al B |
| **hCoV-19/Fujian/2021BD-VIR-SPE-0496/2021** | EPI_ISL_15756090 | | Fujian Center for Disease Control and Prevention | Fujian Center for Disease Control and Prevention | Lin Qi et al |
| **hCoV-19/CzechRepublic/NRL_10695/2021** | EPI_ISL_3425762 | | National Institute of Public Health | National Institute of Public Health | Helena Jiřincova et al |
| **hCoV-19/Russia/VOR-RII-MH41129S/2021** | EPI_ISL_8109376 | | Voronezh Regional Clinical Consultative and Diagnostic Center | WHO National Influenza Centre Russian Federation | Andrey Komissarov et al |
| **hCoV-19/India/GJ-INSACOG-GBRC1481/2021** | EPI_ISL_3246248 | OM884047.1 | GMERS Medical College, Valsad | Gujarat Biotechnology Research Centre | Nitin Savaliya et al |
| **hCoV-19/Sudan/ILRI_M03126/2021** | EPI_ISL_17971644 | | Epidemiology Lab, National Public Health | International Livestock Research Institute | Samuel O. Oyola et al |
| **hCoV-19/Sudan/ILRI_M03092/2021** | EPI_ISL_17971612 | | Epidemiology Lab, National Public Health | International Livestock Research Institute | Samuel O. Oyola et al |
| **hCoV-19/Venezuela/Mir2Lab3/2021** | EPI_ISL_6989445 | | IVIC | Laboratorio de Virología Molecular | Rossana C Jaspe et al |
| **hCoV-19/Luxembourg/LNS5903291/2021** | EPI_ISL_5392421 | | BioneXt Lab | Laboratoire national de sante, Microbiology, Microbial Genomics Platform | Anke Wienecke-Baldacchino et al |
| **hCoV-19/Venezuela/Anz6539/2021** | EPI_ISL_10728903 | | INHRR | Laboratorio de Virología Molecular | Rossana C Jaspe et al |
| **hCoV-19/Andorra/AND-235_212521294601_COV-GC/2021** | EPI_ISL_4949311 | | Laboratori d'anàlisis clíniques, Hospital Nostra Senyora de Meritxell | LBM de CHU de Toulouse, Hôpitaux de Toulouse | Rendon M et al |
| **hCoV-19/Aruba/AW-RIVM-66093/2021** | EPI_ISL_5967752 | | National Institute for Public Health and the Environment (RIVM) | National Institute for Public Health and the Environment (RIVM) | Adam Meijer et al |
| **hCoV-19/Curacao/CW-RIVM-76910/2021** | EPI_ISL_8586810 | | National Institute for Public Health and the Environment (RIVM) | National Institute for Public Health and the Environment (RIVM) | Adam Meijer et al |
| **hCoV-19/Netherlands/GR-RIVM-60840/2021** | EPI_ISL_4841557 | | National Institute for Public Health and the Environment (RIVM) | National Institute for Public Health and the Environment (RIVM) | Adam Meijer et al |
| **hCoV-19/Liechtenstein/FL-Risch-21B2212042/2021** | EPI_ISL_6937235 | | Dr. Risch Ostschweiz AG | Dr Risch Laboratory | Nadia Wohlwend et al |
| **hCoV-19/Reunion/PIMIT_21322/2021** | EPI_ISL_9373950 | | REUNILAB | UMR PIMIT | Dr David A Wilkinson et al |
| **hCoV-19/Romania/B-18240/2021** | EPI_ISL_3259603 | | National Institute of Infectious Diseases-Prof. Dr. Matei Bals Molecular Diagnostics Laboratory | National Institute of Infectious Diseases-Prof. Dr. Matei Bals Molecular Diagnostics Laboratory | Corina Casangiu et al |
| **hCoV-19/Armenia/IMB12-19/2021** | EPI_ISL_9018795 | | National Center of Disease Control and Prevention of the Republic of Armenia, Davidyants Laboratories, Yerevan, Armenia | Institute of Molecular Biology NAS RA, Republic of Armenia, Department of Bioengineering, BioinformaticsInstitute and Molecular Biology IBMPh RAU, Republic of Armenia | Arsen Arakelyan et al |
| **hCoV-19/Bulgaria/22BG-EU_014736_PI136/2022** | EPI_ISL_10443876 | | National Center of Infectious and Parasitic Diseases | National Center of Infectious and Parasitic Diseases | Alexiev et al |
| **hCoV-19/Curacao/CW-RIVM-76467/2021** | EPI_ISL_8015587 | | National Institute for Public Health and the Environment (RIVM) | National Institute for Public Health and the Environment (RIVM) | Adam Meijer et al |
| **hCoV-19/Zhejiang/ZPCDCP-33/2021** | EPI_ISL_17261535 | | Zhejiang Provincial Center for Disease Control and Prevention | Zhejiang Provincial Center for Disease Control and Prevention | Yi Sun et al |
| **hCoV-19/EquatorialGuinea/57930/2021** | EPI_ISL_4601590 | | Laboratorio de Investigaciones de Baney | Swiss Tropical and Public Health Institute | Salome Hosch et al |
| **hCoV-19/EquatorialGuinea/61992/2021** | EPI_ISL_4601591 | | Laboratorio de Investigaciones de Baney | Swiss Tropical and Public Health Institute | Salome Hosch et al |
| **hCoV-19/Albania/IPH-DR-0076/2021** | EPI_ISL_17637039 | | Unit of Lab Surveillance of emmergent respiratory diseases, Institute of Public Health | Unit of Lab Surveillance of emmergent respiratory diseases, Institute of Public Health | Victor M Corman et al |
| **hCoV-19/Israel/SMC-7015890/2021** | EPI_ISL_6086471 | | Shamir Medical Center (Asaf Harofe) | Shamir Medical Center (Asaf Harofe) | Nir Rainy et al |
| **hCoV-19/Suriname/SR-964/2022** | EPI_ISL_11283826 | | Central Laboratory, Bureau of Public Health (BOG) and Academic Hospital Paramaribo | Erasmus Medical Center | Bas B Oude Munnink et al |
| **hCoV-19/Suriname/SR-862/2021** | EPI_ISL_7957574 | | Central Laboratory, Bureau of Public Health (BOG) and Academic Hospital Paramaribo | Erasmus Medical Center | Bas B Oude Munnink et al |
| **hCoV-19/Monaco/IPP28141/2021** | EPI_ISL_4740991 | | Labo Analyses Med | National Reference Center for Viruses of Respiratory Infections, Institut Pasteur, Paris | Marion Barbet et al A |
| **hCoV-19/Uruguay/DLSP_p2295/2021** | EPI_ISL_12532653 | | Departamento Laboratorios de Salud P/blica (DLSP) Ministerio de Salud P/blica | Departamento Laboratorios de Salud P/blica (DLSP) Ministerio de Salud P/blica | Cortinas et al |
| **hCoV-19/Uruguay/DLSP_p2301/2021** | EPI_ISL_11883610 | | Departamento Laboratorios de Salud Pública (DLSP) Ministerio de Salud Pública | Departamento Laboratorios de Salud Pública (DLSP) Ministerio de Salud Pública | Cortinas et al |
| **hCoV-19/Denmark/DCGC-191354/2021** | EPI_ISL_5734110 | OW443712.1 | Department of Bacteria, Parasites and Fungi, Statens Serum Institut, Copenhagen, Denmark | Statens Serum Institut Bioinformatics and Microbial Genomics | Povilas Matusevicius on behalf of the Danish Covid-19 Genome Consortium et al |
| **hCoV-19/Luxembourg/LNS6059959/2021** | EPI_ISL_7378642 | | Centre Hospitalier du Nord | Laboratoire national de sante, Microbiology, Microbial Genomics Platform | Anke Wienecke-Baldacchino et al |
| **hCoV-19/CzechRepublic/NRL-18869/2021** | EPI_ISL_8481011 | | National Institute of Public Health | National Institute of Public Health | Helena Jiřincova et al |
| **hCoV-19/Laos/LOMWRU-0826/2021** | EPI_ISL_16017398 | | National Center for Laboratory and Epidemiology (NCLE) | Microbiology Laboratory, Lao-Oxford-Mahosot Hospital-Wellcome Trust Research Unit (LOMWRU), Mahosot Hospital | Manila Souksavanh et al |
| **hCoV-19/Laos/LOMWRU-0449/2021** | EPI_ISL_15022092 | OP474032.1 | LOMWRU | LOMWRU/Microbiology Laboratory, Mahosot Hospital | Malavanh Vongsouvath et al |
| **hCoV-19/Yunnan/IVDC-561/2022** | EPI_ISL_11873924 | | Department of Acute Infectious Diseases Control and Prevention, Yunnan Center for Disease Control and Prevention | National Institute for Viral Disease Control and Prevention, China CDC | Meiling Zhang et al |
| **hCoV-19/Thailand/Chonburi_SEQ19681/2021** | EPI_ISL_6594701 | | Regional Medical Sciences Center 6 Chonburi | National Institute of Health, Department of Medical Sciences, Ministry of Public Health, Thailand | Pilailuk Okada et al |
| **hCoV-19/Cambodia/888913/2021** | EPI_ISL_7711176 | | Virology Unit, Institut Pasteur du Cambodge | Virology Unit, Institut Pasteur du Cambodge | Leakhena Pum et al |
| **hCoV-19/Cambodia/866333/2021** | EPI_ISL_6227871 | | Virology Unit, Institut Pasteur du Cambodge | Virology Unit, Institut Pasteur du Cambodge | Leakhena Pum et al |
| **hCoV-19/India/DL-NCDC-470929/2021** | EPI_ISL_11050105 | | National Centre for Disease Control (NCDC) Biotechnology Division, Delhi - INSACOG | NCDC Delhi, Biotechnology Division -INSACOG | Robin Marwal et al |
| **hCoV-19/Nepal/EPN-K2137/2021** | EPI_ISL_13331230 | | Koshi Hospital | Center for Molecular Dynamics Nepal | Epidemic Intelligence Team et al |
| **hCoV-19/India/TN-SEQ_5096_S196_R1_001/2021** | EPI_ISL_6033374 | | Dr S raju, Director of Public Health and Preventive Medicine | inStem NCBS – INSACOG | Uma Ramakrishnan et al |
| **hCoV-19/India/SK-INSACOG-1122500090112/2021** | EPI_ISL_4504609 | | INSACOG-Sikkim | National Institute of Biomedical Genomics – INSACOG | Arindam Maitra et al A |
| **hCoV-19/India/LD-ICMR-NIV-INSACOG-GSEQ-5435/2021** | EPI_ISL_4950419 | | ICMR-National Institute of Virology - INSACOG | NIV Influenza | Potdar et al B |
| **hCoV-19/India/JH-ILSGS08993/2021** | EPI_ISL_7877727 | | MGMMC, Jamshedpur | Institute of Life Sciences-INSACOG | Sunil K. Raghav et al |
| **hCoV-19/India/UT-NCDC-5809204/2021** | EPI_ISL_11052989 | | National Centre for Disease Control (NCDC) Biotechnology Division, Delhi - INSACOG | NCDC Delhi, Biotechnology Division -INSACOG | Mahesh S Dhar et al |
| **hCoV-19/Sweden/SUB004197/2021** | EPI_ISL_4130926 | | Clinical Microbiology, Infection Prevention and Control | Section for Molecular Diagnostics | Björn Hallström et al |
| **hCoV-19/Bangladesh/icddrb-GSB17-023/2021** | EPI_ISL_4196681 | | Rangamati Medical College, Rangamati | Virology Laboratory, International Centre for Diarrhoeal Disease Research, Bangladesh (ICDDR,B) | Mojnu Miah et al |
| **hCoV-19/Bangladesh/ideSHi-IR-2107/2022** | EPI_ISL_16180628 | | International Centre for Diarrhoeal Disease Research, Bangladesh (icddr,b) | Institute for Developing Science and Health Initiatives (ideSHi) | Rahman S et al |
| **hCoV-19/Bangladesh/icddrb-GSB11-130/2021** | EPI_ISL_8408948 | | Kushtia Medical College, Kushtia | Virology Laboratory, International Centre for Diarrhoeal Disease Research, Bangladesh (ICDDR,B) | Mojnu Miah et al |
| **hCoV-19/Curacao/CW-RIVM-55546/2021** | EPI_ISL_4076373 | | National Institute for Public Health and the Environment (RIVM) | National Institute for Public Health and the Environment (RIVM) | Adam Meijer et al |
| **hCoV-19/Iran/50160/2021** | EPI_ISL_15755397 | | COVID-19 National Reference Laboratory, Pasteur Institute of Iran | COVID-19 National Reference Laboratory, Pasteur Institute of Iran | Mahsa Tavakoli et al |
| **hCoV-19/India/AS-INSACOG-1870500227695/2021** | EPI_ISL_9056262 | | INSACOG-Assam | National Institute of Biomedical Genomics – INSACOG | Arindam Maitra et al A |
| **hCoV-19/India/AP-CCMB-CIB2429/2021** | EPI_ISL_8270031 | | AP SSO | CSIR-Centre for Cellular and Molecular Biology-INSACOG | Lamuk Zaveri et al |
| **hCoV-19/India/UT-NCDC-610996/2021** | EPI_ISL_11064844 | | National Centre for Disease Control (NCDC) Biotechnology Division, Delhi - INSACOG | NCDC Delhi, Biotechnology Division -INSACOG | Kalaiarasan Ponnusamy et al |
| **hCoV-19/India/MN-INSACOG-1425600197752/2021** | EPI_ISL_6885482 | | INSACOG-Manipur | National Institute of Biomedical Genomics – INSACOG | Arindam Maitra et al A |
| **hCoV-19/India/HP-NCDC-5909281/2021** | EPI_ISL_11053425 | | National Centre for Disease Control (NCDC) Biotechnology Division, Delhi - INSACOG | NCDC Delhi, Biotechnology Division -INSACOG | Kalaiarasan Ponnusamy et al |
| **hCoV-19/India/HP-NCDC-6109280/2021** | EPI_ISL_11064895 | | National Centre for Disease Control (NCDC) Biotechnology Division, Delhi - INSACOG | NCDC Delhi, Biotechnology Division -INSACOG | Kalaiarasan Ponnusamy et al |
| **hCoV-19/Panama/GMI-PAM85505/2021** | EPI_ISL_11696303 | | Gorgas Memorial Institute of Health Studies | Gorgas Memorial Institute of Health Studies | Claudia Gonzalez et al |
| **hCoV-19/Benin/MTMC_35_094/2021** | EPI_ISL_17425672 | | Centre Hospitalier et Universitaire de Zone Abomey-Calavi | Molecular Malaria Laboratory, Mahidol Oxford Tropical Medicine Research Unit | Aurel Allabi et al |
| **hCoV-19/India/GJ-INSACOG-GBRC2865/2021** | EPI_ISL_6639403 | | Unipath Specialty Laboratory Limited, Vadodara | Gujarat Biotechnology Research Centre | Sonal Sharma et al |
| **hCoV-19/Bulgaria/21BG-EU_009713_PI92/2021** | EPI_ISL_7327137 | | National Center of Infectious and Parasitic Diseases | National Center of Infectious and Parasitic Diseases | Alexiev et al |
| **hCoV-19/Nepal/EPN-K2476/2021** | EPI_ISL_12956320 | | Koshi Hospital | Center for Molecular Dynamics Nepal | Epidemic Intelligence Team et al |
| **hCoV-19/India/UT-NCDC-6209337/2021** | EPI_ISL_11055265 | | National Centre for Disease Control (NCDC) Biotechnology Division, Delhi - INSACOG | NCDC Delhi, Biotechnology Division -INSACOG | Kalaiarasan Ponnusamy et al |
| **hCoV-19/Myanmar/DSMRC-054/2021** | EPI_ISL_8920487 | | DSMRC | Defence Services Medical Research Centre (DSMRC) | Khine Zaw Oo et al |
| **hCoV-19/Bahrain/21990097246_S31_L001/2021** | EPI_ISL_6266516 | OL441403.1 | Communicable Disease Laboratory, Public Health Directorate | Communicable Disease Laboratory, Public Health Directorate | Alwazii et al |
| **hCoV-19/India/JH-ILSGS12170/2021** | EPI_ISL_7878156 | | Department of Pathology, Tata Main Hospital, Jamshedpur | Institute of Life Sciences-INSACOG | Sunil K. Raghav et al |
| **hCoV-19/India/MN-IBSD-IW-PS-39/2021** | EPI_ISL_10794442 | | Babina Diagnostics | INSACOG-IBSD Manipur | Nanaocha Sharma et al |
| **hCoV-19/Taiwan/NTU-HCH-066/2021** | EPI_ISL_14189428 | | Department of Laboratory Medicine, National Taiwan University Hospital Hsin-Chu Hospital, Taiwan | Research Center for Emerging Viral Infections, Chang Gung University, Taiwan | Pei-Lan Shao et al |
| **hCoV-19/India/DH-ICMR-NIV-INSACOG-GSEQ-7602/2021** | EPI_ISL_9503816 | | ICMR-National Institute of Virology - INSACOG | NIV Influenza | Dr. Varsha Potdar et al |
| **hCoV-19/India/HP-NCDC-780945/2021** | EPI_ISL_11059151 | | National Centre for Disease Control (NCDC) Biotechnology Division, Delhi - INSACOG | NCDC Delhi, Biotechnology Division -INSACOG | Kalaiarasan Ponnusamy et al |
| **hCoV-19/India/NL-IBSD-KHM-P8-86/2021** | EPI_ISL_10123074 | | NHAK | INSACOG-IBSD Manipur | Dr. Nanaocha Sharma et al |
| **hCoV-19/Norway/3839/2020** | EPI_ISL_591012 | | Department of Medical Microbiology - section Molde, Molde Hospital | Norwegian Institute of Public Health, Department of Virology | Kathrine Stene-Johansen et al |
| **hCoV-19/Libya/421067109/2020** | EPI_ISL_17682085 | | Libyan Biotechnology Research Center | Libyan Biotechnology Research Center | Inas Alhudiri et al |
| **hCoV-19/Russia/MOW-CSP-3-2-R_S6/2020** | EPI_ISL_8907761 | | Chief Federal State Budgetary Healthcare Institution "Centre of Hygiene and Epidemiology" of the Federal Medical Biological Agency, Central laboratory of microbiological and clinical diagnostic tests | Federal State Budgetary Institution "Centre for Strategic Planning and Management of Biomedical Health Risks" of the Federal Medical Biological Agency, Laboratory for advanced molecular diagnostics of human diseases. | Mikhaylov et al |
| **hCoV-19/India/MH-NIV-65861/2020** | EPI_ISL_479569 | | NIV Influenza | NIV Influenza | Potdar et al B |
| **hCoV-19/Bangladesh/DNAS-RAB-Isl-114/2020** | EPI_ISL_605904 | MT581432.1 | NGS Lab, DNA SOLUTION LTD. | NGS Lab, DNA SOLUTION LTD. | Khan et al |
| **hCoV-19/Egypt/NRC-6290/2020** | EPI_ISL_8193879 | | Center of Scientific Excellence for Influenza Viruses (CSEIV), National Research Centre | Center of Scientific Excellence for Influenza Viruses (CSEIV), National Research Centre | Rabeh El-Shesheny et al |
| **hCoV-19/Philippines/PH-RITM-706/2020** | EPI_ISL_10943003 | | CALAMBA CITY HEALTH OFFICE | Research Institute for Tropical Medicine | Inez Andrea Medado et al |
| **hCoV-19/Philippines/PH-PGC-01803/2020** | EPI_ISL_2156215 | | Cagayan Valley Medical Center Molecular Laboratory | Philippine Genome Center | Francis A. Tablizo et al |
| **hCoV-19/Australia/WA257/2020** | EPI_ISL_594171 | | PathWest Laboratory Medicine WA | PathWest Laboratory Medicine WA Microbial Surveillance Unit | PathWest Laboratory Medicine WA Microbial Surveillance Unit et al |

| | | | | | |
|---|---|---|---|---|---|
| **hCoV-19/Senegal/1902/2020** | EPI_ISL_8525488 | | Institut Pasteur de Dakar | Institut Pasteur de Dakar | Diallo Amadou et al |
| **hCoV-19/Sichuan/SC-MY-001/2020** | EPI_ISL_661271 | | Research platform for Transfusion-transmitted Disease, Institute of Blood Transfusion, Chinese Academy of Medical Sciences | Research platform for Transfusion-transmitted Disease, Institute of Blood Transfusion, Chinese Academy of Medical Sciences | He et al |
| **hCoV-19/India/TG-CCMB-M697/2020** | EPI_ISL_495225 | | CSIR-Centre for Cellular and Molecular Biology | CSIR-Centre for Cellular and Molecular Biology | Pratheusa Maccha et al |
| **hCoV-19/Bahrain/090000572_S16_L001/2020** | EPI_ISL_632285 | MW228280.1 | Communicable Disease Laboratory, Public Health Directorate | Communicable Disease Laboratory, Public Health Directorate | AlWasti et al |
| **hCoV-19/India/KA-CS1026/2020** | EPI_ISL_811155 | | Dharwad | CSIR Institute of Genomics and Integrative Biology | Dr. Shivarudrapp B Bhairappanavar et al |
| **hCoV-19/India/MH-NEERI-NGP-29586/2021** | EPI_ISL_1360327 | | CSIR-National Environmental Engineering Research Institute | CSIR-Centre for Cellular and Molecular Biology - INSACOG | Onkar Kulkarni et al |
| **hCoV-19/India/MH-NEERI-NGP-25677/2020** | EPI_ISL_1916382 | | CSIR-National Environmental Engineering Research Institute | CSIR-Centre for Cellular and Molecular Biology - INSACOG | Lamuk Zaveri et al |
| **hCoV-19/Bahrain/023335731/2020** | EPI_ISL_14133030 | MW655535.1 | Communicable Disease Laboratory, Public Health Directorate | Communicable Disease Laboratory, Public Health Directorate | Alwasti et al |
| **hCoV-19/India/MH-ACTREC-534/2020** | EPI_ISL_700180 | | Hematopathology Laboratory, ACTREC, TMC | Hematopathology Laboratory, ACTREC, TMC | Hematopathology Laboratory et al |
| **hCoV-19/Bahrain/920457374_S4/2020** | EPI_ISL_632901 | MW242650.1 | Communicable Disease Laboratory, Public Health Directorate | Communicable Disease Laboratory, Public Health Directorate | AlAbbas et al |
| **hCoV-19/Bahrain/341013928_S4_L001/2020** | EPI_ISL_678263 | MW332215.1 | Communicable Disease Laboratory, Public Health Directorate | Communicable Disease Laboratory, Public Health Directorate | Alwasti et al |
| **hCoV-19/Canada/SK-NML-17304/2020** | EPI_ISL_2883403 | | SK-Roy Romanow Provincial Laboratory | National Microbiology Laboratory (NML) | Anna Majer et al |
| **hCoV-19/India/GJ-INSACOG-GBRC1616/2021** | EPI_ISL_3843679 | OM915402.1 | VRDL,Goverment Medical College Surat | Gujarat Biotechnology Research Centre | Janvi Raval et al |
| **hCoV-19/Senegal/9439/2020** | EPI_ISL_486861 | | Institut Pasteur Dakar | Institut Pasteur de Dakar | Ndongo Dia et al |
| **hCoV-19/Poland/Masovia-14790/2020** | EPI_ISL_754910 | | Laboratory Diagnostics and Clinical Immunology of Developmental Age, Medical University of Warsaw | genXone SA, Research & Development Laboratory; The Faculty of Mathematics, Informatics and Mechanics of the University of Warsaw | Maciej Sykulski et al |
| **hCoV-19/Japan/PG-9989/2020** | EPI_ISL_902712 | | Pathogen Genomics Center, National Institute of Infectious Diseases | Pathogen Genomics Center, National Institute of Infectious Diseases | Tsuyoshi Sekizuka et al B |
| **hCoV-19/Japan/PG-8751/2020** | EPI_ISL_686347 | | Pathogen Genomics Center, National Institute of Infectious Diseases | Pathogen Genomics Center, National Institute of Infectious Diseases | Tsuyoshi Sekizuka et al B |
| **hCoV-19/Japan/PG-8173/2020** | EPI_ISL_687272 | | Pathogen Genomics Center, National Institute of Infectious Diseases | Pathogen Genomics Center, National Institute of Infectious Diseases | Tsuyoshi Sekizuka et al B |
| **hCoV-19/Japan/PG-11540/2020** | EPI_ISL_898403 | | Pathogen Genomics Center, National Institute of Infectious Diseases | Pathogen Genomics Center, National Institute of Infectious Diseases | Tsuyoshi Sekizuka et al B |
| **hCoV-19/Japan/PG-15543/2020** | EPI_ISL_894359 | | Department of Infectious Diseases, Kobe Institute of Health | Pathogen Genomics Center, National Institute of Infectious Diseases | Tsuyoshi Sekizuka et al B |
| **hCoV-19/Japan/PG-19294/2021** | EPI_ISL_1128028 | | Chiba Prefectural Institute of Public Health | Pathogen Genomics Center, National Institute of Infectious Diseases | Tsuyoshi Sekizuka et al B |
| **hCoV-19/Japan/PG-19458/2020** | EPI_ISL_1123572 | | Gunma Prefectural Institute of Public Health and Environmental Sciences | Pathogen Genomics Center, National Institute of Infectious Diseases | Tsuyoshi Sekizuka et al B |
| **hCoV-19/Japan/PG-3700/2020** | EPI_ISL_691953 | | Pathogen Genomics Center, National Institute of Infectious Diseases | Pathogen Genomics Center, National Institute of Infectious Diseases | Tsuyoshi Sekizuka et al B |
| **hCoV-19/Botswana/BOT2254/2020** | EPI_ISL_560390 | | National Health Laboratory | Botswana Institute for Technology Research and Innovation | Kefentse Arnold Tumedi et al |
| **hCoV-19/India/UT-AR8/2020** | EPI_ISL_508205 | | All india institute of Medical Sciences Rishikesh | National Institute of Biomedical Genomics | Arindam Maitra et al A |
| **hCoV-19/Israel/13075882/2020** | EPI_ISL_447442 | | Clinical Microbiology Laboratory, Sheba Medical Center | Stern Lab | Stern Lab et al (https://dx.doi.org/10.1101/2020.05.21.20104521) |
| **hCoV-19/Mexico/JAL-UANL-002/2020** | EPI_ISL_1091261 | | Laboratorio de Apoyo a la Vigilancia e Investigación Epidemiológica - CIBO | Laboratorio de Infectología Molecular, Departamento de Bioquímica y Medicina Molecular,Facultad de Medicina - Universidad Autónoma de Nuevo León | Kame A. Galán-Huerta et al |
| **hCoV-19/Japan/PG-0969/2020** | EPI_ISL_684579 | | Pathogen Genomics Center, National Institute of Infectious Diseases | Pathogen Genomics Center, National Institute of Infectious Diseases | Tsuyoshi Sekizuka et al B |
| **hCoV-19/Bulgaria/21/2020** | EPI_ISL_480302 | | National Reference Laboratory "Influenza and acute respiratory diseases" | NRL-HIV | Ivan Ivanov et al |
| **hCoV-19/Israel/CVL_S13-1/2020** | EPI_ISL_2786923 | | Central Virology Laboratory, Ministry of Health | Central Virology Laboratory, Ministry of Health | Neta S. Zuckerman et al |
| **hCoV-19/NorthMacedonia/883/2020** | EPI_ISL_677674 | | General Hospital - Prilep | Research Center for Genetic Engineering and Biotechnology "Georgi D. Efremov" , Macedonian Academy of Sciences and Arts | RCGEB - MASA et al |
| **hCoV-19/Brazil/BA-FIOCRUZ-PVM9457/2020** | EPI_ISL_14291041 | | Plataforma de Vigilancia Molecular (PVM) - FIOCRUZ/BA | Plataforma de Vigilancia Molecular (PVM) - FIOCRUZ/BA | Laise de Moraes et al |
| **hCoV-19/India/HP-ICMR-NIV-INSACOG-SEQ-2125/2020** | EPI_ISL_15038475 | | ICMR-National Institute of Virology - INSACOG | NIV Influenza | Dr. Varsha Potdar et al |
| **hCoV-19/Turkey/DEN-PAU19/2020** | EPI_ISL_708194 | | Pamukkale University Hospital | Pamukkale University Department of Medical Genetics | Onur TOKGUN et al. |
| **hCoV-19/Israel/HMO-VL-2429879/2020** | EPI_ISL_2096775 | | Hadassah Medical Center Clinical Virology Laboratory, Hadassah Ein Kerem | Hadassah Hebrew University Viral Sequencing Group, Hadassah Hebrew University Medical Center | Hadar Golan Berman et al |
| **hCoV-19/Palestine/AAS36/2020** | EPI_ISL_1273095 | MW737843.1 | Biochemistry and Molecular Biology Department-Faculty of Medicine, Al-Quds University | Biochemistry and Molecular Biology Department-Faculty of Medicine, Al-Quds University | Ereqat et al |
| **hCoV-19/Israel/HMO-VL-2497871/2020** | EPI_ISL_2096776 | | Hadassah Medical Center Clinical Virology Laboratory, Hadassah Ein Kerem | Hadassah Hebrew University Viral Sequencing Group, Hadassah Hebrew University Medical Center | Hadar Golan Berman et al |
| **hCoV-19/Israel/CVL-n26134/2020** | EPI_ISL_514289 | | Israel Central Virology laboratory | Israel Central Virology laboratory | Neta Zuckerman et al |
| **hCoV-19/Palestine/49/2020** | EPI_ISL_596532 | | Palestinian Ministry of Health | Molecular Genetics Lab | Nouar Qutob et al |
| **hCoV-19/Palestine/AAS27/2020** | EPI_ISL_739660 | MW411948.1 | Al-Quds Nutrition and Health Research Institute, Al-Quds University | Al-Quds Nutrition and Health Research Institute, Al-Quds University | Nasereddin et al |
| **hCoV-19/Japan/PG-26066/2021** | EPI_ISL_1428301 | | Chiba Prefectural Institute of Public Health | Pathogen Genomics Center, National Institute of Infectious Diseases | Tsuyoshi Sekizuka et al B |
| **hCoV-19/Japan/PG-4623/2020** | EPI_ISL_689760 | | Fukuoka Institute of Health and Environmental Sciences | Pathogen Genomics Center, National Institute of Infectious Diseases | Tsuyoshi Sekizuka et al B |
| **hCoV-19/Japan/PG-16355/2020** | EPI_ISL_1128359 | | Pathogen Genomics Center, National Institute of Infectious Diseases | Pathogen Genomics Center, National Institute of Infectious Diseases | Tsuyoshi Sekizuka et al B |
| **hCoV-19/Japan/PG-16902/2020** | EPI_ISL_1128955 | | Pathogen Genomics Center, National Institute of Infectious Diseases | Pathogen Genomics Center, National Institute of Infectious Diseases | Tsuyoshi Sekizuka et al B |
| **hCoV-19/Fujian/TH-H210159/2021** | EPI_ISL_1663017 | | Fujian Center for Disease Control and Prevention | Fujian Center for Disease Control and Prevention | Lin Qi et al |
| **hCoV-19/Japan/PG-14667/2020** | EPI_ISL_892599 | | Saitama Prefectural Institute of Public Health | Pathogen Genomics Center, National Institute of Infectious Diseases | Tsuyoshi Sekizuka et al B |
| **hCoV-19/Japan/PG-7273/2020** | EPI_ISL_688376 | | Saitama Prefectural Institute of Public Health | Pathogen Genomics Center, National Institute of Infectious Diseases | Tsuyoshi Sekizuka et al B |
| **hCoV-19/Cambodia/101372/2020** | EPI_ISL_1040028 | | Original detection - Virology Unit, Institut Pasteur du Cambodge; Sequencing - US National Institute of Allergy and Infectious Diseases Cambodia | Virology Unit, Institut Pasteur du Cambodge | Jennifer Bohl et al |
| **hCoV-19/Mongolia/UB-53319/2020** | EPI_ISL_9879602 | | National Center for Communicable Diseases (NCCD) National Influenza Center | National Center for Communicable Diseases (NCCD) National Influenza Center | Naranzul Tsedenbal et al |
| **hCoV-19/Japan/PG-26509/2021** | EPI_ISL_1430917 | | Pathogen Genomics Center, National Institute of Infectious Diseases | Pathogen Genomics Center, National Institute of Infectious Diseases | Tsuyoshi Sekizuka et al B |
| **hCoV-19/Japan/PG-11631/2020** | EPI_ISL_898494 | | Pathogen Genomics Center, National Institute of Infectious Diseases | Pathogen Genomics Center, National Institute of Infectious Diseases | Tsuyoshi Sekizuka et al B |
| **hCoV-19/Guangxi/NAN.20-8/2020** | EPI_ISL_6951092 | | Molecular Diagnostics Lab, Affiliated Infectious Disease Hospital of Nanning, Guangxi Medical University | Key Laboratory of Infectious Diseases,Nanning Health Commission. | JianYan Lin et al |
| **hCoV-19/Togo/TGS4/2020** | EPI_ISL_1508958 | | Institut National d'hygiène | Unité Mixte Internationale TransVIHMI (UMI 233 IRD – U1175 INSERM - Université de Montpellier) IRD (Institut de recherche pour le développement) | Mounerou SALOU et al |
| **hCoV-19/Mongolia/UB-12770/2020** | EPI_ISL_9879585 | | National Center for Communicable Diseases (NCCD) National Influenza Center | National Center for Communicable Diseases (NCCD) National Influenza Center | Naranzul Tsedenbal et al |
| **hCoV-19/Mexico/PUE-InDRE-IBT-28747-NC/2020** | EPI_ISL_3463545 | | Instituto de Diagnostico y Referencia Epidemiologicos InDRE_RNLSP | Instituto de Biotecnología de la UNAM | Authors from IBT et al |
| **hCoV-19/Mexico/OAX-InDRE_266/2020** | EPI_ISL_1054966 | | Instituto de Diagnostico y Referencia Epidemiologicos INDRE_RNLSP | Instituto de Diagnostico y Referencia Epidemiologicos (INDRE) | Claudia Wong-Arambula et al |
| **hCoV-19/Georgia/Tb-s083/2020** | EPI_ISL_13300281 | | National Center for Disease Control and Public Health (NCDC) of Georgia. | Department for Virology, Molecular Biology and Genome Research, R. G. Lugar Center for Public Health Research, National Center for Disease Control and Public Health (NCDC) of Georgia. | Giorgi Tomashvili et al |
| **hCoV-19/Georgia/Tb-RNGS013/2020** | EPI_ISL_1921879 | | Department for Virology, Molecular Biology and Genome Research, R. G. Lugar Center for Public Health Research, National Center for Disease Control and Public Health (NCDC) of Georgia. | Department for Virology, Molecular Biology and Genome Research, R. G. Lugar Center for Public Health Research, National Center for Disease Control and Public Health (NCDC) of Georgia. | Giorgi Tomashvili et al |
| **hCoV-19/Georgia/Tb-RNGS001/2020** | EPI_ISL_1921858 | | Department for Virology, Molecular Biology and Genome Research, R. G. Lugar Center for Public Health Research, National Center for Disease Control and Public Health (NCDC) of Georgia. | Department for Virology, Molecular Biology and Genome Research, R. G. Lugar Center for Public Health Research, National Center for Disease Control and Public Health (NCDC) of Georgia. | Giorgi Tomashvili et al |
| **hCoV-19/Georgia/Tb-R030/2020** | EPI_ISL_3076898 | | Department for Virology, Molecular Biology and Genome Research, R. G. Lugar Center for Public Health Research, National Center for Disease Control and Public Health (NCDC) of Georgia. | Department for Virology, Molecular Biology and Genome Research, R. G. Lugar Center for Public Health Research, National Center for Disease Control and Public Health (NCDC) of Georgia. | Giorgi Tomashvili et al |
| **hCoV-19/SouthAfrica/R13452-20/2020** | EPI_ISL_622940 | | National Institute for Communicable Diseases of the National Health Laboratory Service | National Institute for Communicable Diseases of the National Health Laboratory Service | Allam M et al |
| **hCoV-19/Botswana/R8B5_BHP_58650/2020** | EPI_ISL_2233456 | | Botswana Harvard HIV Reference Laboratory | Botswana Harvard HIV Reference Laboratory | Sikhulile Wonderful T. Choga et al |
| **hCoV-19/Botswana/R8B4_BHP_AAB8004/2020** | EPI_ISL_1516828 | | Botswana Harvard HIV Reference Laboratory | Botswana Harvard HIV Reference Laboratory | Sikhulile Wonderful T. Choga et al |
| **hCoV-19/Botswana/R8B40_BHP_AAB8009/2020** | EPI_ISL_1516856 | | Botswana Harvard HIV Reference Laboratory | Botswana Harvard HIV Reference Laboratory | Sikhulile Dorcas Maruapula et al |
| **hCoV-19/Peru/LIM-INS-470/2020** | EPI_ISL_1111368 | | Laboratorio de Referencia Nacional de Virus Respiratorio. Instituto Nacional de Salud Perú | Laboratorio de Referencia Nacional de Enteropatógenos. Instituto Nacional de Salud del Perú | Ronnie Gavilan Chavez et al |
| **hCoV-19/Peru/LIM-INS-362/2020** | EPI_ISL_1111260 | | Laboratorio de Referencia Nacional de Virus Respiratorio. Instituto Nacional de Salud Perú | Laboratorio de Referencia Nacional de Enteropatógenos. Instituto Nacional de Salud del Perú | Ronnie Gavilan Chavez et al |
| **hCoV-19/Peru/AMA-UPCH-0266/2020** | EPI_ISL_729912 | | Instituto de Medicina Tropical, Universidad Nacional Toribio Rodríguez de Mendoza de Amazonas | Laboratorio de Genómica Microbiana, Universidad Peruana Cayetano Heredia | Pablo Tsukayama et al |
| **hCoV-19/Peru/AMA-UPCH-0260/2020** | EPI_ISL_729906 | | Instituto de Medicina Tropical, Universidad Nacional Toribio Rodríguez de Mendoza de Amazonas | Laboratorio de Genómica Microbiana, Universidad Peruana Cayetano Heredia | Pablo Tsukayama et al |
| **hCoV-19/Egypt/PHARCO-ARMY-27/2021** | EPI_ISL_1936346 | MW829328.1 | Main Chemical Laboratories Egypt Army | Main Chemical Laboratories Egypt Army | Sherine Helmy et al |
| **hCoV-19/Italy/SIC-186413300/2021** | EPI_ISL_2731295 | | Molecular Hepatology Laboratory AOU Messina | Molecular Hepatology Laboratory AOU Messina | Giuseppina Raffa et al |
| **hCoV-19/Chile/RM-157643/2020** | EPI_ISL_746678 | | Genetica Molecular and Subdepartamento de Virologia ISP Chile | Instituto de Salud Publica de Chile | Javier Tognarelli et al |
| **hCoV-19/Chile/Santiago-PUC_MVL_0347/2020** | EPI_ISL_801780 | MW365048.1 | Laboratory of Molecular Virology, Pontificia Universidad Católica de Chile | MSHS Pathogen Surveillance Program | Leonardo I. Almonacid et al |
| **hCoV-19/Angola/KRISP-K009709/2020** | EPI_ISL_1347898 | | Instituto Nacional de Investigação em Saúde | KRISP, KZN Research Innovation and Sequencing Platform | Morais J et al |
| **hCoV-19/Peru/un-CDC-4033267-001/2020** | EPI_ISL_2382733 | MZ323732.1 | NAMRU-6 | Genomics and Discovery, Respiratory Viruses Branch, Division of Viral Diseases, Centers for Disease Control and Prevention | Anna Kelleher et al |
| **hCoV-19/Peru/un-CDC-3959175-001/2020** | EPI_ISL_2379243 | MZ323731.1 | NAMRU-6 | Genomics and Discovery, Respiratory Viruses Branch, Division of Viral Diseases, Centers for Disease Control and Prevention | Anna Kelleher et al |
| **hCoV-19/Peru/CAL-INS-743/2020** | EPI_ISL_1563627 | | Laboratorio de Referencia Nacional de Biotecnología y Biología Molecular. Instituto Nacional de Salud Peru | Laboratorio de Referencia Nacional de Biotecnología y Biología Molecular. Instituto Nacional de Salud Peru | Carlos Padilla Rojas et al |
| **hCoV-19/Ecuador/157876_S14/2020** | EPI_ISL_18283639 | | INSPI | INSPI - Charité | Alfredo Bruno Caicedo et al |
| **hCoV-19/Mozambique/INS-K011806/2020** | EPI_ISL_2396911 | | Instituto Nacional de Saude (INS), Mozambique | KRISP, KZN Research Innovation and Sequencing Platform | Nalia Ismael et al |
| **hCoV-19/Aruba/AW-RIVM-11400/2021** | EPI_ISL_1014657 | | National Institute for Public Health and the Environment (RIVM) | National Institute for Public Health and the Environment (RIVM) | Adam Meijer et al |
| **hCoV-19/Ireland/WH-NVRL-83IRL31298/2021** | EPI_ISL_875446 | | National Virus Reference Laboratory | National Virus Reference Laboratory | Michael Carr et al |
| **hCoV-19/SouthAfrica/KRISP-K002311/2020** | EPI_ISL_515687 | | NHLS-IALCH | KRISP, KZN Research Innovation and Sequencing Platform | Giandhari J et al |
| **hCoV-19/Serbia/NoviSad-158472905/2020** | EPI_ISL_676590 | | Scientific Veterinary Institute Novi Sad | Veterinary Specialized Institute "Kraljevo", Serbia | Vidanovic et al |
| **hCoV-19/Serbia/Valjevo-596B15-1909/2020** | EPI_ISL_6959656 | | Directorate for national reference laboratories | Center for forensic and applied molecular genetics, Faculty of Biology, University of Belgrade | D.Keckarević et al |
| **hCoV-19/Serbia/Beograd-RNK_c26049-2010/2020** | EPI_ISL_6930644 | | Department of Biocides and Medical Ecology | Center for forensic and applied molecular genetics, Faculty of Biology, University of Belgrade | D.Keckarević et al |
| **hCoV-19/Serbia/Kraljevo-02092507/2020** | EPI_ISL_582533 | | Veterinary Specialized Institute "Kraljevo", Serbia | Veterinary Specialized Institute "Kraljevo", Serbia | Vidanovic et al |
| **hCoV-19/Egypt/ARMY-213/2021** | EPI_ISL_1936104 | | Main Chemical Laboratories Egypt Army | Main Chemical Laboratories Egypt Army | Mohamed Seadawy et al |
| **hCoV-19/Argentina/INEI104530/2021** | EPI_ISL_2158728 | | Servicio Virosis Respiratorias-Departamento Virología-INEI | Instituto Nacional Enfermedades Infecciosas C.G.Malbran | Baumeister E. et al |
| **hCoV-19/Argentina/INEI113365/2021** | EPI_ISL_7753854 | | Servicio Virosis Respiratorias-Departamento Virología-INEI | Instituto Nacional Enfermedades Infecciosas C.G.Malbran | Baumeister E. et al |
| **hCoV-19/Argentina/PAIS-A0583/2021** | EPI_ISL_18290150 | | Laboratorio de Virología del Hospital de Niños Dr. Ricardo Gutierrez | Área de Secuenciación del Laboratorio de Virología del Hospital de Niños Dr. Ricardo Gutierrez on behalf of 'Proyecto Argentino Interinstitucional de genomica de SARS-CoV-2' (PAIS Consortium) | Nabaes Jodar et al |
| **hCoV-19/SaintKittsandNevis/72432/2021** | EPI_ISL_2756582 | | The Caribbean Public Health Agency | Carrington Lab, Department of Preclinical Sciences, Faculty of Medical Sciences, The University of the West Indies, St Augustine Campus | Nikita S. D. Sahadeo et al |
| **hCoV-19/Argentina/PAIS-C0254/2021** | EPI_ISL_7956482 | | Laboratorio de Virología Molecular - Hospital Blas L. Dubarry | Insituto Nacional de Tecnología Agropecuaria, CICVyA, Instituto de Biotecnología, IABIMO-CONICET, Unidad de Genómica | Distéfano et al |
| **hCoV-19/Panama/PA739543-GMI/2021** | EPI_ISL_11686205 | | Gorgas Memorial Institute of Health Studies | Gorgas Memorial Institute of Health Studies | Claudia Gonzalez et al |
| **hCoV-19/Panama/GMI-PASEQ590/2021** | EPI_ISL_11695725 | | Gorgas Memorial Institute of Health Studies | Gorgas Memorial Institute of Health Studies | Claudia Gonzalez et al |
| **hCoV-19/Chile/RM-68797/2021** | EPI_ISL_1534599 | | Genetica Molecular and Subdepartamento de Virologia ISP Chile | Instituto de Salud Publica de Chile | Javier Tognarelli et al |
| **hCoV-19/Mexico/QUE-InDRE_FB16443_S2829/2021** | EPI_ISL_2598001 | | LESP Queretaro | Instituto de Diagnostico y Referencia Epidemiologicos (INDRE) | Claudia Wong-Arambula et al |
| **hCoV-19/Ecuador/NIC-INSPI-343836/2021** | EPI_ISL_4496984 | | INSPI-CRN DE INFLUENZA Y OTROS VIRUS RESPIRATORIOS | NIC-INSPI | Alfredo Bruno et al |
| **hCoV-19/Peru/C02m42/2021** | EPI_ISL_3050726 | | Dirección regional de salud del Callao (DIRESA-CALLAO) | Centro de Investigaciones Tecnológicas, Biomédicas y Medioambientales (CITBM) | M. Sovero et al |
| **hCoV-19/HongKong/HKPU-00808/2020** | EPI_ISL_2713077 | | Department of Health Technology and Informatics, The Hong Kong Polytechnic University | Department of Health Technology and Informatics, The Hong Kong Polytechnic University | Gilman Kit-Hang Siu et al |
| **hCoV-19/Gambia/0547/2020** | EPI_ISL_471164 | | MRCG at LSHTM Genomics lab | MRCG at LSHTM Genomics lab | Sesay et al |
| **hCoV-19/Croatia/Zagreb-22/2020** | EPI_ISL_710570 | | University Hospital Dubrava | Ruder Boškovic Institute; Forensic Science Centre Ivan Vučetić; University of Zagreb Faculty of Science | Robert Belužić et al |
| **hCoV-19/Serbia/Belgrade-2972550801/2021** | EPI_ISL_1017696 | | Institute of Virology, Vaccines and Sera ''Torlak'' | Institute of microbiology and Immunology, Faculty of Medicine, University of Belgrade | Knezevic et al |
| **hCoV-19/Mongolia/Selenge-50797/2020** | EPI_ISL_9879599 | | National Center for Communicable Diseases (NCCD) National Influenza Center | National Center for Communicable Diseases (NCCD) National Influenza Center | Naranzul Tsedenbal et al |
| **hCoV-19/Beijing/BJCDC-220/2020** | EPI_ISL_1117378 | | Beijing Center for Disease Prevention and Control | Beijing Center for Disease Prevention and Control | Yang Pan et al |
| **hCoV-19/Mongolia/Darkhan-52609/2020** | EPI_ISL_9879601 | | National Center for Communicable Diseases (NCCD) National Influenza Center | National Center for Communicable Diseases (NCCD) National Influenza Center | Naranzul Tsedenbal et al |
| **hCoV-19/Mongolia/30/2020** | EPI_ISL_1805697 | | National Center for Communicable Diseases (NCCD) National Influenza Center | National Centre for Disease Control (NCDC) National Influenza Center | Naranzul Ts et al |
| **hCoV-19/Togo/NMIMR-TG-TGS-015-21/2020** | EPI_ISL_2001061 | MZ149969.1 | Immunology, Noguchi Memorial Institute for Medical Research | Immunology, Noguchi Memorial Institute for Medical Research | Halatoko et al |
| **hCoV-19/Estonia/Cov1369/2020** | EPI_ISL_2642891 | OU278770.1 | SYNLAB Eesti OÜ | Department of Microbiology, Institute of Biomedicine and Translational Medicine, University of Tartu | Radko Avi et al |
| **hCoV-19/Guinea/Conakry_IPG002/2020** | EPI_ISL_2245780 | | Institut Pasteur de Guinée | Institut Pateur de Dakar | Grayo Solene et al |
| **hCoV-19/Senegal/SN-IR5-001991/2020** | EPI_ISL_1827947 | | Institute for Health Research, Epidemiological Surveillance and Training (IRESSEF) | Abbott Laboratories | Souleymane Mboup et al |
| **hCoV-19/Senegal/SN-THI-THD-0293/2021** | EPI_ISL_13307538 | | Iressef Genomics Lab | IHU Mediterranée Infection, Genomics lab | Anna Julienne Selbé NDIAYE et al |
| **hCoV-19/Algeria/135160/2020** | EPI_ISL_15928096 | | NIC, Viral Respiratory Unit | NIC, Viral Respiratory Unit | Mohamed Amine Beloufa et al |
| **hCoV-19/Iceland/11222/2021** | EPI_ISL_3930176 | | The National University Hospital of Iceland | deCODE genetics | Daniel F Gudbjartsson et al B |
| **hCoV-19/Poland/415601621_diag1_gx1/2020** | EPI_ISL_875983 | | Laboratory of Molecular Biology, Diagnostyka sp. z o.o. | genXone SA, Research & Development Laboratory | Maciej Sykulski et al |
| **hCoV-19/Greece/6638/2020** | EPI_ISL_2301774 | | Laboratory of Molecular Biology, Mamatsio General Hospital of Kozani | Greek Genome Center, Biomedical Research Foundation of the Academy of Athens (BRFAA) | Emmanouil Athanasiadis et al |
| **hCoV-19/Iceland/3245/2020** | EPI_ISL_826937 | OU163669.1 | The National University Hospital of Iceland | deCODE genetics | Daniel F Gudbjartsson et al B |
| **hCoV-19/DominicanRepublic/UNIBE193/2020** | EPI_ISL_9671428 | | Instituto de Medicina Tropical & Salud Global, Universidad Iberoamericana (UNIBE) | Instituto de Medicina Tropical & Salud Global, Universidad Iberoamericana (UNIBE) | Paulino-Ramírez et al |
| **hCoV-19/SaudiArabia/KAIMRC-S611/2021** | EPI_ISL_3799776 | MZ895780.1 | Clinical Microbiology Lab, King Abdulaziz Medical City, Ministry of National Guard – Health Affairs, Riyadh, Saudi Arabia | Infectious Diseases Research, King Abdullah International Medical Research Center (KAIMRC), Riyadh, Saudi Arabia | Alghoribi et al |
| **hCoV-19/SaudiArabia/KAIMRC-S606/2021** | EPI_ISL_3799775 | MZ895778.1 | Clinical Microbiology Lab, King Abdulaziz Medical City, Ministry of National Guard – Health Affairs, Riyadh, Saudi Arabia | Infectious Diseases Research, King Abdullah International Medical Research Center (KAIMRC), Riyadh, Saudi Arabia | Alghoribi et al |
| **hCoV-19/SaudiArabia/KAIMRC-S586/2021** | EPI_ISL_3799876 | MZ895768.1 | Clinical Microbiology Lab, King Abdulaziz Medical City, Ministry of National Guard – Health Affairs, Riyadh, Saudi Arabia | Infectious Diseases Research, King Abdullah International Medical Research Center (KAIMRC), Riyadh, Saudi Arabia | Alghoribi et al |
| **hCoV-19/CostaRica/INC-0089/2020** | EPI_ISL_527757 | | Area De Salud Goicoechea 1 | Inciensa, Instituto Costarricense de Investigación y Enseñanza en Nutrición y Salud | Francisco Duarte et al |
| **hCoV-19/CostaRica/INC-0215/2020** | EPI_ISL_914821 | | HOSPITAL DR. ENRIQUE BALTODANO BRICEÑO | Inciensa, Instituto Costarricense de Investigación y Enseñanza en Nutrición y Salud | Francisco Duarte et al |
| **hCoV-19/CostaRica/INC-0214/2021** | EPI_ISL_914820 | | AREA DE SALUD SAN JUAN-SAN DIEGO-CONCEPCION 2 | Inciensa, Instituto Costarricense de Investigación y Enseñanza en Nutrición y Salud | Francisco Duarte et al |
| **hCoV-19/CostaRica/HNN-0368/2020** | EPI_ISL_1517412 | | Inciensa, Instituto Costarricense de Investigación y Enseñanza en Nutrición y Salud | Inciensa, Instituto Costarricense de Investigación y Enseñanza en Nutrición y Salud | Cristian Peréz-Corrales et al |
| **hCoV-19/CostaRica/CABANA-CR-UCR-INCIENSA-ITV-UBA-Uda-UGA-IMSS-UPC-1931-648486/2021** | EPI_ISL_7711624 | | Hospital Mexico | Universidad de Costa Rica e Instituto Costarricense de Investigacion y Ensenanza en Nutricion y Salud | Brenes Hebleen et al |
| **hCoV-19/India/MZ-ZMC3076A/2020** | EPI_ISL_1419087 | | INSACOG-Mizoram | National Institute of Biomedical Genomics – INSACOG | Arindam Maitra et al A |
| **hCoV-19/India/OR-ILS07/2020** | EPI_ISL_1255308 | | Institute of Life Sciences | Institute of Life Sciences | Sanchari Chatterjee et al |
| **hCoV-19/India/MZ-ZMC3334/2020** | EPI_ISL_1419105 | | INSACOG-Mizoram | National Institute of Biomedical Genomics – INSACOG | Arindam Maitra et al A |
| **hCoV-19/India/MN-ICMR-NIV-INSACOG-SEQ-2405/2020** | EPI_ISL_15038506 | | ICMR-National Institute of Virology - INSACOG | NIV Influenza | Dr. Varsha Potdar et al |
| **hCoV-19/Nepal/2097/2020** | EPI_ISL_754068 | | Nepal Korea Friendship Municipality Hospital | Nepal Health Research Council | Pradip Gyanwali et al |
| **hCoV-19/India/JH-ICMR-NIV-INSACOG-SEQ-2570/2020** | EPI_ISL_15038573 | | ICMR-National Institute of Virology - INSACOG | NIV Influenza | Dr. Varsha Potdar et al |
| **hCoV-19/Bhutan/RCDC0028-SV200584/2020** | EPI_ISL_16984389 | | AFRIMS | AFRIMS | Chonticha Klungthong et al |
| **hCoV-19/India/MZ-ZMC6230G/2020** | EPI_ISL_1419104 | | INSACOG-Mizoram | National Institute of Biomedical Genomics – INSACOG | Arindam Maitra et al A |
| **hCoV-19/India/MZ-ZMC3432H/2020** | EPI_ISL_1419095 | | INSACOG-Mizoram | National Institute of Biomedical Genomics – INSACOG | Arindam Maitra et al A |
| **hCoV-19/India/SK-1122700009269/2020** | EPI_ISL_1419124 | | INSACOG-Sikkim | National Institute of Biomedical Genomics – INSACOG | Arindam Maitra et al A |
| **hCoV-19/India/UP-MRUBHU3/2021** | EPI_ISL_862051 | | Multidisciplinary Research Unit, DHR-ICMR, Institute of Medical Sciences, Banaras Hindu University | Multidisciplinary Research Unit, DHR-ICMR, Institute of Medical Sciences, Banaras Hindu University | Priyoneel Basu et al |
| **hCoV-19/India/WB-1931500690592/2020** | EPI_ISL_1419446 | | INSACOG-WB | National Institute of Biomedical Genomics – INSACOG | Arindam Maitra et al A |
| **hCoV-19/India/CT-ICMR-NIV-INSACOG-SEQ-1460/2020** | EPI_ISL_15038561 | | ICMR-National Institute of Virology - INSACOG | NIV Influenza | Dr. Varsha Potdar et al |
| **hCoV-19/India/HR-ICMR-NIV-INSACOG-SEQ-376/2020** | EPI_ISL_15038480 | | ICMR-National Institute of Virology - INSACOG | NIV Influenza | Dr. Varsha Potdar et al |
| **hCoV-19/India/JH-ICMR-NIV-INSACOG-SEQ-2592/2020** | EPI_ISL_15038483 | | ICMR-National Institute of Virology - INSACOG | NIV Influenza | Dr. Varsha Potdar et al |
| **hCoV-19/India/TR-AGMC-R2074/2020** | EPI_ISL_2433258 | | INSACOG-Tripura | National Institute of Biomedical Genomics – INSACOG | Arindam Maitra et al A |
| **hCoV-19/India/WB-1930600111245/2020** | EPI_ISL_1419832 | | INSACOG-WB | National Institute of Biomedical Genomics – INSACOG | Arindam Maitra et al A |
| **hCoV-19/India/WB-S2/2020** | EPI_ISL_430468 | | ICMR-National Institute of Cholera and Enteric Diseases | National Institute of Biomedical Genomics | Arindam Maitra et al B (https://dx.doi.org/10.1007/s12038-020-00046-1) |
| **hCoV-19/TrinidadandTobago/25889/2020** | EPI_ISL_1490252 | | The Caribbean Public Health Agency | Carrington Lab, Department of PreClinical Sciences, Faculty of Medical Sciences, The University of the West Indies | Nikita S. D. Sahadeo et al |
| **hCoV-19/Nigeria/ISTH-E0220/2020** | EPI_ISL_14605232 | | Irrua Specialist Teaching Hospital, ISTH | "Irrua Specialist Teaching Hospital, ISTH, and the Bernhard Nocht Institute for Tropical Medicine, BNITM" | "Sylvanus Okogbenin et al |
| **hCoV-19/Nigeria/CV629/2020** | EPI_ISL_941278 | | Nigeria Centre for Disease Control (NCDC) | African Centre of Excellence for Genomics of Infectious Diseases (ACEGID), Redeemer's University | Oluniyi P.E. et al |
| **hCoV-19/Paraguay/33623/2020** | EPI_ISL_1340753 | | Departamento de Virologia, Laboratorio Central de Salud Pública, Avenida Venezuela y Teniente Escurra, Asunción, Paraguay | Laboratory of Respiratory Viruses and Measles, Oswaldo Cruz Institute, FIOCRUZ | Paola Resende et al |
| **hCoV-19/Uruguay/DLSP_648/2020** | EPI_ISL_11799567 | | Departamento Laboratorios de Salud Pública (DLSP) Ministerio de Salud Pública | Departamento Laboratorios de Salud Pública (DLSP) Ministerio de Salud Pública | Cortinas et al |
| **hCoV-19/Uruguay/UY-651/2020** | EPI_ISL_480339 | | Microbial Genomics Laboratory, Institut Pasteur de Montevideo | Microbial Genomics Laboratory, Institut Pasteur de Montevideo | Cecilia Salazar et al |
| **hCoV-19/Paraguay/WOR12903A607/2020** | EPI_ISL_6944512 | ON017193.1 | Laboratorio Central de Salud Publica | Viral Special Pathogens Branch, Centers for Disease Control and Prevention | Cynthia Vázquez et al |
| **hCoV-19/Brazil/AC-FIOCRUZ-34704/2020** | EPI_ISL_13498151 | | Laboratorio Rodolphe Merieux Rio Branco | Laboratory of Respiratory Viruses and Measles, Oswaldo Cruz Institute, FIOCRUZ | Paola Resende et al |
| **hCoV-19/Chile/BI-253479/2020** | EPI_ISL_1167849 | | Genetica Molecular and Subdepartamento de Virologia ISP Chile | Instituto de Salud Publica de Chile | Javier Tognarelli et al |
| **hCoV-19/Argentina/INEI100871/2021** | EPI_ISL_2104826 | | Servicio Virosis Respiratorias-Departamento Virología-INEI | Instituto Nacional Enfermedades Infecciosas C.G.Malbran | Baumeister E. et al |
| **hCoV-19/Argentina/PAIS-E0043/2020** | EPI_ISL_1395921 | | Laboratorio Central de la Ciudad de Santa Fe | Grupo de Genómica y Bioinformática del Instituto de Investigación de la Cadena Láctea CONICET-INTA on behalf of 'Proyecto Argentino Interinstitucional de genomica de SARS-CoV-2' (PAIS Consortium) | Eberhardt et al |
| **hCoV-19/Argentina/PAIS-H0109/2020** | EPI_ISL_1396420 | | Centro de Tecnología en Salud Pública de la Universidad Nacional de Rosario | Laboratorio Mixto de Biotecnología Acuática (LMBA) on behalf of 'Proyecto Argentino Interinstitucional de genomica de SARS-CoV-2' (PAIS Consortium) | Joaquín Ezpeleta et al |
| **hCoV-19/Brazil/SP-966/2020** | EPI_ISL_861659 | | PS e Maternidade Nair Fonseca Leitao Arantes | Instituto Adolfo Lutz, Interdiciplinary Procedures Center, Strategic Laboratory | Claudio Tavares Sacchi et al |
| **hCoV-19/Ecuador/174174_S54/2020** | EPI_ISL_18283537 | | INSPI | INSPI - Charité | Alfredo Bruno Caicedo et al |

| | | | | | |
|---|---|---|---|---|---|
| **hCoV-19/Qatar/QA-WCMQ_FD18163191/2020** | EPI_ISL_1714459 | | Ministry of Public Health / Hamad Medical Corporation | Weill Cornell Medical College - Qatar (WCM-Q), Genomics Core Laboratory / Qatar Genome Project (QGP) | WCMQ: Ayeda A. Ahmed et al |
| **hCoV-19/Brazil/SE-FIOCRUZ-1887/2021** | EPI_ISL_2660683 | | Laboratorio Central de Saude Publica do Estado de Sergipe (LACEN/SE) | Laboratory of Respiratory Viruses and Measles, Oswaldo Cruz Institute, FIOCRUZ | Paola Resende et al |
| **hCoV-19/Paraguay/631690/2021** | EPI_ISL_6944373 | ON017403.1 | Laboratorio Central de Salud Publica | Viral Special Pathogens Branch, Centers for Disease Control and Prevention | Cynthia Vázquez et al |
| **hCoV-19/Brazil/un-UW-31908/2020** | EPI_ISL_17745154 | | Laboratory Medicine and Pathology, University of Washington | Laboratory Medicine and Pathology, University of Washington | Roychoudhury et al |
| **hCoV-19/Brazil/MT-L120-CD7813/2021** | EPI_ISL_2344240 | | Instituto Butantan | Instituto de Medicina Tropical de Sao Paulo | Brazil-UK Centre for Arbovirus Discovery Diagnosis Genomics et al |
| **hCoV-19/Brazil/SP-IB_RP6848/2021** | EPI_ISL_14204562 | | Projeto S | Instituto Butantan | Dimas Tadeu Covas et al |
| **hCoV-19/Paraguay/631910/2021** | EPI_ISL_6944392 | ON017419.1 | Laboratorio Central de Salud Publica | Viral Special Pathogens Branch, Centers for Disease Control and Prevention | Cynthia Vázquez et al |
| **hCoV-19/Paraguay/33610/2020** | EPI_ISL_1340751 | | Departamento de Virologia, Laboratorio Central de Salud Pública, Avenida Venezuela y Teniente Escurra, Asunción, Paraguay | Laboratory of Respiratory Viruses and Measles, Oswaldo Cruz Institute, FIOCRUZ | Paola Resende et al |
| **hCoV-19/Philippines/PH-PGC-01060/2020** | EPI_ISL_2105418 | | Research Institute for Tropical Medicine, Inc. (RITM) | Philippine Genome Center | Francis A. Tablizo et al |
| **hCoV-19/Paraguay/631454/2021** | EPI_ISL_6944345 | ON017375.1 | Laboratorio Central de Salud Publica | Viral Special Pathogens Branch, Centers for Disease Control and Prevention | Cynthia Vázquez et al |
| **hCoV-19/Uruguay/CUY1-000001-UYTT/2021** | EPI_ISL_2965573 | | Laboratorio de Biología Molecular del Sanatorio Americano | Centro de Innovación en Vigilancia Epidemiológica (CiVE), Institut Pasteur Montevideo, Uruguay | Natalia Rego et al |
| **hCoV-19/Mauritius/P6913/2021** | EPI_ISL_13833237 | | Virology Department, Central Health Laboratory | The Francis Crick Institute | Ramuth M et al |
| **hCoV-19/Uruguay/UY-NYULH2060/2021** | EPI_ISL_2427638 | | Laboratorio de Biología Molecular Médica Uruguaya | Departments of Pathology and Medicine, New York University School of Medicine | Maria Victoria Elizondo et al |
| **hCoV-19/UnitedArabEmirates/2582/2020** | EPI_ISL_859663 | | BTC, Khalifa University | BTC, Khalifa University | Al Safar et al |
| **hCoV-19/BurkinaFaso/7990/2021** | EPI_ISL_2289109 | | Laboratoire de Microbiologie CHU Souro Sanou | Laboratoire bacteriologie virologie CHUSS | Yacouba Sawadogo et al |
| **hCoV-19/Qingdao/IVDC-01-01/2020** | EPI_ISL_850949 | | National Institute for Viral Disease Control and Prevention, China CDC | National Institute for Viral Disease Control and Prevention, China CDC | Xiang Zhao et al |
| **hCoV-19/BosniaandHerzegovina/VFS-UNSA-LMGFI009/2020** | EPI_ISL_955155 | | University of Sarajevo, Veterinary Faculty, Laboratory for Molecular Diagnostic and Research Laboratory | University of Sarajevo, Veterinary Faculty, Laboratory for Molecular Diagnostic and Research Laboratory | Goletić T. et al |
| **hCoV-19/Gabon/K0361/2020** | EPI_ISL_11522712 | | Centre de Recherches Medicales de Lambarene | Department of Emerging Infectious Diseases, Institute of Tropical Medicine, Nagasaki University | Haruka Abe et al |
| **hCoV-19/Gabon/1833-33A/2020** | EPI_ISL_8886170 | OM731973.1 | Centre Interdisciplinaire de Recherches Médicales de Franceville (CIRMF) | Institut Hospitalo-Universitaire - Méditerranée Infection | Sonia Etenna Lekana-Douki et al |
| **hCoV-19/Gabon/S0274/2020** | EPI_ISL_11522684 | | Centre de Recherches Medicales de Lambarene | Department of Emerging Infectious Diseases, Institute of Tropical Medicine, Nagasaki University | Haruka Abe et al |
| **hCoV-19/Gabon/R0703/2020** | EPI_ISL_11522777 | | Centre de Recherches Medicales de Lambarene | Department of Emerging Infectious Diseases, Institute of Tropical Medicine, Nagasaki University | Haruka Abe et al |
| **hCoV-19/Gabon/V0242/2020** | EPI_ISL_11522784 | | Centre de Recherches Medicales de Lambarene | Department of Emerging Infectious Diseases, Institute of Tropical Medicine, Nagasaki University | Haruka Abe et al |
| **hCoV-19/Oman/520278568/2020** | EPI_ISL_2925612 | | Oman-NIC | Oman-National Influenza Center | Samiha Al Kharusi et al |
| **hCoV-19/UnitedArabEmirates/16B/2020** | EPI_ISL_528690 | | Alsafar - Khalifa University Abu Dhabi | Alsafar - Khalifa University Abu Dhabi | Andreas Henschel et al |
| **hCoV-19/Madagascar/IPM-00608/2021** | EPI_ISL_1660300 | | Virology Unit, Institut Pasteur de Madagascar | Virology Unit, Institut Pasteur de Madagascar | Marion Barbet et al A |
| **hCoV-19/Jordan/AM-ALSR-5138/2020** | EPI_ISL_730377 | | Biolab Diagnostic Laboratories | Andersen lab at Scripps Research | Issa Abu-Dayyeh et al |
| **hCoV-19/Jordan/SEARCH-5460/2020** | EPI_ISL_755247 | | Biolab Diagnostic Laboratories | Andersen lab at Scripps Research | Issa Abu-Dayyeh et al |
| **hCoV-19/Jordan/Biolab0068/2021** | EPI_ISL_2617459 | | Biolab Diagnostic Laboratories | Biolab Diagnostic Laboratories | Issa Abu-Dayyeh et al |
| **hCoV-19/Jordan/IR-ALSR-5200/2020** | EPI_ISL_730438 | | Biolab Diagnostic Laboratories | Andersen lab at Scripps Research | Issa Abu-Dayyeh et al |
| **hCoV-19/env/Liaoning/Pollock-IVDC-30-05/2020** | EPI_ISL_2193221 | | National Institute for Viral Disease Control and Prevention, China CDC | National Institute for Viral Disease Control and Prevention, China CDC | Huilai Ma1& et al |
| **hCoV-19/env/Liaoning/Dalian-IVDC-Pollock-03-06/2020** | EPI_ISL_2170894 | | National Institute for Viral Disease Control and Prevention, China CDC | National Institute for Viral Disease Control and Prevention, China CDC | Huilai Ma1& et al |
| **hCoV-19/Liaoning/Case-3-IVDC-27-08/2020** | EPI_ISL_2177398 | | National Institute for Viral Disease Control and Prevention, China CDC | National Institute for Viral Disease Control and Prevention, China CDC | Huilai Ma1& et al |
| **hCoV-19/Liaoning/IVDC-01/2020** | EPI_ISL_498692 | | National Institute for Viral Disease Control and Prevention, China CDC | National Institute for Viral Disease Control and Prevention, China CDC | Xiang Zhao et al |
| **hCoV-19/Liaoning/IVDC-YK-214/2021** | EPI_ISL_9301317 | | Liaoning Center for Disease Control and Prevention | National Institute for Viral Disease Control and Prevention, China CDC | Haibo Sun et al |
| **hCoV-19/Lu-an/5053/2021** | EPI_ISL_2132218 | | Microbiology Department, Lu`an Center for Disease Control and Prevention | Microbiology Department, Lu`an Center for Disease Control and Prevention | Chen Zhichao et al |
| **hCoV-19/Paraguay/239808/2021** | EPI_ISL_13147992 | | Laboratorio CYRLAB | Laboratorio Central de Salud Pública | Sandra Gonzalez et al |
| **hCoV-19/Venezuela/Ara105/2021** | EPI_ISL_15104842 | | INHRR | Laboratorio de Virologia Molecular | Rossana C Jaspe et al |
| **hCoV-19/Venezuela/Bol4715/2021** | EPI_ISL_2628036 | MZ611956.1 | INHRR | Laboratorio de Virología Molecular | Rossana C Jaspe et al |
| **hCoV-19/TrinidadandTobago/64267/2021** | EPI_ISL_2658392 | | The Caribbean Public Health Agency | Carrington Lab, Department of Preclinical Sciences, Faculty of Medical Sciences, The University of the West Indies, St Augustine Campus | Nikita S. D. Sahadeo et al |
| **hCoV-19/TrinidadandTobago/58605/2021** | EPI_ISL_8939202 | | Trinidad Public Health Laboratory | Carrington Lab, Department of Preclinical Sciences, Faculty of Medical Sciences, The University of the West Indies, St Augustine Campus | Nikita S. D. Sahadeo et al |
| **hCoV-19/TrinidadandTobago/46813/2021** | EPI_ISL_3610772 | | Trinidad Public Health Laboratory | Carrington Lab, Department of Preclinical Sciences, Faculty of Medical Sciences, The University of the West Indies, St Augustine Campus | Nikita S. D. Sahadeo et al |
| **hCoV-19/Haiti/63230037/2021** | EPI_ISL_4029006 | | Laboratoire National de Santé Publique – LNSP | Genomics and Proteomics Departament, Gorgas Memorial Institute For Health Studies | César Roberto Conde Pereira et al |
| **hCoV-19/Aruba/AW-RIVM-41532/2021** | EPI_ISL_2787924 | | National Institute for Public Health and the Environment (RIVM) | National Institute for Public Health and the Environment (RIVM) | Adam Meijer et al |
| **hCoV-19/Peru/CAJ-INS-1795/2021** | EPI_ISL_3376010 | | Laboratorio de Referencial Nacional de Virus Respiratorios | Laboratorio de Referencial Nacional de Virus Respiratorios | Carlos Padilla Rojas et al |
| **hCoV-19/Brazil/SP-IB_100113/2021** | EPI_ISL_1966158 | | COMPLEXO HOSPITALAR OURO VERDE DE CAMPINAS | Instituto Butantan / Mendelics | Instituto Butantan: Dimas Tadeu Covas et al |
| **hCoV-19/USA/ID-CDC-LC0073079/2021** | EPI_ISL_4377689 | OK232871.1 | Laboratory Corporation of America | Centers for Disease Control and Prevention Division of Viral Diseases, Pathogen Discovery | Dakota Howard et al |
| **hCoV-19/Peru/LAM-INS-10890/2021** | EPI_ISL_8143839 | | Laboratorio de Referencia Nacional de Virus Respiratorios. Centro Nacional de Salud Publica. Instituto Nacional de Salud Peru. | Laboratorio de Referencia Nacional de Virus Respiratorios. Centro Nacional de Salud Publica. Instituto Nacional de Salud Peru. | Carlos Padilla Rojas et al |
| **hCoV-19/Peru/LIM-INS-10966/2021** | EPI_ISL_8143842 | | Laboratorio de Referencia Nacional de Virus Respiratorios. Centro Nacional de Salud Publica. Instituto Nacional de Salud Peru. | Laboratorio de Referencia Nacional de Virus Respiratorios. Centro Nacional de Salud Publica. Instituto Nacional de Salud Peru. | Carlos Padilla Rojas et al |
| **hCoV-19/Chile/RM-70789/2021** | EPI_ISL_1633516 | | Genetica Molecular and Subdepartamento de Virologia ISP Chile | Instituto de Salud Publica de Chile | Javier Tognarelli et al |
| **hCoV-19/Argentina/INEI111303/2021** | EPI_ISL_6694837 | | Servicio Virosis Respiratorias-Departamento Virología-INEI | Instituto Nacional Enfermedades Infecciosas C.G.Malbran | Baumeister E. et al |
| **hCoV-19/Chile/ML-102815/2021** | EPI_ISL_2663419 | | Genetica Molecular and Subdepartamento de Virologia ISP Chile | Instituto de Salud Publica de Chile | Karen Orostica et al |
| **hCoV-19/Paraguay/WDR12903A838/2021** | EPI_ISL_6944588 | ON017269.1 | Laboratorio Central de Salud Publica | Viral Special Pathogens Branch, Centers for Disease Control and Prevention | Cynthia Vázquez et al |
| **hCoV-19/Guangdong/IVDC-05-02/2021** | EPI_ISL_1969692 | | Guangdong Provincial Center for Disease Control and Prevention | National Institute for Viral Disease Control and Prevention, China CDC | Yao Hu et al |
| **hCoV-19/Canada/NL-NML-62014/2021** | EPI_ISL_2484584 | | NL-Dr. Leonard A. Miller Centre for Health Services | National Microbiology Laboratory (NML) | Anna Majer et al |
| **hCoV-19/Canada/BC-BCCDC-122791/2021** | EPI_ISL_3601774 | | British Columbia Centre For Disease Control | BCCDC Public Health Laboratory | Prystajecky Natalie et al |
| **hCoV-19/Brazil/SP-IB_114728/2021** | EPI_ISL_5647450 | | LABORATORIO MUNICIPAL DE PIRACICABA | Instituto Butantan | Dimas Tadeu Covas et al |
| **hCoV-19/Luxembourg/LNS5806719/2021** | EPI_ISL_3148178 | | Laboratoires d'analyses medicales - Ketterthill | Laboratoire national de sante, Microbiology, Microbial Genomics Platform | Anke Wienecke-Baldacchino et al |
| **hCoV-19/Mexico/YUC-InDRE_FB25259_S5616/2021** | EPI_ISL_3769145 | | LESP Yucatan | Instituto de Diagnostico y Referencia Epidemiologicos (INDRE) | Claudia Wong-Arambula et al |
| **hCoV-19/Suriname/SR-446/2021** | EPI_ISL_3462687 | | Central Laboratory, Bureau of Public Health (BOG) and Academic Hospital Paramaribo | Erasmus Medical Center | Bas B Oude Munnink et al |
| **hCoV-19/Suriname/SR-363/2021** | EPI_ISL_3462543 | | Central Laboratory, Bureau of Public Health (BOG) and Academic Hospital Paramaribo | Erasmus Medical Center | Bas B Oude Munnink et al |
| **hCoV-19/Uruguay/DLSP_p754/2021** | EPI_ISL_11799605 | | Departamento Laboratorios de Salud Pública (DLSP) Ministerio de Salud Pública | Departamento Laboratorios de Salud Pública (DLSP) Ministerio de Salud Pública | Cortinas et al |
| **hCoV-19/Uruguay/DLSP_4302/2021** | EPI_ISL_12532629 | | Departamento Laboratorios de Salud P?blica (DLSP) Ministerio de Salud P?blica | Departamento Laboratorios de Salud P?blica (DLSP) Ministerio de Salud P?blica | Cortinas et al |
| **hCoV-19/Peru/LOR-FPY01506/2021** | EPI_ISL_4772372 | | NAMRU-6 | NAMRU-6 | Gilda Troncos et al |
| **hCoV-19/USA/NM-CDC-LC0058820/2021** | EPI_ISL_2184905 | MZ254007.1 | Laboratory Corporation of America | Centers for Disease Control and Prevention Division of Viral Diseases, Pathogen Discovery | Dakota Howard et al |
| **hCoV-19/Chile/RM-0263-04985941/2021** | EPI_ISL_2467275 | | Clinica INDISA | "Facultad de Ciencias de la Vida, UNAB" | "Claudio Meneses et al |
| **hCoV-19/Brazil/AM-FIOCRUZ-COVAC08304/2021** | EPI_ISL_4030349 | | Laboratorio de Ecologia de Doencas Transmissiveis na Amazonia, Instituto Leonidas e Maria Deane - Fiocruz Amazonia | Laboratorio de Ecologia de Doencas Transmissiveis na Amazonia, Instituto Leonidas e Maria Deane - Fiocruz Amazonia | Marcus Vinicius Guimarães de Lacerda et al |
| **hCoV-19/Uruguay/DLSP_3735/2021** | EPI_ISL_12532630 | | Departamento Laboratorios de Salud P?blica (DLSP) Ministerio de Salud P?blica | Departamento Laboratorios de Salud P?blica (DLSP) Ministerio de Salud P?blica | Cortinas et al |
| **hCoV-19/Bolivia/SAN-CABANA-PE-UPCH-2258/2021** | EPI_ISL_8600640 | | Laboratorio de Diagnóstico e Investigación BIOSCIENCE SRL | Laboratorio de Genómica Microbiana, Universidad Peruana Cayetano Heredia | Esdenka Pérez et al |
| **hCoV-19/Venezuela/Mon1645/2021** | EPI_ISL_11629629 | | IVIC | Laboratorio de Virologia Molecular | Rossana C Jaspe et al |
| **hCoV-19/Chile/RM-134835/2021** | EPI_ISL_4005248 | | Genetica Molecular and Subdepartamento de Virologia ISP Chile | Instituto de Salud Publica de Chile | Karen Orostica et al |
| **hCoV-19/Colombia/ANT-PRO-LDSP367/2021** | EPI_ISL_4820605 | | HPTU | Laboratorio Departamental de Salud Publica de Antioquia | Idabely Betancur Ortiz et al |
| **hCoV-19/USA/OK-QDX-3311/2020** | EPI_ISL_884404 | MW522470.1 | Infectious Diseases, Quest Diagnostics | Infectious Diseases, Quest Diagnostics | Rosenthal et al |
| **hCoV-19/Ireland/D-NVRL-20Z16503/2020** | EPI_ISL_981264 | | National Virus Reference Laboratory | Irish Coronavirus Sequencing Consortium - Hellsworks | Sachin Chalapati et al |
| **hCoV-19/Colombia/COR-U13/2020** | EPI_ISL_2339864 | | Universidad de Cordoba, Montería, Colombia | Centro de Investigaciones en Microbiología y Biotecnología-UR (CIMBIUR), Facultad de Ciencias Naturales, Universidad del Rosario, Bogotá, Colombia | Salim Mattar et al |
| **hCoV-19/Venezuela/Mir3290/2021** | EPI_ISL_16895329 | | Instituto Nacional de Higiene Rafael Rangel | Laboratorio de Virologia Molecular | Rossana C Jaspe et al |
| **hCoV-19/Bangladesh/DNAS_DRCIM_isl_6300/2020** | EPI_ISL_774908 | | Designated Reference Institute for Chemical Measurements (DRICM) | DNA SOLUTION LTD. | Md. Imran Khan et al |
| **hCoV-19/Bangladesh/ECHD_00508/2020** | EPI_ISL_2350800 | LC647135.1 | Molecular Lab, Evercare hospital Dhaka | International Institute for Zoonosis Control, Hokkaido university | Junya Yamagishi et al |
| **hCoV-19/Bangladesh/CHRF-0468/2020** | EPI_ISL_625462 | | Child Health Research Foundation | Child Health Research Foundation | Senjuti Saha et al |
| **hCoV-19/Australia/WA190/2020** | EPI_ISL_596753 | | PathWest Laboratory Medicine WA | PathWest Laboratory Medicine WA Microbial Surveillance Unit | PathWest Laboratory Medicine WA Microbial Surveillance Unit et al |
| **hCoV-19/Bangladesh/BCSIR-NILMRC_426/2020** | EPI_ISL_603247 | | National Institute of Laboratory Medicine and Referral Center | Genomic Research Lab, BCSIR | Md. Saddam Hossain et al |
| **hCoV-19/Bangladesh/CHRF-0147/2021** | EPI_ISL_1360423 | | Child Health Research Foundation | Child Health Research Foundation | CHRF Bangladesh Genomics Team et al |
| **hCoV-19/UnitedArabEmirates/3880/2020** | EPI_ISL_859887 | | BTC, Khalifa University | BTC, Khalifa University | Al Safar et al |
| **hCoV-19/BurkinaFaso/CV1891/2021** | EPI_ISL_4255115 | | Centre Hospitalier Universitaire Yalgado OUEDRAOGO | Africa Centre for Excellence for Genomics of Infectious Diseases (ACEGID), Redeemer's University | Olawoye et al |
| **hCoV-19/Canada/NB-NML-3282/2020** | EPI_ISL_961608 | | Hôpital Georges L. Dumont | National Microbiology Laboratory (NML) | Anna Majer et al |
| **hCoV-19/Canada/NB-NML-3280/2020** | EPI_ISL_961607 | | Hôpital Georges L. Dumont | National Microbiology Laboratory (NML) | Anna Majer et al |
| **hCoV-19/Uzbekistan/Tashkent-CGB-32/2020** | EPI_ISL_1477049 | | Sanitary-Epidemiological And Public Health Department Of Tashkent Region, Uzbekistan | Center of Genomics and bioinformatics, Bioinformatics laboratory | Mirzakamol S Ayubov et al |
| **hCoV-19/Ghana/WACCBIP-GS367/2020** | EPI_ISL_1255240 | | West African Centre for Cell Biology of Infectious Pathogens (WACCBIP), University of Ghana, Accra, Ghana | West African Centre for Cell Biology of Infectious Pathogens (WACCBIP), University of Ghana, Volta Road, Legon-Accra, Ghana | Collins M. Morang´a et al |
| **hCoV-19/Tunisia/6895/2020** | EPI_ISL_763065 | MW428253.1 | 1-Laboratory of Microbiology, National Reference Lab, Charles Nicolle Hospital; 2-University of Tunis El Manar, Faculty of Medicine of Tunis, LR99ES09, Tunis, Tunisia | 1-Clinical and Experimental Pharmacology Lab, LR16SP02, National Center of Pharmacovigilance, University of Tunis El Manar, Tunis, Tunisia. 2-Neurodegenerative diseases and psychiatric troubles, LR18SP03, Razi Hospital, University of Tunis El Manar, Tunis, Tunisia. 3- Ministry of Health, National Observatory of New and Emerging Diseases, 1006, Tunis, Tunisia | Ilhem Boutiba-Ben Boubaker et al |
| **hCoV-19/Venezuela/VEN-89312/2020** | EPI_ISL_476702 | | Incubadora Venezolana de Ciencia, Venezuela | Incubadora Venezolana de Ciencia, Venezuela / Instituto Nacional de Salud, Bogotá, Colombia / Grupo de Investigaciones Microbiológicas-UR (GIMUR), Departamento de Biología, Facultad de Ciencias Naturales, Universidad del Rosario, Bogotá, Colombia / Icahn School of Medicine at Mount Sinai, New York, USA | Alberto Paniz-Mondolfi et al |
| **hCoV-19/Madagascar/IPM-11980/2020** | EPI_ISL_3716442 | | Virology Unit, Institut Pasteur de Madagascar | Institut Pateur de Dakar | Norosoa Razanajatovo et al |
| **hCoV-19/Oman/520259611/2020** | EPI_ISL_1532298 | | Oman-National Influenza Center | Biotechnology & OMICs Laboratory | Abdul Latif Khan et al |
| **hCoV-19/Oman/205042822/2020** | EPI_ISL_492020 | | Oman-NIC | Department of Microbiology and Immunology-SQUH | Fahad Zadjali et al |
| **hCoV-19/Oman/520279487/2020** | EPI_ISL_1532305 | | Oman-National Influenza Center | Biotechnology & OMICs Laboratory | Abdul Latif Khan et al |
| **hCoV-19/Oman/B-7106/2020** | EPI_ISL_491145 | | Oman-National Influenza Center | Biotechnology & OMICs Laboratory | Samiha Al-Kharusi et al |
| **hCoV-19/Oman/205033293/2020** | EPI_ISL_457983 | | Oman-NIC | Department of Microbiology and Immunology-SQUH | Fahad Zadjali et al |
| **hCoV-19/Oman/13564202/2020** | EPI_ISL_1819072 | | Oman-NIC | Oman-National Influenza Center-Department of Microbiology and Immunology-SQUH | Samira Al-Maruqi et al |
| **hCoV-19/Oman/205038906/2020** | EPI_ISL_492001 | | Oman-NIC | Department of Microbiology and Immunology-SQUH | Fahad Zadjali et al |
| **hCoV-19/Greece/60988/2020** | EPI_ISL_2232122 | | Central Public Health Laboratory, National Public Health Organization | Greek Genome Center, Biomedical Research Foundation of the Academy of Athens (BRFAA) | Emmanouil Athanasiadis et al |
| **hCoV-19/Ecuador/UEES-11445/2021** | EPI_ISL_1443666 | | Omics Sciences Laboratory | Omics Sciences Laboratory | Dery Andrade Molina et al |
| **hCoV-19/Latvia/883/2020** | EPI_ISL_1312394 | | E. Gulbja Laboratorija | Latvian Biomedical Research and Study Centre | Janis Pjatkovskis et al |
| **hCoV-19/BosniaandHerzegovina/VFS-UNSA-LMGFI006/2020** | EPI_ISL_955148 | | University of Sarajevo, Veterinary Faculty, Laboratory for Molecular Diagnostic and Research Laboratory | University of Sarajevo, Veterinary Faculty, Laboratory for Molecular Diagnostic and Research Laboratory | Goletić T. et al |
| **hCoV-19/Switzerland/BS-UHB-42501845/2020** | EPI_ISL_931104 | | Clinical Virology | Clinical Bacteriology | Tim Roloff et al |
| **hCoV-19/Austria/MUW_1327056/2020** | EPI_ISL_913095 | | Center for Virology | Center for Virology | Jeremy V. Camp et al |
| **hCoV-19/Switzerland/ZH-ETHZ-300401/2020** | EPI_ISL_603367 | | Viollier AG | Department of Biosystems Science and Engineering, ETH Zürich | Christian Beisel et al |
| **hCoV-19/Azerbaijan/un-ChVir24287_59/2020** | EPI_ISL_9214590 | | Ministry of Health of the Republic of Azerbaijan | Charité Universitätsmedizin Berlin, Institut für Virologie | Victor M Corman et al |
| **hCoV-19/Azerbaijan/un-ChVir6070/2020** | EPI_ISL_9214449 | | Ministry of Health of the Republic of Azerbaijan | Charité Universitätsmedizin Berlin, Institut für Virologie | Victor M Corman et al |
| **hCoV-19/Ghana/WACCBIP-GS477/2020** | EPI_ISL_2422525 | | West African Centre for Cell Biology of Infectious Pathogen, University of Ghana, Legon | WACCBIP, University of Ghana, Volta Road, Legon, Accra | Collins M. Morang´a et al |
| **hCoV-19/Mongolia/UB-15104/2020** | EPI_ISL_9879589 | | National Center for Communicable Diseases (NCCD) National Influenza Center | National Center for Communicable Diseases (NCCD) National Influenza Center | Naranzul Tsedenbal et al |
| **hCoV-19/Iran/Gilan-NICB1-58/2020** | EPI_ISL_959281 | | National Influenza Center, Virology Department | National Influenza Center | NZ Shafiei Jandaghi et al |
| **hCoV-19/Philippines/PH-RITM-495/2020** | EPI_ISL_10942927 | | LAS PI | Research Institute for Tropical Medicine | Inez Andrea Medado et al |
| **hCoV-19/Ecuador/EC-SD-119429/2020** | EPI_ISL_826827 | | INSPI-CRN DE INFLUENZA Y OTROS VIRUS RESPIRATORIOS | Instituto de Salud Publica de Chile | Javier Tognarelli et al |
| **hCoV-19/SaudiArabia/KAUST-JEDDAH1325/2020** | EPI_ISL_677932 | OV079428.1 | Pathogen Genomics Lab King Abdullah University of Science and Technology(KAUST) | Pathogen Genomics Lab King Abdullah University of Science and Technology(KAUST) | Sara Mfarrej et al |
| **hCoV-19/Tunisia/F1689/2020** | EPI_ISL_11172673 | | Laboratory of Clinical Virology, Institut Pasteur de Tunis | Laboratory of Clinical Virology, Institut Pasteur de Tunis | Haifa Jmel et al |
| **hCoV-19/Niger/ZR-306/2020** | EPI_ISL_2281764 | | Centre de Recherche Médicale et Sanitaire (CERMES) | Institut Pasteur de Dakar | Lagare Adamou et al |
| **hCoV-19/Russia/KAM-RII-MH123545/2020** | EPI_ISL_1039051 | | Kamchatka Regional Children's Infectious Diseases Hospital | WHO National Influenza Centre Russian Federation | Andrey Komissarov et al |
| **hCoV-19/USA/NY-PRL-2021_0706_01K11/2020** | EPI_ISL_2924008 | | Pandemic Response Lab - NYC | Pandemic Response Lab, R&D | Henry Lee et al |
| **hCoV-19/USA/NC-UNCC-000131/2020** | EPI_ISL_1937974 | | UNC Charlotte COVID-19 Testing Lab | UNC Charlotte Environmental Monitoring Laboratory | Cynthia Gibas et al |
| **hCoV-19/Serbia/KosovskaMitrovica-A07438-3004/2020** | EPI_ISL_6959696 | | Huo-yan national laboratory for molecular detection of infectious agents Belgrade | Center for forensic and applied molecular genetics, Faculty of Biology, University of Belgrade | D.Keckarević et al |
| **hCoV-19/NorthMacedonia/1692/2021** | EPI_ISL_946913 | | Laboratory of Virology and Molecular Diagnostics | Institute of Public Health of Republic of North Macedonia Laboratory of Virology and Molecular Diagnostics | Maja Kuzmanovska et al |
| **hCoV-19/NorthMacedonia/CoV-16442/2020** | EPI_ISL_933566 | | General Hospital - Bitola | Research Center for Genetic Engineering and Biotechnology "Georgi D. Efremov" , Macedonian Academy of Sciences and Arts | RCGEB - MASA et al |
| **hCoV-19/Serbia/Ritopek-B89189-1208/2020** | EPI_ISL_6842233 | | Huo-yan national laboratory for molecular detection of infectious agents Belgrade | Center for forensic and applied molecular genetics, Faculty of Biology, University of Belgrade | D.Keckarević et al |
| **hCoV-19/Thailand/A21952/2020** | EPI_ISL_3892178 | | Port Health and Quarantine, Suvarnabhumi Airport | Thai Red Cross Emerging Infectious Diseases Clinical Center and Faculty of Medicine, Chulalongkorn University | Rome Buathong et al |
| **hCoV-19/Serbia/C20229/2020** | EPI_ISL_1056974 | | Institute of Biocides and Medical Ecology, Belgarde, Serbia | Virology Department Institute of Microbiology and Immunology Faculty of Medicine University of Belgrade | Banko Ana et al |
| **hCoV-19/NorthMacedonia/15060/2020** | EPI_ISL_910326 | | General Hospital - Strumica | Research Center for Genetic Engineering and Biotechnology "Georgi D. Efremov" , Macedonian Academi | RCGEB - MASA et al |
| **hCoV-19/China/AP62/2020** | EPI_ISL_2779638 | | State Key Laboratory of Emerging Infectious Diseases, The University of Hong Kong | The University of Hong Kong | Lifeng Li et al |
| **hCoV-19/Armenia/UW-ARM49/2020** | EPI_ISL_1854843 | MZ230721.1 | National Center of Disease Control and Prevention of the Republic of Armenia | UW Virology Lab | Pavitra Roychoudhury et al |
| **hCoV-19/Kazakhstan/53468/2020** | EPI_ISL_1335044 | | Laboratory of virology, National center of expertise | RSE "National Center for Biotechnology" and RSE "National Center of Expertise" | Shevtsov Alexandr et al |
| **hCoV-19/Hebei/IVDC-04-05/2021** | EPI_ISL_796019 | | Hebei Provincial Center for Disease Control and Prevention, Shijiazhuang, Hebei Province; National Institute for Viral Disease Control and Prevention, China CDC | Hebei Provincial Center for Disease Control and Prevention, Shijiazhuang, Hebei Province; National Institute for Viral Disease Control and Prevention, China CDC | Shunxiang Qi et al |
| **hCoV-19/Russia/StPetersburg-RII16735/2020** | EPI_ISL_507270 | | WHO National Influenza Centre Russian Federation | WHO National Influenza Centre Russian Federation | Andrey Komissarov et al |
| **hCoV-19/Turkey/24_Ankara_GUMV_50471/2020** | EPI_ISL_984985 | | Gazi University Faculty of Medicine, Medical Virology Laboratory | Gazi University Faculty of Medicine, Medical Virology Laboratory | Erdem Şahin et al |
| **hCoV-19/Bulgaria/21BG-NC_A000009/2020** | EPI_ISL_2081878 | | National Center of Infectious and Parasitic Diseases | National Center of Infectious and Parasitic Diseases | Alexiev et al |
| **hCoV-19/Lebanon/MDB_58/2020** | EPI_ISL_12470091 | | Rafik Hariri University Hospital | Khazen Lab | Rola Husni et al |
| **hCoV-19/Madagascar/IPM-02027/2021** | EPI_ISL_1660280 | | Virology Unit, Institut Pasteur de Madagascar | Virology Unit, Institut Pasteur de Madagascar | Christian Ranavoson et al |
| **hCoV-19/Sweden/20-21500/2020** | EPI_ISL_582778 | | Uppsala klinisk mikrobiologi | The Public Health Agency of Sweden | Anna-Malin Linde et al |
| **hCoV-19/Qingdao/IVDC-02-10/2020** | EPI_ISL_591271 | | National Institute for Viral Disease Control and Prevention, China CDC | National Institute for Viral Disease Control and Prevention, China CDC | Huilai Ma et al |
| **hCoV-19/Ireland/D-NVRL-AIIDV0432v2-3/2020** | EPI_ISL_768760 | | AIID | Irish Coronavirus Sequencing Consortium - National Virus Reference Laboratory | Michael Carr et al |
| **hCoV-19/NewZealand/21CH0007/2021** | EPI_ISL_1082255 | | Canterbury Health Laboratories | Institute of Environmental Science and Research (ESR) | Paula scholes et al |
| **hCoV-19/Morocco/FMP_77/2021** | EPI_ISL_1116592 | | Laboratoire central de Virologie | Laboratoire de Biotechnologie | Hakima Kabbaj et al |
| **hCoV-19/Nepal/DH_002/2020** | EPI_ISL_2171041 | | Dhulikhel Hospital, Kathmandu University Hospital | Molecular and Genomics Research Lab, Dhulikhel Hospital, Kathmandu University Hospital | Rajeev Shrestha et al |
| **hCoV-19/Heilongjiang/HU-AUMC-000648/2021** | EPI_ISL_1678790 | | Department of Medical Microbiology & Infection prevention, Amsterdam University Medical Centers location AMC | Department of Medical Microbiology & Infection prevention, Amsterdam University Medical Centers location AMC | Matthijs Welkers et al |
| **hCoV-19/Mauritius/155354/2021** | EPI_ISL_3231389 | | Virology Department,Central Health Laboratory | UMR PIMIT | Manoj SS et al |
| **hCoV-19/Mauritius/181965/2021** | EPI_ISL_13830541 | | Virology Department, Central Health Laboratory | The Francis Crick Institute | Ramuth M et al |
| **hCoV-19/Uganda/UG373/2021** | EPI_ISL_2346387 | | MRC/UVRI & LSHTM Uganda Research Unit, Central Public Health Laboratories | MRC/UVRI & LSHTM Uganda Research Unit, Central Public Health Laboratories | Matthew Cotten et al |
| **hCoV-19/Liechtenstein/FL-Risch-2142909658/2021** | EPI_ISL_3570072 | | Dr. Risch Ostschweiz AG | Microbiology, Dr. Risch | Nadia Wohlwend et al |
| **hCoV-19/Beijing/BJCDC-067/2021** | EPI_ISL_8607167 | | Beijing Center for Disease Control and Prevention | Beijing Center for Disease Control and Prevention | Hui Xu et al |
| **hCoV-19/Gabon/CC0432/2021** | EPI_ISL_11522677 | | Centre de Recherches Medicales de Lambarene | Department of Emerging Infectious Diseases, Institute of Tropical Medicine, Nagasaki University | Haruka Abe et al |
| **hCoV-19/Gabon/EE0525/2021** | EPI_ISL_11522596 | | Centre de Recherches Medicales de Lambarene | Department of Emerging Infectious Diseases, Institute of Tropical Medicine, Nagasaki University | Haruka Abe et al |
| **hCoV-19/Guinea/CERFIG-24250/2021** | EPI_ISL_4205889 | | Clinical specimen | Centre de Recherche et de Formation en Infectiologie Guinée | Alpha Kabinet KEITA et al |
| **hCoV-19/Guinea/CERFIG-24665/2021** | EPI_ISL_4205890 | | Clinical specimen | Centre de Recherche et de Formation en Infectiologie Guinée | Alpha Kabinet KEITA et al |
| **hCoV-19/Uganda/UG481/2021** | EPI_ISL_2690458 | | MRC/UVRI & LSHTM Uganda Research Unit, Central Public Health Laboratories, Rakai Health Sciences Program | MRC/UVRI & LSHTM Uganda Research Unit, Central Public Health Laboratories | Matthew Cotten et al |
| **hCoV-19/Zhejiang/ZPCDCP-03/2021** | EPI_ISL_17261505 | | Zhejiang Provincial Center for Disease Control and Prevention | Zhejiang Provincial Center for Disease Control and Prevention | Yi Sun et al |
| **hCoV-19/Gabon/GG0216/2021** | EPI_ISL_11522604 | | Centre de Recherches Medicales de Lambarene | Department of Emerging Infectious Diseases, Institute of Tropical Medicine, Nagasaki University | Haruka Abe et al |
| **hCoV-19/Mongolia/UB-11361/2020** | EPI_ISL_9879583 | | National Center for Communicable Diseases (NCCD) National Influenza Center | National Center for Communicable Diseases (NCCD) National Influenza Center | Naranzul Tsedenbal et al |
| **hCoV-19/Madagascar/IPM-10090/2020** | EPI_ISL_3716411 | | Virology Unit, Institut Pasteur de Madagascar | Institut Pateur de Dakar | Norosoa Razanajatovo et al |
| **hCoV-19/Finland/19MR1cF7/2020** | EPI_ISL_757340 | | Department of Virology and Immunology, University of Helsinki and Helsinki University Hospital, Huslab Finland | Department of Virology, Faculty of Medicine, University of Helsinki, Helsinki, Finland | Teemu Smura et al |
| **hCoV-19/Madagascar/IPM-27446/2020** | EPI_ISL_3716570 | | Virology Unit, Institut Pasteur de Madagascar | Institut Pateur de Dakar | Norosoa Razanajatovo et al |

| | | | | | |
|---|---|---|---|---|---|
| **hCoV-19/Turkey/IST-GLAB-143/2020** | EPI_ISL_735280 | | Genomic Laboratory (GLAB) (Conjoint lab of Health Directorate of Istanbul and Istanbul Technical University) | Genomic Laboratory (GLAB), Istanbul Technical University | Ilker Karacan et al |
| **hCoV-19/India/TG-RFCH01806_CIB5537/2020** | EPI_ISL_7468209 | | CSIR-Centre for Cellular and Molecular Biology | CSIR-Centre for Cellular and Molecular Biology | Amareshwar Vodapalli et al |
| **hCoV-19/Mexico/SEARCH-100622/2021** | EPI_ISL_2231516 | | Centro de Diagnostico COVID-19 UABC Tijuana | Andersen lab at Scripps Research | SEARCH Alliance San Diego with Idanya Rubi Serafin Higuera et al |
| **hCoV-19/Guatemala/ASI2089/2021** | EPI_ISL_5926971 | | Hospital General San Juan de Dios | Asociación de Salud Integral / Clínica Familiar "Luis Ángel García" | Eduardo Arathon et al |
| **hCoV-19/Seychelles/G9497/2020** | EPI_ISL_4880612 | OK377421.1 | Seychelles Public Health Laboratory | KEMRI-Wellcome Trust Research Programme,Kilifi | Githinji G. et al |
| **hCoV-19/Cyprus/BMVUCY283/2020** | EPI_ISL_7210360 | | Famagusta General Hospital | Laboratory of Biotechnology and Molecular Virology, University of Cyprus | Leondios G. Kostrikis et al. |
| **hCoV-19/SouthKorea/KDCA0637/2020** | EPI_ISL_850319 | | Division of Emerging Infectious Diseases, Bureau of Infectious Diseases Diagnosis Control, Korea Disease Control and Prevention Agency | Division of Emerging Infectious Diseases, Bureau of Infectious Diseases Diagnosis Control, Korea Disease Control and Prevention Agency | Ae Kyung Park et al |
| **hCoV-19/CzechRepublic/IAB20_016_015/2020** | EPI_ISL_889345 | | The University Hospital Brno | Institute of Applied Biotechnologies a.s. | Petr Klempt et al |
| **hCoV-19/Vietnam/KH-55/2020** | EPI_ISL_17373280 | | CDC Khánh Hòa, Vietnam | Department of Microbiology and Immunology, Pasteur Institute in Nha Trang | Trinh Hoang Long et al |
| **hCoV-19/India/RJ-ICMR-NIV-INSACOG-SEQ-2070/2020** | EPI_ISL_15038527 | | ICMR-National Institute of Virology - INSACOG | NIV Influenza | Dr. Varsha Potdar et al |
| **hCoV-19/Mongolia/UB-NCCD-29341/2021** | EPI_ISL_18254358 | | National Center for Communicable Diseases | Laboratory of Molecular biology, Institute of Biomedical Sciences, Mongolian National University of Medical Sciences | Munkhtuya Erendereg et al |
| **hCoV-19/Mongolia/UB-NCCD-184401/2021** | EPI_ISL_18262179 | | National Center for Communicable Diseases | Laboratory of Molecular biology, Institute of Biomedical Sciences, Mongolian National University of Medical Sciences | Munkhtuya Erendereg et al |
| **hCoV-19/Mongolia/UB-NCCD-258612/2021** | EPI_ISL_18134639 | | National Center for Communicable Diseases | Laboratory of Molecular biology, Institute of Biomedical Sciences, Mongolian National University of Medical Sciences | Munkhtuya Erendereg et al |
| **hCoV-19/Singapore/1101/2020** | EPI_ISL_596491 | | National Public Health Laboratory, National Centre for Infectious Diseases | National Public Health Laboratory, National Centre for Infectious Diseases | Tze Minn Mak et al |
| **hCoV-19/India/WB-1931300201452/2021** | EPI_ISL_1419224 | | INSACOG-WB | National Institute of Biomedical Genomics – INSACOG | Arindam Maitra et al A |
| **hCoV-19/India/MZ-ZMC478J/2020** | EPI_ISL_1419067 | | INSACOG-Mizoram | National Institute of Biomedical Genomics – INSACOG | Arindam Maitra et al A |
| **hCoV-19/India/KL-GENESCOV00586/2020** | EPI_ISL_7214935 | | INSACOG-IGIB and Kerala State SARS-COV-2 Genome Surveillance Programme (GENESCOV2) | INSACOG and GENESCOV2 at CSIR Institute of Genomics and Integrative Biology | INSACOG-IGIB et al |
| **hCoV-19/India/un-IRSHA-CD210922/2021** | EPI_ISL_2714694 | MW969752.1 | Communicable Diseases, Interactive Research School for Health Affairs (IRSHA) | Communicable Diseases, Interactive Research School for Health Affairs (IRSHA) | Shrivastava et al |
| **hCoV-19/Malaysia/IMR_WI100/2020** | EPI_ISL_718269 | | Institute for Medical Research, Infectious Disease Research Centre, National Institutes of Health, Ministry of Health Malaysia | Institute for Medical Research, Infectious Disease Research Centre, National Institutes of Health, Ministry of Health Malaysia | Suppiah J et al |
| **hCoV-19/Brunei/7/2020** | EPI_ISL_962878 | | National Virology Reference Laboratory | National Public Health Laboratory, National Centre for Infectious Diseases | Tze Minn Mak et al |
| **hCoV-19/India/KL-GENESCOV02211/2021** | EPI_ISL_7217826 | | INSACOG-IGIB and Kerala State SARS-COV-2 Genome Surveillance Programme (GENESCOV2) | INSACOG and GENESCOV2 at CSIR Institute of Genomics and Integrative Biology | INSACOG-IGIB et al |
| **hCoV-19/India/TN-CLRI-CIC0644/2020** | EPI_ISL_9232385 | | CSIR-Central Leather Research Institute | CSIR-Centre for Cellular and Molecular Biology-INSACOG | Lamuk Zaveri et al |
| **hCoV-19/India/OR-ACTREC-688/2020** | EPI_ISL_700313 | | Hematopathology Laboratory, ACTREC, TMC | Hematopathology Laboratory, ACTREC, TMC | Hematopathology Laboratory et al |
| **hCoV-19/India/TN-ICMR-NIV-INSACOG-SEQ-4049/2020** | EPI_ISL_15038533 | | ICMR-National Institute of Virology - INSACOG | NIV Influenza | Dr. Varsha Potdar et al |
| **hCoV-19/India/OR-ILSGS00072/2020** | EPI_ISL_1063888 | | Immunogenomics lab, Institute of Life Sciences, Bhubaneswar | Immunogenomics lab, Institute of Life Sciences, Bhubaneswar | Sunil K. Raghav et al |
| **hCoV-19/India/PY-SEQ159_S28_R1_001/2021** | EPI_ISL_1939842 | | JIPMER Puducherry | inStem NCBS - INSACOG | Uma Ramakrishnan Dasaradhi Palakodeti Aswin SaiNarain et al |
| **hCoV-19/UnitedArabEmirates/4306/2020** | EPI_ISL_860066 | | BTC, Khalifa University | BTC, Khalifa University | Al Safar et al |
| **hCoV-19/env/Beijing/IVDC-03-06/2020** | EPI_ISL_469256 | | National Institute for Viral Disease Control and Prevention, China CDC | National Institute for Viral Disease Control and Prevention, China CDC | Xiang Zhao et al |
| **hCoV-19/Niger/CS656/2021** | EPI_ISL_2281768 | | Centre de Recherche Médicale et Sanitaire (CERMES) | Institut Pasteur de Dakar | Lagare Adamou et al |
| **hCoV-19/Ecuador/179320_S63/2021** | EPI_ISL_18283509 | | INSPI | INSPI - Charité | Alfredo Bruno Caicedo et al |
| **hCoV-19/Kyrgyzstan/ChVir-34188/2020** | EPI_ISL_6335246 | | Department of Disease Prevention and State Sanitary and Epidemiological Surveillance of the Ministry of Health of the Kyrgyz Republic | Charité Universitätsmedizin Berlin, Institut für Virologie | Victor M Corman et al |
| **hCoV-19/Romania/DJ14962/2020** | EPI_ISL_5917807 | | Medical Genetics Laboratory, Regional Centre of Medical Genetics, Emergency County Hospital Craiova | Medical Genetics Laboratory, Regional Centre of Medical Genetics, Emergency County Hospital Craiova | Anca-Lelia (Riza) Costache et al |
| **hCoV-19/Cyprus/BMVUCY304/2020** | EPI_ISL_7210347 | | Famagusta General Hospital | Laboratory of Biotechnology and Molecular Virology, University of Cyprus | Leondios G. Kostrikis et al. |
| **hCoV-19/Spain/GA-EPI-HULA-182/2020** | EPI_ISL_18630530 | | Servicio de Microbiología. Hospital Lucus Augusti | CHUVI / EPICOVIGAL Consortium | María del Pilar Gallego on behalf of EPICOVIGAL Consortium et al |
| **hCoV-19/Ecuador/69732_S38/2020** | EPI_ISL_18283254 | | INSPI | INSPI - Charité | Alfredo Bruno Caicedo et al |
| **hCoV-19/BurkinaFaso/2-041-1649/2020** | EPI_ISL_660492 | | Laboratoire de Microbiologie CHU Souro Sanou | Centre Muraz | Abdoul-Salam Ouedraogo et al |
| **hCoV-19/Russia/AST-RII-MH4459S/2020** | EPI_ISL_733200 | | Pathogenic Microorganisms Variability Laboratory | WHO National Influenza Centre Russian Federation | Andrey Komissarov et al |
| **hCoV-19/Jiangsu/WX-820/2020** | EPI_ISL_16138012 | | Center Laboratory of Health Quarantine, Nanjing Customs | Center Laboratory of Health Quarantine, Nanjing Customs | Geng et al |
| **hCoV-19/Lithuania/T_25/2020** | EPI_ISL_904944 | | Vilnius University Hospital Santaros Klinikos, Vilnius University | Institute of Biotechnology, Life Sciences Center, Vilnius University | Emilija Vasiliunaite et al |
| **hCoV-19/Kyrgyzstan/NRL-25169/2020** | EPI_ISL_13285368 | | Republican Reference Laboratory, National TB center, Kyrgyzstan | Republican Reference Laboratory, National TB center, Kyrgyzstan | Gulmira Kalmambetova et al |
| **hCoV-19/Kyrgyzstan/NRL-43587/2021** | EPI_ISL_13285376 | | Republican Reference Laboratory, National TB center, Kyrgyzstan | Republican Reference Laboratory, National TB center, Kyrgyzstan | Gulmira Kalmambetova et al |
| **hCoV-19/Urumqi/IVDC-01-01/2020** | EPI_ISL_850951 | | National Institute for Viral Disease Control and Prevention, China CDC | National Institute for Viral Disease Control and Prevention, China CDC | Xiang Zhao et al |
| **hCoV-19/Zhejiang/SX0715/2020** | EPI_ISL_497950 | | Shaoxing CDC | Zhejiang Provincial Center for Disease Control and Prevention | Yin Chen et al |
| **hCoV-19/Russia/SPE-CSP-922_S52/2020** | EPI_ISL_8904171 | | Chief Federal State Budgetary Healthcare Institution "Centre of Hygiene and Epidemiology" of the Federal Medical Biological Agency, Central laboratory of microbiological and clinical diagnostic tests | Federal State Budgetary Institution "Centre for Strategic Planning and Management of Biomedical Health Risks" of the Federal Medical Biological Agency, Laboratory for advanced molecular diagnostics of human diseases. | Mikhaylov et al |
| **hCoV-19/Latvia/1157/2020** | EPI_ISL_1312668 | | Centrālā Laboratorija | Latvian Biomedical Research and Study Centre | Janis Pjalkovskis et al |
| **hCoV-19/Tunisia/X6185/2020** | EPI_ISL_11172706 | | Laboratory of Clinical Virology, Institut Pasteur de Tunis | Laboratory of Clinical Virology, Institut Pasteur de Tunis | Wasfi Fares et al |
| **hCoV-19/Tunisia/Tunis-Y-1294/2020** | EPI_ISL_18516169 | | Laboratoire de Virologie Clinique, Institut Pasteur de Tunis | Laboratoire de Virologie Clinique, Institut Pasteur de Tunis | Haddad-Boubaker Sondes et al |
| **hCoV-19/Azerbaijan/un-ChVir24287_71/2020** | EPI_ISL_9214612 | | Ministry of Health of the Republic of Azerbaijan | Charité Universitätsmedizin Berlin, Institut für Virologie | Victor M Corman et al |
| **hCoV-19/Taiwan/06-109-064485/2020** | EPI_ISL_12665831 | | Virology Lab,Department of Pathology, National Cheng Kung University Hospital | Virology Lab,Department of Pathology, National Cheng Kung University Hospital | Huey-Pin Tsai et al |
| **hCoV-19/Armenia/IMB1-4/2021** | EPI_ISL_1139149 | | National Center of Disease Control and Prevention of the Republic of Armenia | Institute of Molecular Biology NAS RA, Republic of Armenia, Department of Bioengineering, BioinformaticsInstitute and Molecular Biology IBMPh RAU, Republic of Armenia | Arsen Arakelyan et al |
| **hCoV-19/Israel/CVL-928ngs/2020** | EPI_ISL_889141 | | Israel Central Virology laboratory | Israel National Consortium for SARS-CoV-2 sequencing | Neta Zuckerman et al |
| **hCoV-19/USA/AK-PHL6791/2021** | EPI_ISL_1049504 | MZ233464.1 | Alaska State Virology Laboratory | Alaska State Virology Laboratory | Stephanie DeRonde et al |
| **hCoV-19/Turkey/HSGM-B11546/2021** | EPI_ISL_2403282 | | Ministry of Health Turkey | Ministry of Health Turkey | Fatma Bayrakdar et al |
| **hCoV-19/Heilongjiang/IVDC-HEB-267/2021** | EPI_ISL_9910218 | | Heilongjiang Center for Disease Control and Prevention | National Institute for Viral Disease Control and Prevention, China CDC | Ming Yang et al |
| **hCoV-19/Heilongjiang/IVDC-SH/240/2021** | EPI_ISL_9910191 | | Heilongjiang Center for Disease Control and Prevention | National Institute for Viral Disease Control and Prevention, China CDC | Qi Liang et al |
| **hCoV-19/India/TG-CCMB-BM780/2021** | EPI_ISL_1838827 | | CSIR-Centre for Cellular and Molecular Biology | CSIR-Centre for Cellular and Molecular Biology-INSACOG | Payel Mukherjee et al B |
| **hCoV-19/Jordan/Biolab0202/2021** | EPI_ISL_5529836 | | Biolab Diagnostic Laboratories | Biolab Diagnostic Laboratories | Issa Abu-Dayyeh et al |
| **hCoV-19/India/RJ-CCMB-CIA6262/2021** | EPI_ISL_3453515 | | RUHS College of Medical Sciences | CSIR-Centre for Cellular and Molecular Biology - INSACOG | Dr. Gaurav Dalela et al |
| **hCoV-19/Congo/ITM-069-6277/2021** | EPI_ISL_1857281 | | Fondation Congolaise pour la recherche medicale (FCRM), Francine Ntoumi | Institute of Tropical Medicine | Prof. Francine Ntoumi et al |
| **hCoV-19/UnitedArabEmirates/BTC-5110/2021** | EPI_ISL_5142875 | | BTC, Khalifa University | BTC, Khalifa University | Habiba Al Safar et al |
| **hCoV-19/India/MP-NCDC-1009293/2021** | EPI_ISL_2459720 | | National Centre for Disease Control (NCDC) Biotechnology Division, Delhi | NCDC Delhi, Biotechnology Division INSACOG | Kalaiarasan Ponnusamy et al |
| **hCoV-19/Romania/BZ-15451/2021** | EPI_ISL_2566464 | | Department of Public Health Buzau | National Institute of Infectious Diseases-Prof. Dr. Matei Bals Molecular Diagnostics Laboratory | Corina Casangiu et al |
| **hCoV-19/England/CAMC-10C033D/2021** | EPI_ISL_908717 | OD948436.1 | Lighthouse Lab in Cambridge | Wellcome Sanger Institute for the COVID-19 Genomics UK (COG-UK) Consortium | Rob Howes et al |
| **hCoV-19/Greece/210187/2021** | EPI_ISL_2365272 | | Laboratory of Clinical Virology | Greek Genome Center, Biomedical Research Foundation of the Academy of Athens (BRFAA) | Emmanouil Athanasiadis et al |
| **hCoV-19/Albania/210901055/2020** | EPI_ISL_1299864 | | Unit of lab surveillance of viral emerging diseases, National Lab of Influenza | Respiratory Virus Unit, National Infection Service, Public Health England | PHE Covid Sequencing Team et al |
| **hCoV-19/Taiwan/NTU76/2021** | EPI_ISL_13618346 | | Department of Laboratory Medicine, National Taiwan University Hospital | Department of Clinical Laboratory Sciences and Medical Biotechnology, National Taiwan University | Shiou-Hwei Yeh et al |
| **hCoV-19/France/OCC-HCL021071180201/2021** | EPI_ISL_1934990 | | CHU TOULOUSE | CNR Virus des Infections Respiratoires - France SUD | Antonin Bal et al |
| **hCoV-19/Georgia/Tb-S354/2021** | EPI_ISL_3076935 | | Department for Virology, Molecular Biology and Genome Research, R. G. Lugar Center for Public Health Research, National Center for Disease Control and Public Health (NCDC) of Georgia. | Department for Virology, Molecular Biology and Genome Research, R. G. Lugar Center for Public Health Research, National Center for Disease Control and Public Health (NCDC) of Georgia. | Giorgi Tomashvili et al |
| **hCoV-19/India/ML-1727400000081/2021** | EPI_ISL_2547488 | | INSACOG-MEGHALAYA | National Institute of Biomedical Genomics – INSACOG | Arindam Maitra et al A |
| **hCoV-19/Kyrgyzstan/NRL-41589/2021** | EPI_ISL_13285375 | | Republican Reference Laboratory, National TB center, Kyrgyzstan | Republican Reference Laboratory, National TB center, Kyrgyzstan | Gulmira Kalmambetova et al |
| **hCoV-19/Ireland/D-NVRL-84IRL01786/2021** | EPI_ISL_959774 | | National Virus Reference Laboratory | National Virus Reference Laboratory | Michael Carr et al |
| **hCoV-19/Algeria/33008/2021** | EPI_ISL_12156751 | | NIC, Viral Respiratory Unit | NIC, Viral Respiratory Unit | Sarah Belkalem et al |
| **hCoV-19/Cyprus/BMV670/2021** | EPI_ISL_16373277 | | Department of Molecular Virology, Cyprus Institute of Neurology and Genetics | Laboratory of Biotechnology and Molecular Virology, University of Cyprus | Leondios G. Kostrikis et al. |
| **hCoV-19/Cyprus/BMV517/2021** | EPI_ISL_16368277 | | Nicosia General Hospital | Laboratory of Biotechnology and Molecular Virology, University of Cyprus | Leondios G. Kostrikis et al. |
| **hCoV-19/Luxembourg/LNS5741525/2021** | EPI_ISL_2400622 | | Laboratoires d'analyses medicales - Ketterthill | Laboratoire national de sante, Microbiology, Microbial Genomics Platform | Anke Wienecke-Baldacchino et al |
| **hCoV-19/India/TR-AGMC-R12953/2021** | EPI_ISL_2433272 | | INSACOG-Tripura | National Institute of Biomedical Genomics – INSACOG | Arindam Maitra et al A |
| **hCoV-19/Kuwait/KU-12093/2021** | EPI_ISL_2429126 | | Mubarak Al-Kabeer Hospital | Virology Unit, Department of Microbiology, Faculty of Medicine, Kuwait University | Nada Madi et al |
| **hCoV-19/Bahrain/631201947/2021** | EPI_ISL_14133008 | MW672567.1 | Communicable Disease Laboratory, Public Health Directorate | Communicable Disease Laboratory, Public Health Directorate | Alwasti et al |
| **hCoV-19/India/ML-1727400000071/2021** | EPI_ISL_2547478 | | INSACOG-MEGHALAYA | National Institute of Biomedical Genomics – INSACOG | Arindam Maitra et al A |
| **hCoV-19/India/UK-NCDC-160963/2021** | EPI_ISL_2505130 | | National Centre for Disease Control (NCDC) Biotechnology Division, Delhi | NCDC Delhi, Biotechnology Division INSACOG | Robin Marwal et al |
| **hCoV-19/Italy/CAM-TIGEM-IZSM-COLLI-5525/2021** | EPI_ISL_1307801 | | Lab. Microbiologia e Virologia - Cotugno - A.O. dei Colli | TIGEM | Antonio Grimaldi Patrizia Annunziata Francesco Panariello Biancamaria Pierri Claudia Tiberio Valentina Bouche Chiara Colantuono Maria Concetta Cuomo Denise Di Concilio Lucio Di Filippo Anna Manfredi Marcello Salvi Antonio Limone Luigi Atripaldi Pellegrino Cerino Andrea Ballabio Davide Cacchiarelli et al |
| **hCoV-19/Bahrain/341702536/2021** | EPI_ISL_1660461 | MW947232.1 | Communicable Disease Laboratory, Public Health Directorate | Communicable Disease Laboratory, Public Health Directorate | Alwasti et al |
| **hCoV-19/Moldova/un-ChVir25014/2021** | EPI_ISL_3446541 | | WHO/Country Office in the Republic of Moldova | Charité Universitätsmedizin Berlin, Institut für Virologie | Victor M Corman et al |
| **hCoV-19/Pakistan/NIH-688675X/2021** | EPI_ISL_1732274 | | Department of Virology | Department of Virology | Massab Umair et al |
| **hCoV-19/Israel/CVL-933ngs/2020** | EPI_ISL_889144 | | Israel Central Virology laboratory | Israel National Consortium for SARS-CoV-2 sequencing | Neta Zuckerman et al |
| **hCoV-19/Gabon/CERMEL-JJ0691/2021** | EPI_ISL_5031219 | | Centre de Recherches Médicales de Lambaréné | Centre de Recherches Médicales de Lambaréné | Moustapha Nzamba Maloum et al |
| **hCoV-19/Gibraltar/EXP_05092021-1235128-75/2021** | EPI_ISL_3989030 | | Gibraltar Health Authority Lab | Gibraltar Health Authority Covid-19 Laboratory | Dr Nicholas Cortes et al |
| **hCoV-19/Jordan/Biolab0264/2021** | EPI_ISL_6268678 | | Biolab Diagnostic Laboratories | Biolab Diagnostic Laboratories | Issa Abu-Dayyeh et al |
| **hCoV-19/Finland/THL-202112080/2021** | EPI_ISL_2507885 | | Islab, Etelä-Savon aluelaboratorio | Expert Microbiology, National Institute for Health and Welfare | Soile Blomqvist et al |
| **hCoV-19/Russia/MOW-CSP-LQ-1470_S32/2021** | EPI_ISL_18025621 | | Chief Federal State Budgetary Healthcare Institution "Centre of Hygiene and Epidemiology" of the Federal Medical Biological Agency, Central laboratory of microbiological and clinical diagnostic tests | Federal State Budgetary Institution "Centre for Strategic Planning and Management of Biomedical Health Risks" of the Federal Medical Biological Agency, Laboratory for advanced molecular diagnostics of human diseases | Mikhaylov et al |
| **hCoV-19/CentralAfricanRepublic/24264/2021** | EPI_ISL_2281246 | | Insitut Pasteur Bangui | Institut Pasteur de Dakar | Marie-Astrid Vernet et al |
| **hCoV-19/Uganda/UG453/2021** | EPI_ISL_2346419 | | MRC/UVRI & LSHTM Uganda Research Unit, Central Public Health Laboratories | MRC/UVRI & LSHTM Uganda Research Unit, Central Public Health Laboratories | Matthew Cotten et al |
| **hCoV-19/Armenia/IMB3-11/2021** | EPI_ISL_1718304 | | National Center of Disease Control and Prevention of the Republic of Armenia | Institute of Molecular Biology NAS RA, Republic of Armenia, Department of Bioengineering, BioinformaticsInstitute and Molecular Biology IBMPh RAU, Republic of Armenia | Arsen Arakelyan et al |
| **hCoV-19/Bahrain/341709279/2021** | EPI_ISL_1660463 | MW947234.1 | Communicable Disease Laboratory, Public Health Directorate | Communicable Disease Laboratory, Public Health Directorate | Alwasti et al |
| **hCoV-19/Palestine/AAS78/2021** | EPI_ISL_2107526 | MZ207874.1 | Medical Laboratory Sciences, Arab American University | Medical Laboratory Sciences, Arab American University | Al-Jawabreh et al |
| **hCoV-19/Libya/EMC-11/2021** | EPI_ISL_2860646 | | Public Health Reference Laboratory | Erasmus Medical Center | Omar Elahmer-Abdulla Bashein- Rahma Algheriani-Ahlam Alarif et al |
| **hCoV-19/Oman/rega-OM-106/2021** | EPI_ISL_4062899 | | Sultan Qaboos University Hospital, Department of Microbiology & Immunology, Molecular Biology Section | KU Leuven, Rega Institute, Clinical and Epidemiological Virology | Tony Wawina-Bokalanga et al |
| **hCoV-19/Netherlands/NB-MVD-CWGS2100316/2021** | EPI_ISL_1661273 | | Microvida | Microvida | Suzan D. Pas et al |
| **hCoV-19/India/SK-1122700010136/2021** | EPI_ISL_2151083 | | INSACOG-Sikkim | National Institute of Biomedical Genomics – INSACOG | Arindam Maitra et al A |
| **hCoV-19/Montenegro/CO-00357_MNE000_2204592102/2021** | EPI_ISL_3550685 | | Laboratory for Molecular Diagnostics, IPHMN | Eurofins Genomics Europe Sequencing GmbH | Danijela Vujošević et al |
| **hCoV-19/Israel/CVL-9030/2021** | EPI_ISL_2084059 | | Israel Central Virology laboratory | Israel National Consortium for SARS-CoV-2 sequencing | Neta Zuckerman et al |
| **hCoV-19/Palestine/AAS51/2021** | EPI_ISL_1502571 | MW858238.1 | Biochemistry and Molecular Biology Department-Faculty of Medicine, Al-Quds University | Biochemistry and Molecular Biology Department-Faculty of Medicine, Al-Quds University | Nasereddin et al |
| **hCoV-19/Jordan/UK6/2020** | EPI_ISL_904016 | | Molecular Biology and Virology lab, Faculty of Veterinary Medicine, Jordan University of Science and Technology | Molecular Biology and Virology lab, Faculty of Veterinary Medicine, Jordan University of Science and Technology | Mohammad Hussien Alboom et al |
| **hCoV-19/Portugal/PT4048/2021** | EPI_ISL_1172008 | | H Divino Espirito Santo - Ponta Delgada | Instituto Nacional de Saude (INSA) | Borges et al |
| **hCoV-19/USA/PA-CDC-QDX26343982/2021** | EPI_ISL_2929480 | MZ716322.1 | Quest Diagnostics Incorporated | Centers for Disease Control and Prevention Division of Viral Diseases, Pathogen Discovery | Dakota Howard et al |
| **hCoV-19/Senegal/192720/2021** | EPI_ISL_14598630 | | Institut Pasteur de Dakar | Institut Pasteur de Dakar | Diallo Amadou et al |
| **hCoV-19/France/OCC-CHU-TLS-210771381601/2021** | EPI_ISL_2366696 | | CHU Purpan - Laboratoire de Virologie - Institut Fédératif de Biologie | CHU Purpan - Laboratoire de Virologie - Institut Fédératif de Biologie | Latour J. et al |
| **hCoV-19/Norway/14989/2021** | EPI_ISL_2927881 | | Department of Medical Microbiology, St. Olavs hospital | Norwegian Institute of Public Health, Department of Virology | Kathrine Stene-Johansen et al |
| **hCoV-19/Norway/OUS-05765/2021** | EPI_ISL_4118555 | | Oslo University Hospital, Department of Microbiology | Norwegian Institute of Public Health, Department of Virology | Mona Holberg-Petersen et al |
| **hCoV-19/CaboVerde/Praia_4732/2021** | EPI_ISL_1123282 | | Virology Laboratory of Praia | Institut Pasteur de Dakar | Kevin Sanders et al |
| **hCoV-19/CaboVerde/Praia_4731/2021** | EPI_ISL_1711646 | | Virology Laboratory of Praia | Institut Pasteur de Dakar | Sanders Kevin et al |
| **hCoV-19/Iraq/KR-SEARCH-119952/2021** | EPI_ISL_12277936 | | University of Duhok COVID-19 Center for Research and Diagnosis | Andersen lab at Scripps Research | SEARCH Alliance with Jassim Mohammed Abdo et al |
| **hCoV-19/GuineaBissau/JPLNSP0035/2021** | EPI_ISL_2110853 | | Reitor da Universidade Jean Piaget Guine-Bissau | MRCG at LSHTM, Genomics lab | Aladje Balde et al |
| **hCoV-19/Estonia/Cov11286/2021** | EPI_ISL_2643049 | OU278143.1 | SYNLAB Eesti OÜ | Department of Microbiology, Institute of Biomedicine and Translational Medicine, University of Tartu | Radko Avi et al |
| **hCoV-19/Estonia/16043025/2021** | EPI_ISL_2601439 | | Synlab Eesti OÜ | 1. Laboratory of Communicable Diseases (Estonia); 2. Eurofins Genomics Europe Sequencing GmbH | Liidia Dotsenko et al. |
| **hCoV-19/Iran/GRC-TI-01-132-Tehran-IRAN/2021** | EPI_ISL_6258790 | | Molecular diagnostics laboratory, West Tehran Health Center, Health Deputy of Iran University of Medical Sciences, Tehran, Iran | Genetics Research Center, University of Social Welfare and Rehabilitation Sciences | Zohreh Fattahi et al |
| **hCoV-19/USA/CA-CDC-IBX382653181106/2021** | EPI_ISL_2089299 | MZ193151.1 | Infinity Biologix | Centers for Disease Control and Prevention Division of Viral Diseases, Pathogen Discovery | Dakota Howard et al |
| **hCoV-19/Belgium/UMONS-P02-04519-01/2021** | EPI_ISL_2687967 | | National Platform bis UMONS/Jolimont | National Platform bis UMONS/Jolimont | François Dufrasne et al |
| **hCoV-19/Norway/13383/2021** | EPI_ISL_2491511 | | Ostfold Hospital Trust - Kalnes, Centre for Laboratory Medicine, Section for gene technology and infection serology | Norwegian Institute of Public Health, Department of Virology | Kathrine Stene-Johansen et al |
| **hCoV-19/SaintLucia/LCA47987/2020** | EPI_ISL_872196 | | Caribbean Public Health Agency | Carrington Lab, Department of PreClinical Sciences, Faculty of Medical Sciences, The University of the West Indies | Nikita S. D. Sahadeo et al |
| **hCoV-19/Myanmar/DSMRC014/2021** | EPI_ISL_2595725 | MZ397174.1 | DSMRC | DSMRC | Khine Zaw Oo et al |
| **hCoV-19/Romania/MS-17854/2021** | EPI_ISL_3259553 | | Emergency County Hospital Targu Mures | National Institute of Infectious Diseases-Prof. Dr. Matei Bals Molecular Diagnostics Laboratory | Corina Casangiu et al |
| **hCoV-19/Beijing/IVDC-04/2021** | EPI_ISL_1121993 | | Emergency department of COVID-19, Beijing Ditan Hospital | National Institute for Viral Disease Control and Prevention | Yang Song2* et al |
| **hCoV-19/Djibouti/NAMRU3_D364/2021** | EPI_ISL_2858657 | MZ520059.1 | Naval Medical Research Unit No. 3 | Naval Medical Research Center Biological Defense Research Directorate | Catherine E. Arnold et al |
| **hCoV-19/Somalia/NAMRU3-Mog-10/2021** | EPI_ISL_2533882 | MZ394629.1 | Naval Medical Research Unit No. 3 | Naval Medical Research Center Biological Defense Research Directorate | Catherine E. Arnold et al |
| **hCoV-19/NorthMacedonia/27591/2021** | EPI_ISL_1669996 | | Laboratory of virology and molecular diagnostics, Institute of Public Health | Laboratory of virology and molecular diagnostics, Institute of Public Health | Kuzmanovska M et al |
| **hCoV-19/Iraq/DSeq-C14/2021** | EPI_ISL_4574544 | | Shar Hospital | DeepSeq Nottingham | Paywast Jamal Jalal et al |
| **hCoV-19/Norway/OUS-09050/2021** | EPI_ISL_4122731 | | Oslo University Hospital, Department of Microbiology | Norwegian Institute of Public Health, Department of Virology | Mona Holberg-Petersen et al |
| **hCoV-19/Croatia/5118/2021** | EPI_ISL_2674756 | | Hrvatski zavod za javno zdravstvo | Hrvatski zavod za javno zdravstvo | Irena Tabain et al |
| **hCoV-19/NorthMacedonia/41667/2021** | EPI_ISL_3545683 | | Laboratory of virology and molecular diagnostics, Institute of Public Health | Laboratory of virology and molecular diagnostics, Institute of Public Health | Kuzmanovska M et al |
| **hCoV-19/Ghana/WACCBIP-GS203/2021** | EPI_ISL_1255132 | | West African Centre for Cell Biology of Infectious Pathogens (WACCBIP), University of Ghana, Accra, Ghana | West African Centre for Cell Biology of Infectious Pathogens (WACCBIP), University of Ghana, Volta Road, Legon-Accra, Ghana | Collins M. Morang'a et al |
| **hCoV-19/Iran/GRC-T-I-11-117/2021** | EPI_ISL_2795678 | | Iran University of Medical Sciences, Tehran, Iran | Genetics Research Center, University of Social Welfare and Rehabilitation Sciences | Zohreh Fattahi et al |
| **hCoV-19/Lithuania/LTU000_INMEDICA_447860/2021** | EPI_ISL_2615054 | | UAB InMedica | National Public Health Surveillance Laboratory | Lukas Zemaitis et al |
| **hCoV-19/Lithuania/S21D1179/2021** | EPI_ISL_1756804 | | UAB Medicina Practica Laboratory | Vilnius University Hospital Santaros Klinikos, Center of Laboratory Medicine | Dovile Ezerskyte et al |
| **hCoV-19/Afghanistan/2870x0145_23822/2021** | EPI_ISL_4572796 | | Afghanistan | WRAIR | Irina Maljkovic Berry et al |
| **hCoV-19/CzechRepublic/UMTM218816/2021** | EPI_ISL_2403226 | | Institute of Molecular and Translational Medicine / Laboratory of Experimental Medicine, Faculty of Medicine and Dentistry, Palacky University and University Hospital Olomouc | Institute of Molecular and Translational Medicine / Laboratory of Experimental Medicine, Faculty of Medicine and Dentistry, Palacky University | Rastislav Slavkovský et al |
| **hCoV-19/NorthMacedonia/39285/2021** | EPI_ISL_2839459 | | General Hospital - Kumanovo | Research Center for Genetic Engineering and Biotechnology "Georgi D. Efremov" , Macedonian Academy of Sciences and Arts | Aleksandar J. Dimovski et al |
| **hCoV-19/Croatia/5635/2021** | EPI_ISL_2673970 | | Zavod za javno zdravstvo Zagrebačke Županije | Hrvatski zavod za javno zdravstvo | Irena Tabain et al |
| **hCoV-19/Maldives/MAV16830/2021** | EPI_ISL_3452729 | | Indira Gandhi Memorial Hospital | Indira Gandhi Memorial Hospital | Dr. Milza Abdul Muhsin et al |
| **hCoV-19/Ecuador/USFQ-1816/2021** | EPI_ISL_3506145 | | Institute of Microbiology, Universidad San Francisco de Quito | Institute of Microbiology, Universidad San Francisco de Quito | Fernanda Zurita et al |
| **hCoV-19/Ecuador/NIC-INSPI-338157/2021** | EPI_ISL_3068184 | | INSPI-CRN DE INFLUENZA Y OTROS VIRUS RESPIRATORIOS | NIC-INSPI | Alfredo Bruno et al |
| **hCoV-19/Guinea/CERFIG-19469/2021** | EPI_ISL_4205876 | | Clinical specimen | Centre de Recherche et de Formation en Infectiologie Guinée | Alpha Kabinet KEITA et al |
| **hCoV-19/Italy/CAM-AMES_34_38/2021** | EPI_ISL_1229683 | | ASL Napoli 1 Centro | AMES Centro Polidiagnostico Strumentale S.r.l. | "Giovanni Savarese et al |
| **hCoV-19/Albania/211060667/2021** | EPI_ISL_1299884 | | Unit of lab surveillance of viral emerging diseases, National Lab of Influenza | Respiratory Virus Unit, National Infection Service, Public Health England | PHE Covid Sequencing Team et al |
| **hCoV-19/Japan/PG-42365/2021** | EPI_ISL_2336544 | | SARS-CoV-2 testing team, National Institute of Infectious Diseases | Pathogen Genomics Center, National Institute of Infectious Diseases | Tsuyoshi Sekizuka et al B |
| **hCoV-19/Japan/PG-47263/2021** | EPI_ISL_2810902 | | Department of Infectious Diseases, Kobe Institute of Health | Department of Infectious Diseases, Kobe Institute of Health | Ryohei Nomoto et al |
| **hCoV-19/Japan/PG-44143/2021** | EPI_ISL_2337875 | | SARS-CoV-2 testing team, National Institute of Infectious Diseases | Pathogen Genomics Center, National Institute of Infectious Diseases | Tsuyoshi Sekizuka et al B |
| **hCoV-19/Japan/PG-70905/2021** | EPI_ISL_3190519 | | Pathogen Genomics Center, National Institute of Infectious Diseases | Pathogen Genomics Center, National Institute of Infectious Diseases | Tsuyoshi Sekizuka et al B |
| **hCoV-19/Moldova/un-ChVir25023/2021** | EPI_ISL_3446544 | | WHO/Country Office in the Republic of Moldova | Charité Universitätsmedizin Berlin, Institut für Virologie | Victor M Corman et al |
| **hCoV-19/SouthAfrica/NICD-DNC12671/2021** | EPI_ISL_2013049 | | Lancet Laboratories | National Institute for Communicable Diseases of the National Health Laboratory Service | Amoako DG et al |
| **hCoV-19/Vietnam/HNa_Alpha_610/2021** | EPI_ISL_17016448 | | National Hospital for Tropical Diseases | Oxford University Clinical Research Unit, Hanoi, Vietnam | Pham Ngoc Thach et al |
| **hCoV-19/Sweden/8542520450/2021** | EPI_ISL_2794994 | | The Public Health Agency of Sweden | The Public Health Agency of Sweden | Maximilian Riess et al |
| **hCoV-19/Finland/7291/2021** | EPI_ISL_3014552 | | Department of Virology and Immunology, University of Helsinki and Helsinki University Hospital, Huslab Finland | Department of Virology, Faculty of Medicine, University of Helsinki, Helsinki, Finland | Teemu Smura et al |
| **hCoV-19/India/HR-IGIB-2_117938/2021** | EPI_ISL_2880749 | | INSACOG Surveillance | INSACOG at CSIR Institute of Genomics and Integrative Biology | INSACOG et al |
| **hCoV-19/Spain/HUB00720/2021** | EPI_ISL_2352001 | | Hospital Universitari Bellvitge | Microbiology Department | Sara Marti et al |
| **hCoV-19/Spain/HUD-79232065/2021** | EPI_ISL_2228077 | | Microbiology Department, University Hospital Donostia | Microbiology Department, University Hospital Donostia | Marimon JM et al |
| **hCoV-19/Latvia/2105069019/2021** | EPI_ISL_2798814 | | Centrālā Laboratorija | Riga East Clinical University Hospital, National Microbiology Reference Laboratory | Reinis Vangravs et al |
| **hCoV-19/Curacao/CW-RIVM-27028/2021** | EPI_ISL_1792381 | | National Institute for Public Health and the Environment (RIVM) | National Institute for Public Health and the Environment (RIVM) | Adam Meijer et al |
| **hCoV-19/Brazil/RS-FIOCRUZ-14218/2021** | EPI_ISL_2661750 | | Laboratorio Central de Saude Publica do Estado do Rio Grande do Sul (LACEN-RS) | Laboratory of Respiratory Viruses and Measles, Oswaldo Cruz Institute, FIOCRUZ | Paola Resende et al |

| | | | | | |
|---|---|---|---|---|---|
| **hCoV-19/Slovenia/08-103548-MB/2021** | EPI_ISL_2001731 | | National Laboratory for Health, Environment and Food, OMM, Maribor | NLZOH (National Laboratory for Health, Environment and Food) / CISLD (Clinical Institute of Special Laboratory Diagnostics), University Children's Hospital, University Medical Center Ljubljana | Sandra Janezic et al |
| **hCoV-19/Jordan/Biolab0140/2021** | EPI_ISL_4053487 | | Biolab Diagnostic Laboratories | Biolab Diagnostic Laboratories | Issa Abu-Dayyeh et al |
| **hCoV-19/Kuwait/JAH2851/2021** | EPI_ISL_2673282 | | MOH - Jaber Al-Ahmad Hospital (Innovation Research Laboratory) | MOH - Jaber Al-Ahmad Hospital (Innovation Research Laboratory) | Salman Al-Sabah et al |
| **hCoV-19/Kuwait/JAH2882/2021** | EPI_ISL_2672785 | | MOH - Jaber Al-Ahmad Hospital (Innovation Research Laboratory) | MOH - Jaber Al-Ahmad Hospital (Innovation Research Laboratory) | Salman Al-Sabah et al |
| **hCoV-19/Netherlands/ZH-RIVM-33313/2021** | EPI_ISL_2303466 | | National Institute for Public Health and the Environment (RIVM) | National Institute for Public Health and the Environment (RIVM) | Adam Meijer et al |
| **hCoV-19/Kenya/C79725/2021** | EPI_ISL_1039228 | OL914612.1 | KEMRI-Wellcome Trust Research Programme/KEMRI-CGMR-C Kilifi | KEMRI-Wellcome Trust Research Programme/KEMRI-CGMR-C Kilifi | Githinji et al |
| **hCoV-19/Bonaire/BQ-RIVM-35129/2021** | EPI_ISL_2405585 | | National Institute for Public Health and the Environment (RIVM) | National Institute for Public Health and the Environment (RIVM) | Adam Meijer et al |
| **hCoV-19/Bonaire/BQ-RIVM-42379/2021** | EPI_ISL_2981899 | | National Institute for Public Health and the Environment (RIVM) | National Institute for Public Health and the Environment (RIVM) | Adam Meijer et al |
| **hCoV-19/Curacao/CW-RIVM-16507/2021** | EPI_ISL_1288141 | | National Institute for Public Health and the Environment (RIVM) | National Institute for Public Health and the Environment (RIVM) | Adam Meijer et al |
| **hCoV-19/Curacao/CW-RIVM-23348/2021** | EPI_ISL_1596229 | | National Institute for Public Health and the Environment (RIVM) | National Institute for Public Health and the Environment (RIVM) | Adam Meijer et al |
| **hCoV-19/Coted-Ivoire/BKE-1880-bc28/2021** | EPI_ISL_1365024 | | Molecular diagnostic unit for viral haemorrhagic fevers and emerging viruses, Bouaké CHU Laboratory | Molecular diagnostic unit for viral haemorrhagic fevers and emerging viruses, Bouaké CHU Laboratory | Chantal Akoua-Koffi et al |
| **hCoV-19/Curacao/CW-RIVM-16333/2021** | EPI_ISL_1233007 | | National Institute for Public Health and the Environment (RIVM) | National Institute for Public Health and the Environment (RIVM) | Adam Meijer et al |
| **hCoV-19/Bonaire/BQ-RIVM-27386/2021** | EPI_ISL_1792360 | | National Institute for Public Health and the Environment (RIVM) | National Institute for Public Health and the Environment (RIVM) | Adam Meijer et al |
| **hCoV-19/Aruba/AW-RIVM-40490/2021** | EPI_ISL_2673215 | | National Institute for Public Health and the Environment (RIVM) | National Institute for Public Health and the Environment (RIVM) | Adam Meijer et al |
| **hCoV-19/Curacao/CW-RIVM-35048/2021** | EPI_ISL_2405504 | | National Institute for Public Health and the Environment (RIVM) | National Institute for Public Health and the Environment (RIVM) | Adam Meijer et al |
| **hCoV-19/Estonia/Cov11331/2021** | EPI_ISL_2643012 | OU370788.1 | SYNLAB Eesti OÜ | Department of Microbiology, Institute of Biomedicine and Translational Medicine, University of Tartu | Radko Avi et al |
| **hCoV-19/Morocco/INH-480/2021** | EPI_ISL_18263512 | | NIC at National Institute of Hygiene (NIH) | National Institute of Hygiene | Hassan Ihazmade et al |
| **hCoV-19/Coted-Ivoire/COV-BKE2273-bc25/2021** | EPI_ISL_1662501 | | Molecular diagnostic unit for viral haemorrhagic fevers and emerging viruses, Bouaké CHU Laboratory | Molecular diagnostic unit for viral haemorrhagic fevers and emerging viruses, Bouaké CHU Laboratory | Chantal Akoua-Koffi et al |
| **hCoV-19/USA/NV-CDC-LC0044002/2021** | EPI_ISL_1684845 | MZ039474.1 | Laboratory Corporation of America | Centers for Disease Control and Prevention Division of Viral Diseases, Pathogen Discovery | Dakota Howard et al |
| **hCoV-19/Togo/C6959/2021** | EPI_ISL_1524720 | | Laboratoire Biolim/FSS/UL | Unité Mixte Internationale TransVIHMI (UMI 233 IRD – U1175 INSERM - Université de Montpellier) IRD (Institut de recherche pour le développement) | Mounerou SALOU et al |
| **hCoV-19/USA/IN-GD-SID-C21165026/2021** | EPI_ISL_2696528 | MZ405237.1 | Laboratory, Gravity Diagnostics | Laboratory, Gravity Diagnostics | Diagnostics et al |
| **hCoV-19/USA/MI-CDC-QDX26076106/2021** | EPI_ISL_2868962 | MZ530173.1 | Quest Diagnostics Incorporated | Centers for Disease Control and Prevention Division of Viral Diseases, Pathogen Discovery | Dakota Howard et al |
| **hCoV-19/Nigeria/BCVL-21978/2021** | EPI_ISL_4026351 | | Biorepository and Clinical Virology Laboratory | Northwestern University - Center for Pathogen Genomics and Microbial Evolution | Ramon Lorenzo-Redondo et al |
| **hCoV-19/SaoTomeandPrincipe/06/2021** | EPI_ISL_10322959 | | LNR-TB | LNR-TB | Rosa Neto et al |
| **hCoV-19/Dominica/65658/2021** | EPI_ISL_2631439 | | The Caribbean Public Health Agency | Carrington Lab, Department of PreClinical Sciences, Faculty of Medical Sciences, The University of the West Indies, St Augustine Campus | Nikita S. D. Sahadeo et al |
| **hCoV-19/Kenya/C89001/2021** | EPI_ISL_2602496 | OL914811.1 | KEMRI-Wellcome Trust Research Programme,Kilifi | KEMRI-Wellcome Trust Research Programme,Kilifi | Githinji G. et al |
| **hCoV-19/Oman/182621/2020** | EPI_ISL_766569 | | Oman-National Influenza Center | Oman-National Influenza Center | Samiha Al-Kharusi et al |
| **hCoV-19/Colombia/ANT-UdeA-21001965v/2021** | EPI_ISL_4549188 | | Grupo Inmunovirología, Universidad de Antioquia | Grupo Inmunovirología, Universidad de Antioquia | Wbeimar Aguilar-Jimenez et al |
| **hCoV-19/Bulgaria/21BG-EU_000173_PI2/2021** | EPI_ISL_1233613 | | National Center for Infectious and Parasitic Diseases (NCIPD) | National Center for Infectious and Parasitic Diseases (NCIPD) | Alexiev et al |
| **hCoV-19/India/HR-IGIB-9_118052/2021** | EPI_ISL_2880907 | | INSACOG Surveillance | INSACOG at CSIR Institute of Genomics and Integrative Biology | INSACOG et al |
| **hCoV-19/Australia/NSW1545/2021** | EPI_ISL_1904452 | | South Eastern Area Laboratory Services (SEALS) | NSW Health Pathology - Institute of Clinical Pathology and Medical Research; Westmead Hospital; University of Sydney | CIDM-PH et al. |
| **hCoV-19/India/HP-NCDC-160946/2021** | EPI_ISL_2505124 | | National Centre for Disease Control (NCDC) Biotechnology Division, Delhi | NCDC Delhi, Biotechnology Division INSACOG | Robin Marwal et al |
| **hCoV-19/India/PB-NCDC-1409157/2021** | EPI_ISL_2460353 | | National Centre for Disease Control (NCDC) Biotechnology Division, Delhi | NCDC Delhi, Biotechnology Division INSACOG | Manoj K Singh et al |
| **hCoV-19/India/PB-NCDC-1409188/2021** | EPI_ISL_2460358 | | National Centre for Disease Control (NCDC) Biotechnology Division, Delhi | NCDC Delhi, Biotechnology Division INSACOG | Manoj K Singh et al |
| **hCoV-19/Canada/UN-24124/2021** | EPI_ISL_12387214 | | Switch Health | Public Health Agency of Canada (PHAC) National Microbiology Laboratory | Anna Majer et al |
| **hCoV-19/BosniaandHerzegovina/VFS-UNSA-LMGFI024/2020** | EPI_ISL_1016690 | | University of Sarajevo, Veterinary Faculty, Laboratory for Molecular Diagnostic and Research Laboratory | University of Sarajevo, Veterinary Faculty, Laboratory for Molecular Diagnostic and Research Laboratory | Goletic T. et al |
| **hCoV-19/Bulgaria/21BG-EU_002124_PI20/2021** | EPI_ISL_2180080 | | National Center of Infectious and Parasitic Diseases | National Center of Infectious and Parasitic Diseases | Alexiev et al |
| **hCoV-19/Belize/CML-124/2021** | EPI_ISL_4296428 | | Central Medical Laboratory | Baylor College of Medicine | Alexander Kneubehl et al |
| **hCoV-19/Slovakia/UKBA-708/2021** | EPI_ISL_875527 | | Institute of Virology, Biomedical Research Center of the Slovak Academy of Sciences, Bratislava | Faculty of Natural Sciences, Comenius University, Bratislava | Broňa Brejová et al |
| **hCoV-19/CzechRepublic/UMTM_187890/2021** | EPI_ISL_18241839 | | Institute of Molecular and Translational Medicine / Laboratory of Experimental Medicine, Faculty of Medicine and Dentistry, Palacky University | Institute of Molecular and Translational Medicine / Laboratory of Experimental Medicine, Faculty of Medicine and Dentistry, Palacky University | Rastislav Slavkovský et al |
| **hCoV-19/Austria/CeMM4949/2021** | EPI_ISL_2324175 | | Klinikum Wels-Grieskirchen | Bergthaler laboratory, CeMM Research Center for Molecular Medicine of the Austrian Academy of Sciences | Lukas Endler et al |
| **hCoV-19/Australia/VIC18220/2021** | EPI_ISL_1033148 | | Microbiological Diagnostic Unit - Public Health Laboratory (MDU-PHL) | MDU-PHL | Seemann T. et al |
| **hCoV-19/Germany/BW-RKI-I-151173/2021** | EPI_ISL_2262830 | | SYNLAB MVZ Leinfelden-Echterdingen | Robert Koch Institute | ? |
| **hCoV-19/Germany/SN-RKI-I-176330/2021** | EPI_ISL_2632950 | | Medizinisches Labor Ostsachsen MVZ GbR Nebenbetriebsstätte Dresden | Robert Koch Institute | ? |
| **hCoV-19/Jiangsu/JS03/2021** | EPI_ISL_16405160 | | Jiangsu Provincial Center for Disease Control and Prevention | Jiangsu Provincial Center for Disease Control and Prevention | Lunbiao Cui et al |
| **hCoV-19/Spain/AN-CBA-02887/2021** | EPI_ISL_15643667 | | Servicio de Microbiologia, Hospital Universitario Virgen del Rocio, Sevilla, Spain | Clinical Bioinformatics Area, Fundación Progreso y Salud | María Lara on behalf of SARS-CoV-2 whole genome sequencing circuit of Andalusia et al |
| **hCoV-19/Ireland/WH-NVRL-30IRL74713/2021** | EPI_ISL_2533704 | | National Virus Reference Laboratory | National Virus Reference Laboratory | Zoe Yandle et al |
| **hCoV-19/CzechRepublic/NRL_8204/2021** | EPI_ISL_2726979 | | National Institute of Public Health | National Institute of Public Health | Helena Jirincova et al |
| **hCoV-19/Lebanon/QIB-88/2021** | EPI_ISL_3343310 | OU600507.1 | National Influenza Center | Quadram Institute Bioscience | Georgi Merhi et al |
| **hCoV-19/Lebanon/QIB-103/2021** | EPI_ISL_3343315 | OU600497.1 | National Influenza Center | Quadram Institute Bioscience | Georgi Merhi et al |
| **hCoV-19/Lebanon/QIB-779/2021** | EPI_ISL_3343959 | OU599540.1 | National Influenza Center | Quadram Institute Bioscience | Georgi Merhi et al |
| **hCoV-19/Luxembourg/LNS9620023/2021** | EPI_ISL_3334791 | | Laboratoires Reunis, 38 Rue Hiel, 6131 Junglinster, Luxembourg | Laboratoire national de sante, Microbiology, Microbial Genomics Platform | Anke Wienecke-Baldacchino et al |
| **hCoV-19/Denmark/DCGC-112200/2021** | EPI_ISL_2666463 | | Department of Bacteria, Parasites and Fungi, Statens Serum Institut, Copenhagen, Denmark | Statens Serum Institut Bioinformatics and Microbial Genomics | Povilas Matusevicius on behalf of the Danish Covid-19 Genome Consortium et al |
| **hCoV-19/Lebanon/QIB-843/2021** | EPI_ISL_3344017 | OU597191.1 | National Influenza Center | Quadram Institute Bioscience | Georgi Merhi et al |
| **hCoV-19/Jordan/Biolab0303/2021** | EPI_ISL_6268705 | | Biolab Diagnostic Laboratories | Biolab Diagnostic Laboratories | Issa Abu-Dayyeh et al |
| **hCoV-19/Bahrain/341726177/2021** | EPI_ISL_1660467 | MW947244.1 | Communicable Disease Laboratory, Public Health Directorate | Communicable Disease Laboratory, Public Health Directorate | Alwasti et al |
| **hCoV-19/HongKong/CM21000264-2/2021** | EPI_ISL_5937310 | | Temporary Specimen Collection Centre at the AsiaWorld-Expo | Hong Kong Department of Health | Alan K.L. Tsang et al |
| **hCoV-19/SouthKorea/KDCA4285/2021** | EPI_ISL_2332557 | | Division of Emerging Infectious Diseases, Bureau of Infectious Diseases Diagnosis Control, Korea Disease Control and Prevention Agency | Division of Emerging Infectious Diseases, Bureau of Infectious Diseases Diagnosis Control, Korea Disease Control and Prevention Agency | Ae Kyung Park et al |
| **hCoV-19/Oman/rega-OM-56/2021** | EPI_ISL_4062616 | | Sultan Qaboos University Hospital, Department of Microbiology & Immunology, Molecular Biology Section | KU Leuven, Rega Institute, Clinical and Epidemiological Virology | Tony Wawina-Bokalanga et al |
| **hCoV-19/Oman/rega-OM-66/2021** | EPI_ISL_4062626 | | Sultan Qaboos University Hospital, Department of Microbiology & Immunology, Molecular Biology Section | KU Leuven, Rega Institute, Clinical and Epidemiological Virology | Tony Wawina-Bokalanga et al |
| **hCoV-19/Kuwait/JAH3101604/2021** | EPI_ISL_2556117 | | MOH - Jaber Al-Ahmad Hospital (Innovation Research Laboratory) | MOH - Jaber Al-Ahmad Hospital (Innovation Research Laboratory) | Salman Al-Sabah et al |
| **hCoV-19/Cambodia/431455/2021** | EPI_ISL_1969687 | | Virology Unit, Institut Pasteur du Cambodge | Virology Unit, Institut Pasteur du Cambodge | Jurre Y Siegers et al |
| **hCoV-19/Cambodia/01-2105201900/2021** | EPI_ISL_2343221 | | Banteay Meanchey Rapid Response Team | Virology Unit, Institut Pasteur du Cambodge | Jurre Y Siegers et al |
| **hCoV-19/Azerbaijan/Aghayev-10/2021** | EPI_ISL_2161031 | | National Hematology and Transfusiology Center | National Hematology and Transfusiology Center | Aghayev AR et al |
| **hCoV-19/India/PY-SEQ165_S34_R1_001/2021** | EPI_ISL_1939848 | | JIPMER Puducherry | inStem NCBS - INSACOG | Uma Ramakrishnan Dasaradhi Palakodeti Aswin SaiNarain et al |
| **hCoV-19/Gibraltar/211000769/2021** | EPI_ISL_1299843 | | Microbiology, Department of Pathology, St. Bernard's Hospital, Gibraltar Health Authority | Respiratory Virus Unit, National Infection Service, Public Health England | PHE Covid Sequencing Team et al |
| **hCoV-19/India/HR-IGIB-27_A_H640/2021** | EPI_ISL_2880738 | | INSACOG Surveillance | INSACOG at CSIR Institute of Genomics and Integrative Biology | INSACOG et al |
| **hCoV-19/BosniaandHerzegovina/AGCOB4239/2021** | EPI_ISL_2029122 | | General Hospital "Abdulah Nakas" | Alea Genetic Centre | Rijad Konjhodzic et al |
| **hCoV-19/Germany/MV-RKI-I-138373/2021** | EPI_ISL_2123150 | | Centogene; Dr. Bauer Laboratoriums GmbH | Robert Koch Institute | ? |
| **hCoV-19/Sweden/20249670P/2021** | EPI_ISL_2794568 | | The Public Health Agency of Sweden | The Public Health Agency of Sweden | Maximilian Riess et al |
| **hCoV-19/SriLanka/CMC118026/2021** | EPI_ISL_2803295 | | Centre for Dengue Research and AICBU, Department of Immunology and Molecular Medicine | Centre for Dengue Research and AICBU, Department of Immunology and Molecular Medicine | Chandima Jeewandara et al |
| **hCoV-19/Shulan/IVDC-01-01/2020** | EPI_ISL_850947 | | National Institute for Viral Disease Control and Prevention, China CDC | National Institute for Viral Disease Control and Prevention, China CDC | Xiang Zhao et al |
| **hCoV-19/Qatar/QA-QU_08-A4/2020** | EPI_ISL_1713042 | | Ministry of Public Health / Hamad Medical Corporation | Biomedical Research Center (BRC), Qatar University / Qatar Genome Project (QGP) | BRC: Fatiha M. Benslimane et al |
| **hCoV-19/Peru/LIM-INS-744/2020** | EPI_ISL_1137481 | | Laboratorio de Referencia Nacional de Virus Respiratorio. Instituto Nacional de Salud Perú | Laboratorio de Referencia Nacional de Biotecnología y Biología Molecular. Instituto Nacional de Salud Perú | Carlos Padilla Rojas et al |
| **hCoV-19/Maldives/MAV14010/2021** | EPI_ISL_3452719 | | Indira Gandhi Memorial Hospital | Indira Gandhi Memorial Hospital | Dr. Milza Abdul Muhsin et al |
| **hCoV-19/Russia/MOW-RII-MH16102S/2021** | EPI_ISL_1919488 | | HELIX LLC | WHO National Influenza Centre Russian Federation | Andrey Komissarov et al |
| **hCoV-19/Heilongjiang/IVDC-HH/235/2020** | EPI_ISL_9910186 | | Heilongjiang Center for Disease Control and Prevention | National Institute for Viral Disease Control and Prevention, China CDC | Wei Sun et al |
| **hCoV-19/Algeria/96626/2020** | EPI_ISL_12156733 | | NIC, Viral Respiratory Unit | NIC, Viral Respiratory Unit | Sarah Belkalem et al |
| **hCoV-19/Mexico/TAB-InDRE_446/2021** | EPI_ISL_1168524 | | Instituto de Diagnostico y Referencia Epidemiologicos INDRE_RNLSP | Instituto de Diagnostico y Referencia Epidemiologicos (INDRE) | Claudia Wong-Arambula et al |
| **hCoV-19/Mexico/CMX-INER-IBT-146/2020** | EPI_ISL_1302381 | OK435506.1 | Instituto Nacional de Enfermedades Respiratorias (INER) | Instituto de Biotecnología de la UNAM | Authors from IBT et al |
| **hCoV-19/Turkey/IST-GLAB-341/2020** | EPI_ISL_735356 | | Genomic Laboratory (GLAB) (Conjoint lab of Health Directorate of Istanbul and Istanbul Technical University) | Genomic Laboratory (GLAB), Istanbul Technical University | Ilker Karacan et al |
| **hCoV-19/Algeria/G35155-8850/2020** | EPI_ISL_766864 | | NIC Viral Respiratory Unit - Institut Pasteur of Algeria | National Reference Center for Viruses of Respiratory Infections, Institut Pasteur, Paris | Mélanie Albert et al |
| **hCoV-19/Turkey/CTF_19092020_191/2020** | EPI_ISL_2047926 | | IU-Cerrahpasa, Cerrahpasa School of Medicine, COVID-19 Lab | IU-Cerrahpasa, Cerrahpasa School of Medicine, COVID-19 Lab | Mert Kuskucu et al |
| **hCoV-19/Algeria/96150/2020** | EPI_ISL_12156732 | | NIC, Viral Respiratory Unit | NIC, Viral Respiratory Unit | Sarah Belkalem et al |
| **hCoV-19/Serbia/Lipljan-005360911/2020** | EPI_ISL_904010 | | Veterinary Specialized Institute Kraljevo | Veterinary Specialized Institute "Kraljevo", Serbia | Vidanovic et al |
| **hCoV-19/USA/IL-IDPH-I-000605/2020** | EPI_ISL_848433 | | Illinois Department of Public Health | Gagnon Lab, Southern Illinois University | Keith Gagnon et al |
| **hCoV-19/Guinea/GUI0090/2021** | EPI_ISL_4274049 | | Service des Maladies Infectieuses | Northwestern University - Center for Pathogen Genomics and Microbial Evolution | Ramon Lorenzo-Redondo et al |
| **hCoV-19/TrinidadandTobago/14119/2021** | EPI_ISL_1498262 | | The Caribbean Public Health Agency | Carrington Lab, Department of PreClinical Sciences, Faculty of Medical Sciences, The University of the West Indies | Nikita S. D. Sahadeo et al |
| **hCoV-19/SierraLeone/nmimr-SLE--SLSEQ047/2021** | EPI_ISL_15283210 | | Immunology, Noguchi Memorial Institute for Medical Research | Immunology, Noguchi Memorial Institute for Medical Research | Campbell et al |
| **hCoV-19/SierraLeone/CPHRL-094/2021** | EPI_ISL_4755979 | | China P3 Laboratory | Central Public Health Reference Laboratory | Allan K.O. Campbell et al |
| **hCoV-19/Mexico/CHH_INCMNSZ_CS-147039/2021** | EPI_ISL_13984122 | | Centro de Investigacion Integral MEDIVEST SC | Instituto Nacional de Ciencias Medicas y Nutricion Salvador Zubiran, Departamento de Infectologia | Miriam Arciniega Fuentes et al |
| **hCoV-19/USA/OH-21AM-084CO02953/2021** | EPI_ISL_2200088 | | The Ohio State University Applied Microbiology Services Laboratory | The Ohio State University Applied Microbiology Services Laboratory | Seth A. Faith PhD et al |
| **hCoV-19/Mexico/GUA-InDRE_F12432_S1333/2021** | EPI_ISL_1661499 | | LESP Guanajuato | Instituto de Diagnostico y Referencia Epidemiologicos (INDRE) | Claudia Wong-Arambula et al |
| **hCoV-19/Honduras/1946/2021** | EPI_ISL_2648258 | | Laboratorio Nacional de Vigilancia de la Salud – Sección de Virologia | Genomics and Proteomics Departament, Gorgas Memorial Institute For Health Studies | Elda Martínez et al |
| **hCoV-19/Mexico/un-UW-1902618/2021** | EPI_ISL_17757834 | | Laboratory Medicine and Pathology, University of Washington | Laboratory Medicine and Pathology, University of Washington | Roychoudhury et al |
| **hCoV-19/Mexico/GRO-InDRE_FB17786_S4179/2021** | EPI_ISL_2920750 | | LESP Guerrero | Instituto de Diagnostico y Referencia Epidemiologicos (INDRE) | Claudia Wong-Arambula et al |
| **hCoV-19/Mexico/YUC_LANGEBIO_IMSS_07539-NC/2021** | EPI_ISL_2970068 | | Unidad de Investigacion Medica de Yucatan (UIMY) | Unidad de Genomica Avanzada | Consorcio Mexicano de Vigilancia Genomica (CoViGen-Mex). Authors (in alphabetical order): Julio Elias Alvarado-Yaah et al |
| **hCoV-19/Mexico/VER-InDRE_169/2020** | EPI_ISL_913970 | | Instituto de Diagnostico y Referencia Epidemiologicos INDRE_RNLSP | Instituto de Diagnostico y Referencia Epidemiologicos (INDRE) | Claudia Wong-Arambula et al |
| **hCoV-19/Mexico/SLP_LANGEBIO_IMSS_0443/2021** | EPI_ISL_1662195 | | Laboratorio Central de Epidemiologia (LCE) | Unidad de Genomica Avanzada | Consorcio Mexicano de Vigilancia Genomica (CoViGen-Mex). Authors (in alphabetical order): Julio Elias Alvarado-Yaah et al |
| **hCoV-19/Mexico/YUC-InDRE_FB13182_S1581/2021** | EPI_ISL_1821153 | | LESP Yucatan | Instituto de Diagnostico y Referencia Epidemiologicos (INDRE) | Claudia Wong-Arambula et al |
| **hCoV-19/Russia/CHE-CSP-Cov-19_S33/2020** | EPI_ISL_8908058 | | Chief Federal State Budgetary Healthcare Institution "Centre of Hygiene and Epidemiology" of the Federal Medical Biological Agency, Central laboratory of microbiological and clinical diagnostic tests | Federal State Budgetary Institution "Centre for Strategic Planning and Management of Biomedical Health Risks" of the Federal Medical Biological Agency, Laboratory for advanced molecular diagnostics of human diseases. | Mikhaylov et al |
| **hCoV-19/Brunei/7122175428/2022** | EPI_ISL_15710366 | | National Virology Reference Laboratory | Microbial Genomic Services | Zainun Zaini et al |
| **hCoV-19/Australia/WA13361/2023** | EPI_ISL_17661363 | | PathWest Laboratory Medicine WA | PathWest Laboratory Medicine WA Microbial Surveillance Unit | PathWest Laboratory Medicine WA Microbial Surveillance Unit et al |
| **hCoV-19/NewZealand/23YA0805/2023** | EPI_ISL_16954279 | | Medlab Central (PN) | Institute of Environmental Science and Research (ESR) | Sarah Cockerton et al |
| **hCoV-19/NewZealand/ESR-XA4231/2023** | EPI_ISL_18047905 | | Waikato DHB Laboratory | Institute of Environmental Science and Research (ESR) | Sarah Cockerton et al |
| **hCoV-19/HongKong/VB23156220/2023** | EPI_ISL_18306760 | | Queen Mary Hospital | Hong Kong Department of Health | Alan K.L. Tsang et al |
| **hCoV-19/Australia/SA691145/2023** | EPI_ISL_18453812 | | SA Pathology | SA Pathology | Caitlin Selway et al |
| **hCoV-19/Moldova/Nr-17961/2022** | EPI_ISL_16017096 | | Virology Laboratory, NAPH, Moldova | Virology Laboratory, NAPH, Moldova | Olga Burduniuc et al |
| **hCoV-19/Germany/HH-RKI-I-426417/2021** | EPI_ISL_8674726 | | SYNLAB MVZ Hamburg | Robert Koch Institute | ? |
| **hCoV-19/Brazil/MG-FIOCRUZ-863/2022** | EPI_ISL_9747760 | | Laboratorio Municipal de Biologia Molecular | Instituto Rene Rachou / Fiocruz Minas | Thais Silva et al |
| **hCoV-19/Congo/RC-415/2021** | EPI_ISL_10023530 | | Molecular Lab, National Laboratory of Public Health | Pathogen Genomics Lab, National Institute for Biomedical Research (INRB) | Placide Mbala-Kingebeni et al B |
| **hCoV-19/Palestine/QIBPS-OM_5_S8/2022** | EPI_ISL_10550533 | | Al-Quds University | Quadram Institute Bioscience | Abdelmajeed Nasereddin et al |
| **hCoV-19/Senegal/22197316/2021** | EPI_ISL_14481782 | | Institut Pasteur de Dakar | Institut Pasteur de Dakar | Diallo Amadou et al |
| **hCoV-19/India/UT-ILBS-WGS2935/2022** | EPI_ISL_10630432 | | MAX HOSPITAL | ILBS | Ekta Gupta et al |
| **hCoV-19/Sweden/2A62215830099/2022** | EPI_ISL_10594522 | | The Public Health Agency of Sweden | The Public Health Agency of Sweden | Maximilian Riess et al |
| **hCoV-19/Madagascar/LA2M-179994/2021** | EPI_ISL_13729330 | | Laboratoire d'Analyses Médicales Malagasy (LA2M) | Virology Unit, Institut Pasteur de Madagascar | Vololoniaina Raharinosy et al |
| **hCoV-19/Kenya/ILRI_COVM01645/2021** | EPI_ISL_9093437 | | International Livestock Research Institute | International Livestock Research Institute | Samuel O. Oyola et al |
| **hCoV-19/Guinea/CERFIG-34031/2022** | EPI_ISL_14578567 | | CERFIG | CERFIG | Haby DIALLO et al |
| **hCoV-19/Senegal/SN-DKR-DIAM-4842/2021** | EPI_ISL_7571605 | | Iressef Genomics lab | IRESSEF | Souleymane MBOUP et al |
| **hCoV-19/Argentina/PAIS-E0729/2022** | EPI_ISL_10637273 | | Laboratorio Central de la Ciudad de Santa Fe | Grupo de Genómica y Bioinformática del Instituto de Investigación de la Cadena Láctea CONICET-INTA on behalf of 'Proyecto Argentino Interinstitucional de genomica de SARS-CoV-2' (PAIS Consortium) | Eberhardt et al |
| **hCoV-19/Venezuela/NE2311/2022** | EPI_ISL_10239604 | | IVIC | Laboratorio de Virologia Molecular | Rossana C Jaspe et al |
| **hCoV-19/Canada/MB-CPL-367037/2022** | EPI_ISL_10654840 | | MB-Cadham Provincial laboratory | Manitoba Cadham Provincial Laboratory | David Alexander et al |
| **hCoV-19/Uruguay/DLSP_577/2022** | EPI_ISL_17227679 | | Departamento Laboratorios de Salud Publica (DLSP) Ministerio de Salud Publica | Departamento Laboratorios de Salud Publica (DLSP) Ministerio de Salud Publica | Cortinas et al |
| **hCoV-19/Curacao/CW-RIVM-88422/2022** | EPI_ISL_10346014 | | National Institute for Public Health and the Environment (RIVM) | National Institute for Public Health and the Environment (RIVM) | Adam Meijer et al |
| **hCoV-19/Panama/M122900-GMI/2021** | EPI_ISL_11854323 | | Gorgas Memorial Institute of Health Studies | Gorgas Memorial Institute of Health Studies | Claudia Gonzalez et al |
| **hCoV-19/Mexico/JAL_InDRE_FB4142_E14313927569_S11212/2022** | EPI_ISL_9749739 | | LESP Jalisco | Instituto de Diagnostico y Referencia Epidemiologicos (INDRE) | Claudia Wong-Arambula et al |
| **hCoV-19/Bulgaria/22BG-EU_017400_PI164/2022** | EPI_ISL_11734777 | | National Center of Infectious and Parasitic Diseases | National Center of Infectious and Parasitic Diseases | Alexiev et al |
| **hCoV-19/Curacao/CW-RIVM-89999/2022** | EPI_ISL_10899255 | | National Institute for Public Health and the Environment (RIVM) | National Institute for Public Health and the Environment (RIVM) | Adam Meijer et al |
| **hCoV-19/Canada/QC-L00460809001/2022** | EPI_ISL_12299897 | | Laboratoire de santé publique du Québec | Laboratoire de santé publique du Québec | Sandrine Moreira et al |
| **hCoV-19/Guatemala/3179-LNS/2022** | EPI_ISL_13080307 | | Laboratorio Nacional de Salud (LNS), Guatemala | Laboratorio Nacional de Salud (LNS), Guatemala | Mendoza L et al |
| **hCoV-19/Benin/MTMC_05_109/2022** | EPI_ISL_17425649 | | Centre Hospitalier et Universitaire de Zone Abomey-Calavi | Molecular Malaria Laboratory, Mahidol Oxford Tropical Medicine Research Unit | Aurel Allabi et al |
| **hCoV-19/EastTimor/898261/2022** | EPI_ISL_14710807 | | Laboratório Nacional em Pesquisas Microbiológicas | Centro de Sequenciamento Genômico | Soraia Antunes et al |
| **hCoV-19/Macau/NMA9990/2021** | EPI_ISL_14710810 | | Laboratório Nacional em Pesquisas Microbiológicas | Centro de Sequenciamento Genômico | Soraia Antunes et al |
| **hCoV-19/UnitedArabEmirates/BTC-10816/2022** | EPI_ISL_16800098 | | Khalifa University Center for Biotechnology | Khalifa University Center for Biotechnology | AlSafar et al. |
| **hCoV-19/Lesotho/N36081/2022** | EPI_ISL_12956136 | | NATIONAL REFERENCE LABORATORIES | National Institute for Communicable Diseases of the National Health Laboratory Service | Maphalala G et al |
| **hCoV-19/Lesotho/N36097/2022** | EPI_ISL_12956150 | | NATIONAL REFERENCE LABORATORIES | National Institute for Communicable Diseases of the National Health Laboratory Service | Maphalala G et al |
| **hCoV-19/Finland/R-1399/2022** | EPI_ISL_12416492 | | Department of Virology and Immunology, University of Helsinki and Helsinki University Hospital, Huslab Finland | Department of Virology, Faculty of Medicine, University of Helsinki, Helsinki, Finland | Teemu Smura et al |
| **hCoV-19/Lebanon/LEBN-GS-85/2022** | EPI_ISL_14605844 | | American University of Beirut Medical Center | Melhem/Matar lab | Nada M. Melhem et al |
| **hCoV-19/DominicanRepublic/DR-Yale-21433/2022** | EPI_ISL_15096319 | | Instituto de Medicina Tropical & Salud Global Universidad Iberoamericana | Grubaugh Lab - Yale School of Public Health | Chantal Vogels et al |
| **hCoV-19/Ghana/NMIMR-NCOV-21-61240/2021** | EPI_ISL_11765262 | | Noguchi Memorial Institute for Medical Research, University of Ghana, Legon, Ghana | Noguchi Memorial Institute for Medical Research, University of Ghana, Legon, Ghana | Quaneeta Mohktar et al |
| **hCoV-19/Myanmar/DSMRC-063/2021** | EPI_ISL_8920492 | | DSMRC | Defence Services Medical Research Centre (DSMRC) | Khine Zaw Oo et al |
| **hCoV-19/Slovakia/BA_22_00010261/2022** | EPI_ISL_10774887 | | Biomedical Centre Martin, Jessenius Faculty of Medicine in Martin, Comenius University | Laboratory of Genomics and Bioinformatics, Comenius University Science Park | Dušan Loderer et al |
| **hCoV-19/Malta/MDxMDHO126/2022** | EPI_ISL_9776828 | | Molecular Diagnostics Pathology Department Mater Dei Hospital Malta | Molecular Diagnostics Pathology Department Mater Dei Hospital Malta | G Zahra et al |
| **hCoV-19/Zambia/CHAZ-CHS0739/2021** | EPI_ISL_10222421 | | CHAZ Laboratory | CHAZ Laboratory | Shempela. D et al |
| **hCoV-19/Bonaire/BQ-RIVM-87375/2022** | EPI_ISL_10345976 | | National Institute for Public Health and the Environment (RIVM) | National Institute for Public Health and the Environment (RIVM) | Adam Meijer et al |
| **hCoV-19/Peru/ARE-INS-16521/2022** | EPI_ISL_12256512 | | Laboratorio de Referencia Nacional de Virus Respiratorios. Centro Nacional de Salud Publica. Instituto Nacional de Salud Peru. | Laboratorio de Referencia Nacional de Virus Respiratorios. Centro Nacional de Salud Publica. Instituto Nacional de Salud Peru. | Carlos Padilla Rojas et al |
| **hCoV-19/India/GA-ICMR-NIV-INSACOG-384403/2021** | EPI_ISL_9494651 | | ICMR-National Institute of Virology - INSACOG | NIV Influenza | Dr. Varsha Potdar et al |
| **hCoV-19/Poland/WSSEGorzow-22S0036/2021** | EPI_ISL_8643670 | | Diagnostyka Sp. z o. o. Wrocław | Wojewodzka Stacja Sanitarno-Epidemiologiczna w Gorzowie Wielkopolskim | Elzbieta Justynska et al |
| **hCoV-19/Canada/NL-PHML-344743/2021** | EPI_ISL_13078886 | | NL-Dr. Leonard A. Miller Centre for Health Services | Newfoundland and Labrador - Eastern Health | Anna Majer et al |
| **hCoV-19/Guatemala/1057-LNSGua/2022** | EPI_ISL_9801667 | | Laboratorio Nacional de Salud (LNS), Guatemala | Laboratorio Nacional de Salud (LNS), Guatemala | Mendoza L et al |
| **hCoV-19/Beijing/BJCDC-260/2022** | EPI_ISL_16610203 | | Beijing Center for Disease Control and Prevention | Beijing Center for Disease Control and Prevention | Zhaomin Feng et al |
| **hCoV-19/Singapore/2479/2022** | EPI_ISL_11019906 | | National Public Health Laboratory, National Centre for Infectious Diseases | National Public Health Laboratory, National Centre for Infectious Diseases | Ding Yichen et al |
| **hCoV-19/Aruba/AW-RIVM-82598/2022** | EPI_ISL_9184723 | | National Institute for Public Health and the Environment (RIVM) | National Institute for Public Health and the Environment (RIVM) | Adam Meijer et al |

| | | | | | |
|---|---|---|---|---|---|
| **hCoV-19/Aruba/AW-RIVM-82613/2022** | EPI_ISL_9184738 | | National Institute for Public Health and the Environment (RIVM) | National Institute for Public Health and the Environment (RIVM) | Adam Meijer et al |
| **hCoV-19/Cameroon/CERI-KRISP-K040916/2021** | EPI_ISL_12650898 | | National Public Health Laboratory, Ministry of Public Health, Yaounde, Messa, Cameroon | CERI, Centre for Epidemic Response and Innovation, Stellenbosch University and KRISP, KZN Research Innovation and Sequencing Platform, UKZN. | Dr Essomba Rene Ghislain et al |
| **hCoV-19/Zimbabwe/S_3083_B49/2021** | EPI_ISL_9184708 | | National Microbilogy Reference Laboratory | National Microbilogy Reference Laboratory | Jasper Chimedza et al |
| **hCoV-19/Lesotho/Ls-73895/2021** | EPI_ISL_13101650 | | National Reference Laboratory - Ministry of Health Maseru Lesotho | National Reference Laboratory - Ministry of Health Maseru Lesotho | Marelebogile Pooe et al |
| **hCoV-19/Nigeria/LSB238/2022** | EPI_ISL_14056947 | | Lagos State Biobank | Lagos State Biobank | Akin Abayomi et al |
| **hCoV-19/Netherlands/NH-inBiome-210610/2022** | EPI_ISL_8783336 | | Public Health Laboratory, Public Health Service, Amsterdam, The Netherlands | inBiome | Martine Bos et al |
| **hCoV-19/HongKong/HK-HKPU-12880/2022** | EPI_ISL_12417511 | | Department of Health Technology and Informatics, The Hong Kong Polytechnic University | Department of Health Technology and Informatics, The Hong Kong Polytechnic University | Gilman Kit-Hang Siu et al |
| **hCoV-19/Morocco/611/2022** | EPI_ISL_13421944 | | NIC at NIH | National Institute of Hygiene | Hassan Ihazmade et al |
| **hCoV-19/Zhejiang/ZPCDCP-62/2022** | EPI_ISL_17261564 | | Zhejiang Provincial Center for Disease Control and Prevention | Zhejiang Provincial Center for Disease Control and Prevention | Yi Sun et al |
| **hCoV-19/Canada/NL-PHML-378797/2022** | EPI_ISL_10926089 | | NL-Dr. Leonard A. Miller Centre for Health Services | Newfoundland and Labrador - Eastern Health | Anna Majer et al |
| **hCoV-19/Taiwan/NTU-HCH-100/2022** | EPI_ISL_14189462 | | Department of Laboratory Medicine, National Taiwan University Hospital Hsin-Chu Hospital, Taiwan | Research Center for Emerging Viral Infections, Chang Gung University, Taiwan | Pei-Lan Shao et al |
| **hCoV-19/Mali/MTMC_07_346/2022** | EPI_ISL_17425653 | | Hospital Of Mali | Molecular Malaria Laboratory, Mahidol Oxford Tropical Medicine Research Unit | Samba Sow et al |
| **hCoV-19/Kenya/KNPHLGEN127/2022** | EPI_ISL_18098360 | | Mbaghadhi County Hospital | Kenya NGMSL | Leonard Kingrawa et al |
| **hCoV-19/Brazil/TO-LACENTO-170503397/2022** | EPI_ISL_12690811 | | Laboratório Central de Saúde Pública do Estado do Tocantins (LACEN/TO) | Laboratório Central de Saúde Pública do Estado do Tocantins (LACEN/TO) | Ueric José Borges de Souza et al |
| **hCoV-19/Australia/QLD0x00CE72/2022** | EPI_ISL_10975816 | | Pathology Queensland and Forensic Scientific Services | PHV-FSS | Chenwei Wang on behalf of Q-PHIRE Genomics et al |
| **hCoV-19/Australia/ACT3012/2022** | EPI_ISL_10010141 | | ACT Pathology | Schwessinger Lab | Austin Bird et al |
| **hCoV-19/Ethiopia/ILRI_COVM02877/2021** | EPI_ISL_11294988 | | Addis Ababa University | International Livestock Research Institute | Abay Sisay et al |
| **hCoV-19/Australia/NSW-ICPMR-20385/2022** | EPI_ISL_9821215 | | Pathology West - NSW Health Pathology | NSW Health Pathology - Institute of Clinical Pathology and Medical Research; Westmead Hospital; University of Sydney | Arnott A. et al |
| **hCoV-19/Andorra/AND-253_220681345301/2022** | EPI_ISL_12349794 | | Laboratori d'anàlisis clíniques, Hospital Nostra Senyora de Meritxell | LBM de CHU de Toulouse, Hôpitaux de Toulouse | Rendon M et al |
| **hCoV-19/India/GJ-INSACOG-GBRC4048/2021** | EPI_ISL_8744178 | | Unipath Speciality Laboratory Limited, Ahmedabad | Gujarat Biotechnology Research Centre | Jaykumar Rangani et al |
| **hCoV-19/Zambia/ZMB-128376/2021** | EPI_ISL_7988149 | | University of Zambia, School of Veterinary Medicine | UNZAVET and PATH | Mulenga Mwenda-Chimfwembe et al |
| **hCoV-19/Indonesia/JK-GS-GSILab-858673/2022** | EPI_ISL_8443087 | | RS ST CAROLUS JAKARTA | Genomik Solidaritas Indonesia Laboratorium | Dian Pitaloka Rajagukguk et al |
| **hCoV-19/Oman/5225286/2022** | EPI_ISL_16073631 | | Oman-NIC | Oman-NIC | Samiha Al Kharusi et al |
| **hCoV-19/Tonga/22CV3411/2022** | EPI_ISL_18236467 | | Vaiola Hospital | Institute of Environmental Science and Research (ESR) | Senisaleti Pasikala et al |
| **hCoV-19/Serbia/Belgrade-406121_0101/2022** | EPI_ISL_9467235 | | Institute of Virology, Vaccines and Sera "Torlak" | Institute of microbiology and Immunology, Faculty of Medicine, University of Belgrade | Knezevic et al |
| **hCoV-19/Eswatini/N22763/2021** | EPI_ISL_9439186 | | Good Shepard Hospital | National Institute for Communicable Diseases of the National Health Laboratory Service | Maphalala G et al |
| **hCoV-19/Iraq/KR-SEARCH-120469/2022** | EPI_ISL_12604522 | | University of Duhok COVID-19 Center for Research and Diagnosis | Andersen lab at Scripps Research | SEARCH Alliance with Jassim Mohammed Abdo et al |
| **hCoV-19/Djibouti/NAMRU3_H932/2021** | EPI_ISL_9712325 | OM635175.1 | Naval Medical Research Unit No. 3 | Naval Medical Research Center Biological Defense Research Directorate | Logan J. Voegtly et al |
| **hCoV-19/Georgia/Tb-s3057/2022** | EPI_ISL_8878829 | | JSC Megalab | Department for Virology, Molecular Biology and Genome Research, R. G. Lugar Center for Public Health Research, National Center for Disease Control and Public Health (NCDC) of Georgia. | Giorgi Tomashvili et al |
| **hCoV-19/Djibouti/NAMRU3_C233/2022** | EPI_ISL_11533216 | ON099649.1 | Naval Medical Research Unit No. 3 | Naval Medical Research Center Biological Defense Research Directorate | Logan J. Voegtly et al |
| **hCoV-19/India/PY-ILBS-WGS3843/2022** | EPI_ISL_10661129 | | RVRDL-JIPMER | ILBS | Ekta Gupta et al |
| **hCoV-19/Cameroon/CERI-KRISP-K040710/2022** | EPI_ISL_12749887 | | National Public Health Laboratory, Ministry of Public Health, Yaounde | CERI, Centre for Epidemic Response and Innvoation, Stellenbosch University and KRISP, KZN Research Innovation and Sequencing Platform, UKZN. | Ghislain ER et al |
| **hCoV-19/Ethiopia/ILRI_COVM02834/2022** | EPI_ISL_11294976 | | Addis Ababa University | International Livestock Research Institute | Abay Sisay et al |
| **hCoV-19/Israel/ICH-741110535/2022** | EPI_ISL_8519600 | | Clinical Microbiology Laboratory, Tel Aviv Sourasky Medical Center | Clinical Microbiology Laboratory, Tel Aviv Sourasky Medical Center | Katya Levytskyi et al |
| **hCoV-19/Hubei/IVDC-553/2022** | EPI_ISL_11873916 | | Hubei Provincial Center for Disease Control and Prevention | National Institute for Viral Disease Control and Prevention, China CDC | Bin Fang et al |
| **hCoV-19/Shandong/WCDC-128/2022** | EPI_ISL_18070460 | | Weihai Center for Disease Control and Prevention | Weihai Center for Disease Control and Prevention | Xiang Li et al |
| **hCoV-19/SouthKorea/KDCA34964/2022** | EPI_ISL_12129863 | | Division of Emerging Infectious Diseases, Bureau of Infectious Diseases Diagnosis Control, Korea Disease Control and Prevention Agency | Division of Emerging Infectious Diseases, Bureau of Infectious Diseases Diagnosis Control, Korea Disease Control and Prevention Agency | Jeong-Ah Kim et al |
| **hCoV-19/Montenegro/CO-00777_MNE000_1604552202/2022** | EPI_ISL_12345050 | | Laboratory for Molecular Diagnostics/JPHMN | Eurofins Genomics Europe Sequencing GmbH | Danijela Vujošević et al |
| **hCoV-19/NorthMacedonia/IPH-MKD-143/2022** | EPI_ISL_13032691 | | Laboratory of Virology - Institute of Public Health Skopje, Macedonia | Laboratory of Virology - Institute of Public Health Skopje, Macedonia | Teodora Buzharova et al |
| **hCoV-19/Sichuan/IVDC-LZ-588/2022** | EPI_ISL_11905853 | | Sichuan Center of Disease Control and Prevention | National Institute for Viral Disease Control and Prevention, China CDC | Jianan Xu et al |
| **hCoV-19/Cameroon/LNSP-001-34703/2022** | EPI_ISL_17104890 | | NPHL-Cameroon | National Public Health Laboratory, Cameroon | Dr Essomba Rene Ghislain et al |
| **hCoV-19/DRC/INRB-RDC-0298/2022** | EPI_ISL_17190910 | | Laboratoire Rodolphe Mérieux (INRB Goma) | Pathogen Genomics Lab, National Institute for Biomedical Research (INRB) | Placide Mbala-Kingebeni et al B |
| **hCoV-19/Azerbaijan/un-ChWr28865/2022** | EPI_ISL_15714284 | | Xususi Tehlukeli Infeksiyalara Nezaret Merkezi | Charité Universitätsmedizin Berlin, Institut für Virologie | Victor M Corman et al |
| **hCoV-19/SouthSudan/UG1177/2021** | EPI_ISL_10512583 | | South Sudan Ministry of Health, WHO South Sudan, MRC/UVRI & LSHTM Uganda Research Unit | MRC/UVRI & LSHTM Uganda Research Unit, South Sudan Ministry of Health, WHO South Sudan | Abe G Abias et al |
| **hCoV-19/Jordan/MoH-1013/2021** | EPI_ISL_8189873 | | Ministry of Health - Central Labs | Ministry of Health - Central Labs | Dr. Mahmoud Ghazo et al |
| **hCoV-19/Egypt/CPHL-EGY22656/2022** | EPI_ISL_16299767 | | CPHL_MOH,EGYPT | CPHL | Wael H. Roshdy et al |
| **hCoV-19/Kyrgyzstan/NRL-90518/2022** | EPI_ISL_13711460 | | National Virological Laboratory of the Department of Disease Prevention and State Sanitary and Epidemiological Surveillance | Republican Reference Laboratory, National TB center, Kyrgyzstan | Gulmira Kalmambetova et al |
| **hCoV-19/Israel/CVL-9001528/2022** | EPI_ISL_12095082 | | Public health laboratories Tel Aviv | Israel Central Virology laboratory | Neta Zuckerman et al |
| **hCoV-19/Pakistan/AKU-JAN-RUN-MIN-15/2021** | EPI_ISL_9861357 | | Department of Pathology and Laboratory Medicine, AKUH Laboratories, Karachi, Pakistan | Department of Pathology and Laboratory Medicine, AKUH Laboratories, Karachi, Pakistan | A.R. Bukhari et al |
| **hCoV-19/Bolivia/SAN-UPCH-3346/2021** | EPI_ISL_13845427 | | Laboratorio de Diagnóstico e Investigación BIOSCIENCE SRL | Laboratorio de Genómica Microbiana, Universidad Peruana Cayetano Heredia | Esdenka Pérez et al |
| **hCoV-19/Nigeria/BCVL_GHRU_32946_S41/2022** | EPI_ISL_16248832 | | Biorepository Clinical Virology Laboratory, College of Medicine | Global Health Research Unit for the Genomic Surveillance of Antimicrobial Resistance, University of Ibadan | Fowotade AA et al |
| **hCoV-19/Serbia/Belgrade-K6874144_2302/2022** | EPI_ISL_11704798 | | Laboratory for COVID19 diagnostics, Clinical Centre of Serbia | Institute of microbiology and Immunology, Faculty of Medicine, University of Belgrade | Knezevic et al |
| **hCoV-19/Germany/HB-RKI-I-457697/2022** | EPI_ISL_9208986 | | Sonic - MVZ Medizinisches Labor Bremen GmbH | Robert Koch Institute | ? |
| **hCoV-19/UnitedArabEmirates/BTC-10796/2022** | EPI_ISL_16800079 | | Khalifa University Center for Biotechnology | Khalifa University Center for Biotechnology | AlSafar et al. |
| **hCoV-19/UnitedArabEmirates/BTC-10819/2022** | EPI_ISL_16800101 | | Khalifa University Center for Biotechnology | Khalifa University Center for Biotechnology | AlSafar et al. |
| **hCoV-19/UnitedArabEmirates/BTC-10815/2022** | EPI_ISL_16800097 | | Khalifa University Center for Biotechnology | Khalifa University Center for Biotechnology | AlSafar et al. |
| **hCoV-19/Chile/VS-UV-368/2022** | EPI_ISL_10251497 | | Laboratorio de Genomica y Laboratorio de Virologia Clinica, Universidad de Valparaiso | Laboratorio de Genomica y Laboratorio de Virologia Clinica, Universidad de Valparaiso | Javier Tognarelli et al |
| **hCoV-19/Vietnam/PIHCM_083/2022** | EPI_ISL_9416724 | | Pasteur Institute in Ho Chi Minh city | Department of Microbiology and Immunology | Hang Pham Thi Thu et al |
| **hCoV-19/Peru/CAJ-INS-14000/2022** | EPI_ISL_10454176 | | Laboratorio de Referencia Nacional de Virus Respiratorios. Centro Nacional de Salud Publica. Instituto Nacional de Salud Peru. | Laboratorio de Referencia Nacional de Virus Respiratorios. Centro Nacional de Salud Publica. Instituto Nacional de Salud Peru. | Carlos Padilla Rojas et al |
| **hCoV-19/Mexico/SON-IBT_IMSS_6634/2022** | EPI_ISL_11202173 | | Centro de Investigación Biomédica de Occidente (CIBO) | Instituto de Biotecnologia de la UNAM | Consorcio Mexicano de Vigilancia Genómica (CoViGen-Mex). Authors (in alphabetical order): Julio Elias Alvarado-Yaah et al |
| **hCoV-19/Chile/RM-35412/2022** | EPI_ISL_11565172 | | Genetica Molecular and Subdepartamento de Virologia ISP Chile | Instituto de Salud Pública de Chile | Karen Oroztica et al |
| **hCoV-19/Mexico/BCS_InDRE_FB13182_E03316125404_S1308 8/2022** | EPI_ISL_11412971 | | LESP Baja California Sur | Instituto de Diagnostico y Referencia Epidemiologicos (INDRE) | Claudia Wong-Arambula et al |
| **hCoV-19/Beijing/Omicron-1/2021** | EPI_ISL_16138082 | | Key Laboratory of Human Diseases Comparative Medicine, Institute of Laboratory Animal Science | Key Laboratory of Human Diseases Comparative Medicine, Institute of Laboratory Animal Science | Qin et al |
| **hCoV-19/Liaoning/IVDC-589/2022** | EPI_ISL_11905854 | | Dalian Municipal Center for Disease Control and Prevention | National Institute for Viral Disease Control and Prevention, China CDC | Xingying Lang et al |
| **hCoV-19/Tianjin/TJCDC-0036/2022** | EPI_ISL_16923077 | | Tianjin Centers for Disease Control and Prevention | Tianjin Centers for Disease Control and Prevention | Li Xiaoyan et al |
| **hCoV-19/Tianjin/TJCDC-0243/2022** | EPI_ISL_16923277 | | Tianjin Centers for Disease Control and Prevention | Tianjin Centers for Disease Control and Prevention | Li Xiaoyan et al |
| **hCoV-19/TrinidadandTobago/130276/2022** | EPI_ISL_11998857 | | Trinidad Public Health Laboratory | Carrington Lab, Department of Preclinical Sciences, Faculty of Medical Sciences, The University of the West Indies, St Augustine Campus | Nikita S. D. Sahadeo et al |
| **hCoV-19/Guyana/2421-GMI/2022** | EPI_ISL_11764502 | | National Public Health Reference Laboratory, Ministry of Health, | Genomics and Proteomics Departament, Gorgas Memorial Institute For Health Studies | Claudia Gonzalez et al |
| **hCoV-19/Guyana/2520-GMI/2022** | EPI_ISL_11764503 | | National Public Health Reference Laboratory, Ministry of Health, | Genomics and Proteomics Departament, Gorgas Memorial Institute For Health Studies | Claudia Gonzalez et al |
| **hCoV-19/Guyana/2409-GMI/2022** | EPI_ISL_11764501 | | National Public Health Reference Laboratory, Ministry of Health, | Genomics and Proteomics Departament, Gorgas Memorial Institute For Health Studies | Claudia Gonzalez et al |
| **hCoV-19/Vietnam/PIHCM_096/2022** | EPI_ISL_9416693 | | Pasteur Institute in Ho Chi Minh city | Department of Microbiology and Immunology | Thang Minh Cao et al |
| **hCoV-19/Myanmar/DSMRC-067/2022** | EPI_ISL_8920500 | | DSMRC | Defence Services Medical Research Centre (DSMRC) | Khine Zaw Oo et al |
| **hCoV-19/Belize/CML-472/2022** | EPI_ISL_13766397 | | Central Medical Laboratory | Central Medical Laboratory | Roberto Melendez et al |
| **hCoV-19/Ecuador/NIC-INSPI-112307/2022** | EPI_ISL_11760559 | | INSPI-CRN DE INFLUENZA Y OTROS VIRUS RESPIRATORIOS | INSPI-CRN DE INFLUENZA Y OTROS VIRUS RESPIRATORIOS | Alfredo Bruno et al |
| **hCoV-19/Heilongjiang/IVDC-549/2022** | EPI_ISL_11873912 | | HeiLongjiang Provincial Center For Disease Control And Prevention | National Institute for Viral Disease Control and Prevention, China CDC | Qi Liang et al |
| **hCoV-19/Canada/ON-PHL-22-13587/2022** | EPI_ISL_11500107 | | Public Health Ontario Laboratory | Public Health Ontario Laboratory | Vanessa G Allen et al |
| **hCoV-19/Uzbekistan/3_1465-CAT/2022** | EPI_ISL_15941881 | | SI"Zangiota No.1 and 2 special hospital for the treatment of patients with coronovirus infection" | Center for Advanced Technologies | Dalimova D.A. et al |
| **hCoV-19/Zhejiang/ZPCDCP-42/2021** | EPI_ISL_17261544 | | Zhejiang Provincial Center for Disease Control and Prevention | Zhejiang Provincial Center for Disease Control and Prevention | Yi Sun et al |
| **hCoV-19/USA/WA-PHL-013719/2022** | EPI_ISL_9918855 | | Washington State Department of Health Public Health Laboratories | Washington State Department of Health Public Health Laboratories | Drew MacKellar et al |
| **hCoV-19/Madagascar/LA2M-207488/2022** | EPI_ISL_13729358 | | Laboratoire d'Analyses Médicales Malagasy (LA2M) | Virology Unit, Institut Pasteur de Madagascar | Vololoniaina Raharinosy et al |
| **hCoV-19/Estonia/18806064/2022** | EPI_ISL_10444801 | | Synlab Eesti OÜ | 1. Laboratory of Communicable Diseases (Estonia); 2. Eurofins Genomics Europe Sequencing GmbH | Abroi A. et al |
| **hCoV-19/Ireland/WD-NVRL-S22IRL00073514/2022** | EPI_ISL_11548783 | | National Virus Reference Laboratory | National Virus Reference Laboratory | Zoe Yandle et al |
| **hCoV-19/USA/GA-CDC-LC0462170/2022** | EPI_ISL_9037856 | OM358370.1 | Laboratory Corporation of America | Centers for Disease Control and Prevention Division of Viral Diseases, Pathogen Discovery | Dakota Howard et al |
| **hCoV-19/U.S.VirginIslands/VI-CDC-2-5571419/2022** | EPI_ISL_10497770 | OM865541.1 | VI-US Virgin Islands Department of Health | Centers for Disease Control and Prevention Division of Viral Diseases, Pathogen Discovery | Mili Sheth et al |
| **hCoV-19/USA/KY-CDC-LC0531962/2022** | EPI_ISL_10271269 | OM788214.1 | Laboratory Corporation of America | Centers for Disease Control and Prevention Division of Viral Diseases, Pathogen Discovery | Dakota Howard et al |
| **hCoV-19/USA/AL-CDC-ASC210689669/2022** | EPI_ISL_9934215 | OM844377.1 | Aegis Sciences Corporation | Centers for Disease Control and Prevention Division of Viral Diseases, Pathogen Discovery | Dakota Howard et al |
| **hCoV-19/Albania/IPH-263012_2/2022** | EPI_ISL_16859981 | | Institute of Public Health | Institute of Public Health | Iris Hasibra et al |
| **hCoV-19/Bulgaria/22BG-EU_016764_PI157/2022** | EPI_ISL_11734177 | | National Center of Infectious and Parasitic Diseases | National Center of Infectious and Parasitic Diseases | Alexiev et al |
| **hCoV-19/SouthKorea/KDCA32158/2022** | EPI_ISL_11149246 | | Division of Emerging Infectious Diseases, Bureau of Infectious Diseases Diagnosis Control, Korea Disease Control and Prevention Agency | Division of Emerging Infectious Diseases, Bureau of Infectious Diseases Diagnosis Control, Korea Disease Control and Prevention Agency | Jeong-Ah Kim et al |
| **hCoV-19/InnerMongolia/IVDC-596/2022** | EPI_ISL_12030361 | | The center for disease control and prevention of Hulunbeier | National Institute for Viral Disease Control and Prevention, China CDC | Zhansong Zhu et al |
| **hCoV-19/Nicaragua/CNDR-667/2022** | EPI_ISL_17065328 | | Centro Nacional de Diagnóstico y Referencia | CNDR, Departamento de Virología | Angel Balmaseda et al |
| **hCoV-19/Azerbaijan/un-ChWr28819/2022** | EPI_ISL_15714253 | | Xususi Tehlukeli Infeksiyalara Nezaret Merkezi | Charité Universitätsmedizin Berlin, Institut für Virologie | Victor M Corman et al |
| **hCoV-19/NewZealand/22CV1505/2022** | EPI_ISL_10186714 | | MedLab South Ltd Nelson | Institute of Environmental Science and Research (ESR) | Sarah Cockerton et al |
| **hCoV-19/Hebei/IVDC-545/2022** | EPI_ISL_11873908 | | Hebei Provincial Center for Disease Control and Prevention | National Institute for Viral Disease Control and Prevention, China CDC | Shiyou Liu et al |
| **hCoV-19/Mongolia/UB-45265/2022** | EPI_ISL_14713866 | | National Center for Communicable Diseases (NCCD) National Influenza Center | National Center for Communicable Diseases (NCCD) National Influenza Center | Naranzul Tsedenbal et al |
| **hCoV-19/USA/WI-Mayo23601/2022** | EPI_ISL_13677963 | | Dept. Clinical Microbiology/Mayo Clinic | Dept. Clinical Microbiology/Mayo Clinic | Pritt Bobbi Norgan Andrew et al |
| **hCoV-19/Algeria/26888/2022** | EPI_ISL_13282959 | | Service microbiologie - EHU Oran | Virology Departement | Zahida Mehdi et al |
| **hCoV-19/SriLanka/aicbu4610/2022** | EPI_ISL_10104379 | | Allergy, Immunology and Cell Biology Unit (AICBU) | Allergy, Immunology and Cell Biology Unit (AICBU) | Chandima Jeewandara et al |
| **hCoV-19/USA/LA-TUSOM-220205_S36/2022** | EPI_ISL_9817530 | | Tulane University School of Medicine | Tulane University School of Medicine | Di Tian et al |
| **hCoV-19/Croatia/29498/2022** | EPI_ISL_10447239 | | Zavod za javno zdravstvo Šibensko- Kninske županije | Hrvatski zavod za javno zdravstvo | Irena Tabain et al |
| **hCoV-19/Gansu/IVDC-577/2022** | EPI_ISL_11873940 | | Gansu Provincial Center for Disease Control and Prevention | National Institute for Viral Disease Control and Prevention, China CDC | Baodi Li et al |
| **hCoV-19/Canada/NB-CHUDGLD-280222-MM00180R/2022** | EPI_ISL_11169513 | | New Brunswick - Vitalité Health Network | New Brunswick - Vitalité Health Network | Chacko S. et al |
| **hCoV-19/India/WB-INSACOG-1930301985134/2022** | EPI_ISL_13159160 | | INSACOG-WB | National Institute of Biomedical Genomics – INSACOG | Arindam Maitra et al A |
| **hCoV-19/UnitedArabEmirates/BTC-10829/2022** | EPI_ISL_16800107 | | Khalifa University Center for Biotechnology | Khalifa University Center for Biotechnology | AlSafar et al. |
| **hCoV-19/Ethiopia/CO_GX_439/2022** | EPI_ISL_17615496 | | Ethiopian Public Health Institute | EPHI Pathogen Genomic Sequence Lab | Woldaregay Erku Abegaz et al |
| **hCoV-19/Chongqing/YBCDC-0147/2022** | EPI_ISL_16829067 | | Chongqing Yubei District Center for Disease Control and Prevention | Chongqing Yubei District Center for Disease Control and Prevention | Jin Yan et al |
| **hCoV-19/Venezuela/Mer777/2021** | EPI_ISL_15165729 | | INHRR | Laboratorio de Virología Molecular | Rossana C Jaspe et al |
| **hCoV-19/Tianjin/TJCDC-0026/2021** | EPI_ISL_16922286 | | Tianjin Centers for Disease Control and Prevention | Tianjin Centers for Disease Control and Prevention | Li Xiaoyan et al |
| **hCoV-19/DRC/GOM-036/2022** | EPI_ISL_11450689 | | The Rodolphe Mérieux Laboratory of Goma (INRB) | Pathogen Sequencing Lab, National Institute for Biomedical Research (INRB) | Placide Mbala-Kingebeni et al B |
| **hCoV-19/Russia/MOW-FMVL-DZ-29064936912/2022** | EPI_ISL_15877383 | | N.F. Gamaleya National Research Center for Epidemiology and Microbiology of the Ministry of Health of the Russian Federation / Moscow Healthcare Department | N.F. Gamaleya National Research Center for Epidemiology and Microbiology of the Ministry of Health of the Russian Federation / Moscow Healthcare Department | Vladimir A. Gushchin et al |
| **hCoV-19/Mexico/SON_IBT_IMSS_6414/2022** | EPI_ISL_10436366 | | Centro de Investigación Biomédica de Occidente (CIBO) | Instituto de Biotecnologia de la UNAM | Consorcio Mexicano de Vigilancia Genómica (CoViGen-Mex). Authors (in alphabetical order): Julio Elias Alvarado-Yaah et al |
| **hCoV-19/USA/CT-JAX-2209-002/2022** | EPI_ISL_10707295 | | The Jackson Laboratory | The Jackson Laboratory | Long E et al |
| **hCoV-19/USA/ID-IBL-787489/2022** | EPI_ISL_9520562 | | Idaho Bureau of Laboratories | IBL | "R. Beukelman et al |
| **hCoV-19/Shanghai/IVDC-613/2022** | EPI_ISL_12461214 | | Shanghai Municipal Center for Disease Control & Prevention | National Institute for Viral Disease Control and Prevention, China CDC | Hui Jiang et al |
| **hCoV-19/Zhejiang/ZPCDCP-71/2022** | EPI_ISL_17261573 | | Zhejiang Provincial Center for Disease Control and Prevention | Zhejiang Provincial Center for Disease Control and Prevention | Yi Sun et al |
| **hCoV-19/USA/MS-USAFSAM-S10413/2021** | EPI_ISL_8976252 | | US Air Force School of Aerospace Medicine | US Air Force School of Aerospace Medicine | Anthony Fries et al |
| **hCoV-19/Palestine/28_2201_511_S3/2022** | EPI_ISL_12375043 | | Midecare | Al-Quds University | Abedelmajeed Nasereddin et al |
| **hCoV-19/Argentina/INEI117898/2022** | EPI_ISL_10705521 | | Servicio Virosis Respiratorias-Departamento Virología-INEI | Instituto Nacional Enfermedades Infecciosas C.G.Malbran | Baumeister E. et al |
| **hCoV-19/Ecuador/NIC-INSPI-397480/2022** | EPI_ISL_9903558 | | INSPI-CRN DE INFLUENZA Y OTROS VIRUS RESPIRATORIOS | INSPI-CRN DE INFLUENZA Y OTROS VIRUS RESPIRATORIOS | Alfredo Bruno et al |
| **hCoV-19/Uruguay/DLSP_701/2022** | EPI_ISL_12532643 | | Departamento Laboratorios de Salud P.blica (DLSP) Ministerio de Salud P.blica | Departamento Laboratorios de Salud P.blica (DLSP) Ministerio de Salud P.blica | Cortinas et al |
| **hCoV-19/Liechtenstein/FL-Risch-2231002314/2022** | EPI_ISL_11254595 | OW213860.1 | Dr. Risch Ostschweiz AG | Dr Risch Laboratory | Nadia Wohlwend et al |
| **hCoV-19/Jiangxi/IVDC-569/2022** | EPI_ISL_11873932 | | Jiangxi Provincial Center for Disease Control and Prevention | National Institute for Viral Disease Control and Prevention, China CDC | Tian Gong et al |
| **hCoV-19/Mexico/OAX-INMEGEN-56-333/2022** | EPI_ISL_9792182 | | SALUD DIGNA | Instituto Nacional de Medicina Genomica | Castro-Tarda A et al |
| **hCoV-19/Uruguay/DLSP_406/2022** | EPI_ISL_12532627 | | Departamento Laboratorios de Salud P.blica (DLSP) Ministerio de Salud P.blica | Departamento Laboratorios de Salud P.blica (DLSP) Ministerio de Salud P.blica | Cortinas et al |
| **hCoV-19/Colombia/ANT-CWOHC-VG-SEC3342/2022** | EPI_ISL_12572713 | | Universidad Nacional de Colombia - Laboratorio Genómico One Health | Universidad Nacional de Colombia - Laboratorio Genómico One Health | Isabel Moreno López et al |
| **hCoV-19/Panama/M223257-GMI/2022** | EPI_ISL_13105272 | | Gorgas Memorial Institute of Health Studies | Gorgas Memorial Institute of Health Studies | Claudia Gonzalez et al |
| **hCoV-19/Peru/JUN-INS-15613/2022** | EPI_ISL_14254425 | | Laboratorio de Referencia Nacional de Virus Respiratorios. Centro Nacional de Salud Publica. Instituto Nacional de Salud Peru. | Laboratorio de Referencia Nacional de Virus Inmunoprevenibles. Centro Nacional de Salud Publica. Instituto Nacional de Salud Peru. | Carlos Padilla Rojas et al |

| | | | | | |
|---|---|---|---|---|---|
| **hCoV-19/Mexico/TLA_InDRE_FB13724_E29116483248_S13130/2022** | EPI_ISL_12137693 | | LESP Tlaxcala | Instituto de Diagnostico y Referencia Epidemiologicos (INDRE) | Claudia Wong-Arambula et al |
| **hCoV-19/Mexico/MEX_CIAD-IMSS_202201045769/2022** | EPI_ISL_11816892 | | Laboratorio Central de Epidemiología (LCE) | Microbial Genomics Laboratory | Consorcio Mexicano de Vigilancia Genómica (CoViGen-Mex). Authors (in alphabetical order): Julio Elias Alvarado-Yaah et al |
| **hCoV-19/Venezuela/Voy1393/2022** | EPI_ISL_15165708 | | IVIC | Laboratorio de Virologia Molecular | Rossana C Jaspe et al |
| **hCoV-19/Guatemala/INC-LNS-272/2022** | EPI_ISL_12543187 | | Laboratorio Nacional de Salud (LNS), Guatemala | Inciensa, Instituto Costarricense de Investigación y Enseñanza en Nutrición y Salud | César Conde et al |
| **hCoV-19/Guatemala/4581-LNS/2022** | EPI_ISL_13081519 | | Laboratorio Nacional de Salud (LNS), Guatemala | Laboratorio Nacional de Salud (LNS), Guatemala | Mendoza L et al |
| **hCoV-19/Guatemala/4607-LNS/2022** | EPI_ISL_13081536 | | Laboratorio Nacional de Salud (LNS), Guatemala | Laboratorio Nacional de Salud (LNS), Guatemala | Mendoza L et al |
| **hCoV-19/Guatemala/46468-LNS/2022** | EPI_ISL_13096547 | | Laboratorio Nacional de Salud (LNS), Guatemala | Laboratorio Nacional de Salud (LNS), Guatemala | Mendoza L et al |
| **hCoV-19/Peru/CUS-INS-20576/2022** | EPI_ISL_13806348 | | Laboratorio de Referencia Regional de Cusco. Dirección Regional de Salud de Cusco. | Laboratorio de Referencia Nacional de Virus Respiratorios. Centro Nacional de Salud Publica. Instituto Nacional de Salud Peru. | José Santa Cruz Quispe et al |
| **hCoV-19/CzechRepublic/NAT-22-28272/2022** | EPI_ISL_14026714 | | Faculty Hospital Bulovka, Department of Clinical Microbiology | Charles University, Faculty of Science, BIOCEV, OMICS Genomics | Vladimír Hampl et al |
| **hCoV-19/USA/NM-CDC-LC0881717/2022** | EPI_ISL_15103092 | OP519166.1 | Laboratory Corporation of America | Centers for Disease Control and Prevention Division of Viral Diseases, Pathogen Discovery | Dakota Howard et al |
| **hCoV-19/PuertoRico/PR-CVL-010336/2022** | EPI_ISL_16360060 | | Laboratorio Clinico y Referencia M. Landron | PRCVSI | Lorraine D Rodriguez-Rivera et al |
| **hCoV-19/Venezuela/Mir093/2022** | EPI_ISL_15014583 | | IVIC | Laboratorio de Virologia Molecular | Rossana C Jaspe et al |
| **hCoV-19/ElSalvador/879-23-LNSP/2022** | EPI_ISL_18401353 | | Hospital ISSS Regional San Miguel SM | Laboratorio Nacional de Salud Publica | Laboratorio de Secuenciacion y Genomica del LNSP-INS et al |
| **hCoV-19/Peru/HUV-INS-31997/2022** | EPI_ISL_15806100 | | Laboratorio de Referencia Nacional de Virus Inmunoprevenibles. Centro Nacional de Salud Publica. Instituto Nacional de Salud Peru. | Laboratorio de Referencia Nacional de Virus Inmunoprevenibles. Centro Nacional de Salud Publica. Instituto Nacional de Salud Peru. | Carlos Padilla Rojas et al |
| **hCoV-19/Kazakhstan/496/2022** | EPI_ISL_13822823 | | National Center of Expertise in Nur-Sultan city | National center of Expertise, National center for Biotechnology | Abdaliyev Askar et al |
| **hCoV-19/Aruba/AW-RIVM-110218/2022** | EPI_ISL_14773666 | | National Institute for Public Health and the Environment (RIVM) | National Institute for Public Health and the Environment (RIVM) | Adam Meijer et al |
| **hCoV-19/HongKong/HK-HKPU-PUN525512090/2022** | EPI_ISL_15679986 | | Department of Health Technology and Informatics, The Hong Kong Polytechnic University | Department of Health Technology and Informatics, The Hong Kong Polytechnic University | Gilman Kit-Hang Siu et al |
| **hCoV-19/HongKong/HK-HKPU-PUN52062723A/2022** | EPI_ISL_15681011 | | Department of Health Technology and Informatics, The Hong Kong Polytechnic University | Department of Health Technology and Informatics, The Hong Kong Polytechnic University | Gilman Kit-Hang Siu et al |
| **hCoV-19/Aruba/AW-RIVM-106908/2022** | EPI_ISL_14239332 | | National Institute for Public Health and the Environment (RIVM) | National Institute for Public Health and the Environment (RIVM) | Adam Meijer et al |
| **hCoV-19/USA/IA-ASC-210776708/2022** | EPI_ISL_14953655 | | Aegis Sciences Corporation | Aegis Sciences Corporation | Cyndi Clark et al |
| **hCoV-19/Zhejiang/ZPCDCP-115/2022** | EPI_ISL_17261617 | | Zhejiang Provincial Center for Disease Control and Prevention | Zhejiang Provincial Center for Disease Control and Prevention | Yi Sun et al |
| **hCoV-19/Peru/LIM-INS-32082/2022** | EPI_ISL_15806185 | | Laboratorio de Referencia Nacional de Virus Inmunoprevenibles. Centro Nacional de Salud Publica. Instituto Nacional de Salud Peru. | Laboratorio de Referencia Nacional de Virus Inmunoprevenibles. Centro Nacional de Salud Publica. Instituto Nacional de Salud Peru. | Carlos Padilla Rojas et al |
| **hCoV-19/Belize/CML-802/2022** | EPI_ISL_18053246 | | Central Medical Laboratory | Central Medical Laboratory | Roberto Melendez et al |
| **hCoV-19/Mexico/OAX_LANGEBIO_IMSS_13722/2022** | EPI_ISL_16861616 | | Laboratorio Central de Epidemiologia | Unidad de Genomica Avanzada | Consorcio Mexicano de Vigilancia Genomica (CoViGen-Mex). Authors (in last name alphabetical order): Julio Elias Alvarado-Yaah et al |
| **hCoV-19/Taiwan/NCKUH-823622174375/2022** | EPI_ISL_15188010 | | Virology Lab,Department of Pathology, National Cheng Kung University Hospital | Virology Lab,Department of Pathology, National Cheng Kung University Hospital | Huey-Pin Tsai et al |
| **hCoV-19/Denmark/DCGC-598116/2022** | EPI_ISL_15453557 | | Department of Bacteria, Parasites and Fungi, Statens Serum Institut, Copenhagen, Denmark | Statens Serum Institut Bioinformatics and Microbial Genomics | Povilas Matusevicius on behalf of the Danish Covid-19 Genome Consortium et al |
| **hCoV-19/Chile/RM-133282/2022** | EPI_ISL_16641199 | | Genetica Molecular and Subdepartamento de Virologia ISP Chile | Instituto de Salud Publica de Chile | Mariela Guajardo et al |
| **hCoV-19/Poland/Kce-4489/2022** | EPI_ISL_14433838 | | Medyczne Laboratorium Diagnostyczne, Szpital Miejski nr 4 w Gliwicach Sp. z.o.o | Wojewódzka Stacja Sanitarno-Epidemiologiczna w Katowicach | Beata Rozwadowska et al |
| **hCoV-19/Bermuda/153710/2022** | EPI_ISL_14824145 | | Department of Health | CARPHA | Nikhella Winter on behalf of CARPHA et al |
| **hCoV-19/Slovakia/BB_22_00001798/2022** | EPI_ISL_13734485 | | Regional Public Health Authority with the seat in Banská Bystrica, Slovakia | Laboratory of Genomics and Bioinformatics, Comenius University Science Park | Lucia Maďarová et al |
| **hCoV-19/Russia/MOW-PMVL-DZ-25213125630/2022** | EPI_ISL_15865246 | | N.F. Gamaleya National Research Center for Epidemiology and Microbiology of the Ministry of Health of the Russian Federation / Moscow Healthcare Department | N.F. Gamaleya National Research Center for Epidemiology and Microbiology of the Ministry of Health of the Russian Federation / Moscow Healthcare Department | Vladimir A. Gushchin et al |
| **hCoV-19/Cameroon/LNSP-004-HCY-54708-CO19/2022** | EPI_ISL_18002341 | | NPHL-Cameroon | National Public Health Laboratory, Cameroon | Dr Essomba Rene Ghislain et al |
| **hCoV-19/Brazil/RJ-NVBS27132GENOV852182270715/2022** | EPI_ISL_16465534 | | DASA | DASA | Rodrigo Guarischi et al |
| **hCoV-19/Russia/BA-RII-MH125184S/2022** | EPI_ISL_16902734 | | Beloretsk Central District Clinical Hospital | WHO National Influenza Centre Russian Federation | Andrey Komissarov et al |
| **hCoV-19/Guatemala/8150-LNS/2022** | EPI_ISL_14524774 | | Laboratorio Nacional de Salud (LNS), Guatemala | Laboratorio Nacional de Salud (LNS), Guatemala | Mendoza L et al |
| **hCoV-19/Colombia/BOY-INS-VG-25426/2022** | EPI_ISL_15784988 | | CLINICA UNIVERSITARIA COLOMBIA | Instituto Nacional de Salud- Dirección de Investigación en Salud Pública | Katherine Laiton-Donato et al |
| **hCoV-19/Venezuela/MirLab7A/2022** | EPI_ISL_15014556 | | IVIC | Laboratorio de Virologia Molecular | Rossana C Jaspe et al |
| **hCoV-19/Honduras/FM-001/2022** | EPI_ISL_17562436 | | Laboratorio Nacional de Virologìa (SESAL),Honduras | Laboratorio Nacional de Virologìa (SESAL),Honduras | Dra. Soany Avilez et al |
| **hCoV-19/DominicanRepublic/2032655-LNSPDD/2022** | EPI_ISL_14151670 | | Laboratorio Nacional de Salud Pública Dr. Defilló | Laboratorio Nacional de Salud Pública Dr. Defilló | Isaac Miguel Sánchez et al |
| **hCoV-19/USA/NE-NM_L357260156/2022** | EPI_ISL_15288405 | | Nebraska Public Health Laboratory | NPHL COVID-19 Response Team | Paulino et al |
| **hCoV-19/Vietnam/PIHCM_2600/2022** | EPI_ISL_15058932 | | Pasteur Institute in Ho Chi Minh city | Department of Microbiology and Immunology | Nhung Vu Pham Hong et al |
| **hCoV-19/India/LA-INSACOG-NII-1196/2022** | EPI_ISL_14097805 | | COVID-19 Laboratory Chuchot SNM Hospital | National Institute of Immunology | Aneeshkumar Arimbasseri et al |
| **hCoV-19/Bulgaria/22BG-EU_021485_PI202/2022** | EPI_ISL_14889851 | | National Center of Infectious and Parasitic Diseases | National Center of Infectious and Parasitic Diseases | Alexiev et al |
| **hCoV-19/BosniaandHerzegovina/AGC3383/2022** | EPI_ISL_13804556 | | PZU SANASA | Alea Genetic Centre | Pecar D et al |
| **hCoV-19/Guangdong/GDCDC-22689/2022** | EPI_ISL_17495558 | | Guangdong Provincial Center for Disease Prevention and Control | Guangdong Provincial Center for Disease Prevention and Control | Liang et al |
| **hCoV-19/Hainan/SY332/2022** | EPI_ISL_15941044 | | Central & Clinical Laboratory of Sanya People's Hospital | Beijing Chao-Yang Hospital | Xiaoxia Wang et al |
| **hCoV-19/Peru/CUS-INS-26056/2022** | EPI_ISL_14816841 | | Laboratorio de Referencia Regional de Cusco. Dirección Regional de Salud de Cusco. | Laboratorio de Referencia Nacional de Virus Inmunoprevenibles. Centro Nacional de Salud Publica. Instituto Nacional de Salud Peru. | José Santa Cruz Quispe et al |
| **hCoV-19/Israel/SMC-7101406/2022** | EPI_ISL_14687100 | | Shamir Medical Center (Asaf Harofe) | Shamir Medical Center (Asaf Harofe) | Nir Rainy et al |
| **hCoV-19/CzechRepublic/BC_30082022_24/2022** | EPI_ISL_14846579 | | Nemocnice Strakonice | Institute of Parasitology, Biology Centre, CAS Branišovská 31, 37005, Českeké Budějovice, Czech | Martin Kolisko et al |
| **hCoV-19/Curacao/CW-RIVM-114105/2022** | EPI_ISL_15374717 | | National Institute for Public Health and the Environment (RIVM) | National Institute for Public Health and the Environment (RIVM) | Adam Meijer et al |
| **hCoV-19/Poland/WSSEGDA-1878/2022** | EPI_ISL_14624987 | | Uniwersyteckie Centrum Kliniczne | Wojewódzka Stacja Sanitarno-Epidemiologiczna w Gdańsku | Gabriela Rutkowska et al |
| **hCoV-19/Qatar/QA.QU_18.54.F4/2022** | EPI_ISL_16076632 | | Ministry of Public Health / Hamad Medical Corporation | Biomedical Research Center (BRC), Qatar University / Qatar Genome Project (QGP) | BRC: Heba A. Al-Khatib et al |
| **hCoV-19/NorthMacedonia/IPH-MKD-9460/2022** | EPI_ISL_13866711 | | Laboratory of Virology - Institute of Public Health Skopje, Macedonia | Laboratory of Virology - Institute of Public Health Skopje, Macedonia | Teodora Buzharova et al |
| **hCoV-19/Mexico/CHH_LANGEBIO_IMSS_9397/2022** | EPI_ISL_13803045 | | Centro de Investigacion Biomedica del Noreste (CIBIN) | Unidad de Genomica Avanzada | Consorcio Mexicano de Vigilancia Genomica (CoViGen-Mex). Authors (in alphabetical order): Julio Elias Alvarado-Yaah et al |
| **hCoV-19/Austria/LB-R00090-S110/2022** | EPI_ISL_14019456 | | Lifebrain Covid Labor GmbH | Lifebrain Covid Labor GmbH | Filip Sima et al |
| **hCoV-19/Romania/NIPH-215161/2022** | EPI_ISL_15812485 | | National Institute of Public Health Laboratory | National Institute of Public Health Laboratory | Mihaela Leustean et al |
| **hCoV-19/Germany/BW-RKI-I-985562/2022** | EPI_ISL_15148988 | | MVZ Labor Dr. Limbach & Kollegen GbR | Robert Koch Institute | ? |
| **hCoV-19/Mexico/GUA_CIAD-IMSS_202201023497/2022** | EPI_ISL_13983110 | | Centro de Investigacion Biomedica de Occidente (CIBO) | Microbial Genomics Laboratory | Consorcio Mexicano de Vigilancia Genómica (CoViGen-Mex). Authors (in alphabetical order): Julio Elias Alvarado-Yaah et al |
| **hCoV-19/Mongolia/GS-75734/2022** | EPI_ISL_14935776 | | National Center for Communicable Diseases (NCCD) National Influenza Center | National Center for Communicable Diseases (NCCD) National Influenza Center | Bayasgalan Namuutsetseg et al |
| **hCoV-19/Croatia/41659/2022** | EPI_ISL_14814632 | | Klinika za infektivne bolesti "Dr. Fran Mihaljević" | Hrvatski zavod za javno zdravstvo | Irena Tabain et al |
| **hCoV-19/NewZealand/22XA3529/2022** | EPI_ISL_15322540 | | LabPLUS | Institute of Environmental Science and Research (ESR), Kenepuru Science Centre | Sarah Cockerton et al |
| **hCoV-19/SriLanka/aicbu5942/2022** | EPI_ISL_14508350 | | Allergy, Immunology and Cell Biology Unit (AICBU) | Allergy, Immunology and Cell Biology Unit (AICBU) | Chandima Jeewandara et al |
| **hCoV-19/SriLanka/aicbu5934/2022** | EPI_ISL_14508342 | | Allergy, Immunology and Cell Biology Unit (AICBU) | Allergy, Immunology and Cell Biology Unit (AICBU) | Chandima Jeewandara et al |
| **hCoV-19/Albania/IPH-279413/2022** | EPI_ISL_17295695 | | Respiratory emmergent viruses laboratory, Institute of Public Health, | Respiratory emmergent viruses laboratory, Institute of Public Health, | Iris Hasibra et al |
| **hCoV-19/ElSalvador/893-23-LNSP/2022** | EPI_ISL_18401361 | | Unidad Médica ISSS La Unión | Laboratorio Nacional de Salud Publica | Laboratorio de Secuenciacion y Genomica del LNSP-INS et al |
| **hCoV-19/Belize/CML-743/2022** | EPI_ISL_15397432 | | Central Medical Laboratory | Central Medical Laboratory | Roberto Melendez et al |
| **hCoV-19/Japan/TKYkbm57894/2022** | EPI_ISL_16078997 | | Tokyo Metropolitan Institute of Public Health | Tokyo Metropolitan Institute of Public Health | Akane Negishi et al |
| **hCoV-19/Peru/PIU-INS-33724/2022** | EPI_ISL_15944625 | | Laboratorio de Referencia Regional de Piura. Dirección Regional de Salud de Piura. | Laboratorio de Referencia Nacional de Virus Inmunoprevenibles. Centro Nacional de Salud Publica. Instituto Nacional de Salud Peru. | Carlos Enrique Holguin Mauricci et al |
| **hCoV-19/Israel/SMC-7092466/2022** | EPI_ISL_14159641 | | Shamir Medical Center (Asaf Harofe) | Shamir Medical Center (Asaf Harofe) | Nir Rainy et al |
| **hCoV-19/France/NAQ-HCL722004200201/2022** | EPI_ISL_15809884 | | LAM BIOPOLE ORTHEZ | Maladies infectieuses, Hospices Civils de Lyon | Antonin Bal et al |
| **hCoV-19/Israel/SMC-7107025/2022** | EPI_ISL_15669708 | | Shamir Medical Center (Asaf Harofe) | Shamir Medical Center (Asaf Harofe) | Nir Rainy et al |
| **hCoV-19/UnitedArabEmirates/UZB_5482022071117/2022** | EPI_ISL_13840347 | | Microbiology Department - University Hospital Brussel | Microbiology Department - University Hospital Brussel | Thomas Demuyser et al |
| **hCoV-19/CaboVerde/FG-AG0102/2022** | EPI_ISL_14721357 | | Virology Laboratory of Fogo | Instituto Nacional de Saúde Pública (INSP) | Kevin Sanders et al |
| **hCoV-19/Greece/28231/2022** | EPI_ISL_14096207 | | National Public Health Organization | National Public Health Organization | Kyriaki Tryfinopoulou et al |
| **hCoV-19/USA/MT-MTPHL-4050245/2022** | EPI_ISL_14908201 | | Montana Public Health Laboratory | Montana Public Health Laboratory | Joy Ritter et al |
| **hCoV-19/USA/KS-CDC-QDX41380775/2022** | EPI_ISL_15217466 | OP567865.1 | Quest Diagnostics Incorporated | Centers for Disease Control and Prevention Division of Viral Diseases, Pathogen Discovery | Dakota Howard et al |
| **hCoV-19/Peru/LIM-INS-28031/2022** | EPI_ISL_15116045 | | Laboratorio de Referencia Nacional de Virus Inmunoprevenibles. Centro Nacional de Salud Publica. Instituto Nacional de Salud Peru. | Laboratorio de Referencia Nacional de Virus Inmunoprevenibles. Centro Nacional de Salud Publica. Instituto Nacional de Salud Peru. | Carlos Padilla Rojas et al |
| **hCoV-19/Angola/CERI-KRISP-K051244/2022** | EPI_ISL_16567227 | | Africa_CDC_Angola | CERI, Centre for Epidemic Response and Innovation, Stellenbosch University and KRISP, KZN Research Innovation and Sequencing Platform, UKZN. | Morais J et al |
| **hCoV-19/Bonaire/BQ-RIVM-112822/2022** | EPI_ISL_15216346 | | National Institute for Public Health and the Environment (RIVM) | National Institute for Public Health and the Environment (RIVM) | Adam Meijer et al |
| **hCoV-19/Mexico/MEX_IBT_IMSS_9742/2022** | EPI_ISL_15248566 | | Laboratorio Central de Epidemiología (LCE) | Instituto de Biotecnología de la UNAM | Consorcio Mexicano de Vigilancia Genómica (CoViGen-Mex). Authors (in alphabetical order): Julio Elias Alvarado-Yaah et al |
| **hCoV-19/Israel/SMC-7106341/2022** | EPI_ISL_15474141 | | Shamir Medical Center (Asaf Harofe) | Shamir Medical Center (Asaf Harofe) | Nir Rainy et al |
| **hCoV-19/CostaRica/INC-7689-772197/2022** | EPI_ISL_15708596 | | HOSPITAL DR. MAX PERALTA JIMENEZ | Inciensa, Instituto de Investigación y Enseñanza en Nutrición y Salud | Francisco Duarte et al |
| **hCoV-19/Paraguay/507493/2022** | EPI_ISL_15664980 | | Hospital Nacional de Itaugua | Laboratorio Central de Salud Pública | Sandra González et al |
| **hCoV-19/Zhejiang/ZPCDCP-94/2022** | EPI_ISL_17261596 | | Zhejiang Provincial Center for Disease Control and Prevention | Zhejiang Provincial Center for Disease Control and Prevention | Yi Sun et al |
| **hCoV-19/Singapore/11026/2022** | EPI_ISL_14810708 | | National Public Health Laboratory, National Centre for Infectious Diseases | National Public Health Laboratory, National Centre for Infectious Diseases | Grace Ngan et al |
| **hCoV-19/Australia/VIC-VIDRL-00003755/2022** | EPI_ISL_17827599 | | Melbourne Pathology | Victorian Infectious Disease Reference Laboratory | Aziz et al |
| **hCoV-19/Paraguay/546370/2022** | EPI_ISL_17632015 | | Laboratorio Hospital Barrio Obrero | Laboratorio Central de Salud Pública | Sandra González et al |
| **hCoV-19/Brazil/AM-FIOCRUZ-ILMD2205885/2022** | EPI_ISL_15603504 | | Laboratório Central de Saúde Pública do Amazonas - LACEN-AM | Laboratório de Ecologia de Doenças Transmissíveis na Amazônia, Instituto Leônidas e Maria Deane - Fiocruz Amazônia | Dejanane Silva et al |
| **hCoV-19/Brazil/AM-FIOCRUZ-ILMD2300112/2023** | EPI_ISL_17013701 | | Laboratório Central de Saúde Pública do Amazonas - LACEN-AM | Laboratório de Ecologia de Doenças Transmissíveis na Amazônia, Instituto Leônidas e Maria Deane - Fiocruz Amazônia | Dejanane Silva et al |
| **hCoV-19/CostaRica/INC-7435-771172/2022** | EPI_ISL_15415302 | | HOSPITAL SAN VICENTE DE PAUL | Inciensa, Instituto de Investigación y Enseñanza en Nutrición y Salud | Francisco Duarte et al |
| **hCoV-19/Zimbabwe/78107/2022** | EPI_ISL_16848176 | | National Microbiology Reference Laboratory | National Microbiology Reference Laboratory | Air Comodore Dr. J Chimedza et al |
| **hCoV-19/SouthAfrica/NICD-N51029/2022** | EPI_ISL_16473786 | | Dr George Mukhari Laboratory | National Institute for Communicable Diseases of the National Health Laboratory Service | Everatt J et al |
| **hCoV-19/Zimbabwe/78684/2023** | EPI_ISL_17324897 | | National Microbiology Reference Laboratory | National Microbiology Reference Laboratory | Air Comodore Dr. J Chimedza et al |
| **hCoV-19/Zimbabwe/78701/2023** | EPI_ISL_17324896 | | National Microbiology Reference Laboratory | National Microbiology Reference Laboratory | Air Comodore Dr. J Chimedza et al |
| **hCoV-19/Zimbabwe/78691/2022** | EPI_ISL_17324898 | | National Microbiology Reference Laboratory | National Microbiology Reference Laboratory | Air Comodore Dr. J Chimedza et al |
| **hCoV-19/Zimbabwe/78692/2022** | EPI_ISL_17324894 | | National Microbiology Reference Laboratory | National Microbiology Reference Laboratory | Air Comodore Dr. J Chimedza et al |
| **hCoV-19/Paraguay/528821/2022** | EPI_ISL_15572034 | | Laboratorio Hospital de Barrio Obrero | Laboratorio Central de Salud Pública | Sandra González et al |
| **hCoV-19/Georgia/Tb-s6646/2022** | EPI_ISL_16601472 | | Department for Virology, Molecular Biology and Genome Research, R. G. Lugar Center for Public Health Research, National Center for Disease Control and Public Health (NCDC) of Georgia. | Department for Virology, Molecular Biology and Genome Research, R. G. Lugar Center for Public Health Research, National Center for Disease Control and Public Health (NCDC) of Georgia. | Giorgi Tomashvili et al |
| **hCoV-19/Ukraine/71252/2022** | EPI_ISL_14503448 | | Reference laboratory for diagnostics of HIV/AIDS, virological and especially dangerous pathogens | Reference laboratory for diagnostics of HIV/AIDS, virological and especially dangerous pathogens | Iryna Demchyshyna et al |
| **hCoV-19/NewZealand/22YA2627/2022** | EPI_ISL_15277876 | | Pathlab Tauranga | Institute of Environmental Science and Research (ESR) | Sarah Cockerton et al |
| **hCoV-19/Botswana/R150B83_BHP_1061955/2022** | EPI_ISL_13766553 | | Botswana Harvard HIV Reference Laboratory | Botswana Harvard HIV Reference Laboratory | Sikhulile Moyo et al |
| **hCoV-19/Venezuela/Voy2055/2022** | EPI_ISL_13985221 | | IVIC | Laboratorio de Virologia Molecular | Rossana C Jaspe et al |
| **hCoV-19/Mexico/SON_InDRE_FB22015_E26319329701_S15313/2022** | EPI_ISL_15528548 | | LESP Sonora | Instituto de Diagnostico y Referencia Epidemiologicos (INDRE) | Claudia Wong-Arambula et al |
| **hCoV-19/USA/NV-CDC-QDX40313439/2022** | EPI_ISL_14742909 | OP337850.1 | Quest Diagnostics Incorporated | Centers for Disease Control and Prevention Division of Viral Diseases, Pathogen Discovery | Dakota Howard et al |
| **hCoV-19/Paraguay/522448/2022** | EPI_ISL_15571854 | | Centro Medico La Costa | Laboratorio Central de Salud Pública | Sandra González et al |
| **hCoV-19/Israel/ICH-741187594/2022** | EPI_ISL_15176377 | | Clinical Microbiology Laboratory, Tel Aviv Sourasky Medical Center | Clinical Microbiology Laboratory, Tel Aviv Sourasky Medical Center | Katya Levytskyi et al |
| **hCoV-19/Gabon/CERMEL-PP0455/2022** | EPI_ISL_14064982 | | Centre de Recherches Médicales de Lambaréné | Centre de Recherches Médicales de Lambaréné | Moustapha Nzamba Maloum et al |
| **hCoV-19/Gabon/CERMEL-PP0477/2022** | EPI_ISL_14064987 | | Centre de Recherches Médicales de Lambaréné | Centre de Recherches Médicales de Lambaréné | Moustapha Nzamba Maloum et al |
| **hCoV-19/Djibouti/NAMRU3_H038/2022** | EPI_ISL_15748318 | | Naval Medical Research Unit No. 3 | Naval Medical Research Center Biological Defense Research Directorate | J. Alexander Chitty et al |
| **hCoV-19/Finland/THL-202218220/2022** | EPI_ISL_14936696 | | Islab, Pohjois-Savon aluelaboratorio | Expert Microbiology, National Institute for Health and Welfare | Soile Blomqvist et al |
| **hCoV-19/Slovenia/235063/2022** | EPI_ISL_16234159 | | Institute of Microbiology and Immunology, Faculty of Medicine, University of Ljubljana | Institute of Microbiology and Immunology, Faculty of Medicine, University of Ljubljana | Alen Suljič et al |
| **hCoV-19/Canada/AB-ABPHL-90679/2023** | EPI_ISL_16706324 | | Alberta Precision Labs (APL) | Alberta Precision Labs (APL) | Buss E et al |
| **hCoV-19/India/GA-ICMR-NIV-G-12872/2022** | EPI_ISL_16353051 | | ICMR-National Institute of Virology - INSACOG | NIV Influenza | Dr. Varsha Potdar et al |
| **hCoV-19/Colombia/DC-INS-VG-25624/2022** | EPI_ISL_15826585 | | CLINICA UNIVERSITARIA COLOMBIA - LABORATORIO ESPECIALIZADO | Instituto Nacional de Salud- Dirección de Investigación en Salud Pública | Katherine Laiton-Donato et al |
| **hCoV-19/Poland/WSSEGorzow-23S0341/2023** | EPI_ISL_17859573 | | Laboratorium Analiz Lekarskich Alab Poznań (punkt Grudzieniec) | Wojewodzka Stacja Sanitarno-Epidemiologiczna w Gorzowie Wielkopolskim | Elzbieta Justynska et al |
| **hCoV-19/Aruba/AW-RIVM-121162/2022** | EPI_ISL_16296107 | | National Institute for Public Health and the Environment (RIVM) | National Institute for Public Health and the Environment (RIVM) | Adam Meijer et al |
| **hCoV-19/Israel/ICH-741194910/2022** | EPI_ISL_16395607 | | Clinical Microbiology Laboratory, Tel Aviv Sourasky Medical Center | Clinical Microbiology Laboratory, Tel Aviv Sourasky Medical Center | Katya Levytskyi et al |
| **hCoV-19/Romania/NIPH-218310/2023** | EPI_ISL_17699525 | | St Parascheva Iasi Clinical Hospital for Infectious Diseases | National Institute of Public Health Laboratory | Mihaela Leustean et al |

| | | | | | |
|---|---|---|---|---|---|
| **hCoV-19/Iceland/L-1896/2022** | EPI_ISL_15452975 | | Landspitali Department of Clinical Microbiology | Landspitali Department of Clinical Microbiology | Freyja Valsdottir et al |
| **hCoV-19/Djibouti/NAMRU3_H294/2022** | EPI_ISL_17612204 | | Naval Medical Research Unit No. 3 | Naval Medical Research Command Biological Defense Research Directorate | Shiva Rawat et al |
| **hCoV-19/Algeria/39419/2022** | EPI_ISL_15637961 | | NIC, Viral Respiratory Unit | NIC, Viral Respiratory Unit | Baya Hedjal et al |
| **hCoV-19/India/GJ-GBRC10081/2022** | EPI_ISL_16303696 | | Green Cross Genetics Lab Pvt Ltd, Ahmedabad | Gujarat Biotechnology Research Centre-INSACOG | Zarna Patel et al |
| **hCoV-19/Sweden/03_SE600_000200512541N2/2022** | EPI_ISL_16549441 | | Akademiska Sjukhuset, Clinical Microbiology and Hospital Hygiene | Akademiska Sjukhuset, Clinical Microbiology and Hospital Hygiene | Patrik Ellström et al |
| **hCoV-19/Paraguay/573595/2022** | EPI_ISL_17678222 | | IPS - Hospital Geriatrico Gerardo Boungermini | Laboratorio Central de Salud Pública | Sandra González et al |
| **hCoV-19/Finland/THL-01509/2023** | EPI_ISL_16816080 | | Huslab Finland | Expert Microbiology, National Institute for Health and Welfare | Erika Lindh et al |
| **hCoV-19/Greece/NPHO-4064/2023** | EPI_ISL_17668246 | | National Public Health Organization | National Public Health Organization | Kyriaki Tryfinopoulou et al |
| **hCoV-19/SouthKorea/KDCA211388/2023** | EPI_ISL_16902063 | | Division of Emerging Infectious Diseases, Bureau of Infectious Diseases Diagnosis Control, Korea Disease Control and Prevention Agency | Division of Emerging Infectious Diseases, Bureau of Infectious Diseases Diagnosis Control, Korea Disease Control and Prevention Agency | Jeong-Ah Kim et al |
| **hCoV-19/Venezuela/Voy4631/2022** | EPI_ISL_16958052 | | LabGenomik | Laboratorio de Virologia Molecular | Rossana C Jaspe et al |
| **hCoV-19/Colombia/DC-INS-VG-25628/2022** | EPI_ISL_15826589 | | CLINICA UNIVERSITARIA COLOMBIA - LABORATORIO ESPECIALIZADO | Instituto Nacional de Salud- Dirección de Investigación en Salud Pública | Katherine Laiton-Donato et al |
| **hCoV-19/USA/MT-MTPHL-4073195/2022** | EPI_ISL_16137492 | | Montana Public Health Laboratory | Montana Public Health Laboratory | Joy Ritter et al |
| **hCoV-19/CentralAfricanRepublic/LNBCSP111/2023** | EPI_ISL_18210115 | | Laboratoire National de Biologie Clinique et de Santé Publique | Laboratoire National de Biologie Clinique et de Santé Publique | POUNGINZA Simon et al |
| **hCoV-19/Canada/QC-L00566080001/2022** | EPI_ISL_16306878 | | Laboratoire de santé publique du Québec | Laboratoire de santé publique du Québec | Sandrine Moreira et al |
| **hCoV-19/Mongolia/UB-98946/2022** | EPI_ISL_15911440 | | National Center for Communicable Diseases (NCCD) National Influenza Center | National Center for Communicable Diseases (NCCD) National Influenza Center | Bayasgalan Namuutsetseg et al |
| **hCoV-19/Mongolia/UB-2893/2023** | EPI_ISL_17502971 | | National Center for Communicable Diseases (NCCD) National Influenza Center | National Center for Communicable Diseases (NCCD) National Influenza Center | Bayasgalan Namuutsetseg et al |
| **hCoV-19/Japan/TKYnat13438/2023** | EPI_ISL_17590238 | | Tokyo Metropolitan Institute of Public Health | Tokyo Metropolitan Institute of Public Health | Akane Negishi et al |
| **hCoV-19/Taiwan/NCKUH-061023000908/2023** | EPI_ISL_17280364 | | Virology Lab,Department of Pathology, National Cheng Kung University Hospital | Virology Lab,Department of Pathology, National Cheng Kung University Hospital | Huey-Pin Tsai et al |
| **hCoV-19/Ghana/WACCBIP-GS4267/2022** | EPI_ISL_16698913 | | WACCBIP, Volta Road, University of Ghana, Accra, Ghana | WACCBIP, Volta Road, University of Ghana, Accra, Ghana | Collins M. Morang'a et al |
| **hCoV-19/USA/NM-NMDOH-2022145242/2022** | EPI_ISL_16130103 | | Curative | New Mexico Department of Health Scientific Laboratory | Ellie Larence et al |
| **hCoV-19/Italy/FVG-PN-65342343/2022** | EPI_ISL_15814158 | | SC (UCO) Igiene e Sanità Pubblica, ASUGI, Trieste | ARGO Laboratorio Genomica ed Epigenomica | Licastro D et al |
| **hCoV-19/Georgia/Tb-sNCDC6598/2022** | EPI_ISL_16458644 | | Department for Virology, Molecular Biology and Genome Research, R. G. Lugar Center for Public Health Research, National Center for Disease Control and Public Health (NCDC) of Georgia. | Department for Virology, Molecular Biology and Genome Research, R. G. Lugar Center for Public Health Research, National Center for Disease Control and Public Health (NCDC) of Georgia. | Giorgi Tomashvili et al |
| **hCoV-19/Myanmar/DMR-22121/2023** | EPI_ISL_17162794 | | Department of Medical Research | Department of Medical Research | Myat Htut Nyunt et al |
| **hCoV-19/Myanmar/DMR-22115/2023** | EPI_ISL_17162789 | | Department of Medical Research | Department of Medical Research | Myat Htut Nyunt et al |
| **hCoV-19/Mexico/SIN_CIAD-MZT_S9036/2023** | EPI_ISL_17562430 | | Hospital Sharp | Microbial Genomics Laboratory | Julio Elías Alvarado-Yaah et al |
| **hCoV-19/DJI/NAMRU3_H726/2022** | EPI_ISL_17675099 | | Naval Medical Research Unit No. 3 | Naval Medical Research Command Biological Defense Research Directorate | Shiva Rawat et al |
| **hCoV-19/Paraguay/557924/2022** | EPI_ISL_17632309 | | Laboratorio Central de Salud Pública | Laboratorio Central de Salud Pública | Sandra González et al |
| **hCoV-19/Morocco/NIH-1260/2023** | EPI_ISL_18263156 | | NIC at NIH | National Institute of Hygiene | Hassan Ihazmade et al |
| **hCoV-19/Mali/CICM-28149/2022** | EPI_ISL_17401207 | | CICM, Bamako | Malaria Research and Training Center | Antoine Dara et al |
| **hCoV-19/Mali/CICM-27999/2022** | EPI_ISL_15508460 | | CICM, Bamako | Malaria Research and Training Center | Antoine Dara et al |
| **hCoV-19/England/BRBR-327128D6/2023** | EPI_ISL_17372842 | | Berkshire and Surrey Pathology Services Lighthouse Laboratory | Wellcome Sanger Institute for the COVID-19 Genomics UK (COG-UK) Consortium | Berkshire et al |
| **hCoV-19/CaboVerde/ST-AG0867/2022** | EPI_ISL_16761442 | | Virology Laboratory of Praia | Instituto Nacional de Saúde Pública (INSP) | Kevin Sanders et al |
| **hCoV-19/Togo/TOG-GOL-22-6007/2022** | EPI_ISL_17512187 | | Laboratoire National de Référence LNR/CHU-SO | Laboratoire de Biologie Moléculaire et d'Immunologie (BIOLIM/FSS/UL), Faculté des Sciences de la Santé, Université de Lomé | A.A. KONOU et al |
| **hCoV-19/Canada/QC-L00562884001/2022** | EPI_ISL_16190082 | | Laboratoire de santé publique du Québec | Laboratoire de santé publique du Québec | Sandrine Moreira et al |
| **hCoV-19/CostaRica/INC-10098-783636/2023** | EPI_ISL_17100593 | | HOSPITAL DR. FERNANDO ESCALANTE PRADILLA | Inciensa, Instituto de Investigación y Enseñanza en Nutrición y Salud | Francisco Duarte et al |
| **hCoV-19/CotedIvoire/IPCI-DVE-COV836316/2022** | EPI_ISL_15813179 | | Departement of Epidemic Viruses | Departement of Epidemic Viruses Institut Pasteur of CotedIvoire Abidjan | Hervé Albéric Adje Kadjo et al |
| **hCoV-19/Canada/SK-RRPL-606349/2023** | EPI_ISL_17271024 | | SK-Roy Romanow Provincial Laboratory | Saskatchewan - Roy Romanow Provincial Laboratory (RRPL) | Ryan McDonald et al |
| **hCoV-19/Colombia/DC-LSPSDS-336683/2022** | EPI_ISL_16026016 | | CONGLOMERADO - SUB RED NORTE | LSPSDS | Johana Hernandez Toloza et al |
| **hCoV-19/Peru/LIM-INS-32180/2022** | EPI_ISL_15806283 | | Laboratorio de Referencia Nacional de Virus Inmunoprevenibles. Centro Nacional de Salud Publica. Instituto Nacional de Salud Peru. | Laboratorio de Referencia Nacional de Virus Inmunoprevenibles. Centro Nacional de Salud Publica. Instituto Nacional de Salud Peru. | Carlos Padilla Rojas et al |
| **hCoV-19/Brazil/MG-NVBS25577GENOV101220868601/2022** | EPI_ISL_16297614 | | DASA | DASA | Rodrigo Guarischi et al |
| **hCoV-19/Brazil/SC-FIOCRUZ-269/2023** | EPI_ISL_16967226 | | Laboratorio Central de Saude Publica do Estado de Santa Catarina (LACEN/SC) | Laboratory of Respiratory Viruses and Measles, Oswaldo Cruz Institute, FIOCRUZ | Paola Resende et al |
| **hCoV-19/Spain/MD-HRYC-12612554/2023** | EPI_ISL_16831055 | | Servicio de Microbiologia Hospital Ramon y Cajal | Servicio de Microbiologia Hospital Ramon y Cajal | Ponce-Alonso M et al |
| **hCoV-19/Iceland/L-3288/2023** | EPI_ISL_16497499 | | Landspitali Department of Clinical Microbiology | Landspitali Department of Clinical Microbiology | Freyja Valsdottir et al |
| **hCoV-19/Palestine/526722/2023** | EPI_ISL_17485346 | | Molecular Diagnostic Laboraory (CPHL) - Palestinian Ministry of Health (MOH) | Molecular Diagnostic Laboraory-CPHL | Shtayeh Issa et al |
| **hCoV-19/Botswana/BHP-B03-3450p9017/2023** | EPI_ISL_17773021 | | Victus Global Botswana, Plot 132, Independence Avenue Road, Unit 3G and National Health laboratory, Gaborone | Botswana Harvard HIV Reference Laboratory | Chawangwa Modongo et al |
| **hCoV-19/Eswatini/N52540/2023** | EPI_ISL_17096029 | | Manzania | National Institute for Communicable Diseases of the National Health Laboratory Service | Maphalala G et al |
| **hCoV-19/Guinea/CERFIG-38144/2022** | EPI_ISL_15581447 | | Centre de recherche et de formation en infectiologie de Guinée | Centre de recherche et de formation en infectiologie de Guinée | Thibaut Armel Cherif GNIMADI et al |
| **hCoV-19/Malaysia/UiTM2454/2023** | EPI_ISL_17039356 | | Hospital Raja Permaisuri Bainun | iPROMISE, UiTM | Mohd Zaki Salleh et al |
| **hCoV-19/NewZealand/22XA5799/2022** | EPI_ISL_16050926 | | Waikato DHB Laboratory | Institute of Environmental Science and Research (ESR) | Sarah Cockerton et al |
| **hCoV-19/Coted-Ivoire/IPCI-DVE-E750265/2023** | EPI_ISL_16894718 | | Departement of Epidemic Viruses | Departement of Epidemic Viruses Institut Pasteur of Cote d'Ivoire Abidjan | Hervé Albéric Adje Kadjo et al |
| **hCoV-19/Mali/CICM-216/2022** | EPI_ISL_17401212 | | CICM, Bamako | Malaria Research and Training Center | Antoine Dara et al |
| **hCoV-19/Colombia/DC-LSPSDS-349752/2022** | EPI_ISL_16428496 | | MUESTRA CONGLOMERADO - SUB RED SUR OCCIDENTE | LSPSDS | Johana Hernandez Toloza et al |
| **hCoV-19/Mexico/VER_LANGEBIO_IMSS_0722/2022** | EPI_ISL_16204989 | | Laboratorio Central de Epidemiologia (LCE) | Unidad de Genomica Avanzada | Consorcio Mexicano de Vigilancia Genomica (CoViGen-Mex). Authors (in last name alphabetical order): Julio Elias Alvarado-Yaah et al |
| **hCoV-19/Uruguay/DLSP_3958/2022** | EPI_ISL_16378325 | | Departamento Laboratorios de Salud Pública (DLSP) Ministerio de Salud Pública | Departamento Laboratorios de Salud Pública (DLSP) Ministerio de Salud Pública | Cortinas et al |
| **hCoV-19/Uruguay/DLSP_MU86480/2023** | EPI_ISL_18041146 | | Departamento Laboratorios de Salud Pública (DLSP) Ministerio de Salud Pública | Departamento Laboratorios de Salud Pública (DLSP) Ministerio de Salud Pública | Cortinas et al |
| **hCoV-19/Kuwait/AH-USAFSAM-S21266/2023** | EPI_ISL_17250749 | | US Air Force School of Aerospace Medicine | US Air Force School of Aerospace Medicine | Anthony Fries et al |
| **hCoV-19/Romania/IS-28357/2022** | EPI_ISL_17540844 | | Department of Public Health Iasi | National Institute of Infectious Diseases-Prof. Dr. Matei Bals Molecular Diagnostics Laboratory | Corina Casangiu et al |
| **hCoV-19/Romania/MS-1503/2023** | EPI_ISL_18261639 | | Department of Public Health Mures | National Institute of Infectious Diseases-Prof. Dr. Matei Bals Molecular Diagnostics Laboratory | Simona Paraschiv et al |
| **hCoV-19/Spain/AB-HGUA-60664041/2023** | EPI_ISL_16925973 | | Hospital General Universitario Albacete | Hospital General Universitario Albacete | Lorena Robles-Fonseca et al |
| **hCoV-19/NorthMacedonia/IPH-MKD-134R/2023** | EPI_ISL_18216601 | | Laboratory of Virology - Institute of Public Health Skopje, Macedonia | Laboratory of Virology - Institute of Public Health Skopje, Macedonia | Teodora Buzharova et al |
| **hCoV-19/USA/ME-CDC-VSX-A064811/2023** | EPI_ISL_17278604 | | Helix | Centers for Disease Control and Prevention Division of Viral Diseases, Pathogen Discovery | Dakota Howard et al |
| **hCoV-19/Paraguay/548874/2022** | EPI_ISL_17632079 | | Instituto de Medicina Tropical | Laboratorio Central de Salud Pública | Sandra González et al |
| **hCoV-19/Canada/un-KVI300536/2023** | EPI_ISL_17013265 | | TIEHH Bioterrorism Response Laboratory | TIEHH Bioterrorism Response Laboratory | Adriana Estrada et al |
| **hCoV-19/India/KA-ICMR-VRDL-MYS-81721-JSS/2022** | EPI_ISL_16199601 | | VRDL-MMC&RI-INSACOG | NIV Influenza | Dr. Amrutha Kumari et al |
| **hCoV-19/Egypt/CPHL-EGY22709/2022** | EPI_ISL_16299818 | | CPHL,MOH,EGYPT | CPHL | Wael H. Roshdy et al |
| **hCoV-19/Japan/PG-544657/2023** | EPI_ISL_18066127 | | Medical Microbiology and Zoology Section, Okinawa Prefectural Institute of Health and Environment | Medical Microbiology and Zoology Section, Okinawa Prefectural Institute of Health and Environment | Tetsuya Kakita et al |
| **hCoV-19/Congo/FCRM-203-17837/2022** | EPI_ISL_15334249 | | Fondation Congolaise pour la recherche medicale (FCRM), Francine Ntoumi | Fondation Congolaise pour la Recherche Médicale | Mfoutou Mapanguy Claujens Chastel et al |
| **hCoV-19/DRC/INRB-RDC-1352/2022** | EPI_ISL_15689158 | | Pathogen Genomics Lab, National Institute for Biomedical Research (INRB) | Pathogen Genomics Lab, National Institute for Biomedical Research (INRB) | Placide Mbala-Kingebeni et al B |
| **hCoV-19/DRC/INRB-RDC-1356/2022** | EPI_ISL_15689157 | | Pathogen Genomics Lab, National Institute for Biomedical Research (INRB) | Pathogen Genomics Lab, National Institute for Biomedical Research (INRB) | Placide Mbala-Kingebeni et al B |
| **hCoV-19/Japan/PG-441538/2023** | EPI_ISL_16825812 | | Pathogen Genomics Center, National Institute of Infectious Diseases | Pathogen Genomics Center, National Institute of Infectious Diseases | Tsuyoshi Sekizuka et al B |
| **hCoV-19/Cameroon/V-25321/2022** | EPI_ISL_18352134 | | CENTRE PASTEUR DU CAMEROUN | CENTRE PASTEUR DU CAMEROUN | Njouom Richard et al |
| **hCoV-19/CentralAfricanRepublic/LNBCSP022/2022** | EPI_ISL_17423776 | | Laboratoire National de Biologie Clinique et de Santé Publique | Laboratoire National de Biologie Clinique et de Santé Publique | Simplice SOMBOT NDICKY et al |
| **hCoV-19/Ecuador/NIC-INSPI-8119971/2022** | EPI_ISL_16579063 | | INSPI-CRN DE INFLUENZA Y OTROS VIRUS RESPIRATORIOS-INTERLAB | INSPI-CRN DE INFLUENZA Y OTROS VIRUS RESPIRATORIOS | Alfredo Bruno et al |
| **hCoV-19/Nigeria/NCDC-NR-GL-0041/2023** | EPI_ISL_17038852 | | National Reference Laboratory, Nigeria Centre for Disease Control | National Reference Laboratory, Nigeria Centre for Disease Control | Dr Ndodo Nnaemeka et al |
| **hCoV-19/Brazil/ES-LACENES-321345931/2023** | EPI_ISL_16816223 | | Central Laboratory of Public Health of the Espirito Santo - Lacen ES | Central Laboratory of Public Health of the Espirito Santo - Lacen ES | Felipe Donateli Gatti et al |
| **hCoV-19/Italy/VEN-IZSVe-22RS8399-5_VR/2022** | EPI_ISL_16235627 | | AOUIVR | Istituto Zooprofilattico Sperimentale delle Venezie | Adelaide Milani et al |
| **hCoV-19/India/AS-NEIST-R41-CID2165/2022** | EPI_ISL_16414094 | | BARUA CLINICAL LABORATORY, Jorhat | Centre for Infectious Diseases, CSIR-NEIST | Emon Kalyan et al |
| **hCoV-19/Tunisia/G-4445/2023** | EPI_ISL_17383736 | | Laboratoire de Virologie Clinique, Institut Pasteur de Tunis | Laboratoire de Virologie Clinique, Institut Pasteur de Tunis | Samar Ben Halima et al |
| **hCoV-19/Germany/SN-RKI-I-1131655/2023** | EPI_ISL_17518671 | | Bioscientia Labor Wermsdorf | Robert Koch Institute | ? |
| **hCoV-19/Bahrain/CDL-770104517/2022** | EPI_ISL_17081897 | | Communicable Disease Laboratory | Communicable Disease Laboratory | Marhoon et al |
| **hCoV-19/Jiangsu/CLHQ-001/2022** | EPI_ISL_16137907 | | Center Laboratory of Health Quarantine, Nanjing Customs | Center Laboratory of Health Quarantine, Nanjing Customs | Geng et al |
| **hCoV-19/Jiangsu/JITHCC_159/2022** | EPI_ISL_16127647 | | Center Laboratory of Health Quarantine, Nanjing Customs District P.R. China | Center Laboratory of Health Quarantine, Nanjing Customs District P.R. China | Geng et al |
| **hCoV-19/Argentina/INEI123755/2022** | EPI_ISL_17800140 | | Servicio Virosis Respiratorias-Departamento Virología-INEI C.G.Malbran | Instituto Nacional Enfermedades Infecciosas C.G.Malbran | Baumeister E. et al |
| **hCoV-19/Austria/AGES-1166778/2022** | EPI_ISL_16551051 | | Covid Fighters | AGES IMED Vienna | Lippert K. et al |
| **hCoV-19/Namibia/NIP0054/2022** | EPI_ISL_17018670 | | NIP_Windhoek_Reference laboratory | NIP_Windhoek_Reference laboratory | Iyaloo Konstantinus et al |
| **hCoV-19/Panama/M266260-GMI/2022** | EPI_ISL_16018892 | | Gorgas Memorial Institute of Health Studies | Gorgas Memorial Institute of Health Studies | Claudia Gonzalez et al |
| **hCoV-19/Brazil/SP-NVBS27932GENOV852285635480/2022** | EPI_ISL_16618733 | | DASA | DASA | Rodrigo Guarischi et al |
| **hCoV-19/NorthernIreland/NIRE-01fbc4/2022** | EPI_ISL_16391949 | | Regional Virus Laboratory, Belfast Health and Social Care Trust; and: Genomics Core Technology Unit, Queen's University Belfast. | COVID-19 Genomics UK (COG-UK) Consortium | [Regional Virus Laboratory et al |
| **hCoV-19/Estonia/32191/2023** | EPI_ISL_18944789 | | West Tallinn Central Hospital | The Laboratory of Communicable Diseases, Republic of Estonia Health Board | Janne Pullat et al |
| **hCoV-19/England/BRBR-326B3FE2/2023** | EPI_ISL_16879805 | | Berkshire and Surrey Pathology Services Lighthouse Laboratory | Wellcome Sanger Institute for the COVID-19 Genomics UK (COG-UK) Consortium | Berkshire et al |
| **hCoV-19/CaboVerde/SV0096/2023** | EPI_ISL_17525391 | | Virology Laboratory of São Vicente | Instituto Nacional de Saúde Pública (INSP) | Kevin Sanders et al |
| **hCoV-19/Uruguay/DLSP_0118/2023** | EPI_ISL_16650526 | | Departamento Laboratorios de Salud Pública (DLSP) Ministerio de Salud Pública | Departamento Laboratorios de Salud Pública (DLSP) Ministerio de Salud Pública | Cortinas et al |
| **hCoV-19/Italy/APU_UNIBA_BA918/2023** | EPI_ISL_16585303 | | Laboratorio di Epidemiologia Molecolare e Sanità Pubblica-Policlinico Bari | University of Bari Biomedical Sciences and Human Oncology | Chironna Maria et al |
| **hCoV-19/Hungary/100829-UPL23-000706/2023** | EPI_ISL_17347311 | | Institute of Laboratory Medicine, Semmelweis University and Epidemiology and Surveillance Centre, Semmelweis University | iBioscience Kft. | Gergő Túri et al |
| **hCoV-19/Argentina/22110-183/2023** | EPI_ISL_17517805 | | Laboratorio Central, Ministerio de Salud Córdoba | Departamento Laboratorio Central and Unidad Operativa Centro Nacional de Genómica y Bioinformática | Castro et al |
| **hCoV-19/Greece/NPHO-4809/2023** | EPI_ISL_17667877 | | National Public Health Organization | National Public Health Organization | Kyriaki Tryfinopoulou et al |
| **hCoV-19/Switzerland/VD-CHUV-GEN16463/2023** | EPI_ISL_17497452 | | Hôpital Cantonal de Fribourg - HFR | Laboratory of genomics and metagenomics, Institute of Microbiology, University Hospital Centre and University of Lausanne | Trestan Pillonel et al |
| **hCoV-19/Eswatini/N47835/2022** | EPI_ISL_14501279 | | Mbabane Government Hospital | National Institute for Communicable Diseases of the National Health Laboratory Service | Maphalala G et al |
| **hCoV-19/Ethiopia/CO_GX_438/2022** | EPI_ISL_17616427 | | Ethiopian Public Health Institute | EPHI Pathogen Genomic Sequence Lab | Woldaregay Erku Abegaz et al |
| **hCoV-19/Ethiopia/CO_GX_440/2022** | EPI_ISL_17615497 | | Ethiopian Public Health Institute | EPHI Pathogen Genomic Sequence Lab | Woldaregay Erku Abegaz et al |
| **hCoV-19/Eswatini/N52466/2022** | EPI_ISL_17096012 | | Good Sherperd Hospital | National Institute for Communicable Diseases of the National Health Laboratory Service | Maphalala G et al |
| **hCoV-19/Zimbabwe/77855/2022** | EPI_ISL_16724339 | | National Microbiology Reference Laboratory | National Microbiology Reference Laboratory | Air Comodore Dr. J Chimedza et al |
| **hCoV-19/Chile/AN-130190/2022** | EPI_ISL_16158142 | | Genetica Molecular and Subdepartamento de Virologia ISP Chile | Instituto de Salud Publica de Chile | Constanza Campano et al |
| **hCoV-19/Japan/PG-411061/2022** | EPI_ISL_16209935 | | Chiba Prefectural Institute of Public Health | Division of Virology and Zoology, Chiba Prefectural Institute of Public Health | Yuki Fujinuma et al |
| **hCoV-19/Georgia/Tb-s6609/2022** | EPI_ISL_16601451 | | Department for Virology, Molecular Biology and Genome Research, R. G. Lugar Center for Public Health Research, National Center for Disease Control and Public Health (NCDC) of Georgia. | Department for Virology, Molecular Biology and Genome Research, R. G. Lugar Center for Public Health Research, National Center for Disease Control and Public Health (NCDC) of Georgia. | Giorgi Tomashvili et al |
| **hCoV-19/USA/SD-CDC-2-6374936/2022** | EPI_ISL_15655657 | | SD Public Health Laboratory | Centers for Disease Control and Prevention Division of Viral Diseases, Pathogen Discovery | Mili Sheth et al |
| **hCoV-19/Belize/CML-808/2022** | EPI_ISL_18053252 | | Central Medical Laboratory | Central Medical Laboratory | Roberto Melendez et al |
| **hCoV-19/HongKong/HK-HKPU-PUN521794490/2022** | EPI_ISL_17581690 | | Department of Health Technology and Informatics, The Hong Kong Polytechnic University | Department of Health Technology and Informatics, The Hong Kong Polytechnic University | Gilman Kit-Hang Siu et al |
| **hCoV-19/Austria/AGES-1166741/2022** | EPI_ISL_16551038 | | Covid Fighters | AGES IMED Vienna | Lippert K. et al |
| **hCoV-19/Tianjin/TJCDC-0958/2022** | EPI_ISL_16922433 | | Tianjin Centers for Disease Control and Prevention | Tianjin Centers for Disease Control and Prevention | Li Xiaoyan et al |
| **hCoV-19/Brunei/7122130898/2022** | EPI_ISL_14290744 | | National Virology Reference Laboratory | Microbial Genomic Services | Zainun Zaini et al |
| **hCoV-19/Guangxi/GCDC-BS-2211/2022** | EPI_ISL_17104933 | | Baise Center for Disease Control and Prevention | Guangxi Zhuang Autonomous Region Center for Disease Control and Prevention | Zan Chen et al |
| **hCoV-19/Mexico/NAY_CIAD-IMSS_01032264/2022** | EPI_ISL_14901739 | | Centro de Investigacion Biomedica de Occidente (CIBO) | Microbial Genomics Laboratory | Consorcio Mexicano de Vigilancia Gen√≥mica (CoViGen-Mex). Authors (in alphabetical order): Julio Elias Alvarado-Yaah et al |
| **hCoV-19/Brazil/SP-NVBS23314GENOV829651816323/2022** | EPI_ISL_15096296 | | DASA | DASA | Rodrigo Guarischi et al |
| **hCoV-19/Australia/NSW-ICPMR-32339/2022** | EPI_ISL_14617392 | | Pathology West - NSW Health Pathology | Institute Of Clinical Pathology And Medical Research (ICPMR), Westmead Hospital | Arnott A. et al |
| **hCoV-19/CostaRica/INC-7021-768239/2022** | EPI_ISL_15125118 | | AREA DE SALUD CARRILLO | Inciensa, Instituto de Investigación y Enseñanza en Nutrición y Salud | Francisco Duarte et al |
| **hCoV-19/Chongqing/YBCDC-0208/2022** | EPI_ISL_17197581 | | Chongqing Yubei District Center for Disease Control and Prevention | Chongqing Yubei District Center for Disease Control and Prevention | Jin Yan et al |
| **hCoV-19/Angola/CERI-KRISP-K050480/2022** | EPI_ISL_16311878 | | Africa_CDC_Angola | CERI, Centre for Epidemic Response and Innovation, Stellenbosch University and KRISP, KZN Research Innovation and Sequencing Platform, UKZN. | Morais J et al |
| **hCoV-19/India/TG-CDFD-SR-169/2022** | EPI_ISL_15153398 | | Apollo Hospital | CDFD-INSACOG | Asmita Gupta et al |

| | | | | | |
|---|---|---|---|---|---|
| **hCoV-19/India/un-134-ALP-10868/2022** | EPI_ISL_14018294 | | LMMD, RGCB | LMMD-RGCB-INSACOG | Dr Radha Krishnan Nair et al |
| **hCoV-19/Qatar/QA.QU_18.58.E1/2022** | EPI_ISL_16076856 | | Ministry of Public Health / Hamad Medical Corporation | Biomedical Research Center (BRC), Qatar University / Qatar Genome Project (QGP) | BRC: Heba A. Al-Khatib et al |
| **hCoV-19/Kenya/ILRI_COVM04374/2022** | EPI_ISL_15059450 | | International Livestock Research Institute | International Livestock Research Institute | Samuel O. Oyola et al |
| **hCoV-19/Georgia/Tb-s5396/2022** | EPI_ISL_14437527 | | JSC Megalab | Department for Virology, Molecular Biology and Genome Research, R. G. Lugar Center for Public Health Research, National Center for Disease Control and Public Health (NCDC) of Georgia. | Giorgi Tomashvili et al |
| **hCoV-19/Bahrain/Bahrain/R15902413496/2022** | EPI_ISL_15252380 | ON848962.1 | Communicable Disease Laboratory, Public Health Directorate | Communicable Disease Laboratory, Public Health Directorate | Marhoon et al |
| **hCoV-19/Philippines/PH-RITM-1453/2022** | EPI_ISL_13304974 | | BSL-SNRL | Research Institute for Tropical Medicine | Francisco Gerardo M. Polotan et al |
| **hCoV-19/Kenya/C117983/2022** | EPI_ISL_13988831 | OP037110.1 | KEMRI-Wellcome Trust Research Programme,Kilifi | KEMRI-Wellcome Trust Research Programme,Kilifi | Githinji G. et al |
| **hCoV-19/SriLanka/aicbu6160/2022** | EPI_ISL_15903990 | | Allergy, Immunology and Cell Biology Unit (AICBU) | Allergy, Immunology and Cell Biology Unit (AICBU) | Chandima Jeewandara et al |
| **hCoV-19/Guangdong/GPCDCP-5/2022** | EPI_ISL_16127645 | | Guangdong Provincial Center for Disease Control and Prevention | Key Laboratory of Human Diseases Comparative Medicine, Institute of Laboratory Animal Science, CAMS&PUMC | Bao et al |
| **hCoV-19/India/GJ-INSACOG-GBRC8265/2022** | EPI_ISL_14436859 | | Supratech Micropath Diagnostics & Research laborat | Gujarat Biotechnology Research Centre | Dinesh Kumar et al |
| **hCoV-19/India/TN-CDFD-A-234/2022** | EPI_ISL_14796261 | | Directorate of Public Health and Preventive Medicine | CDFD-INSACOG | Asmita Gupta et al |
| **hCoV-19/Yunnan/SR01/2022** | EPI_ISL_13867424 | | Department of Acute Infectious Diseases Control and Prevention, Yunnan Provincial Center for Disease Control and Prevention | Department of Acute Infectious Diseases Control and Prevention, Yunnan Provincial Center for Disease Control and Prevention | Meiling Zhang et al |
| **hCoV-19/Brunei/7122146678/2022** | EPI_ISL_14987984 | | National Virology Reference Laboratory | Microbial Genomic Services | Zainun Zaini et al |
| **hCoV-19/Djibouti/NAMRU3_F684/2022** | EPI_ISL_15091093 | OP508861.1 | Naval Medical Research Unit No. 3 | Naval Medical Research Center Biological Defense Research Directorate | J. Alexander Chitty et al |
| **hCoV-19/Qatar/QA.QU_18.51.A9/2022** | EPI_ISL_16076565 | | Ministry of Public Health / Hamad Medical Corporation | Biomedical Research Center (BRC), Qatar University / Qatar Genome Project (QGP) | BRC: Heba A. Al-Khatib et al |
| **hCoV-19/Beijing/BJCDC-1897/2022** | EPI_ISL_16634011 | | Beijing Center for Disease Control and Prevention | Beijing Center for Disease Control and Prevention | Shujuan Cui et al |
| **hCoV-19/Israel/ICH-741179413/2022** | EPI_ISL_14248293 | | Clinical Microbiology Laboratory, Tel Aviv Sourasky Medical Center | Clinical Microbiology Laboratory, Tel Aviv Sourasky Medical Center | Katya Levytskyi et al |
| **hCoV-19/Germany/NW-RKI-I-920345/2022** | EPI_ISL_14395001 | | Eurofins MVZ Labor Gelsenkirchen | Robert Koch Institute | ? |
| **hCoV-19/Israel/ICH-741183510/2022** | EPI_ISL_14458261 | | Clinical Microbiology Laboratory, Tel Aviv Sourasky Medical Center | Clinical Microbiology Laboratory, Tel Aviv Sourasky Medical Center | Katya Levytskyi et al |
| **hCoV-19/USA/MS-UMMC-M990C5-537882/2022** | EPI_ISL_15389768 | | University of Mississippi Medical Center, Department of Pathology | University of Mississippi Medical Center, Molecular and Genomics Core Facility | Raima Sen et al |
| **hCoV-19/Shanghai/SJTU-234013/2022** | EPI_ISL_16327349 | | Shanghai Public Health Clinical Center, Fudan University | Shanghai Institute of Hematology, National Research Center for Translational Medicine, State Key Laboratory of Medical Genomics, Ruijin Hospital Affiliated to Shanghai Jiao Tong University (SJTU) School of Medicine | Gang Lu et al |
| **hCoV-19/SouthAfrica/NICD-N45612/2022** | EPI_ISL_14223622 | | ROB FERREIRA LABORATORY | National Institute for Communicable Diseases of the National Health Laboratory Service | Amoako DG et al |
| **hCoV-19/Denmark/DCGC-592619/2022** | EPI_ISL_15347092 | | Department of Bacteria, Parasites and Fungi, Statens Serum Institut, Copenhagen, Denmark | Statens Serum Institut Bioinformatics and Microbial Genomics | Povilas Matusevicius on behalf of the Danish Covid-19 Genome Consortium et al |
| **hCoV-19/Tianjin/TJCDC-1356/2022** | EPI_ISL_16922832 | | Tianjin Centers for Disease Control and Prevention | Tianjin Centers for Disease Control and Prevention | Li Xiaoyan et al |
| **hCoV-19/Belize/GMI-9321491/2022** | EPI_ISL_16921720 | | Central Medical Laboratory | Gorgas Memorial Institute of Health Studies | Aldo Sosa et al |
| **hCoV-19/Palau/PW-H2318953/2022** | EPI_ISL_16557524 | | Laboratory Services, Palau Ministry of Health | State Laboratories Division, Hawaii State Department of Health | Pamela O'Brien et al |
| **hCoV-19/Spain/GA-CHUVI-22001918/2022** | EPI_ISL_15834254 | | Microbiology Department, Complexo Hospitalario Universitario de Vigo | Microbiology Department, Complexo Hospitalario Universitario de Vigo | Cabrera JJ et al |
| **hCoV-19/USA/CO-CDPHE-2103460591/2022** | EPI_ISL_14286725 | OP268924.1 | Colorado Department of Public Health and Environment | Colorado Department of Public Health and Environment | Laura Bankers et al |
| **hCoV-19/Vietnam/NHTD-OUCRU3533/2022** | EPI_ISL_15887623 | | National Hospital for Tropical Diseases | Oxford University Clinical Research Unit, Hanoi, Vietnam | Pham Ngoc Thach et al |
| **hCoV-19/Montenegro/CO-00931_MNE000_2379302202/2022** | EPI_ISL_14840206 | | Laboratory for Molecular Diagnostics,IPHMN | Eurofins Genomics Europe Sequencing GmbH | Danijela Vujošević et al |
| **hCoV-19/Brunei/7722005479/2022** | EPI_ISL_13602543 | | National Virology Reference Laboratory | Microbial Genomic Services | Zainun Zaini et al |
| **hCoV-19/NewZealand/22YA1871/2022** | EPI_ISL_14634412 | | Wellington SCL | Institute of Environmental Science and Research (ESR) | Sarah Cockerton et al |
| **hCoV-19/Indonesia/JI-ITD-58848NT/2022** | EPI_ISL_15686528 | | Premier Hospital Surabaya | Institute of Tropical Disease, Universitas Airlangga | Aldise M Nastri et al |
| **hCoV-19/India/GA-ICMR-NIV-INSACOG-GSEQ-11657/2022** | EPI_ISL_14149536 | | ICMR-National Institute of Virology - INSACOG | NIV Influenza | Dr. Varsha Potdar et al |
| **hCoV-19/Mexico/CMX_InDRE_FB22091_E09319412618_S15352/2022** | EPI_ISL_15528576 | | LESP Mexico City | Instituto de Diagnostico y Referencia Epidemiologicos (INDRE) | Claudia Wong-Arambula et al |
| **hCoV-19/USA/NE-NPHL22-18532/2022** | EPI_ISL_14485986 | | Nebraska Public Health Laboratory | NPHL COVID-19 Response Team | Paulino et al |
| **hCoV-19/Switzerland/BE-ETHZ-37816218/2022** | EPI_ISL_13700917 | OX181283.1 | Viollier AG | Department of Biosystems Science and Engineering, ETH Zürich | Andrea Patrizia Salzmann et al |
| **hCoV-19/Moldova/un-ChVir29215/2022** | EPI_ISL_14295892 | | National Agency for Public Health , Ministry of Health of the Republic of Moldova | Charité Universitätsmedizin Berlin, Institut für Virologie | Victor M Corman et al |
| **hCoV-19/NewZealand/22YA1822/2022** | EPI_ISL_14637291 | | Waikato DHB Laboratory | Institute of Environmental Science and Research (ESR) | Sarah Cockerton et al |
| **hCoV-19/NewZealand/22YA3062/2022** | EPI_ISL_15276710 | | Taranaki Pathology Services | Institute of Environmental Science and Research (ESR) | Sarah Cockerton et al |
| **hCoV-19/SouthKorea/KDCA108085/2022** | EPI_ISL_14784888 | | Division of Emerging Infectious Diseases, Bureau of Infectious Diseases Diagnosis Control, Korea Disease Control and Prevention Agency | Division of Emerging Infectious Diseases, Bureau of Infectious Diseases Diagnosis Control, Korea Disease Control and Prevention Agency | Jeong-Ah Kim et al |
| **hCoV-19/USA/UT-UPHL-221118938164/2022** | EPI_ISL_15870428 | | Utah Public Health Laboratory | Utah Public Health Laboratory | Erin L. Young et al |
| **hCoV-19/Shandong/WCDC-168/2022** | EPI_ISL_18070500 | | Weihai Center for Disease Control and Prevention | Weihai Center for Disease Control and Prevention | Xiang Li et al |
| **hCoV-19/NewZealand/22YA3867/2022** | EPI_ISL_15957549 | | Medlab Central (PN) | Institute of Environmental Science and Research (ESR) | Sarah Cockerton et al |
| **hCoV-19/Beijing/BJCDC-2073/2022** | EPI_ISL_16634167 | | Beijing Center for Disease Control and Prevention | Beijing Center for Disease Control and Prevention | Shujuan Cui et al |
| **hCoV-19/Jiangsu/JITHCC_05/2022** | EPI_ISL_16127648 | | Center Laboratory of Health Quarantine, Nanjing Customs District P.R. China | Center Laboratory of Health Quarantine, Nanjing Customs District P.R. China | Geng et al |
| **hCoV-19/Georgia/Tb-s6420/2022** | EPI_ISL_16200650 | | Central Republican Hospital | Department for Virology, Molecular Biology and Genome Research, R. G. Lugar Center for Public Health Research, National Center for Disease Control and Public Health (NCDC) of Georgia. | Giorgi Tomashvili et al |
| **hCoV-19/NewZealand/22CH6187/2022** | EPI_ISL_13764497 | | Southern Community Labs Dunedin | Institute of Environmental Science and Research (ESR), Kenepuru Science Centre | Sarah Cockerton et al |
| **hCoV-19/Shandong/WCDC-189/2022** | EPI_ISL_18070521 | | Weihai Center for Disease Control and Prevention | Weihai Center for Disease Control and Prevention | Xiang Li et al |
| **hCoV-19/Canada/BC-BCCDC-546309/2022** | EPI_ISL_16402002 | | British Columbia Centre For Disease Control | B.C. Centre for Disease Control Public Health Laboratory | Natalie Prystajecky et al |
| **hCoV-19/Bulgaria/22BG-EU_022736_PI211/2022** | EPI_ISL_17445994 | | National Center of Infectious and Parasitic Diseases | National Center of Infectious and Parasitic Diseases | Alexiev et al |
| **hCoV-19/Scotland/EDB72626/2022** | EPI_ISL_15612466 | | Virology Department, Royal Infirmary of Edinburgh, NHS Lothian / School of Biological Sciences, University of Edinburgh | COVID-19 Genomics UK (COG-UK) Consortium | McHugh M et al |
| **hCoV-19/Hungary/MBL-18229/2022** | EPI_ISL_17033708 | | Hungarian Defence Forces Medical Centre COVID Laboratory | Hungarian Defence Forces Medical Centre Epidemiological and Scientific Research Institute | Csaba István Pereszlényi et al |
| **hCoV-19/Curacao/CW-RIVM-118263/2022** | EPI_ISL_15913631 | | National Institute for Public Health and the Environment (RIVM) | National Institute for Public Health and the Environment (RIVM) | Adam Meijer et al |
| **hCoV-19/Montenegro/CO-01061_MNE000_2874942202/2022** | EPI_ISL_17244806 | | Laboratory for Molecular Diagnostics,IPHMN | Eurofins Genomics Europe Sequencing GmbH | Danijela Vujošević et al |
| **hCoV-19/Croatia/HZJZ_1962/2022** | EPI_ISL_16486415 | | Nastavni zavod za javno zdravstvo Primorsko- Goranske županije | Hrvatski zavod za javno zdravstvo | Irena Tabain et al |
| **hCoV-19/USA/ND-NDDH-21574/2022** | EPI_ISL_16320481 | | North Dakota Department of Health, Public Health Laboratory | North Dakota Department of Health, Public Health Laboratory | Lisa Wingerter et al |
| **hCoV-19/Jiangxi/JXNCDC-065/2023** | EPI_ISL_17495460 | | Jiangxi Province Center for Disease Control and Prevention | Jiangxi Province Center for Disease Control and Prevention | Wu et al |
| **hCoV-19/Hainan/SY639/2022** | EPI_ISL_17805472 | | Central & Clinical Laboratory of Sanya People's Hospital | Beijing Chao-Yang Hospital | Xiaoxia Wang et al |
| **hCoV-19/Shanxi/SXCDC-0128/2022** | EPI_ISL_16853832 | | Shanxi Provincial Center for Disease Control and Prevention | Shanxi Provincial Center for Disease Control and Prevention | Junyu Zhao et al |
| **hCoV-19/InnerMongolia/IVDC-632/2022** | EPI_ISL_16277971 | | The Center for Disease Control and Prevention of Inner Mongolia | National Institute for Viral Disease Control and Prevention, China CDC | Xiaoling Tian et al |
| **hCoV-19/InnerMongolia/IVDC-634/2022** | EPI_ISL_16277972 | | Baotou City Center for Disease Control and Prevention | National Institute for Viral Disease Control and Prevention, China CDC | Meng Gao et al |
| **hCoV-19/InnerMongolia/IVDC-292/2023** | EPI_ISL_16635309 | | The Center for Disease Control and Prevention of Inner Mongolia | China Center for Disease Control and Prevention, National Institute for Viral Disease Control and Prevention | Yanbo Bai et al |
| **hCoV-19/DominicanRepublic/2097866-LNSPDD/2022** | EPI_ISL_16022335 | | Laboratorio Nacional de Salud Pública Dr. Defilló | Laboratorio Nacional de Salud Pública Dr. Defilló | Isaac Miguel Sánchez et al |
| **hCoV-19/Liaoning/LNCDC-131-7/2023** | EPI_ISL_17775085 | | Institute for Communicable and Infectious Diseases Prevention and Control, Liaoning Center for Disease Control and Prevention | Institute for Communicable and Infectious Diseases Prevention and Control, Liaoning Center for Disease Control and Prevention | Mao et al |
| **hCoV-19/Beijing/BJCDC-2585/2022** | EPI_ISL_16642044 | | Beijing Center for Disease Control and Prevention | Beijing Center for Disease Control and Prevention | Fu Li et al |
| **hCoV-19/InnerMongolia/IVDC-641/2022** | EPI_ISL_16277978 | | The Center for Disease Control and Prevention of Inner Mongolia | National Institute for Viral Disease Control and Prevention, China CDC | Wuyihan et al |
| **hCoV-19/Liaoning/LNCDC-028-8/2022** | EPI_ISL_17183560 | | Liaoning Center for Disease Control and Prevention | Liaoning Center for Disease Control and Prevention | Lingling Mao et al |
| **hCoV-19/Guangxi/FCGCDC-030103/2023** | EPI_ISL_18330818 | | Institute of Acute Communicable Disease Prevention and Control, Guangxi Zhuang Autonomous Region Center for Disease Control and Prevention | Institute of Acute Communicable Disease Prevention and Control, Guangxi Zhuang Autonomous Region Center for Disease Control and Prevention | Huang et al |
| **hCoV-19/Shanghai/SCDC-XG03-1915/2023** | EPI_ISL_17488905 | | Shanghai Municipal Center for Disease Control & Prevention | Shanghai Municipal Center for Disease Control & Prevention | Zhang et al B |
| **hCoV-19/Guangxi/BHCDC-032214/2023** | EPI_ISL_18330875 | | Institute of Acute Communicable Disease Prevention and Control, Guangxi Zhuang Autonomous Region Center for Disease Control and Prevention | Institute of Acute Communicable Disease Prevention and Control, Guangxi Zhuang Autonomous Region Center for Disease Control and Prevention | Huang et al |
| **hCoV-19/Zhejiang/ZJCDC-564/2023** | EPI_ISL_17487938 | | Zhejiang Provincial Center for Disease Control and Prevention | Zhejiang Provincial Center for Disease Control and Prevention | Zhou et al |
| **hCoV-19/Hunan/HNCDC-2357/2023** | EPI_ISL_17487025 | | Hunan Provincial Center For Disease Control And Prevention | Hunan Provincial Center For Disease Control And Prevention | Xiang Xingyu et al |
| **hCoV-19/Beijing/BJCDC-0618/2023** | EPI_ISL_17620413 | | Institute for Infectious Disease and Endemic Disease Control, Beijing Center for Disease Control and Prevention | Institute for Infectious Disease and Endemic Disease Control, Beijing Center for Disease Control and Prevention | Pan et al |
| **hCoV-19/Anhui/WHCDC-117/2023** | EPI_ISL_17494011 | | Wuhu Center for Disease Control and Prevention | Wuhu Center for Disease Control and Prevention | Zha et al |
| **hCoV-19/Hubei/LHCDC-0133/2023** | EPI_ISL_17545760 | | Luohe Center for Disease Control and Prevention | Luohe Center for Disease Control and Prevention | Zhang et al B |
| **hCoV-19/Hebei/LF-23/2022** | EPI_ISL_17031544 | | Hebei Provincial Center for Disease Control and Prevention | Hebei Provincial Center for Disease Control and Prevention | Shiyou Liu et al |
| **hCoV-19/India/MH-ICMR-NIV-INSACOG-G-12920/2022** | EPI_ISL_16528823 | | ICMR-National Institute of Virology - INSACOG | NIV Influenza | Dr. Varsha Potdar et al |
| **hCoV-19/Yunnan/IVDC-604/2023** | EPI_ISL_16646070 | | Yunnan Center for Disease Control and Prevention | No.158 Dongsi Street, Xishan District, Kunming City, Yunnan Province, China | Zhang Meiling et al |
| **hCoV-19/China/CHFU-SHR-11/2022** | EPI_ISL_17959230 | | Children's Hospital of Fudan University | Children's Hospital of Fudan University | Ao et al |
| **hCoV-19/Beijing/BJCDC-0586/2023** | EPI_ISL_17162776 | | Beijing Center for Disease Control and Prevention | Beijing Center for Disease Control and Prevention | Fu Li et al |
| **hCoV-19/Tianjin/TJCDC-022311/2023** | EPI_ISL_17495267 | | Tiianjin centers for disease control and prevention | Tiianjin centers for disease control and prevention | Zou et al |
| **hCoV-19/China/CHFU-SHR-21/2023** | EPI_ISL_17959220 | | Children's Hospital of Fudan University | Children's Hospital of Fudan University | Ao et al |
| **hCoV-19/Taiwan/F77859/2023** | EPI_ISL_16723892 | | Centers for Disease Control, R.O.C. (Taiwan) | Centers for Disease Control, R.O.C. (Taiwan) | Yu-Chi Lin et al |
| **hCoV-19/Liaoning/LNCDC-086-1/2023** | EPI_ISL_17485729 | | Liaoning Center for Disease Control and Prevention | Liaoning Center for Disease Control and Prevention | Lingling Mao et al |
| **hCoV-19/Qinghai/QCDC-HSZ-99/2023** | EPI_ISL_17280138 | | Qinghai Red Cross Hospital | Qinghai Center for Disease Prevention and Control | Youju Lei et al |
| **hCoV-19/Jilin/JLCDC-th0041/2023** | EPI_ISL_17494268 | | Jilin Provincial Center for Disease Control and Prevention | Jilin Provincial Center for Disease Control and Prevention | Wu et al |
| **hCoV-19/Fujian/FPCDCP-ZZ31372/2022** | EPI_ISL_16283989 | | Fujian Provincial Center for Disease Control and Prevention | Fujian Provincial Center for Disease Control and Prevention | Lin Qi et al |
| **hCoV-19/Shaanxi/SXCDC-X008/2022** | EPI_ISL_16709470 | | Xi'an Center for Disease Control and Prevention | Shaanxi Provincial Center for Disease Control and Prevention | Yao Meng et al |
| **hCoV-19/Shanxi/SXCDC-0203/2023** | EPI_ISL_16911641 | | Shanxi Provincial Center for Disease Control and Prevention | Shanxi Provincial Center for Disease Control and Prevention | Junyu Zhao et al |
| **hCoV-19/Anhui/FYCDC-032/2023** | EPI_ISL_16679338 | | Fuyang Center for Disease Control and Prevention | Fuyang Center for Disease Control and Prevention | Zhangkuihu et al |
| **hCoV-19/HongKong/VK23009507/2023** | EPI_ISL_17187490 | | Pamela Youde Nethersole Eastern Hospital | Hong Kong Department of Health | Alan K.L. Tsang et al |
| **hCoV-19/Jiangsu/SZCDC-104/2022** | EPI_ISL_16698461 | | Jiangsu Provincial Center for Disease Control and Prevention | Jiangsu Provincial Center for Disease Control and Prevention | Li Zhifeng et al |
| **hCoV-19/Ningxia/IVDC-455/2023** | EPI_ISL_16635125 | | Ningxia Center for Disease Control and Prevention | China Center for Disease Control and Prevention, National Institute for Viral Disease Control and Prevention | Zhonglan Wu et al |
| **hCoV-19/Sichuan/SCDC-0075/2022** | EPI_ISL_17489250 | | Sichuan Center of Disease Control and Prevention | Sichuan Center of Disease Control and Prevention | Tao Zhang et al |
| **hCoV-19/Shanxi/TYCDC-002/2023** | EPI_ISL_17493233 | | Taiyuan Center for Disease Control and Prevention | Taiyuan Center for Disease Control and Prevention | Wang et al |
| **hCoV-19/Shannxi/SXCDC-0147/2023** | EPI_ISL_17492745 | | Shaanxi Provincial Center for Disease Control and Prevention | Shaanxi Provincial Center for Disease Control and Prevention | Meng Yao et al |
| **hCoV-19/Heilongjiang/HLJCDC-1593/2022** | EPI_ISL_16598357 | | Heilongjiang Provincial Center for Disease Control and Prevention, Department for Viral Disease Control and Prevention | Heilongjiang Provincial Center for Disease Control and Prevention, Department for Viral Disease Control and Prevention | Xu Jun et al |
| **hCoV-19/Shandong/WCDC-147/2023** | EPI_ISL_18070810 | | Weihai Center for Disease Control and Prevention | Weihai Center for Disease Control and Prevention | Xiang Li et al |
| **hCoV-19/Fujian/FCDC-QZBF006/2023** | EPI_ISL_17489157 | | Fujian Provincial Center for Disease Control and Prevention | Fujian Provincial Center for Disease Control and Prevention | Lin Qi et al |
| **hCoV-19/Hainan/CDC-045/2023** | EPI_ISL_16719075 | | Hainan Provincial Center for Disease Control and Prevention | Hainan Provincial Center for Disease Control and Prevention | Sun Chuyang et al |
| **hCoV-19/Zhejiang/ZJCDC-743/2023** | EPI_ISL_17501575 | | Zhejiang Provincial Center For Disease Control and Prevention | Zhejiang Provincial Center For Disease Control and Prevention | Zhou et al |
| **hCoV-19/Jiangsu/JCDC-095/2023** | EPI_ISL_17494016 | | Jiangsu Provincial Center for Disease Control and Prevention | Jiangsu Provincial Center for Disease Control and Prevention | Fan et al |
| **hCoV-19/InnerMongolia/NMCDC-CF-001/2023** | EPI_ISL_17789932 | | Immunization Branch, Inner Mongolia Center for Disease Control and Prevention | Immunization Branch, Inner Mongolia Center for Disease Control and Prevention | Tian et al |
| **hCoV-19/Tianjin/TJCDC-0407-2/2023** | EPI_ISL_17488802 | | Tianjin Centers for Disease Control and Prevention | Tianjin Centers for Disease Control and Prevention | Xiaoyan Li et al |
| **hCoV-19/Fujian/FCDC-QZ-S017/2023** | EPI_ISL_17547546 | | Institute for the Prevention and Control of Infectious Diseases, Fujian Provincial Center for Disease Control and Prevention | Institute for the Prevention and Control of Infectious Diseases, Fujian Provincial Center for Disease Control and Prevention | Lin Qi et al |
| **hCoV-19/Tianjin/TJCDC-033134/2023** | EPI_ISL_17494631 | | Tianjin Centers for Disease Control and Prevention | Tianjin Centers for Disease Control and Prevention | Li et al |
| **hCoV-19/Beijing/BJCDC-0910/2023** | EPI_ISL_17784939 | | Institute for Infectious Disease and Endemic Disease Control, Beijing Center for Disease Control and Preventio | Institute for Infectious Disease and Endemic Disease Control, Beijing Center for Disease Control and Preventio | Pan et al |
| **hCoV-19/Heilongjiang/HLJCDC-1172/2023** | EPI_ISL_17739618 | | Institute for Viral Disease Control and Prevention, Department for Viral Disease Control and Prevention, Heilongjiang Provincial Center for Disease Control and Prevention | Institute for Viral Disease Control and Prevention, Department for Viral Disease Control and Prevention, Heilongjiang Provincial Center for Disease Control and Prevention | Xu Jun et al |
| **hCoV-19/Shanxi/TYCDC-115/2023** | EPI_ISL_17951285 | | Microbiology Department, Taiyuan Center for Disease Control and Prevention | Microbiology Department, Taiyuan Center for Disease Control and Prevention | Wang et al |
| **hCoV-19/Shanxi/TYCDC-143/2023** | EPI_ISL_17951282 | | Microbiology Department, Taiyuan Center for Disease Control and Prevention | Microbiology Department, Taiyuan Center for Disease Control and Prevention | Wang et al |
| **hCoV-19/India/KA-15281_S38/2022** | EPI_ISL_15194105 | | DSUUdupi | inStem_NCBS_Strand Life Sciences (Supported by IN COVID support, FZE LLC) | Dasaradhi Palakodeti et al |
| **hCoV-19/India/TN-CDFD-J-144/2022** | EPI_ISL_13342189 | | Directorate of Public Health and Preventive Medicine | CDFD-INSACOG | Asmita Gupta et al |
| **hCoV-19/USA/IN-CDC-STM-57EN4QSJJ/2022** | EPI_ISL_14830953 | OP370692.1 | Helix | Centers for Disease Control and Prevention Division of Viral Diseases, Pathogen Discovery | Dakota Howard et al |
| **hCoV-19/Bolivia/CBBA-PUC_MVL_3354/2022** | EPI_ISL_16765654 | | Laboratorio de Biología Molecular, IIBISMED Facultad de Medicina, Universidad Mayor de San Simón. | Laboratory of Molecular Virology, School of Medicine, Pontificia Universidad Catolica de Chile | Catalina Pardo-Roa et al |
| **hCoV-19/Bahamas/23622-GMI/2022** | EPI_ISL_17039707 | | Reference Laboratory of the Ministry of Health | Genomics and Proteomics Departament, Gorgas Memorial Institute For Health Studies | Indira Martins et al |
| **hCoV-19/SaintLucia/168227/2022** | EPI_ISL_16166902 | | Ezra Long Laboratory Victoria Hospital | CARPHA | Carlos Rampersad on behalf of CARPHA et al |

| | | | | | |
|---|---|---|---|---|---|
| **hCoV-19/Uganda/MRC-UVRI-C2301040/2022** | EPI_ISL_16613341 | | MRC/UVRI & LSHTM Uganda Research Unit, Uganda Virus Research Institute | MRC/UVRI & LSHTM Uganda Research Unit, Uganda Virus Research Institute | Deogratius Ssemwanga et al |
| **hCoV-19/Ethiopia/BETIN-21356071780/2022** | EPI_ISL_18904534 | | Health Biotechnology, Bio and Emerging Technology Institute | Health Biotechnology, Bio and Emerging Technology Institute | Bitew et al |
| **hCoV-19/India/TG-CDFD-GH-554/2022** | EPI_ISL_14445812 | | Gandhi Medical College | CDFD-INSACOG | Asmita Gupta et al |
| **hCoV-19/Egypt/CPHL-23526/2023** | EPI_ISL_18115713 | | Central Public Health Laboratories, Ministry of Health and Population | Central Public Health Laboratories, Ministry of Health and Population | Wael H. Roshdy et al |
| **hCoV-19/Beijing/BJCDC-3220/2022** | EPI_ISL_16350939 | | Beijing Center for Disease Control and Prevention | Beijing Center for Disease Control and Prevention | Yang Pan et al |
| **hCoV-19/Tunisia/574/2022** | EPI_ISL_15298418 | | 1-Laboratory of Microbiology and Biochemistry, Aziza Othmena Hospital; 2-University of Tunis ElManar, Faculty of Medicine of Tunis, LR16SP01, Tunis, Tunisia | Clinical and Experimental Pharmacology Lab, LR16SP02, National Center of Pharmacovigilance, University of Tunis El Manar, Tunis, Tunisia. 2- Ministry of Health, National Observatory of New and Emerging Diseases, 1006, Tunis, Tunisia | Sana Ferjani et al |
| **hCoV-19/Italy/CAM-TIGEM-IZSM-COLLI-44681/2022** | EPI_ISL_16984739 | | Ospedale Santa Maria delle Grazie | TIGEM | Antonio Grimaldi Patrizia Annunziata Francesco Panariello Claudia TiberioTonya Fusco Michela Daniele Valentina Bouche Ilaria Cimmino Chiara Colantuono Lucio Di Filippo Anna Manfredi Marcello Salvi Antonio Limone Luigi Atripaldi Andrea Ballabio Davide Cacchiarelli et al |
| **hCoV-19/Guangxi/GCDC-BS-2210/2022** | EPI_ISL_17104932 | | Baise Center for Disease Control and Prevention | Guangxi Zhuang Autonomous Region Center for Disease Control and Prevention | Zan Chen et al |
| **hCoV-19/Laos/LOMWRU-0589/2022** | EPI_ISL_15973677 | | LOMWRU | LOMWRU/Microbiology Laboratory, Mahosot Hospital | Manila Souksavanh et al |
| **hCoV-19/SaudiArabia/KAUST161/2022** | EPI_ISL_15826950 | | King Faisal Specialist Hospital & Research Center | Pathogen Genomics Lab King Abdullah University of Science and Technology (KAUST) | Sabir A Adoub et al |
| **hCoV-19/Kazakhstan/NLA-29_27/2022** | EPI_ISL_17454137 | | Republican Diagnostic Center, University Medical Center, Nazarbayev University | Center for Life Sciences, National Laboratory Astana, Nazarbayev University | Ulykbek Kairov et al |
| **hCoV-19/Russia/KYA-RII-MH122319S/2022** | EPI_ISL_16336513 | | Krasnoyarsk Interdistrict Clinical Emergency Hospital named after N. S. Karpovich | WHO National Influenza Centre Russian Federation | Andrey Komissarov et al |
| **hCoV-19/Ukraine/71896/2022** | EPI_ISL_14923324 | | Reference laboratory for diagnostics of HIV/AIDS, virological and especially dangerous pathogens | Reference laboratory for diagnostics of HIV/AIDS, virological and especially dangerous pathogens | Iryna Demchyshyna et al |
| **hCoV-19/USA/NC-UNC-7721/2022** | EPI_ISL_14796487 | | Clinical Molecular Microbiology Laboratory, UNC Hospitals | Jeremy Wang | Jeremy Wang et al |
| **hCoV-19/Thailand/NIC_CRI_FRS2438/2022** | EPI_ISL_15126587 | | MaechanHospital | National Institute of Health, Department of Medical Sciences, Ministry of Public Health, Thailand | Pilailuk Okada et al |
| **hCoV-19/Bahrain/22020471554/2022** | EPI_ISL_15251476 | OP295557.1 | Communicable Disease Laboratory, Public Health Directorate | Communicable Disease Laboratory, Public Health Directorate | Alwasti et al |
| **hCoV-19/Lebanon/SG-1/2022** | EPI_ISL_16583841 | | Saint George Hospital University Medical Center | Melhem/Matar lab | Nada M. Melhem et al |
| **hCoV-19/Shandong/WCDC-148/2022** | EPI_ISL_18070480 | | Weihai Center for Disease Control and Prevention | Weihai Center for Disease Control and Prevention | Xiang Li et al |
| **hCoV-19/Qatar/QA.QU_18.51.G6/2022** | EPI_ISL_16076555 | | Ministry of Public Health / Hamad Medical Corporation | Biomedical Research Center (BRC), Qatar University / Qatar Genome Project (QGP) | BRC: Heba A. Al-Khatib et al |
| **hCoV-19/Shandong/WCDC-158/2022** | EPI_ISL_18070490 | | Weihai Center for Disease Control and Prevention | Weihai Center for Disease Control and Prevention | Xiang Li et al |
| **hCoV-19/Japan/TKYkbm50149/2022** | EPI_ISL_15188664 | | Tokyo Metropolitan Institute of Public Health | Tokyo Metropolitan Institute of Public Health | Akane Negishi et al |
| **hCoV-19/Laos/LOMWRU-0512/2022** | EPI_ISL_15655972 | | LOMWRU | LOMWRU/Microbiology Laboratory, Mahosot Hospital | Manila Souksavanh et al |
| **hCoV-19/Kazakhstan/AST-921/2022** | EPI_ISL_15757840 | | National Center of Expertise, Branch in Astana city | National center of Expertise, National center for Biotechnology | Abdaliyev Askar et al |
| **hCoV-19/SouthKorea/KDCA146539/2022** | EPI_ISL_15535052 | | Division of Emerging Infectious Diseases, Bureau of Infectious Diseases Diagnosis Control, Korea Disease Control and Prevention Agency | Division of Emerging Infectious Diseases, Bureau of Infectious Diseases Diagnosis Control, Korea Disease Control and Prevention Agency | Jeong-Ah Kim et al |
| **hCoV-19/Laos/LOMWRU-0354/2022** | EPI_ISL_13747284 | OP159674.1 | LOMWRU | LOMWRU/Microbiology Laboratory, Mahosot Hospital | Manila Souksavanh et al |
| **hCoV-19/Morocco/IPM20428897/2022** | EPI_ISL_15251958 | | Institut Pasteur du Maroc, Medical Virology and BSL3 Laboratory | Institut Pasteur du Maroc, Centre de Séquençage Génomique | EL Hamouchi Adil et al |
| **hCoV-19/Russia/SAM-RII-MH73147S/2022** | EPI_ISL_14901472 | | Tolyatti Municipal Clinical Hospital No 5 | WHO National Influenza Centre Russian Federation | Andrey Komissarov et al |
| **hCoV-19/Ukraine/71938/2022** | EPI_ISL_15348940 | | Reference laboratory for diagnostics of HIV/AIDS, virological and especially dangerous pathogens | Reference laboratory for diagnostics of HIV/AIDS, virological and especially dangerous pathogens | Iryna Demchyshyna et al |
| **hCoV-19/Bahrain/CDL-750904577/2022** | EPI_ISL_17081909 | | Communicable Disease Laboratory | Communicable Disease Laboratory | Marhoon et al |
| **hCoV-19/Lithuania/MB46553/2022** | EPI_ISL_17977417 | | Lietuvos sveikatos mokslų universiteto Kauno ligoninė | National Public Health Surveillance Laboratory | Lukas Zemaitis et al |
| **hCoV-19/SaudiArabia/KAUST116/2022** | EPI_ISL_15826907 | | King Faisal Specialist Hospital & Research Center | Pathogen Genomics Lab King Abdullah University of Science and Technology (KAUST) | Sabir A Adoub et al |
| **hCoV-19/Jilin/JLCDC-59072/2022** | EPI_ISL_16713322 | | Jilin Center for Disease Control and Prevention | Jilin Center for Disease Control and Prevention | Wu Donglin et al |
| **hCoV-19/SouthKorea/KDCA180197/2022** | EPI_ISL_16339594 | | Division of Emerging Infectious Diseases, Bureau of Infectious Diseases Diagnosis Control, Korea Disease Control and Prevention Agency | Division of Emerging Infectious Diseases, Bureau of Infectious Diseases Diagnosis Control, Korea Disease Control and Prevention Agency | Jeong-Ah Kim et al |
| **hCoV-19/Serbia/K948598-11/2022** | EPI_ISL_16317156 | | Huo-yan national laboratory for molecular detection of infectious agents Nis | Institute of Molecular Genetics and Genetic Engineering University of Belgrade | Valentina Djordjevic et al |
| **hCoV-19/Serbia/L266648-01/2023** | EPI_ISL_17250968 | | Huo-yan national laboratory for molecular detection of infectious agents Nis | Institute of Molecular Genetics and Genetic Engineering University of Belgrade | Valentina Djordjevic et al |
| **hCoV-19/Serbia/K500006-10/2022** | EPI_ISL_16317050 | | Huo-yan national laboratory for molecular detection of infectious agents Nis | Institute of Molecular Genetics and Genetic Engineering University of Belgrade | Valentina Djordjevic et al |
| **hCoV-19/Thailand/TUC-L22-53806/2022** | EPI_ISL_15363605 | | Mae Sot General Hospital | Thailand MOPH - U.S. CDC Collaboration (TUC) | Thidathip Wongsurawat et al |
| **hCoV-19/Myanmar/DSMRC-111-7/2022** | EPI_ISL_14241971 | OP159818.1 | Defence Services Medical Research Center | DSMRC | Khine Zaw OO et al |
| **hCoV-19/Mexico/SIN_IBT_IMSS_9850/2022** | EPI_ISL_15671279 | | Centro de Investigacion Biomedica de Occidente (CIBO) | Instituto de Biotecnologia de la UNAM | Consorcio Mexicano de Vigilancia Genomica (CoViGen-Mex). Julio Elias Alvarado-Yaah et al |
| **hCoV-19/SouthAfrica/NICD-N49529/2022** | EPI_ISL_15259373 | | LANCET | National Institute for Communicable Diseases of the National Health Laboratory Service | Everatt J et al |
| **hCoV-19/Uruguay/DLSP_3556/2022** | EPI_ISL_16389521 | | Departamento Laboratorios de Salud Pública (DLSP) Ministerio de Salud Pública | Departamento Laboratorios de Salud Pública (DLSP) Ministerio de Salud Pública | Cortinas et al |
| **hCoV-19/Tianjin/TJCDC-1050/2022** | EPI_ISL_16922475 | | Tianjin Centers for Disease Control and Prevention | Tianjin Centers for Disease Control and Prevention | Li Xiaoyan et al |
| **hCoV-19/Pakistan/IDRL-AKU-KHI-79135-S15/2022** | EPI_ISL_15962614 | | "Clinical Lab, Aga Khan University Hospital" | "Infectious Diseases Research Laboratory, Department of Pediatrics and Child Health" | "Imran Nisar et al |
| **hCoV-19/Pakistan/NIH-B112-S2/2022** | EPI_ISL_15850306 | | Department of Virology, National Institute of Health, Islamabad, Pakistan | Department of Virology, National Institute of Health, Islamabad, Pakistan | Massab Umair et al |
| **hCoV-19/Romania/NIPH-216483/2022** | EPI_ISL_16550589 | | National Institute of Public Health Laboratory | National Institute of Public Health Laboratory | Mihaela Leustean et al |
| **hCoV-19/Iraq/BG-USAFSAM-S20126/2022** | EPI_ISL_15889752 | | US Air Force School of Aerospace Medicine | US Air Force School of Aerospace Medicine | Anthony Fries et al |
| **hCoV-19/Norway/32307/2022** | EPI_ISL_15742464 | | University Hospital of Northern Norway, Department for Microbiology and Infectious Disease Control | Norwegian Institute of Public Health, Department of Virology | Kathrine Stene-Johansen et al |
| **hCoV-19/CoteDIvoire/IPCI-DVE-COV826635/2022** | EPI_ISL_15908258 | | Departement of Epidemic Viruses, Institut Pasteur of Cote d'Ivoire Abidjan | Departement of Epidemic Viruses, Institut Pasteur of Cote d'Ivoire Abidjan | Hervé Albéric Adje Kadjo et al |
| **hCoV-19/Chongqing/YBCDC-0115/2022** | EPI_ISL_16613442 | | Chongqing Yubei District Center for Disease Control and Prevention | Chongqing Yubei District Center for Disease Control and Prevention | Jin Yan et al |
| **hCoV-19/Beijing/BJCDC-1738/2022** | EPI_ISL_16614675 | | Beijing Center for Disease Control and Prevention | Beijing Center for Disease Control and Prevention | Zhaomin Feng et al |
| **hCoV-19/Croatia/HZJZ_2854/2023** | EPI_ISL_17247606 | | Nastavni zavod za javno zdravstvo Primorsko- Goranske županije | Hrvatski zavod za javno zdravstvo | Irena Tabain et al |
| **hCoV-19/Georgia/Tb-s7407/2022** | EPI_ISL_17182891 | | Adjara Public Health Center | Department for Virology, Molecular Biology and Genome Research, R. G. Lugar Center for Public Health Research, National Center for Disease Control and Public Health (NCDC) of Georgia. | Giorgi Tomashvili et al |
| **hCoV-19/SaudiArabia/KAUST92/2022** | EPI_ISL_15826883 | | King Faisal Specialist Hospital & Research Center | Pathogen Genomics Lab King Abdullah University of Science and Technology (KAUST) | Sabir A Adoub et al |
| **hCoV-19/Sweden/C-03_SE600_000200449722N2/2022** | EPI_ISL_17713029 | | Akademiska Sjukhuset, Clinical Microbiology and Hospital Hygiene | Akademiska Sjukhuset, Clinical Microbiology and Hospital Hygiene | Jonathan Haars et al |
| **hCoV-19/Indonesia/KI-GS-NIHRD-GSILab-1107941/2022** | EPI_ISL_14605152 | | Lab Klinik Khatulistiwa | Genomik Solidaritas Indonesia Laboratorium | Dian Pitaloka Rajagukguk et al |
| **hCoV-19/Morocco/IPM20429271/2022** | EPI_ISL_15251965 | | Institut Pasteur du Maroc, Medical Virology and BSL3 Laboratory | Institut Pasteur du Maroc, Centre de Séquençage Génomique | EL Hamouchi Adil et al |
| **hCoV-19/Zhejiang/ZPCDCP-121/2022** | EPI_ISL_17261623 | | Zhejiang Provincial Center for Disease Control and Prevention | Zhejiang Provincial Center for Disease Control and Prevention | Yi Sun et al |
| **hCoV-19/USA/FL-CDC-STM-S77PTDD8Z/2022** | EPI_ISL_15260261 | OP590968.1 | Helix | Centers for Disease Control and Prevention Division of Viral Diseases, Pathogen Discovery | Dakota Howard et al |
| **hCoV-19/Zhejiang/ZPCDCP-126/2022** | EPI_ISL_17261628 | | Zhejiang Provincial Center for Disease Control and Prevention | Zhejiang Provincial Center for Disease Control and Prevention | Yi Sun et al |
| **hCoV-19/Montenegro/CO-00892_MNE000_2173162202/2022** | EPI_ISL_14509126 | | Laboratory for Molecular Diagnostics,IPHMN | Eurofins Genomics Europe Sequencing GmbH | Danijela Vujošević et al |
| **hCoV-19/Egypt/CPHL-EGY22927/2022** | EPI_ISL_17423966 | | CPHL,MOH,EGYPT | Central Public Health Laboratories (CPHL) | Wael H. Roshdy et al |
| **hCoV-19/Tunisia/540/2022** | EPI_ISL_15298411 | | 1-Laboratory of Microbiology and Biochemistry, Aziza Othmena Hospital; 2-University of Tunis ElManar, Faculty of Medicine of Tunis, LR16SP01, Tunis, Tunisia | Clinical and Experimental Pharmacology Lab, LR16SP02, National Center of Pharmacovigilance, University of Tunis El Manar, Tunis, Tunisia. 2- Ministry of Health, National Observatory of New and Emerging Diseases, 1006, Tunis, Tunisia | Sana Ferjani et al |
| **hCoV-19/Egypt/CPHL-EGY23784/2022** | EPI_ISL_18139076 | | CPHL,MOH,EGYPT | CPHL,MOH,EGYPT | Wael H. Roshdy et al |
| **hCoV-19/Uzbekistan/47_1808-CAT/2022** | EPI_ISL_15941918 | | SI"Zangiota No.1 and 2 special hospital for the treatment of patients with coronovirus infection" | Center for Advanced Technologies | Dalimova D.A. et al |
| **hCoV-19/Uzbekistan/43_1819-CAT/2022** | EPI_ISL_15941914 | | SI"Zangiota No.1 and 2 special hospital for the treatment of patients with coronovirus infection" | Center for Advanced Technologies | Dalimova D.A. et al |
| **hCoV-19/Georgia/Tb-s5942/2022** | EPI_ISL_14997081 | | Central Republican Hospital | Department for Virology, Molecular Biology and Genome Research, R. G. Lugar Center for Public Health Research, National Center for Disease Control and Public Health (NCDC) of Georgia. | Giorgi Tomashvili et al |
| **hCoV-19/Georgia/Tb-s5903/2022** | EPI_ISL_14791459 | | Kutaisi Public Health and Safe Environment Center | Department for Virology, Molecular Biology and Genome Research, R. G. Lugar Center for Public Health Research, National Center for Disease Control and Public Health (NCDC) of Georgia. | Giorgi Tomashvili et al |
| **hCoV-19/Israel/SMC-7101324/2022** | EPI_ISL_14687071 | | Shamir Medical Center (Asaf Harofe) | Shamir Medical Center (Asaf Harofe) | Nir Rainy et al |
| **hCoV-19/India/MP-AIIMS_B-ICMR-INSACOG-WGS-1567/2022** | EPI_ISL_15063809 | | AIIMS Bhopal VRDL/Translational Medicine - INSACOG | AIIMS Bhopal VRDL/Translational Medicine - INSACOG | Anvita G. Malhotra et al |
| **hCoV-19/Chongqing/YBCDC-0215/2022** | EPI_ISL_17197587 | | Chongqing Yubei District Center for Disease Control and Prevention | Chongqing Yubei District Center for Disease Control and Prevention | Jin Yan et al |
| **hCoV-19/Cambodia/2214828/2022** | EPI_ISL_14561942 | | National Institute of Public Health | National Institute of Public Health | Chhorvann CHHEA et al |
| **hCoV-19/Guangxi/GCDC-BS-2219/2022** | EPI_ISL_17104935 | | Baise Center for Disease Control and Prevention | Guangxi Zhuang Autonomous Region Center for Disease Control and Prevention | Zan Chen et al |
| **hCoV-19/Israel/SMC-7105570/2022** | EPI_ISL_15405138 | | Shamir Medical Center (Asaf Harofe) | Shamir Medical Center (Asaf Harofe) | Nir Rainy et al |
| **hCoV-19/Hungary/120815-UPL23-000245/2022** | EPI_ISL_16724425 | | Institute of Laboratory Medicine, Semmelweis University and Epidemiology and Surveillance Centre, Semmelweis University | iBioscience Kft. | Gergő Túri et al |
| **hCoV-19/CzechRepublic/NRL-s1095/2023** | EPI_ISL_17544588 | | Masaryk Hospital | National Institute of Public Health | Helena Jirincova et al |
| **hCoV-19/Mongolia/UB-97529/2022** | EPI_ISL_15607943 | | National Center for Communicable Diseases (NCCD) National Influenza Center | National Center for Communicable Diseases (NCCD) National Influenza Center | Bayasgalan Namuutsetseg et al |
| **hCoV-19/Pakistan/IDRL-AKU-KHI-79284-S6/2022** | EPI_ISL_16362571 | | "Clinical Lab, Aga Khan University Hospital" | "Infectious Diseases Research Laboratory, Department of Pediatrics and Child Health" | "Waqasuddin Khan et al |
| **hCoV-19/Austria/LB-R001009-S181/2022** | EPI_ISL_15671029 | | Lifebrain Covid Labor GmbH | Lifebrain Covid Labor GmbH | Filip Sima et al |
| **hCoV-19/USA/ND-NDDH-21455/2022** | EPI_ISL_16132133 | | North Dakota Department of Health, Public Health Laboratory | North Dakota Department of Health, Public Health Laboratory | Lisa Wingerter et al |
| **hCoV-19/Zhejiang/ZJCDC-380/2023** | EPI_ISL_17488115 | | Zhejiang Provincial Center for Disease Control and Prevention | Zhejiang Provincial Center for Disease Control and Prevention | Zhou et al |
| **hCoV-19/Portugal/INSA-PT45655/2023** | EPI_ISL_16825059 | | Hospital Fernando Fonseca | Instituto Nacional de Saude Doutor Ricardo Jorge | Borges et al |
| **hCoV-19/Lebanon/EH-1/2022** | EPI_ISL_16583794 | | Elias Hrawi Governmental Hospital | Melhem/Matar lab | Nada M. Melhem et al |
| **hCoV-19/Croatia/40157/2022** | EPI_ISL_14571278 | | Zavod za javno zdravstvo Međimurske županije | Hrvatski zavod za javno zdravstvo | Irena Tabain et al |
| **hCoV-19/Singapore/9880/2022** | EPI_ISL_14440612 | | National Public Health Laboratory, National Centre for Infectious Diseases | National Public Health Laboratory, National Centre for Infectious Diseases | Grace Ngan et al |
| **hCoV-19/Jordan/s25/2022** | EPI_ISL_14770454 | | MOH-Central Lab | MOH-Central Lab | Dr.Mahmood Al Gazo et al |
| **hCoV-19/Tianjin/TJCDC-1111/2022** | EPI_ISL_16922513 | | Tianjin Centers for Disease Control and Prevention | Tianjin Centers for Disease Control and Prevention | Li Xiaoyan et al |
| **hCoV-19/CostaRica/INC-6424-765561/2022** | EPI_ISL_14593582 | | AREA DE SALUD CAÑAS | Inciensa, Instituto de Investigación y Enseñanza en Nutrición y Salud | Francisco Duarte et al |
| **hCoV-19/Syria/Northwest-00351/2022** | EPI_ISL_14987038 | | EWARN's Lab | Ministry of Health Turkey | Yasir Elferruh et al |
| **hCoV-19/Vietnam/PIHCM_2810/2022** | EPI_ISL_16526960 | | Pasteur Institute in Ho Chi Minh city | Pasteur Institute in Ho Chi Minh city | Hang Pham Thi Thu et al |
| **hCoV-19/Indonesia/LA-GS-NIHRD-GSILab-1122566/2022** | EPI_ISL_15232179 | | Dinkes Provinsi Lampung (Labkes Provinsi) | Genomik Solidaritas Indonesia Laboratorium | Aisyi Sakina Rifani et al |
| **hCoV-19/Estonia/24000031872/2022** | EPI_ISL_15652871 | | Synlab Eesti OÜ | 1. Laboratory of Communicable Diseases (Estonia); 2. Eurofins Genomics Europe Sequencing GmbH | Abroi A. et al |
| **hCoV-19/Georgia/Tb-s7397/2022** | EPI_ISL_17182882 | | Adjara Public Health Center | Department for Virology, Molecular Biology and Genome Research, R. G. Lugar Center for Public Health Research, National Center for Disease Control and Public Health (NCDC) of Georgia. | Giorgi Tomashvili et al |
| **hCoV-19/Qatar/QU.18.64.A12/2022** | EPI_ISL_15922694 | | Ministry of Public Health / Hamad Medical Corporation | Biomedical Research Center (BRC), Qatar University / Qatar Genome Project (QGP) | BRC: Heba A. Al-Khatib et al |
| **hCoV-19/USA/MD-CDC-LC0894692/2022** | EPI_ISL_15399597 | | Laboratory Corporation of America | Centers for Disease Control and Prevention Division of Viral Diseases, Pathogen Discovery | Dakota Howard et al |
| **hCoV-19/Singapore/12550/2022** | EPI_ISL_15209215 | | National Public Health Laboratory, National Centre for Infectious Diseases | National Public Health Laboratory, National Centre for Infectious Diseases | Grace Ngan et al |
| **hCoV-19/Georgia/Tb-s6419/2022** | EPI_ISL_16200649 | | Central Republican Hospital | Department for Virology, Molecular Biology and Genome Research, R. G. Lugar Center for Public Health Research, National Center for Disease Control and Public Health (NCDC) of Georgia. | Giorgi Tomashvili et al |
| **hCoV-19/Shandong/SDCDC-0759/2023** | EPI_ISL_16758111 | | Jinan Center for Disease Control and Prevention | Shandong Provincial Center for Disease Control and Prevention | Yan Li et al |
| **hCoV-19/Shandong/SDCDC-1837/2022** | EPI_ISL_16604393 | | Jinan Municipal Center for Disease Control and Prevention | Shandong Provincial Center for Disease Control and Prevention | Yan Li et al |
| **hCoV-19/Taiwan/NCKUH-061022012873/2022** | EPI_ISL_16298556 | | Virology Lab,Department of Pathology, National Cheng Kung University Hospital | Virology Lab,Department of Pathology, National Cheng Kung University Hospital | Huey-Pin Tsai et al |
| **hCoV-19/SouthKorea/KDCA159926/2022** | EPI_ISL_15906732 | | Division of Emerging Infectious Diseases, Bureau of Infectious Diseases Diagnosis Control, Korea Disease Control and Prevention Agency | Division of Emerging Infectious Diseases, Bureau of Infectious Diseases Diagnosis Control, Korea Disease Control and Prevention Agency | Jeong-Ah Kim et al |
| **hCoV-19/Romania/B-28323/2022** | EPI_ISL_17540854 | | National Institute of Infectious Diseases-Prof. Dr. Matei Bals Molecular Diagnostics Laboratory | National Institute of Infectious Diseases-Prof. Dr. Matei Bals Molecular Diagnostics Laboratory | Corina Casangiu et al |
| **hCoV-19/Egypt/CPHL-23600/2023** | EPI_ISL_18115518 | | Central Public Health Laboratories, Ministry of Health and Population | Central Public Health Laboratories, Ministry of Health and Population | Wael H. Roshdy et al |
| **hCoV-19/Shandong/SDCDC-3025/2022** | EPI_ISL_16604462 | | Jinan Municipal Center for Disease Control and Prevention | Shandong Provincial Center for Disease Control and Prevention | Yan Li et al |
| **hCoV-19/Sichuan/NCCDC-6/2023** | EPI_ISL_17563621 | | Nanchong Center of Disease Control and Prevention | Microbiology laboratory, Sichuan Center of Disease Control and Prevention | Sheng et al |
| **hCoV-19/Jiangsu/CDC-JS75/2022** | EPI_ISL_17791060 | | Jiangsu Provincial Center for Disease Control & Prevention | Jiangsu Provincial Center for Disease Control & Prevention | Kangchen Zhao et al |
| **hCoV-19/Beijing/BJCDC-2272/2022** | EPI_ISL_16634307 | | Beijing Center for Disease Control and Prevention | Beijing Center for Disease Control and Prevention | Shujuan Cui et al |
| **hCoV-19/Henan/CDC-004/2022** | EPI_ISL_16610102 | | Kaifeng No.2 People's Hospital | Kaifeng Disease Prevention and Control Center | Huaijie Hu et al |
| **hCoV-19/Qinghai/QHCDC-1/2022** | EPI_ISL_16713429 | | Qinghai Center for Disease Prevention and Control | Qinghai Center for Disease Prevention and Control | Youju Lei et al |
| **hCoV-19/Qinghai/QHCDC-HB2Y08/2023** | EPI_ISL_16713424 | | Haibei Second People's Hospital | Qinghai Center for Disease Control and Prevention | Youju Lei et al |
| **hCoV-19/Tianjin/TJCDC-1541/2022** | EPI_ISL_16922976 | | Tianjin Centers for Disease Control and Prevention | Tianjin Centers for Disease Control and Prevention | Li Xiaoyan et al |
| **hCoV-19/Guizhou/GZGYCDC-112/2023** | EPI_ISL_17501625 | | Experimental Center, Guizhou Provincial Center for Disease Control and Prevention | Experimental Center, Guizhou Provincial Center for Disease Control and Prevention | Wan et al |
| **hCoV-19/Tianjin/TJCDC-001/2022** | EPI_ISL_16481836 | | Tianjin Centers for Disease Control and Prevention | Tianjin Centers for Disease Control and Prevention | Xiaoyan Li et al |
| **hCoV-19/Qinghai/QCDC-QL-10/2022** | EPI_ISL_16723097 | | Qilian Center for Disease Prevention and Control | Qinghai Center for Disease Prevention and Control | Youju Lei et al |
| **hCoV-19/Hunan/CCDC-51/2023** | EPI_ISL_16598721 | | Changsha Center for Disease Control and Prevention | Changsha Center for Disease Control and Prevention | Zheng Huang et al |
| **hCoV-19/Guangxi/GCDC-NN-123002/2022** | EPI_ISL_17088669 | | Nanning Center for Disease Control and Prevention | Guangxi Zhuang Autonomous Region Center for Disease Control and Prevention | Jianqiu Qin et al |

| | | | | | |
|---|---|---|---|---|---|
| **hCoV-19/Kazakhstan/NLA-30_33/2022** | EPI_ISL_17454338 | | Clinical Diagnostic Laboratory "Olymp" | Center for Life Sciences, National Laboratory Astana, Nazarbayev University | Ulykbek Kairov et al |
| **hCoV-19/Mongolia/UB-93433/2022** | EPI_ISL_15295384 | | National Center for Communicable Diseases (NCCD) National Influenza Center | National Center for Communicable Diseases (NCCD) National Influenza Center | Bayasgalan Namuutsetseg et al |
| **hCoV-19/Estonia/RC0714255/2022** | EPI_ISL_15019299 | | Synlab Eesti OÜ | 1. Laboratory of Communicable Diseases (Estonia); 2. Eurofins Genomics Europe Sequencing GmbH | Abroi A. et al |
| **hCoV-19/Kazakhstan/NLA-30_31/2022** | EPI_ISL_17454307 | | Republican Diagnostic Center, University Medical Center, Nazarbayev University | Center for Life Sciences, National Laboratory Astana, Nazarbayev University | Ulykbek Kairov et al |
| **hCoV-19/Xinjiang/XJCDCHM-007/2023** | EPI_ISL_17501696 | | Institute for the Prevention and Control of Infectious Diseases, Xinjiang Center for Disease Control and Prevention | Institute for the Prevention and Control of Infectious Diseases, Xinjiang Center for Disease Control and Prevention | Ma et al |
| **hCoV-19/Fujian/FCDC-XM010668/2023** | EPI_ISL_17488502 | | Fujian Provincial Center for Disease Control and Prevention | Fujian Provincial Center for Disease Control and Prevention | Lin et al |
| **hCoV-19/Sichuan/GZCDC-2303028/2023** | EPI_ISL_17309147 | | Ganzi Center of Disease Control and Prevention | Sichuan Center of Disease Control and Prevention | Hailiang Qi et al |
| **hCoV-19/Yunnan/IVDC-1391/2022** | EPI_ISL_16641641 | | Yunnan Center for Disease Control and Prevention | China Center for Disease Control and Prevention, National Institute for Viral Disease Control and Prevention | ZhangMeiling et al |
| **hCoV-19/Yunnan/YNCDC-0164/2023** | EPI_ISL_17493264 | | Yunnan Center for Disease Control and Prevention | Yunnan Center for Disease Control and Prevention | Zhang et al B |
| **hCoV-19/Qinghai/QCDC-ETYY-67/2023** | EPI_ISL_17584966 | | Microbiological testing center, Qinghai Center for Disease Prevention and Control | Microbiological testing center, Qinghai Center for Disease Prevention and Control | Li et al |
| **hCoV-19/Gansu/GSCDC-253/2023** | EPI_ISL_17150499 | | Gansu Provincial Centre for Disease Control and Prevention | Gansu Provincial Centre for Disease Control and Prevention | Baodi Li et al |
| **hCoV-19/Ningxia/IVDC-1307/2022** | EPI_ISL_16635071 | | Ningxia Center for Disease Control and Prevention | China Center for Disease Control and Prevention, National Institute for Viral Disease Control and Prevention | Zhonglan Wu et al |
| **hCoV-19/Fujian/FCDC-FJPNT01888/2022** | EPI_ISL_17489453 | | Fujian Center for Disease Control and Prevention | Fujian Center for Disease Control and Prevention | Lin et al |
| **hCoV-19/Gansu/GSCDC-029/2022** | EPI_ISL_16702248 | | Gansu Provincial Center for Disease Control and Prevention | Gansu Provincial Centre for Disease Control and Prevention | Baodi Li et al |
| **hCoV-19/Gansu/GSCDC-288/2023** | EPI_ISL_17150534 | | Gansu Provincial Centre for Disease Control and Prevention | Gansu Provincial Centre for Disease Control and Prevention | Baodi Li et al |
| **hCoV-19/Gansu/GSCDC-230/2022** | EPI_ISL_17150476 | | Gansu Provincial Centre for Disease Control and Prevention | Gansu Provincial Centre for Disease Control and Prevention | Baodi Li et al |
| **hCoV-19/Gansu/GSCDC-0102/2023** | EPI_ISL_17494997 | | Gansu Provincial Centre for Disease Control and Prevention | Gansu Provincial Centre for Disease Control and Prevention | Li et al |
| **hCoV-19/Chongqing/CQCDC-1381/2023** | EPI_ISL_17495305 | | Chongqing Center for Disease Control and Prevetion | Chongqing Center for Disease Control and Prevetion | Ye et al |
| **hCoV-19/Chongqing/CQCDC-1379/2023** | EPI_ISL_17495307 | | Chongqing Center for Disease Control and Prevetion | Chongqing Center for Disease Control and Prevetion | Ye et al |
| **hCoV-19/Jiangxi/JXCDC-659/2022** | EPI_ISL_16597085 | | Jiangxi Province Center for Disease Control and Prevention | Jiangxi Province Center for Disease Control and Prevention | JianXiong Li et al |
| **hCoV-19/Hainan/HCDC-252/2023** | EPI_ISL_16968133 | | Hainan Provincial Center for Disease Control and Prevention | Hainan Provincial Center for Disease Control and Prevention | Li Dandan et al |
| **hCoV-19/Henan/NYCDC-059/2023** | EPI_ISL_17186817 | | Nanyang Second People's Hospital | Nanyang Center for Disease Control and Prevention | ShiRui et al |
| **hCoV-19/Shaanxi/SCDC-0187/2023** | EPI_ISL_17494927 | | Shaanxi Provincial Center for Disease Control and Prevention | Shaanxi Provincial Center for Disease Control and Prevention | Meng et al |
| **hCoV-19/Tibet/XZCDC_0125/2023** | EPI_ISL_18624456 | | Inspection and verification office, Center for Disease Control and Prevention of Tibet Autonomous Region | Inspection and verification office, Center for Disease Control and Prevention of Tibet Autonomous Region | Hong et al |
| **hCoV-19/Guizhou/IVDC-1268/2022** | EPI_ISL_16636176 | | Guizhou Provincial Center for Disease Control and Prevention | China Center for Disease Control and Prevention, National Institute for Viral Disease Control and Prevention | ZhaoMingwu et al |
| **hCoV-19/Shandong/WCDC-157/2023** | EPI_ISL_18070820 | | Weihai Center for Disease Control and Prevention | Weihai Center for Disease Control and Prevention | Xiang Li et al |
| **hCoV-19/Heilongjiang/HLJCDC-0728/2023** | EPI_ISL_17492714 | | Department for Viral Disease Control and Prevention, Heilongjiang Provincial Center for Disease Control and Prevention | Department for Viral Disease Control and Prevention, Heilongjiang Provincial Center for Disease Control and Prevention | Xu Jun et al |
| **hCoV-19/Xinjiang/XJCDCCJ-020/2023** | EPI_ISL_17501705 | | Institute for the Prevention and Control of Infectious Diseases, Xinjiang Center for Disease Control and Prevention | Institute for the Prevention and Control of Infectious Diseases, Xinjiang Center for Disease Control and Prevention | Ma et al |
| **hCoV-19/Hubei/HCDC-WH0015/2022** | EPI_ISL_17487832 | | Hubei Provincial Center for Disease Control and Prevention | Hubei Provincial Center for Disease Control and Prevention | Yongzhong Jiang et al |
| **hCoV-19/Fujian/FCDC-202303S037/2023** | EPI_ISL_17262032 | | Fujian Provincial Center for Disease Control and Prevention | Fujian Provincial Center for Disease Control and Prevention | Lin Qi et al |
| **hCoV-19/Hubei/HCDC-0260/2022** | EPI_ISL_17487124 | | Hubei Provincial Center for Disease Control and Prevention | Hubei Provincial Center for Disease Control and Prevention | Yongzhong Jiang et al |
| **hCoV-19/Jiangxi/NCCDC-020/2023** | EPI_ISL_16597050 | | Nanchang City Center for Disease Control and Prevention | Nanchang City Center for Disease Control and Prevention | Qian Ke et al |
| **hCoV-19/Hubei/HCDC-WH0576/2023** | EPI_ISL_17487683 | | Hubei Provincial Center for Disease Control and Prevention | Hubei Provincial Center for Disease Control and Prevention | Yongzhong Jiang et al |
| **hCoV-19/Gansu/TSCDC-051103/2023** | EPI_ISL_17684474 | | Laboratory Department, Tianshui Center for Disease Control and Prevention | Laboratory Department, Tianshui Center for Disease Control and Prevention | Wenjing et al |
| **hCoV-19/Hubei/HCDC-753/2023** | EPI_ISL_17182643 | | Jingzhou Center for Disease Control and Prevention | Hubei Provincial Center for Disease Control and Prevention | Maoyi Chen et al |
| **hCoV-19/Hubei/HCDC-YC7/2022** | EPI_ISL_16617188 | | Yichang Center for Disease Control and Prevention | Hubei Provincial Center for Disease Control and Prevention | Jing Ma et al |
| **hCoV-19/Xinjiang/IVDC-1205/2022** | EPI_ISL_16636155 | | Xinjiang Center for Disease Control and Prevention | China Center for Disease Control and Prevention, National Institute for Viral Disease Control and Prevention | Xin Ma et al |
| **hCoV-19/Zhejiang/IVDC-837/2022** | EPI_ISL_16635541 | | Zhejiang Center for Disease Control and Prevention | China Center for Disease Control and Prevention, National Institute for Viral Disease Control and Prevention | Jun Li et al |
| **hCoV-19/Henan/NYCDC-009/2023** | EPI_ISL_16709099 | | Nanyang First People's Hospital | Nanyang Center for Disease Control and Prevention | Shi Rui et al |
| **hCoV-19/Jiangxi/NCCDC-094/2023** | EPI_ISL_17494151 | | Nanyang City Center for Disease Control and Prevention | Nanyang City Center for Disease Control and Prevention | Shi et al |
| **hCoV-19/Guangdong/GPIPH-2022A_XG35178/2022** | EPI_ISL_18924569 | | Guangdong Provincial Institution of Public Health, Guangdong Provinical Center for Disease Control and Prevention | Guangdong Provinical Center for Disease Control and Prevention | Zhonghua Zheng et al |
| **hCoV-19/Shanghai/IVDC-699/2023** | EPI_ISL_16636004 | | Shanghai Municipal Center For Disease Control and Prevention | China Center for Disease Control and Prevention, National Institute for Viral Disease Control and Prevention | JiangHui et al |
| **hCoV-19/Guangdong/GCDC-00173/2022** | EPI_ISL_16711855 | | Guangdong Provincial Center for Disease Control and Prevention | Guangdong Provincial Center for Disease Control and Prevention | Zhonghua Zheng et al |
| **hCoV-19/Guangdong/GDCDC-SZ-0028/2023** | EPI_ISL_17170915 | | Guangdong Provincial Center for Disease Control and Prevention | Guangdong Provincial Center for Disease Control and Prevention | Chang Zhang et al |
| **hCoV-19/Guizhou/IVDC-374/2023** | EPI_ISL_16636226 | | Guizhou Provincial Center for Disease Control and Prevention | China Center for Disease Control and Prevention, National Institute for Viral Disease Control and Prevention | ZhaoMingwu et al |
| **hCoV-19/Guangdong/GPIPH-GDCDC_CZ_0074/2023** | EPI_ISL_18926060 | | Guangdong Provincial Institution of Public Health, Guangdong Provinical Center for Disease Control and Prevention | Guangdong Provinical Center for Disease Control and Prevention | Zhonghua Zheng et al |
| **hCoV-19/Hainan/HCDC-436/2023** | EPI_ISL_17205158 | | Danzhou Center for Disease Control and Prevention | Hainan Provincial Center for Disease Control and Prevention | Liao Xuejing et al |
| **hCoV-19/Sichuan/SCDCP-4316/2022** | EPI_ISL_16306988 | | Sichuan Center of Disease Control and Prevention | Sichuan Center of Disease Control and Prevention | Xiaozhen Ma et al |
| **hCoV-19/Chongqing/CQCDC-5326/2022** | EPI_ISL_16698201 | | Chongqing Municipal Center for Disease Control and Prevention | Chongqing Municipal Center for Disease Control and Prevention | Ye Sheng et al |
| **hCoV-19/Chongqing/WZCDC-118/2022** | EPI_ISL_16698312 | | Wanzhou District Center for Disease Control and Prevention | Chongqing Municipal Center for Disease Control and Prevention | Ye Sheng et al |
| **hCoV-19/Jiangsu/XZCDC-38/2023** | EPI_ISL_16698432 | | Jiangsu Provincial Center for Disease Control and Prevention | Jiangsu Provincial Center for Disease Control and Prevention | Li Zhifeng et al |
| **hCoV-19/Chongqing/CQCDC-0124/2023** | EPI_ISL_16698293 | | Chongqing Municipal Center for Disease Control and Prevention | Chongqing Municipal Center for Disease Control and Prevention | Ye Sheng et al |
| **hCoV-19/Xinjiang/IVDC-235/2023** | EPI_ISL_16635948 | | Xinjiang Center for Disease Control and Prevention | China Center for Disease Control and Prevention, National Institute for Viral Disease Control and Prevention | Xin Ma et al |
| **hCoV-19/Guangxi/GCDC-FCG-010513/2023** | EPI_ISL_17088757 | | Fangchenggang Center for Disease Control and Prevention | Guangxi Zhuang Autonomous Region Center for Disease Control and Prevention | Weihui Pan et al |
| **hCoV-19/Zhejiang/ZJU-S84/2023** | EPI_ISL_16899217 | | Zhejiang University School of Medicine, First Affiliated Hospital, National Clinical Research Center for Infectious Diseases, State Key Laboratory for Diagnosis and Treatment of Infectious Diseases | Zhejiang University School of Medicine, First Affiliated Hospital, National Clinical Research Center for Infectious Diseases, State Key Laboratory for Diagnosis and Treatment of Infectious Diseases | Hangping Yao et al |
| **hCoV-19/Sichuan/CDCDC-230294/2023** | EPI_ISL_17104812 | | Chengdu Center of Disease Control and Prevention | Sichuan Center of Disease Control and Prevention | Heng Chen et al |
| **hCoV-19/Yunnan/YCDC-35/2023** | EPI_ISL_17273685 | | Department of Acute Infectious Diseases Control and Prevention, Yunnan Center for Disease Control and Prevention | Department of Acute Infectious Diseases Control and Prevention, Yunnan Center for Disease Control and Prevention | Meiling Zhang et al |
| **hCoV-19/Yunnan/YNCDC-0467/2023** | EPI_ISL_17633016 | | The department for prevention and control of acute infectious diseases | The department for prevention and control of acute infectious diseases | Zhang et al B |
| **hCoV-19/Jilin/JLCDC-57293/2022** | EPI_ISL_16598660 | | Jilin Center for Disease Control and Prevention | Jilin Center for Disease Control and Prevention | Wu Donglin et al |
| **hCoV-19/Guangdong/GPIPH-XG_ZH_22284/2022** | EPI_ISL_18924572 | | Guangdong Provincial Institution of Public Health, Guangdong Provinical Center for Disease Control and Prevention | Guangdong Provinical Center for Disease Control and Prevention | Zhonghua Zheng et al |
| **hCoV-19/Anhui/BZCDC-16/2023** | EPI_ISL_17227544 | | Bozhou Municipal Center for Disease Control & Prevention | Bozhou Municipal Center for Disease Control & Prevention | Qingqiao et al |
| **hCoV-19/Anhui/HFCDC-1/2022** | EPI_ISL_16599033 | | Hefei Center for Disease Control and Prevention | Hefei Center for Disease Control and Prevention | Song Yongqing et al |
| **hCoV-19/Hebei/CD-16/2023** | EPI_ISL_17031858 | | Hebei Provincial Center for Disease Control and Prevention | Hebei Provincial Center for Disease Control and Prevention | Shiyou Liu et al |
| **hCoV-19/InnerMongolia/IVDC-1256/2022** | EPI_ISL_16635282 | | The Center for Disease Control and Prevention of Inner Mongolia | China Center for Disease Control and Prevention, National Institute for Viral Disease Control and Prevention | Yanbo Bai et al |
| **hCoV-19/Liaoning/LNCDC-029-2/2023** | EPI_ISL_17183562 | | Liaoning Center for Disease Control and Prevention | Liaoning Center for Disease Control and Prevention | Lingling Mao et al |
| **hCoV-19/Jiangsu/SZCDC-207/2023** | EPI_ISL_17072162 | | Suzhou CDC Lab | Jiangsu Provincial Center for Disease Control and Prevention | Zefeng Dong et al |
| **hCoV-19/Shaanxi/SXCDC-A023/2023** | EPI_ISL_16709516 | | Ankang Center for Disease Control and Prevention | Shaanxi Provincial Center for Disease Control and Prevention | Bin Chen et al |
| **hCoV-19/Hunan/HuNCDC-1916/2022** | EPI_ISL_16390373 | | Loudi Center for Disease Control and Prevention | Hunan Provincial Center for Disease Control and Prevention | Chenshuai et al |
| **hCoV-19/Tianjin/TJCDC-002/2022** | EPI_ISL_16481837 | | Tianjin Centers for Disease Control and Prevention | Tianjin Centers for Disease Control and Prevention | Xiaoyan Li et al |
| **hCoV-19/Jilin/JLCDC-59476/2023** | EPI_ISL_16713275 | | Jilin Center for Disease Control and Prevention | Jilin Center for Disease Control and Prevention | Wu Donglin et al |
| **hCoV-19/Heilongjiang/HLJCDC-0279/2023** | EPI_ISL_17199735 | | Department for Viral Disease Control and Prevention, Heilongjiang Provincial Center for Disease Control and Prevention | Department for Viral Disease Control and Prevention, Heilongjiang Provincial Center for Disease Control and Prevention | Xiafengyang et al |
| **hCoV-19/Liaoning/SYCDC-0017/2023** | EPI_ISL_17493403 | | Shenyang Center for Disease Control and Prevention | Shenyang Center for Disease Control and Prevention | Sun et al |
| **hCoV-19/Guangxi/GCDC-LZ-120712/2022** | EPI_ISL_17088781 | | Liuzhou Center for Disease Control and Prevention | Guangxi Zhuang Autonomous Region Center for Disease Control and Prevention | Liujun Chen et al |
| **hCoV-19/Hebei/SJZ-3/2022** | EPI_ISL_17031529 | | Hebei Provincial Center for Disease Control and Prevention | Hebei Provincial Center for Disease Control and Prevention | Shiyou Liu et al |
| **hCoV-19/Shanghai/IVDC-1987/2022** | EPI_ISL_16634880 | | Shanghai Municipal Center For Disease Control and Prevention | China Center for Disease Control and Prevention, National Institute for Viral Disease Control and Prevention | JiangHui et al |
| **hCoV-19/Tianjin/TJCDC-058/2023** | EPI_ISL_16597944 | | Tianjin Centers for Disease Control and Prevention | Tianjin Centers for Disease Control and Prevention | Xiaoyan Li et al |
| **hCoV-19/Sichuan/SCDC-10/2023** | EPI_ISL_16737897 | | Sichuan Center for Disease Control and Prevention | Sichuan Center for Disease Control and Prevention | Jianan Xu et al |
| **hCoV-19/Heilongjiang/HLJCDC-0030/2023** | EPI_ISL_16598382 | | Heilongjiang Provincial Center for Disease Control and Prevention, Department for Viral Disease Control and Prevention | Heilongjiang Provincial Center for Disease Control and Prevention, Department for Viral Disease Control and Prevention | Xu Jun et al |
| **hCoV-19/Hebei/ZJK-62/2023** | EPI_ISL_17031739 | | Hebei Provincial Center for Disease Control and Prevention | Hebei Provincial Center for Disease Control and Prevention | Shiyou Liu et al |
| **hCoV-19/Qinghai/QCDC-QHSYY-84/2023** | EPI_ISL_17075751 | | Qinghai Provincial People's Hospital | Qinghai Center for Disease Prevention and Control | Youju Lei et al |
| **hCoV-19/Henan/XCDC-030803/2023** | EPI_ISL_17488890 | | Xinxiang Center for Disease Control and Prevention | Xinxiang Center for Disease Control and Prevention | Ma Jianmin et al |
| **hCoV-19/Beijing/BJCDC-0762/2023** | EPI_ISL_17592206 | | Institute for Infectious Disease and Endemic Disease Control, Beijing Center for Disease Control and Prevention | Institute for Infectious Disease and Endemic Disease Control, Beijing Center for Disease Control and Prevention | Yang Pan et al |
| **hCoV-19/InnerMongolia/NMCDC-ULQ-001/2023** | EPI_ISL_17785595 | | Immunization Branch, Inner Mongolia Center for Disease Control and Prevention | Immunization Branch, Inner Mongolia Center for Disease Control and Prevention | Li et al |
| **hCoV-19/Shanxi/TYCDC-043/2023** | EPI_ISL_17493987 | | Taiyuan Center for Disease Control and Prevention | Taiyuan Center for Disease Control and Prevention | Wang et al |
| **hCoV-19/Kenya/SS8362/2022** | EPI_ISL_12905592 | ON603278.1 | Pathologist Lancet Kenya | KEMRI-Wellcome Trust Research Programme,Kilifi | Githinji G. et al |
| **hCoV-19/Bangladesh/icddrb-G142063/2022** | EPI_ISL_11204830 | | Virology Laboratory, International Centre for Diarrhoeal Disease Research, Bangladesh (ICDDR,B) | Virology Laboratory, International Centre for Diarrhoeal Disease Research, Bangladesh (ICDDR,B) | Mohammad Jubair et al |
| **hCoV-19/India/AP-CCMB-CID2222/2022** | EPI_ISL_13286056 | | AP SSO | CSIR-Centre for Cellular and Molecular Biology-INSACOG | Pramada Prasad et al |
| **hCoV-19/Malaysia/IMR_OS2055/2022** | EPI_ISL_14195936 | | Institute for Medical Research, Infectious Disease Research Centre, National Institutes of Health, Ministry of Health Malaysia | Institute for Medical Research, Infectious Disease Research Centre, National Institutes of Health, Ministry of Health Malaysia | Kalyanasundram J et al |
| **hCoV-19/Malaysia/IMR_OS1806/2022** | EPI_ISL_14195690 | | Institute for Medical Research, Infectious Disease Research Centre, National Institutes of Health, Ministry of Health Malaysia | Institute for Medical Research, Infectious Disease Research Centre, National Institutes of Health, Ministry of Health Malaysia | Kalyanasundram J et al |
| **hCoV-19/India/JK-NCDC-7709373/2021** | EPI_ISL_11062734 | | National Centre for Disease Control (NCDC) Biotechnology Division, Delhi - INSACOG | NCDC Delhi, Biotechnology Division -INSACOG | Kalaiarasan Ponnusamy et al |
| **hCoV-19/Panama/CDED23099-GMI/2022** | EPI_ISL_16027400 | | Centro de Diagnostico de Enfermedades Desatendidas | Gorgas Memorial Institute of Health Studies | Eduardo Aguirre et al |
| **hCoV-19/India/MH-ICMR-NIV-INSACOG-10347/2022** | EPI_ISL_13079842 | | ICMR-National Institute of Virology - INSACOG | NIV Influenza | Dr. Varsha Potdar et al |
| **hCoV-19/India/HP-ICMR-MCL-22-4333-10723/2022** | EPI_ISL_13906935 | | Indian Council of Medical Research-National Institute of Virology, Microbial Containment Complex | Indian Council of Medical Research-National Institute of Virology, Microbial Containment Complex | Pragya Yadav et al |
| **hCoV-19/India/AN-ICMR-NIV-INSACOG-GSEQ-10043/2022** | EPI_ISL_10801582 | | ICMR-National Institute of Virology - INSACOG | NIV Influenza | Dr. Varsha Potdar et al |
| **hCoV-19/Mozambique/CERI-KRISP-K037911/2022** | EPI_ISL_11621077 | | Instituto Nacional de Saude (INS), Mozambique | CERI, Centre for Epidemic Response and Innovation, Stellenbosch University and KRISP, KZN Research Innovation and Sequencing Platform, UKZN. | Nalia Ismael et al |
| **hCoV-19/Oman/5225281/2022** | EPI_ISL_16073626 | | Oman-NIC | Oman-NIC | Samiha Al Kharusi et al |
| **hCoV-19/India/UP-NCDC-8109117/2022** | EPI_ISL_11074864 | | National Centre for Disease Control (NCDC) Biotechnology Division, Delhi - INSACOG | NCDC Delhi, Biotechnology Division -INSACOG | Kalaiarasan Ponnusamy et al |
| **hCoV-19/India/BR-NCDC-7809256/2021** | EPI_ISL_11064084 | | National Centre for Disease Control (NCDC) Biotechnology Division, Delhi - INSACOG | NCDC Delhi, Biotechnology Division -INSACOG | Kalaiarasan Ponnusamy et al |
| **hCoV-19/India/AP-CCMB-CIC4697/2022** | EPI_ISL_11001396 | | AP SSO | CSIR-Centre for Cellular and Molecular Biology-INSACOG | Valli Nagalakshmi Undamatla et al |
| **hCoV-19/India/SK-INSACOG-1122500126136/2022** | EPI_ISL_10434469 | | INSACOG-Sikkim | National Institute of Biomedical Genomics – INSACOG | Arindam Maitra et al A |
| **hCoV-19/India/KA-SLS-31104/2022** | EPI_ISL_11790056 | | | Strand Life Sciences (supported by IN COVID Support, FZE LLC) | Anson Kunjumon George et al |
| **hCoV-19/India/HP-INSACOG/NII-235/2022** | EPI_ISL_11262593 | | Dr RPGMC Tanda | National Institute of Immunology | Dr. G. Aneeshkumar Arimbasseri et al |
| **hCoV-19/India/KA-0902BENU006/2022** | EPI_ISL_10065720 | | CBR,IISc | CBR,IISc | Bratati Kahali et al |
| **hCoV-19/India/WB-INSACOG-1930900399628/2022** | EPI_ISL_12748920 | | INSACOG-WB | National Institute of Biomedical Genomics | Arindam Maitra et al A |
| **hCoV-19/India/RJ-SMS-ICMR-INSACOG-TS-12945/2022** | EPI_ISL_11881850 | | SMS MEDICAL COLLEGE,JAIPUR | NIV Influenza | Bharti Malhotra et al |
| **hCoV-19/India/MZ-INSACOG-1526600094191/2022** | EPI_ISL_13159267 | | INSACOG-Mizoram | National Institute of Biomedical Genomics – INSACOG | Arindam Maitra et al A |
| **hCoV-19/Bangladesh/CHRF-GSB04-153/2022** | EPI_ISL_11704654 | | Child Health Research Foundation | Child Health Research Foundation | CHRF Bangladesh Genomics Team et al |
| **hCoV-19/India/MZ-INSACOG-1526101090572/2022** | EPI_ISL_13876773 | | INSACOG-Mizoram | National Institute of Biomedical Genomics – INSACOG | Arindam Maitra et al A |
| **hCoV-19/India/KL-ILSGS17589/2022** | EPI_ISL_14749055 | | ILS, Bhubaneswar | Institute of Life Sciences-INSACOG | Sunil K. Raghav et al |
| **hCoV-19/India/RJ-SMS-ICMR-INSACOG-TS-2641/2022** | EPI_ISL_11610260 | | SMS MEDICAL COLLEGE,JAIPUR | NIV Influenza | Bharti Malhotra et al |
| **hCoV-19/India/AN-ICMR-NIV-INSACOG-GSEQ-10050/2022** | EPI_ISL_10801590 | | ICMR-National Institute of Virology - INSACOG | NIV Influenza | Dr. Varsha Potdar et al |
| **hCoV-19/USA/IA-CDC-LC0574018/2022** | EPI_ISL_12168002 | ON295350.1 | Laboratory Corporation of America | Centers for Disease Control and Prevention Division of Viral Diseases, Pathogen Discovery | Dakota Howard et al |
| **hCoV-19/Nepal/EPN-B1395/2022** | EPI_ISL_13330930 | | Bheri Hospital | Center for Molecular Dynamics Nepal | Epidemic Intelligence Team et al |
| **hCoV-19/India/WB-INSACOG-1931502934078/2022** | EPI_ISL_10123738 | | INSACOG-WB | National Institute of Biomedical Genomics – INSACOG | Arindam Maitra et al A |
| **hCoV-19/Yunnan/IVDC-563/2022** | EPI_ISL_11873926 | | Department of Acute Infectious Diseases Control and Prevention, Yunnan Center for Disease Control and Prevention | National Institute for Viral Disease Control and Prevention, China CDC | Meiling Zhang et al |
| **hCoV-19/India/AP-CCMB-CIC8741/2022** | EPI_ISL_12305399 | | AP SSO | CSIR-Centre for Cellular and Molecular Biology-INSACOG | Sumedha Avadhanula et al |
| **hCoV-19/USA/WY-WYPHL-22039634/2022** | EPI_ISL_12226277 | | Wyoming Public Health Laboratory | Wyoming Public Health Laboratory | Jim Mildenberger et al |
| **hCoV-19/India/WB-INSACOG-1930500403784/2022** | EPI_ISL_13408892 | | INSACOG-WB | National Institute of Biomedical Genomics – INSACOG | Arindam Maitra et al A |
| **hCoV-19/NewZealand/22CV6899/2022** | EPI_ISL_13764619 | | Wellington SCL (WN) | Institute of Environmental Science and Research (ESR) | Sarah Cockerton et al |
| **hCoV-19/Bangladesh/IEDCR-JIMC-182/2022** | EPI_ISL_14869954 | | Institute of Epidemiology, Disease Control and Research (IEDCR) | Institute of Epidemiology, Disease Control and Research (IEDCR) | Manjur Hossain Khan et al |

| | | | | | |
|---|---|---|---|---|---|
| **hCoV-19/India/AN-ICMR-NIV-INSACOG-GSEQ-10052/2022** | EPI_ISL_10801592 | | ICMR-National Institute of Virology - INSACOG | NIV Influenza | Dr. Varsha Potdar et al |
| **hCoV-19/England/BRBR-3A7F546/2022** | EPI_ISL_11274408 | OW269402.1 | Berkshire and Surrey Pathology Services Lighthouse Laboratory | Wellcome Sanger Institute for the COVID-19 Genomics UK (COG-UK) Consortium | Berkshire et al |
| **hCoV-19/India/WB-INSACOG-1930400988433/2022** | EPI_ISL_13158941 | | INSACOG-WB | National Institute of Biomedical Genomics – INSACOG | Arindam Maitra et al A |
| **hCoV-19/Philippines/PGCV_0010_0404/2022** | EPI_ISL_12852728 | | METRO ILOILO HOSPITAL AND MEDICAL CENTER | PHILIPPINE GENOME CENTER VISAYAS | Dr. Cynthia P. Saloma et al |
| **hCoV-19/Myanmar/DSMRC-078/2022** | EPI_ISL_11149537 | ON038755.1 | Defence Services Medical Research Centre (DSMRC) | Defence Services Medical Research Centre (DSMRC) | Khine Zaw Oo et al |
| **hCoV-19/Shandong/IVDC-585/2022** | EPI_ISL_11905850 | | Shandong Center for Disease Control and Prevention | National Institute for Viral Disease Control and Prevention, China CDC | Mingxiao Yao et al |
| **hCoV-19/Myanmar/DSMRC-090/2022** | EPI_ISL_11149564 | ON038763.1 | DSMRC | Defence Services Medical Research Centre (DSMRC) | Khine Zaw Oo et al |
| **hCoV-19/India/DL-ILBS-WGS2315/2021** | EPI_ISL_8769910 | | MAHAJAN IMAGING | ILBS | Ekta Gupta et al |
| **hCoV-19/India/NL-IBSD-DMP-P13-164/2021** | EPI_ISL_11465733 | | DSO, DIMAPUR | INSACOG-IBSD Manipur | Dr. Nanaocha Sharma et al |
| **hCoV-19/India/KL-INSACOG-1931502632525/2021** | EPI_ISL_9472839 | | INSACOG-WB | National Institute of Biomedical Genomics – INSACOG | Arindam Maitra et al A |
| **hCoV-19/India/OR-ILSGS14645/2022** | EPI_ISL_10433694 | | ILS, Bhubaneswar | Institute of Life Sciences-INSACOG | Sunil K. Raghav et al |
| **hCoV-19/India/OR-ILSGS14699/2022** | EPI_ISL_10433742 | | ILS, Bhubaneswar | Institute of Life Sciences-INSACOG | Sunil K. Raghav et al |
| **hCoV-19/India/AR-CSIR-NEIST-INSACOG-NEIST23-CID0530/2022** | EPI_ISL_9974974 | | TRIHMS, Naharlagun | Centre for Infectious Diseases, CSIR-NEIST | Romi Wahengbam et al |
| **hCoV-19/India/CT-ILSGS15423/2022** | EPI_ISL_10436723 | | GMC, Raigarh | Institute of Life Sciences-INSACOG | Sunil K. Raghav et al |
| **hCoV-19/India/AP-CCMB-CIC8128/2022** | EPI_ISL_11333822 | | AP SSO | CSIR-Centre for Cellular and Molecular Biology-INSACOG | Malini Nemalikanti et al |
| **hCoV-19/Bangladesh/icddrb-GAC2-381/2022** | EPI_ISL_13477152 | | Virology Laboratory, International Centre for Diarrhoeal Disease Research, Bangladesh (ICDDR,B) | Virology Laboratory, International Centre for Diarrhoeal Disease Research, Bangladesh (ICDDR,B) | Mohammad Jubair et al |
| **hCoV-19/SouthAfrica/NICD-N33618/2022** | EPI_ISL_9963219 | | AMPATH | National Institute for Communicable Diseases of the National Health Laboratory Service | Amoako DG et al |
| **hCoV-19/Poland/2462WSN2022_wsseol/2022** | EPI_ISL_10687025 | | WSSE w Bydgoszczy | Wojewodzka Stacja Sanitarno-Epidemiologiczna w Olsztynie, Laboratorium Badan Epidemiologiczno-Klinicznych | Sylwia Krzetowska et al |
| **hCoV-19/Zambia/CHAZ-CHS2034/2022** | EPI_ISL_13228475 | | ZNPHL | CHAZ Laboratory | Shempela. D et al |
| **hCoV-19/Turkey/HSGM-GS4240/2022** | EPI_ISL_13203212 | | Ministry of Health Turkey | Ministry of Health Turkey | Fatma Bayrakdar et al |
| **hCoV-19/Mexico/GRO-INMEGEN-68-17/2022** | EPI_ISL_12401387 | | SALUD DIGNA | Instituto Nacional de Medicina Genomica | Cedro-Tanda A et al |
| **hCoV-19/Bahrain/22020076982/2022** | EPI_ISL_14063582 | ON197716.1 | Communicable Disease Laboratory, Public Health Directorate | Communicable Disease Laboratory, Public Health Directorate | Marhoon et al |
| **hCoV-19/Thailand/NIC_BKK_SJ0538/2022** | EPI_ISL_16061473 | | Siriraj Hospital | National Institute of Health, Department of Medical Sciences, Ministry of Public Health, Thailand | Pilailuk Okada et al |
| **hCoV-19/Tunisia/14107/2022** | EPI_ISL_15298442 | | 1-Laboratory of Microbiology, National Reference Lab, Charles Nicolle Hospital; 2-University of Tunis ElManar, Faculty of Medicine of Tunis, LR99ES09, Tunis, Tunisia | Clinical and Experimental Pharmacology Lab, LR16SP02, National Center of Pharmacovigilance, University of Tunis El Manar, Tunis, Tunisia. 2- Ministry of Health, National Observatory of New and Emerging Diseases, 1006, Tunis, Tunisia | Sana Ferjani et al |
| **hCoV-19/Kuwait/Jaber-20148241/2022** | EPI_ISL_13477523 | | Jaber Al Ahmad Hospital | Virology Lab, Jaber Al Ahmad Hospital | Dr. Zahrah Buhamad et al |
| **hCoV-19/Philippines/PH-PGC-113044/2022** | EPI_ISL_13169216 | | PHILIPPINE GENERAL HOSPITAL (PGH) | Philippine Genome Center | Cynthia P. Saloma et al |
| **hCoV-19/Indonesia/JK-CW62-0437/2022** | EPI_ISL_11587089 | | RS Paru dr M Goenawan Partowidigdo Cisarua | National Quality Control Laboratory of Drug and Food | Sri Utaminingsih et al |
| **hCoV-19/Switzerland/FR-ETHZ-37082303/2022** | EPI_ISL_12130079 | OW566675.1 | Viollier AG | Department of Biosystems Science and Engineering, ETH Zürich | Christian Beisel et al |
| **hCoV-19/Mexico/CMX_InDRE_FB14337_E09316859587_S13439/2022** | EPI_ISL_13192167 | | LESP Mexico City | Instituto de Diagnostico y Referencia Epidemiologicos (INDRE) | Claudia Wong-Arambula et al |
| **hCoV-19/Georgia/Tb-s5257/2022** | EPI_ISL_13884652 | | National Center for Disease Control and Public Health (NCDC) of Georgia. | Department for Virology, Molecular Biology and Genome Research, R. G. Lugar Center for Public Health Research, National Center for Disease Control and Public Health (NCDC) of Georgia. | Giorgi Tomashvili et al |
| **hCoV-19/Qatar/DA-USAFSAM-S15229/2022** | EPI_ISL_12185174 | | US Air Force School of Aerospace Medicine | US Air Force School of Aerospace Medicine | Anthony Fries et al |
| **hCoV-19/Botswana/R102B53_BHP_985655/2022** | EPI_ISL_10967592 | | Botswana Harvard HIV Reference Laboratory | Botswana Harvard HIV Reference Laboratory | Sikhulile Moyo et al |
| **hCoV-19/Lebanon/MA-3/2022** | EPI_ISL_16583752 | | Deir Saydet El Maaounat Hospital | Melhem/Matar lab | Nada M. Melhem et al |
| **hCoV-19/Botswana/R107B09_BHP_AAC78597/2022** | EPI_ISL_12090017 | | Nyangabgwe Referral Hospital Laboratory | Botswana Harvard HIV Reference Laboratory | Sikhulile Moyo et al |
| **hCoV-19/Myanmar/DSMRC-097/2022** | EPI_ISL_11149643 | ON038769.1 | DSMRC | Defence Services Medical Research Centre (DSMRC) | Khine Zaw Oo et al |
| **hCoV-19/Sweden/4073556260/2022** | EPI_ISL_11844796 | | The Public Health Agency of Sweden | The Public Health Agency of Sweden | Emmi Andersson et al |
| **hCoV-19/India/GJ-INSACOG-GBRC5795/2022** | EPI_ISL_10749083 | | Supratech Micropath Diagnostics & Research laboratory Ahmedabad | Gujarat Biotechnology Research Centre | Zarna Patel et al |
| **hCoV-19/NewZealand/22CV1851/2022** | EPI_ISL_10187689 | | PathLab Bay of Plenty | Institute of Environmental Science and Research (ESR) | Sarah Cockerton et al |
| **hCoV-19/CookIslands/CV4662/2022** | EPI_ISL_18209498 | | Cook Islands Ministry of Health | Institute of Environmental Science and Research (ESR) | Theresa Tatuava et al |
| **hCoV-19/NewZealand/22CV6909/2022** | EPI_ISL_13764306 | | Wellington SCL (WN) | Institute of Environmental Science and Research (ESR), Kenepuru Science Centre | Sarah Cockerton et al |
| **hCoV-19/Nepal/NPHL-S-1471/2022** | EPI_ISL_12140454 | | Decode Genomics and Research Center | National Public Health Laboratory | National Public Health Laboartory Team/Decode Genomics et al |
| **hCoV-19/Georgia/Tb-s4409/2022** | EPI_ISL_9879875 | | R. G. Lugar Center for Public Health Research, National Center for Disease Control and Public Health (NCDC) of Georgia. | Department for Virology, Molecular Biology and Genome Research, R. G. Lugar Center for Public Health Research, National Center for Disease Control and Public Health (NCDC) of Georgia. | Giorgi Tomashvili et al |
| **hCoV-19/India/PB-VRDLGMCPTA-ICMR-INSACOG-623903604/2022** | EPI_ISL_14148162 | | Virus Research and Diagnostic Laboratory | NIV Influenza | Dr.Rupinder Bakshi et al |
| **hCoV-19/India/DH-ICMR-NIV-INSACOG-GSEQ-8891/2022** | EPI_ISL_9968841 | | ICMR-National Institute of Virology - INSACOG | NIV Influenza | Dr. Varsha Potdar et al |
| **hCoV-19/India/LA-INSACOG/NII-1155/2022** | EPI_ISL_13966912 | | COVID-19 Laboratory Chuchot SNM Hospital | National Institute of Immunology | Aneeshkumar Arimbasseri et al |
| **hCoV-19/India/PY-SEQ_12988_S270_R1_001/2022** | EPI_ISL_10729556 | | JIPMER | inStem, NCBS-INSACOG | Uma Ramakrishnan et al |
| **hCoV-19/India/MH-ICMR-NIV-INSACOG-10411/2022** | EPI_ISL_13079794 | | ICMR-National Institute of Virology - INSACOG | NIV Influenza | Dr. Varsha Potdar et al |
| **hCoV-19/Kuwait/Jaber-20276227/2022** | EPI_ISL_14175978 | | Mubarak Al-Kabeer Hospital | Virology Lab, Jaber Al Ahmad Hospital | Dr. Zahrah Buhamad et al |
| **hCoV-19/India/GJ-INSACOG-GBRC6569/2022** | EPI_ISL_11789535 | | Supratech Micropath Diagnostics & Research laboratory Ahmedabad | Gujarat Biotechnology Research Centre | Akhilesh Modi et al |
| **hCoV-19/India/AP-CCMB-CIC5052/2022** | EPI_ISL_11001712 | | AP SSO | CSIR-Centre for Cellular and Molecular Biology-INSACOG | Amareshwar Vodapalli et al |
| **hCoV-19/Chongqing/YBCDC-0068/2022** | EPI_ISL_16523420 | | Chongqing Yubei District Center for Disease Control and Prevention | Chongqing Yubei District Center for Disease Control and Prevention | Jin Yan et al |
| **hCoV-19/India/GJ-INSACOG-GBRC5908/2022** | EPI_ISL_10749187 | | Supratech Micropath Diagnostics & Research laboratory Ahmedabad | Gujarat Biotechnology Research Centre | Bhadreshsinh Gohil et al |
| **hCoV-19/India/BR-INSACOG-GBRC5890/2022** | EPI_ISL_10749172 | | In Vitro Speciality Laboratory, Ahmedabad | Gujarat Biotechnology Research Centre | Disha Vora et al |
| **hCoV-19/India/LD-ICMR-NIV-INSACOG-GSEQ-9977/2022** | EPI_ISL_10801681 | | ICMR-National Institute of Virology - INSACOG | NIV Influenza | Dr. Varsha Potdar et al |
| **hCoV-19/India/UP-ICMR-INSACOG-KGMU-ZY39/352/2022** | EPI_ISL_12492767 | | Department of Microbiology,King George's Medical College | NIV Influenza | Prof. Amita Jain et al |
| **hCoV-19/Cyprus/BMV6692/2022** | EPI_ISL_18479915 | | Medicover Genetics | Laboratory of Biotechnology and Molecular Virology, University of Cyprus | Leondios G. Kostrikis et al. |
| **hCoV-19/Australia/NSW-SAVID-11507/2022** | EPI_ISL_12666382 | | Area of Virology, Serology and Virology Division (SAViD), New South Wales Health Pathology Randwick | Virology Research Laboratory; Area of Virology, Serology and Virology Division (SAViD), New South Wales Health Pathology Randwick | Foster et al |
| **hCoV-19/Canada/NB-CHUDGLD-120422-MM01072R/2022** | EPI_ISL_12227982 | | New Brunswick - Vitalite Health Network | New Brunswick - Vitalite Health Network | Chacko S. et al |
| **hCoV-19/Australia/NSW-ICPMR-30524/2022** | EPI_ISL_14285173 | | 4Cyte Pathology | Institute of Clinical Pathology and Medical Research, Westmead Hospital | Arnott A. et al |
| **hCoV-19/Estonia/RC0707395/2022** | EPI_ISL_13805370 | | Synlab Eesti OÜ | 1. Laboratory of Communicable Diseases (Estonia); 2. Eurofins Genomics Europe Sequencing GmbH | Abroi A. et al |
| **hCoV-19/Curacao/CW-RIVM-100456/2022** | EPI_ISL_12953382 | | National Institute for Public Health and the Environment (RIVM) | National Institute for Public Health and the Environment (RIVM) | Adam Meijer et al |
| **hCoV-19/Mongolia/UB-60533/2022** | EPI_ISL_14713973 | | National Center for Communicable Diseases (NCCD) National Influenza Center | National Center for Communicable Diseases (NCCD) National Influenza Center | Naranzul Tsedenbal et al |
| **hCoV-19/Mongolia/UB-64059/2022** | EPI_ISL_14724028 | | National Center for Communicable Diseases (NCCD) National Influenza Center | National Center for Communicable Diseases (NCCD) National Influenza Center | Naranzul Tsedenbal et al |
| **hCoV-19/Israel/ICH-741218723/2022** | EPI_ISL_12376291 | | Clinical Microbiology Laboratory, Tel Aviv Sourasky Medical Center | Clinical Microbiology Laboratory, Tel Aviv Sourasky Medical Center | Orly Eshel et al |
| **hCoV-19/Uganda/517/2022** | EPI_ISL_13952139 | | Central Public Health Laboratories | Central Public Health Laboratories | Alisen Ayitewala et al |
| **hCoV-19/Kenya/C117524/2022** | EPI_ISL_13460285 | ON844294.1 | KEMRI-Wellcome Trust Research Programme | KEMRI-Wellcome Trust Research Programme | Githinji G. et al |
| **hCoV-19/HongKong/VM21048494/2021** | EPI_ISL_9643316 | | North Lantau Hospital | Hong Kong Department of Health | Alan K.L. Tsang et al |
| **hCoV-19/Philippines/PGCV_004_0154/2022** | EPI_ISL_12852521 | | WESTERN VISAYAS MEDICAL CENTER SUB NATIONAL LABORATORY | PHILIPPINE GENOME CENTER VISAYAS | Dr. Cynthia P. Saloma et al |
| **hCoV-19/Uruguay/DLSP_1759/2022** | EPI_ISL_17227697 | | Departamento Laboratorios de Salud Publica (DLSP) Ministerio de Salud Publica | Departamento Laboratorios de Salud Publica (DLSP) Ministerio de Salud Publica | Cortinas et al |
| **hCoV-19/Philippines/PH-PGC-113211/2022** | EPI_ISL_13169110 | | SOUTHERN PHILIPPINES MEDICAL CENTER | Philippine Genome Center | Cynthia P. Saloma et al |
| **hCoV-19/Vietnam/PIHCM_705/2022** | EPI_ISL_11058602 | | Pasteur Institute in Ho Chi Minh city | Department of Microbiology and Immunology | Hang Pham Thi Thu et al |
| **hCoV-19/Guangxi/GCDC-BS-2208/2022** | EPI_ISL_17104931 | | Baise Center for Disease Control and Prevention | Guangxi Zhuang Autonomous Region Center for Disease Control and Prevention | Zan Chen et al |
| **hCoV-19/Libya/79742-96/2022** | EPI_ISL_13505286 | | Reference Lab for Public Health, NCDC | Reference Lab for Public Health, NCDC | Abdulla Bashein et al |
| **hCoV-19/Brunei/9222123474/2022** | EPI_ISL_12261224 | | Departmenrt of Laboratory Services, National Virology Reference Laboratory | Microbial Genomic Services, Clinical Molecular Diagnostic Laboratory For Infectious Disease | Zainun Zaini et al |
| **hCoV-19/Brunei/7122068766/2022** | EPI_ISL_12261241 | | Departmenrt of Laboratory Services, National Virology Reference Laboratory | Microbial Genomic Services, Clinical Molecular Diagnostic Laboratory For Infectious Disease | Zainun Zaini et al |
| **hCoV-19/Laos/LOMWRU-0238/2022** | EPI_ISL_13094353 | OP159580.1 | LOMWRU | LOMWRU/Microbiology Laboratory, Mahosot Hospital | Manila Souksavanh et al |
| **hCoV-19/SouthKorea/KDCA90475/2022** | EPI_ISL_14017180 | | Division of Emerging Infectious Diseases, Bureau of Infectious Diseases Diagnosis Control, Korea Disease Control and Prevention Agency | Division of Emerging Infectious Diseases, Bureau of Infectious Diseases Diagnosis Control, Korea Disease Control and Prevention Agency | Jeong-Ah Kim et al |
| **hCoV-19/Vietnam/NHTD-OUCRU3121/2022** | EPI_ISL_14691876 | | National Hospital for Tropical Diseases | Oxford University Clinical Research Unit, Hanoi, Vietnam | Pham Ngoc Thach et al |
| **hCoV-19/Malaysia/UNIMAS-B17189/2022** | EPI_ISL_18889726 | | Borneo Medical Centre (Kuching) | Institute of Health and Community Medicine | Chan Chia Jui et al |
| **hCoV-19/Vietnam/BMH-200822-41/2022** | EPI_ISL_14685759 | | CDC Ha Noi | Bachmai Hospital | Phuong Truong et al |
| **hCoV-19/Taiwan/NTU-HCH-133/2022** | EPI_ISL_14189495 | | Department of Laboratory Medicine, National Taiwan University Hospital Hsin-Chu Hospital, Taiwan | Research Center for Emerging Viral Infections, Chang Gung University, Taiwan | Pei-Lan Shao et al |
| **hCoV-19/Taiwan/NTU266o/2022** | EPI_ISL_16047950 | | Department of Laboratory Medicine, National Taiwan University Hospital | Department of Clinical Laboratory Sciences and Medical Biotechnology, National Taiwan University | Shiou-Hwei Yeh et al |
| **hCoV-19/Taiwan/NTU-HCH-1877/2022** | EPI_ISL_14191231 | | Department of Laboratory Medicine, National Taiwan University Hospital Hsin-Chu Hospital, Taiwan | Research Center for Emerging Viral Infections, Chang Gung University, Taiwan | Pei-Lan Shao et al |
| **hCoV-19/Taiwan/NTU271/2022** | EPI_ISL_15908758 | | Department of Laboratory Medicine, National Taiwan University Hospital | Department of Clinical Laboratory Sciences and Medical Biotechnology, National Taiwan University | Shiou-Hwei Yeh et al |
| **hCoV-19/Taiwan/NCKUH-061022009875/2022** | EPI_ISL_18851391 | | Virology Lab,Department of Pathology, National Cheng Kung University Hospital | Virology Lab,Department of Pathology, National Cheng Kung University Hospital | Huey-Pin Tsai et al |
| **hCoV-19/Taiwan/NCKUH-823622126990/2022** | EPI_ISL_18716454 | | Virology Lab,Department of Pathology, National Cheng Kung University Hospital | Virology Lab,Department of Pathology, National Cheng Kung University Hospital | Huey-Pin Tsai et al |
| **hCoV-19/Guangdong/GPCDCP-2/2022** | EPI_ISL_16127643 | | Guangdong Provincial Center for Disease Control and Prevention | Key Laboratory of Human Diseases Comparative Medicine, Institute of Laboratory Animal Science, CAMS&PUMC | Bao et al |
| **hCoV-19/Cambodia/VIRU22-03972/2022** | EPI_ISL_11023877 | | National Institute of Public Health | Virology Unit, Institut Pasteur du Cambodge | Leakhena Pum et al |
| **hCoV-19/Cambodia/934362/2022** | EPI_ISL_12089783 | | Virology Unit, Institut Pasteur du Cambodge | Virology Unit, Institut Pasteur du Cambodge | Leakhena Pum et al |
| **hCoV-19/Brunei/9222117777/2022** | EPI_ISL_12261198 | | Departmenrt of Laboratory Services, National Virology Reference Laboratory | Microbial Genomic Services, Clinical Molecular Diagnostic Laboratory For Infectious Disease | Zainun Zaini et al |
| **hCoV-19/Brunei/7122067613/2022** | EPI_ISL_11515057 | | National Virology Reference Laboratory | Microbial Genomic Services | Zainun Zaini et al |
| **hCoV-19/USA/AK-CDC-2-5699073/2022** | EPI_ISL_11101318 | ON001282.1 | "AK State Public Health Lab, State Health Department" | Centers for Disease Control and Prevention Division of Viral Diseases, Pathogen Discovery | Mili Sheth et al |
| **hCoV-19/Mongolia/UB-58546/2022** | EPI_ISL_14713979 | | National Center for Communicable Diseases (NCCD) National Influenza Center | National Center for Communicable Diseases (NCCD) National Influenza Center | Naranzul Tsedenbal et al |
| **hCoV-19/Ghana/WACCBIP-GS2988/2022** | EPI_ISL_13392339 | | WACCBIP, University of Ghana | WACCBIP, University of Ghana | Collins M. Morang'a et al |
| **hCoV-19/Djibouti/NAMRU3_E501/2022** | EPI_ISL_13719655 | ON952000.1 | Naval Medical Research Unit No. 3 | Naval Medical Research Center Biological Defense Research Directorate | Catherine E. Arnold et al |
| **hCoV-19/Australia/VIC58342/2022** | EPI_ISL_12979537 | | Microbiological Diagnostic Unit - Public Health Laboratory (MDU-PHL) | Microbiological Diagnostic Unit - Public Health Laboratory (MDU-PHL) | Seemann et al |
| **hCoV-19/Malaysia/UNIMAS-B17591/2022** | EPI_ISL_12510237 | | Rejang Medical Centre (Sibu) | Institute of Health and Community Medicine | Chan Chia Jui et al |
| **hCoV-19/Libya/80894-93/2022** | EPI_ISL_13505289 | | Abushosha Triage Venter | Reference Lab for Public Health, NCDC | Abdulla Bashein et al |
| **hCoV-19/Canada/MB-CPL-421008/2022** | EPI_ISL_12692136 | | MB-Cadham Provincial laboratory | Manitoba Cadham Provincial Laboratory | David Alexander et al |
| **hCoV-19/Indonesia/KB-WGS-FKUNTAN-12469/2022** | EPI_ISL_13985491 | | Tanjungpura University Hospital | Faculty of Medicine, Tanjungpura University | Andriani et al |
| **hCoV-19/Canada/MB-CPL-421105/2022** | EPI_ISL_12692218 | | MB-Cadham Provincial laboratory | Manitoba Cadham Provincial Laboratory | David Alexander et al |
| **hCoV-19/Guam/GU-H2212866/2022** | EPI_ISL_13765558 | | Guam Public Health Laboratory | State Laboratories Division, Hawaii State Department of Health | Pamela O'Brien et al |
| **hCoV-19/Singapore/5504/2022** | EPI_ISL_13058494 | | National Public Health Laboratory, National Centre for Infectious Diseases | National Public Health Laboratory, National Centre for Infectious Diseases | Benny Yeo et al |
| **hCoV-19/Zhejiang/ZPCDCP-87/2022** | EPI_ISL_17261589 | | Zhejiang Provincial Center for Disease Control and Prevention | Zhejiang Provincial Center for Disease Control and Prevention | Yi Sun et al |
| **hCoV-19/Taiwan/NTU-HCH-697/2022** | EPI_ISL_14190057 | | Department of Laboratory Medicine, National Taiwan University Hospital Hsin-Chu Hospital, Taiwan | Research Center for Emerging Viral Infections, Chang Gung University, Taiwan | Pei-Lan Shao et al |
| **hCoV-19/Malaysia/MKAK_CL_22_28899/2022** | EPI_ISL_13517535 | | National Public Health Laboratory | National Public Health Laboratory | Yu Kie A/P Chem et al |
| **hCoV-19/Guangdong/GDCDC-20370/2022** | EPI_ISL_17494681 | | Guangdong Provincial Center for Disease Prevention and Control | Guangdong Provincial Center for Disease Prevention and Control | Liang et al |
| **hCoV-19/Philippines/PGCV_0047_5019/2022** | EPI_ISL_16995865 | | Western Visayas Medical Center Sub National Laboratory | PHILIPPINE GENOME CENTER VISAYAS | Dr. Cynthia P. Saloma et al |
| **hCoV-19/Japan/PG-453039/2023** | EPI_ISL_17058453 | | Tottori Prefectural Institute of Public Health and Environmental Science | Tottori Prefectural Institute of Public Health and Environmental Science | Kenkichi Hayashibara et al |
| **hCoV-19/MarshallIslands/KWA-H2320313/2023** | EPI_ISL_17182281 | | Ebeye Department of Health Services, Leiroj Hospital | State Laboratories Division, Hawaii State Department of Health | Pamela O'Brien et al |
| **hCoV-19/HongKong/HKU-230518-370/2023** | EPI_ISL_17685269 | | Department of Microbiology, The University of Hong Kong | Department of Microbiology, The University of Hong Kong | Kelvin K.W. To et al |
| **hCoV-19/HongKong/HK-HKPU-PUN534631323/2022** | EPI_ISL_17581967 | | Department of Health Technology and Informatics, The Hong Kong Polytechnic University | Department of Health Technology and Informatics, The Hong Kong Polytechnic University | Gilman Kit-Hang Siu et al |
| **hCoV-19/NorthernMarianaIslands/MP-CDC-2-6731089/2022** | EPI_ISL_16650487 | | Commonwealth Healthcare Center | Centers for Disease Control and Prevention Division of Viral Diseases, Pathogen Discovery | Mili Sheth et al |
| **hCoV-19/Singapore/R2NP2/2023** | EPI_ISL_16735319 | | National Public Health Laboratory, National Centre for Infectious Diseases | National Public Health Laboratory, National Centre for Infectious Diseases | Yichen Ding et al |
| **hCoV-19/Philippines/PGCV_0060_5362/2023** | EPI_ISL_18551728 | | Western Visayas Medical Center | PHILIPPINE GENOME CENTER VISAYAS | Dr. Cynthia P. Saloma et al |
| **hCoV-19/India/TR-INSACOG-634636413/2022** | EPI_ISL_10004951 | | INSACOG-Tripura | National Institute of Biomedical Genomics – INSACOG | Arindam Maitra et al A |
| **hCoV-19/Guangxi/IVDC-574/2022** | EPI_ISL_11873937 | | Guangxi Zhuang Autonomous Region Center for Disease Prevention and Control | National Institute for Viral Disease Control and Prevention, China CDC | Yu Ju et al |
| **hCoV-19/India/RJ-SMS-ICMR-INSACOG-TS-10141/2022** | EPI_ISL_10524207 | | SMS MEDICAL COLLEGE,JAIPUR | NIV Influenza | Bharti Malhotra et al |
| **hCoV-19/Canada/un-NML-378488/2022** | EPI_ISL_13288827 | | LifeLabs BC | National Microbiology Laboratory (NML) | Anna Majer et al |
| **hCoV-19/India/RJ-SMS-ICMR-INSACOG-TS-12026/2022** | EPI_ISL_10924197 | | SMS MEDICAL COLLEGE,JAIPUR | NIV Influenza | Bharti Malhotra et al |

| | | | | | |
|---|---|---|---|---|---|
| **hCoV-19/Myanmar/DSMRC-094/2022** | EPI_ISL_11149639 | ON038767.1 | DSMRC | Defence Services Medical Research Centre (DSMRC) | Khine Zaw Oo et al |
| **hCoV-19/Tianjin/TJCDC-0625/2022** | EPI_ISL_16923904 | | Tianjin Centers for Disease Control and Prevention | Tianjin Centers for Disease Control and Prevention | Li Xiaoyan et al |
| **hCoV-19/India/BR-INSACOG-GBRC7291/2022** | EPI_ISL_13697107 | | Pangenomics International Pvt. Ltd, Ahmedabad | Gujarat Biotechnology Research Centre | Sonal Sharma et al |
| **hCoV-19/India/WB-INSACOG-2032701965728/2022** | EPI_ISL_13536494 | | INSACOG-WB | National Institute of Biomedical Genomics – INSACOG | Arindam Maitra et al A |
| **hCoV-19/TrinidadandTobago/145526/2022** | EPI_ISL_13586159 | | Trinidad Public Health Laboratory | CARPHA | Carlos Rampersad on behalf of CARPHA et al |
| **hCoV-19/Maldives/MAV124802/2022** | EPI_ISL_9738920 | | Indira Gandhi Memorial Hospital | Indira Gandhi Memorial Hospital | D. Fathmath Nazla Rafeeq et al |
| **hCoV-19/India/MH-KasturbaMCGM-ICMR-INSACOG-WG3576/2022** | EPI_ISL_11664656 | | Kasturba Hospital Molecular Lab | NIV Influenza | Jayanthi Shastri et al |
| **hCoV-19/India/CT-ILSGS17473/2022** | EPI_ISL_14770239 | | GMC, Ambikapur | Institute of Life Sciences-INSACOG | Sunil K. Raghav et al |
| **hCoV-19/Iraq/BG-USAFSAM-S20132/2022** | EPI_ISL_15889755 | | US Air Force School of Aerospace Medicine | US Air Force School of Aerospace Medicine | Anthony Fries et al |
| **hCoV-19/Bhutan/RCDC0164/2022** | EPI_ISL_17967084 | | Royal Centre for Disease Control | Royal Centre for Disease Control | Sonam Gyeltshen et al |
| **hCoV-19/India/RJ-SMS-ICMR-INSACOG-TS-13879/2022** | EPI_ISL_13577808 | | SMS MEDICAL COLLEGE,JAIPUR | NIV Influenza | Bharti Malhotra et al |
| **hCoV-19/India/TG-CDFD-GH-13/2022** | EPI_ISL_14196415 | | Gandhi Medical College | CDFD-INSACOG | Asmita Gupta et al |
| **hCoV-19/India/GA-ICMR-NIV-INSACOG-GSEQ-11247/2022** | EPI_ISL_14149561 | | ICMR-National Institute of Virology - INSACOG | NIV Influenza | Dr. Varsha Potdar et al |
| **hCoV-19/India/UP-ICMR-INSACOG-KGMU-ZY55-068/2022** | EPI_ISL_12784401 | | Department of Microbiology,King George's Medical College | NIV Influenza | Prof. Amita Jain et al |
| **hCoV-19/India/RJ-SMS-ICMR-INSACOG-TS-14263/2022** | EPI_ISL_13576347 | | SMS MEDICAL COLLEGE,JAIPUR | NIV Influenza | Bharti Malhotra et al |
| **hCoV-19/India/GA-ICMR-NIV-INSACOG-GSEQ-11251/2022** | EPI_ISL_14149564 | | ICMR-National Institute of Virology - INSACOG | NIV Influenza | Dr. Varsha Potdar et al |
| **hCoV-19/India/DL-ILBSWGS6336/2022** | EPI_ISL_14290499 | | P BHASIN | ILBS/INSACOG | Ekta Gupta et al |
| **hCoV-19/India/SK-INSACOG-663798651/2022** | EPI_ISL_14303985 | | INSACOG-Sikkim | National Institute of Biomedical Genomics – INSACOG | Arindam Maitra et al A |
| **hCoV-19/India/RJ-SMS-ICMR-INSACOG-TS-15405/2022** | EPI_ISL_14356286 | | SMS MEDICAL COLLEGE,JAIPUR | NIV Influenza | Bharti Malhotra et al |
| **hCoV-19/India/GJ-INSACOG-GBRC6913/2022** | EPI_ISL_12751818 | | Supratech Micropath Diagnostics & Research laboratory Ahmedabad | Gujarat Biotechnology Research Centre | Zarna Patel et al |
| **hCoV-19/India/CG-AIIMSR-VRDL-INSACOG-0011/2022** | EPI_ISL_13728142 | | State Virus Research and Diagnostic Laboratory (VRDL), AIIMS Raipur | State Virus Research and Diagnostic Laboratory (VRDL), AIIMS Raipur | Pushpendra Singh et al |
| **hCoV-19/India/un-SPHL-605/2022** | EPI_ISL_17757788 | | Public Health, State Public Health Laboratory Chennai | Public Health, State Public Health Laboratory Chennai | Raju et al |
| **hCoV-19/India/DL-AIIMS-C-TN73076/2022** | EPI_ISL_13825875 | | COVID-CBNAAT | Genomics Lab; CCRF; AIIMS | Urvashi B. Singh et al |
| **hCoV-19/India/AS-NEIST-INSACOG-R38-CID2099/2022** | EPI_ISL_15332263 | | IDSP, Kamrup Metro (SRL) | Centre for Infectious Diseases, CSIR-NEIST | Rina Yumnam et al |
| **hCoV-19/India/MP-AIIMS_B-ICMR-INSACOG-WGS-1255/2022** | EPI_ISL_14842356 | | AIIMS Bhopal VRDL / Translational Medicine - INSACOG | AIIMS Bhopal VRDL/Translational Medicine - INSACOG | Anvita G. Malhotra et al |
| **hCoV-19/Bhutan/RCDC0165/2022** | EPI_ISL_17967085 | | Royal Centre for Disease Control | Royal Centre for Disease Control | Sonam Gyeltshen et al |
| **hCoV-19/India/MN-INSACOG-1425600254235/2022** | EPI_ISL_14303359 | | INSACOG-Manipur | National Institute of Biomedical Genomics – INSACOG | Arindam Maitra et al A |
| **hCoV-19/Guatemala/7692-LNS/2022** | EPI_ISL_15887327 | | Laboratorio Nacional de Salud (LNS), Guatemala | Laboratorio Nacional de Salud (LNS), Guatemala | Mendoza L et al |
| **hCoV-19/Guatemala/4716-LNS/2022** | EPI_ISL_13096601 | | Laboratorio Nacional de Salud (LNS), Guatemala | Laboratorio Nacional de Salud (LNS), Guatemala | Mendoza L et al |
| **hCoV-19/Guatemala/8455-LNS/2022** | EPI_ISL_14766843 | | Laboratorio Nacional de Salud (LNS), Guatemala | Laboratorio Nacional de Salud (LNS), Guatemala | Mendoza L et al |
| **hCoV-19/Mexico/TLA_InDRE_FB14866_E29217109312_S13634/2022** | EPI_ISL_13454120 | | LESP Tlaxcala | Instituto de Diagnostico y Referencia Epidemiologicos (INDRE) | Claudia Wong-Arambula et al |
| **hCoV-19/Philippines/PGCV_0016_0449/2022** | EPI_ISL_13131978 | | WESTERN VISAYAS MEDICAL CENTER SUB NATIONAL LABORATORY | PHILIPPINE GENOME CENTER VISAYAS | Dr. Cynthia P. Saloma et al |
| **hCoV-19/Guangdong/GPCDCP-2-12/2022** | EPI_ISL_16127644 | | Guangdong Provincial Center for Disease Control and Prevention | Key Laboratory of Human Diseases Comparative Medicine, Institute of Laboratory Animal Science, CAMS&PUMC | Bao et al |
| **hCoV-19/Belize/CML-580/2022** | EPI_ISL_14048092 | | Central Medical Laboratory | Central Medical Laboratory | Roberto Melendez et al |
| **hCoV-19/Mexico/BCS_IBT_IMSS_7829/2022** | EPI_ISL_13405498 | | Laboratorio Central de Epidemiología (LCE) | Instituto de Biotecnología de la UNAM | Consorcio Mexicano de Vigilancia Genómica (CoViGen-Mex). Authors (in alphabetical order): Julio Elias Alvarado-Yaah et al |
| **hCoV-19/Belize/CML-704/2022** | EPI_ISL_15397467 | | Central Medical Laboratory | Central Medical Laboratory | Roberto Melendez et al |
| **hCoV-19/Mexico/TLA_CIAD-IMSS_202201067059/2022** | EPI_ISL_13983270 | | Laboratorio Central de Epidemiologia (LCE) | Microbial Genomics Laboratory | Consorcio Mexicano de Vigilancia Genómica (CoViGen-Mex). Authors (in alphabetical order): Julio Elias Alvarado-Yaah et al |
| **hCoV-19/USA/MO-CDC-QDX38430420/2022** | EPI_ISL_13898309 | OP008405.1 | Quest Diagnostics Incorporated | Centers for Disease Control and Prevention Division of Viral Diseases, Pathogen Discovery | Dakota Howard et al |
| **hCoV-19/Curacao/CW-RIVM-103900/2022** | EPI_ISL_13760635 | | National Institute for Public Health and the Environment (RIVM) | National Institute for Public Health and the Environment (RIVM) | Adam Meijer et al |
| **hCoV-19/SintMaarten/SX-RIVM-104613/2022** | EPI_ISL_13854441 | | National Institute for Public Health and the Environment (RIVM) | National Institute for Public Health and the Environment (RIVM) | Adam Meijer et al |
| **hCoV-19/USA/KY-CDC-LC0749094/2022** | EPI_ISL_13807873 | ON976152.1 | Laboratory Corporation of America | Centers for Disease Control and Prevention Division of Viral Diseases, Pathogen Discovery | Dakota Howard et al |
| **hCoV-19/Venezuela/Car026/2022** | EPI_ISL_15975698 | | INHRR | Laboratorio de Virologia Molecular | Rossana C Jaspe et al |
| **hCoV-19/Singapore/3705/2022** | EPI_ISL_11982707 | | National Public Health Laboratory, National Centre for Infectious Diseases | National Public Health Laboratory, National Centre for Infectious Diseases | Yichen Ding et al |
| **hCoV-19/India/TN-CDFD-W2-142/2022** | EPI_ISL_10634026 | | Directorate Of Public Health And Preventive Medicine | INSACOG-Centre for DNA Fingerprinting and Diagnostics | Asmita Gupta et al |
| **hCoV-19/Kuwait/Jaber-20124569/2022** | EPI_ISL_11800551 | | Jaber Al Ahmad Al Sabah Hospital | Virology Lab, Jaber Al Ahmad Al Sabah Hospital | Dr. Zahrah Buhamad et al |
| **hCoV-19/India/DH-ICMR-NIV-INSACOG-G-11199/2022** | EPI_ISL_13989078 | | ICMR-National Institute of Virology - INSACOG | NIV Influenza | Dr. Varsha Potdar et al |
| **hCoV-19/India/GA-ICMR-12390/2022** | EPI_ISL_15074145 | | ICMR-National Institute of Virology - INSACOG | NIV Influenza | Dr. Varsha Potdar et al |
| **hCoV-19/India/MP-AIIMS_B-ICMR-INSACOG-WGS-1009/2022** | EPI_ISL_14842239 | | AIIMS Bhopal VRDL / Translational Medicine - INSACOG | AIIMS Bhopal VRDL/Translational Medicine - INSACOG | Anvita G. Malhotra et al |
| **hCoV-19/SouthKorea/KDCA63711/2022** | EPI_ISL_13137694 | | Division of Emerging Infectious Diseases, Bureau of Infectious Diseases Diagnosis Control, Korea Disease Control and Prevention Agency | Division of Emerging Infectious Diseases, Bureau of Infectious Diseases Diagnosis Control, Korea Disease Control and Prevention Agency | Jeong-Ah Kim et al |
| **hCoV-19/Tunisia/IPT-S-2869/2022** | EPI_ISL_17731628 | | Laboratoire de Virologie Clinique, Institut Pasteur de Tunis | Laboratoire de Virologie Clinique, Institut Pasteur de Tunis | Samar Ben Halima et al |
| **hCoV-19/Australia/NSW-ICPMR-24234/2022** | EPI_ISL_12405888 | | Pathology North - Hunter - NSW Health Pathology | NSW Health Pathology - Institute of Clinical Pathology and Medical Research; Westmead Hospital; University of Sydney | Arnott A. et al |
| **hCoV-19/India/GA-ICMR-NIV-INSACOG-GSEQ-11254/2022** | EPI_ISL_14149566 | | ICMR-National Institute of Virology - INSACOG | NIV Influenza | Dr. Varsha Potdar et al |
| **hCoV-19/Israel/SMC-7066375/2022** | EPI_ISL_12882740 | | Shamir Medical Center (Asaf Harofe) | Shamir Medical Center (Asaf Harofe) | Nir Rainy et al |
| **hCoV-19/India/PY-SLS-41195_S195_R1_001/2022** | EPI_ISL_15293919 | | JIPMER | inStem_NCBS_Strand Life Sciences (Supported by IN COVID support, FZE LLC) | Dasaradhi Palakodeti et al |
| **hCoV-19/India/JH-MCL-6757_11328/2022** | EPI_ISL_16477666 | | Indian Council of Medical Research-National Institute of Virology, Microbial Containment Complex | Indian Council of Medical Research-National Institute of Virology, Microbial Containment Complex | Pragya D. Yadav et al |
| **hCoV-19/India/MH-ICMR-NIV-INSACOG-G-11876/2022** | EPI_ISL_14405508 | | ICMR-National Institute of Virology - INSACOG | NIV Influenza | Dr. Varsha Potdar et al |
| **hCoV-19/India/BR-INSACOG-1930302083380/2022** | EPI_ISL_14890769 | | INSACOG-WB | National Institute of Biomedical Genomics – INSACOG | Arindam Maitra et al A |
| **hCoV-19/India/WB-INSACOG-1931503215316/2022** | EPI_ISL_13877198 | | INSACOG-WB | National Institute of Biomedical Genomics – INSACOG | Arindam Maitra et al A |
| **hCoV-19/India/GA-ICMR-NIV-INSACOG-G-12163/2022** | EPI_ISL_14861606 | | ICMR-National Institute of Virology - INSACOG | NIV Influenza | Dr. Varsha Potdar et al |
| **hCoV-19/India/MN-INSACOG-1425500178674/2022** | EPI_ISL_14303418 | | INSACOG-Manipur | National Institute of Biomedical Genomics – INSACOG | Arindam Maitra et al A |
| **hCoV-19/India/TG-CDFD-GH-193/2022** | EPI_ISL_14196550 | | Gandhi Medical College | CDFD-INSACOG | Asmita Gupta et al |
| **hCoV-19/USA/CO-CDC-MMB14878197/2022** | EPI_ISL_12297960 | ON345929.1 | Mako Medical | Centers for Disease Control and Prevention Division of Viral Diseases, Pathogen Discovery | Dakota Howard et al |
| **hCoV-19/Panama/1352-GMI/2022** | EPI_ISL_15224210 | | Gorgas Memorial Institute of Health Studies | Gorgas Memorial Institute of Health Studies | Claudia Gonzalez et al |
| **hCoV-19/CaboVerde/ST1283/2022** | EPI_ISL_14076726 | | Virology Laboratory of Praia | Instituto Nacional de Saúde Pública (INSP) | Kevin Sanders et al |
| **hCoV-19/SriLanka/aicbu4176/2022** | EPI_ISL_9285951 | | Allergy, Immunology and Cell Biology Unit (AICBU) | Allergy, Immunology and Cell Biology Unit (AICBU) | Chandima Jeewandara et al |
| **hCoV-19/Guangdong/IVDC-598/2022** | EPI_ISL_12030363 | | Guangdong Provincial Center for Disease Control and Prevention | National Institute for Viral Disease Control and Prevention, China CDC | Baisheng Li et al |
| **hCoV-19/Hunan/IVDC-555/2022** | EPI_ISL_11873918 | | Hunan Provincial Center for Disease Control and Prevention | National Institute for Viral Disease Control and Prevention, China CDC | Xingyu Xiang et al |
| **hCoV-19/Shanghai/IVDC-611/2022** | EPI_ISL_12461212 | | Shanghai Municipal Center for Disease Control & Prevention | National Institute for Viral Disease Control and Prevention, China CDC | Hui Jiang et al |
| **hCoV-19/Guangdong/IVDC-599/2022** | EPI_ISL_12030364 | | Guangdong Provincial Center for Disease Control and Prevention | National Institute for Viral Disease Control and Prevention, China CDC | Baisheng Li et al |
| **hCoV-19/Shaanxi/IVDC-560/2022** | EPI_ISL_11873923 | | Shaanxi Provincial Center for Disease Control and Prevention | National Institute for Viral Disease Control and Prevention, China CDC | Shen Li et al |
| **hCoV-19/Jiangsu/IVDC-580/2022** | EPI_ISL_11905845 | | Jiangsu Provincial Center for Disease Control and Prevention | National Institute for Viral Disease Control and Prevention, China CDC | Shenjiao Wang et al |
| **hCoV-19/Shanghai/IVDC-616/2022** | EPI_ISL_12461217 | | Shanghai Municipal Center for Disease Control & Prevention | National Institute for Viral Disease Control and Prevention, China CDC | Hui Jiang et al |
| **hCoV-19/Tianjin/TJCDC-0682/2022** | EPI_ISL_16923939 | | Tianjin Centers for Disease Control and Prevention | Tianjin Centers for Disease Control and Prevention | Li Xiaoyan et al |
| **hCoV-19/Anhui/CDC-3341/2022** | EPI_ISL_17167874 | | Lu`an Municipal Center for Disease Control and Prevention - Microbiological Laboratory | Lu`an Municipal Center for Disease Control and Prevention - Microbiological Laboratory | Yang et al |
| **hCoV-19/Jiangxi/JXCDC-18/2022** | EPI_ISL_12241081 | | Jiaxing Center for Disease Control and Prevention | Jiaxing Center for Disease Control and Prevention | Zhongwen Chen et al |
| **hCoV-19/Xinjiang/IVDC-576/2022** | EPI_ISL_11873939 | | The Center for Disease Control and Prevention of Xinjiang Uygur Autonomous Region | National Institute for Viral Disease Control and Prevention, China CDC | Xin Ma et al |
| **hCoV-19/Jiangxi/IVDC-557/2022** | EPI_ISL_11873920 | | Jiangxi Provincial Center for Disease Control and Prevention | National Institute for Viral Disease Control and Prevention, China CDC | Tian Gong et al |
| **hCoV-19/Anhui/WHCDC-008/2022** | EPI_ISL_12901920 | | Wuhu Center for Disease Control and Prevention | Wuhu Center for Disease Control and Prevention | Gongfei Wang et al |
| **hCoV-19/Anhui/WHCDC-004/2022** | EPI_ISL_12901916 | | Wuhu Center for Disease Control and Prevention | Wuhu Center for Disease Control and Prevention | Gongfei Wang et al |
| **hCoV-19/Hainan/IVDC-608/2022** | EPI_ISL_12030373 | | Hainan Provincial Center for Disease Control and Prevention | National Institute for Viral Disease Control and Prevention, China CDC | Yinggui Cai et al |
| **hCoV-19/Zhejiang/NBCDC_20220315_HuYibo/2022** | EPI_ISL_16137942 | | Virus research department, Ningbo Municipal Center for Disease Control and Prevention | Virus research department, Ningbo Municipal Center for Disease Control and Prevention | Mao et al |
| **hCoV-19/Yunnan/IVDC-568/2022** | EPI_ISL_11873931 | | Department of Acute Infectious Diseases Control and Prevention, Yunnan Center for Disease Control and Prevention | National Institute for Viral Disease Control and Prevention, China CDC | Meiling Zhang et al |
| **hCoV-19/Sichuan/IVDC-LS-587/2022** | EPI_ISL_11905852 | | Sichuan Center of Disease Control and Prevention | National Institute for Viral Disease Control and Prevention, China CDC | Teng Qi et al |
| **hCoV-19/Beijing/BJCDC-1641/2022** | EPI_ISL_16633951 | | Beijing Center for Disease Control and Prevention | Beijing Center for Disease Control and Prevention | Fu Li et al |
| **hCoV-19/Beijing/BJCDC-1233/2022** | EPI_ISL_16614304 | | Beijing Center for Disease Control and Prevention | Beijing Center for Disease Control and Prevention | Fu Li et al |
| **hCoV-19/Shanghai/2-43-s28/2022** | EPI_ISL_16137916 | | Microbiology, The Second Military Medical University | Microbiology, The Second Military Medical University | Tang et al |
| **hCoV-19/Beijing/BJCDC-659/2022** | EPI_ISL_16557220 | | Beijing Center for Disease Control and Prevention | Beijing Center for Disease Control and Prevention | Zhaomin Feng et al |
| **hCoV-19/Romania/IS-20010/2022** | EPI_ISL_15488908 | | Department of Public Health Iasi | National Institute of Infectious Diseases-Prof. Dr. Matei Bals Molecular Diagnostics Laboratory | Corina Casangiu et al |
| **hCoV-19/Turkey/HSGM-IST-01862/2022** | EPI_ISL_13113257 | | Ministry of Health Turkey | Ministry of Health Turkey | Gülay Korukluoğlu et al |
| **hCoV-19/India/TG-CDFD-ncov-21310/2022** | EPI_ISL_11021949 | | Institute of Preventive Medicine | CDFD-INSACOG | Asmita Gupta et al |
| **hCoV-19/India/TN-CDFD-W5-289/2022** | EPI_ISL_11930866 | | Directorate of Public Health and Preventive Medicine | CDFD-INSACOG | Asmita Gupta et al |
| **hCoV-19/India/TG-CDFD-MMG_435/2022** | EPI_ISL_11931174 | | Centre for DNA Fingerprinting and Diagnostics | CDFD-INSACOG | Asmita Gupta et al |
| **hCoV-19/Guizhou/IVDC-578/2022** | EPI_ISL_11905843 | | Guizhou Provincial Center for Disease Control and Prevention | National Institute for Viral Disease Control and Prevention, China CDC | Yonghu Wan et al |
| **hCoV-19/SouthAfrica/NHLS-UCT-LA-Z962/2022** | EPI_ISL_12221559 | ON324380.1 | Lancet | NHLS/UCT | Arash Iranzadeh et al |
| **hCoV-19/HongKong/HK-HKPU-PUU037208984/2022** | EPI_ISL_13140464 | | Department of Health Technology and Informatics, The Hong Kong Polytechnic University | Department of Health Technology and Informatics, The Hong Kong Polytechnic University | Gilman Kit-Hang Siu et al |
| **hCoV-19/India/RJ-SMS-ICMR-INSACOG-TS-12573/2022** | EPI_ISL_11881633 | | SMS MEDICAL COLLEGE,JAIPUR | NIV Influenza | Bharti Malhotra et al |
| **hCoV-19/Brazil/PR-IMTSP-CD32021/2022** | EPI_ISL_16960869 | | DB Diagnosticos do Brasil | Instituto de Medicina Tropical de Sao Paulo | Ester Cerdeira Sabino et al |
| **hCoV-19/Jilin/IVDC-556/2022** | EPI_ISL_11873919 | | Jilin Province Center for Disease Control and Prevention | National Institute for Viral Disease Control and Prevention, China CDC | DonglinWu et al |
| **hCoV-19/Russia/MOW-CRIE-7735371896/2022** | EPI_ISL_12955897 | | Molecular diagnostic laboratory of Federal Budget Institution of Science "Central Research Institute of Epidemiology" of The Federal Service on Customers' Rights Protection and Human Well-being Surveillance | Group of Genomics and Postgenomic Technologies of Central Research Institute of Epidemiology | Kaptelova V.V. et al |
| **hCoV-19/SriLanka/aicbu5379/2022** | EPI_ISL_11438061 | | Allergy, Immunology and Cell Biology Unit (AICBU) | Allergy, Immunology and Cell Biology Unit (AICBU) | Chandima Jeewandara et al |
| **hCoV-19/Curacao/CW-RIVM-95627/2022** | EPI_ISL_11840187 | | National Institute for Public Health and the Environment (RIVM) | National Institute for Public Health and the Environment (RIVM) | Adam Meijer et al |
| **hCoV-19/Morocco/724/2022** | EPI_ISL_13289800 | | NIH-Rabat | NIC at National Institute of Hygiene | Hassan Ihazmade et al |
| **hCoV-19/Oman/7228735/2022** | EPI_ISL_14241487 | | Oman-NIC | Oman-NIC | Samiha Al Kharusi et al |
| **hCoV-19/Henan/IVDC-548/2022** | EPI_ISL_11873911 | | Henan Center for Disease Control and Prevention | National Institute for Viral Disease Control and Prevention, China CDC | Hongxia Ma et al |
| **hCoV-19/Malaysia/IMR_OS927/2022** | EPI_ISL_12983219 | | BP Healthcare Group | Institute for Medical Research, Infectious Disease Research Centre, National Institutes of Health, Ministry of Health Malaysia | Kalyanasundram J et al |
| **hCoV-19/Bonaire/BQ-RIVM-95657/2022** | EPI_ISL_11840165 | | National Institute for Public Health and the Environment (RIVM) | National Institute for Public Health and the Environment (RIVM) | Adam Meijer et al |
| **hCoV-19/Bonaire/BQ-RIVM-101066/2022** | EPI_ISL_12953992 | | National Institute for Public Health and the Environment (RIVM) | National Institute for Public Health and the Environment (RIVM) | Adam Meijer et al |
| **hCoV-19/SriLanka/aicbu5037/2022** | EPI_ISL_10994029 | | Allergy, Immunology and Cell Biology Unit (AICBU) | Allergy, Immunology and Cell Biology Unit (AICBU) | Chandima Jeewandara et al |
| **hCoV-19/India/MH-ICMR-NIV-INSACOG-GSEQ-10026/2022** | EPI_ISL_10801565 | | ICMR-National Institute of Virology - INSACOG | NIV Influenza | Dr. Varsha Potdar et al |
| **hCoV-19/Jiangsu/NJ_07/2022** | EPI_ISL_16138060 | | Center Laboratory of Health Quarantine, Nanjing Customs | Center Laboratory of Health Quarantine, Nanjing Customs | Geng et al |
| **hCoV-19/SaudiArabia/KAU_CEBR_C56/2022** | EPI_ISL_15668770 | | Alborg Diagnostics | Centre of Excellence in Bionanoscience Research | Ahmed Bouna et al |
| **hCoV-19/Pakistan/NIH-B97-S17/2022** | EPI_ISL_15112027 | | Department of Virology | Department of Virology | Massab Umair et al |
| **hCoV-19/Poland/WSSEGDA-1347/2022** | EPI_ISL_12055081 | | Diagnostyka Sp. z o.o. Gdańsk | Wojewódzka Stacja Sanitarno-Epidemiologiczna w Gdańsku | Gabriela Rutkowska et al |
| **hCoV-19/CaboVerde/ST5891/2022** | EPI_ISL_13525887 | | Virology Laboratory of Praia | Instituto Nacional de Saúde Pública | Kevin Sanders et al |
| **hCoV-19/Uruguay/DLSP_2432/2022** | EPI_ISL_16385382 | | Departamento Laboratorios de Salud Pública (DLSP) Ministerio de Salud Pública | Departamento Laboratorios de Salud Pública (DLSP) Ministerio de Salud Pública | Cortinas et al |
| **hCoV-19/Pakistan/NIH-B38-S7/2022** | EPI_ISL_11114542 | | Department of Virology | Department of Virology | Massab Umair et al |
| **hCoV-19/Malaysia/MKAK_22_69815/2022** | EPI_ISL_12099754 | | National Public Health Laboratory | National Public Health Laboratory | Yu Kie A/P Chem et al |
| **hCoV-19/Mongolia/GS-33884/2022** | EPI_ISL_14807746 | | National Center for Communicable Diseases (NCCD) National Influenza Center | National Center for Communicable Diseases (NCCD) National Influenza Center | Bayasgalan Namuutsetseg et al |
| **hCoV-19/Mongolia/UB-602338/2022** | EPI_ISL_14713975 | | National Center for Communicable Diseases (NCCD) National Influenza Center | National Center for Communicable Diseases (NCCD) National Influenza Center | Naranzul Tsedenbal et al |
| **hCoV-19/Cyprus/BMV6540/2022** | EPI_ISL_18479062 | | Limassol General Hospital | Laboratory of Biotechnology and Molecular Virology, University of Cyprus | Leondios G. Kostrikis et al. |
| **hCoV-19/Netherlands/OV-DR-GD-03507031/2022** | EPI_ISL_12293900 | | Royal GD | Royal GD | Remco Dijkman et al |

| | | | | | |
|---|---|---|---|---|---|
| **hCoV-19/India/KL-DL-AIIMS-COVTN71885/2022** | EPI_ISL_12610369 | | COVID-CBNAAT; Department of Microbiology; AIIMS; New Delhi | Genomics Lab; CCRF; AIIMS; New Delhi_INSACOG | Urvashi B. Singh et al |
| **hCoV-19/NewZealand/22CV4987/2022** | EPI_ISL_13437569 | | Wellington SCL (WN) | Institute of Environmental Science and Research (ESR) | Sarah Cockerton et al |
| **hCoV-19/Shandong/WCDC-028/2022** | EPI_ISL_18070360 | | Weihai Center for Disease Control and Prevention | Weihai Center for Disease Control and Prevention | Xiang Li et al |
| **hCoV-19/USA/FL-CDC-LC0571786/2022** | EPI_ISL_12167967 | ON295315.1 | Laboratory Corporation of America | Centers for Disease Control and Prevention Division of Viral Diseases, Pathogen Discovery | Dakota Howard et al |
| **hCoV-19/Chongqing/YBCDC-0110/2022** | EPI_ISL_16613438 | | Chongqing Yubei District Center for Disease Control and Prevention | Chongqing Yubei District Center for Disease Control and Prevention | Jin Yan et al |
| **hCoV-19/Georgia/Tb-s5219/2022** | EPI_ISL_13480538 | | National Center for Disease Control and Public Health (NCDC) of Georgia. | Department for Virology, Molecular Biology and Genome Research, R. G. Lugar Center for Public Health Research, National Center for Disease Control and Public Health (NCDC) of Georgia. | Giorgi Tomashvili et al |
| **hCoV-19/USA/OR-CDC-LC0608440/2022** | EPI_ISL_12656502 | ON499953.1 | Laboratory Corporation of America | Centers for Disease Control and Prevention Division of Viral Diseases, Pathogen Discovery | Dakota Howard et al |
| **hCoV-19/Canada/PE-NML-434029/2022** | EPI_ISL_15722442 | | PE-QEH Dept of Lab Medicine | National Microbiology Laboratory (NML) | Vanessa Arseneau et al |
| **hCoV-19/Bonaire/BQ-RIVM-100087/2022** | EPI_ISL_12782645 | | National Institute for Public Health and the Environment (RIVM) | National Institute for Public Health and the Environment (RIVM) | Adam Meijer et al |
| **hCoV-19/SaudiArabia/KAUST260/2022** | EPI_ISL_15827045 | | King Faisal Specialist Hospital & Research Center | Pathogen Genomics Lab King Abdullah University of Science and Technology (KAUST) | Sabir A Adoub et al |
| **hCoV-19/Montenegro/CO-00892_MNE000_1927902202/2022** | EPI_ISL_14491756 | | Laboratory for Molecular Diagnostics,IPHMN | Eurofins Genomics Europe Sequencing GmbH | Danijela Vujošević et al |
| **hCoV-19/Montenegro/CO-00892_MNE000_1920702202/2022** | EPI_ISL_14491127 | | Laboratory for Molecular Diagnostics,IPHMN | Eurofins Genomics Europe Sequencing GmbH | Danijela Vujošević et al |
| **hCoV-19/India/MZ-INSACOG-1526100691460/2022** | EPI_ISL_13876770 | | INSACOG-Mizoram | National Institute of Biomedical Genomics – INSACOG | Arindam Maitra et al A |
| **hCoV-19/Lithuania/LTU000_VULSK_220501120173/2022** | EPI_ISL_12771681 | | Viesoji istaiga Vilniaus universiteto ligonine Santaros klinikos | National Public Health Surveillance Laboratory | Lukas Zemaitis et al |
| **hCoV-19/Aruba/AW-RIVM-101154/2022** | EPI_ISL_12954080 | | National Institute for Public Health and the Environment (RIVM) | National Institute for Public Health and the Environment (RIVM) | Adam Meijer et al |
| **hCoV-19/Aruba/AW-RIVM-126320/2022** | EPI_ISL_16963548 | | National Institute for Public Health and the Environment (RIVM) | National Institute for Public Health and the Environment (RIVM) | Adam Meijer et al |
| **hCoV-19/Curacao/CW-RIVM-101113/2022** | EPI_ISL_12954039 | | National Institute for Public Health and the Environment (RIVM) | National Institute for Public Health and the Environment (RIVM) | Adam Meijer et al |
| **hCoV-19/Curacao/CW-RIVM-101615/2022** | EPI_ISL_13144589 | | National Institute for Public Health and the Environment (RIVM) | National Institute for Public Health and the Environment (RIVM) | Adam Meijer et al |
| **hCoV-19/Australia/TAS001798/2022** | EPI_ISL_12386001 | | Royal Hobart Hospital | Royal Hobart Hospital | J Williamson et al |
| **hCoV-19/BosniaandHerzegovina/AGC3217/2022** | EPI_ISL_13804565 | | Alea Genetic Centre | Alea Genetic Centre | Pecar D et al |
| **hCoV-19/USA/DC-CDC-LC0629508/2022** | EPI_ISL_12889383 | ON592723.1 | Laboratory Corporation of America | Centers for Disease Control and Prevention Division of Viral Diseases, Pathogen Discovery | Dakota Howard et al |
| **hCoV-19/Pakistan/NIH-B49-GOV-S18/2022** | EPI_ISL_12902394 | | Department of Virology, Armed Forces Institute of Pathology (AFIP), Rawalpindi, Pakistan | Department of Virology, National Institute of Health, Islamabad, Pakistan | Massab Umair et al |
| **hCoV-19/Laos/LOMWRU-0578/2022** | EPI_ISL_15973666 | | LOMWRU | LOMWRU/Microbiology Laboratory, Mahosot Hospital | Manila Souksavanh et al |
| **hCoV-19/Bulgaria/22BG-EU_020858_PI197/2022** | EPI_ISL_14889505 | | National Center of Infectious and Parasitic Diseases | National Center of Infectious and Parasitic Diseases | Alexiev et al |
| **hCoV-19/India/MZ-INSACOG-1526101089213/2022** | EPI_ISL_13159398 | | INSACOG-Mizoram | National Institute of Biomedical Genomics – INSACOG | Arindam Maitra et al A |
| **hCoV-19/Russia/SPE-RII-21723S/2022** | EPI_ISL_13145393 | | WHO National Influenza Centre Russian Federation | WHO National Influenza Centre Russian Federation | Andrey Komissarov et al |
| **hCoV-19/Uruguay/DLSP_2156/2022** | EPI_ISL_16385391 | | Departamento Laboratorios de Salud Pública (DLSP) Ministerio de Salud Pública | Departamento Laboratorios de Salud Pública (DLSP) Ministerio de Salud Pública | Cortinas et al |
| **hCoV-19/Kazakhstan/516/2022** | EPI_ISL_13822834 | | National Center of Expertise in Nur-Sultan city | National center of Expertise, National center for Biotechnology | Shevtsov Alexandr et al |
| **hCoV-19/Tianjin/TJCDC-0755/2022** | EPI_ISL_16923520 | | Tianjin Centers for Disease Control and Prevention | Tianjin Centers for Disease Control and Prevention | Li Xiaoyan et al |
| **hCoV-19/Ireland/D-Enfer-COV150222027_E4/2022** | EPI_ISL_10585101 | | Enfer | Enfer | Elaine M. Kenny et al |
| **hCoV-19/Montenegro/CO-00688_MNE000_1633892202/2022** | EPI_ISL_12280269 | | Laboratory for Molecular Diagnostics,IPHMN | Eurofins Genomics Europe Sequencing GmbH | Danijela Vujošević et al |
| **hCoV-19/Beijing/BJCDC-444/2022** | EPI_ISL_13858991 | | Beijing Center for Disease Control and Prevention | Beijing Center for Disease Control and Prevention | Shujuan Cui et al |
| **hCoV-19/Maldives/MAV172364/2022** | EPI_ISL_10994841 | | Kulhudufushi Regional Hospital Lab | Kulhudufushi Regional Hospital Lab | D. Fathmath Nazla Rafeeq et al |
| **hCoV-19/India/TN-CDFD-J-1097/2022** | EPI_ISL_13668094 | | Directorate of Public Health and Preventive Medicine | CDFD-INSACOG | Asmita Gupta et al |
| **hCoV-19/Barbados/142989/2022** | EPI_ISL_13446655 | | Best-dos Santos Public Health Lab | CARPHA | Carlos Rampersad on behalf of CARPHA et al |
| **hCoV-19/Singapore/6627/2022** | EPI_ISL_13360588 | | National Public Health Laboratory, National Centre for Infectious Diseases | National Public Health Laboratory, National Centre for Infectious Diseases | Yichen Ding et al |
| **hCoV-19/Singapore/8923/2022** | EPI_ISL_14101297 | | National Public Health Laboratory, National Centre for Infectious Diseases | National Public Health Laboratory, National Centre for Infectious Diseases | Benny Yeo et al |
| **hCoV-19/Georgia/Tb-s5326/2022** | EPI_ISL_13884667 | | Humanity Lab | Department for Virology, Molecular Biology and Genome Research, R. G. Lugar Center for Public Health Research, National Center for Disease Control and Public Health (NCDC) of Georgia. | Giorgi Tomashvili et al |
| **hCoV-19/Bangladesh/icddrb-GSB14-268/2022** | EPI_ISL_13408130 | | Shaheed Ziaur Rahman Medical College, Bogura | Virology Laboratory, International Centre for Diarrhoeal Disease Research, Bangladesh (ICDDR,B) | Mohammad Jubair et al |
| **hCoV-19/India/UP-ICMR-INSACOG-KGMU-ZY32/523/2022** | EPI_ISL_12492491 | | Department of Microbiology,King George's Medical College | NIV Influenza | Prof. Amita Jain et al |
| **hCoV-19/India/AS-NEIST-INSACOG-NEIST29-CID0791/2022** | EPI_ISL_13354412 | | IDSP, Kamrup Metro (VRDL, GMCH) | Centre for Infectious Diseases, CSIR-NEIST | Romi Wahengbam et al |
| **hCoV-19/India/PB-VRDLGMCPTA-ICMR-INSACOG-729517643/2022** | EPI_ISL_12784924 | | Virus Research and Diagnostic Laboratory | NIV Influenza | Dr.Rupinder Bakshi et al |
| **hCoV-19/India/AS-NEIST-INSACOG-NEIST28-CID0786/2022** | EPI_ISL_13354407 | | IDSP, Kamrup Metro (VRDL, GMCH) | Centre for Infectious Diseases, CSIR-NEIST | Romi Wahengbam et al |
| **hCoV-19/India/SK-INSACOG-1122500131944/2022** | EPI_ISL_14303999 | | INSACOG-Sikkim | National Institute of Biomedical Genomics – INSACOG | Arindam Maitra et al A |
| **hCoV-19/India/LA-INSACOG-NII-739/2022** | EPI_ISL_13409363 | | COVID-19 Laboratory Chuchot SNM Hospital | National Institute of Immunology | Dr. G. Aneeshkumar Arimbasseri et al |
| **hCoV-19/India/MP-AIIMS_B-ICMR-INSACOG-WGS-906/2022** | EPI_ISL_13574755 | | AIIMS Bhopal VRDL/Translational Medicine - INSACOG | NIV Influenza | Anvita G. Malhotra et al |
| **hCoV-19/Malaysia/MGVI_GS1235/2022** | EPI_ISL_13329633 | | Hospital Ampang | Malaysia Genome and Vaccine Institute | Mohd Noor Mat Isa et al |
| **hCoV-19/Norway/14268/2022** | EPI_ISL_11656386 | | University Hospital of Northern Norway, Department for Microbiology and Infectious Disease Control | Norwegian Institute of Public Health, Department of Virology | Kathrine Stene-Johansen et al |
| **hCoV-19/DominicanRepublic/1858328-LNSPDD/2022** | EPI_ISL_10398029 | | Laboratorio Nacional de Salud Pública Dr. Defilló | Laboratorio Nacional de Salud Pública Dr. Defilló | Isaac Miguel Sánchez et al |
| **hCoV-19/USA/WA-PHL-020816/2022** | EPI_ISL_12969423 | | Northwest Laboratory | Washington State Department of Health Public Health Laboratories | Drew MacKellar et al |
| **hCoV-19/Albania/UZB_548202204116/2022** | EPI_ISL_12086771 | | Microbiology Department - University Hospital Brussel | Microbiology Department - University Hospital Brussel | Thomas Demuyser et al |
| **hCoV-19/NorthMacedonia/IPH-MKD-5598/2022** | EPI_ISL_13032665 | | Laboratory of Virology - Institute of Public Health Skopje, Macedonia | Laboratory of Virology - Institute of Public Health Skopje, Macedonia | Teodora Buzharova et al |
| **hCoV-19/Norway/OUS-23060/2022** | EPI_ISL_11811047 | | Oslo University Hospital, Department of Microbiology | Norwegian Institute of Public Health, Department of Virology | Mona Holberg-Petersen et al |
| **hCoV-19/Switzerland/VS-ETHZ-37155304/2022** | EPI_ISL_12339528 | OW577271.1 | Viollier AG | Department of Biosystems Science and Engineering, ETH Zürich | Andrea Patrizia Salzmann et al |
| **hCoV-19/Austria/LB-R00083-S122/2022** | EPI_ISL_13445714 | | Lifebrain Covid Labor GmbH | Lifebrain Covid Labor GmbH | Filip Sima et al |
| **hCoV-19/Yunnan/Dali02/2022** | EPI_ISL_14770175 | | The Centers for Disease Control and Prevention in Dali Bai Autonomous Prefecture, Yunnan province | Department of Acute Infectious Diseases Control and Prevention, Yunnan Provincial Center for Disease Control and Prevention | Duan feiyun et al |
| **hCoV-19/Yunnan/Dali01/2022** | EPI_ISL_14770174 | | The Centers for Disease Control and Prevention in Dali Bai Autonomous Prefecture, Yunnan province | Yunnan Provincial Center for Disease Control and Prevention | Duan feiyun et al |
| **hCoV-19/Netherlands/ZH-NMDL-01336/2022** | EPI_ISL_11244313 | | Eurofins-NMDL | Eurofins-NMDL | Anne Vogel et al |
| **hCoV-19/Germany/HH-RKI-I-746917/2022** | EPI_ISL_12357188 | | MVZ Labor Dr. Fenner und Kollegen (Standort Hamburg) | Robert Koch Institute | ? |
| **hCoV-19/Scotland/QEUH-3DFD756/2022** | EPI_ISL_12647656 | OW961449.1 | Lighthouse Lab in Glasgow | Wellcome Sanger Institute for the COVID-19 Genomics UK (COG-UK) Consortium | Harper VanSteenhouse et al |
| **hCoV-19/Denmark/DCGC-509553/2022** | EPI_ISL_12609335 | OX192833.1 | Department of Bacteria, Parasites and Fungi, Statens Serum Institut, Copenhagen, Denmark | Statens Serum Institut Bioinformatics and Microbial Genomics | Povilas Matusevicius on behalf of the Danish Covid-19 Genome Consortium et al |
| **hCoV-19/CzechRepublic/NAT-22-26810/2022** | EPI_ISL_12804235 | | Faculty Hospital Bulovka, Department of Clinical Microbiology | Charles University, Faculty of Science, BIOCEV, OMICS Genomics | Vladimír Hampl et al |
| **hCoV-19/Brazil/SP-IB_156955/2022** | EPI_ISL_13020883 | | HOSPITAL FORNECEDORES DE CANA DE PIRACICABA | Instituto Butantan | Dimas Tadeu Covas et al |
| **hCoV-19/Lithuania/LSMULKKGMMK44C23/2022** | EPI_ISL_12171015 | | UAB InMedica | Hospital of Lithuanian University of Health Sciences (LSMU) Kaunas Clinics | Rasa Ugenskiene et al |
| **hCoV-19/Ecuador/USFQ-3327/2022** | EPI_ISL_12590933 | | Institute of Microbiology, Universidad San Francisco de Quito | Institute of Microbiology, Universidad San Francisco de Quito | Belén Prado-Vivar et al |
| **hCoV-19/USA/ID-IBL-828080/2022** | EPI_ISL_13453636 | | Idaho Bureau of Laboratories | Idaho Bureau of Laboratories | "R. Beukelman et al |
| **hCoV-19/Ukraine/70868/2022** | EPI_ISL_14051628 | | Reference laboratory for diagnostics of HIV/AIDS, virological and especially dangerous pathogens | Reference laboratory for diagnostics of HIV/AIDS, virological and especially dangerous pathogens | Iryna Demchyshyna et al |
| **hCoV-19/CanaryIslands/HUGCDN-02682538/2022** | EPI_ISL_12144238 | | Servicio Microbiología H.U. Dr. Negrín | Servicio Microbiología H.U. Dr. Negrín | Francisco Javier Chamizo López Ana Bordes Benítez et al |
| **hCoV-19/Aruba/AW-RIVM-101173/2022** | EPI_ISL_12954099 | | National Institute for Public Health and the Environment (RIVM) | National Institute for Public Health and the Environment (RIVM) | Adam Meijer et al |
| **hCoV-19/Belgium/UZA-UA-CV8366479362/2022** | EPI_ISL_12683795 | | Platform BIS UZA/UAntwerpen | Labo Klinische Biologie, UZA | Jasmine Coppens et al |
| **hCoV-19/Sweden/2A68213430099/2022** | EPI_ISL_13184481 | | The Public Health Agency of Sweden | The Public Health Agency of Sweden | Emmi Andersson et al |
| **hCoV-19/CzechRepublic/UMTM484904/2022** | EPI_ISL_13756487 | | Institute of Molecular and Translational Medicine / Laboratory of Experimental Medicine, Faculty of Medicine and Dentistry, Palacky University | Institute of Molecular and Translational Medicine / Laboratory of Experimental Medicine, Faculty of Medicine and Dentistry, Palacky University | Rastislav Slavkovský et al |
| **hCoV-19/CzechRepublic/33SVAN43392/2022** | EPI_ISL_12919185 | | Covid laboratory, Department of Microbiology, St. Anne's University Hospital Brno | Covid laboratory, Department of Microbiology, St. Anne's University Hospital Brno | Milada Dvořáčková et al |
| **hCoV-19/Moldova/G0014/2022** | EPI_ISL_13159676 | | MedExpert | MedExpert | Vadim Nirca et al |
| **hCoV-19/Panama/CDED23130-GMI/2022** | EPI_ISL_16027402 | | Centro de Diagnostico de Enfermedades Desatendidas | Gorgas Memorial Institute of Health Studies | Eduardo Aguirre et al |
| **hCoV-19/Colombia/CUN-IAVH-VG-18750/2022** | EPI_ISL_12666254 | | Colsanitas Keralty | Instituto de Investigación de Recursos Biológicos Alexander von Humboldt | Nicolás D. Franco-Sierra et al |
| **hCoV-19/Mexico/CHP_InDRE_FB14070_E07316786946_S13225/2022** | EPI_ISL_12739336 | | LESP Chiapas | Instituto de Diagnostico y Referencia Epidemiologicos (INDRE) | Claudia Wong-Arambula et al |
| **hCoV-19/Mexico/YUC_InDRE_FB14297_E31316907901_S13418/2022** | EPI_ISL_13192155 | | LESP Yucatan | Instituto de Diagnostico y Referencia Epidemiologicos (INDRE) | Claudia Wong-Arambula et al |
| **hCoV-19/TrinidadandTobago/S22-20506/2022** | EPI_ISL_11998950 | | Trinidad Public Health Laboratory | Carrington Lab, Department of Preclinical Sciences, Faculty of Medical Sciences, The University of the West Indies, St Augustine Campus | Nikita S. D. Sahadeo et al |
| **hCoV-19/Mexico/TAM_InDRE_FB14655_E28117036809_S13587/2022** | EPI_ISL_13454073 | | LESP Tamaulipas | Instituto de Diagnostico y Referencia Epidemiologicos (INDRE) | Claudia Wong-Arambula et al |
| **hCoV-19/Mexico/TLA_InDRE_FB14057_E29316747590_S13228/2022** | EPI_ISL_12739379 | | LESP Tlaxcala | Instituto de Diagnostico y Referencia Epidemiologicos (INDRE) | Claudia Wong-Arambula et al |
| **hCoV-19/Norway/OUS-21041/2022** | EPI_ISL_10861841 | | Oslo University Hospital, Department of Microbiology | Norwegian Institute of Public Health, Department of Virology | Mona Holberg-Petersen et al |
| **hCoV-19/Norway/OUS-24817/2022** | EPI_ISL_12982839 | | Oslo University Hospital, Department of Microbiology | Norwegian Institute of Public Health, Department of Virology | Mona Holberg-Petersen et al |
| **hCoV-19/Bonaire/BQ-RIVM-101080/2022** | EPI_ISL_12954006 | | National Institute for Public Health and the Environment (RIVM) | National Institute for Public Health and the Environment (RIVM) | Adam Meijer et al |
| **hCoV-19/Montenegro/CO-00688_MNE000_1622362202/2022** | EPI_ISL_12280145 | | Laboratory for Molecular Diagnostics,IPHMN | Eurofins Genomics Europe Sequencing GmbH | Danijela Vujošević et al |
| **hCoV-19/Croatia/36186/2022** | EPI_ISL_12686438 | | Hrvatski zavod za javno zdravstvo | Hrvatski zavod za javno zdravstvo | Irena Tabain et al |
| **hCoV-19/Bonaire/BQ-RIVM-100069/2022** | EPI_ISL_12782627 | | National Institute for Public Health and the Environment (RIVM) | National Institute for Public Health and the Environment (RIVM) | Adam Meijer et al |
| **hCoV-19/Finland/THL-202212700/2022** | EPI_ISL_12916706 | | Islab, Pohjois-Savon aluelaboratorio | Expert Microbiology, National Institute for Health and Welfare | Soile Blomqvist et al |
| **hCoV-19/Peru/LIM-INS-18582/2022** | EPI_ISL_13408418 | | Laboratorio de Referencia Nacional de Virus Respiratorios. Centro Nacional de Salud Publica. Instituto Nacional de Salud Peru. | Laboratorio de Referencia Nacional de Virus Respiratorios. Centro Nacional de Salud Publica. Instituto Nacional de Salud Peru. | Carlos Padilla Rojas et al |
| **hCoV-19/Uzbekistan/4_1466-CAT/2022** | EPI_ISL_15941882 | | SI"Zangiota No.1 and 2 special hospital for the treatment of patients with coronovirus infection" | Center for Advanced Technologies | Dalimova D.A. et al |
| **hCoV-19/Tunisia/4273/2022** | EPI_ISL_15022516 | | Laboratory of Microbiology, La Rabta Hospital | Clinical and Experimental Pharmacology Lab, LR16SP02, National Center of Pharmacovigilance, University of Tunis El Manar, Tunis, Tunisia. 2- Ministry of Health, National Observatory of New and Emerging Diseases, 1006, Tunis, Tunisia | Sana Ferjani et al |
| **hCoV-19/Tunisia/S-2607/2022** | EPI_ISL_16185981 | | Laboratoire de Virologie Clinique, Institut Pasteur de Tunis | Laboratoire de Virologie Clinique, Institut Pasteur de Tunis | Chouikha Anissa et al |
| **hCoV-19/India/DH-ICMR-NIV-INSACOG-GSEQ-9705/2022** | EPI_ISL_9973098 | | ICMR-National Institute of Virology - INSACOG | NIV Influenza | Dr. Varsha Potdar et al |
| **hCoV-19/Jilin/JSY-CC5/2022** | EPI_ISL_18435548 | | Changchun Veterinary Research Institute, Chinese Academy of Agricultural Sciences, Changchun, China | Changchun Veterinary Research Institute, Chinese Academy of Agricultural Sciences, Changchun, China | Yuewei Gao et al |
| **hCoV-19/Fujian/2022XG-2040/2022** | EPI_ISL_12046160 | | Fujian Center for Disease Control and Prevention | Fujian Center for Disease Control and Prevention | Lin Qi et al |
| **hCoV-19/Hubei/IVDC-552/2022** | EPI_ISL_11873915 | | Hubei Provincial Center for Disease Control and Prevention | National Institute for Viral Disease Control and Prevention, China CDC | Bin Fang et al |
| **hCoV-19/Chongqing/IVDC-610/2022** | EPI_ISL_12030375 | | Chongqing Center for disease control and prevertion | National Institute for Viral Disease Control and Prevention, China CDC | Sheng Ye et al |
| **hCoV-19/Liaoning/IVDC-583/2022** | EPI_ISL_11905848 | | Liaoning Center for Disease Prevention and Control | National Institute for Viral Disease Control and Prevention, China CDC | Lingling Mao et al |
| **hCoV-19/India/GA-ICMR-NIV-INSACOG-GSEQ-11250/2022** | EPI_ISL_14149530 | | ICMR-National Institute of Virology - INSACOG | NIV Influenza | Dr. Varsha Potdar et al |
| **hCoV-19/SriLanka/aicbu5511/2022** | EPI_ISL_13232325 | | Allergy, Immunology and Cell Biology Unit (AICBU) | Allergy, Immunology and Cell Biology Unit (AICBU) | Chandima Jeewandara et al |
| **hCoV-19/Lesotho/N47909/2022** | EPI_ISL_14952352 | | National Reference Laboratories | National Institute for Communicable Diseases of the National Health Laboratory Service | Maphalala G et al |
| **hCoV-19/Iraq/CPHL-180122-18/2022** | EPI_ISL_9517284 | | Central Public Health Laboratory | Central Public Health Laboratory | Dr. Hani Mousa Al-Akabi et al |
| **hCoV-19/SouthAfrica/NICD-N35503/2022** | EPI_ISL_11270141 | | AMPATH | National Institute for Communicable Diseases of the National Health Laboratory Service | Amoako DG et al |
| **hCoV-19/Peru/LIM-INS-16743/2022** | EPI_ISL_12590973 | | Laboratorio de Referencia Nacional de Virus Respiratorios. Centro Nacional de Salud Publica. Instituto Nacional de Salud Peru. | Laboratorio de Referencia Nacional de Virus Respiratorios. Centro Nacional de Salud Publica. Instituto Nacional de Salud Peru. | Carlos Padilla Rojas et al |
| **hCoV-19/Namibia/N43550/2022** | EPI_ISL_14503120 | | Namibia Institute of Pathology (NIP) | National Institute for Communicable Diseases of the National Health Laboratory Service | Van Rooyen G et al |
| **hCoV-19/Niger/CE2871/2022** | EPI_ISL_13285099 | | CERMES | CERMES | Lagare Adamou et al |
| **hCoV-19/CotedIvoire/IPCI-DVE-COV800899/2022** | EPI_ISL_15813211 | | Departement of Epidemic Viruses | Departement of Epidemic Viruses Institut Pasteur of CotedIvoire Abidjan | Hervé Albéric Adje Kadjo et al |
| **hCoV-19/Togo/TOG-5461/2022** | EPI_ISL_17075678 | | LNR/CHU-SO | BIOLIM/FSS-UL | A.A. KONOU et al |
| **hCoV-19/Niger/CE-5361/2022** | EPI_ISL_13285208 | | CERMES | CERMES | Lagare Adamou et al |
| **hCoV-19/USA/TN-ASC-210768528/2022** | EPI_ISL_15224050 | | Aegis Sciences Corporation | Aegis Sciences Corporation | Cyndi Clark et al |
| **hCoV-19/Cameroon/LNSP-004-SW06469/2022** | EPI_ISL_18002424 | | NPHL-Cameroon | National Public Health Laboratory, Cameroon | Dr Essomba Rene Ghislain et al |
| **hCoV-19/Uruguay/DLSP_2945/2022** | EPI_ISL_16389503 | | Departamento Laboratorios de Salud Pública (DLSP) Ministerio de Salud Pública | Departamento Laboratorios de Salud Pública (DLSP) Ministerio de Salud Pública | Cortinas et al |
| **hCoV-19/Malaysia/IMR_OS4233/2022** | EPI_ISL_14977183 | | Institute for Medical Research, Infectious Disease Research Centre, National Institutes of Health, Ministry of Health Malaysia | Institute for Medical Research, Infectious Disease Research Centre, National Institutes of Health, Ministry of Health Malaysia | Kalyanasundram J et al |
| **hCoV-19/Uruguay/DLSP_3260/2022** | EPI_ISL_16376115 | | Departamento Laboratorios de Salud Pública (DLSP) Ministerio de Salud Pública | Departamento Laboratorios de Salud Pública (DLSP) Ministerio de Salud Pública | Cortinas et al |
| **hCoV-19/Mozambique/INS-PMB0600984/2022** | EPI_ISL_14015058 | | Biotechnology and Genetics Laboratory - Instituto Nacional de Saúde | Biotechnology and Genetics Laboratory - Instituto Nacional de Saúde | Sofia Viegas et al |
| **hCoV-19/Lesotho/N47924/2022** | EPI_ISL_14952322 | | National Reference Laboratories | National Institute for Communicable Diseases of the National Health Laboratory Service | Maphalala G et al |
| **hCoV-19/NewZealand/22YA4760/2022** | EPI_ISL_16050371 | | Wellington SCL | Institute of Environmental Science and Research (ESR) | Sarah Cockerton et al |

| | | | | | |
|---|---|---|---|---|---|
| **hCoV-19/Lesotho/N47911/2022** | EPI_ISL_14952334 | | National Reference Laboratories | National Institute for Communicable Diseases of the National Health Laboratory Service | Maphalala G et al |
| **hCoV-19/Ireland/D-NVRL-S22IRL00367722/2022** | EPI_ISL_14025939 | | National Virus Reference Laboratory | National Virus Reference Laboratory | Zoe Yandle et al |
| **hCoV-19/Mozambique/INS-PCS0022563/2022** | EPI_ISL_14015050 | | Biotechnology and Genetics Laboratory - Instituto Nacional de Saúde | Biotechnology and Genetics Laboratory - Instituto Nacional de Saúde | Sofia Viegas et al |
| **hCoV-19/Djibouti/NAMRU3_H151/2022** | EPI_ISL_15748337 | | Naval Medical Research Unit No. 3 | Naval Medical Research Center Biological Defense Research Directorate | J. Alexander Chitty et al |
| **hCoV-19/Liechtenstein/FL-Risch-0927R13854/2022** | EPI_ISL_15285515 | | Dr. Risch Ostschweiz AG | Dr Risch Laboratory | Nadia Wohlwend et al |
| **hCoV-19/Ecuador/USFQ-3946/2022** | EPI_ISL_14208892 | | Institute of Microbiology, Universidad San Francisco de Quito | Institute of Microbiology, Universidad San Francisco de Quito | Belén Prado-Vivar et al |
| **hCoV-19/USA/MS-UMMC-S257P06-523423/2022** | EPI_ISL_14425902 | OP208902.1 | University of Mississippi Medical Center, Department of Pathology | University of Mississippi Medical Center, Molecular and Genomics Core Facility | Ashley C. Johnson et al |
| **hCoV-19/DominicanRepublic/2038376-LNSPDD/2022** | EPI_ISL_14276614 | | Laboratorio Nacional de Salud Pública Dr. Defilló | Laboratorio Nacional de Salud Pública Dr. Defilló | Isaac Miguel Sánchez et al |
| **hCoV-19/USA/LA-EVTL17429/2022** | EPI_ISL_15762632 | | Willis-Knighton Medical Center Hospital Laboratory | LSUHS Emerging Viral Threat Laboratory | Lorie M. Atkins et al |
| **hCoV-19/Peru/LIM-INS-31113/2022** | EPI_ISL_15723610 | | Laboratorio de Referencia Nacional de Virus Inmunoprevenibles. Centro Nacional de Salud Publica. Instituto Nacional de Salud Peru. | Laboratorio de Referencia Nacional de Virus Inmunoprevenibles. Centro Nacional de Salud Publica. Instituto Nacional de Salud Peru. | Carlos Padilla Rojas et al |
| **hCoV-19/DominicanRepublic/2079810-LNSPDD/2022** | EPI_ISL_15647848 | | Laboratorio Nacional de Salud Pública Dr. Defilló | Laboratorio Nacional de Salud Pública Dr. Defilló | Isaac Miguel Sánchez et al |
| **hCoV-19/Zhejiang/ZPCDCP-146/2022** | EPI_ISL_17261648 | | Zhejiang Provincial Center for Disease Control and Prevention | Zhejiang Provincial Center for Disease Control and Prevention | Yi Sun et al |
| **hCoV-19/DominicanRepublic/2064316-LNSPDD/2022** | EPI_ISL_15164628 | | Laboratorio Nacional de Salud Pública Dr. Defilló | Laboratorio Nacional de Salud Pública Dr. Defilló | Isaac Miguel Sánchez et al |
| **hCoV-19/DominicanRepublic/2066705-LNSPDD/2022** | EPI_ISL_15164641 | | Laboratorio Nacional de Salud Pública Dr. Defilló | Laboratorio Nacional de Salud Pública Dr. Defilló | Isaac Miguel Sánchez et al |
| **hCoV-19/USA/IN-CDC-LC0686513/2022** | EPI_ISL_13308287 | ON769605.1 | Laboratory Corporation of America | Centers for Disease Control and Prevention Division of Viral Diseases, Pathogen Discovery | Dakota Howard et al |
| **hCoV-19/Eswatini/N47806/2022** | EPI_ISL_14501254 | | The Luke Commission | National Institute for Communicable Diseases of the National Health Laboratory Service | Maphalala G et al |
| **hCoV-19/Ethiopia/BETIN-21356071823/2022** | EPI_ISL_18904549 | | Health Biotechnology, Bio and Emerging Technology Institute | Health Biotechnology, Bio and Emerging Technology Institute | Bitew et al |
| **hCoV-19/DRC/INRB-RDC-776/2022** | EPI_ISL_14404659 | | Pathogen Genomics Lab, National Institute for Biomedical Research (INRB) | Pathogen Genomics Lab, National Institute for Biomedical Research (INRB) | Placide Mbala-Kingebeni et al B |
| **hCoV-19/Senegal/DKR-UVR-1361/2022** | EPI_ISL_15321646 | | Institut Pasteur de Dakar | Institut Pasteur de Dakar | Ndongo Dia et al |
| **hCoV-19/DominicanRepublic/2010121-LNSPDD/2022** | EPI_ISL_13811387 | | Laboratorio Nacional de Salud Pública Dr. Defilló | Laboratorio Nacional de Salud Pública Dr. Defilló | Isaac Miguel Sánchez et al |
| **hCoV-19/SouthAfrica/SU-NHLS_4828/2022** | EPI_ISL_12705855 | | PathCare, Cape Town | Division of Medical Virology, National Health Laboratory Service (NHLS), Tygerberg Hospital / Stellenbosch University | Jean Maritz et al |
| **hCoV-19/Cyprus/BMV7986/2022** | EPI_ISL_18476596 | | Ammochostos General Hospital | Laboratory of Biotechnology and Molecular Virology, University of Cyprus | Leondios G. Kostrikis et al. |
| **hCoV-19/Guangdong/SZTH-039/2022** | EPI_ISL_17008292 | | Shenzhen Third People's Hospital | Shenzhen Key Laboratory of Pathogen and Immunity | Lin Cheng et al |
| **hCoV-19/Uganda/547/2022** | EPI_ISL_13955175 | | Mulago National Referral Hospital | Central Public Health Laboratories | Alisen Ayitewala et al |
| **hCoV-19/Mozambique/INS-PNV0082367/2022** | EPI_ISL_15808035 | | Biotechnology and Genetics Laboratory - Instituto Nacional de Saúde | Biotechnology and Genetics Laboratory - Instituto Nacional de Saúde | Sofia Viegas et al |
| **hCoV-19/Uganda/663/2022** | EPI_ISL_15718863 | | | Central Public Health Laboratories | Alisen Ayitewala et al |
| **hCoV-19/Botswana/R131B64_BHP_1058644/2022** | EPI_ISL_13345360 | | Palapye Primary Hospital Laboratory | Botswana Harvard HIV Reference Laboratory | Sikhulile Moyo et al |
| **hCoV-19/Comoros/SS10399/2022** | EPI_ISL_15585870 | | Institut National de Recherche pour l'Agriculture, la Peche et l'Environnement (INRAPE),Comoros | KEMRI-Wellcome Trust Research Programme,Kilifi | Githinji G. et al |
| **hCoV-19/Montenegro/CO-00892_MNE000_2063892202/2022** | EPI_ISL_14509027 | | Laboratory for Molecular Diagnostics,IPHMN | Eurofins Genomics Europe Sequencing GmbH | Danijela Vujošević et al |
| **hCoV-19/Cameroon/LNSP-003-82159/2022** | EPI_ISL_17604103 | | NPHL-Cameroon | National Public Health Laboratory, Cameroon | Dr Essomba Rene Ghislain et al |
| **hCoV-19/DRC/INRB-RDC-1106/2022** | EPI_ISL_14815703 | | Laboratoire Rodolphe Mérieux (INRB Goma) | Pathogen Genomics Lab, National Institute for Biomedical Research (INRB) | Placide Mbala-Kingebeni et al B |
| **hCoV-19/Peru/LAL-INS-22507/2022** | EPI_ISL_14241646 | | Laboratorio de Referencia Nacional de Virus Inmunoprevenibles. Centro Nacional de Salud Publica. Instituto Nacional de Salud Peru. | Laboratorio de Referencia Nacional de Virus Inmunoprevenibles. Centro Nacional de Salud Publica. Instituto Nacional de Salud Peru. | Carlos Padilla Rojas et al |
| **hCoV-19/Chile/RM-92900/2022** | EPI_ISL_14264416 | | Genetica Molecular and Subdepartamento de Virologia ISP Chile | Instituto de Salud Publica de Chile | Constanza Campano et al |
| **hCoV-19/USA/NM-UNM-ED03075/2022** | EPI_ISL_14418858 | OP564265.1 | Univeristy of New Mexico Hospital | Center for Global Health, University of New Mexico Health Sciences Center | Daryl Domman et al |
| **hCoV-19/Venezuela/Dtt999/2022** | EPI_ISL_15975690 | | IVIC | Laboratorio de Virologia Molecular | Rossana C Jaspe et al |
| **hCoV-19/Mexico/29886/2022** | EPI_ISL_17658374 | | Laboratory of Respiratory Viruses and Measles, Oswaldo Cruz Institute, FIOCRUZ | Laboratory of Respiratory Viruses and Measles, Oswaldo Cruz Institute, FIOCRUZ | Paola Resende et al |
| **hCoV-19/Congo/BZV-LNSP-.09/2022** | EPI_ISL_16275212 | | Laboratoire National de Santé Publique de Brazzaville | Laboratoire National de Santé Publique de Brazzaville | Felix KOUKOUIKILA-KOUSSOUNDA et al |
| **hCoV-19/Fujian/FCDC-XM011705HG/2024** | EPI_ISL_18805103 | | Institute for the Prevention and Control of Infectious Diseases, Fujian Provincial Center for Disease Control and Prevention | Institute for the Prevention and Control of Infectious Diseases, Fujian Provincial Center for Disease Control and Prevention | Qi Lin et al |
| **hCoV-19/India/GJ-INSACOG-GBRC13661/2023** | EPI_ISL_18694676 | | Sterling Accuris Diagnostic, Mehsana | Gujarat Biotechnology Research Centre-INSACOG | Priyank Chavda et al |
| **hCoV-19/Uruguay/DLSP_Mu56/2024** | EPI_ISL_18899296 | | Departamento Laboratorios de Salud Pública (DLSP) Ministerio de Salud Pública | Departamento Laboratorios de Salud Pública (DLSP) Ministerio de Salud Pública | Cortinas et al |
| **hCoV-19/Brazil/PB-002480-IEC/2024** | EPI_ISL_18902084 | | Central Public Health Laboratory of the State Paraiba | Evandro Chagas Institute | Mirleide Cordeiro dos Santos et al |
| **hCoV-19/Russia/SPE-RII-MH165548S/2024** | EPI_ISL_18845578 | | Saint Petersburg Dermatovenerologic Dispensary No 1 | WHO National Influenza Centre Russian Federation | Andrey Komissarov et al |
| **hCoV-19/India/GJ-INSACOG-GBRC13805/2024** | EPI_ISL_18856262 | | Pathocare Pathology Laboratory, Vadodara | Gujarat Biotechnology Research Centre-INSACOG | Rameshchandra Pandit et al |
| **hCoV-19/Oman/CPHL_52338551/2023** | EPI_ISL_18815421 | | Central Public Health Laboratories-Sultanat of Oman | Central Public Health Laboratories-Sultanat of Oman | Samiha Al Kharusi et al |
| **hCoV-19/Laos/IPL-78506/2023** | EPI_ISL_18877133 | | National Centre for Laboratory and Epidemiology (NCLE) | Institut Pasteur du Laos | Kedkeo INTAVONG et al |
| **hCoV-19/SouthAfrica/SU-NHLS_6733/2023** | EPI_ISL_18706512 | | PathCare Reference Laboratory, PathCare Park, N1 City, Cape Town | Division of Medical Virology, National Health Laboratory Service (NHLS), Tygerberg Hospital / Stellenbosch University | Tongai Maponga et al |
| **hCoV-19/Oman/CPHL_7237542/2023** | EPI_ISL_18815416 | | Central Public Health Laboratories-Sultanat of Oman | Central Public Health Laboratories-Sultanat of Oman | Samiha Al Kharusi et al |
| **hCoV-19/USA/CA-CDC-LC1073875/2023** | EPI_ISL_18621446 | | Laboratory Corporation of America | Centers for Disease Control and Prevention Division of Viral Diseases, Pathogen Discovery | Dakota Howard et al |
| **hCoV-19/Laos/IPL-78507/2023** | EPI_ISL_18877134 | | National Centre for Laboratory and Epidemiology (NCLE) | Institut Pasteur du Laos | Kedkeo INTAVONG et al |
| **hCoV-19/Vietnam/HCMC-S05-M189/2023** | EPI_ISL_18909563 | | Oxford University Clinical Research Unt | Oxford University Clinical Research Unt | Nguyen Van Vinh Chau et al |
| **hCoV-19/Austria/AGES-AZ-1661246/2023** | EPI_ISL_18499932 | | Zentrum für Virologie / Medizinische Universität Wien | AGES IMED Vienna | Blaschitz M. et al |
| **hCoV-19/USA/NJ-CDC-QDX88429436/2023** | EPI_ISL_18644431 | | Quest Diagnostics Incorporated | Centers for Disease Control and Prevention Division of Viral Diseases, Pathogen Discovery | Dakota Howard et al |
| **hCoV-19/India/un-SPHL-44/2023** | EPI_ISL_18904625 | | Public Health, State Public Health Laboratory Chennai | Public Health, State Public Health Laboratory Chennai | Raju et al |
| **hCoV-19/Lebanon/AU-7189/2024** | EPI_ISL_18939699 | | American University of Beirut Medical Center | Center for Infectious Diseases Research (CIDR) | Ghassan Dbaibo et al |
| **hCoV-19/India/TG-CDFD-GMC-1103/2024** | EPI_ISL_18884613 | | Gandhi Medical College | CDFD | Asmita Gupta et al |
| **hCoV-19/India/KA-RFNB-21218/2024** | EPI_ISL_18944587 | | Aster Clinical Labs Llp | National Centre for Biological Sciences, TIFR - Rockefeller Foundation | Vinay Kumar et al |
| **hCoV-19/Ireland/LK-UHL-673928/2023** | EPI_ISL_18549615 | | University Hospital Limerick | Molecular Laboratory, Serology and Virology at Pathology Department | Diane Slater-Kinghorn et al |
| **hCoV-19/Bulgaria/NC-023963_Run46/2023** | EPI_ISL_18945201 | | National Center of Infectious and Parasitic Diseases | National Center of Infectious and Parasitic Diseases | Ivelina Trifonova et al |
| **hCoV-19/Kenya/HF232128/2023** | EPI_ISL_18788222 | | KEMRI-Wellcome Trust Research Programme,Kilifi | KEMRI-Wellcome Trust Research Programme,Kilifi | Lambisia A. et al |
| **hCoV-19/Cambodia/KSH240006/2024** | EPI_ISL_18814612 | | National Institute of Public Health | National Institute of Public Health | Chhorvann CHHEA et al |
| **hCoV-19/Venezuela/Dtto_VP_A/2023** | EPI_ISL_18679956 | | IVIC | Laboratorio de Virologia Molecular | Rossana C Jaspe et al |
| **hCoV-19/Germany/MV-RKI-I-1142131/2024** | EPI_ISL_18896477 | | Limbach - MVZ Labor Westmecklenburg Schmudlach-Oswald-Kettermann & Kollegen | Robert Koch Institute | Antje Schwarz et al |
| **hCoV-19/Belgium/ULG-301676/2023** | EPI_ISL_18673898 | | CHR de La Citadelle | GIGA Medical Genomics | Keith Durkin et al |
| **hCoV-19/Sichuan/CDCDC-19-CX160/2024** | EPI_ISL_18947795 | | Chengdu Center of Disease Control and Prevention | Microbiology laboratory, Sichuan Center of Disease Control and Prevention | Xie et al |
| **hCoV-19/Botswana/R235B025_BHP_124002152I/2024** | EPI_ISL_18800047 | | Sidilega Private Hospital Laboratory | Botswana Harvard HIV Reference Laboratory | Sikhulile Moyo et al |
| **hCoV-19/India/un-SPHL-46/2023** | EPI_ISL_18904596 | | Public Health, State Public Health Laboratory Chennai | Public Health, State Public Health Laboratory Chennai | Raju et al |
| **hCoV-19/Namibia/NIP0117/2024** | EPI_ISL_18884736 | | NIP_Windhoek_Reference laboratory | NIP_Windhoek_Reference laboratory | Ndahafa Frans et al |
| **hCoV-19/Germany/BE-ChVir45211/2023** | EPI_ISL_18624804 | | Charité Universitätsmedizin Berlin, Institut für Virologie/Labor Berlin | Charité Universitätsmedizin Berlin, Institut für Virologie | Victor M Corman et al |
| **hCoV-19/Fujian/FCDC-FZ1512078001/2023** | EPI_ISL_18556238 | | Institute for the Prevention and Control of Infectious Diseases, Fujian Provincial Center for Disease Control and Prevention | Institute for the Prevention and Control of Infectious Diseases, Fujian Provincial Center for Disease Control and Prevention | Qi Lin et al |
| **hCoV-19/Switzerland/VD-CHUV-GEN18815/2023** | EPI_ISL_18686344 | | CHUV | Laboratory of genomics and metagenomics, Institute of Microbiology, University Hospital Centre and University of Lausanne | Trestan Pillonel et al |
| **hCoV-19/Australia/NSW-SAVID-16562/2024** | EPI_ISL_18918410 | | Area of Virology, Serology and Virology Division (SAViD), New South Wales Health Pathology Randwick | Virology Research Laboratory; Area of Virology, Serology and Virology Division (SAViD), New South Wales Health Pathology Randwick | Foster et al |
| **hCoV-19/Brunei/MGS-7124000881/2024** | EPI_ISL_18939056 | | National Virology Reference Laboratory | Microbial Genomic Services | Zainun Zaini et al |
| **hCoV-19/Ireland/G-GUH-N2142/2023** | EPI_ISL_18722263 | | GUH | UHG | Wendy Brennan et al |
| **hCoV-19/Netherlands/ZH-RIVM-137715/2024** | EPI_ISL_18878192 | | National Institute for Public Health and the Environment (RIVM) | National Institute for Public Health and the Environment (RIVM) | Adam Meijer et al |
| **hCoV-19/Ireland/G-GUH-N2353/2024** | EPI_ISL_18933873 | | GUH | UHG | Wendy Brennan et al |
| **hCoV-19/Taiwan/CDC-F90690/2024** | EPI_ISL_18838506 | | Centers for Disease Control, R.O.C. (Taiwan) | Centers for Disease Control, R.O.C. (Taiwan) | Yu-Chi Lin et al |
| **hCoV-19/ElSalvador/60-24-LNSP/2024** | EPI_ISL_18899104 | | Hospital Santa Ana | Laboratorio Nacional de Salud Publica. | Laboratorio Nacional de Salud Publica de El Salvador - Instituto Nacional de Salud et al |
| **hCoV-19/Poland/Kce-277/2024** | EPI_ISL_18918586 | | LABORATORIUM BADAŃ KLINICZNYCH WSSE w OPOLU | Wojewódzka Stacja Sanitarno-Epidemiologiczna w Katowicach | Beata Rozwadowska et al |
| **hCoV-19/Slovakia/ruvztn-32541/2024** | EPI_ISL_18889679 | | Regional Public Health Authority with the seat in Trenčín, Slovakia | Laboratory of Genomics and Bioinformatics, Comenius University Science Park | Nikola Slivková et al |
| **hCoV-19/Curacao/CW-RIVM-137929/2024** | EPI_ISL_18937293 | | National Institute for Public Health and the Environment (RIVM) | National Institute for Public Health and the Environment (RIVM) | Adam Meijer et al |
| **hCoV-19/Romania/B-563812/2024** | EPI_ISL_18910777 | | Cantacuzino National Military-Medical Institute, Viral Respiratory Infections Laboratory | Cantacuzino National Military-Medical Institute, Viral Respiratory Infections Laboratory | Catalina Pascu et al |
| **hCoV-19/USA/KS-CDC-QDX92231752/2024** | EPI_ISL_18920614 | | Quest Diagnostics Incorporated | Centers for Disease Control and Prevention Division of Viral Diseases, Pathogen Discovery | Dakota Howard et al |
| **hCoV-19/Liaoning/LNCDC-057-1/2024** | EPI_ISL_18930360 | | Institute for Communicable and Infectious Diseases Prevention and Control, Liaoning Center for Disease Control and Prevention | Institute for Communicable and Infectious Diseases Prevention and Control, Liaoning Center for Disease Control and Prevention | Mao et al |
| **hCoV-19/Denmark/DCGC-670636/2024** | EPI_ISL_18850036 | | Department of Bacteria, Parasites and Fungi, Statens Serum Institut, Copenhagen, Denmark | Statens Serum Institut Bioinformatics and Microbial Genomics | Povilas Matusevicius on behalf of the Danish Covid-19 Genome Consortium et al |
| **hCoV-19/Anhui/BZCDC-216/2024** | EPI_ISL_18941269 | | Clinical laboratory, Bozhou Municipal Center for Disease Control & Prevention | Clinical laboratory, Bozhou Municipal Center for Disease Control & Prevention | Qingqiao et al |
| **hCoV-19/Thailand/NIC_KBI_118/2024** | EPI_ISL_18854154 | | Regional Medical Sciences Center 11/1 Phuket | National Institute of Health, Department of Medical Sciences, Ministry of Public Health, Thailand | Pilailuk Okada et al |
| **hCoV-19/Indonesia/JK-GSILab-OUCRU-1156257/2024** | EPI_ISL_18909384 | | OUCRU | Genomik Solidaritas Indonesia Laboratorium | Rohyani et al |
| **hCoV-19/Israel/SMC-7116141/2023** | EPI_ISL_18683470 | | Shamir Medical Center (Asaf Harofe) | Shamir Medical Center (Asaf Harofe) | Nir Rainy et al |
| **hCoV-19/Vietnam/HCMC-S01-M189/2023** | EPI_ISL_18909559 | | Oxford University Clinical Research Unt | Oxford University Clinical Research Unt | Nguyen Van Vinh Chau et al |
| **hCoV-19/Fujian/FCDC-FZLG0492/2024** | EPI_ISL_18930376 | | Institute for the Prevention and Control of Infectious Diseases, Fujian Provincial Center for Disease Control and Prevention | Institute for the Prevention and Control of Infectious Diseases, Fujian Provincial Center for Disease Control and Prevention | Qi Lin et al |
| **hCoV-19/Malaysia/C19UMB4683/2024** | EPI_ISL_18935690 | | Hospital Melaka | UKM Medical Molecular Biology Institute (UMBI) | Rahman Jamal et al |
| **hCoV-19/Singapore/R6MQ36/2024** | EPI_ISL_18831192 | | National Public Health Laboratory, National Centre for Infectious Diseases | National Public Health Laboratory, National Centre for Infectious Diseases | Yichen Ding et al |
| **hCoV-19/England/CLIMB-CM7YGERJ/2023** | EPI_ISL_18688125 | | Respiratory Virus Unit, Microbiology Services Colindale, Public Health England | UK Health Security Agency (UKHSA) | PHE Covid Sequencing Team et al |
| **hCoV-19/Luxembourg/LNS7803004/2023** | EPI_ISL_18727960 | | Hospital Center Luxembourg | Laboratoire National de Santé Microbiology | Anke Wienecke-Baldacchino et al |
| **hCoV-19/TrinidadandTobago/433178/2024** | EPI_ISL_18880311 | | Trinidad Public Health Laboratory | Carrington Lab, Department of Preclinical Sciences, Faculty of Medical Sciences, The University of the West Indies, St Augustine Campus | Nikita S. D. Sahadeo et al |
| **hCoV-19/Kosovo/48_Jan/2024** | EPI_ISL_18825170 | | NIPH Kosovo | Department of Microbiology, National Institute of Public Health of Kosovo | Xhevat Jakupi et al |
| **hCoV-19/USA/NE-NPHL24-284/2024** | EPI_ISL_18892240 | | Nebraska Public Health Laboratory | NPHL COVID-19 Response Team | Paulino et al |
| **hCoV-19/TrinidadandTobago/433313/2024** | EPI_ISL_18880314 | | Trinidad Public Health Laboratory | Carrington Lab, Department of Preclinical Sciences, Faculty of Medical Sciences, The University of the West Indies, St Augustine Campus | Nikita S. D. Sahadeo et al |
| **hCoV-19/Senegal/25576845/2023** | EPI_ISL_18713710 | | Institut Pasteur de Dakar | Institut Pasteur de Dakar | Jallow et al |
| **hCoV-19/Ukraine/84611/2024** | EPI_ISL_18919249 | | Reference laboratory for diagnostics of HIV/AIDS, virological and especially dangerous pathogens | Reference laboratory for diagnostics of HIV/AIDS, virological and especially dangerous pathogens | Iryna Demchyshyna et al |
| **hCoV-19/Senegal/D755741010/2024** | EPI_ISL_18783719 | | Iressef Genomics lab | Institute for Health Research - Epidemiological Surveillance and Training (IRESSEF) | Souleymane MBOUP et al |
| **hCoV-19/Oman/CPHL_7236365/2023** | EPI_ISL_18690692 | | Central Public Health Laboratories-Sultanat of Oman | Central Public Health Laboratories-Sultanat of Oman | Yathrib Al Zakwani et al |
| **hCoV-19/Pakistan/IDRL-AKU-KHI-ML0009-S12/2024** | EPI_ISL_18745112 | | Clinical Lab, Aga Khan University Hospital | Infectious Diseases Research Laboratory, Department of Pediatrics and Child Health | Samiah Kanwar et al |
| **hCoV-19/Sichuan/LSZCDC-20231222-19/2024** | EPI_ISL_18947828 | | Liangshan Center of Disease Control and Prevention | Microbiology laboratory, Sichuan Center of Disease Control and Prevention | Chen et al |
| **hCoV-19/Bangladesh/IEDCR-1625075/2024** | EPI_ISL_18851601 | | Institute of Epidemiology, Disease Control and Research (IEDCR) | Institute of Epidemiology, Disease Control and Research (IEDCR) | Manjur Hossain Khan et al |
| **hCoV-19/NewZealand/24YA0303/2023** | EPI_ISL_18851109 | | Pathlab Tauranga | Institute of Environmental Science and Research (ESR) | Sarah Cockerton et al |
| **hCoV-19/Chile/MA-13695/2024** | EPI_ISL_18946081 | | Genetica Molecular and Subdepartamento de Virologia ISP Chile | Instituto de Salud Publica de Chile | Mariela Guajardo et al |
| **hCoV-19/USA/KS-USAFSAM-S22180/2024** | EPI_ISL_18871268 | | US Air Force School of Aerospace Medicine | US Air Force School of Aerospace Medicine | Anthony Fries et al |
| **hCoV-19/Brazil/SC-FIOCRUZ-435/2024** | EPI_ISL_18934139 | | Laboratorio Central de Saude Publica do Estado de Santa Catarina (LACEN/SC) | Instituto Oswaldo Cruz FIOCRUZ - Laboratory of Respiratory Viruses and Measles (LVRS) | Paola Resende et al |
| **hCoV-19/Hubei/HCDC-XY3801/2024** | EPI_ISL_18941314 | | Xiangyang Center for Disease Control and Prevention | Hubei Provincial Center for Disease Control and Prevention Center of Pathogenic Microorganism Detection | Yongzhong Jiang et al |
| **hCoV-19/France/IDF-IPP02788/2024** | EPI_ISL_18913130 | | Labo Analyses Med | National Reference Center for Viruses of Respiratory Infections, Institut Pasteur, Paris | Léa Avon et al |
| **hCoV-19/Denmark/DCGC-670795/2024** | EPI_ISL_18938963 | | Department of Bacteria, Parasites and Fungi, Statens Serum Institut, Copenhagen, Denmark | Statens Serum Institut Bioinformatics and Microbial Genomics | Povilas Matusevicius on behalf of the Danish Covid-19 Genome Consortium et al |
| **hCoV-19/Yunnan/YNCDC-0016/2024** | EPI_ISL_18876921 | | The department for prevention and control of acute infectious diseases, Yunnan Center for Disease Control and Prevention | The department for prevention and control of acute infectious diseases, Yunnan Center for Disease Control and Prevention | Zhang et al B |
| **hCoV-19/Shanghai/SCDC-HG1124-4023/2023** | EPI_ISL_18634016 | | Microbe lab, Shanghai Municipal Center for Disease Control & Prevention | Microbe lab, Shanghai Municipal Center for Disease Control & Prevention | Chen et al |
| **hCoV-19/Chongqing/YBCDC-0094/2024** | EPI_ISL_18926253 | | Chongqing Yubei District Center for Disease Control and Prevention | Chongqing Yubei District Center for Disease Control and Prevention | Jin Yan et al |
| **hCoV-19/Chongqing/YBCDC-S6701/2024** | EPI_ISL_18944031 | | Chongqing Yubei District Center for Disease Control and Prevention | Chongqing Yubei District Center for Disease Control and Prevention | Jin Yan et al |
| **hCoV-19/Hubei/HCDC-WH3623/2023** | EPI_ISL_18797879 | | Wuhan Center for Disease Control and Prevention | Hubei Provincial Center for Disease Control and Prevention Center of Pathogenic Microorganism Detection | Yongzhong Jiang et al |
| **hCoV-19/Fujian/FCDC-XM010303HG/2023** | EPI_ISL_18792236 | | Institute for the Prevention and Control of Infectious Diseases, Fujian Provincial Center for Disease Control and Prevention | Institute for the Prevention and Control of Infectious Diseases, Fujian Provincial Center for Disease Control and Prevention | Qi Lin et al |
| **hCoV-19/Taiwan/NCKUH-823624000363/2024** | EPI_ISL_18819963 | | Virology Lab,Department of Pathology, National Cheng Kung University Hospital | Virology Lab,Department of Pathology, National Cheng Kung University Hospital | Huey-Pin Tsai et al |
| **hCoV-19/Oman/CPHL_7237265/2023** | EPI_ISL_18690688 | | Central Public Health Laboratories-Sultanat of Oman | Central Public Health Laboratories-Sultanat of Oman | Yathrib Al Zakwani et al |
| **hCoV-19/Senegal/25577078/2024** | EPI_ISL_18877940 | | Institut Pasteur de Dakar | Institut Pasteur de Dakar | Jallow et al |

| | | | | | |
|---|---|---|---|---|---|
| **hCoV-19/Senegal/D918538040/2024** | EPI_ISL_18783733 | | Iressef Genomics lab | Institute for Health Research - Epidemiological Surveillance and Training (IRESSEF) | Souleymane MBOUP et al |
| **hCoV-19/Ireland/CO-CUH-SC0002/2024** | EPI_ISL_18737416 | | Cork University Hospital | Cork University Hospital | Catherine Dempsey et al |
| **hCoV-19/Ireland/D-BH-270027240/2024** | EPI_ISL_18870258 | | Microbiology Beaumont Hospital | Microbiology Beaumont Hospital | Aisling Richmond et al |
| **hCoV-19/Kuwait/Jaber-20804427/2023** | EPI_ISL_18699717 | | Public Health | Virology Lab, Jaber Al Ahmad Hospital | Dr. Zahrah Buhamad et al |
| **hCoV-19/Pakistan/NIH-B164-S3/2023** | EPI_ISL_18724709 | | Department of Virology, Armed Forces Institute of Pathology (AFIP) | Department of Virology, National Institute of Health, Islamabad, Pakistan | Massab Umair et al |
| **hCoV-19/India/TG-CDFD-GMC-1114/2023** | EPI_ISL_18884624 | | Gandhi Medical College | CDFD | Asmita Gupta et al |
| **hCoV-19/India/CG-AIIMSR-VRDL-INSACOG-1945/2024** | EPI_ISL_18745589 | | State Virus Research and Diagnostic Laboratory (VRDL), Department of Microbiology, All India Institute of Medical Sciences | State Virus Research and Diagnostic Laboratory (VRDL), Department of Microbiology, All India Institute of Medical Sciences | Pushpendra Singh et al |
| **hCoV-19/Denmark/DCGC-665513/2023** | EPI_ISL_18617339 | | Department of Bacteria, Parasites and Fungi, Statens Serum Institut, Copenhagen, Denmark | Statens Serum Institut Bioinformatics and Microbial Genomics | Povilas Matusevicius on behalf of the Danish Covid-19 Genome Consortium et al |
| **hCoV-19/DominicanRepublic/2172584-LNSPDD/2024** | EPI_ISL_18864410 | | Laboratorio Nacional de Salud Pública Dr. Defilló | Laboratorio Nacional de Salud Pública Dr. Defilló | Isaac Miguel Sánchez et al |
| **hCoV-19/DominicanRepublic/2172611-LNSPDD/2024** | EPI_ISL_18864409 | | Laboratorio Nacional de Salud Pública Dr. Defilló | Laboratorio Nacional de Salud Pública Dr. Defilló | Isaac Miguel Sánchez et al |
| **hCoV-19/DominicanRepublic/2169643-LNSPDD/2024** | EPI_ISL_18794951 | | Laboratorio Nacional de Salud Pública Dr. Defilló | Laboratorio Nacional de Salud Pública Dr. Defilló | Isaac Miguel Sánchez et al |
| **hCoV-19/Malaysia/C19UMB4684/2024** | EPI_ISL_18935691 | | Hospital Melaka | UKM Medical Molecular Biology Institute (UMBI) | Rahman Jamal et al |
| **hCoV-19/Indonesia/JK-NIHRD-W03919/2023** | EPI_ISL_18673512 | | RSUP Persahabatan | National Institute of Health Research and Development | Arie Ardiansyah Nugraha et al |
| **hCoV-19/Curacao/CW-RIVM-136991/2023** | EPI_ISL_18824221 | | National Institute for Public Health and the Environment (RIVM) | National Institute for Public Health and the Environment (RIVM) | Adam Meijer et al |
| **hCoV-19/Shanghai/SCDC-HG1220-4244/2023** | EPI_ISL_18769894 | | Microbe lab, Shanghai Municipal Center for Disease Control & Prevention | Microbe lab, Shanghai Municipal Center for Disease Control & Prevention | Chen et al |
| **hCoV-19/Anhui/BZCDC-222/2024** | EPI_ISL_18941273 | | Clinical laboratory, Bozhou Municipal Center for Disease Control & Prevention | Clinical laboratory, Bozhou Municipal Center for Disease Control & Prevention | Qingqiao et al |
| **hCoV-19/Guadeloupe/IPP03571/2024** | EPI_ISL_18945128 | | Outre Mer | National Reference Center for Viruses of Respiratory Infections, Institut Pasteur, Paris | Léa Avon et al |
| **hCoV-19/Chile/BI-11724/2024** | EPI_ISL_18938110 | | Genetica Molecular and Subdepartamento de Virologia ISP Chile | Instituto de Salud Publica de Chile | Mariela Guajardo et al |
| **hCoV-19/Japan/PG-591428/2024** | EPI_ISL_18884282 | | Chiba Prefectural Institute of Public Health | Division of Virology and Zoology, Chiba Prefectural Institute of Public Health | Yuki Fujinuma et al |
| **hCoV-19/Hubei/HCDC-SY3742/2024** | EPI_ISL_18916141 | | Shiyan Center for Disease Control and Prevention | Hubei Provincial Center for Disease Control and Prevention Center of Pathogenic Microorganism Detection | Yongzhong Jiang et al |
| **hCoV-19/NewZealand/24YA1384/2024** | EPI_ISL_18923639 | | Medlab Central (PN) | Institute of Environmental Science and Research (ESR) | Sarah Cockerton et al |
| **hCoV-19/Portugal/PT50569/2024** | EPI_ISL_18880918 | | H Santarem | Instituto Nacional de Saude Doutor Ricardo Jorge (INSA) | Borges et al |
| **hCoV-19/Switzerland/TI-EOC-22_28631394/2024** | EPI_ISL_18833867 | | Laboratorio di Microbiologia | Laboratorio di Microbiologia, Istituto di Medicina di Laboratorio EOLAB, Ente Ospedaliero Cantonale | Martinetti Lucchini Gladys et al |
| **hCoV-19/England/CLIMB-CM7YR8T6/2023** | EPI_ISL_18763846 | | Respiratory Virus Unit, Microbiology Services Colindale, Public Health England | UK Health Security Agency (UKHSA) | Angie Lackenby on behalf of PHE Covid Sequencing Team et al |
| **hCoV-19/Bonaire/BQ-RIVM-136349/2023** | EPI_ISL_18801708 | | National Institute for Public Health and the Environment (RIVM) | National Institute for Public Health and the Environment (RIVM) | Adam Meijer et al |
| **hCoV-19/SouthKorea/KDCA283064/2024** | EPI_ISL_18823842 | | Division of Emerging Infectious Diseases, Bureau of Infectious Diseases Diagnosis Control, Korea Disease Control and Prevention Agency | Division of Emerging Infectious Diseases, Bureau of Infectious Diseases Diagnosis Control, Korea Disease Control and Prevention Agency | Jeong-Ah Kim et al |
| **hCoV-19/Australia/TAS006718/2023** | EPI_ISL_18744346 | | Royal Hobart Hospital | Royal Hobart Hospital | J Williamson et al |
| **hCoV-19/France/BRE-IPP03484/2024** | EPI_ISL_18945099 | | Labo Analyses Med | National Reference Center for Viruses of Respiratory Infections, Institut Pasteur, Paris | Léa Avon et al |
| **hCoV-19/Brazil/MS-FIOCRUZ-FMS153/2024** | EPI_ISL_18853750 | | LACEN/MS | Fiocruz Mato Grosso do Sul on behalf of Fiocruz Genomic Surveillance Network | Zoraida del Carmen Fernandez Grillo et al |
| **hCoV-19/NewZealand/24YA1093/2024** | EPI_ISL_18923801 | | CHL Hastings | Institute of Environmental Science and Research (ESR) | Sarah Cockerton et al |
| **hCoV-19/Austria/AGES-AZ-1679757/2024** | EPI_ISL_18789989 | | Department f. Virologie d. Med. Universität Wien | AGES IMED Vienna | Posselt S. et al |
| **hCoV-19/Belgium/Sciensano-LS-S0975/2024** | EPI_ISL_18918679 | | AZ Sint-Jan Brugge-Oostende AV | Sciensano, Viral Diseases Laboratory | François Dufrasne et al |
| **hCoV-19/Oman/CPHL_7241053/2024** | EPI_ISL_18931750 | | Central Public Health Laboratories-Sultanat of Oman | Central Public Health Laboratories-Sultanat of Oman | Samiha Al Kharusi et al |
| **hCoV-19/USA/HI-TAMC_3352/2024** | EPI_ISL_18941894 | | Tripler Army Medical Center | Tripler Army Medical Center, Dept of Clinical Investigation, | Keith S. K. Fong et al |
| **hCoV-19/India/DL-ILBS-WGS6703/2022** | EPI_ISL_14584566 | | STAR IMAGING & PATH LAB | ILBS/INSACOG | Ekta Gupta et al |
| **hCoV-19/India/GJ-INSACOG-GBRC8719/2022** | EPI_ISL_15058292 | | In Vitro Speciality Laboratory, Ahmedabad | Gujarat Biotechnology Research Centre | Priyank Chavda et al |
| **hCoV-19/India/OR-ILSGS19034/2022** | EPI_ISL_14601929 | | Nayagarh, DHH | Institute of Life Sciences-INSACOG | Sunil K. Raghav et al |
| **hCoV-19/India/MZ-1526101101465/2022** | EPI_ISL_16183119 | | INSACOG-Mizoram | National Institute of Biomedical Genomics - INSACOG | Arindam Maitra et al A |
| **hCoV-19/Bhutan/RCDC0169/2022** | EPI_ISL_17967089 | | Royal Centre for Disease Control | Royal Centre for Disease Control | Sonam Gyeltshen et al |
| **hCoV-19/India/TN-ICMR-SPHL-2035/2022** | EPI_ISL_16241401 | | State Public Health Laboratory,Chennai -TN - INSACOG | State Public Health Laboratory,Chennai -TN - INSACOG | Selvavinayagam Sivaprakasam et al |
| **hCoV-19/India/HR-INSACOG-NII-882/2022** | EPI_ISL_13521499 | | RTPCR Lab Panchkula | National Institute of Immunology | Dr. G. Aneeshkumar Arimbasseri et al |
| **hCoV-19/India/PB-VRDLGMCPTA-ICMR-INSACOG-GS0000261/2022** | EPI_ISL_14410225 | | Virus Research and Diagnostic Laboratory | NIV Influenza | Dr.Rupinder Bakshi et al |
| **hCoV-19/India/MP-AIIMS_B-ICMR-INSACOG-WGS-1464/2022** | EPI_ISL_15063749 | | AIIMS Bhopal VRDL/Translational Medicine - INSACOG | AIIMS Bhopal VRDL/Translational Medicine - INSACOG | Anvita G. Malhotra et al |
| **hCoV-19/India/MP-AIIMS_B-ICMR-INSACOG-WGS-1185/2022** | EPI_ISL_14842320 | | AIIMS Bhopal VRDL / Translational Medicine - INSACOG | AIIMS Bhopal VRDL/Translational Medicine - INSACOG | Anvita G. Malhotra et al |
| **hCoV-19/India/MG-PML-RFCH07398_SVGS519/2022** | EPI_ISL_18050773 | | Military Hospital | Pasteur Molecular Laboratory | Phibangipan Warjri et al |
| **hCoV-19/India/HP-INSACOG/NII-1104/2022** | EPI_ISL_13964686 | | Pt JLNGMCH Chamba | National Institute of Immunology | Aneeshkumar Arimbasseri et al |
| **hCoV-19/India/WB-INSACOG-1931300706784/2022** | EPI_ISL_14891496 | | INSACOG-WB | National Institute of Biomedical Genomics – INSACOG | Arindam Maitra et al A |
| **hCoV-19/India/TG-CDFD-GH-740/2022** | EPI_ISL_14445936 | | Gandhi Medical College | CDFD-INSACOG | Asmita Gupta et al |
| **hCoV-19/India/SK-INSACOG-753695834/2022** | EPI_ISL_14303969 | | INSACOG-Sikkim | National Institute of Biomedical Genomics – INSACOG | Arindam Maitra et al A |
| **hCoV-19/India/TR-AGMC/SO1/2022** | EPI_ISL_16183154 | | INSACOG-Tripura | National Institute of Biomedical Genomics - INSACOG | Arindam Maitra et al A |
| **hCoV-19/India/WB-INSACOG-1931503297345/2022** | EPI_ISL_14891590 | | INSACOG-WB | National Institute of Biomedical Genomics – INSACOG | Arindam Maitra et al A |
| **hCoV-19/Chile/AT-123754/2022** | EPI_ISL_15792458 | | Genetica Molecular and Subdepartamento de Virologia ISP Chile | Instituto de Salud Publica de Chile | Constanza Campano et al |
| **hCoV-19/Vietnam/OUCRU-S12-M160/2022** | EPI_ISL_17198132 | | Oxford University Clinical Research Unt | Oxford University Clinical Research Unt | Nguyen Van Vinh Chau et al |
| **hCoV-19/Vietnam/31159/2022** | EPI_ISL_15937765 | | Oxford University Clinical Research Unt | Oxford University Clinical Research Unt | Nguyen Van Vinh Chau et al |
| **hCoV-19/India/TG-CDFD-SR-178/2022** | EPI_ISL_15153405 | | Apollo Hospital | CDFD-INSACOG | Asmita Gupta et al |
| **hCoV-19/India/SK-1122500139469/2022** | EPI_ISL_16183052 | | INSACOG-Sikkim | National Institute of Biomedical Genomics - INSACOG | Arindam Maitra et al A |
| **hCoV-19/India/un-SPHL-1110/2022** | EPI_ISL_17757732 | | Public Health, State Public Health Laboratory Chennai | Public Health, State Public Health Laboratory Chennai | Raju et al |
| **hCoV-19/India/CG-AIIMSR-VRDL-INSACOG-0814/2022** | EPI_ISL_14668613 | | State Virus Research and Diagnostic Laboratory (VRDL), AIIMS Raipur | State Virus Research and Diagnostic Laboratory (VRDL), AIIMS Raipur | Pushpendra Singh et al |
| **hCoV-19/India/JK-SK-1033466/2022** | EPI_ISL_17448659 | | SKIMS-VRDL, Srinagar | NIV Influenza | Dr. Bashir Ahmad Fomda et al |
| **hCoV-19/Nepal/NPHL-S-1902/2022** | EPI_ISL_15430107 | | Kailali Hospital Pvt. Ltd. | National Public Health Laboratory | National Public Health Laboratory Team/Dhangadi Hospital et al |
| **hCoV-19/India/RJ-SMS-ICMR-INSACOG-TS-16647/2022** | EPI_ISL_15308482 | | SMS MEDICAL COLLEGE,JAIPUR | SMS MEDICAL COLLEGE,JAIPUR | Bharti Malhotra et al |
| **hCoV-19/India/RJ-INSACOG-AIIMSJ-213849/2022** | EPI_ISL_15309259 | | Regional Level Viral Research and Diagnostic Laboratory | Regional Level Viral Research and Diagnostic Laboratory | Dr. NAVEEN PRAKASH BOKOLIA et al |
| **hCoV-19/India/RJ-SMS-ICMR-INSACOG-US-39/2023** | EPI_ISL_17661874 | | SMS MEDICAL COLLEGE,JAIPUR | SMS MEDICAL COLLEGE,JAIPUR | Bharti Malhotra et al |
| **hCoV-19/India/RJ-SMS-ICMR-INSACOG-US-32/2023** | EPI_ISL_17661869 | | SMS MEDICAL COLLEGE,JAIPUR | SMS MEDICAL COLLEGE,JAIPUR | Bharti Malhotra et al |
| **hCoV-19/Bahrain/CDL-23020065558/2023** | EPI_ISL_17081404 | | Communicable Disease Laboratory | Communicable Disease Laboratory | Marhoon et al |
| **hCoV-19/USA/CO-CDPHE-2103748693/2022** | EPI_ISL_16606224 | | Colorado Department of Public Health and Environment | Colorado Department of Public Health and Environment | Laura Bankers et al |
| **hCoV-19/Germany/NW-RKI-I-1093532/2023** | EPI_ISL_16696402 | | MVZ Labor Krone GbR | Robert Koch Institute | ? |
| **hCoV-19/Romania/MS-28025/2022** | EPI_ISL_17540805 | | Department of Public Health Mures | National Institute of Infectious Diseases-Prof. Dr. Matei Bals Molecular Diagnostics Laboratory | Corina Casangiu et al |
| **hCoV-19/Thailand/DMSc-15048/2022** | EPI_ISL_17186406 | | Regional Medical Sciences Center 6 Chonburi | Medical Genomic Centre,Medical Life Sciences Institute,Department of Medical Sciences, Ministry of Public Health, Thailand | Surakameth Mahasirimongkol et al |
| **hCoV-19/Laos/IPL-77936/2023** | EPI_ISL_17670083 | | NCLE | Institut Pasteur du Laos | Thonglakhone XAYBOUNSOU et al |
| **hCoV-19/England/QEUH-32654F2F/2022** | EPI_ISL_16406139 | | Lighthouse Lab in Glasgow | Wellcome Sanger Institute for the COVID-19 Genomics UK (COG-UK) Consortium | Harper VanSteenhouse et al |
| **hCoV-19/Singapore/14835/2022** | EPI_ISL_15831955 | | National Public Health Laboratory, National Centre for Infectious Diseases | National Public Health Laboratory, National Centre for Infectious Diseases | Benny Yeo et al |
| **hCoV-19/Guangdong/GDCDC-030/2023** | EPI_ISL_17489850 | | Guangdong Provincial Center for Disease Control and Prevention | Guangdong Provincial Center for Disease Control and Prevention | Zhang et al B |
| **hCoV-19/Vietnam/BMH-507300/2022** | EPI_ISL_16073623 | | Bachmai Hospital | Bachmai Hospital | Phuong Truong et al |
| **hCoV-19/Vietnam/OUCRU-S43-M160/2022** | EPI_ISL_17198157 | | Oxford University Clinical Research Unt | Oxford University Clinical Research Unt | Nguyen Van Vinh Chau et al |
| **hCoV-19/Cambodia/275684/2022** | EPI_ISL_16272800 | | National Institute of Public Health | National Institute of Public Health | Chhorvann CHHEA et al |
| **hCoV-19/Thailand/NIH-16-1350-4-02/2023** | EPI_ISL_17597616 | | Thatphanom Crown Prince Hospital | National Institute of Health of Thailand (Thai-NIH) | Sumonmal Uttayamakul et al |
| **hCoV-19/Taiwan/CDC-F78504/2023** | EPI_ISL_16996282 | | Centers for Disease Control, R.O.C. (Taiwan) | Centers for Disease Control, R.O.C. (Taiwan) | Yu-Chi Lin et al |
| **hCoV-19/India/UT-INSACOG-GBRC9028/2022** | EPI_ISL_15309356 | | Supratech Micropath Diagnostics & Research laborat | Gujarat Biotechnology Research Centre | Jinal M. Thakor et al |
| **hCoV-19/India/MH-RFIP15641/2022** | EPI_ISL_15440583 | | BJMC | RF-IISER Pune | Krishanpal Karmodiya et al |
| **hCoV-19/Cambodia/939879/2022** | EPI_ISL_16230726 | | Virology Unit, Institut Pasteur du Cambodge | Virology Unit, Institut Pasteur du Cambodge | Leakhena Pum et al |
| **hCoV-19/Romania/DJ-543454/2022** | EPI_ISL_16941604 | | Clinical Emergency County Hospital, Craiova | Cantacuzino National Military-Medical Institute, Viral Respiratory Infections Laboratory | Catalina Pascu et al |
| **hCoV-19/USA/AK-PHL24045/2022** | EPI_ISL_16909717 | | Alaska State Virology Laboratory | Alaska State Virology Laboratory | Lisa Smith et al |
| **hCoV-19/Bahrain/CDL-22350085522/2022** | EPI_ISL_17081831 | | Communicable Disease Laboratory | Communicable Disease Laboratory | Marhoon et al |
| **hCoV-19/Taiwan/CDC-F79105/2023** | EPI_ISL_17308730 | | Centers for Disease Control, R.O.C. (Taiwan) | Centers for Disease Control, R.O.C. (Taiwan) | Yu-Chi Lin et al |
| **hCoV-19/Thailand/NIC_KKN_749/2023** | EPI_ISL_17062169 | | Regional Medical Sciences Center 7 Khon Kaen | National Institute of Health, Department of Medical Sciences, Ministry of Public Health, Thailand | Pilailuk Okada et al |
| **hCoV-19/Mauritius/N50927/2022** | EPI_ISL_16412678 | | CENTRAL HEALTH LABORATORY | National Institute for Communicable Diseases of the National Health Laboratory Service | Magalutcheemee R et al |
| **hCoV-19/Taiwan/CDC-F78625/2023** | EPI_ISL_17079914 | | Centers for Disease Control, R.O.C. (Taiwan) | Centers for Disease Control, R.O.C. (Taiwan) | Yu-Chi Lin et al |
| **hCoV-19/HongKong/HKU-750/2022** | EPI_ISL_16348220 | | Department of Microbiology, The University of Hong Kong | Department of Microbiology, The University of Hong Kong | Kelvin K.W. To et al |
| **hCoV-19/HongKong/HKU-230518-406/2023** | EPI_ISL_17685305 | | Department of Microbiology, The University of Hong Kong | Department of Microbiology, The University of Hong Kong | Kelvin K.W. To et al |
| **hCoV-19/Malaysia/TIDREC-NSC2594/2023** | EPI_ISL_17544334 | | Hospital Tuanku Fauziah, Perlis | Tropical Infectious Diseases Research & Education Centre (TIDREC), Universiti Malaya | Kim-Kee Tan et al |
| **hCoV-19/Cambodia/295891/2023** | EPI_ISL_17597722 | | National Institute of Public Health | National Institute of Public Health | Chhorvann CHHEA et al |
| **hCoV-19/SouthKorea/KDCA218306/2023** | EPI_ISL_17172022 | | Division of Emerging Infectious Diseases, Bureau of Infectious Diseases Diagnosis Control, Korea Disease Control and Prevention Agency | Division of Emerging Infectious Diseases, Bureau of Infectious Diseases Diagnosis Control, Korea Disease Control and Prevention Agency | Jeong-Ah Kim et al |
| **hCoV-19/Chongqing/YBCDC-0679/2023** | EPI_ISL_17858254 | | Chongqing Yubei District Center for Disease Control and Prevention | Chongqing Yubei District Center for Disease Control and Prevention | Jin Yan et al |
| **hCoV-19/Liaoning/LNCDC-190-3/2023** | EPI_ISL_17850718 | | Institute for Communicable and Infectious Diseases Prevention and Control, Liaoning Center for Disease Control and Prevention | Institute for Communicable and Infectious Diseases Prevention and Control, Liaoning Center for Disease Control and Prevention | Mao et al |
| **hCoV-19/Shanghai/SCDC-599/2023** | EPI_ISL_17994458 | | Microbe lab, Shanghai Municipal Center for Disease Control & Prevention | Microbe lab, Shanghai Municipal Center for Disease Control & Prevention | Zhang et al B |
| **hCoV-19/Liaoning/LNCDC-222-2/2023** | EPI_ISL_17972879 | | Institute for Communicable and Infectious Diseases Prevention and Control, Liaoning Center for Disease Control and Prevention | Institute for Communicable and Infectious Diseases Prevention and Control, Liaoning Center for Disease Control and Prevention | Mao et al |
| **hCoV-19/India/MP-AIIMS_B-ICMR-1808/2022** | EPI_ISL_15961902 | | AIIMS Bhopal VRDL/Translational Medicine - INSACOG | AIIMS Bhopal VRDL/Translational Medicine - INSACOG | Anvita G. Malhotra et al |
| **hCoV-19/Kuwait/batch2-Sample72/2022** | EPI_ISL_16201335 | | Jaber Hospital, MOH, Kuwait | Genetics & Bioinformatics Dept., Dasman Diabetes Insitute | Fahd Al-Mulla et al |
| **hCoV-19/Pakistan/IDRL-AKU-KHI-79242-S8/2022** | EPI_ISL_16362569 | | "Clinical Lab, Aga Khan University Hospital" | "Infectious Diseases Research Laboratory, Department of Pediatrics and Child Health" | "Samiah Kanwar et al |
| **hCoV-19/Vietnam/PIHCM_3379/2022** | EPI_ISL_16342386 | | Tay Ninh | Pasteur Institute in Ho Chi Minh city | Hieu Minh Le et al |
| **hCoV-19/India/MZ-1572800055876/2022** | EPI_ISL_16183187 | | INSACOG-Mizoram | National Institute of Biomedical Genomics - INSACOG | Arindam Maitra et al A |
| **hCoV-19/India/un-SPHL-1048/2022** | EPI_ISL_17757725 | | Public Health, State Public Health Laboratory Chennai | Public Health, State Public Health Laboratory Chennai | Raju et al |
| **hCoV-19/India/RJ-SMS-ICMR-INSACOG-TS-16539/2022** | EPI_ISL_15308433 | | SMS MEDICAL COLLEGE,JAIPUR | SMS MEDICAL COLLEGE,JAIPUR | Bharti Malhotra et al |
| **hCoV-19/India/SK-306555/2022** | EPI_ISL_16182838 | | INSACOG-Sikkim | National Institute of Biomedical Genomics - INSACOG | Arindam Maitra et al A |
| **hCoV-19/India/AP-CCMB-CID4716/2022** | EPI_ISL_15933734 | | AP SSO | CSIR-Centre for Cellular and Molecular Biology-INSACOG | Sumedha Avadhanula et al |
| **hCoV-19/Nepal/NPHL-S-1832/2022** | EPI_ISL_15313028 | | Shree Birendra Hospital | National Public Health Laboratory | National Public Health Laboratory Team/Shree Birendra Hospital et al |
| **hCoV-19/Bhutan/RCDC-0018/2022** | EPI_ISL_16984439 | | AFRIMS | AFRIMS | Chonticha Klungthong et al |
| **hCoV-19/Niue/ESR-YA1227/2023** | EPI_ISL_18284539 | | Niue Health Department | Institute of Environmental Science and Research (ESR) | Edgar Akauola et al |
| **hCoV-19/Australia/NSW-ICPMR-42967/2023** | EPI_ISL_17173335 | | Histopath | NSW Health Pathology - Institute of Clinical Pathology and Medical Research; Westmead Hospital; University of Sydney | Arnott A. et al |
| **hCoV-19/India/WB-1930302170257/2023** | EPI_ISL_17349761 | | School of Tropical Medicine | National Institute of Biomedical Genomics | Arindam Maitra et al A |
| **hCoV-19/Bhutan/RCDC-0026/2023** | EPI_ISL_16984447 | | AFRIMS | AFRIMS | Chonticha Klungthong et al |
| **hCoV-19/Kuwait/B3-S63/2022** | EPI_ISL_16436880 | | Jaber Hospital, MOH, Kuwait | Genetics & Bioinformatics Dept., Dasman Diabetes Insitute | Fahd Al-Mulla et al |
| **hCoV-19/India/un-SPHL-1020/2022** | EPI_ISL_17757716 | | Public Health, State Public Health Laboratory Chennai | Public Health, State Public Health Laboratory Chennai | Raju et al |
| **hCoV-19/Kuwait/Jaber-20442948/2022** | EPI_ISL_15983213 | | I'm Negative Laboratory | Virology Lab, Jaber Al Ahmad Hospital | Dr. Zahrah Buhamad et al |
| **hCoV-19/Mauritius/N50775/2022** | EPI_ISL_16412550 | | CENTRAL HEALTH LABORATORY | National Institute for Communicable Diseases of the National Health Laboratory Service | Magalutcheemee R et al |
| **hCoV-19/India/GJ-INSACOG-GBRC8931/2022** | EPI_ISL_15309437 | | AMC MET Medical College, Ahmedabad (LG Hospital) | Gujarat Biotechnology Research Centre | Chaitanya G. Joshi et al |
| **hCoV-19/India/SK-306196/2022** | EPI_ISL_16183008 | | INSACOG-Sikkim | National Institute of Biomedical Genomics - INSACOG | Arindam Maitra et al A |
| **hCoV-19/SriLanka/aicbu6450/2023** | EPI_ISL_17699456 | | Allergy, Immunology and Cell Biology Unit (AICBU) | Allergy, Immunology and Cell Biology Unit (AICBU) | Chandima Jeewandara et al |
| **hCoV-19/Israel/SMC-7114244/2023** | EPI_ISL_18062580 | | Shamir Medical Center (Asaf Harofe) | Shamir Medical Center (Asaf Harofe) | Nir Rainy et al |
| **hCoV-19/NewZealand/23YA6509/2023** | EPI_ISL_18561940 | | Medlab Central (PN) | Institute of Environmental Science and Research (ESR) | Sarah Cockerton et al |

| | | | | | |
|---|---|---|---|---|---|
| **hCoV-19/NewZealand/23ZA6963/2023** | EPI_ISL_18670840 | | Medlab Central (PN) | Institute of Environmental Science and Research (ESR) | Sarah Cockerton et al |
| **hCoV-19/NewZealand/ESR-YA3166/2023** | EPI_ISL_17962625 | | Medlab South | Institute of Environmental Science and Research (ESR) | Sarah Cockerton et al |
| **hCoV-19/Portugal/PT48510/2023** | EPI_ISL_17954728 | | CHU Sao Joao, Porto | Instituto Nacional de Saude Doutor Ricardo Jorge (INSA) | Borges et al |
| **hCoV-19/Turkey/HSGM-AHSL-16/2023** | EPI_ISL_18422108 | | Ministry of Health Turkey, Adana Public Health Lab. | Ministry of Health Türkiye, Public Health Directorate, National Virology Reference Laboratory | Süleyman Yalcin et al |
| **hCoV-19/HongKong/VK23009699/2023** | EPI_ISL_17433589 | | Tseung Kwan O Hospital | Hong Kong Department of Health | Alan K.L. Tsang et al |
| **hCoV-19/Switzerland/ZH-UZH-IMV-3ba690ec/2023** | EPI_ISL_17075996 | | Stadtspital Waid | Institute of Medical Virology, University of Zurich | Verena Kufner et al |
| **hCoV-19/Yunnan/YCDC-12/2022** | EPI_ISL_16750664 | | Yunnan Center for Disease Control and Prevention, Department of Acute Infectious Diseases Control and Prevention | Yunnan Center for Disease Control and Prevention, Department of Acute Infectious Diseases Control and Prevention | Meiling Zhang et al |
| **hCoV-19/Greece/C2095/2022** | EPI_ISL_17391676 | | National Public Health Organization | National Public Health Organization | Kyriaki Tryfinopoulou et al |
| **hCoV-19/Poland/0086WSN2023_wsseol/2023** | EPI_ISL_16954816 | | WSSE w Olsztynie | Wojewodzka Stacja Sanitarno-Epidemiologiczna w Olsztynie, Laboratorium Badan Epidemiologiczno-Klinicznych | Sylwia Krzetowska et al |
| **hCoV-19/Lithuania/MB-01240164/2023** | EPI_ISL_16963639 | | Viesoji istaiga Jurininku sveikatos prieziuros centras | National Public Health Surveillance Laboratory | Lukas Zemaitis et al |
| **hCoV-19/Belgium/Briant-12304531601/2023** | EPI_ISL_17309495 | | Heilig hart Lier | Imelda Ziekenhuis | Johan Frans et al |
| **hCoV-19/HongKong/VB23015380/2023** | EPI_ISL_17187466 | | Queen Elizabeth Hospital | Hong Kong Department of Health | Alan K.L. Tsang et al |
| **hCoV-19/India/GA-ICMR-NIV-G-12871/2022** | EPI_ISL_16353082 | | ICMR-National Institute of Virology - INSACOG | NIV Influenza | Dr. Varsha Potdar et al |
| **hCoV-19/India/GA-ICMR-NIV-INSACOG-GSEQ-12545/2022** | EPI_ISL_15854754 | | ICMR-National Institute of Virology - INSACOG | NIV Influenza | Dr. Varsha Potdar et al |
| **hCoV-19/India/un-SPHL-1139/2022** | EPI_ISL_17757734 | | Public Health, State Public Health Laboratory Chennai | Public Health, State Public Health Laboratory Chennai | Raju et al |
| **hCoV-19/India/MH-KasturbaMCGM-ICMR-INSACOG-WG5304/2022** | EPI_ISL_15760045 | | Kasturba Hospital | GeSCOPE, Kasturba Hospital | Jayanthi Shastri et al |
| **hCoV-19/SouthAfrica/NICD-N51401/2022** | EPI_ISL_16338587 | | TAMBO MEMORIAL LABORATORY | National Institute for Communicable Diseases of the National Health Laboratory Service | Everatt J et al |
| **hCoV-19/Mozambique/INS-PCS0029779/2023** | EPI_ISL_17253469 | | Biotechnology and Genetics Laboratory - Instituto Nacional de Saúde | Biotechnology and Genetics Laboratory - Instituto Nacional de Saúde | Sofia Viegas et al |
| **hCoV-19/India/SK-784328232/2022** | EPI_ISL_17349773 | | STNM Hospital | National Institute of Biomedical Genomics | Arindam Maitra et al A |
| **hCoV-19/India/SK-1122500141402/2022** | EPI_ISL_17349748 | | STNM Hospital | National Institute of Biomedical Genomics | Arindam Maitra et al A |
| **hCoV-19/India/WB-1931503362743/2022** | EPI_ISL_16182889 | | INSACOG-WB | National Institute of Biomedical Genomics - INSACOG | Arindam Maitra et al A |
| **hCoV-19/India/SK-785579286/2022** | EPI_ISL_17349767 | | STNM Hospital | National Institute of Biomedical Genomics | Arindam Maitra et al A |
| **hCoV-19/India/WB-1931503358260/2022** | EPI_ISL_16182911 | | INSACOG-WB | National Institute of Biomedical Genomics - INSACOG | Arindam Maitra et al A |
| **hCoV-19/India/SK-1122500141151/2022** | EPI_ISL_16182863 | | INSACOG-Sikkim | National Institute of Biomedical Genomics - INSACOG | Arindam Maitra et al A |
| **hCoV-19/India/MH-KasturbaMCGM-ICMR-INSACOG-WG5292/2022** | EPI_ISL_15760033 | | Kasturba Hospital | GeSCOPE, Kasturba Hospital | Jayanthi Shastri et al |
| **hCoV-19/Philippines/PGCV_0047_4976/2022** | EPI_ISL_16995827 | | Corazon Locsin Montelibano Memorial Regional Hospital | PHILIPPINE GENOME CENTER VISAYAS | Dr. Cynthia P. Saloma et al |
| **hCoV-19/Philippines/PGCV_0038_3817/2022** | EPI_ISL_16019350 | | Western Visayas Medical Center Sub National Laboratory | PHILIPPINE GENOME CENTER VISAYAS | Dr. Cynthia P. Saloma et al |
| **hCoV-19/India/WB-1931503341822/2022** | EPI_ISL_15465686 | | INSACOG-WB | National Institute of Biomedical Genomics – INSACOG | Arindam Maitra et al A |
| **hCoV-19/India/NA-GS20116/2022** | EPI_ISL_16487004 | | ILS, Bhubaneswar | Institute of Life Sciences-INSACOG | Sunil K. Raghav et al |
| **hCoV-19/India/GJ-INSACOG-GBRC10041/2022** | EPI_ISL_16276872 | | Supratech Micropath Diagnostics & Research laborat | Gujarat Biotechnology Research Centre-INSACOG | Bhumika Patel et al |
| **hCoV-19/Malawi/10486/2022** | EPI_ISL_17812887 | | Karonga District Hospital Laboratory | National Genomic Sequencing Reference Laboratory | Kagoli M et al |
| **hCoV-19/Malawi/NGRL-10527/2022** | EPI_ISL_17443394 | | Karonga District Hospital Laboratory | National Genomics Reference Laboratory | Kagoli M et al |
| **hCoV-19/India/WB-1931503371100/2022** | EPI_ISL_16182992 | | INSACOG-WB | National Institute of Biomedical Genomics - INSACOG | Arindam Maitra et al A |
| **hCoV-19/Russia/BA-RII-MH125944S/2022** | EPI_ISL_16894663 | | Gemotest Medical Laboratory | WHO National Influenza Centre Russian Federation | Andrey Komissarov et al |
| **hCoV-19/India/TG-CDFD-SR-227/2022** | EPI_ISL_16158730 | | Apollo Hospital | CDFD-INSACOG | Asmita Gupta et al |
| **hCoV-19/India/un-ICMR-MCL-H-7062-11193/2022** | EPI_ISL_17422736 | | Indian Council of Medical Research-National Institute of Virology, Microbial Containment Complex | Indian Council of Medical Research-National Institute of Virology, Microbial Containment Complex | Yadav et al |
| **hCoV-19/India/un-ICMR-MCL-H-7062-11243/2022** | EPI_ISL_17422738 | | Indian Council of Medical Research-National Institute of Virology, Microbial Containment Complex | Indian Council of Medical Research-National Institute of Virology, Microbial Containment Complex | Yadav et al |
| **hCoV-19/India/MH_BJCOG_INSACOG_illumina_Run_8_5064/2023** | EPI_ISL_16946788 | | National Institute of Virology, Pune | Center for Genomics, Department of Microbiology, BJ Government Medical College and Sassoon Hospitals, Pune - 411001, Maharashtra, India | Dr Rajesh Karyakarte et al |
| **hCoV-19/Thailand/NIC_UBN_689/2023** | EPI_ISL_17017003 | | Regional Medical Sciences Center 10 Ubonratchathani | Department of Medical Sciences, National Institute of Health, Ministry of Public Health, Thailand | Pilailuk Okada et al |
| **hCoV-19/Ecuador/USFQ-4803/2023** | EPI_ISL_17250784 | | Institute of Microbiology, Universidad San Francisco de Quito | Institute of Microbiology, Universidad San Francisco de Quito | Mateo Carvajal et al |
| **hCoV-19/Ecuador/USFQ-4725/2023** | EPI_ISL_16609173 | | Institute of Microbiology, Universidad San Francisco de Quito | Institute of Microbiology, Universidad San Francisco de Quito | Rommel Guevara et al |
| **hCoV-19/India/KA-SLS-53028/2023** | EPI_ISL_16939623 | | Strand Life Sciences [supported by IN COVID Support FZE LLC (Blockchain For Impact)] | Strand Life Sciences [supported by IN COVID Support FZE LLC (Blockchain For Impact)] | Harsha Pulleri Kandi et al |
| **hCoV-19/India/MH-ICMR-NIV-G-14649/2023** | EPI_ISL_18209948 | | ICMR-National Institute of Virology - INSACOG | NIV Influenza | Dr. Varsha Potdar et al |
| **hCoV-19/SouthAfrica/NICD-N55591/2023** | EPI_ISL_17859317 | | AMPATH | National Institute for Communicable Diseases of the National Health Laboratory Service | Everatt J et al |
| **hCoV-19/India/GJ-INSACOG-GBRC13544/2023** | EPI_ISL_18136388 | | Pathocare Pathology Laboratory, Vadodara | Gujarat Biotechnology Research Centre-INSACOG | Apurvasinh Puvar et al |
| **hCoV-19/Iceland/L-4225/2023** | EPI_ISL_17980584 | | Landspitali Department of Clinical Microbiology | Landspitali Department of Clinical Microbiology | Freyja Valsdottir et al |
| **hCoV-19/Brunei/MGS-7123066262/2023** | EPI_ISL_18484807 | | National Virology Reference Laboratory | Microbial Genomic Services | Zainun Zaini et al |
| **hCoV-19/India/GA-ICMR-NIV-G-13465/2023** | EPI_ISL_17445809 | | ICMR-National Institute of Virology - INSACOG | NIV Influenza | Dr. Varsha Potdar et al |
| **hCoV-19/India/GJ-GBRC10304/2023** | EPI_ISL_17190105 | | BJ Medical College, Ahmedabad | Gujarat Biotechnology Research Centre-INSACOG | Apurvasinh Puvar et al |
| **hCoV-19/India/CT-ILS-GS20510/2023** | EPI_ISL_17730180 | | Pt. JNM Medical College, Raipur | Institute of Life Sciences-INSACOG | Sunil K. Raghav et al |
| **hCoV-19/Singapore/R8NP15/2023** | EPI_ISL_17539918 | | National Public Health Laboratory, National Centre for Infectious Diseases | National Public Health Laboratory, National Centre for Infectious Diseases | Benny Yeo et al |
| **hCoV-19/India/MH-ICMR-NIV-G-14644/2023** | EPI_ISL_18209944 | | ICMR-National Institute of Virology - INSACOG | NIV Influenza | Dr. Varsha Potdar et al |
| **hCoV-19/Shanghai/SCDC-HG0802-2905-2/2023** | EPI_ISL_18105646 | | Microbe lab, Shanghai Municipal Center for Disease Control & Prevention | Microbe lab, Shanghai Municipal Center for Disease Control & Prevention | Zhang et al B |
| **hCoV-19/USA/MO-CDC-VSX-A065827/2023** | EPI_ISL_17550724 | | Helix | Centers for Disease Control and Prevention Division of Viral Diseases, Pathogen Discovery | Dakota Howard et al |
| **hCoV-19/Taiwan/CDC-F78794/2023** | EPI_ISL_18687198 | | Centers for Disease Control, R.O.C. (Taiwan) | Centers for Disease Control, R.O.C. (Taiwan) | Yu-Chi Lin et al |
| **hCoV-19/India/MH-INSACOG-RF-WT-NCL-5134/2023** | EPI_ISL_18740603 | | Health Accurate | NCL, Pune | Anu Raghunathan et al |
| **hCoV-19/Anhui/HNCDC-62/2023** | EPI_ISL_17801889 | | Microbiology lab, Huainan City Center for Disease Control and Prevention | Huainan City Center for Disease Control and Prevention | Liu Jiang et al |
| **hCoV-19/Poland/0752WSN2023_wsseol/2023** | EPI_ISL_18243835 | | WSSE w Bydgoszczy | Wojewodzka Stacja Sanitarno-Epidemiologiczna w Olsztynie, Laboratorium Badan Epidemiologiczno-Klinicznych | Sylwia Krzetowska et al |
| **hCoV-19/India/AS-DIB-1571/2023** | EPI_ISL_18414641 | | Regional VRDL ICMR-RMRC-NER Dibrugarh Assam India | Regional VRDL ICMR-RMRC-NER Dibrugarh Assam India | B Borkakoty et al |
| **hCoV-19/Argentina/INEI126008/2023** | EPI_ISL_18422387 | | Servicio Virosis Respiratorias-Departamento Virología-INEI | Instituto Nacional Enfermedades Infecciosas C.G.Malbran | Baumeister E. et al |
| **hCoV-19/India/JK-SGR-MAR-15/2023** | EPI_ISL_17554841 | | SKIMS-VRDL, Srinagar | NIV Influenza | Dr. Bashir Ahmad Fomda et al |
| **hCoV-19/Bahrain/PHD-0902/2023** | EPI_ISL_17744983 | | Communicable Disease Laboratory, Public Health Directorate | Communicable Disease Laboratory, Public Health Directorate | Marhoon et al |
| **hCoV-19/Honduras/FM-070/2023** | EPI_ISL_18107671 | | Laboratorio Nacional de Virologìa (SESAL) | Laboratorio Nacional de Virologìa (SESAL) | Dra. Soany Avilez et al |
| **hCoV-19/Kuwait/Jaber-20535806/2023** | EPI_ISL_18694824 | | Jahra Hospital | Virology Lab, Jaber Al Ahmad Hospital | Dr. Zahrah Buhamad et al |
| **hCoV-19/Malaysia/MKAI-6789517/2023** | EPI_ISL_18253917 | | Public Health Laboratory Ipoh | Disease Section, Public Health Laboratory Ipoh | MH. Chew et al |
| **hCoV-19/India/KA-21381/2023** | EPI_ISL_18340414 | | District Surveillance Unit | Institute For Stem Cell Science and Regenerative Medicine | Dasaradhi Palakodeti et al |
| **hCoV-19/Australia/VIC-VIDRL-00011987/2023** | EPI_ISL_18219016 | | Royal Melbourne Hospital - Microbiology Department | Victorian Infectious Disease Reference Laboratory | Aziz et al |
| **hCoV-19/Philippines/PH-VSMMC-7-245/2023** | EPI_ISL_18611971 | | Cebu City Medical Center | Vicente Sotto Memorial Medical Center | Reynette Christine J. Ligaray et al |
| **hCoV-19/Philippines/PH-VSMMC-7-365/2023** | EPI_ISL_18944500 | | VICENTE SOTTO MEMORIAL MEDICAL CENTER | VICENTE SOTTO MEMORIAL MEDICAL CENTER | Reynette Christine J. Ligaray et al |
| **hCoV-19/TrinidadandTobago/UWI-428416/2023** | EPI_ISL_18050339 | | Trinidad Public Health Laboratory | Carrington Lab, Department of Preclinical Sciences, Faculty of Medical Sciences, The University of the West Indies, St Augustine Campus | Nikita S. D. Sahadeo et al |
| **hCoV-19/Poland/WSSEGorzow-23S0346/2023** | EPI_ISL_18255402 | | WSSE w Poznaniu. Laboratorium Mikrobiologii i Parazytologii | Wojewodzka Stacja Sanitarno-Epidemiologiczna w Gorzowie Wielkopolskim | Elzbieta Justynska et al |
| **hCoV-19/ElSalvador/1986-LNSP/2023** | EPI_ISL_18615351 | | Hospital El Salvador San Salvador Ss | Laboratorio Nacional de Salud Publica. | Laboratorio Nacional de Salud Publica de El Salvador - Instituto Nacional de Salud et al |
| **hCoV-19/Kuwait/Jaber-20723908/2023** | EPI_ISL_18699294 | | Jahra Hospital | Virology Lab, Jaber Al Ahmad Hospital | Dr. Zahrah Buhamad et al |
| **hCoV-19/Singapore/R55MR61/2023** | EPI_ISL_18127430 | | National Public Health Laboratory, National Centre for Infectious Diseases | National Public Health Laboratory, National Centre for Infectious Diseases | Yichen Ding et al |
| **hCoV-19/Malaysia/C19UMB4232/2023** | EPI_ISL_18135449 | | Hospital Melaka | UKM Medical Molecular Biology Institute (UMBI) | Rahman Jamal et al |
| **hCoV-19/Tunisia/416/2023** | EPI_ISL_18227530 | | Laboratory of Microbiology,Farhat Hached Hospital | 1-Laboratory of Microbiology, National Reference Lab, Charles Nicolle Hospital; 2-University of Tunis ElManar, Faculty of Medicine of Tunis, LR99ES09, Tunis, Tunisia | Zaineb Hamzaoui et al |
| **hCoV-19/India/MH-ICMR-NIV-G-14699/2023** | EPI_ISL_18248367 | | ICMR-National Institute of Virology - INSACOG | NIV Influenza | Dr. Varsha Potdar et al |
| **hCoV-19/CzechRepublic/NRL_s2253/2023** | EPI_ISL_18612329 | | University Hospital in Pilsen | National Institute of Public Health | Helena Jirincova et al |
| **hCoV-19/Romania/219442/2023** | EPI_ISL_18932904 | | Cluj-Napoca Clinical Hospital for Infectious Diseases | National Institute of Public Health Laboratory | Mihaela Leustean et al |
| **hCoV-19/Georgia/Tb-s8138/2023** | EPI_ISL_18698445 | | Department for Virology, Molecular Biology and Genome Research, R. G. Lugar Center for Public Health Research, National Center for Disease Control and Public Health (NCDC) of Georgia. | Department for Virology, Molecular Biology and Genome Research, R. G. Lugar Center for Public Health Research, National Center for Disease Control and Public Health (NCDC) of Georgia. | Giorgi Tomashvili et al |
| **hCoV-19/India/KA-RFNB-20002/2023** | EPI_ISL_17720597 | | Apollo Health And Lifestyle Ltd | National Centre for Biological Sciences, TIFR - Rockefeller Foundation | Vinay Kumar et al |
| **hCoV-19/Brunei/MGS-7123058094/2023** | EPI_ISL_18272604 | | National Virology Reference Laboratory | Microbial Genomic Services | Zainun Zaini et al |
| **hCoV-19/Georgia/Tb-s8077/2023** | EPI_ISL_18413147 | | Department for Virology, Molecular Biology and Genome Research, R. G. Lugar Center for Public Health Research, National Center for Disease Control and Public Health (NCDC) of Georgia. | Department for Virology, Molecular Biology and Genome Research, R. G. Lugar Center for Public Health Research, National Center for Disease Control and Public Health (NCDC) of Georgia. | Giorgi Tomashvili et al |
| **hCoV-19/Oman/CPHL_52330354/2023** | EPI_ISL_18458154 | | Central Public Health Laboratories-Sultanat of Oman | Central Public Health Laboratories-Sultanat of Oman | Yathrib Al Zakwani et al |
| **hCoV-19/Montenegro/201369/2023** | EPI_ISL_18215545 | | Laboratory for Molecular Diagnostics,IPHMN | Department for Molecular Diagnostics, Centre for Medical Microbiology, Institute of Public Health of Montenegro | Danijela Vujošević et al |
| **hCoV-19/Bangladesh/SIRS_ideSHi_3230909009/2023** | EPI_ISL_18560370 | | International Centre for Diarrhoeal Disease Research, Bangladesh (icddr,b) | Institute for Developing Science and Health Initiatives (ideSHi) | Saikt Rahman et al |
| **hCoV-19/India/UP-ICMR-NIV-G-14616/2023** | EPI_ISL_18120313 | | Indian Council of Medical Research, National Institute of Virology | National Institute of Virology - Influenza | Dr. Varsha Potdar et al |
| **hCoV-19/Kenya/2022037566/2023** | EPI_ISL_18516321 | | DLSP KEMRI/CDC NAIROBI Lab | DLSP KEMRI/CDC NAIROBI Lab | Lydia Mwasi et al |
| **hCoV-19/USA/SD-SDPHL-3318/2024** | EPI_ISL_18886472 | | South Dakota Public Health Laboratory | South Dakota Public Health Laboratory | Chris Carlson et al |
| **hCoV-19/Kenya/HM27287/2023** | EPI_ISL_18788164 | | KEMRI-Wellcome Trust Research Programme,Kilifi | KEMRI-Wellcome Trust Research Programme,Kilifi | Lambisia A. et al |
| **hCoV-19/Bangladesh/icddrb_TND_05_1077/2022** | EPI_ISL_15801473 | | International Centre for Diarrhoeal Disease Research, Bangladesh (icddr,b) | Institute for Developing Science and Health Initiatives (ideSHi) | Habib T et al |
| **hCoV-19/SaudiArabia/KAUST603/2022** | EPI_ISL_17088920 | | King Faisal Specialist Hospital & Research Center | Pathogen Genomics Lab King Abdullah University of Science and Technology (KAUST) | Sabir A Adoub et al |
| **hCoV-19/India/MH-ICMR-NIV-INSACOG-NIVSARI-22-3098/2022** | EPI_ISL_15854988 | | ICMR-National Institute of Virology - INSACOG | NIV Influenza | Dr. Varsha Potdar et al |
| **hCoV-19/Bangladesh/icddrb-TND-06-1232/2022** | EPI_ISL_16528796 | | International Centre for Diarrhoeal Disease Research, Bangladesh (icddr,b) | Institute for Developing Science and Health Initiatives (ideSHi) | Rahman S et al |
| **hCoV-19/India/DL-ILBS7401/2022** | EPI_ISL_16549296 | | LIFELINE LABORATORY | ILBS/INSACOG | Ekta Gupta et al |
| **hCoV-19/Pakistan/NIH-B115-S7/2023** | EPI_ISL_16394564 | | Department of Virology, National Institute of Health, Islamabad, Pakistan | Department of Virology, National Institute of Health, Islamabad, Pakistan | Massab Umair et al |
| **hCoV-19/India/TR-AGMC/SO18/2022** | EPI_ISL_16182925 | | INSACOG-Tripura | National Institute of Biomedical Genomics - INSACOG | Arindam Maitra et al A |
| **hCoV-19/DominicanRepublic/2100538-LNSPDD/2022** | EPI_ISL_16022338 | | Laboratorio Nacional de Salud Pública Dr. Defilló | Laboratorio Nacional de Salud Pública Dr. Defilló | Isaac Miguel Sánchez et al |
| **hCoV-19/DominicanRepublic/2125820-LNSPDD/2023** | EPI_ISL_16533197 | | Laboratorio Nacional de Salud Pública Dr. Defilló | Laboratorio Nacional de Salud Pública Dr. Defilló | Isaac Miguel Sánchez et al |
| **hCoV-19/DRC/INRB-RDC-0102/2022** | EPI_ISL_17190902 | | Pathogen Genomics Lab, National Institute for Biomedical Research (INRB) | Pathogen Genomics Lab, National Institute for Biomedical Research (INRB) | Placide Mbala-Kingebeni et al B |
| **hCoV-19/India/WB-1931503372882/2022** | EPI_ISL_16183189 | | INSACOG-WB | National Institute of Biomedical Genomics - INSACOG | Arindam Maitra et al A |
| **hCoV-19/Bolivia/BG430/2023** | EPI_ISL_17818562 | | LABORATORIO BIOGENESIS S.R.L.(SANTA CRUZ/SCZ) | CENTRO NACIONAL DE ENFERMEDADES TROPICALES (CENETROP) | Loayza et al |
| **hCoV-19/Bolivia/BMD25/2023** | EPI_ISL_17818575 | | LABORATORIO "CORPLUSMEDIK SRL" (SANTA CRUZ/SCZ) | CENTRO NACIONAL DE ENFERMEDADES TROPICALES (CENETROP) | Loayza et al |
| **hCoV-19/India/MH_BJCOG_INSACOG_illumina_Run_8_5065/2023** | EPI_ISL_16946789 | | National Institute of Virology, Pune | Center for Genomics, Department of Microbiology, BJ Government Medical College and Sassoon Hospitals, Pune - 411001, Maharashtra, India | Dr Rajesh Karyakarte et al |
| **hCoV-19/Mexico/CAM-LANGEBIO_IMSS_14024/2023** | EPI_ISL_16968400 | | Unidad de Investigacion Medica de Yucatan | Unidad de Genomica Avanzada | Julio Elias Alvarado-Yaah et al |
| **hCoV-19/Russia/MOW-RII-MH127784S/2023** | EPI_ISL_17106201 | | Gemotest Medical Laboratory | WHO National Influenza Centre Russian Federation | Andrey Komissarov et al |
| **hCoV-19/Chongqing/YBCDC-0181/2022** | EPI_ISL_17197573 | | Chongqing Yubei District Center for Disease Control and Prevention | Chongqing Yubei District Center for Disease Control and Prevention | Jin Yan et al |
| **hCoV-19/India/DL-ILBS7436/2022** | EPI_ISL_16549249 | | LIFECELL INTERNATIONAL PVT LTD | ILBS/INSACOG | Ekta Gupta et al |
| **hCoV-19/India/RJ-SMS-ICMR-INSACOG-TS-17497/2022** | EPI_ISL_17661847 | | SMS MEDICAL COLLEGE,JAIPUR | SMS MEDICAL COLLEGE,JAIPUR | Bharti Malhotra et al |
| **hCoV-19/India/RJ-SMS-ICMR-TS-17308/2022** | EPI_ISL_17654502 | | SMS MEDICAL COLLEGE,JAIPUR | SMS MEDICAL COLLEGE,JAIPUR | Bharti Malhotra et al |
| **hCoV-19/Uganda/747/2022** | EPI_ISL_15909071 | | Central Public Health Laboratories | Central Public Health Laboratories | Alisen Ayitewala et al |
| **hCoV-19/Mexico/DUR_InDRE_S15877/2023** | EPI_ISL_16889452 | | Laboratorio Estatal de Salud Pública (LESP) Durango | Instituto de Diagnostico y Referencia Epidemiologicos (INDRE) | Claudia Wong-Arambula et al |
| **hCoV-19/Guatemala/526-LNS/2023** | EPI_ISL_16958546 | | Laboratorio Nacional de Salud (LNS), Guatemala | Laboratorio Nacional de Salud (LNS), Guatemala | Mendoza L et al |
| **hCoV-19/ElSalvador/58645-88-LNSP/2022** | EPI_ISL_18110620 | | Laboratorio Nacional de Salud Publica | Laboratorio Nacional de Salud Publica | Laboratorio de Secuenciacion y genomica del LNSP-INS et al |
| **hCoV-19/Bahrain/CDL-440105110/2022** | EPI_ISL_17081948 | | Communicable Disease Laboratory | Communicable Disease Laboratory | Marhoon et al |
| **hCoV-19/Myanmar/DMR-22109/2022** | EPI_ISL_17162784 | | Department of Medical Research | Department of Medical Research | Myat Htut Nyunt et al |
| **hCoV-19/Guatemala/9485-LNS/2022** | EPI_ISL_16207943 | | Laboratorio Nacional de Salud (LNS), Guatemala | Laboratorio Nacional de Salud (LNS), Guatemala | Mendoza L et al |
| **hCoV-19/Indonesia/JB-WJHL-ITB-W05359/2023** | EPI_ISL_17736548 | | RSUD Al Ihsan | West Java Health Laboratory; School of Life Sciences and Technology, Institut Teknologi Bandung | Azzania Fibriani et al |
| **hCoV-19/Malaysia/IMR_CV05369/2023** | EPI_ISL_16947128 | | KUALA LUMPUR HOSPITAL | Institute for Medical Research, Infectious Disease Research Centre, National Institutes of Health, Ministry of Health Malaysia | Kamel K et al |

| | | | | | |
|---|---|---|---|---|---|
| **hCoV-19/Panama/CDED24977-GMI/2023** | EPI_ISL_17977446 | | Centro de Diagnostico de Enfermedades Desatendidas | Gorgas Memorial Institute of Health Studies | Eduardo Aguirre et al |
| **hCoV-19/India/KA-ICMR-NIV-G-14579/2023** | EPI_ISL_18120284 | | Indian Council of Medical Research, National Institute of Virology | National Institute of Virology - Influenza | Dr. Varsha Potdar et al |
| **hCoV-19/India/HP-INSACOG-NII-1539/2023** | EPI_ISL_16649090 | | CRI Kasauli | National Institute of Immunology | Aneeshkumar Arimbasseri et al |
| **hCoV-19/India/GA-ICMR-NIV-G-12865/2022** | EPI_ISL_16353049 | | ICMR-National Institute of Virology - INSACOG | NIV Influenza | Dr. Varsha Potdar et al |
| **hCoV-19/Brunei/7723000012/2023** | EPI_ISL_17061017 | | National Virology Reference Laboratory, Department of Laboratory Services | Clinical Molecular Diagnostic Laboratory For Infectious Disease, Department of Laboratory Services, Microbial Genomic Services | Zainun Zaini et al |
| **hCoV-19/Guatemala/232-LNS/2023** | EPI_ISL_16735842 | | Laboratorio Nacional de Salud (LNS) | Laboratorio Nacional de Salud (LNS) | Mendoza L et al |
| **hCoV-19/Brunei/7122196596/2022** | EPI_ISL_16477593 | | National Virology Reference Laboratory | Microbial Genomic Services | Zainun Zaini et al |
| **hCoV-19/Iceland/L-4151/2023** | EPI_ISL_17716122 | | Landspitali Department of Clinical Microbiology | Landspitali Department of Clinical Microbiology | Freyja Valsdottir et al |
| **hCoV-19/Portugal/PT47136/2023** | EPI_ISL_17484151 | | CH Porto - H Sto Antonio | Instituto Nacional de Saude Doutor Ricardo Jorge (INSA) | Borges et al |
| **hCoV-19/Shanxi/TYCDC442/2023** | EPI_ISL_18495373 | | Microbiology Department, Taiyuan Center for Disease Control and Prevention | Microbiology Department, Taiyuan Center for Disease Control and Prevention | Wang et al |
| **hCoV-19/Shanxi/TYCDC382/2023** | EPI_ISL_18495313 | | Microbiology Department, Taiyuan Center for Disease Control and Prevention | Microbiology Department, Taiyuan Center for Disease Control and Prevention | Wang et al |
| **hCoV-19/Yunnan/YNCDC-0549/2023** | EPI_ISL_17683967 | | The department for prevention and control of acute infectious diseases, Yunnan Center for Disease Control and Prevention | The department for prevention and control of acute infectious diseases, Yunnan Center for Disease Control and Prevention | Zhang et al B |
| **hCoV-19/Shanghai/SCDC-126-2/2023** | EPI_ISL_17494198 | | Shanghai Municipal Center for Disease Control and Prevention | Shanghai Municipal Center for Disease Control and Prevention | Zhang et al B |
| **hCoV-19/Algeria/5414/2023** | EPI_ISL_18090026 | | NIC, Viral Respiratory Unit | NIC, Viral Respiratory Unit | Baya Hedjal et al |
| **hCoV-19/Mexico/CHP_IBT_IMSS_11503/2023** | EPI_ISL_18588976 | | Laboratorio Central de Epidemiologia (LCE) | Instituto de Biotecnologia de la UNAM | Julio Elias Alvarado-Yaah et al |
| **hCoV-19/Guinea/CERFIG-38439/2023** | EPI_ISL_17418960 | | Centre de recherche et de fomation en infectiologie de Guinee (CERFIG) | Centre de recherche et de fomation en infectiologie de Guinee (CERFIG) | Thibaut Armel Cherif GNIMADI et al |
| **hCoV-19/Italy/CAM-TIGEM-IZSM-COLLI-46602/2023** | EPI_ISL_17483082 | | Ospedale Santa Maria delle Grazie | The Telethon Institute of Genetics and Medicine (TIGEM) | Antonio Grimaldi Patrizia Annunziata Francesco Panariello Claudia Tiberio Tonya Fusco Valentina Bouche Chiara Colantuono Lucio Di Filippo Anna Manfredi Marcello Salvi Angela De Monaco Antonio Sorrentino Mariarosaria Granata Antonio Limone Luigi Atripaldi Andrea Ballabio Davide Cacchiarelli et al |
| **hCoV-19/DRC/INRB-DRC-0567/2023** | EPI_ISL_17976348 | | Laboratoire Rodolphe Mérieux (INRB Goma) | Pathogen Genomics Lab, National Institute for Biomedical Research (INRB) | Placide Mbala-Kingebeni et al B |
| **hCoV-19/Togo/TOG-GOL-6016/2023** | EPI_ISL_17831204 | | Laboratoire National de Référence (LNR), Centre Hospitalier Universitaire Sylvanus Olympio (CHU-SO) | Laboratoire BIOLIM / Faculté des Sciences de la Santé, Université de Lomé (FSS-UL) | A.A. Konou et al |
| **hCoV-19/Ghana/NMIMR-NCOV-23-042/2023** | EPI_ISL_17635315 | | Noguchi Memorial Institute for Medical Research, University of Ghana, Legon, Ghana | Noguchi Memorial Institute for Medical Research, University of Ghana, Legon, Ghana | Quaneeta Mohktar et al |
| **hCoV-19/Togo/TOG-GOL-6003/2023** | EPI_ISL_17830872 | | Laboratoire National de Référence (LNR), Centre Hospitalier Universitaire Sylvanus Olympio (CHU-SO) | Laboratoire BIOLIM / Faculté des Sciences de la Santé, Université de Lomé (FSS-UL) | A.A. Konou et al |
| **hCoV-19/Guinea/IPGui-28519/2023** | EPI_ISL_18050124 | | Institut Pasteur de Guinée | Institut Pasteur de Guinée | Reine ANGUINZE et al |
| **hCoV-19/Guinea/IPGui-28520/2023** | EPI_ISL_18050125 | | Institut Pasteur de Guinée | Institut Pasteur de Guinée | Reine ANGUINZE et al |
| **hCoV-19/Mali/CICM-28888/2023** | EPI_ISL_17474575 | | CICM, Bamako | Malaria Research and Training Center | Antoine Dara et al |
| **hCoV-19/BurkinaFaso/CHUSS_16949/2023** | EPI_ISL_17319108 | | Laboratoire de Microbiologie CHU Souro Sanou | Laboratoire de Microbiologie CHU Souro Sanou | Yacouba Sawadogo et al |
| **hCoV-19/BurkinaFaso/CHUSS_16955/2023** | EPI_ISL_17319107 | | Laboratoire de Microbiologie CHU Souro Sanou | Laboratoire de Microbiologie CHU Souro Sanou | Yacouba Sawadogo et al |
| **hCoV-19/Mali/CICM-28814/2023** | EPI_ISL_17474568 | | CICM, Bamako | Malaria Research and Training Center | Antoine Dara et al |
| **hCoV-19/Nigeria/NCDC-AFI-00126/2023** | EPI_ISL_18110459 | | National Reference Laboratory | National Reference Laboratory, Nigeria Centre for Disease Control | Dr Ndodo Nnaemeka et al |
| **hCoV-19/Togo/TOG-GOL-6032/2023** | EPI_ISL_17970742 | | Laboratoire National de Référence (LNR), Centre Hospitalier Universitaire Sylvanus Olympio (CHU-SO) | Laboratoire BIOLIM / Faculté des Sciences de la Santé, Université de Lomé (FSS-UL) | A.A. Konou et al |
| **hCoV-19/Nigeria/NCDC-NRL-GL-0287/2023** | EPI_ISL_18593990 | | FCT | National Reference Laboratory, Nigeria Centre for Disease Control and Prevention | Olusola Anuoluwapo Akanbi et al |
| **hCoV-19/Brunei/7123014310/2023** | EPI_ISL_17107272 | | National Virology Reference Laboratory, Departmenrt of Laboratory Services | Microbial Genomic Services, Clinical Molecular Diagnostic Laboratory For Infectious Disease, Department of Laboratory Services | Zainun Zaini et al |
| **hCoV-19/Brunei/7123010782/2023** | EPI_ISL_17090695 | | National Virology Reference Laboratory - Departmenrt of Laboratory Services | Clinical Molecular Diagnostic Laboratory For Infectious Disease - Microbial Genomic Services | Zainun Zaini et al |
| **hCoV-19/India/WB-1930302185947/2023** | EPI_ISL_18109187 | | INSACOG-WB | National Institute of Biomedical Genomics - INSACOG | Arindam Maitra et al A |
| **hCoV-19/Taiwan/CDC-F78859/2023** | EPI_ISL_18672882 | | Centers for Disease Control, R.O.C. (Taiwan) | Centers for Disease Control, R.O.C. (Taiwan) | Yu-Chi Lin et al |
| **hCoV-19/Iceland/L-4056/2023** | EPI_ISL_17529055 | | Landspitali Department of Clinical Microbiology | Landspitali Department of Clinical Microbiology | Freyja Valsdottir et al |
| **hCoV-19/Barbados/181555/2023** | EPI_ISL_18137631 | | Best-dos Santos Public Health Lab | The Caribbean Public Health Agency (CARPHA) | Chariesse Dos Santos on behalf of CARPHA et al |
| **hCoV-19/Venezuela/AraXMLD/2023** | EPI_ISL_18059774 | | LabGENOMIK | Laboratorio de Virologia Molecular | Rossana C Jaspe et al |
| **hCoV-19/Brazil/SC-FIOCRUZ-3771/2023** | EPI_ISL_17658334 | | Laboratorio Central de Saude Publica do Estado de Santa Catarina (LACEN/SC) | Laboratory of Respiratory Viruses and Measles, Oswaldo Cruz Institute, FIOCRUZ | Paola Resende et al |
| **hCoV-19/Chile/MA-62154/2023** | EPI_ISL_18335024 | | Genetica Molecular and Subdepartamento de Virologia ISP Chile | Instituto de Salud Publica de Chile | Mariela Guajardo et al |
| **hCoV-19/Bolivia/CENETROP-008/2023** | EPI_ISL_18038197 | | Hospital El Torno | Centro Nacional de Enfermedades Tropicales | Loayza et al |
| **hCoV-19/Mexico/SIN_InDRE_FB3975/2023** | EPI_ISL_18296071 | | LESP Sinaloa | Instituto de Diagnostico y Referencia Epidemiologicos (INDRE) | Abril Rodriguez-Maldonado et al |
| **hCoV-19/SintMaarten/SX-RIVM-130638/2023** | EPI_ISL_18098334 | | National Institute for Public Health and the Environment (RIVM) | National Institute for Public Health and the Environment (RIVM) | Adam Meijer et al |
| **hCoV-19/India/KA-ICMR-NIV-G-14591/2023** | EPI_ISL_18120293 | | Indian Council of Medical Research, National Institute of Virology | National Institute of Virology - Influenza | Dr. Varsha Potdar et al |
| **hCoV-19/ElSalvador/1028-LNSP/2023** | EPI_ISL_18401382 | | Secretaría de Estado SS Ministerio de Salud | Laboratorio Nacional de Salud Publica | Laboratorio de Secuenciacion y Genomica del LNSP-INS et al |
| **hCoV-19/Guatemala/604-LNS/2023** | EPI_ISL_17277062 | | Laboratorio Nacional de Salud (LNS), Guatemala | Laboratorio Nacional de Salud (LNS), Guatemala | Mendoza L et al |
| **hCoV-19/Mexico/GUA_InDRE_FB1821/2023** | EPI_ISL_17713995 | | LESP Guanajuato | Instituto de Diagnostico y Referencia Epidemiologicos (INDRE) | Claudia Wong-Arambula et al |
| **hCoV-19/DRC/INRB-0529/2023** | EPI_ISL_17788180 | | Pathogen Genomics Lab, National Institute for Biomedical Research (INRB) | Pathogen Genomics Lab, National Institute for Biomedical Research (INRB) | Placide Mbala-Kingebeni et al B |
| **hCoV-19/Guatemala/1129-LNS/2023** | EPI_ISL_17954110 | | Dirección del Laboratorio Nacional de Salud (LNS), Guatemala | Dirección del Laboratorio Nacional de Salud (LNS), Guatemala | Mendoza L et al |
| **hCoV-19/Mexico/DUR_IBT_IMSS_009188_CIBIN_NC/2023** | EPI_ISL_18589778 | | Centro de Investigacion Biomedica del Noreste (CIBIN) | Instituto de Biotecnologia de la UNAM | Julio Elias Alvarado-Yaah et al |
| **hCoV-19/India/MH-ICMR-NIV-G-14526/2023** | EPI_ISL_18052815 | | ICMR-National Institute of Virology | NIV Influenza | Dr. Varsha Potdar et al |
| **hCoV-19/Pakistan/NIH-AHF-S35/2023** | EPI_ISL_18642781 | | Department of Virology, National Institute of Health, Islamabad, Pakistan | Department of Virology, National Institute of Health, Islamabad, Pakistan | Massab Umair et al |
| **hCoV-19/Pakistan/NIH-B120-S10/2023** | EPI_ISL_17156318 | | Department of Virology, National Institute of Health, Islamabad, Pakistan | Department of Virology, National Institute of Health, Islamabad, Pakistan | Massab Umair et al |
| **hCoV-19/Beijing/BJCDC-2781/2023** | EPI_ISL_18438180 | | Institute for Infectious Disease and Endemic Disease Control, Beijing Center for Disease Control and Prevention | Institute for Infectious Disease and Endemic Disease Control, Beijing Center for Disease Control and Prevention | Pan et al |
| **hCoV-19/Pakistan/NIH-AHF-S27/2023** | EPI_ISL_18445466 | | Department of Virology, National Institute of Health, Islamabad, Pakistan | Department of Virology, National Institute of Health, Islamabad, Pakistan | Massab Umair et al |
| **hCoV-19/HongKong/VB23250718/2023** | EPI_ISL_18760242 | | Prince of Wales Hospital | Hong Kong Department of Health | Alan K.L. Tsang et al |
| **hCoV-19/USA/GA-CDC-QDX48586893/2023** | EPI_ISL_17478153 | | Quest Diagnostics Incorporated | Centers for Disease Control and Prevention Division of Viral Diseases, Pathogen Discovery | Dakota Howard et al |
| **hCoV-19/Botswana/R231B91_1114449/22023** | EPI_ISL_18282215 | | National Health Laboratory | Botswana Harvard HIV Reference Laboratory | Sikhulile Moyo et al |
| **hCoV-19/Guinea/IPGui-28521/2023** | EPI_ISL_18050126 | | Institut Pasteur de Guinée | Institut Pasteur de Guinée | Reine ANGUINZE et al |
| **hCoV-19/Senegal/25576369/2023** | EPI_ISL_18713709 | | Institut Pasteur de Dakar | Institut Pasteur de Dakar | Jallow et al |
| **hCoV-19/ElSalvador/1163-LNSP/2023** | EPI_ISL_18401383 | | Hospital Nacional San Rafael | Laboratorio Nacional de Salud Publica | Laboratorio de Secuenciacion y Genomica del LNSP-INS et al |
| **hCoV-19/Mexico/COL_InDRE_FB3788/2023** | EPI_ISL_18295950 | | LESP Colima | Instituto de Diagnostico y Referencia Epidemiologicos (INDRE) | Abril Rodriguez-Maldonado et al |
| **hCoV-19/Mexico/PUE_InDRE_FB3999/2023** | EPI_ISL_18296048 | | LESP Puebla | Instituto de Diagnostico y Referencia Epidemiologicos (INDRE) | Abril Rodriguez-Maldonado et al |
| **hCoV-19/Ecuador/GENSBIO-INSPI-SG-167/2023** | EPI_ISL_17722429 | | Laboratorio de Influenza y Otros Virus Respiratorios, INSPI-CZ9 | Centro de Referencia Nacional de Genomica, Secuenciacion y Bioinformatica GENSBIO, INSPI-CZ9 | Andres Carrazco-Montalvo* et al |
| **hCoV-19/Ecuador/USFQ-4814/2023** | EPI_ISL_17525324 | | Institute of Microbiology, Universidad San Francisco de Quito | Institute of Microbiology, Universidad San Francisco de Quito | Rommel Guevara et al |
| **hCoV-19/Uruguay/DLSP_727/2023** | EPI_ISL_18041135 | | Departamento Laboratorios de Salud Pública (DLSP) Ministerio de Salud Pública | Departamento Laboratorios de Salud Pública (DLSP) Ministerio de Salud Pública | Cortinas et al |
| **hCoV-19/Argentina/INEI125673/2023** | EPI_ISL_18095180 | | Servicio Virosis Respiratorias-Departamento Virología-INEI | Instituto Nacional Enfermedades Infecciosas C.G.Malbran | Baumeister E. et al |
| **hCoV-19/Uruguay/DLSP_1239/2023** | EPI_ISL_18041142 | | Departamento Laboratorios de Salud Pública (DLSP) Ministerio de Salud Pública | Departamento Laboratorios de Salud Pública (DLSP) Ministerio de Salud Pública | Cortinas et al |
| **hCoV-19/Argentina/INEI125536/2023** | EPI_ISL_17801367 | | Servicio Virosis Respiratorias-Departamento Virología-INEI C.G.Malbran | Instituto Nacional Enfermedades Infecciosas C.G.Malbran | Baumeister E. et al |
| **hCoV-19/Bolivia/CENETROP-017/2023** | EPI_ISL_18038206 | | Laboratorio de Biología Molecular Vacaflores&Bartos s.r.l. Plexus | Centro Nacional de Enfermedades Tropicales | Loayza et al |
| **hCoV-19/Argentina/INEI125774/2023** | EPI_ISL_18422426 | | Servicio Virosis Respiratorias-Departamento Virología-INEI | Instituto Nacional Enfermedades Infecciosas C.G.Malbran | Baumeister E. et al |
| **hCoV-19/DominicanRepublic/STGO2-LNSPDD/2023** | EPI_ISL_17818998 | | Laboratorio Nacional de Salud Pública Dr. Defilló | Laboratorio Nacional de Salud Pública Dr. Defilló | Isaac Miguel Sánchez et al |
| **hCoV-19/Hungary/101139-UPL23-000720/2023** | EPI_ISL_17347322 | | Institute of Laboratory Medicine, Semmelweis University and Epidemiology and Surveillance Centre, Semmelweis University | iBioscience Kft. | Gergő Túri et al |
| **hCoV-19/Kosovo/15_NIPHK/2023** | EPI_ISL_17300046 | | Department of Microbiology, National Institute of Public Health of Kosovo | Department of Microbiology, National Institute of Public Health of Kosovo | Xhevat Jakupi et al |
| **hCoV-19/USA/NY-NYCPHL-016833/2023** | EPI_ISL_18865277 | | Pathline Labs | New York City Public Health Laboratory | Jade Wang et al |
| **hCoV-19/Bahrain/CDL-730502449/2023** | EPI_ISL_17081297 | | Communicable Disease Laboratory | Communicable Disease Laboratory | Marhoon et al |
| **hCoV-19/CaboVerde/ST-0069/2023** | EPI_ISL_17813776 | | Virology Laboratory of Praia | Instituto Nacional de Saúde Pública (INSP) | Kevin Sanders et al |
| **hCoV-19/CaboVerde/ST-AG0056/2023** | EPI_ISL_17813739 | | Virology Laboratory of Praia | Instituto Nacional de Saúde Pública (INSP) | Kevin Sanders et al |
| **hCoV-19/Bolivia/1012/2023** | EPI_ISL_17818558 | | Clínica Médica MEDICMEL | CENTRO NACIONAL DE ENFERMEDADES TROPICALES (CENETROP) | Loayza et al |
| **hCoV-19/Lithuania/MB0328_0218/2023** | EPI_ISL_17622034 | | Viesoji istaiga KLAIPEDOS SENAMIESCIO PIRMINES SVEIKATOS PRIEZIUROS CENTRAS | National Public Health Surveillance Laboratory | Lukas Zemaitis et al |
| **hCoV-19/Guatemala/1211-LNS/2023** | EPI_ISL_17954169 | | Dirección del Laboratorio Nacional de Salud (LNS), Guatemala | Dirección del Laboratorio Nacional de Salud (LNS), Guatemala | Mendoza L et al |
| **hCoV-19/Denmark/DCGC-647295/2023** | EPI_ISL_17787537 | | Department of Bacteria, Parasites and Fungi, Statens Serum Institut, Copenhagen, Denmark | Statens Serum Institut Bioinformatics and Microbial Genomics | Povilas Matusevicius on behalf of the Danish Covid-19 Genome Consortium et al |
| **hCoV-19/Iceland/L-4050/2023** | EPI_ISL_17529049 | | Landspitali Department of Clinical Microbiology | Landspitali Department of Clinical Microbiology | Freyja Valsdottir et al |
| **hCoV-19/Iran/EE176/2023** | EPI_ISL_18443006 | | Mashhad University of Medical Sciences | COVID-19 National Reference Laboratory, Pasteur Institute of Iran | Zahra Ahmadi et al |
| **hCoV-19/Austria/AGES-1114510/2023** | EPI_ISL_16951665 | | Covid Fighters | AGES IMED Vienna | Lippert K. et al |
| **hCoV-19/NewZealand/23XA2562/2023** | EPI_ISL_17740132 | | Pathlab Rotorua | Institute of Environmental Science and Research (ESR) | Sarah Cockerton et al |
| **hCoV-19/Mexico/NLE_LANGEBIO_IMSS_15151/2023** | EPI_ISL_18365251 | | Centro de Investigación Biomédica del Noreste (CIBIN) | Unidad de Genomica Avanzada | Authors (in last name alphabetical order): Julio Elias Alvarado-Yaah et al |
| **hCoV-19/Haiti/LNSP89260021/2023** | EPI_ISL_18360182 | | Laboratoire National de Santé Publique | Laboratoire National de Santé Publique | Rosette Demar Calixte et al |
| **hCoV-19/USA/NH-NHPHL-2303020025/2023** | EPI_ISL_18527258 | | State of New Hampshire Public Health Laboratories | State of New Hampshire Public Health Laboratories | Chris Benton et al |
| **hCoV-19/Palestine/527738/2023** | EPI_ISL_17671795 | | CPHL-MOH | Molecular Diagnostic Laboraory-CPHL | Shtayeh Issa et al |
| **hCoV-19/Colombia/DC-INS-VG-29054/2023** | EPI_ISL_17652715 | | Instituto De Diagnostico Medico Idime | Instituto Nacional de Salud, Dirección de Investigación en Salud Pública | Katherine Laiton-Donato et al |
| **hCoV-19/Peru/LIM-INS-45040/2023** | EPI_ISL_17773051 | | Laboratorio de Referencia Nacional de Virus Inmunoprevenibles. Centro Nacional de Salud Publica. Instituto Nacional de Salud Peru. | Laboratorio de Referencia Nacional de Virus Inmunoprevenibles. Centro Nacional de Salud Publica. Instituto Nacional de Salud Peru. | Carlos Padilla Rojas et al |
| **hCoV-19/Peru/LIM-INS-45745/2023** | EPI_ISL_18143283 | | Laboratorio de Referencia Nacional de Virus Inmunoprevenibles. Centro Nacional de Salud Publica. Instituto Nacional de Salud Peru. | Laboratorio de Referencia Nacional de Virus Inmunoprevenibles. Centro Nacional de Salud Publica. Instituto Nacional de Salud Peru. | Carlos Padilla Rojas et al |
| **hCoV-19/Norway/4066/2023** | EPI_ISL_17512550 | | University Hospital of Northern Norway, Department for Microbiology and Infectious Disease Control | Norwegian Institute of Public Health, Department of Virology | Kathrine Stene-Johansen et al |
| **hCoV-19/Mexico/YUC_LANGEBIO_IMSS_14356/2023** | EPI_ISL_18259722 | | Unidad de Investigacion Medica de Yucatan (UIMY) | Unidad de Genomica Avanzada | Authors (in last name alphabetical order): Julio Elias Alvarado-Yaah et al |
| **hCoV-19/Mexico/SON_LANGEBIO_IMSS_14634/2023** | EPI_ISL_18260137 | | Laboratorio Central de Epidemiologia (LCE) | Unidad de Genomica Avanzada | Authors (in last name alphabetical order): Julio Elias Alvarado-Yaah et al |
| **hCoV-19/Philippines/PH-PGCMIN2752/2023** | EPI_ISL_17819162 | | Southern Philippines Medical Center | Philippine Genome Center Mindanao | Joel Hassan G. Tolentino et al |
| **hCoV-19/NewZealand/23ZA1828/2023** | EPI_ISL_17504234 | | Canterbury Health Laboratories (CHL) | Institute of Environmental Science and Research (ESR) | Sarah Cockerton et al |
| **hCoV-19/Guinea/IPGui-28514/2023** | EPI_ISL_18050122 | | Institut Pasteur de Guinée | Institut Pasteur de Guinée | Reine ANGUINZE et al |
| **hCoV-19/env/Liechtenstein/CeMM35236/2023** | EPI_ISL_17499230 | | Universität Innsbruck, Institut für Mikrobiologie | Bergthaler laboratory, CeMM Research Center for Molecular Medicine of the Austrian Academy of Sciences | Lukas Endler et al |
| **hCoV-19/Qinghai/QHCDC-ETYY-189/2023** | EPI_ISL_17729979 | | Microbiological testing center, Qinghai Center for Disease Prevention and Control | Microbiological testing center, Qinghai Center for Disease Prevention and Control | He et al |
| **hCoV-19/DominicanRepublic/2144764-LNSPDD/2023** | EPI_ISL_17818903 | | Laboratorio Nacional de Salud Pública Dr. Defilló | Laboratorio Nacional de Salud Pública Dr. Defilló | Isaac Miguel Sánchez et al |
| **hCoV-19/USA/SD-CDC-2-7134224/2023** | EPI_ISL_18031727 | | SD Public Health Laboratory | Centers for Disease Control and Prevention Division of Viral Diseases, Pathogen Discovery | Mili Sheth et al |
| **hCoV-19/Thailand/NIC_MKM_5209/2023** | EPI_ISL_17785930 | | Regional Medical Sciences Center 7 Khon Kaen | National Institute of Health, Department of Medical Sciences, Ministry of Public Health, Thailand | Pilailuk Okada et al |
| **hCoV-19/USA/WV-WVU-WV137220/2023** | EPI_ISL_18319775 | | Reynolds Memorial Hospital | WVU and Marshall University Combined Genomics Core Facilities | James Denvir et al |
| **hCoV-19/Venezuela/Dtto2566/2023** | EPI_ISL_18494947 | | INHRR | Laboratorio de Virologia Molecular | Rossana C Jaspe et al |
| **hCoV-19/DominicanRepublic/2144624-LNSPDD/2023** | EPI_ISL_17818884 | | Laboratorio Nacional de Salud Pública Dr. Defilló | Laboratorio Nacional de Salud Pública Dr. Defilló | Isaac Miguel Sánchez et al |
| **hCoV-19/NewZealand/ESR-ZA3517/2023** | EPI_ISL_18047872 | | ESR | Institute of Environmental Science and Research (ESR) | Sarah Cockerton et al |
| **hCoV-19/Croatia/HZJZ_4851/2023** | EPI_ISL_18248903 | | Klinika za infektivne bolesti "Dr. Fran Mihaljević" | Hrvatski zavod za javno zdravstvo | Irena Tabain et al |
| **hCoV-19/Peru/LIM-INS-45261/2023** | EPI_ISL_17852732 | | Laboratorio de Referencia Nacional de Virus Inmunoprevenibles. Centro Nacional de Salud Publica. Instituto Nacional de Salud Peru. | Laboratorio de Referencia Nacional de Virus Inmunoprevenibles. Centro Nacional de Salud Publica. Instituto Nacional de Salud Peru. | Carlos Padilla Rojas et al |
| **hCoV-19/Peru/LIM-INS-45985/2023** | EPI_ISL_18251859 | | Laboratorio de Referencia Nacional de Virus Inmunoprevenibles, Centro Nacional de Salud Publica, Instituto Nacional de Salud Peru | Laboratorio de Referencia Nacional de Virus Inmunoprevenibles, Centro Nacional de Salud Publica, Instituto Nacional de Salud Peru | Carlos Padilla Rojas et al |
| **hCoV-19/Peru/LIM-INS-46083/2023** | EPI_ISL_18251957 | | Laboratorio de Referencia Nacional de Virus Inmunoprevenibles, Centro Nacional de Salud Publica, Instituto Nacional de Salud Peru | Laboratorio de Referencia Nacional de Virus Inmunoprevenibles, Centro Nacional de Salud Publica, Instituto Nacional de Salud Peru | Carlos Padilla Rojas et al |
| **hCoV-19/Peru/LIM-INS-46140/2023** | EPI_ISL_18252013 | | Laboratorio de Referencia Nacional de Virus Inmunoprevenibles, Centro Nacional de Salud Publica, Instituto Nacional de Salud Peru | Laboratorio de Referencia Nacional de Virus Inmunoprevenibles, Centro Nacional de Salud Publica, Instituto Nacional de Salud Peru | Carlos Padilla Rojas et al |
| **hCoV-19/Mexico/SIN_InDRE_FB4806/2023** | EPI_ISL_18785800 | | Laboratorio Estatal de Salud Publica Sinaloa | Instituto de Diagnostico y Referencia Epidemiologicos (INDRE) | Abril Rodriguez-Maldonado et al |
| **hCoV-19/Paraguay/LERAP_43-3/2023** | EPI_ISL_18630901 | | Laboratorio Epidemiológico Regional Alto Paraná | Laboratorio Epidemiológico Regional Alto Paraná | Daniela Poullion et al |
| **hCoV-19/Japan/5GA-529/2023** | EPI_ISL_18390813 | | Department of Laboratory Medicine, Shinshu University Hospital | Department of Laboratory Medicine, Shinshu University Hospital | Shohei Shigeto et al |
| **hCoV-19/Paraguay/LERP-08-2-09/2023** | EPI_ISL_18396196 | | Laboratorio Epidemiológico Regional Alto Paraná | Laboratorio Epidemiológico Regional Alto Paraná | Daniela Poullion et al |
| **hCoV-19/Hungary/103379-UPL24-000565/2023** | EPI_ISL_18883059 | | Institute of Laboratory Medicine, Semmelweis University and Epidemiology and Surveillance Centre, Semmelweis University | iBioscience Kft. | Gergő Túri et al |
| **hCoV-19/Paraguay/LERAP_11-4/2023** | EPI_ISL_18852476 | | Laboratorio Epidemiológico Regional Alto Paraná | Laboratorio Epidemiológico Regional Alto Paraná | Daniela Poullion et al |
| **hCoV-19/Paraguay/731912/2023** | EPI_ISL_18887013 | | HR - PEDRO JUAN CABALLERO | Laboratorio Central de Salud Pública | Sandra González et al |
| **hCoV-19/Chile/ML-5367/2024** | EPI_ISL_18898654 | | Genetica Molecular and Subdepartamento de Virologia ISP Chile | Instituto de Salud Publica de Chile | Mariela Guajardo et al |

| | | | | | |
|---|---|---|---|---|---|
| **hCoV-19/Chile/ML-5372/2024** | EPI_ISL_18898657 | | Genetica Molecular and Subdepartamento de Virologia ISP Chile | Instituto de Salud Publica de Chile | Mariela Guajardo et al |
| **hCoV-19/CanaryIslands/HUGCDN-04309197/2023** | EPI_ISL_18741452 | | Servicio Microbiología H.U. Dr. Negrín | Servicio Microbiología H.U. Dr. Negrín | Eduardo Lagarejos González Laura Floren Zabala Ana Bordes Benítez et al |
| **hCoV-19/Germany/BE-RKI-I-1140546/2023** | EPI_ISL_18599897 | | Nationale Referenzzentrum für Influenza; FG17 | Robert Koch Institute | Ralf Dürrwald et al |
| **hCoV-19/Peru/LIM-INS-47800/2023** | EPI_ISL_18756799 | | Laboratorio de Referencia Nacional de Virus Inmunoprevenibles. Centro Nacional de Salud Publica. Instituto Nacional de Salud Peru. | Laboratorio de Referencia Nacional de Virus Inmunoprevenibles. Centro Nacional de Salud Publica. Instituto Nacional de Salud Peru. | Carlos Padilla Rojas et al |
| **hCoV-19/Chile/RM-1285/2024** | EPI_ISL_18768502 | | Genetica Molecular and Subdepartamento de Virologia ISP Chile | Instituto de Salud Publica de Chile | Mariela Guajardo et al |
| **hCoV-19/Peru/LIM-INS-47223/2023** | EPI_ISL_18611315 | | Laboratorio de Referencia Nacional de Virus Inmunoprevenibles. Centro Nacional de Salud Publica. Instituto Nacional de Salud Peru. | Laboratorio de Referencia Nacional de Virus Inmunoprevenibles. Centro Nacional de Salud Publica. Instituto Nacional de Salud Peru. | Carlos Padilla Rojas et al |
| **hCoV-19/Aruba/AW-RIVM-123087/2022** | EPI_ISL_16445610 | | National Institute for Public Health and the Environment (RIVM) | National Institute for Public Health and the Environment (RIVM) | Adam Meijer et al |
| **hCoV-19/Vietnam/BMH-519064/2023** | EPI_ISL_17719708 | | Thai Binh General Hospital | Bachmai Hospital | Phuong Truong et al |
| **hCoV-19/USA/GA-CDC-QDX46677096/2023** | EPI_ISL_17009686 | | Quest Diagnostics Incorporated | Centers for Disease Control and Prevention Division of Viral Diseases, Pathogen Discovery | Dakota Howard et al |
| **hCoV-19/Argentina/7302-119/2023** | EPI_ISL_17966242 | | Laboratorio Central, Ministerio de Salud Córdoba | Departamento Laboratorio Central and Unidad Operativa Centro Nacional de Genómica y Bioinformática | Castro et al |
| **hCoV-19/USA/MT-MTPHL-4090927/2023** | EPI_ISL_17301104 | | Montana Public Health Laboratory | Montana Public Health Laboratory | Joy Ritter et al |
| **hCoV-19/USA/IN-CDC-QDX46502051/2023** | EPI_ISL_16993151 | | Quest Diagnostics Incorporated | Centers for Disease Control and Prevention Division of Viral Diseases, Pathogen Discovery | Dakota Howard et al |
| **hCoV-19/Curacao/CW-RIVM-130062/2023** | EPI_ISL_17689055 | | National Institute for Public Health and the Environment (RIVM) | National Institute for Public Health and the Environment (RIVM) | Adam Meijer et al |
| **hCoV-19/Mexico/MEX_IBT_IMSS_11583/2023** | EPI_ISL_18588936 | | Laboratorio Central de Epidemiologia (LCE) | Instituto de Biotecnologia de la UNAM | Julio Elias Alvarado-Yaah et al |
| **hCoV-19/Mexico/BCN_InDRE_FB1819/2023** | EPI_ISL_17713953 | | LESP Baja California | Instituto de Diagnostico y Referencia Epidemiologicos (INDRE) | Claudia Wong-Arambula et al |
| **hCoV-19/Jordan/RMSR14S91/2023** | EPI_ISL_16968043 | | Jordan Royal Medical Services Mobile biological lab | Jordan Royal Medical Services Genomics Core Lab | Dr. Rame Hamdi khasawneh et al |
| **hCoV-19/Egypt/CPHL-22873/2023** | EPI_ISL_17427623 | | Central Public Health Laboratories (CPHL) | Central Public Health Laboratories (CPHL) | Wael H. Roshdy et al |
| **hCoV-19/Aruba/AW-RIVM-130924/2023** | EPI_ISL_18210141 | | National Institute for Public Health and the Environment (RIVM) | National Institute for Public Health and the Environment (RIVM) | Adam Meijer et al |
| **hCoV-19/USA/SC-SCDHEC-0232/2023** | EPI_ISL_18207287 | | South Carolina Department of Health and Environmental Control | South Carolina Department of Health and Environmental Control | Freeman et al |
| **hCoV-19/Iceland/L-3497/2023** | EPI_ISL_16720229 | | Landspitali Department of Clinical Microbiology | Landspitali Department of Clinical Microbiology | Freyja Valsdottir et al |
| **hCoV-19/Mexico/NLE_CIAD-IMSS_01008492-CIBIN/2023** | EPI_ISL_18279696 | | Centro de Investigacion Biomedica del Noreste (CIBIN) | Microbial Genomics Laboratory | Consorcio Mexicano de Vigilancia Genomica (CoViGen-Mex). Authors (in last name alphabetical order): Julio Elias Alvarado-Yaah et al |
| **hCoV-19/Mexico/VER_InDRE_FB4123/2023** | EPI_ISL_18424280 | | LESP Veracruz | Instituto de Diagnostico y Referencia Epidemiologicos (INDRE) | Abril Rodriguez-Maldonado et al |
| **hCoV-19/USA/KS-KHEL-13207/2023** | EPI_ISL_16987228 | | Kansas Health and Environmental Lab | Kansas Health and Environmental Lab | Mike Grose et al |
| **hCoV-19/CostaRica/INC-10688-787890/2023** | EPI_ISL_17510676 | | Hospital Dr. Enrique Baltodano Briceño (Liberia/Guanacaste) | Inciensa, Instituto de Investigación y Enseñanza en Nutrición y Salud | Francisco Duarte et al |
| **hCoV-19/CaboVerde/ST-AG0125/2023** | EPI_ISL_18873729 | | Virology Laboratory of Praia | Instituto Nacional de Saúde Pública (INSP) | Kevin Sanders et al |
| **hCoV-19/Bangladesh/IEDCR-2200729/2023** | EPI_ISL_17837265 | | Institute of Epidemiology, Disease Control and Research (IEDCR) | Institute of Epidemiology, Disease Control and Research (IEDCR) | Manjur Hossain Khan et al |
| **hCoV-19/AntiguaandBarbuda/173234/2023** | EPI_ISL_16913070 | | Mount St. John's Medical Centre | CARPHA | Carlos Rampersad on behalf of CARPHA et al |
| **hCoV-19/Colombia/DC-LSPSDS-12447/2023** | EPI_ISL_17703478 | | MUESTRA CONGLOMERADO - SUB RED SUR OCCIDENTE | LSPSDS | Johana Hernandez Toloza et al |
| **hCoV-19/Venezuela/Zul2489/2023** | EPI_ISL_18059754 | | INHRR | Laboratorio de Virologia Molecular | Rossana C Jaspe et al |
| **hCoV-19/Guatemala/INC-LNS-423/2023** | EPI_ISL_18139728 | | Dirección de Laboratorio Nacional de Salud (DLNS), Guatemala | Inciensa, Instituto Costarricense de Investigación y Enseñanza en Nutrición y Salud | Mendoza L et al |
| **hCoV-19/Chile/RM-38156/2023** | EPI_ISL_17831849 | | Genetica Molecular and Subdepartamento de Virologia ISP Chile | Instituto de Salud Publica de Chile | Mariela Guajardo et al |
| **hCoV-19/Romania/B-2851/2023** | EPI_ISL_18261939 | | National Institute of Infectious Diseases-Prof. Dr. Matei Bals Molecular Diagnostics Laboratory | National Institute of Infectious Diseases-Prof. Dr. Matei Bals Molecular Diagnostics Laboratory | Simona Paraschiv et al |
| **hCoV-19/Jordan/MOH-CPHL-06/2023** | EPI_ISL_18166721 | | Ministry of Health (MOH), Central Public Health Laboratory (CPHL), Molecular lab | Ministry of Health (MOH), Central Public Health Laboratory (CPHL) | Dr.Gazo et al |
| **hCoV-19/CostaRica/INC-11476-796404/2023** | EPI_ISL_17953742 | | Area de Salud Cariari [Pococi/Limon] | Inciensa, Instituto de Investigación y Enseñanza en Nutrición y Salud | Francisco Duarte et al |
| **hCoV-19/Kosovo/13_S13/2023** | EPI_ISL_17383547 | | Department of Microbiology, National Institute of Public Health of Kosovo | Department of Microbiology, National Institute of Public Health of Kosovo | Xhevat Jakupi et al |
| **hCoV-19/Chile/CO-23255/2023** | EPI_ISL_17299945 | | Genetica Molecular and Subdepartamento de Virologia ISP Chile | Instituto de Salud Publica de Chile | Mariela Guajardo et al |
| **hCoV-19/Honduras/FM-144/2023** | EPI_ISL_18663653 | | Laboratorio Nacional de Virologìa (SESAL),Honduras | Laboratorio Nacional de Virologìa (SESAL),Honduras | Dra. Soany Avilez et al |
| **hCoV-19/Curacao/CW-RIVM-128976/2023** | EPI_ISL_17468147 | | National Institute for Public Health and the Environment (RIVM) | National Institute for Public Health and the Environment (RIVM) | Adam Meijer et al |
| **hCoV-19/Curacao/CW-RIVM-127280/2023** | EPI_ISL_17202048 | | National Institute for Public Health and the Environment (RIVM) | National Institute for Public Health and the Environment (RIVM) | Adam Meijer et al |
| **hCoV-19/Ireland/LK-NVRL-SIRL00008339/2023** | EPI_ISL_16997354 | | National Virus Reference Laboratory | National Virus Reference Laboratory | Zoe Yandle et al |
| **hCoV-19/India/DL-ILBS-7560/2023** | EPI_ISL_17268952 | | Dr. Dangs Lab | Institute of Liver and Biliary Sciences (ILBS) | Ekta Gupta et al |
| **hCoV-19/USA/MN-CDC-VSX-A062551/2023** | EPI_ISL_17167426 | | Helix | Centers for Disease Control and Prevention Division of Viral Diseases, Pathogen Discovery | Dakota Howard et al |
| **hCoV-19/USA/WA-UW-89061/2023** | EPI_ISL_18207620 | | Laboratory Medicine and Pathology, University of Washington | Laboratory Medicine and Pathology, University of Washington | Roychoudhury et al |
| **hCoV-19/Vietnam/NHTD-S28/2023** | EPI_ISL_18876198 | | EI group, Oxford University Clinical Research Unit | EI group, Oxford University Clinical Research Unit | Nguyen et al |
| **hCoV-19/CaboVerde/SV-0139/2023** | EPI_ISL_17813793 | | Virology Laboratory of São Vicente | Instituto Nacional de Saúde Pública (INSP) | Kevin Sanders et al |
| **hCoV-19/Greece/3301/2023** | EPI_ISL_17392875 | | National Public Health Organization | National Public Health Organization | "Kyriaki Tryfinopoulou et al |
| **hCoV-19/Croatia/HZJZ_3568/2023** | EPI_ISL_17554316 | | Hrvatski zavod za javno zdravstvo | Hrvatski zavod za javno zdravstvo | Irena Tabain et al |
| **hCoV-19/USA/MI-CDC-LC1024428/2023** | EPI_ISL_17237156 | | Laboratory Corporation of America | Centers for Disease Control and Prevention Division of Viral Diseases, Pathogen Discovery | Dakota Howard et al |
| **hCoV-19/Switzerland/BE-IFIK-230327_os_32/2023** | EPI_ISL_17404565 | | Institute for Infectious Diseases | Institute for Infectious Diseases, University of Bern | Stefan Neuenschwander et al |
| **hCoV-19/NewZealand/23XA1730/2023** | EPI_ISL_17422810 | | Medlab South | Institute of Environmental Science and Research (ESR) | Sarah Cockerton et al |
| **hCoV-19/Bonaire/BQ-RIVM-130469/2023** | EPI_ISL_17971226 | | National Institute for Public Health and the Environment (RIVM) | National Institute for Public Health and the Environment (RIVM) | Adam Meijer et al |
| **hCoV-19/Qinghai/QHCDC-QHSYY-566/2023** | EPI_ISL_17988032 | | Microbiological testing center, Qinghai Center for Disease Prevention and Control | Microbiological testing center, Qinghai Center for Disease Prevention and Control | Lei et al |
| **hCoV-19/Japan/PG-568810/2023** | EPI_ISL_18289715 | | Sendai City Institute of Public Health | Sendai City Institute of Public Health | Youko Kano et al |
| **hCoV-19/CostaRica/INC-11848-804842/2023** | EPI_ISL_18284334 | | AREA DE SALUD BARVA (COOPESIBA) [BARVA/HEREDIA] | Inciensa, Instituto de Investigación y Enseñanza en Nutrición y Salud | Francisco Duarte et al |
| **hCoV-19/Ireland/D-SVUH-FAX24467-026/2023** | EPI_ISL_18290907 | | St. Vincent's Microbiology | St. Vincent's Microbiology | Eric Brown et al |
| **hCoV-19/Sweden/Z-6045590245/2023** | EPI_ISL_18558539 | | The Public Health Agency of Sweden | The Public Health Agency of Sweden | Emmi Andersson et al |
| **hCoV-19/Brunei/MGS-7123058180/2023** | EPI_ISL_18272607 | | National Virology Reference Laboratory | Microbial Genomic Services | Zainun Zaini et al |
| **hCoV-19/Singapore/R52MR50/2023** | EPI_ISL_18056583 | | National Public Health Laboratory, National Centre for Infectious Diseases | National Public Health Laboratory, National Centre for Infectious Diseases | Yichen Ding et al |
| **hCoV-19/Palestine/532874/2023** | EPI_ISL_18545864 | | CPHL-MOH | Molecular Diagnostic Laboraory-CPHL | Shtayeh Issa et al |
| **hCoV-19/SouthAfrica/CERI-KRISP-K059837/2023** | EPI_ISL_18607095 | | Pathcare | CERI, Centre for Epidemic Response and Innovation, Stellenbosch University and KRISP, KZN Research Innovation and Sequencing Platform, UKZN. | Jean Maritz et al |
| **hCoV-19/Bulgaria/23BG-NC_023480_R39/2023** | EPI_ISL_18399158 | | National Center of Infectious and Parasitic Diseases | National Center of Infectious and Parasitic Diseases | Alexiev et al |
| **hCoV-19/Kenya/CDC-DLSP-Kisumu-COV235936/2023** | EPI_ISL_18885262 | | DLSP KEMRI/CDC KISUMU Lab | DLSP_CDC_KISUMU | Shirley Lidechi et al |
| **hCoV-19/Japan/PG-556061/2023** | EPI_ISL_18106209 | | Pathogen Genomics Center, National Institute of Infectious Diseases | Pathogen Genomics Center, National Institute of Infectious Diseases | Naobumi Sasaki et al |
| **hCoV-19/SouthKorea/KDCA260834/2023** | EPI_ISL_18236813 | | Division of Emerging Infectious Diseases, Bureau of Infectious Diseases Diagnosis Control, Korea Disease Control and Prevention Agency | Division of Emerging Infectious Diseases, Bureau of Infectious Diseases Diagnosis Control, Korea Disease Control and Prevention Agency | Jeong-Ah Kim et al |
| **hCoV-19/Hubei/HCDC-0978/2023** | EPI_ISL_17615019 | | Xiaogan Center for Disease Control and Prevention | Institute of Health Inspection and Testing, Hubei Provincial Center for Disease Control and Prevention | Jiang et al |
| **hCoV-19/Shanxi/TYCDC231/2023** | EPI_ISL_18495162 | | Microbiology Department, Taiyuan Center for Disease Control and Prevention | Microbiology Department, Taiyuan Center for Disease Control and Prevention | Wang et al |
| **hCoV-19/Guangdong/GDCDC_GZHG_1265/2023** | EPI_ISL_18034549 | | Institute of Pathogenic Microbiology, Guangdong Provincial Center for Disease Control and Prevention | Institute of Pathogenic Microbiology, Guangdong Provincial Center for Disease Control and Prevention | Chang Zhang et al |
| **hCoV-19/Hubei/HCDC-WH2407/2023** | EPI_ISL_18078315 | | Wuhan Center for Disease Control and Prevention | Hubei Provincial Center for Disease Control and Prevention Center of Pathogenic Microorganism Detection | Yongzhong Jiang et al |
| **hCoV-19/Henan/XXCDC-060703/2023** | EPI_ISL_18213307 | | Clinical Laboratory, Xinxiang Center for Disease Control and Prevention | Clinical Laboratory, Xinxiang Center for Disease Control and Prevention | Yu et al |
| **hCoV-19/Fujian/FCDC-FZCX0131/2023** | EPI_ISL_18127150 | | Institute for the Prevention and Control of Infectious Diseases, Fujian Provincial Center for Disease Control and Prevention | Institute for the Prevention and Control of Infectious Diseases, Fujian Provincial Center for Disease Control and Prevention | Qi Lin et al |
| **hCoV-19/Anhui/TLCDC-CX038/2023** | EPI_ISL_18277172 | | Microbiology Laboratory, Tongling City Center for Disease Control and Prevention | Microbiology Laboratory, Tongling City Center for Disease Control and Prevention | Yihua Zhang et al |
| **hCoV-19/China/un-SMMU-31/2023** | EPI_ISL_18169211 | | The Second Military Medical University | The Second Military Medical University | Peng et al |
| **hCoV-19/Shanghai/SCDC-HG0531-1476/2023** | EPI_ISL_17795250 | | Microbe lab, Shanghai Municipal Center for Disease Control & Prevention | Shanghai Municipal Center for Disease Control and Prevention | Zhang et al B |
| **hCoV-19/Shanghai/SCDC-XG09-13776/2023** | EPI_ISL_18320786 | | Microbe lab, Shanghai Municipal Center for Disease Control & Prevention | Microbe lab, Shanghai Municipal Center for Disease Control & Prevention | Zhang et al B |
| **hCoV-19/India/WB-1931800284839/2023** | EPI_ISL_18001542 | | Calcutta School of Tropical Medicine | National Institute of Biomedical Genomics | Arindam Maitra et al A |
| **hCoV-19/India/TR-INSACOG-L23796/2023** | EPI_ISL_18112863 | | Department of Microbiology, Agartala Government Medical College | National Institute of Biomedical Genomics | Arindam Maitra et al A |
| **hCoV-19/India/JK-ICMR-NIV-G-14604/2023** | EPI_ISL_18120304 | | Indian Council of Medical Research, National Institute of Virology | National Institute of Virology - Influenza | Dr. Varsha Potdar et al |
| **hCoV-19/Oman/CPHL_7233030/2023** | EPI_ISL_18214445 | | Central Public Health Laboratories-Sultanat of Oman | Central Public Health Laboratories-Sultanat of Oman | Yathrib Al Zakwani et al |
| **hCoV-19/Tianjn/TJCDC0630-37/2023** | EPI_ISL_17956717 | | Department of Microbiology Detection, Tianjin Centers for Disease Control and Prevention | Tianjin Centers for Disease Control and Prevention | Li et al |
| **hCoV-19/Bhutan/RCDC0189/2023** | EPI_ISL_17967102 | | Royal Centre for Disease Control | Royal Centre for Disease Control | Sonam Gyeltshen et al |
| **hCoV-19/India/CT-ILS-GS21017/2023** | EPI_ISL_17730270 | | GMC, Raigarh | Institute of Life Sciences-INSACOG | Sunil K. Raghav et al |
| **hCoV-19/India/CT-ILS-GS20476/2023** | EPI_ISL_17730160 | | Pt. JNM Medical College, Raipur | Institute of Life Sciences-INSACOG | Sunil K. Raghav et al |
| **hCoV-19/India/JH-INSACOG-GBRC13517/2023** | EPI_ISL_17743149 | | Sterling Accuris Diagnostics | Gujarat Biotechnology Research Centre-INSACOG | Apurvasinh Puvar et al |
| **hCoV-19/Mexico/GUA_IBT_IMSS_11820/2023** | EPI_ISL_18589665 | | Centro de Investigacion Biomedica de Occidente (CIBO) | Instituto de Biotecnologia de la UNAM | Julio Elias Alvarado-Yaah et al |
| **hCoV-19/India/MH-ICMR-NIV-G-14640/2023** | EPI_ISL_18209941 | | ICMR-National Institute of Virology - INSACOG | NIV Influenza | Dr. Varsha Potdar et al |
| **hCoV-19/Nepal/NPHL-S-2429/2023** | EPI_ISL_17591976 | | PPHL-Lumbini | National Public Health Laboratory | Dr Runa Jha et al |
| **hCoV-19/Taiwan/CDC-W29126/2023** | EPI_ISL_18208077 | | Centers for Disease Control, R.O.C. (Taiwan) | Centers for Disease Control, R.O.C. (Taiwan) | Yu-Chi Lin et al |
| **hCoV-19/Singapore/R28MR34/2023** | EPI_ISL_17631445 | | National Public Health Laboratory, National Centre for Infectious Diseases | National Public Health Laboratory, National Centre for Infectious Diseases | Yichen Ding et al |
| **hCoV-19/Nepal/NPHL-S-2449/2023** | EPI_ISL_17620497 | | Trinagar Health Desk (POE) | National Public Health Laboratory | Dr. Runa Jha et al |
| **hCoV-19/Indonesia/JK-NIHRD-W03837/2023** | EPI_ISL_18639165 | | Laboratorium Econolab | National Institute of Health Research and Development | Hartanti Dian Ikawati et al |
| **hCoV-19/Philippines/PH-VSMMC-7-179/2023** | EPI_ISL_18355650 | | Vicente Sotto Memorial Medical Center (VSMMC) | Vicente Sotto Memorial Medical Center | Reynette Christine J. Ligaray et al |
| **hCoV-19/Brunei/MGS-7123050539/2023** | EPI_ISL_17989263 | | National Virology Reference Laboratory | Microbial Genomic Services | Zainun Zaini et al |
| **hCoV-19/Bangladesh/icddrb-G151023/2023** | EPI_ISL_17960110 | | Virology Laboratory, International Centre for Diarrhoeal Disease Research, Bangladesh (ICDDR,B) | Genome Centre, International Centre for Diarrhoeal Disease Research, Bangladesh (ICDDR,B) | Mohammad Jubair et al |
| **hCoV-19/India/CG-AIIMSR-VRDL-INSACOG-1324/2023** | EPI_ISL_17658967 | | State Virus Research and Diagnostic Laboratory (VRDL), Department of Microbiology, All India Institute of Medical Sciences | State Virus Research and Diagnostic Laboratory (VRDL), Department of Microbiology, All India Institute of Medical Sciences | Pushpendra Singh et al |
| **hCoV-19/India/TN-CDFD-A4-155/2023** | EPI_ISL_17714768 | | Directorate of Public Health and Preventive Medicine | CDFD-INSACOG | Asmita Gupta et al |
| **hCoV-19/India/SK-INSACOG-1122500144920/2023** | EPI_ISL_18113136 | | Sir Thutob Namgyal Memorial Hospital | National Institute of Biomedical Genomics | Arindam Maitra et al A |
| **hCoV-19/SouthAfrica/CERI-KRISP-K056064/2023** | EPI_ISL_17820248 | | NHLS | CERI, Centre for Epidemic Response and Innovation, Stellenbosch University and KRISP, KZN Research Innovation and Sequencing Platform, UKZN. | Giandhari J et al |
| **hCoV-19/Oman/CPHL_7232965/2023** | EPI_ISL_18214444 | | Central Public Health Laboratories-Sultanat of Oman | Central Public Health Laboratories-Sultanat of Oman | Yathrib Al Zakwani et al |
| **hCoV-19/Brunei/MGS-7523001162/2023** | EPI_ISL_17775931 | | National Virology Reference Laboratory | Microbial Genomic Services | Zainun Zaini et al |
| **hCoV-19/USA/MA-IVY-G43Q73X1/2023** | EPI_ISL_18433292 | | IVY5 Central Lab, Vanderbilt University Medical Center | Lauring Lab, University of Michigan, Department of Microbiology and Immunology | Baker et al |
| **hCoV-19/Jiangsu/YZCDC-222/2023** | EPI_ISL_17795740 | | Institute for the Prevention and Control of Acute Infectious Diseases | Institute for the Prevention and Control of Acute Infectious Diseases | Jianguang Fu et al |
| **hCoV-19/USA/NC-CDC-QDX83177818/2023** | EPI_ISL_18104186 | | Quest Diagnostics Incorporated | Centers for Disease Control and Prevention Division of Viral Diseases, Pathogen Discovery | Dakota Howard et al |
| **hCoV-19/Fujian/FCDC-FZLG02063/2023** | EPI_ISL_17801789 | | Institute for the Prevention and Control of Infectious Diseases, Fujian Provincial Center for Disease Control and Prevention | Institute for the Prevention and Control of Infectious Diseases, Fujian Provincial Center for Disease Control and Prevention | Qi Lin et al |
| **hCoV-19/Shanghai/SCDC-XG04-2653/2023** | EPI_ISL_17619243 | | Microbe lab, Shanghai Municipal Center for Disease Control & Prevention | Shanghai Municipal Center for Disease Control and Prevention | Zhang et al B |
| **hCoV-19/Singapore/R40MR67/2023** | EPI_ISL_17815712 | | National Public Health Laboratory, National Centre for Infectious Diseases | National Public Health Laboratory, National Centre for Infectious Diseases | Benny Yeo et al |
| **hCoV-19/Zhejiang/SKLID_ZJU-S118/2023** | EPI_ISL_17999262 | | State Key Laboratory for Diagnosis and Treatment of Infectious Diseases, The First Affiliated Hospital, Zhejiang University School of Medicine | State Key Laboratory for Diagnosis and Treatment of Infectious Diseases, The First Affiliated Hospital, Zhejiang University School of Medicine | Hangping Yao et al |
| **hCoV-19/Sichuan/LZCDC--84-WYL/2023** | EPI_ISL_17795032 | | Luzhou Center of Disease Control and Prevention | Microbiology laboratory, Sichuan Center of Disease Control and Prevention | Yu Ai et al |
| **hCoV-19/Jiangxi/JXNC-307/2023** | EPI_ISL_17794763 | | Nanchang Center for Disease Control and Prevention | Institute of Disease Control and Inspection, Jiangxi Province Center for Disease Control and Prevention | Yangbowen Wu et al |
| **hCoV-19/Hubei/HCDC-WH2401/2023** | EPI_ISL_18078309 | | Wuhan Center for Disease Control and Prevention | Hubei Provincial Center for Disease Control and Prevention Center of Pathogenic Microorganism Detection | Yongzhong Jiang et al |
| **hCoV-19/Jiangsu/WXCDC-396/2023** | EPI_ISL_18037379 | | Institute for the Prevention and Control of Acute Infectious Diseases, Jiangsu Provincial Center for Disease Control and Prevention | Institute for the Prevention and Control of Acute Infectious Diseases, Jiangsu Provincial Center for Disease Control and Prevention | Jianguang Fu et al |
| **hCoV-19/India/DL-ILBS-7900/2023** | EPI_ISL_17447400 | | MAHAJAN IMAGING | Institute of Liver and Biliary Sciences | Ekta Gupta et al |
| **hCoV-19/CaboVerde/ST-AG0128/2023** | EPI_ISL_18873725 | | Virology Laboratory of Praia | Instituto Nacional de Saúde Pública (INSP) | Kevin Sanders et al |
| **hCoV-19/India/MH-ICMR-NIV-G-14636/2023** | EPI_ISL_18209937 | | ICMR-National Institute of Virology - INSACOG | NIV Influenza | Dr. Varsha Potdar et al |
| **hCoV-19/CzechRepublic/NRL_s2097/2023** | EPI_ISL_18343543 | | National Institute of Public Health | National Institute of Public Health | Helena Jirincova et al |
| **hCoV-19/India/TR-INSACOG-812075099/2023** | EPI_ISL_18559588 | | INSACOG-Tripura | National Institute of Biomedical Genomics | Arindam Maitra et al A |
| **hCoV-19/India/TR-INSACOG-812434496/2023** | EPI_ISL_18559591 | | INSACOG-Tripura | National Institute of Biomedical Genomics | Arindam Maitra et al A |
| **hCoV-19/Canada/QC-L00662072001/2023** | EPI_ISL_18374595 | | Laboratoire de santé publique du Québec | Laboratoire de santé publique du Québec | Benjamin Delisle et al |
| **hCoV-19/Spain/CL-43859/2023** | EPI_ISL_18367985 | | Hospital Clínico Universitario de Valladolid (HCUV) | Hospital Clínico Universitario de Valladolid (HCUV) | Antonio Orduña-Domingo et al |
| **hCoV-19/Jamaica/184348/2023** | EPI_ISL_18871036 | | National Public Health Laboratory | The Caribbean Public Health Agency (CARPHA) | Nikhella Winter-Reece on behalf of CARPHA et al |
| **hCoV-19/Slovakia/ruvztn_05_00002645/2023** | EPI_ISL_18759802 | | Regional Public Health Authority with the seat in Trenčín, Slovakia | Laboratory of Genomics and Bioinformatics, Comenius University Science Park | Nikola Slivková et al |

| | | | | | |
|---|---|---|---|---|---|
| **hCoV-19/DominicanRepublic/2146631-LNSPDD/2023** | EPI_ISL_17955826 | | Laboratorio Nacional de Salud Pública Dr. Defilló | Laboratorio Nacional de Salud Pública Dr. Defilló | Isaac Miguel Sánchez et al |
| **hCoV-19/Oman/CPHL_7235492/2023** | EPI_ISL_18458172 | | Central Public Health Laboratories-Sultanat of Oman | Central Public Health Laboratories-Sultanat of Oman | Yathrib Al Zakwani et al |
| **hCoV-19/Bhutan/RCDC0194/2023** | EPI_ISL_18161250 | | Royal Centre for Disease Control | Royal Centre for Disease Control | Sonam Gyeltshen et al |
| **hCoV-19/Oman/CPHL_7232691/2023** | EPI_ISL_18214441 | | Central Public Health Laboratories-Sultanat of Oman | Central Public Health Laboratories-Sultanat of Oman | Yathrib Al Zakwani et al |
| **hCoV-19/Nepal/NPHL-S-2433/2023** | EPI_ISL_17591980 | | Central Diagnostic Laboratory | National Public Health Laboratory | Dr Runa Jha et al |
| **hCoV-19/India/JK-St6313/2023** | EPI_ISL_17554826 | | SKIMS-VRDL, Srinagar | NIV Influenza | Dr. Bashir Ahmad Fomda et al |
| **hCoV-19/Myanmar/DMR_171/2023** | EPI_ISL_18066554 | | Department of Medical Research | Department of Medical Research | Myat Htut Nyunt et al |
| **hCoV-19/India/DL-AIIMS149010/2023** | EPI_ISL_18873216 | | Virology Laboratory, AIIMS New Delhi | Virology Laboratory, AIIMS New Delhi | Megha Brijwal et al |
| **hCoV-19/India/DL-AIIMS-COVGE35962/2023** | EPI_ISL_17591373 | | COVID-CBNAAT; Department of Microbiology; AIIMS; New Delhi | Genomics Lab; CCRF; AIIMS; New Delhi_INSACOG | Urvashi B. Singh et al |
| **hCoV-19/India/UP-ICMR-NIV-G-14618/2023** | EPI_ISL_18120315 | | Indian Council of Medical Research, National Institute of Virology | National Institute of Virology - Influenza | Dr. Varsha Potdar et al |
| **hCoV-19/India/DL-AIIMS-COVGE35933/2023** | EPI_ISL_17591358 | | COVID-CBNAAT; Department of Microbiology; AIIMS; New Delhi | Genomics Lab; CCRF; AIIMS; New Delhi_INSACOG | Urvashi B. Singh et al |
| **hCoV-19/Mexico/GRO_InDRE_FB4143/2023** | EPI_ISL_18424285 | | LESP Guerrero | Instituto de Diagnostico y Referencia Epidemiologicos (INDRE) | Abril Rodriguez-Maldonado et al |
| **hCoV-19/India/WB-INSACOG-1931503455000/2023** | EPI_ISL_18559711 | | INSACOG-WB | National Institute of Biomedical Genomics | Arindam Maitra et al A |
| **hCoV-19/India/DL-AIIMS-COVGE35814/2023** | EPI_ISL_17591294 | | COVID-CBNAAT; Department of Microbiology; AIIMS; New Delhi | Genomics Lab; CCRF; AIIMS; New Delhi_INSACOG | Urvashi B. Singh et al |
| **hCoV-19/India/CT-AIIMSR-VRDL-INSACOG-1110/2023** | EPI_ISL_17466375 | | State Virus Research and Diagnostic Laboratory (VRDL), Department of Microbiology, All India Institute of Medical Sciences | State Virus Research and Diagnostic Laboratory (VRDL), Department of Microbiology, All India Institute of Medical Sciences | Pushpendra Singh et al |
| **hCoV-19/SouthKorea/KDCA249064/2023** | EPI_ISL_17983958 | | Division of Emerging Infectious Diseases, Bureau of Infectious Diseases Diagnosis Control, Korea Disease Control and Prevention Agency | Division of Emerging Infectious Diseases, Bureau of Infectious Diseases Diagnosis Control, Korea Disease Control and Prevention Agency | Jeong-Ah Kim et al |
| **hCoV-19/India/SK-ICMR-NIV-G-14592/2023** | EPI_ISL_18120294 | | Indian Council of Medical Research, National Institute of Virology | National Institute of Virology - Influenza | Dr. Varsha Potdar et al |
| **hCoV-19/India/DL-AIIMS-COVGE35837/2023** | EPI_ISL_17591306 | | COVID-CBNAAT; Department of Microbiology; AIIMS; New Delhi | Genomics Lab; CCRF; AIIMS; New Delhi_INSACOG | Urvashi B. Singh et al |
| **hCoV-19/Vietnam/HCMC-M170-S06/2023** | EPI_ISL_18115337 | | Oxford University Clinical Research Unt | Oxford University Clinical Research Unt | Nguyen Van Vinh Chau et al |
| **hCoV-19/Vietnam/HCMC-M177-S27/2023** | EPI_ISL_18115403 | | Oxford University Clinical Research Unt | Oxford University Clinical Research Unt | Nguyen Van Vinh Chau et al |
| **hCoV-19/Mauritius/37605/2023** | EPI_ISL_17849962 | | Central Health Laboratory, Ministry of Health and Wellness , Mauritius | Central Health Laboratory, Ministry of Health and Wellness , Mauritius | Ramuth M et al |
| **hCoV-19/CaboVerde/ST-AG0082/2023** | EPI_ISL_18058930 | | Virology Laboratory of Praia | Instituto Nacional de Saúde Pública (INSP) | Kevin Sanders et al |
| **hCoV-19/Myanmar/DMR_187/2023** | EPI_ISL_18397037 | | Department of Medical Research | Department of Medical Research | Myat Htut Nyunt et al |
| **hCoV-19/Myanmar/DMR_166/2023** | EPI_ISL_18066550 | | Department of Medical Research | Department of Medical Research | Myat Htut Nyunt et al |
| **hCoV-19/India/MZ-1526101122101/2023** | EPI_ISL_18001590 | | Zoram Medical College | National Institute of Biomedical Genomics | Arindam Maitra et al A |
| **hCoV-19/India/TN-CDFD-M3-187/2023** | EPI_ISL_17543952 | | Directorate of Public Health and Preventive Medicine | CDFD-INSACOG | Asmita Gupta et al |
| **hCoV-19/Vietnam/NHTD-OUCRU3807/2023** | EPI_ISL_17623788 | | National Hospital for Tropical Diseases | Oxford University Clinical Research Unit, Hanoi, Vietnam | Pham Ngoc Thach et al |
| **hCoV-19/USA/MI-UM-10051743912/2023** | EPI_ISL_18064441 | | University of Michigan Clinical Microbiology Laboratory | Lauring Lab, University of Michigan, Department of Microbiology and Immunology | Baker et al |
| **hCoV-19/India/JK-ICMR-NIV-G-14605/2023** | EPI_ISL_18120305 | | Indian Council of Medical Research, National Institute of Virology | National Institute of Virology - Influenza | Dr. Varsha Potdar et al |
| **hCoV-19/Bhutan/RCDC0191/2023** | EPI_ISL_18161247 | | Royal Centre for Disease Control | Royal Centre for Disease Control | Sonam Gyeltshen et al |
| **hCoV-19/India/UK-GMC-HLD-06-00239/2023** | EPI_ISL_18908441 | | Multidisciplinary Research Unit (MRU), GMC, Haldwani (UK) | NGS, GMC, Haldwani (UK) -INSACOG | Umesh et al |
| **hCoV-19/Australia/QLD0x01306B/2023** | EPI_ISL_17388899 | | Pathology Queensland and Forensic Scientific Services | PHV-FSS | Chenwei Wang on behalf of Q-PHIRE Genomics et al |
| **hCoV-19/India/JK-6/2023** | EPI_ISL_17554845 | | SKIMS-VRDL, Srinagar | NIV Influenza | Dr. Bashir Ahmad Fomda et al |
| **hCoV-19/Canada/MB-CPL-671237/2023** | EPI_ISL_18608047 | | MB-Cadham Provincial laboratory | Manitoba Cadham Provincial Laboratory | David Alexander et al |
| **hCoV-19/TrinidadandTobago/427293/2023** | EPI_ISL_17975852 | | Trinidad Public Health Laboratory | Carrington Lab, Department of Preclinical Sciences, Faculty of Medical Sciences, The University of the West Indies, St Augustine Campus | Nikita S. D. Sahadeo et al |
| **hCoV-19/TrinidadandTobago/UWI-429141/2023** | EPI_ISL_18139505 | | Trinidad Public Health Laboratory | Carrington Lab, Department of Preclinical Sciences, Faculty of Medical Sciences, The University of the West Indies, St Augustine Campus | Nikita S. D. Sahadeo et al |
| **hCoV-19/Greece/NPHO-6159/2023** | EPI_ISL_18096343 | | National Public Health Organization | National Public Health Organization | Kyriaki Tryfinopoulou et al |
| **hCoV-19/USA/NC-1085-9754/2023** | EPI_ISL_18153020 | | Carolinas HealthCare Systems Laboratory | Wake Forest School of Medicine, Internal Medicine | Kimberly Reeves et al |
| **hCoV-19/Mongolia/Uvs-2689/2023** | EPI_ISL_18788818 | | National Center for Communicable Diseases (NCCD) National Influenza Center | National Center for Communicable Diseases (NCCD) National Influenza Center | Bayasgalan Namuutsetseg et al |
| **hCoV-19/Mongolia/Uvs-1922/2023** | EPI_ISL_18788814 | | National Center for Communicable Diseases (NCCD) National Influenza Center | National Center for Communicable Diseases (NCCD) National Influenza Center | Bayasgalan Namuutsetseg et al |
| **hCoV-19/Canada/MB-CPL-629542/2023** | EPI_ISL_17743664 | | MB-Cadham Provincial laboratory | Manitoba Cadham Provincial Laboratory | David Alexander et al |
| **hCoV-19/India/UP-INSACOG-GBRC11677/2023** | EPI_ISL_17497233 | | Sterling Accuris Diagnostics,Ahmedabad | Gujarat Biotechnology Research Centre-INSACOG | Chaitanya G. Joshi et al |
| **hCoV-19/Xinjiang/XJCDCTC058/2023** | EPI_ISL_17775599 | | Xinjiang CDC | Xinjiang CDC | Ma et al |
| **hCoV-19/USA/KY-CDC-LC1040675/2023** | EPI_ISL_17653070 | | Laboratory Corporation of America | Centers for Disease Control and Prevention Division of Viral Diseases, Pathogen Discovery | Dakota Howard et al |
| **hCoV-19/Laos/IPL-78004/2023** | EPI_ISL_17664303 | | Institut Pasteur du Laos | Institut Pasteur du Laos | Thonglakhone XAYBOUNSOU et al |
| **hCoV-19/Laos/IPL-78010/2023** | EPI_ISL_17664304 | | Institut Pasteur du Laos | Institut Pasteur du Laos | Thonglakhone XAYBOUNSOU et al |
| **hCoV-19/India/MP-AIIMS_B-ICMR-2069/2023** | EPI_ISL_17521499 | | AIIMS Bhopal VRDL/Translational Medicine | AIIMS Bhopal VRDL/Translational Medicine | Anvita G. Malhotra et al |
| **hCoV-19/India/GJ-INSACOG-GBRC12359/2023** | EPI_ISL_17586850 | | Neuberg Labcore Speciality Laboratory - Vadodara | Gujarat Biotechnology Research Centre | Sonal Sharma et al |
| **hCoV-19/Cambodia/h0601082/2023** | EPI_ISL_17811441 | | Virology Unit, Institut Pasteur du Cambodge | Virology Unit, Institut Pasteur du Cambodge | Leakhena Pum et al |
| **hCoV-19/Cambodia/h0616044/2023** | EPI_ISL_17832697 | | Virology Unit, Institut Pasteur du Cambodge | Virology Unit, Institut Pasteur du Cambodge | Leakhena Pum et al |
| **hCoV-19/India/JK-SK-2-1043165/2023** | EPI_ISL_17554909 | | SKIMS-VRDL, Srinagar | NIV Influenza | Dr. Bashir Ahmad Fomda et al |
| **hCoV-19/Guangdong/GDCDC_SZHG_1777/2023** | EPI_ISL_18036355 | | Institute of Pathogenic Microbiology, Guangdong Provincial Center for Disease Control and Prevention | Institute of Pathogenic Microbiology, Guangdong Provincial Center for Disease Control and Prevention | Chang Zhang et al |
| **hCoV-19/Luxembourg/LNS0069147/2023** | EPI_ISL_18391715 | | BioneXt Lab | Laboratoire national de sante, Microbiology, Microbial Genomics Platform | Anke Wienecke-Baldacchino et al |
| **hCoV-19/Switzerland/VS-SNRCI-HUG-42304028/2023** | EPI_ISL_18209725 | | University Hospitals of Geneva, Laboratory of Virology | HUG, Laboratory of Virology and the Health2030 Genome Center | Samuel Cordey et al |
| **hCoV-19/Singapore/R64MR107/2023** | EPI_ISL_18243545 | | National Public Health Laboratory, National Centre for Infectious Diseases | National Public Health Laboratory, National Centre for Infectious Diseases | Grace Ngan et al |
| **hCoV-19/Mexico/SLP_IBT_IMSS_11943/2023** | EPI_ISL_18589553 | | Laboratorio Central de Epidemiologia (LCE) | Instituto de Biotecnologia de la UNAM | Julio Elias Alvarado-Yaah et al |
| **hCoV-19/India/AP-DIB-369965/2023** | EPI_ISL_18098406 | | Regional Virus Research and Diagnostic Laboratory, Indian Council of Medical Research, Regional Medical Research Centre | Regional Virus Research and Diagnostic Laboratory, Indian Council of Medical Research, Regional Medical Research Centre | B Borkakoty et al |
| **hCoV-19/Bhutan/RCDC0190/2023** | EPI_ISL_17967103 | | Royal Centre for Disease Control | Royal Centre for Disease Control | Sonam Gyeltshen et al |
| **hCoV-19/Sweden/AC-459929115118/2023** | EPI_ISL_17742426 | | The Public Health Agency of Sweden | The Public Health Agency of Sweden | Emmi Andersson et al |
| **hCoV-19/India/KA-ICMR-NIV-G-14576/2023** | EPI_ISL_18120281 | | Indian Council of Medical Research, National Institute of Virology | National Institute of Virology - Influenza | Dr. Varsha Potdar et al |
| **hCoV-19/USA/HI-H2323740/2023** | EPI_ISL_18111422 | | State Laboratories Division, Hawaii State Department of Health | State Laboratories Division, Hawaii State Department of Health | Pamela O'Brien et al |
| **hCoV-19/England/CLIMB-CM7YM4ND/2023** | EPI_ISL_18355728 | | Respiratory Virus Unit, Microbiology Services Colindale, Public Health England | UK Health Security Agency (UKHSA) | PHE Covid Sequencing Team et al |
| **hCoV-19/Brunei/MGS-7123075799/2023** | EPI_ISL_18866601 | | National Virology Reference Laboratory | Microbial Genomic Services | Zainun Zaini et al |
| **hCoV-19/Ireland/D-NVRL-GIRL52083/2023** | EPI_ISL_18259359 | | National Virus Reference Laboratory | National Virus Reference Laboratory | Zoe Yandle et al |
| **hCoV-19/Mexico/ZAC_InDRE_FB4752/2023** | EPI_ISL_18785784 | | Laboratorio Estatal de Salud Publica Zacatecas | Instituto de Diagnostico y Referencia Epidemiologicos (INDRE) | Abril Rodriguez-Maldonado et al |
| **hCoV-19/USA/IA-SHL-2356690/2023** | EPI_ISL_18475525 | | State Hygienic Laboratory at the University of Iowa | State Hygienic Laboratory at the University of Iowa | Valerie Reeb et al |
| **hCoV-19/USA/NY-URMC-N9S1L580-1/2023** | EPI_ISL_18666606 | | URMC LABS | URMC LABS | Dwight Hardy et al |
| **hCoV-19/Yunnan/YNCDC-1926/2023** | EPI_ISL_18074911 | | The department for prevention and control of acute infectious diseases, Yunnan Center for Disease Control and Prevention | The department for prevention and control of acute infectious diseases, Yunnan Center for Disease Control and Prevention | Zhang et al B |
| **hCoV-19/Gansu/GSCDC-0362/2023** | EPI_ISL_17998841 | | Gansu Provincial Pathogen Detection Institute, Gansu Provincial Centre for Disease Control and Prevention | Gansu Provincial Pathogen Detection Institute, Gansu Provincial Centre for Disease Control and Prevention | Li et al |
| **hCoV-19/Yunnan/YNCDC-1568/2023** | EPI_ISL_17966450 | | The department for prevention and control of acute infectious diseases, Yunnan Center for Disease Control and Prevention | The department for prevention and control of acute infectious diseases, Yunnan Center for Disease Control and Prevention | Zhang et al B |
| **hCoV-19/Curacao/CW-RIVM-130608/2023** | EPI_ISL_18073849 | | National Institute for Public Health and the Environment (RIVM) | National Institute for Public Health and the Environment (RIVM) | Adam Meijer et al |
| **hCoV-19/NewZealand/ESR-YA4497/2023** | EPI_ISL_18231751 | | LabPlus | Institute of Environmental Science and Research (ESR) | Sarah Cockerton et al |
| **hCoV-19/Ireland/D-CS0408/2023** | EPI_ISL_18221400 | | St. James's Hospital | St James's Hospital | Grier et al |
| **hCoV-19/Norway/8062/2023** | EPI_ISL_18075159 | | Department of Medical Microbiology - section Molde, Molde Hospital | Norwegian Institute of Public Health, Department of Virology | Kathrine Stene-Johansen et al |
| **hCoV-19/SouthAfrica/CERI-KRISP-K058083/2023** | EPI_ISL_18284170 | | PathCare | CERI, Centre for Epidemic Response and Innovation, Stellenbosch University and KRISP, KZN Research Innovation and Sequencing Platform, UKZN. | Maponga T et al |
| **hCoV-19/Laos/IPL-78436/2023** | EPI_ISL_18349883 | | NCLE | Institut Pasteur du Laos | Thonglakhone XAYBOUNSOU et al |
| **hCoV-19/Romania/220547/2023** | EPI_ISL_18932940 | | Cluj-Napoca Clinical Hospital for Infectious Diseases | National Institute of Public Health Laboratory | Mihaela Leustean et al |
| **hCoV-19/Cambodia/PKH230558/2023** | EPI_ISL_18633887 | | National Institute of Public Health | National Institute of Public Health | Chhorvann CHHEA et al |
| **hCoV-19/Tibet/XZCDC_0072/2023** | EPI_ISL_18602790 | | Inspection and verification office, Center for Disease Control and Prevention of Tibet Autonomous Region | Inspection and verification office, Center for Disease Control and Prevention of Tibet Autonomous Region | Hong et al |
| **hCoV-19/Moldova/Nr_30122/2023** | EPI_ISL_17817236 | | Virology Laboratory, NAPH, Moldova | Virology Laboratory, NAPH, Moldova | Olga Burduniuc et al |
| **hCoV-19/Georgia/Tb-s7957/2023** | EPI_ISL_18370627 | | National Center for Disease Control and Public Health (NCDC) of Georgia. | Department for Virology, Molecular Biology and Genome Research, R. G. Lugar Center for Public Health Research, National Center for Disease Control and Public Health (NCDC) of Georgia. | Giorgi Tomashvili et al |
| **hCoV-19/Georgia/Tb-s7760/2023** | EPI_ISL_18263773 | | Department for Virology, Molecular Biology and Genome Research, R. G. Lugar Center for Public Health Research, National Center for Disease Control and Public Health (NCDC) of Georgia. | Department for Virology, Molecular Biology and Genome Research, R. G. Lugar Center for Public Health Research, National Center for Disease Control and Public Health (NCDC) of Georgia. | Giorgi Tomashvili et al |
| **hCoV-19/Brunei/7123022982/2023** | EPI_ISL_17371559 | | National Virology Reference Laboratory | Microbial Genomic Services | Zainun Zaini et al |
| **hCoV-19/India/un-SPHL-1504/2023** | EPI_ISL_17757770 | | Public Health, State Public Health Laboratory Chennai | Public Health, State Public Health Laboratory Chennai | Raju et al |
| **hCoV-19/Bulgaria/23BG-EU_023240_PI215/2023** | EPI_ISL_17820867 | | National Center of Infectious and Parasitic Diseases | National Center of Infectious and Parasitic Diseases | Alexiev et al |
| **hCoV-19/Austria/AGES-1140473/2023** | EPI_ISL_17962938 | | Lifebrain COVID 19 Labor GmbH | AGES IMED Vienna | Polster E. et al |
| **hCoV-19/Thailand/MTM_01_1018/2023** | EPI_ISL_18472914 | | Clinical Trial Unit, Hospital for Tropical Diseases | Molecular Malaria Laboratory, Mahidol Oxford Tropical Medicine Research Unit | William HK Schilling et al |
| **hCoV-19/USA/NH-CDC-LC1042152/2023** | EPI_ISL_17853201 | | Laboratory Corporation of America | Centers for Disease Control and Prevention Division of Viral Diseases, Pathogen Discovery | Dakota Howard et al |
| **hCoV-19/Palestine/528391/2023** | EPI_ISL_17671798 | | CPHL-MOH | Molecular Diagnostic Laboraory-CPHL | Shtayeh Issa et al |
| **hCoV-19/Algeria/4759/2023** | EPI_ISL_18090022 | | NIC, Viral Respiratory Unit | NIC, Viral Respiratory Unit | Baya Hedjal et al |
| **hCoV-19/USA/SD-SDPHL-3015/2023** | EPI_ISL_18534929 | | South Dakota Public Health Laboratory | South Dakota Public Health Laboratory | Chris Carlson et al |
| **hCoV-19/Vietnam/HCMC-S06-M187/2023** | EPI_ISL_18688533 | | Oxford University Clinical Research Unt | Oxford University Clinical Research Unt | Nguyen Van Vinh Chau et al |
| **hCoV-19/Singapore/R75MR71/2023** | EPI_ISL_18428850 | | National Public Health Laboratory, National Centre for Infectious Diseases | National Public Health Laboratory, National Centre for Infectious Diseases | Yichen Ding et al |
| **hCoV-19/Ecuador/GENSBIO-INSPI-1820-2114/2023** | EPI_ISL_18646565 | | Laboratorio de Influenza y Otros Virus Respiratorios, INSPI-CZ9 | Centro de Referencia Nacional de Genomica, Secuenciacion y Bioinformatica GENSBIO, INSPI-CZ9 | Andres Carrazco-Montalvo* et al |
| **hCoV-19/India/GJ-GBRC10569/2023** | EPI_ISL_17270034 | | Apollo Hospital, Ahmedabad | Gujarat Biotechnology Research Centre | Rameshchandra Pandit et al |
| **hCoV-19/Chile/VS-48506/2023** | EPI_ISL_18335165 | | Genetica Molecular and Subdepartamento de Virologia ISP Chile | Instituto de Salud Publica de Chile | Mariela Guajardo et al |
| **hCoV-19/Heilongjiang/HLJCDC-1063/2023** | EPI_ISL_17645626 | | Institute for Viral Disease Control and Prevention, Department for Viral Disease Control and Prevention, Heilongjiang Provincial Center for Disease Control and Prevention | Institute for Viral Disease Control and Prevention, Department for Viral Disease Control and Prevention, Heilongjiang Provincial Center for Disease Control and Prevention | Xu Jun et al |
| **hCoV-19/Vietnam/NHTD-OUCRU3827/2023** | EPI_ISL_17623806 | | National Hospital for Tropical Diseases | Oxford University Clinical Research Unit, Hanoi, Vietnam | Pham Ngoc Thach et al |
| **hCoV-19/SouthKorea/KDCA-252115/2023** | EPI_ISL_18039155 | | Division of Infectious Disease Diagnosis Control, Gyeongnam Regional Center for Disease Control and Prevention, Korea Disease Control and Prevention Agency | Division of Emerging Infectious Diseases, Bureau of Infectious Diseases Diagnosis Control, Korea Disease Control and Prevention Agency | Sungdo Park et al |
| **hCoV-19/Philippines/PH-PGCMIN3305/2023** | EPI_ISL_17948928 | | Southern Philippines Medical Center | Philippine Genome Center Mindanao | Joel Hassan G. Tolentino et al |
| **hCoV-19/Canada/AB-ABPHL-108337/2023** | EPI_ISL_18538918 | | Alberta Precision Labs (APL) | Alberta Precision Labs (APL) | Bu J et al |
| **hCoV-19/Chile/CO-75257/2023** | EPI_ISL_18501353 | | Genetica Molecular and Subdepartamento de Virologia ISP Chile | Instituto de Salud Publica de Chile | Mariela Guajardo et al |
| **hCoV-19/USA/MO-CDC-QDX86356416/2023** | EPI_ISL_18532523 | | Quest Diagnostics Incorporated | Centers for Disease Control and Prevention Division of Viral Diseases, Pathogen Discovery | Dakota Howard et al |
| **hCoV-19/Georgia/Tb-s8081/2023** | EPI_ISL_18413151 | | Department for Virology, Molecular Biology and Genome Research, R. G. Lugar Center for Public Health Research, National Center for Disease Control and Public Health (NCDC) of Georgia. | Department for Virology, Molecular Biology and Genome Research, R. G. Lugar Center for Public Health Research, National Center for Disease Control and Public Health (NCDC) of Georgia. | Giorgi Tomashvili et al |
| **hCoV-19/Georgia/Tb-s8133/2023** | EPI_ISL_18652801 | | Department for Virology, Molecular Biology and Genome Research, R. G. Lugar Center for Public Health Research, National Center for Disease Control and Public Health (NCDC) of Georgia. | Department for Virology, Molecular Biology and Genome Research, R. G. Lugar Center for Public Health Research, National Center for Disease Control and Public Health (NCDC) of Georgia. | Giorgi Tomashvili et al |
| **hCoV-19/Palestine/532679/2023** | EPI_ISL_18218364 | | CPHL-MOH | Molecular Diagnostic Laboraory-CPHL | Shtayeh Issa et al |
| **hCoV-19/Kuwait/Jaber-20536650/2023** | EPI_ISL_18694831 | | Jaber Al Ahmad Hospital | Virology Lab, Jaber Al Ahmad Hospital | Dr. Zahrah Buhamad et al |
| **hCoV-19/USA/MA-CDC-QDX82903570/2023** | EPI_ISL_18072571 | | Quest Diagnostics Incorporated | Centers for Disease Control and Prevention Division of Viral Diseases, Pathogen Discovery | Dakota Howard et al |
| **hCoV-19/Luxembourg/LNS2549061/2023** | EPI_ISL_18391701 | | Hospital Center Emile Mayrisch | Laboratoire national de sante, Microbiology, Microbial Genomics Platform | Anke Wienecke-Baldacchino et al |
| **hCoV-19/DominicanRepublic/2158061-LNSPDD/2023** | EPI_ISL_18077218 | | Laboratorio Nacional de Salud Pública Dr. Defilló | Laboratorio Nacional de Salud Pública Dr. Defilló | Isaac Miguel Sánchez et al |
| **hCoV-19/DominicanRepublic/2160977-LNSPDD/2023** | EPI_ISL_18281823 | | Laboratorio Nacional de Salud Pública Dr. Defilló | Laboratorio Nacional de Salud Pública Dr. Defilló | Isaac Miguel Sánchez et al |
| **hCoV-19/DominicanRepublic/2151610-LNSPDD/2023** | EPI_ISL_17976090 | | Laboratorio Nacional de Salud Pública Dr. Defilló | Laboratorio Nacional de Salud Pública Dr. Defilló | Isaac Miguel Sánchez et al |
| **hCoV-19/Singapore/R27AMR150/2023** | EPI_ISL_17596298 | | National Public Health Laboratory, National Centre for Infectious Diseases | National Public Health Laboratory, National Centre for Infectious Diseases | Grace Ngan et al |
| **hCoV-19/Hubei/HCDC-JM1996/2023** | EPI_ISL_17801825 | | Jingmen Center for Diaease Control and Prevention | Hubei Provincial Center for Disease Control and Prevention Center of Pathogenic Microorganism Detection | Yongzhong Jiang et al |
| **hCoV-19/Bangladesh/ideSHi-IR-3444/2023** | EPI_ISL_16528802 | | International Centre for Diarrhoeal Disease Research, Bangladesh (icddr,b) | Institute for Developing Science and Health Initiatives (ideSHi) | Rahman S et al |
| **hCoV-19/Singapore/R15MR83/2023** | EPI_ISL_17016364 | | National Public Health Laboratory, National Centre for Infectious Diseases | National Public Health Laboratory, National Centre for Infectious Diseases | Benny Yeo et al |
| **hCoV-19/Bahrain/PHD-6346/2023** | EPI_ISL_17744961 | | Communicable Disease Laboratory, Public Health Directorate | Communicable Disease Laboratory, Public Health Directorate | Marhoon et al |
| **hCoV-19/Bahrain/CDL-23970020609/2023** | EPI_ISL_17081351 | | Communicable Disease Laboratory | Communicable Disease Laboratory | Marhoon et al |

| | | | | | |
|---|---|---|---|---|---|
| **hCoV-19/Henan/NYCDC-SDCX146/2023** | EPI_ISL_17696929 | | Nanzhao County People's Hospital | Clinical Laboratory, Nanyang City Center for Disease Control and Prevention | Shi Rui et al |
| **hCoV-19/Laos/IPL-78043/2023** | EPI_ISL_17785486 | | National Center for Laboratory and Epidemiology (NCLE) | Institut Pasteur du Laos | Thonglakhone XAYBOUNSOU et al |
| **hCoV-19/Kuwait/Jaber-20438447/2023** | EPI_ISL_18118632 | | Farwaniya Hospital | Virology Lab, Jaber Al Ahmad Hospital | Dr. Zahrah Buhamad et al |
| **hCoV-19/Henan/XYCDC-0183/2023** | EPI_ISL_18040227 | | Central laboratory, Xinyang Center for Disease Control and Prevention | Central laboratory, Xinyang Center for Disease Control and Prevention | Lv et al |
| **hCoV-19/Egypt/CPHL-EGY24276/2023** | EPI_ISL_18138701 | | CPHL,MOH,EGYPT | CPHL,MOH,EGYPT | Wael H. Roshdy et al |
| **hCoV-19/India/DL-AIIMS-COVGE35734/2023** | EPI_ISL_17591307 | | COVID-CBNAAT; Department of Microbiology; AIIMS; New Delhi | Genomics Lab; CCRF; AIIMS; New Delhi_INSACOG | Urvashi B. Singh et al |
| **hCoV-19/Georgia/Tb-s7620/2023** | EPI_ISL_17876308 | | Department for Virology, Molecular Biology and Genome Research, R. G. Lugar Center for Public Health Research, National Center for Disease Control and Public Health (NCDC) of Georgia. | Department for Virology, Molecular Biology and Genome Research, R. G. Lugar Center for Public Health Research, National Center for Disease Control and Public Health (NCDC) of Georgia. | Nino Berishvili et al |
| **hCoV-19/Kenya/2022024957/2023** | EPI_ISL_18516355 | | DLSP KEMRI/CDC NAIROBI Lab | DLSP KEMRI/CDC NAIROBI Lab | Lydia Mwasi et al |
| **hCoV-19/Fujian/FCDC-XM071203/2023** | EPI_ISL_17999103 | | Institute for the Prevention and Control of Infectious Diseases, Fujian Provincial Center for Disease Control and Prevention, No.386 Chongan Road, Fuzhou, Fujian 350012, China | Institute for the Prevention and Control of Infectious Diseases, Fujian Provincial Center for Disease Control and Prevention, No.386 Chongan Road, Fuzhou, Fujian 350012, China | Qi Lin et al |
| **hCoV-19/Egypt/CPHL-EGY24240/2023** | EPI_ISL_18138668 | | CPHL,MOH,EGYPT | CPHL,MOH,EGYPT | Wael H. Roshdy et al |
| **hCoV-19/Qatar/NIC-10-VI-0099079/2023** | EPI_ISL_18399084 | | Section of Virology and Molecular Biology, Department of Laboratory Medicine and Pathology, Hamad Medical Corporation | Section of Virology and Molecular Biology, Department of Laboratory Medicine and Pathology, Hamad Medical Corporation | Dr. Mohamed Ali Ben Hadj Kacem et al |
| **hCoV-19/Qatar/NIC-25-VI-0118109/2023** | EPI_ISL_18594240 | | Section of Virology and Molecular Biology, Department of Laboratory Medicine and Pathology, Hamad Medical Corporation | Section of Virology and Molecular Biology, Department of Laboratory Medicine and Pathology, Hamad Medical Corporation | Dr. Mohamed Ali Ben Hadj Kacem et al |
| **hCoV-19/Tunisia/234687/2023** | EPI_ISL_18226720 | | Laboratory of Microbiology, La Rabta Hospital | 1-Laboratory of Microbiology, National Reference Lab, Charles Nicolle Hospital; 2-University of Tunis ElManar, Faculty of Medicine of Tunis, LR99ES09, Tunis, Tunisia | Zaineb Hamzaoui et al |
| **hCoV-19/Iceland/L-4695/2023** | EPI_ISL_18439619 | | Landspitali Department of Clinical Microbiology | Landspitali Department of Clinical Microbiology | Freyja Valsdottir et al |
| **hCoV-19/Cambodia/AHC230441/2023** | EPI_ISL_18633631 | | National Institute of Public Health | National Institute of Public Health | Chhorvann CHHEA et al |
| **hCoV-19/Ukraine/79935/2023** | EPI_ISL_18008341 | | Reference laboratory for diagnostics of HIV/AIDS, virological and especially dangerous pathogens | Reference laboratory for diagnostics of HIV/AIDS, virological and especially dangerous pathogens | Iryna Demchyshyna et al |
| **hCoV-19/Cambodia/940021/2023** | EPI_ISL_17736203 | | Virology Unit, Institut Pasteur du Cambodge | Virology Unit, Institut Pasteur du Cambodge | Leakhena Pum et al |
| **hCoV-19/Xinjiang/XJCDCHM_183/2023** | EPI_ISL_18108747 | | Xinjiang center of diease control | Xinjiang center of diease control | Ma et al |
| **hCoV-19/Jilin/JSY-CC8/2023** | EPI_ISL_18908494 | | Changchun Veterinary Research Institute, Chinese Academy of Agricultural Sciences | Changchun Veterinary Research Institute, Chinese Academy of Agricultural Sciences | Yuewei Gao et al |
| **hCoV-19/Malaysia/MGVI_GU0963/2023** | EPI_ISL_17722367 | | Hospital Selayang | Malaysia Genome and Vaccine Institute | Mohd Noor Mat Isa et al |
| **hCoV-19/BosniaandHerzegovina/AGCI847-23/2023** | EPI_ISL_17300879 | | Alea Genetic Centre | Alea Genetic Centre | I Ceko et al |
| **hCoV-19/Oman/723685/2023** | EPI_ISL_17026469 | | Central Public Health Laboratories-Sultanat of Oman | Central Public Health Laboratories-Sultanat of Oman | Yathrib Al Zakwani et al |
| **hCoV-19/Anhui/FYCDC-0284/2023** | EPI_ISL_17744606 | | Disease Control Laboratory, Fuyang Center for Diseases Prevention and Control | Disease Control Laboratory, Fuyang Center for Diseases Prevention and Control | Li et al |
| **hCoV-19/Gansu/QYCDC-813/2023** | EPI_ISL_18495746 | | Central Laboratory, Qingyang Center for Disease Control and Prevention | Central Laboratory, Qingyang Center for Disease Control and Prevention | Zhu et al |
| **hCoV-19/Russia/TYU-RII-MH144144S/2023** | EPI_ISL_18299840 | | Tyumen Regional Hospital No 19 | WHO National Influenza Centre Russian Federation | Andrey Komissarov et al |
| **hCoV-19/Romania/IS-2896/2023** | EPI_ISL_18261945 | | Department of Public Health Iasi | National Institute of Infectious Diseases-Prof. Dr. Matei Bals Molecular Diagnostics Laboratory | Simona Paraschiv et al |
| **hCoV-19/Guangdong/GDCDC_SZ_0829/2023** | EPI_ISL_18035067 | | Institute of Pathogenic Microbiology, Guangdong Provincial Center for Disease Control and Prevention | Institute of Pathogenic Microbiology, Guangdong Provincial Center for Disease Control and Prevention | Chang Zhang et al |
| **hCoV-19/Fujian/FCDC-FZLG02525/2023** | EPI_ISL_17850174 | | Institute for the Prevention and Control of Infectious Diseases, Fujian Provincial Center for Disease Control and Prevention | Institute for the Prevention and Control of Infectious Diseases, Fujian Provincial Center for Disease Control and Prevention | Qi Lin et al |
| **hCoV-19/Hubei/HCDC-JZ1725/2023** | EPI_ISL_17729928 | | Jingzhou Center for Disease Control and Prevention | Hubei Provincial Center for Disease Control and Prevention Center of Pathogenic Microorganism Detection | Yongzhong Jiang et al |
| **hCoV-19/Zhejiang/SKLID_ZJU-33/2023** | EPI_ISL_17714258 | | State Key Laboratory for Diagnosis and Treatment of Infectious Diseases, The First Affiliated Hospital, Zhejiang University School of Medicine | State Key Laboratory for Diagnosis and Treatment of Infectious Diseases, The First Affiliated Hospital, Zhejiang University School of Medicine | Hangping Yao et al |
| **hCoV-19/Ireland/WW-NVRL-SIRL00014507/2023** | EPI_ISL_17376004 | | National Virus Reference Laboratory | National Virus Reference Laboratory | Zoe Yandle et al |
| **hCoV-19/Finland/THL-07332/2023** | EPI_ISL_18347910 | | Nordlab Oulu | Expert Microbiology, National Institute for Health and Welfare | Erika Lindh et al |
| **hCoV-19/Denmark/DCGC-647448/2023** | EPI_ISL_17975282 | | Molekylær Medicinsk Afdeling, Aarhus University Hospital, Aarhus, Denmark | Statens Serum Institut Bioinformatics and Microbial Genomics | Povilas Matusevicius on behalf of the Danish Covid-19 Genome Consortium et al |
| **hCoV-19/Iraq/NRC-T037N/2023** | EPI_ISL_18858302 | | Al Hussein Teaching Hospital | Center of Scientific Excellence for Influenza Viruses (CSEIV),National Research Centre (NRC) | Ahmed.A.Al-Mankhee et al |
| **hCoV-19/HongKong/VB23020350/2023** | EPI_ISL_17433548 | | Kwong Wah Hospital | Hong Kong Department of Health | Alan K.L. Tsang et al |
| **hCoV-19/Romania/IS-CNMMI-549387/2023** | EPI_ISL_17419337 | | Cantacuzino National Military-Medical Institute, Viral Respiratory Infections Laboratory | Cantacuzino National Military-Medical Institute, Viral Respiratory Infections Laboratory | Catalina Pascu et al |
| **hCoV-19/Turkey/HSGM-AHSL-42/2023** | EPI_ISL_18422134 | | Ministry of Health Turkey, Adana Public Health Lab. | Ministry of Health Türkiye, Public Health Directorate, National Virology Reference Laboratory | Süleyman Yalcin et al |
| **hCoV-19/Jiangxi/JXFZ-019/2023** | EPI_ISL_17794268 | | Fuzhou Center for Disease Control and Prevention | Institute of Disease Control and Inspection, Jiangxi Province Center for Disease Control and Prevention | Yangbowen Wu et al |
| **hCoV-19/Beijing/BJCDC-2244/2023** | EPI_ISL_18049434 | | Institute for Infectious Disease and Endemic Disease Control, Beijing Center for Disease Control and Prevention | Institute for Infectious Disease and Endemic Disease Control, Beijing Center for Disease Control and Prevention | Pan et al |
| **hCoV-19/Gansu/QYCDC-0483/2023** | EPI_ISL_18495885 | | Central Laboratory, Qingyang Center for Disease Control and Prevention | Central Laboratory, Qingyang Center for Disease Control and Prevention | Zhu et al |
| **hCoV-19/Norway/6167/2023** | EPI_ISL_17711195 | | Haugesund Hospital, laboratory for Medical Microbiology | Norwegian Institute of Public Health, Department of Virology | Kathrine Stene-Johansen et al |
| **hCoV-19/Russia/MAG-RII-MH138568S/2023** | EPI_ISL_17665556 | | Magadan Regional Dispensary of Phthisiology and Infectious Diseases | WHO National Influenza Centre Russian Federation | Andrey Komissarov et al |
| **hCoV-19/Norway/6637/2023** | EPI_ISL_17770584 | | Unilabs Laboratory Medicine | Norwegian Institute of Public Health, Department of Virology | Kathrine Stene-Johansen et al |
| **hCoV-19/Paraguay/LERAP_23-1/2023** | EPI_ISL_18396027 | | Laboratorio Epidemiológico Regional Alto Paraná | Laboratorio Epidemiológico Regional Alto Paraná | Daniela Poullion et al |
| **hCoV-19/Georgia/Tb-s7678/2023** | EPI_ISL_17885361 | | Department for Virology, Molecular Biology and Genome Research, R. G. Lugar Center for Public Health Research, National Center for Disease Control and Public Health (NCDC) of Georgia. | Department for Virology, Molecular Biology and Genome Research, R. G. Lugar Center for Public Health Research, National Center for Disease Control and Public Health (NCDC) of Georgia. | Giorgi Tomashvili et al |
| **hCoV-19/Xinjiang/XJCDCBOZ-045/2023** | EPI_ISL_17836367 | | Xinjiang center of diease control | Xinjiang center of diease control | Ma et al |
| **hCoV-19/Iran/EF1013/2023** | EPI_ISL_18443024 | | Kermanshah University of Medical Sciences | COVID-19 National Reference Laboratory, Pasteur Institute of Iran | Zahra Ahmadi et al |
| **hCoV-19/Ukraine/80106/2023** | EPI_ISL_18274888 | | Reference laboratory for diagnostics of HIV/AIDS, virological and especially dangerous pathogens | Reference laboratory for diagnostics of HIV/AIDS, virological and especially dangerous pathogens | Iryna Demchyshyna et al |
| **hCoV-19/Beijing/BJCDC-3650/2023** | EPI_ISL_18438264 | | Institute for Infectious Disease and Endemic Disease Control, Beijing Center for Disease Control and Prevention | Institute for Infectious Disease and Endemic Disease Control, Beijing Center for Disease Control and Prevention | Pan et al |
| **hCoV-19/Algeria/44064/2022** | EPI_ISL_16242284 | | NIC, Viral Respiratory Unit | NIC, Viral Respiratory Unit | Baya Hedjal et al |
| **hCoV-19/Mozambique/INS-LSPT0217/2023** | EPI_ISL_17510594 | | Biotechnology and Genetics Laboratory - Instituto Nacional de Saúde | Biotechnology and Genetics Laboratory - Instituto Nacional de Saúde | Sofia Viegas et al |
| **hCoV-19/Egypt/CPHL-23601/2023** | EPI_ISL_18115519 | | Central Public Health Laboratories, Ministry of Health and Population | Central Public Health Laboratories, Ministry of Health and Population | Wael H. Roshdy et al |
| **hCoV-19/Egypt/CPHL-EGY23992/2023** | EPI_ISL_18138786 | | CPHL,MOH,EGYPT | CPHL,MOH,EGYPT | Wael H. Roshdy et al |
| **hCoV-19/env/Germany/BY-LGL-6016165/2023** | EPI_ISL_18412795 | | Bayerisches Landesamt fuer Gesundheit und Lebensmittelsicherheit (LGL) | Bayerisches Landesamt fuer Gesundheit und Lebensmittelsicherheit (LGL) | Alexandra Dangel et al |
| **hCoV-19/Lebanon/HH-2872/2023** | EPI_ISL_18385970 | | Hammoud Hospital University Medical Center (HHUMC) | Center for Infectious Diseases Research (CIDR)- The Global Influenza Hospital Surveillance Network (GIHSN) | Ghassan Dbaibo et al |
| **hCoV-19/Xinjiang/XJCDCBAZ-156/2023** | EPI_ISL_17998932 | | Xinjiang CDC | Xinjiang CDC | Ma et al |
| **hCoV-19/Jiangsu/JCDC-249/2023** | EPI_ISL_17505630 | | Jiangsu Provincial Center for Disease Control and Prevention | Jiangsu Provincial Center for Disease Control and Prevention | Fan et al |
| **hCoV-19/India/DL-ILBS-7476/2023** | EPI_ISL_17483686 | | LIFELINE LABORATORY | ILBS/INSACOG | Ekta Gupta et al |
| **hCoV-19/Indonesia/JT-BBTKLPP-W191-S77B4/2023** | EPI_ISL_17704093 | | Laboratorium RS Soeradji Klaten | BBTKLPP Yogyakarta | Havid Setyawan et al |
| **hCoV-19/India/KL-ICMR-NIV-G-14575/2023** | EPI_ISL_18120280 | | Indian Council of Medical Research, National Institute of Virology | National Institute of Virology - Influenza | Dr. Varsha Potdar et al |
| **hCoV-19/Jiangxi/JXPX-037/2023** | EPI_ISL_17794783 | | Jiangxi Province Center for Disease Control and Prevention | Institute of Disease Control and Inspection, Jiangxi Province Center for Disease Control and Prevention | Yangbowen Wu et al |
| **hCoV-19/Tunisia/7054/2023** | EPI_ISL_18110949 | | 1-Laboratory of Microbiology, National Reference Lab, Charles Nicolle Hospital; 2-University of Tunis ElManar, Faculty of Medicine of Tunis, LR99ES09, Tunis, Tunisia | Clinical and Experimental Pharmacology Lab, LR16SP02, National Center of Pharmacovigilance, University of Tunis El Manar, Tunis, Tunisia. 2- Ministry of Health, National Observatory of New and Emerging Diseases, 1006, Tunis, Tunisia | Zaineb Hamzaoui et al |
| **hCoV-19/Tunisia/9091/2023** | EPI_ISL_18762496 | | 1-Laboratory of Microbiology, National Reference Lab, Charles Nicolle Hospital; 2-University of Tunis ElManar, Faculty of Medicine of Tunis, LR99ES09, Tunis, Tunisia | Clinical and Experimental Pharmacology Lab, LR16SP02, National Center of Pharmacovigilance, University of Tunis El Manar, Tunis, Tunisia. 2- Ministry of Health, National Observatory of New and Emerging Diseases, 1006, Tunis, Tunisia | Zaineb Hamzaoui et al |
| **hCoV-19/Egypt/CPHL-EGY23888/2023** | EPI_ISL_18138977 | | CPHL,MOH,EGYPT | CPHL,MOH,EGYPT | Wael H. Roshdy et al |
| **hCoV-19/Kuwait/AH-USAFSAM-S21326/2023** | EPI_ISL_17501437 | | US Air Force School of Aerospace Medicine | US Air Force School of Aerospace Medicine | Anthony Fries et al |
| **hCoV-19/Greece/NPHO-5733/2023** | EPI_ISL_18096070 | | National Public Health Organization | National Public Health Organization | Kyriaki Tryfinopoulou et al |
| **hCoV-19/India/WB-1930302179883/2023** | EPI_ISL_17812503 | | INSACOG-WB | National Institute of Biomedical Genomics - INSACOG | Arindam Maitra et al A |
| **hCoV-19/Guadeloupe/IPP13611/2023** | EPI_ISL_17705299 | | Hopital | National Reference Center for Viruses of Respiratory Infections, Institut Pasteur, Paris | Léa Avon et al |
| **hCoV-19/Portugal/PT48683/2023** | EPI_ISL_17954734 | | CHU Sao Joao, Porto | Instituto Nacional de Saude Doutor Ricardo Jorge (INSA) | Borges et al |
| **hCoV-19/Bangladesh/IEDCR-2200771/2023** | EPI_ISL_17837239 | | Institute of Epidemiology, Disease Control and Research (IEDCR) | Institute of Epidemiology, Disease Control and Research (IEDCR) | Manjur Hossain Khan et al |
| **hCoV-19/SouthAfrica/NICD-CRDM06622/2023** | EPI_ISL_17885137 | | AMPATH | National Institute for Communicable Diseases of the National Health Laboratory Service | Everatt J et al |
| **hCoV-19/Malaysia/C19UMB4347/2023** | EPI_ISL_18432668 | | Hospital Melaka | UKM Medical Molecular Biology Institute (UMBI) | Rahman Jamal et al |
| **hCoV-19/Taiwan/CDC-F87125/2023** | EPI_ISL_18164780 | | Centers for Disease Control, R.O.C. (Taiwan) | Centers for Disease Control, R.O.C. (Taiwan) | Yu-Chi Lin et al |
| **hCoV-19/SriLanka/aicbu6446/2023** | EPI_ISL_17699453 | | Allergy, Immunology and Cell Biology Unit (AICBU) | Allergy, Immunology and Cell Biology Unit (AICBU) | Chandima Jeewandara et al |
| **hCoV-19/Shanghai/SCDC-HG1019-3824/2023** | EPI_ISL_18475252 | | Microbe lab, Shanghai Municipal Center for Disease Control & Prevention | Microbe lab, Shanghai Municipal Center for Disease Control & Prevention | Chen et al |
| **hCoV-19/Ireland/LK-UHL-674002/2023** | EPI_ISL_18549620 | | University Hospital Limerick | Molecular Laboratory, Serology and Virology at Pathology Department | Diane Slater-Kinghorn et al |
| **hCoV-19/Ukraine/79743/2023** | EPI_ISL_17982490 | | Reference laboratory for diagnostics of HIV/AIDS, virological and especially dangerous pathogens | Reference laboratory for diagnostics of HIV/AIDS, virological and especially dangerous pathogens | Iryna Demchyshyna et al |
| **hCoV-19/Slovenia/17728/2023** | EPI_ISL_18377028 | | Institute of Microbiology and Immunology, Faculty of Medicine, University of Ljubljana | Institute of Microbiology and Immunology, Faculty of Medicine, University of Ljubljana | Alen Suljič et al |
| **hCoV-19/Bulgaria/23BG-NC_023680_R41/2023** | EPI_ISL_18452036 | | National Center of Infectious and Parasitic Diseases | National Center of Infectious and Parasitic Diseases | Ivelina Trifonova et al |
| **hCoV-19/Uruguay/DLSP_HC1/2023** | EPI_ISL_18899089 | | Departamento Laboratorios de Salud Pública (DLSP) Ministerio de Salud Pública | Departamento Laboratorios de Salud Pública (DLSP) Ministerio de Salud Pública | Cortinas et al |
| **hCoV-19/India/DL-ILBS-7564/2023** | EPI_ISL_17268954 | | Dr. Dangs Lab | Institute of Liver and Biliary Sciences (ILBS) | Ekta Gupta et al |
| **hCoV-19/Colombia/DC-INS-VG-30620/2023** | EPI_ISL_18299105 | | SYNLAB COLOMBIA | Instituto Nacional de Salud- Dirección de Investigación en Salud Pública | Katherine Laiton-Donato et al |
| **hCoV-19/Guangdong/GDCDC_SZ_0345/2023** | EPI_ISL_17697704 | | Institute of Pathogenic Microbiology, Guangdong Provincial Center for Disease Control and Prevention | Institute of Pathogenic Microbiology, Guangdong Provincial Center for Disease Control and Prevention | Chang Zhang et al |
| **hCoV-19/Taiwan/CDC-F88476/2023** | EPI_ISL_18488684 | | Centers for Disease Control, R.O.C. (Taiwan) | Centers for Disease Control, R.O.C. (Taiwan) | Yu-Chi Lin et al |
| **hCoV-19/Singapore/R71MR134/2023** | EPI_ISL_18361891 | | National Public Health Laboratory, National Centre for Infectious Diseases | National Public Health Laboratory, National Centre for Infectious Diseases | Grace Ngan et al |
| **hCoV-19/SouthKorea/KDCA254781/2023** | EPI_ISL_18068227 | | Division of Emerging Infectious Diseases, Bureau of Infectious Diseases Diagnosis Control, Korea Disease Control and Prevention Agency | Division of Emerging Infectious Diseases, Bureau of Infectious Diseases Diagnosis Control, Korea Disease Control and Prevention Agency | Jeong-Ah Kim et al |
| **hCoV-19/Portugal/PT48682/2023** | EPI_ISL_17954733 | | CHU Sao Joao, Porto | Instituto Nacional de Saude Doutor Ricardo Jorge (INSA) | Borges et al |
| **hCoV-19/India/KA-ICMR-NIV-G-14577/2023** | EPI_ISL_18120282 | | Indian Council of Medical Research, National Institute of Virology | National Institute of Virology - Influenza | Dr. Varsha Potdar et al |
| **hCoV-19/Shanxi/TYCDC535/2023** | EPI_ISL_18495466 | | Microbiology Department, Taiyuan Center for Disease Control and Prevention | Microbiology Department, Taiyuan Center for Disease Control and Prevention | Wang et al |
| **hCoV-19/Oman/CPHL_7233915/2023** | EPI_ISL_18458150 | | Central Public Health Laboratories-Sultanat of Oman | Central Public Health Laboratories-Sultanat of Oman | Yathrib Al Zakwani et al |
| **hCoV-19/Taiwan/CDC-F89555/2023** | EPI_ISL_18520815 | | Centers for Disease Control, R.O.C. (Taiwan) | Centers for Disease Control, R.O.C. (Taiwan) | Yu-Chi Lin et al |
| **hCoV-19/Zhejiang/SKLID_ZJU-43/2023** | EPI_ISL_18697856 | | State Key Laboratory for Diagnosis and Treatment of Infectious Diseases, The First Affiliated Hospital, Zhejiang University School of Medicine | State Key Laboratory for Diagnosis and Treatment of Infectious Diseases, The First Affiliated Hospital, Zhejiang University School of Medicine | Hangping Yao et al |
| **hCoV-19/Qinghai/QHCDC-HSZ-401/2023** | EPI_ISL_18268267 | | Microbiological testing center, Qinghai Center for Disease Prevention and Control | Microbiological testing center, Qinghai Center for Disease Prevention and Control | Youju Lei et al |
| **hCoV-19/Fujian/FCDC-ND2157/2023** | EPI_ISL_18523045 | | Institute for the Prevention and Control of Infectious Diseases, Fujian Provincial Center for Disease Control and Prevention | Institute for the Prevention and Control of Infectious Diseases, Fujian Provincial Center for Disease Control and Prevention | Qi Lin et al |
| **hCoV-19/Argentina/INEI126481/2024** | EPI_ISL_18930479 | | Servicio Virosis Respiratorias-Departamento Virología-INEI | Instituto Nacional Enfermedades Infecciosas C.G.Malbran | Pontoriero A. et al |
| **hCoV-19/Montenegro/DJ-A/2023** | EPI_ISL_18215550 | | Department for Molecular Diagnostics, Centre for Medical Microbiology, Institute of Public Health of Montenegro | Department for Molecular Diagnostics, Centre for Medical Microbiology, Institute of Public Health of Montenegro | Danijela Vujošević et al |
| **hCoV-19/Germany/BY-MVP-000020618/2023** | EPI_ISL_18264997 | | Max von Pettenkofer Institute, Virology, National Reference Center for Retroviruses, LMU Munich | Laboratory for Functional Genome Analysis; Dept. Genomics; Gene Center of the LMU Munich | Max Muenchhoff et al |
| **hCoV-19/England/CLIMB-CM7YROYC/2023** | EPI_ISL_18098103 | | Respiratory Virus Unit, Microbiology Services Colindale, Public Health England | UK Health Security Agency (UKHSA) | Angie Lackenby on behalf of PHE Covid Sequencing Team et al |
| **hCoV-19/Slovenia/90-001548-NM/2023** | EPI_ISL_18161231 | | National Laboratory for Health, Environment and Food, OMM, Novo mesto | NLZOH (National Laboratory for Health, Environment and Food) | Sandra Janezic et al |
| **hCoV-19/Iceland/L-4326/2023** | EPI_ISL_18094159 | | Landspitali Department of Clinical Microbiology | Landspitali Department of Clinical Microbiology | Freyja Valsdottir et al |
| **hCoV-19/Israel/SMC-7114861/2023** | EPI_ISL_18213449 | | Shamir Medical Center (Asaf Harofe) | Shamir Medical Center (Asaf Harofe) | Nir Rainy et al |
| **hCoV-19/Hungary/102821-UPL-003084/2023** | EPI_ISL_18552526 | | Institute of Laboratory Medicine, Semmelweis University and Epidemiology and Surveillance Centre, Semmelweis University | iBioscience Kft. | Gergő Túri et al |
| **hCoV-19/Bulgaria/23BG_NC_023918_R45/2023** | EPI_ISL_18825928 | | National Center of Infectious and Parasitic Diseases | National Center of Infectious and Parasitic Diseases | Ivelina Trifonova et al |
| **hCoV-19/Turkey/GED-14016_612/2023** | EPI_ISL_18784490 | | Turkish Society of Internal Medicine | Gen Era Diagnostics Corporation Life Science Research and Molecular Diagnostics | Serhat Ünal et al |
| **hCoV-19/Thailand/NIC_BKK_7763/2023** | EPI_ISL_18166621 | | Bureau of the Royal Household | National Institute of Health, Department of Medical Sciences, Ministry of Public Health, Thailand | Pilailuk Okada et al |
| **hCoV-19/Guatemala/DLNS-2375/2023** | EPI_ISL_18475369 | | Dirección del Laboratorio Nacional de Salud (DLNS), Guatemala | Dirección del Laboratorio Nacional de Salud (LNS), Guatemala | Maria Linda Mendoza Prillwitz et al |
| **hCoV-19/Croatia/HZJZ_45962/2024** | EPI_ISL_18913796 | | Klinički bolnički centar Split | Hrvatski zavod za javno zdravstvo | Irena Tabain et al |
| **hCoV-19/Finland/THL-01861/2024** | EPI_ISL_18918971 | | Seinäjoen keskussairaala | Expert Microbiology, National Institute for Health and Welfare | Erika Lindh et al |
| **hCoV-19/Lithuania/MB0912_0125/2023** | EPI_ISL_18393753 | | VIEŠOJI ĮSTAIGA CENTRO POLIKLINIKA | National Public Health Surveillance Laboratory | Martynas Smicius et al |
| **hCoV-19/Laos/IPL-78530/2024** | EPI_ISL_18938889 | | NCLE | Institut Pasteur du Laos | Kedkeo INTAVONG et al |
| **hCoV-19/Chile/ML-84994/2023** | EPI_ISL_18692505 | | Genetica Molecular and Subdepartamento de Virologia ISP Chile | Instituto de Salud Publica de Chile | Mariela Guajardo et al |
| **hCoV-19/Myanmar/DMR_24001/2023** | EPI_ISL_18744368 | | Department of Medical Research | Department of Medical Research | Myat Htut Nyunt et al |
| **hCoV-19/Ecuador/NIC-INSPI-435188/2023** | EPI_ISL_18770128 | | INSPI-CRN DE INFLUENZA Y OTROS VIRUS RESPIRATORIOS | INSPI-CRN DE INFLUENZA Y OTROS VIRUS RESPIRATORIOS | Alfredo Bruno et al |
| **hCoV-19/NewZealand/23YA7490/2023** | EPI_ISL_18670598 | | Pathlab Rotorua | Institute of Environmental Science and Research (ESR) | Sarah Cockerton et al |
| **hCoV-19/Thailand/NIC_BKK_8529/2023** | EPI_ISL_18556654 | | Bumrungrad International Hospital | National Institute of Health, Department of Medical Sciences, Ministry of Public Health, Thailand | Pilailuk Okada et al |
| **hCoV-19/Ecuador/GENSBIO-INSPI-SG-716/2023** | EPI_ISL_18646531 | | Laboratorio de Influenza y Otros Virus Respiratorios, INSPI-CZ9 | Centro de Referencia Nacional de Genomica, Secuenciacion y Bioinformatica GENSBIO, INSPI-CZ9 | Andres Carrazco-Montalvo* et al |
| **hCoV-19/USA/IN-CDC-QDX86946331/2023** | EPI_ISL_18534188 | | Quest Diagnostics Incorporated | Centers for Disease Control and Prevention Division of Viral Diseases, Pathogen Discovery | Dakota Howard et al |

| | | | | | |
|---|---|---|---|---|---|
| **hCoV-19/Norway/10552/2023** | EPI_ISL_18526364 | | Vestfold Hospital, Toensberg, Department of Microbiology | Norwegian Institute of Public Health, Department of Virology | Kathrine Stene-Johansen et al |
| **hCoV-19/USA/DE-DHSS-B1262114/2023** | EPI_ISL_18647157 | | Delaware Public Health Lab | Delaware Public Health Lab | Holly Miller et al |
| **hCoV-19/Ecuador/NIC-INSPI-0006-5970/2024** | EPI_ISL_18879758 | | INSPI-CRN DE INFLUENZA Y OTROS VIRUS RESPIRATORIOS | INSPI-CRN DE INFLUENZA Y OTROS VIRUS RESPIRATORIOS | Alfredo Bruno et al |
| **hCoV-19/Chile/MA-96421/2023** | EPI_ISL_18731460 | | Genetica Molecular and Subdepartamento de Virologia ISP Chile | Instituto de Salud Publica de Chile | Mariela Guajardo et al |
| **hCoV-19/Ireland/D-SVUH-FAX24627-025/2023** | EPI_ISL_18356704 | | St. Vincent's Microbiology | St. Vincent's Microbiology | Eric Brown et al |
| **hCoV-19/Mexico/QUE_InDRE_FB4820/2023** | EPI_ISL_18785805 | | Laboratorio Estatal de Salud Publica Queretaro | Instituto de Diagnostico y Referencia Epidemiologicos (INDRE) | Abril Rodriguez-Maldonado et al |
| **hCoV-19/Slovakia/uvzsr-BA_23_18_00008645/2023** | EPI_ISL_18641961 | | Laboratory of Genomics and Bioinformatics, Comenius University Science Park | Laboratory of Genomics and Bioinformatics, Comenius University Science Park | Tomáš Szemes et al |
| **hCoV-19/Heilongjiang/HLJCDC-1517/2023** | EPI_ISL_17850401 | | Institute for Viral Disease Control and Prevention, Department for Viral Disease Control and Prevention, Heilongjiang Provincial Center for Disease Control and Prevention | Institute for Viral Disease Control and Prevention, Department for Viral Disease Control and Prevention, Heilongjiang Provincial Center for Disease Control and Prevention | Xu Jun et al |
| **hCoV-19/Anhui/HCDC-142/2023** | EPI_ISL_18289739 | | Microbiology lab, Huainan City Center for Disease Control and Prevention | Microbiology lab, Huainan City Center for Disease Control and Prevention | Liu Jiang et al |
| **hCoV-19/Russia/SPE-RII-MH156224S/2023** | EPI_ISL_18737099 | | Saint Petersburg Dermatovenerologic Dispensary No 1 | WHO National Influenza Centre Russian Federation | Andrey Komissarov et al |
| **hCoV-19/Liaoning/LNCDC-415-1/2023** | EPI_ISL_18518301 | | Lingling Mao, Haibo Sun, Shuang Liu, Xiaoting Xu, Xuesheng Liu, Lulu Wang, Bo Wang, Lu Lei, Sinong Sun, Ge Song, Hanyi Guo | Institute for Communicable and Infectious Diseases Prevention and Control, Liaoning Center for Disease Control and Prevention | Mao et al |
| **hCoV-19/Jiangsu/ZJCDC085/2023** | EPI_ISL_18105969 | | Zhenjiang CDC | Institute for the Prevention and Control of Acute Infectious Diseases, Jiangsu Provincial Center for Disease Control and Prevention | Zhou et al |
| **hCoV-19/Sichuan/MYCDC-XGCX212/2023** | EPI_ISL_17978355 | | Mianyang Center of Disease Control and Prevention | Microbiology laboratory, Sichuan Center of Disease Control and Prevention | Yuke Dai et al |
| **hCoV-19/Zhejiang/SKLID_ZJU-S163/2023** | EPI_ISL_18679880 | | State Key Laboratory for Diagnosis and Treatment of Infectious Diseases, The First Affiliated Hospital, Zhejiang University School of Medicine | State Key Laboratory for Diagnosis and Treatment of Infectious Diseases, The First Affiliated Hospital, Zhejiang University School of Medicine | Hangping Yao et al |
| **hCoV-19/Shanxi/TYCDC557/2023** | EPI_ISL_18495488 | | Microbiology Department, Taiyuan Center for Disease Control and Prevention | Microbiology Department, Taiyuan Center for Disease Control and Prevention | Wang et al |
| **hCoV-19/Xinjiang/XJCDCKLMY_294/2023** | EPI_ISL_18505644 | | Disease control, Xinjiang center of diease control | Disease control, Xinjiang center of diease control | Ma et al |
| **hCoV-19/Xinjiang/XJCDCKLMY-261/2023** | EPI_ISL_18320923 | | Xinjiang center of diease control, Jianquan street | Xinjiang center of diease control, Jianquan street | Ma et al |
| **hCoV-19/Henan/XYCDC-0251/2023** | EPI_ISL_18208330 | | Central laboratory, Xinyang Center for Disease Control and Prevention | Central laboratory, Xinyang Center for Disease Control and Prevention | Lv et al |
| **hCoV-19/Sweden/U-R056589/2023** | EPI_ISL_18338768 | | The Public Health Agency of Sweden | The Public Health Agency of Sweden | Emmi Andersson et al |
| **hCoV-19/Sichuan/MYCDC-XGCX444/2023** | EPI_ISL_18518324 | | Mianyang Center of Disease Control and Prevention | Microbiology laboratory, Sichuan Center of Disease Control and Prevention | Dai et al |
| **hCoV-19/Liaoning/LNCDC-275-1/2023** | EPI_ISL_18116286 | | Institute for Communicable and Infectious Diseases Prevention and Control, Liaoning Center for Disease Control and Prevention | Institute for Communicable and Infectious Diseases Prevention and Control, Liaoning Center for Disease Control and Prevention | Mao et al |
| **hCoV-19/Henan/JZCDC-144/2023** | EPI_ISL_18300573 | | Health Test and Surveilance Section, Jiaozuo Center for Disease Control and Prevention | Health Test and Surveilance Section, Jiaozuo Center for Disease Control and Prevention | Zhang et al B |
| **hCoV-19/NewZealand/23YA6112/2023** | EPI_ISL_18562393 | | Wellington SCL | Institute of Environmental Science and Research (ESR) | Sarah Cockerton et al |
| **hCoV-19/Yunnan/YNCDC-3296/2023** | EPI_ISL_18602864 | | The department for prevention and control of acute infectious diseases, Yunnan Center for Disease Control and Prevention | The department for prevention and control of acute infectious diseases, Yunnan Center for Disease Control and Prevention | Zhang et al B |
| **hCoV-19/USA/HI-TAMC_2960/2023** | EPI_ISL_18347037 | | Tripler Army Medical Center | Tripler Army Medical Center, Dept of Clinical Investigation, | Keith S. K. Fong et al |
| **hCoV-19/India/TG-ICMR-NIV-G-14547/2023** | EPI_ISL_18120254 | | Indian Council of Medical Research, National Institute of Virology | National Institute of Virology - Influenza | Dr. Varsha Potdar et al |
| **hCoV-19/Qinghai/QHCDC-HNYY-40/2023** | EPI_ISL_18078206 | | Microbiological testing center, Qinghai Center for Disease Prevention and Control | Microbiological testing center, Qinghai Center for Disease Prevention and Control | Huang et al |
| **hCoV-19/Jilin/JSY-CC9/2023** | EPI_ISL_18908495 | | Changchun Veterinary Research Institute, Chinese Academy of Agricultural Sciences | Changchun Veterinary Research Institute, Chinese Academy of Agricultural Sciences | Yuewei Gao et al |
| **hCoV-19/Sweden/AC-460303272887/2023** | EPI_ISL_18558351 | | The Public Health Agency of Sweden | The Public Health Agency of Sweden | Emmi Andersson et al |
| **hCoV-19/Tibet/XZCDC_0183/2023** | EPI_ISL_18672238 | | Inspection and verification office, Center for Disease Control and Prevention of Tibet Autonomous Region | Inspection and verification office, Center for Disease Control and Prevention of Tibet Autonomous Region | Hong et al |
| **hCoV-19/SouthKorea/KDCA_270640/2023** | EPI_ISL_18503595 | | Division of Emerging Infectious Diseases, Bureau of Infectious Diseases Diagnosis Control, Korea Disease Control and Prevention Agency | Division of Emerging Infectious Diseases, Bureau of Infectious Diseases Diagnosis Control, Korea Disease Control and Prevention Agency | Jeong-Ah Kim et al |
| **hCoV-19/Beijing/BJCDC-4519/2023** | EPI_ISL_18438307 | | Institute for Infectious Disease and Endemic Disease Control, Beijing Center for Disease Control and Prevention | Institute for Infectious Disease and Endemic Disease Control, Beijing Center for Disease Control and Prevention | Pan et al |
| **hCoV-19/Heilongjiang/HLJCDC-2191/2023** | EPI_ISL_18246124 | | Institute for Viral Disease Control and Prevention, Department for Viral Disease Control and Prevention, Heilongjiang Provincial Center for Disease Control and Prevention | Institute for Viral Disease Control and Prevention, Department for Viral Disease Control and Prevention, Heilongjiang Provincial Center for Disease Control and Prevention | Xu Jun et al |
| **hCoV-19/Qatar/DA-USAFSAM-S21612/2023** | EPI_ISL_18566172 | | US Air Force School of Aerospace Medicine | US Air Force School of Aerospace Medicine | Anthony Fries et al |
| **hCoV-19/Shanghai/SCDC-795/2023** | EPI_ISL_18268346 | | Microbe lab, Shanghai Municipal Center for Disease Control & Prevention | Microbe lab, Shanghai Municipal Center for Disease Control & Prevention | Zhang et al B |
| **hCoV-19/Gansu/GSCDC-0518/2023** | EPI_ISL_18634064 | | Gansu Provincial Pathogen Detection Institute, Gansu Provincial Centre for Disease Control and Prevention | Gansu Provincial Pathogen Detection Institute, Gansu Provincial Centre for Disease Control and Prevention | Li et al |
| **hCoV-19/SouthKorea/KDCA264807/2023** | EPI_ISL_18312954 | | Division of Emerging Infectious Diseases, Bureau of Infectious Diseases Diagnosis Control, Korea Disease Control and Prevention Agency | Division of Emerging Infectious Diseases, Bureau of Infectious Diseases Diagnosis Control, Korea Disease Control and Prevention Agency | Jeong-Ah Kim et al |
| **hCoV-19/Bonaire/BQ-RIVM-132688/2023** | EPI_ISL_18463494 | | National Institute for Public Health and the Environment (RIVM) | National Institute for Public Health and the Environment (RIVM) | Adam Meijer et al |
| **hCoV-19/Heilongjiang/HLJCDC-2360/2023** | EPI_ISL_18465128 | | Institute for Viral Disease Control and Prevention, Department for Viral Disease Control and Prevention, Heilongjiang Provincial Center for Disease Control and Prevention | Institute for Viral Disease Control and Prevention, Department for Viral Disease Control and Prevention, Heilongjiang Provincial Center for Disease Control and Prevention | Xu Jun et al |
| **hCoV-19/Henan/ZMDCDC-318/2023** | EPI_ISL_18485235 | | Inspection Division, Zhumadian Center for Disease Control and Prevention | Inspection Division, Zhumadian Center for Disease Control and Prevention | Xu Tao et al |
| **hCoV-19/Gansu/PLCDC-0196/2023** | EPI_ISL_18425065 | | Health Inspection Institute, Pingliang City Center for Disease Control and Prevention | Health Inspection Institute, Pingliang City Center for Disease Control and Prevention | Zhang et al B |
| **hCoV-19/Heilongjiang/HLJCDC-2645/2023** | EPI_ISL_18716344 | | Institute for Viral Disease Control and Prevention, Department for Viral Disease Control and Prevention, Heilongjiang Provincial Center for Disease Control and Prevention | Institute for Viral Disease Control and Prevention, Department for Viral Disease Control and Prevention, Heilongjiang Provincial Center for Disease Control and Prevention | Xu Jun et al |
| **hCoV-19/Chile/AR-82655/2023** | EPI_ISL_18483673 | | Genetica Molecular and Subdepartamento de Virologia ISP Chile | Instituto de Salud Publica de Chile | Mariela Guajardo et al |
| **hCoV-19/Guizhou/GZCDC-ZY-08-5/2023** | EPI_ISL_18217121 | | Experiment Center, Guizhou Provincial Center for Disease Control and Prevention | Experiment Center, Guizhou Provincial Center for Disease Control and Prevention | Jiang et al |
| **hCoV-19/Gansu/BYCDC155/2023** | EPI_ISL_18166109 | | Inspection and testing department, Center for Disease Control and Preventron of Baiyin City | Inspection and testing department, Center for Disease Control and Preventron of Baiyin City | Yang et al |
| **hCoV-19/Tibet/XZCDC_0089/2023** | EPI_ISL_18602831 | | Inspection and verification office, Center for Disease Control and Prevention of Tibet Autonomous Region | Inspection and verification office, Center for Disease Control and Prevention of Tibet Autonomous Region | Hong et al |
| **hCoV-19/Australia/VIC-VIDRL-00012038/2023** | EPI_ISL_18350061 | | Royal Melbourne Hospital - Microbiology Department | Victorian Infectious Disease Reference Laboratory | Aziz et al |
| **hCoV-19/Italy/FVG-TS-65346531/2023** | EPI_ISL_18446734 | | SC (UCO) Igiene e Sanità Pubblica, ASUGI | SC (UCO) Igiene e Sanità Pubblica, ASUGI | Piscianz E et al |
| **hCoV-19/Heilongjiang/HLJCDC-1995/2023** | EPI_ISL_18246198 | | Institute for Viral Disease Control and Prevention, Department for Viral Disease Control and Prevention, Heilongjiang Provincial Center for Disease Control and Prevention | Institute for Viral Disease Control and Prevention, Department for Viral Disease Control and Prevention, Heilongjiang Provincial Center for Disease Control and Prevention | Xu Jun et al |
| **hCoV-19/Australia/VIC-VIDRL-00012194/2023** | EPI_ISL_18488509 | | Royal Melbourne Hospital - Microbiology Department | Victorian Infectious Disease Reference Laboratory | Aziz et al |
| **hCoV-19/Gansu/GSCDC-0914/2023** | EPI_ISL_18683278 | | Gansu Provincial Pathogen Detection Institute, Gansu Provincial Centre for Disease Control and Prevention | Gansu Provincial Pathogen Detection Institute, Gansu Provincial Centre for Disease Control and Prevention | Li et al |
| **hCoV-19/Sichuan/YBCDC-XGCX339/2023** | EPI_ISL_18133993 | | Yibin Center of Disease Control and Prevention | Microbiology laboratory, Sichuan Center of Disease Control and Prevention | Jiang et al |
| **hCoV-19/Liaoning/LNCDC-485-2/2023** | EPI_ISL_18692899 | | Institute for Communicable and Infectious Diseases Prevention and Control, Liaoning Center for Disease Control and Prevention | Institute for Communicable and Infectious Diseases Prevention and Control, Liaoning Center for Disease Control and Prevention | Mao et al |
| **hCoV-19/Qinghai/QHCDC-HSZ-355/2023** | EPI_ISL_18133939 | | Microbiological testing center, Qinghai Center for Disease Prevention and Control | Microbiological testing center, Qinghai Center for Disease Prevention and Control | Zhong et al |
| **hCoV-19/Henan/XYCDC-0407/2023** | EPI_ISL_18523038 | | Central laboratory, Xinyang Center for Disease Control and Prevention | Central laboratory, Xinyang Center for Disease Control and Prevention | Lv et al |
| **hCoV-19/Sichuan/BZCDC-175/2023** | EPI_ISL_18780176 | | Bazhong Center of Disease Control and Prevention | Microbiology laboratory, Sichuan Center of Disease Control and Prevention | Wang et al |
| **hCoV-19/Tibet/LZCDC_001/2024** | EPI_ISL_18876880 | | Microbiology laboratory, Nyingchi Center for Disease Control and Prevention | Microbiology laboratory, Nyingchi Center for Disease Control and Prevention | Ma et al |
| **hCoV-19/Xinjiang/XJCDCKLMY-003/2024** | EPI_ISL_18858721 | | Xinjiang CDC | Xinjiang CDC | Ma et al |
| **hCoV-19/Liaoning/LNCDC-334-1/2023** | EPI_ISL_18320689 | | Institute for Communicable and Infectious Diseases Prevention and Control, Liaoning Center for Disease Control and Prevention | Institute for Communicable and Infectious Diseases Prevention and Control, Liaoning Center for Disease Control and Prevention | Mao et al |
| **hCoV-19/Liaoning/LNCDC-376-3/2023** | EPI_ISL_18425079 | | Institute for Communicable and Infectious Diseases Prevention and Control, Liaoning Center for Disease Control and Prevention | Institute for Communicable and Infectious Diseases Prevention and Control, Liaoning Center for Disease Control and Prevention | Mao et al |
| **hCoV-19/Beijing/BJCDC-4308/2023** | EPI_ISL_18438114 | | Institute for Infectious Disease and Endemic Disease Control, Beijing Center for Disease Control and Prevention | Institute for Infectious Disease and Endemic Disease Control, Beijing Center for Disease Control and Prevention | Pan et al |
| **hCoV-19/NewZealand/23YA4676/2023** | EPI_ISL_18292297 | | Medlab Central (PN) | Institute of Environmental Science and Research (ESR) | Sarah Cockerton et al |
| **hCoV-19/Laos/IPL-78490/2023** | EPI_ISL_18660968 | | National Centre for Laboratory and Epidemiology (NCLE) | Institut Pasteur du Laos | Thonglakhone XAYBOUNSOU et al |
| **hCoV-19/CzechRepublic/NRL-s202/2024** | EPI_ISL_18908228 | | University Hospital Brno | National Institute of Public Health | Timotej Suri et al |
| **hCoV-19/Guizhou/GZCDC-ZY-08-17/2023** | EPI_ISL_18217129 | | Experiment Center, Guizhou Provincial Center for Disease Control and Prevention | Experiment Center, Guizhou Provincial Center for Disease Control and Prevention | Jiang et al |
| **hCoV-19/Singapore/R74MR67/2023** | EPI_ISL_18397622 | | National Public Health Laboratory, National Centre for Infectious Diseases | National Public Health Laboratory, National Centre for Infectious Diseases | Benny Yeo et al |
| **hCoV-19/Brunei/MGS-7123072104/2023** | EPI_ISL_18485089 | | National Virology Reference Laboratory | Microbial Genomic Services | Zainun Zaini et al |
| **hCoV-19/Liaoning/LNCDC-053-4/2024** | EPI_ISL_18930351 | | Institute for Communicable and Infectious Diseases Prevention and Control, Liaoning Center for Disease Control and Prevention | Institute for Communicable and Infectious Diseases Prevention and Control, Liaoning Center for Disease Control and Prevention | Mao et al |
| **hCoV-19/Heilongjiang/HLJCDC-0041/2024** | EPI_ISL_18857580 | | Institute for Viral Disease Control and Prevention, Department for Viral Disease Control and Prevention, Heilongjiang Provincial Center for Disease Control and Prevention | Institute for Viral Disease Control and Prevention, Department for Viral Disease Control and Prevention, Heilongjiang Provincial Center for Disease Control and Prevention | Xu Jun et al |
| **hCoV-19/Hubei/HCDC-XY3253/2023** | EPI_ISL_18438438 | | Xiangyang Center for Disease Control and Prevention | Hubei Provincial Center for Disease Control and Prevention Center of Pathogenic Microorganism Detection | Yongzhong Jiang et al |
| **hCoV-19/Yunnan/YNCDC-3562/2023** | EPI_ISL_18876961 | | The department for prevention and control of acute infectious diseases, Yunnan Center for Disease Control and Prevention | The department for prevention and control of acute infectious diseases, Yunnan Center for Disease Control and Prevention | Zhang et al B |
| **hCoV-19/Shanxi/JCCDC-JCCX0638/2023** | EPI_ISL_18805118 | | Department of Microbiology, Jincheng Centers for Disease Control and prevention | Department of Microbiology, Jincheng Centers for Disease Control and prevention | Zhang et al B |
| **hCoV-19/Anhui/BZCDC-203/2023** | EPI_ISL_18805132 | | Clinical laboratory, Bozhou Municipal Center for Disease Control & Prevention | Clinical laboratory, Bozhou Municipal Center for Disease Control & Prevention | Qingqiao et al |
| **hCoV-19/Liaoning/LNCDC-438-4/2023** | EPI_ISL_18615881 | | Institute for Communicable and Infectious Diseases Prevention and Control, Liaoning Center for Disease Control and Prevention | Institute for Communicable and Infectious Diseases Prevention and Control, Liaoning Center for Disease Control and Prevention | Mao et al |
| **hCoV-19/Shanghai/SCDC-HP01-32/2024** | EPI_ISL_18941349 | | Microbe lab, Shanghai Municipal Center for Disease Control & Prevention | Microbe lab, Shanghai Municipal Center for Disease Control & Prevention | Chen et al |
| **hCoV-19/HongKong/VB23241505/2023** | EPI_ISL_18705758 | | Tuen Mun Hospital | Hong Kong Department of Health | Alan K.L. Tsang et al |
| **hCoV-19/Fujian/FCDC-ND0735/2023** | EPI_ISL_18447513 | | Institute for the Prevention and Control of Infectious Diseases, Fujian Provincial Center for Disease Control and Prevention | Institute for the Prevention and Control of Infectious Diseases, Fujian Provincial Center for Disease Control and Prevention | Qi Lin et al |
| **hCoV-19/HongKong/VB23215412/2023** | EPI_ISL_18526590 | | Yan Chai Hospital | Hong Kong Department of Health | Alan K.L. Tsang et al |
| **hCoV-19/Hubei/HCDC-SY3513/2023** | EPI_ISL_18508539 | | Shiyan Center for Disease Control and Prevention | Hubei Provincial Center for Disease Control and Prevention Center of Pathogenic Microorganism Detection | Yongzhong Jiang et al |
| **hCoV-19/Guizhou/GZSCDC-TR-09-014/2023** | EPI_ISL_18300562 | | Experiment Center, Guizhou Provincial Center for Disease Control and Prevention | Experiment Center, Guizhou Provincial Center for Disease Control and Prevention | Ke et al |
| **hCoV-19/Yunnan/YNCDC-2997/2023** | EPI_ISL_18409394 | | The department for prevention and control of acute infectious diseases, Yunnan Center for Disease Control and Prevention | The department for prevention and control of acute infectious diseases, Yunnan Center for Disease Control and Prevention | Zhang et al B |
| **hCoV-19/Xinjiang/XJCDCBOZ-256/2023** | EPI_ISL_18858697 | | Xinjiang CDC | Xinjiang CDC | Ma et al |
| **hCoV-19/Heilongjiang/HLJCDC-2522/2023** | EPI_ISL_18615802 | | Institute for Viral Disease Control and Prevention, Department for Viral Disease Control and Prevention, Heilongjiang Provincial Center for Disease Control and Prevention | Institute for Viral Disease Control and Prevention, Department for Viral Disease Control and Prevention, Heilongjiang Provincial Center for Disease Control and Prevention | Xu Jun et al |
| **hCoV-19/Hubei/HCDC-TM3296/2023** | EPI_ISL_18438418 | | Tianmen City Center for Disease Control and Prevention | Hubei Provincial Center for Disease Control and Prevention Center of Pathogenic Microorganism Detection | Yongzhong Jiang et al |
| **hCoV-19/Anhui/AQCDC-169/2023** | EPI_ISL_18589864 | | Anqing Center for Disease Control and Prevention lab | Microbiological Testing Department, Anqing Center for Disease Control and Prevention | Yuxin Li et al |
| **hCoV-19/Shanxi/JCCDC-JCCX0598/2023** | EPI_ISL_18602688 | | Department of Microbiology, Jincheng Centers for Disease Control and prevention | Department of Microbiology, Jincheng Centers for Disease Control and prevention | Zhang et al B |
| **hCoV-19/USA/MI-MDHHS-SC44892/2023** | EPI_ISL_18669942 | | Michigan Department of Health and Human Services, Bureau of Laboratories | Michigan Department of Health and Human Services, Bureau of Laboratories | Blankenship HM et al |
| **hCoV-19/Taiwan/CDC-F90435/2023** | EPI_ISL_18941596 | | Centers for Disease Control, R.O.C. (Taiwan) | Centers for Disease Control, R.O.C. (Taiwan) | Yu-Chi Lin et al |
| **hCoV-19/Yunnan/YNCDC-2701/2023** | EPI_ISL_18292834 | | The department for prevention and control of acute infectious diseases, Yunnan Center for Disease Control and Prevention | The department for prevention and control of acute infectious diseases, Yunnan Center for Disease Control and Prevention | Zhang et al B |
| **hCoV-19/Sichuan/ZGCDC-zgxg410/2023** | EPI_ISL_18376573 | | Zigong Center of Disease Control and Prevention | Microbiology laboratory, Sichuan Center of Disease Control and Prevention | Wang et al |
| **hCoV-19/USA/CT-CDC-QDX85012827/2023** | EPI_ISL_18317078 | | Quest Diagnostics Incorporated | Centers for Disease Control and Prevention Division of Viral Diseases, Pathogen Discovery | Dakota Howard et al |
| **hCoV-19/Sichuan/MYCDC-428/2023** | EPI_ISL_18481020 | | Mianyang Center of Disease Control and Prevention | Microbiology laboratory, Sichuan Center of Disease Control and Prevention | Dai et al |
| **hCoV-19/Chile/AR-98749/2023** | EPI_ISL_18768409 | | Genetica Molecular and Subdepartamento de Virologia ISP Chile | Instituto de Salud Publica de Chile | Mariela Guajardo et al |
| **hCoV-19/Jilin/JSY-CC11/2023** | EPI_ISL_18908496 | | Changchun Veterinary Research Institute, Chinese Academy of Agricultural Sciences | Changchun Veterinary Research Institute, Chinese Academy of Agricultural Sciences | Yuewei Gao et al |
| **hCoV-19/Cambodia/IKCM230104/2023** | EPI_ISL_18633891 | | National Institute of Public Health | National Institute of Public Health | Chhorvann CHHEA et al |
| **hCoV-19/Cambodia/PKH230596/2023** | EPI_ISL_18633888 | | National Institute of Public Health | National Institute of Public Health | Chhorvann CHHEA et al |
| **hCoV-19/Henan/XCCDC110102/2023** | EPI_ISL_18535350 | | Clinical Laboratory, Xuchang CDC | Clinical Laboratory, Xuchang CDC | Xia et al |
| **hCoV-19/Henan/HBCDC-CX032/2024** | EPI_ISL_18943907 | | CDC, Hebi Center for Disease Control and Prevention | CDC, Hebi Center for Disease Control and Prevention | Wang Xi et al |
| **hCoV-19/Xinjiang/XJCDCWLMQ-588/2023** | EPI_ISL_18495147 | | Xinjiang CDC | Xinjiang CDC | Ma et al |
| **hCoV-19/Yunnan/YNCDC-3325/2023** | EPI_ISL_18602992 | | The department for prevention and control of acute infectious diseases, Yunnan Center for Disease Control and Prevention | The department for prevention and control of acute infectious diseases, Yunnan Center for Disease Control and Prevention | Zhang et al B |
| **hCoV-19/Pakistan/IDRL-AKU-KHI-5448-S8/2023** | EPI_ISL_18745110 | | Clinical Lab, Aga Khan University Hospital | Infectious Diseases Research Laboratory, Department of Pediatrics and Child Health | Waqasuddin Khan et al |
| **hCoV-19/Ecuador/GENSBIO-INSPI-SG-663/2023** | EPI_ISL_18548348 | | Laboratorio de Influenza y Otros Virus Respiratorios, INSPI-CZ9 | Centro de Referencia Nacional de Genomica, Secuenciacion y Bioinformatica GENSBIO, INSPI-CZ9 | Andres Carrazco-Montalvo* et al |
| **hCoV-19/Cambodia/NPH230894/2023** | EPI_ISL_18633725 | | National Institute of Public Health | National Institute of Public Health | Chhorvann CHHEA et al |
| **hCoV-19/Xinjiang/XJCDCWLMQ-658/2023** | EPI_ISL_18659027 | | Xinjiang CDC | Xinjiang CDC | Ma et al |
| **hCoV-19/Anhui/BZCDC-210/2023** | EPI_ISL_18805139 | | Clinical laboratory, Bozhou Municipal Center for Disease Control & Prevention | Clinical laboratory, Bozhou Municipal Center for Disease Control & Prevention | Qingqiao et al |
| **hCoV-19/Peru/ARE-INS-48305/2024** | EPI_ISL_18872391 | | Laboratorio de Referencia Nacional de Virus Inmunoprevenibles, Centro Nacional de Salud Publica, Instituto Nacional de Salud Peru. | Laboratorio de Referencia Nacional de Virus Inmunoprevenibles, Centro Nacional de Salud Publica, Instituto Nacional de Salud Peru. | Carlos Padilla Rojas et al |
| **hCoV-19/Heilongjiang/HLJCDC-2598/2023** | EPI_ISL_18615862 | | Institute for Viral Disease Control and Prevention, Department for Viral Disease Control and Prevention, Heilongjiang Provincial Center for Disease Control and Prevention | Institute for Viral Disease Control and Prevention, Department for Viral Disease Control and Prevention, Heilongjiang Provincial Center for Disease Control and Prevention | Xu Jun et al |
| **hCoV-19/Gansu/GSCDC-0977/2023** | EPI_ISL_18769806 | | Gansu Provincial Pathogen Detection Institute, Gansu Provincial Centre for Disease Control and Prevention | Gansu Provincial Pathogen Detection Institute, Gansu Provincial Centre for Disease Control and Prevention | Li et al |
